# Computationally scalable regression modeling for ultrahigh-dimensional omics data with ParProx

**DOI:** 10.1101/2021.01.10.426142

**Authors:** Seyoon Ko, Ginny X. Li, Hyungwon Choi, Joong-Ho Won

**Affiliations:** Department of Statistics, Seoul National University, Republic of Korea; Department of Medicine, Yong Loo Lin School of Medicine, National University of Singapore, Singapore

## Abstract

Statistical analysis of ultrahigh-dimensional omics scale data has long depended on univariate hypothesis testing. With growing data features and samples, the obvious next step is to establish multivariable association analysis as a routine method for understanding genotype-phenotype associations. Here we present ParProx, a state-of-the-art implementation to optimize overlapping group lasso regression models for time-to-event and classification analysis, guided by biological priors through coordinated variable selection. ParProx not only enables model fitting for ultrahigh-dimensional data within the architecture for parallel or distributed computing, but also allows users to obtain interpretable regression models consistent with known biological relationships among the independent variables, a feature long neglected in statistical modeling of omics data. We demonstrate ParProx using three different omics data sets of moderate to large numbers of variables, where we use genomic regions and pathways to arrive at sparse regression models comprised of biologically related independent variables. ParProx is naturally applicable to a wide range of studies using ultrahigh-dimensional omics data, ranging from genome-wide association analysis to single cell sequencing studies where multivariable modeling is computationally intractable.

## Introduction

Omics technologies are principal modalities in today’s systems biology and molecular research. Despite significant advances and wide applicability, clinical omics data sets are often modestly sized in a vast majority of biomedical studies, often providing insufficient statistical information for phenotype association analysis or prognostication. Since the arrival of gene expression microarrays in the 1990s, the small sample size in omics data has challenged biostatisticians and bioinformaticians to grapple with the “small *n*, large *p*” problem, which constrains the ability of statistical models or machine learning methods to exploit the multi-dimensional feature space at a sufficient depth. As a consequence, omics data analysis has inevitably depended on univariate hypothesis testing for individual molecular features in combination with *post hoc* multiple testing correction procedures to control overall type I error, further complemented by enrichment analysis of biological processes and pathways in the molecular features not discarded by hypothesis testing. However, hypothesis testing-based analysis does not describe genotype-phenotype relationships in a multivariate space, and statistical analysis results conducted as such often ignore the covariance between functionally related molecules – the underlying structure among data features in omics data sets.

There are well-established approaches to multivariable modeling of high-dimensional data in supervised analysis problems, ranging from regularized linear regression models^1, 2, 3^ to machine learning methods such as decision tree-based random forests (RF)^4^ and support vector machines (SVM)^5^. The spectrum of model complexity in different approaches is often characterized by the trade-off between bias and variance for prediction.^6^ On the one hand, more complex models tend to explain a training data set well, but an overfit model bears large variance in prediction between samples with similar data features in external data sets. On the other hand, simpler models may not fit the training data well compared to more complex machine learning methods, but predictions made by simpler models in independent data sets are more stable and less sensitive to small differences in the training data. With the sample size almost always smaller than the number of molecules measured in omics data, the overfitting problem of complex models is unavoidable in most application of machine learning methods, especially in the model-free approaches. By contrast, simpler methods such as regularized regression models are not flexible enough to capture non-linear relationships, but they produce highly interpretable models with low variance in predictions.

Today, with increasing throughput and decreasing cost of the experimental platforms, we are already transitioning from the era of small *n*, large *p* problems to a time of “large *n*, very large *p*” problems. This phenomenon is perhaps best exemplified by genome-wide association studies with genotypes at millions of loci and with a sample size greater than hundreds of thousands^7^, or multi-omics studies with tens of thousands of tumor biopsies in the Cancer Genome Atlas. (TCGA)^8^ In spite of the emergence of large-sample, high-dimensional omics studies, most data modeling approaches implicitly require in-memory storage of data and computation on central processing units (CPUs), which may be computationally expensive on standard computer hardware. The breaking point is yet to be recognized by those who routinely perform omics data analysis as the computer hardware has improved over time, but this emerging reality poses implementation challenges for future development of biostatistical and bioinformatics tools.

Motivated by these considerations, here we present a new software for fitting regularized linear regression models on high-dimensional clinical omics data, embodying an efficient optimization strategy and optional capability for parallel/distributed computing options. The implementation, called ParProx, fits the group least absolute shrinkage and selection operator (group lasso) regression models for survival analysis or sample classification. During model fitting, variables are regularized by non-overlapping or overlapping group penalties specified by the user and the variables in the same group are penalized jointly to reflect known group information such as pathways and gene ontologies (GO). More importantly, we implemented ParProx in the Julia language to allow for parallel computation with graphics processing units (GPUs) or distributed computing over cloud environments (Amazon Web Services, Google Cloud Platform, Microsoft Azure, etc.) natively, which enable the modeling for ultrahigh-dimensional data sets. We provide detailed protocols and illustration through three representative example data sets below.

## Results

### Overview of applications

We demonstrate regression modeling of clinical omics data using ParProx through three case studies. In the first application, we present a Cox regression analysis for overall survival outcome in 9,707 patients in TCGA, using the somatic mutations as predictor variables.^8, 9^ We create the predictor variables by counting mutations for ~56,000 DNA sequence segments in the codons and regulatory regions, in which mutation counts of individual sequence segments become independent variables (or covariates) and the individual proteins or interacting protein pairs harboring those sequence segments are considered as variable groups for structured regularization. In the second application, we establish a gene expression-based logistic regression model for pathological complete response (pCR) to neoadjuvant chemotherapy for breast cancer, using ~12,000 mRNA-level measurements in 469 patients as covariates and pathways/GO terms as the *overlapping* variable groups for structured regularization.^10, 11, 12, 13^ In the last application, we present a Cox regression analysis of 428 liver cancer patients using DNA methylation status of CpG islands in and out of coding regions as covariates. Each methylation probe represents a CpG island on the genome, and the probes associated with the same gene form a variable group in this case. As certain chromosomal regions are densely populated by multiple protein coding genes, some probes may belong to multiple adjacent genes, creating overlap in the variable groups, i.e. between adjacent genes. The methylation array platform used by TCGA contains as many as >865,000 probes originally, but we have trimmed this data to 289,509 probes for demonstration purposes. Each of these three data sets takes up to 4.3 gigabytes of storage even after trimming. Unless carefully managed, this size of data may cause serious issues in memory allocation, hence reading and modifying data entries, especially when there are overlaps among the variable groups (see **Methods**).

Here we describe each data set in more detail first. The first case study explores a multivariable Cox regression analysis using somatic mutations as predictor of cancer death risk. Since somatic mutations are sparse and not reproducibly detected at predetermined loci in early tumors, statistical analysis associating “locus-level” somatic mutation data and clinical endpoint such as patient survival remains challenging. Alternatively, counts or rates of somatic mutations can be aggregated on predefined polymorphic regions or individual genes. ^14, 15^ However, such “gene-level” analysis fails to retain the resolution required for investigating the potential functional impact of mutations (**Figure 1A**). To address this issue, we have recently proposed a functional region-based association testing approach for exome sequencing data, called Gene-to-Protein-to-Disease (GPD).^16^ GPD separately counts mutations for coding regions pertaining to protein domains and 11 amino acid-long windows surrounding post-translational modification (PTM) sites, and performs univariate statistical analyses with a clinical endpoint. **Figure 1B** illustrates how GPD summarizes mutation counts per protein sequence segments of three different types. These newly organized data can be used as covariates in the regression model.

**Figure 1.**
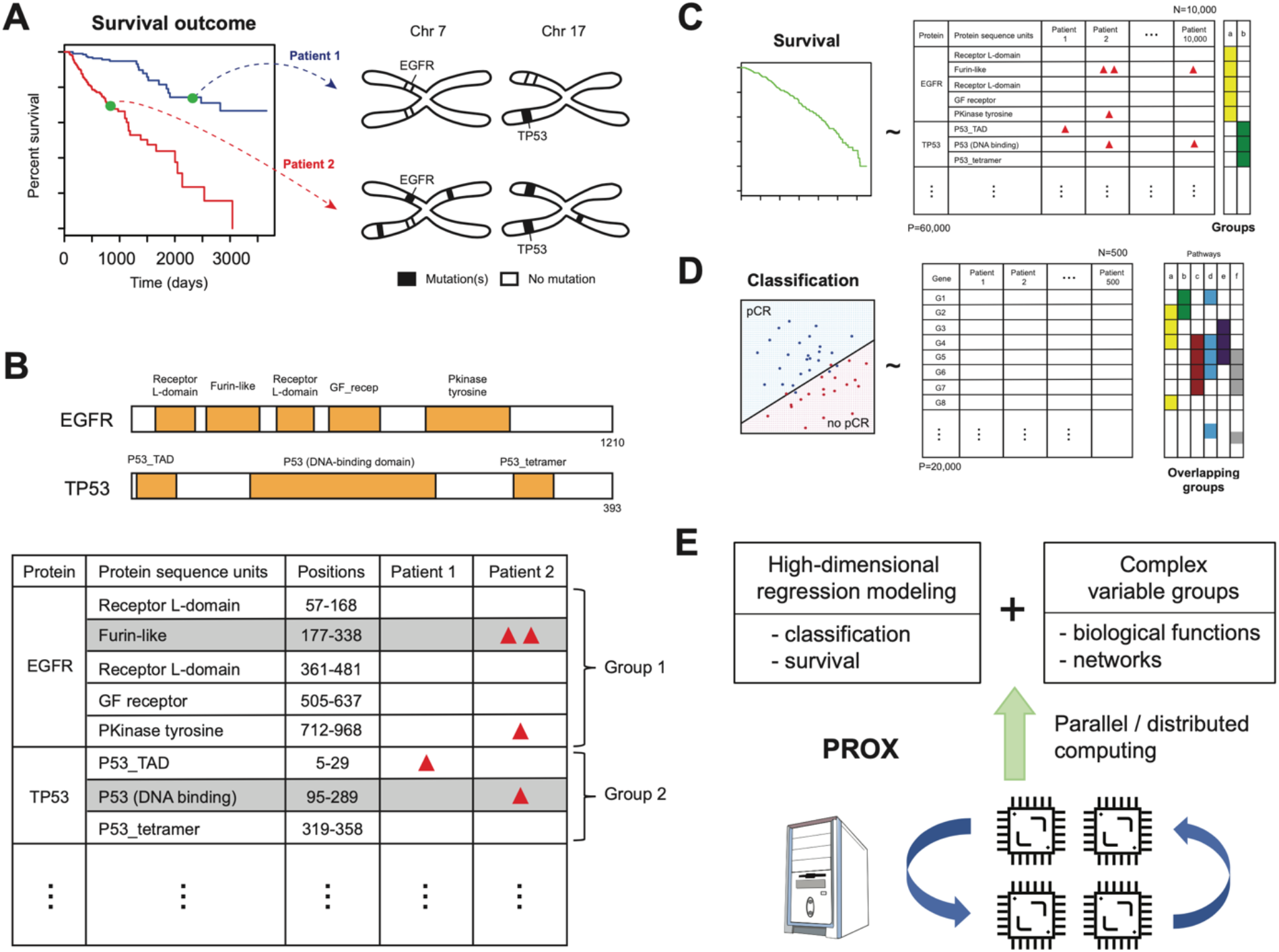
**(A)** Diagram of a hypothetical survival analysis with mutation data in EGFR and TP53 genes, two loci harboring somatic mutations with high frequencies in cancer genomes. Two people with different somatic mutation profiles may have two different cancer death risk. **(B)** The mutation counting method in the Gene-to-Protein-to-Disease (GPD) framework. Mutations are counted by sequence regions encoding functional units of proteins such as protein domains. GPD mapping therefore produces sparse count data with inherent variable group structure, with genes serving as variables groups. **(C-D)** ParProx accommodates high-dimensional Cox regression as well as multi-group logistic regression for classification, with overlapping or non-overlapping group lasso penalties specified by the user. **(E)** ParProx fits overlapping group lasso regression models on large-scale data sets through distributed or parallel computing, if necessary, and handles overlapping variable groups through the proximal gradient optimization algorithm.

Using this framework, we summarized the entire somatic mutation data into a count data set with 9,707 patients and 55,961 protein sequence segments across the human exome. We then fit a pan-cancer Cox regression model with 55,961 variables and group lasso penalties for overall survival of the 9,707 patients using ParProx (**Figure 1C**). Here, all sequence segments in one gene are considered to form a variable group, representing the hypothesis that individual mutations accrued in different protein sequences may have independent functional impact, but those mutations in the same gene are likely to be associated with disease risk of the given individual subject collectively than individually. We also attempted the same regression analysis with group penalties reflecting simultaneous mutations on sequence segments in physically interacting proteins, which produced an interesting outcome that led to selection of no predictor variables. This is discussed in more detail below.

In the second example, we demonstrate that ParProx produces a biologically interpretable logistic regression model of genes associated with pCR to neoadjuvant chemotherapy, a binary outcome determined by expert pathologists (**Figure 1D**). In this conventional “small *n*, large *p*” data example, we use biological pathways and GO terms as variable groups with arbitrary degree of overlap and nesting,^17, 18^ and show that the group lasso regression model optimized by ParProx identifies a sparse prognostic gene signature enriched with specific biological processes, rendering the prognostic model high interpretability over other similarly performing alternatives.

In the third example, we demonstrate ParProx in the context of analyzing data sets with a *p* so large that parallel computing is required to fit a regression model. We fit a Cox regression model with overlapping group lasso penalties on a DNA methylation data set from the liver hepatocellular carcinoma of TCGA. The DNA methylation array platform has probes representing genomic regions of high GC content, and as such, the dimensionality is much higher than other omics data sets where the measurements are often summarized to individual gene level, e.g. gene expression or DNA copy number data. We show that ParProx can perform regularized Cox regression with penalties jointly applied to probes associated with different segments of regulatory and coding regions for individual genes, where there are close to 90,099 variable groups formed over 289,509 variables. We show that the analysis can be completed within a reasonable amount of time using a single GPU, whereas another software package for overlapping group lasso regression analysis could not handle the size of the data.

Through these case studies, we show that ParProx provides a robust and computationally efficient implementation to fit regularized regression models, with the ability to find optimal, interpretable models under the constraint of highly complex group penalties with arbitrary degrees of overlap. This capability opens the door to incorporating prior knowledge on the relationship among predictor variables in the regression modeling (**Figure 1E**). Another important innovation of this implementation is that the model fitting process can be parallelized through distributed or parallel computing environments, if necessary, in case exorbitantly large data need to be analyzed, as illustrated by the third application data set.

### Proximal gradient optimization for logistic and Cox regression models

Before the demonstration, we first describe the computational workflow of ParProx in detail. We first describe this in the typical binary classification setting. The goal of the regression modeling is to understand the influence of *p* covariates *X* = (*X*^1^,…, *X^p^*) on the probability *Pr* (*Y* = 1 | *X*) of a subject belonging to class 1, where the two classes are labeled 0 and 1. In logistic regression, if there are *n* subjects, given the observed label *y_i_* and covariates *x_i_* = (*x_i_*^1^,…, *x_i_^p^*) for each individual *i*, the likelihood of the observed data is modeled based on the assumption that the log odds of the class membership is a linear combination of covariates, yielding the likelihood of the linear combination coefficient *β* = (*β*_1_,…, *β_p_*)

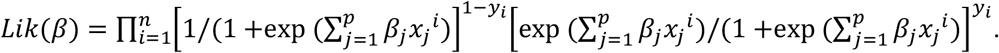

If the number of covariates *p* is large as in omics data, it is customary and also reasonable to assume that only a few independent variables mainly determine the response. This is promoted by multiplying a prior probability to the likelihood that causes all but a few coefficients among (*β*_1_,…, *β_p_*) to be zero, a process known as *regularization*. This prior typically takes the form of *π*(*β*) ∝ *exp* (–*λ*║*β*║), where ║*β*║ is some norm of the coefficient vector *β*. By taking the logarithm, the model fitting procedure amounts to an optimization problem of minimizing

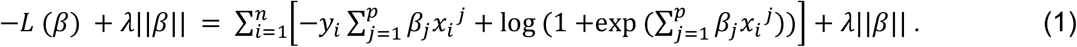

The regularization parameter *λ* is typically selected via cross validation.

In a typical survival analysis setting, the goal is to understand the influence of *p* covariates *X* = (*X*^1^,…, *X^p^*) to the survival probability *S*(*y* | *X*) = *Pr* (*Y* > *y* | *X*) that the survival time *Y* of a subject is longer than time *y*. In the Cox proportional hazards model,^19, 20^ given the *i*-th subject, *i* = 1,…, *n*, with covariates *x*^1^ = (*x_i_*^1^,…, *x_i_^p^*), whose time-to-death *t_i_* or right-censoring time *c_i_* is measured so that the observed survival time is *y_i_* = min(*t_i_, c_i_*), the survival probability is equivalently modeled through the hazard function *h*(*y_i_* | *x_i_*) = −*S′*(*y_i_* | *x_i_*)/*S*(*y_i_* | *x_i_*), where *S’* is the derivative of the survival function *S*:

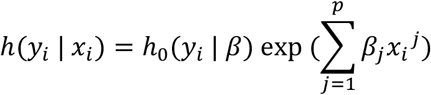

where *h*_0_(*y* | *β*) is the unspecified baseline hazard function. That is, the covariates affect the hazard multiplicatively in such a way that a linear combination of covariates determines the strength of the multiplication. The coefficient of linear combination is denoted by *β* = (*β*_1_,…, *β_p_*). Cox (1972) then proposes to get rid of the unknown baseline hazard in the fitting procedure by maximizing the partial likelihood

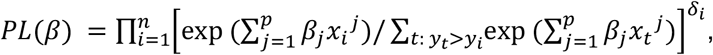

where *δ_i_* is the indicator that is 1 if *t_i_* ≤ *c_i_*, i.e., the death of subject *i* is observed, and 0 otherwise. Like logistic regression, if the number of covariates *p* is large, a prior of the form *π*(*β*) ∝ *exp* (−*λ*║*β*║) is multiplied to the partial likelihood to promote a sparse model. The model fitting procedure then amounts to an optimization problem of minimizing

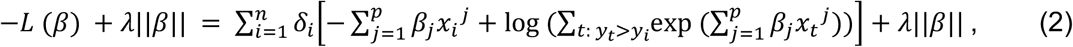

which takes a form of a generalized linear model, similarly to the logistic regression problem (1). The regularization parameter *λ* is typically selected via cross validation.

If this objective function (1) or (2) of the optimization problem were differentiable in *β*, then the typical gradient descent method, which iteratively updates the current estimate of *β* by moving slightly to the opposite direction of the vector of the first-order partial derivatives of the objective function, would eventually yield the correct estimate. Unfortunately, the objective functions (1) and (2) are not differentiable due to the presence of the norm ║*β*║. Nevertheless, the term *L*(*β*) in (1) and (2) is differentiable, hence an extension of the gradient descent method called the *proximal gradient* method can instead be applied. The proximal gradient for (1) or (2) consists of two steps^21^:

1. Compute the gradient ∇*L*(*β*^(*k*)^) of *L*(*β*) at the current estimate *β*^(*k*)^ is the coefficient vector *β*.
2. Update the estimate by the formula

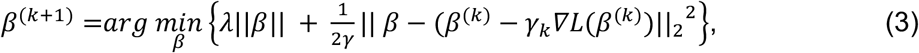

where the ║·║_2_^2^ in the right-hand side of equation (3) is the squared Euclidean norm, i.e., ║,*x*║_2_^2^ = (*x*^1^)^2^ +···+ (*x^p^*)^2^, and the argmin operator refers to the parameter value that minimizes the expression in the right-hand side. The scalar *γ_k_* is the step size that determines how far to move the estimate from the current candidate *β*^(*k*)^. The idea of the proximal gradient method is to approximate *L*(*β*) by a spherically shaped quadratic function tangential to *L*(*β*) at *β*^(*k*)^ and is above it for all other values of *β* and then minimize the approximate objective function. By iteratively doing so, the minimum of the original function (1) or (2) can be found (**Figure 2A**) even when the objective function is not differentiable at the optimal solution. For many choices of the norm ║*β*║, the right-hand side of the second step (3) takes a closed form expression despite its non-differentiability. (This includes the latent group lasso penalty chosen for this paper. See **Methods** below for detailed derivation.) Thus, the whole iterative procedure is almost as simple as the usual gradient descent method. From the point of view of computing, the combination of group lasso penalty and the proximal gradient method enables parallel computation (**Figure 2B**), since both the gradient ∇*L*(*β*) and the closed form expression of (3) can be computed independently for each latent group. Furthermore, within a variable group, each component can also be updated in parallel. An important implication of this doubly parallel feature of our approach in omics analysis is that the data set does not need to reside on a single storage -- *it can be split and stored distributedly in multiple computing devices*. Each device can report the update of the estimate of the regression coefficients in parallel to the master device holding the tally of the objective function, independent of the others. The size of the data to analyze scales linearly with the number of devices, with negligible sacrifice in computing time. This is in a stark contrast with the block coordinate descent method^22^ for optimizing the objectives (1) and (2), which is inherently sequential and requires the whole dataset to be stored in a single device. With the advance of modern high-performance computing infrastructure, such as the GPU, the ability of parallel and distributed computation has become a commodity, and running a parallel algorithm is getting increasingly easier -- ParProx is an instance.

**Figure 2.**
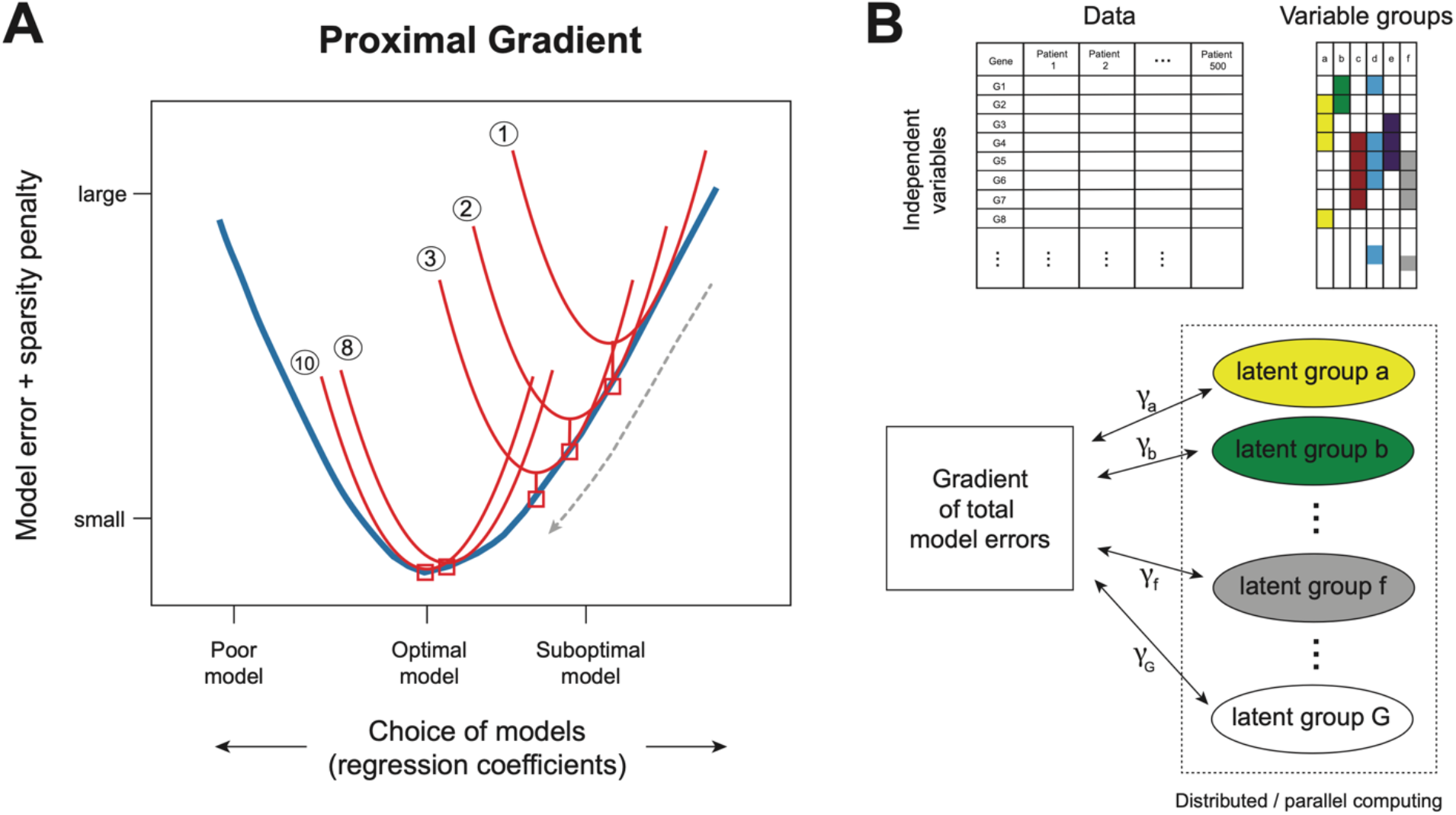
**(A)** Illustration of the proximal gradient optimization method. Thick blue curve: objective function for model selection, which is to be minimized to select the optimal model. Red curves (1 through 10) are surrogate functions of the objective function (blue). Red curve #1 touches the blue curve at the initial point representing a suboptimal model, and it is above the blue curve elsewhere. Each red curve is easy to minimize. At the rightmost red square point, red curve #1 is minimized. Then red curve #2 tangential to the blue curve at this point is constructed and minimized. This process is repeated until the minimizer of a red curve coincides with that of the blue curve (curve #10). The blue curve does not have to be differentiable. **(B)** Parallel computation of the proximal gradient optimization with group penalties. Top: for each group of gene, its latent membership is sought. Each group of genes may belong to multiple latent groups, and each latent group can be assigned to multiple gene groups. Bottom: regression coefficients for the latent groups are distributed in several compute devices and are updated in parallel since the latent groups are independent of each other. Computing the gradient of the objective function requires to gather coefficients from all other devices. Once computed, the gradient information is distributed to all devices to update regression coefficients.

### Pan-cancer survival analysis of somatic mutations using group lasso Cox regression with non-overlapping and overlapping group penalties

We first demonstrate ParProx through survival analysis of somatic exome mutation data in TCGA pan-cancer cohort. As mentioned in the overview, we have mapped all somatic mutations curated by the Pan-Cancer consortium of TCGA to human protein sequence segments (see **Methods** for details). These include (i) protein information units (PIUs) including 26,115 unique protein domains and 1,337 segments surrounding PTM sites, (ii) unannotated regions called linker units (LUs), and non-coding regions (NCUs). Of 10,793 patients, 9,707 tumors had at least one somatic mutation across 55,961 sequence segments. This 9,707 x 55,961 count data matrix, which requires 4.3 gigabytes of memory as double-precision floating-point numbers, was used as the covariates to a Cox proportional hazard regression of all-cause mortality. We set individual proteins as variable groups for regularization (18,250 genes) in the present analysis, but this group structure can be extended with overlap within ParProx, such as genes in the same biological processes, biochemical pathways, or protein complexes as we demonstrate later. In addition to the mutation counts, we have adjusted the model for age at diagnosis, gender, and cancer type without regularization on these coefficients, as they are known cancer death risk factors and the overall survival rates vary widely across different cancers.

The 10-fold cross-validation for the optimization of the regularization parameter took 71 minutes, and the final fit with the optimal regularization parameter took 23 minutes on an Nvidia Titan V. The GPU experiments were run on a workstation with two 2.20GHz 10-core Intel Xeon Silver 4114 CPU with 192GB memory, with four Nvidia Titan V GPUs with 8GB memory each attached. A detailed manual for ParProx analysis of this data set, using CPU and GPU, can be found in the software manual provided as **Supplementary Information**. A similar non-overlapping group lasso regression model could be fitted using the grpreg R package,^3^ and the analysis took 2.8 minutes for solution path calculation with 100 values of the regularization parameter *λ* and 28 minutes for 10-fold cross-validation (iMac desktop with 3.7 GHz 6-core Intel Core i5 processor and 32 GB 2667 MHz DDR4 memory). With non-overlapping group penalties and modest data sizes, this faster speed of grpreg package over ParProx is expected – the coordinate descent algorithm is exceptionally efficient as long as the data can be held by the same device.

Nevertheless, the selected model by the grpreg was counterintuitive in several aspects. First, the selected model included a very small number of variables (**Supplementary Table 1**), with almost all coefficients of mutation harboring segments being negative. Second, the coefficients for cancer type, which adjust for varying relative risk of death in different cancers, were all negative except leukemia (LAML), although there are other cancers that are just as lethal as the baseline cancer (GBM). Third, the selected model excluded the most well-known cancer death-associated protein domains on TP53 and EGFR genes. Put all together, we suspect that this aberrant result may have to do with the default data transformation step (orthonormalization, see **Methods**).

**Figure 3** shows the covariates selected by group lasso regression of ParProx, adjusting the effect of each sequence segment for all others on the death risk. The visualization in **Figures 3A** and **3B** was limited to the selected variables representing protein domains or PTM sites, of absolute values greater than 0.01, and with mutations called in patients of at least 10 different cancer types (see **Supplementary Table 1** for full lists). The group lasso regression selected 2,370 variables (sequence segments) with non-zero coefficients (1131 PIUs, 492 LUs, 747 NCUs). Not surprisingly, P53 domain on TP53 gene, the most commonly mutated protein domain across 34.3% of all tumors (3358 tumors of 31 different types in the pan-cancer cohort), was determined to have the largest deleterious effect on the cancer death risk, adjusting for other somatic mutation events across the genome. Two protein domains on EGFR, namely Furin-like and GF recep IV domains on EGFR, also had comparably large positive coefficients (deleterious), although somatic mutations on these domains of EGFR were observed in specific cancers with much lower frequencies (14 of 33 types). The mutations enriched in the tyrosine protein kinase domain of the BRAF gene, after adjusting for other factors, had protective effect on the death risk.

**Figure 3.**
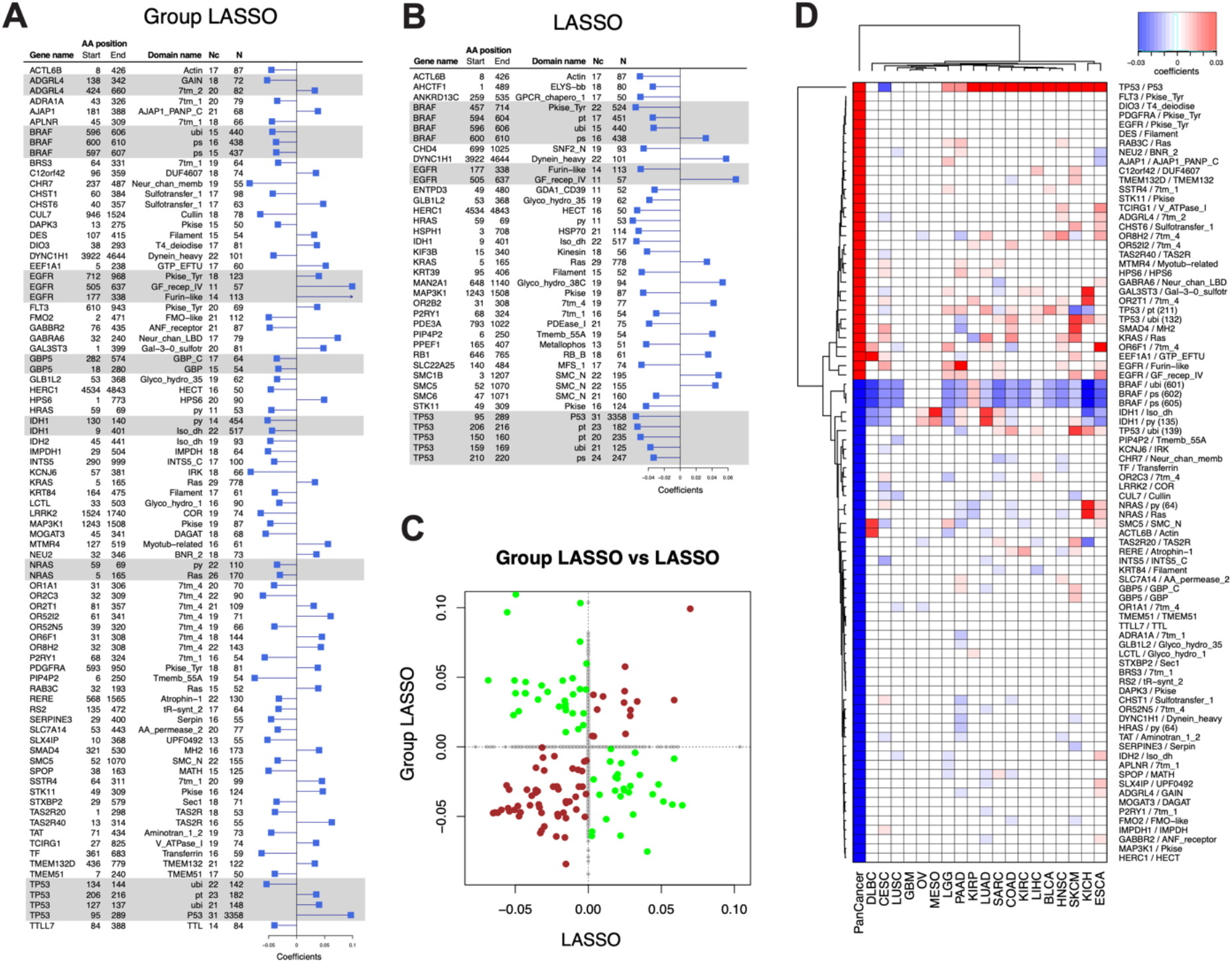
**(A)** Cox regression coefficients from the model with non-overlapping group lasso penalty (with proteins as groups). Sequence segments from the same genes, jointly selected by regularization, are highlighted in gray boxes. The columns on the left-hand side of the barplot show gene identifiers, start and end position on respective protein sequences, protein domain or modification site information, the number of cancers with at least one patient with somatic mutations in the sequence segments, and the total number of patients with mutations in the segment, respectively. The segments are shown in the figure if mutations were detected in at least ten cancers and regression coefficients are >0.03 in absolute value. **(B)** Receiver operating characteristic curves of lasso, group lasso, and RF, and the sensitivity and specificity of SVM, all evaluated using the two test data sets. All four methods perform similarly in the classification of pCR and residual disease. **(C)** Comparison of Cox regression coefficients between the two models. Brown and green dots show sequence segments with selected coefficients with consistent and inconsistent signs, respectively. Gray dots are the sequence segments selected in one of the two models only. **(D)** Comparison of Cox regression coefficients in the group LASSO Cox regression in pan-cancer analysis as well as individual cancer analysis.

We next tested whether ParProx can handle a more complex group penalty structure with overlap. In the analysis above, the membership of sequence segments (PIU, LU, NCU, and PTM site windows) to genes did not have any overlap. This time around, we gathered high confidence protein-protein physical interactions from two representative databases (see **Methods**) and used the membership of sequence segments to pairs of interacting proteins as the variable groups. This analysis tests the hypothesis that simultaneous mutations on two physically binding proteins in the same individual are likely to impact protein functions and thus such simultaneous mutation events have a greater deleterious or protective impact on cancer death risk. The mapping from our data translated into a total of 197,259 overlapping variable groups, many of them sharing the same sequence units (independent variables). In other words, each sequence segment of a protein-coding gene may belong to two or more groups if the protein has multiple interaction partners.

Using a Nvidia Titan V, the analysis took a total of 167 minutes with 10-fold crossvalidation. A similar Cox group lasso analysis could not be performed by grpregOverlap R package,^22^ an add-on to the grpreg package for handling overlapping group penalties, on the iMac desktop computer (“vector memory exhausted” error). Furthermore, the optimal regularization parameter selected by 10-fold cross-validation led to a Cox regression model with no sequence segments. In other words, we did not identify a signature of simultaneous mutations occurring on physically interacting proteins associated with survival, when adjusted for one another. This result can be interpreted in two different ways. It is possible that cancer death risk-associated somatic mutations do not necessarily co-localize in genes encoding protein complex members, especially in the somatic mutation data of early primary tumors at the time of diagnosis. In fact, when we examined co-mutation events in patients with death within five years of followup, only 126 pair of interacting proteins had simultaneous mutations in more than ten such patients. Further restricting to the patients who were deceased within two years, we had only 71 protein interactions with simultaneous mutations (**Supplementary Table 2**).

Alternatively, it is also possible that tumors collected from early diagnosis have a very low probability of harboring functionally consequential mutations on two or more essential members of a protein complex, and such events would not have been observed frequently in the early primary tumor collection of TCGA in the first place. Indeed, when we compared the number of interacting protein pairs with simultaneous mutations across the patients, we observed that only 8,682 out of 133,146 total interaction pairs (6.5%) bear more frequent simultaneous mutations than expected by random co-mutation on any pairs of proteins (more than 10 subjects). (**Supplementary Table 2 & Method**). In either case, an important observation here is that the variable group information for regularized regression representative of a specific biological hypothesis makes a difference in the final model selection, showcasing the importance in informing the variable selection procedure with appropriate biological prior. Furthermore, with complex group penalties, ParProx was able to handle the optimization problem that was not solved by a package built in R, the commonly used statistical analysis environment.

To benchmark the group lasso model, we also ran the same regression with Cox lasso regression, i.e., with *L*_1_ penalty on individual sequence segments but no group-wise regularization ^23, 24^ (**Supplementary Table 1**). To our surprise, the two analyses selected substantially distinct prognostic models (**Figure 3C**). First, the group lasso model selected 861 sequence segments as prognostic signature of cancer death risk, whereas the lasso model selected 288 sequence segments. The deflation in the number of selected sequence segments was expected since group lasso would maintain a sequence segment as predictor as long as there is another sequence segment of prognostic signal in the same protein. Second, the cancer type difference in the cancer death risk was highly inconsistent with the frequency of multi-year death events in the lasso model. For instance, LAML is one of the cancers with the lowest survival rate in TCGA, yet the coefficient representing the difference in mutation-independent death risk between LAML and GBM (baseline cancer in this analysis) was negative, potentially suggesting that the death rate differences were over-adjusted by mutation data in the model fitting. BRCA is a cancer with overall 95% survival rate at five years of follow-up, yet the coefficient for the difference between BRCA and GBM was positive, again producing coefficients with signs inconsistent with the mortality rate differences between the two cancers. Therefore, not accounting for known structure amongst the covariates, i.e. the lasso model, seems to have considerable impact on the interpretability of the final model.

Based on the same selection criteria, the top prognostic sequence segments from the lasso model are shown in **Figure 3B**. It is striking that the four PTM sites (serine/threonine phosphorylation and lysine ubiquitination at respective sites) and P53 domains all have negative coefficients, contradicting the common notion that the somatic mutations in TP53 have a deleterious effect on cancer death risk. The same contradictory result was also observed in the EGFR gene, where lasso regression selected GF recep IV domain and the second Furin-like domain with positive and negative coefficients, respectively. These are undoubtedly counterintuitive results: mutations on P53 have been implicated in cancer incidence risk as well as prognostic outcome in multiple cancers,^25^ whereas somatic mutations in EGFR had a deleterious effect on cancer death risk in cancer type-specific regression analyses for the two cancers (GBM and LUAD, see **Supplementary Table 2**). By contrast, all protein domains in EGFR had positive regression coefficients in the group lasso model fit using ParProx (**Supplementary Table 1**).

We next examined the regression coefficients from the pan-cancer analysis with those models fit on individual cancer data separately. **Figure 3D** shows the heatmap of regularized coefficients obtained from the pan-cancer analysis as well as those from analyses of individual cancer data. As expected, the sign of the coefficients was highly congruent among different analyses, although there were a few exceptions. Hence, we conclude that the pan-cancer survival analysis by the group lasso regression of ParProx successfully pools shared effects of mutations on the risk of cancer death in this data.

### Prognostic gene expression signature of pathological complete response using group lasso regression with overlapping groups

In the next case study, we demonstrate ParProx through a classification problem. Re-analyzing the meta-analysis data of Prat *et al*.,^13^ we aim to identify an mRNA gene expression signature to classify breast cancer patients undergoing chemotherapy with anthracycline and neoadjuvant agents into two groups, i.e., pathological complete response (pCR) and residual disease (RD). Here we use gene expression data sets of 12,307 genes and 469 patients in the training data set^11^ and two test data sets^10, 12^ (*N*=115 and *N*=244), and we use the pathways and GO terms as variable group information in the logistic group lasso regression. The analysis workflow is visually represented in **Figure 4A**.

**Figure 4.**
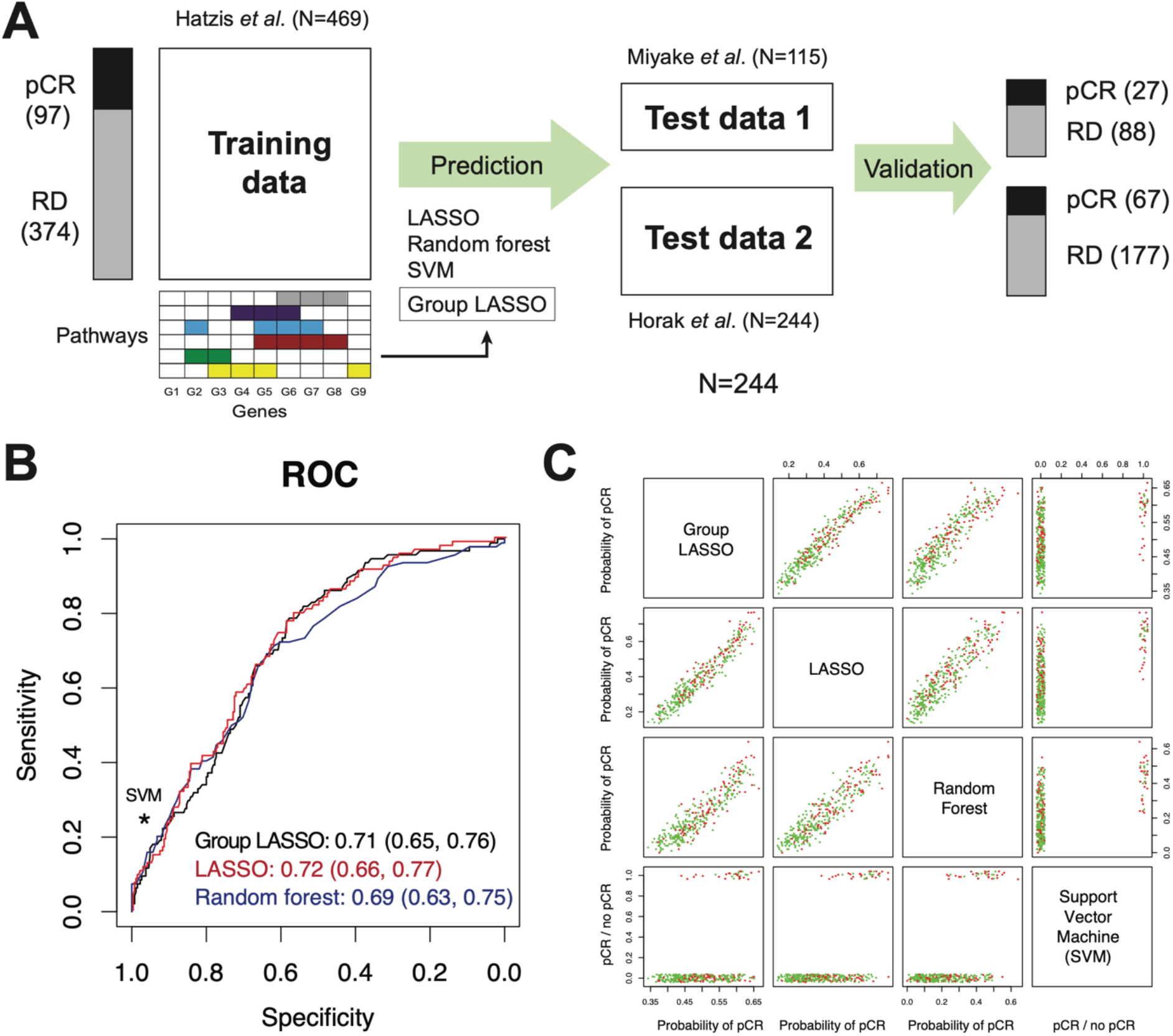
**(A)** Gene expression-based classification analysis of pathological complete response (pCR) in breast cancer. Lasso regression, random forest (RF), and support vector machine (SVM) with radial basis function were used for benchmarking of pathway-based group lasso regression model. **(B)** Receiver operating characteristic curves of lasso, group lasso, and RF, and the sensitivity and specificity of SVM, all evaluated using the two test data sets. All four methods perform similarly in the classification of pCR and residual disease. **(C)** Class probabilities for the samples in the two test data sets reported by the four methods show highly similar results.

Before fitting the regression model, we first carried out the classical univariate analysis by gene-wise hypothesis testing in the training data by Hatzis et al (*N*=469) (see **Methods** for selection criteria). The analysis found as many as 1,825 genes over-expressed and 1,524 genes under-expressed in tumors from patients who achieved pCR compared to those with residual disease (**Supplementary Table 3**). The genes over-expressed in pCR patients showed clear enrichment for biological processes related to cell cycle, DNA repair, cell proliferation and protein folding, whereas the genes under-expressed in pCR patients showed enrichment for less essential pathways such as cilium assembly and extra cellular matrix (ECM) organization. This ‘routine’ analysis via hypothesis testing suggests that the tumors responding to the neoadjuvant agents with pCR have gene expression profiles favoring cell proliferation, while the tumors not achieving pCR with RD do not.

We next built classifiers of pCR using four different methods: logistic regression with lasso penalty, logistic regression with pathway-level overlapping group penalty (ParProx), random forest, and support vector machine (SVM). In ParProx analysis, we used external data resource that combined multiple pathway databases to define variable groups, resulting in 11,734 groups among the 12,307 variables (including those singletons that do not belong to any pathway or GO term).^18^ With smaller size of the data set (12,307 by 469), the analysis was performed within a reasonable amount of time with ParProx on a GPU (7 min for cross-validation, 19 min for final model fitting). A similar analysis could be performed using the grpregOverlap package in R (28 min for cross-validation, 28 min for entire solution path calculation). As shown in **Figure 4A**, we trained the classifiers in the training data by Hatzis *et al*., and made predictions of pCR on the two test data sets. When we compared the area under the curve of the receiver operating characteristic (ROC), the first three methods performed as well as one another (**Figure 4B**), and the predictions from the SVM with radial basis kernel, with cost and gamma parameters optimized through 10-fold cross-validation within the training data, did not perform better than the three methods (scores shown in **Figure 4C**).

Given the highly similar performance metrics across different methods, we next investigated the interpretability of the gene expression signatures. Since the two machine learning methods with greater complexity (RF and SVM) utilize all features in the respective classifiers, we did not pursue interpreting the underlying predictor. Instead, we compared the selected genes between the two logistic regression models with and without group penalties. Logistic regression with lasso penalty selected a total of 289 genes in the predictive signature (182 with positive and 107 negative coefficients, **Supplementary Table 4**). Subsequent pathway enrichment analysis showed that the genes with positive regression coefficients, those contributing to the better chance of pCR, had mild enrichment of mitotic cell cycle and DNA replication genes, whereas the genes with negative coefficients were not particularly enriched in any known pathways other than ECM organization.

By contrast, ParProx analysis incorporating the pathway membership of genes selected a total of 829 genes (481 positive and 348 negative), a larger panel of genes than lasso logistic model above. As stated in the previous case study, this is an expected consequence of using the group penalty, which tends to select genes in the same pathway together if there is a true effect of pathway-wide gene regulation. A clear advantage of the group lasso penalty is that one can rank pathways based on the number of genes with non-zero coefficients (**Figure 5A**). We selected five GO terms and one KEGG pathways with the largest number of genes with non-zero coefficients and large magnitudes in the sum of coefficients, with all six related to one overarching theme and sharing many common genes -- DNA replication during mitotic cell cycle (**Supplementary Table 3**).

**Figure 5.**
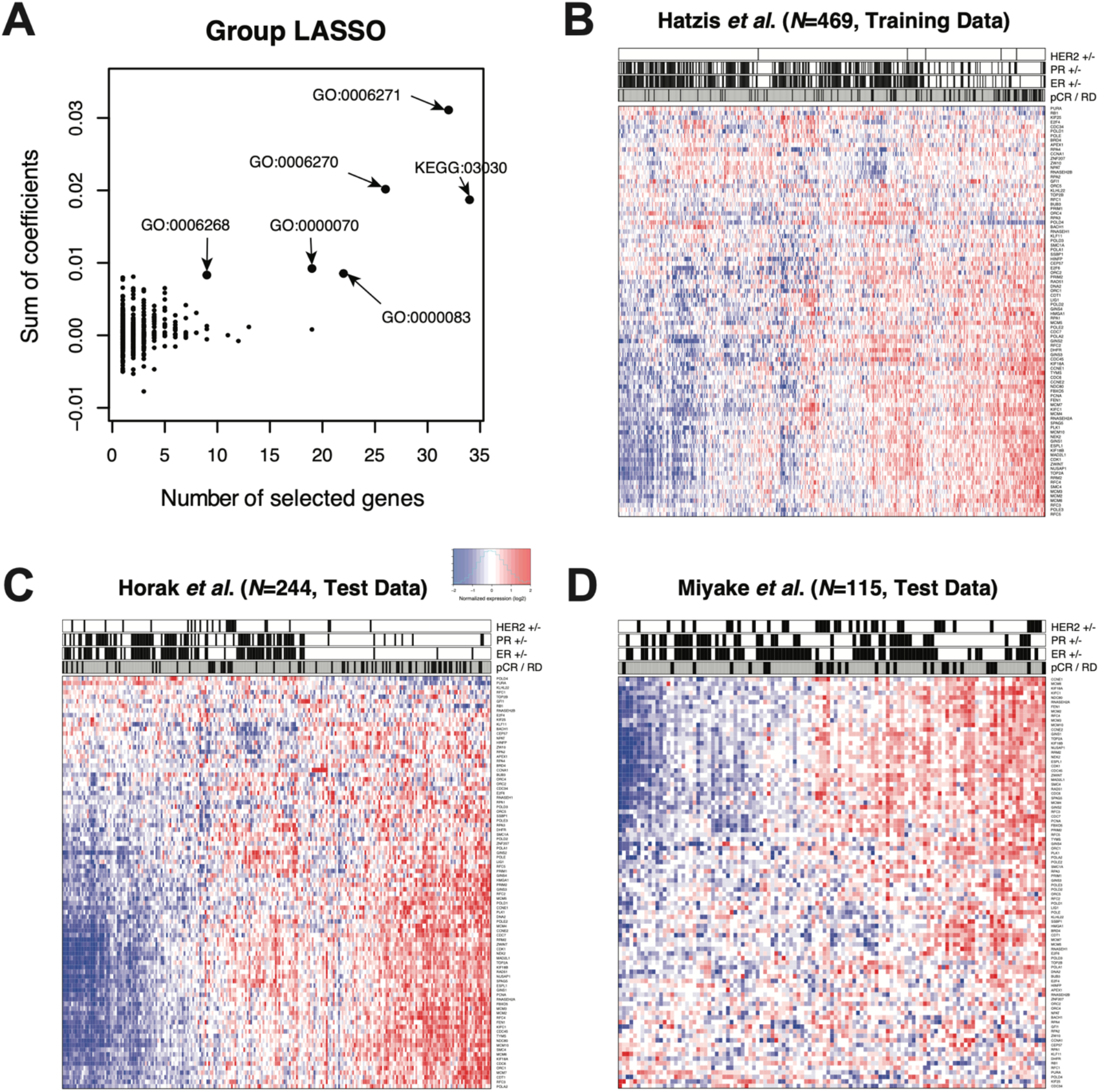
**(A)** Groups (pathways) containing at least one member gene selected with non-zero coefficients in the group lasso logistic regression. The Gene Ontology terms with 9 or more genes with sum of coefficients above 0.05, with positive contribution to the probability of pCR, are shown with arrows. **(B-D)** Heatmaps of the 90 member genes of the selected pathways in the training and test data sets. Each heatmap is annotated in terms of ER, PR, and HER2 status, as well as pathologist-graded pCR status (outcome).

**Figure 5B** shows the gene expression data with each gene normalized by mean expression value, along with immunohistochemistry results of estrogen and progesterone receptors (ER and PR), fluorescence in situ hybridization analysis of HER2, and pCR status. In this training data, it is clear that the tumors with pCR were mostly triple negative tumors (ER-, PR-, and HER2-) as Prat *et al*. initially observed,^13^ indicating that the gene signature obtained by ParProx for a high chance of pCR is negatively associated with the positive hormone receptor status, hence positively associated with the canonical pattern of gene expression regulation for DNA replication and cell cycle progression in the triple negative tumors. Consistent with this, we observed that the gene signature in the test data set showed better concordance with the pCR status of patients in Horak *et al*. with at least half the patients classified under basal-like cancer with majority being triple negative (**Figure 5C**) than the status of patients in Miyake *et al*., where the majority of the patients were ER positive (**Figure 5D**).

In summary, this data example represents a case of logistic regression classifier with high-dimensional feature data with a modest sample size, with the model fitted under a large number of overlapping variable groups. ParProx successfully optimized the objective functions under the constraint of overlapping, complex variable group information. We were able to conduct a similar analysis using an existing R package (grpregOverlap) in comparable computation time but ParProx has the potential to reduce the computation time even further when hardware for parallel or distributed computing are available. In this data, the classification performance is similar to the logistic lasso regression, as well as other machine learning methods including random forest (RF) and support vector machine (SVM) in this data. Among these similarly informative models, however, the lasso logistic regression selected a gene signature devoid of enrichment of particular biological functions associated with the clinical endpoint (pCR), and the two aforementioned machine learning algorithms do not yield interpretation of predictive data features as the regression models do. By contrast, the overlapping group lasso model fitted using ParProx yielded a gene signature indicating improved chance of pCR in patients with DNA repair and cell proliferation genes over-expressed in their tumors.

### Prognostic signature of DNA methylation in liver cancer data – hign-dimensional data

In the third data, we fitted a Cox regression model with overall survival as outcome variable and DNA methylation probes located in distinct genomic positions relative to protein coding genes as predictor variables in a liver cancer data. In this data, even after selecting the probes located near protein coding genes only, the number of data features (*p*) is 289,508, with sample size of *N* = 428. In addition, we treated 90,099 genomic regions representing unique relative positions of probes as variable groups, including TSS1500, TSS200, 5’ UTR, 1^st^ exon, gene body, and 3’ UTR as annotated by the microarray vendor (see **Methods**). We thus guide the Cox regression model fitting to jointly penalize the probes in these genomic segments.

Using the overall survival as clinical endpoint, we first attempted to fit overlapping group Cox regression using grpregOverlap in R, the software was unable to perform model fitting and produce memory allocation errors in multiple desktop computers with at least 8GB RAM and 3.4 GHz quad-core intel i5 CPUs or better. By contrast, ParProx was able to perform the C index-based search of optimal λ value and the final model fit in 458 minutes with parallel computation using a Nvidia Titan V.

ParProx reported an overlapping group lasso Cox model with 393 methylation probes located upstream and along the coding regions of 272 genes (see **Figure 6** and **Supplementary Table 5** for data and regression coefficients, respectively). As in the previous two examples, the group lasso regression model produced an immediately interpretable model. The biological processes enriched in the genes close to the selected CpG island probes included response to stress, negative regulation of transcription from RNA polymerase II, apoptotic process, cell redox homeostasis, and small molecule metabolic process. The model suggests that genes involved in oxidative stress response, metabolism, and gene expression regulation are modulated by DNA methylation differently between patients with longer survival and those with shorter survival.

**Figure 6.**
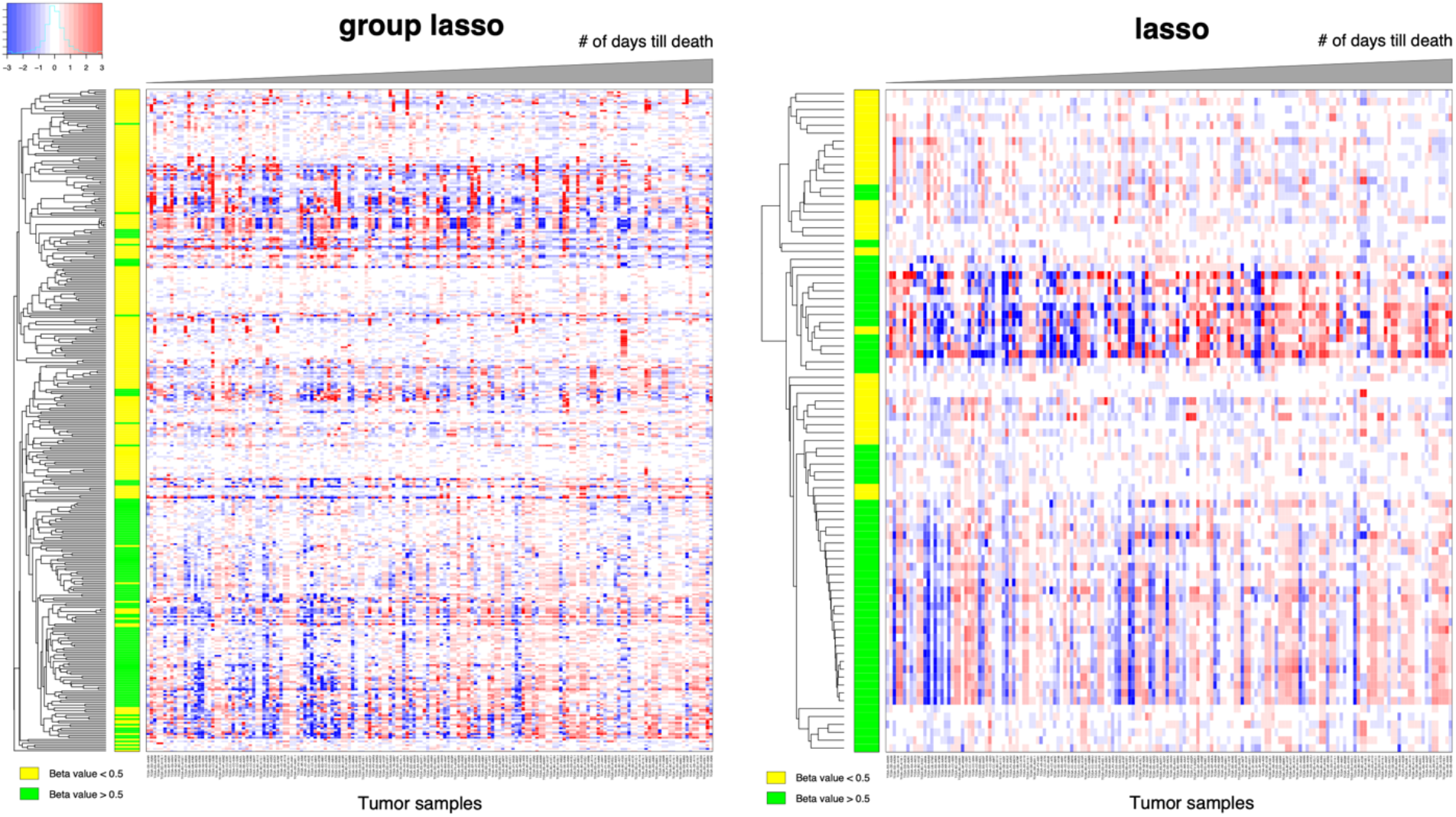
Heatmap of the selected DNA methylation probes in Cox regression models with overlapping group lasso penalty (ParProx) and lasso penalty, showing the samples from the deceased subjects only. Yellow and green color bars indicate that the beta values (percentage of methylation) are above and below 50%, respectively. The tumor samples were ordered in increasing order of the number of days of survival of the donors.

We have benchmarked the model against a Cox regression model with plain lasso penalty, which selected 114 CpG island probes. To our surprise, the probes selected by Cox regression with lasso penalty had poor overlap with the probes selected in the group lasso model, sharing only 16 common probes. In addition, C-index was comparable between the two models: 0.65 (± 0.03) with the group lasso penalty and 0.63 (± 0.03) with the lasso penalty. Similar to the second example above, this result is likely due to the fact that there are a large number of weakly predictive regression models with different predictor variable combinations with comparable degrees of association with overall survival. Among those options, group lasso chose the model that best represents the variable group structure we specified as the modeler. This observation also reaffirms that specification of variable group structure plays an influential role in the selection of data features associated with the clinical endpoint, and ParProx provides the interface to fit these models in ultrahigh-dimensional data sets that would otherwise have been impossible to fit.

## Discussion

In this work, we presented a scalable implementation to fit regression models for survival and classification analysis with structured group penalties representing biological prior information regarding the relationships among the independent variables. The proximal gradient optimization implemented in Julia language can distribute the iterative updates for parallel computation in the case of large-scale data sets, which is the major advance offered by ParProx. We demonstrated the robustness of the implementation in both ‘large *n*, large *p’* case (mutation data example) as well as ‘large *p*, small *n*’ case (gene expression data example) and showed that ParProx can deal with survival regression using a very large-scale data set (*p* = 289,508) using parallel computing with GPU.

In contrast to the conventional differential expression analyses via hypothesis testing, our one-shot regression analysis strategy describes the multivariate relationship between clinical endpoint and high-dimensional molecular data using linear models. Linear models are often thought to be too restrictive to describe complex relationships between genotype and phenotype. However, it has the clear advantage of interpretability of results and low variance of prediction results. In particular, linear models are able to summarize the overall impact of each variable onto the outcome into positive and negative values after accounting for the effect of others, and this directionality of sign is often important for biological interpretation of predictive models. Despite the increasing popularity of machine learning and deep learning methods in omics data analysis, the majority of these methods permitting non-linear classifiers can only tell the importance scores of individual variables, but fail to provide intuitive interpretation of the relationship between the outcome and the variables, as demonstrated in the breast cancer data as well. Within the class of linear models, ParProx provides an efficient solution to enable the challenging overlapping group lasso optimization with ease and it is highly scalable to large data sets.

Assessment of computation time between different implementations of linear models may be affected by the differences in algorithm, choice of grid coordinates for the regularization parameter, convergence criteria, to name a few. Convergence criteria and grid selection are detailed in the **Methods** section. Nonetheless, we emphasize that it is the choice of the algorithm that determines the scalability of the software. The proximal gradient method employed by ParProx is flexible in parallelization over distributed data, hence the computation time improves almost linearly with addition of hardware, e.g., GPU. This is the key contrast to grpreg and grpregOverlap packages implementing the coordinate descent method, which is an inherently sequential algorithm.

## Methods

### Optimization with structured sparsity

Both the logistic and proportional hazards models of the **Results** section admit a negative log-(partial) likelihood *L*(*β*) of the regression coefficient vector *β* = (*β*_1_, *β*_2_,…, *β_p_*) ∈ *R^p^* that is differentiable and convex in *β*, where *R^p^* means the set of *p* real numbers.^26^ Furthermore, the gradient ∇*L* satisfies the Lipschitz condition ║∇*L*(*β*) – ∇*L*(*β′*)║ ≤ *M*║*β* – *β′*║ for some positive constant *M*,^27^ where ║·║ is the Euclidean norm (root-mean-squares of the elements of the vector). In order to encode the prior knowledge, the latent group lasso penalty is defined as follows.^28^ Assume a collection *G* of groups of genes is given. That is, *g* ∈ *G* is a subset of all the gene indexes {1, 2,…, *p*}. Let |*g*| be the number of elements in *g*. Define a |*g*|-dimensional vector *γ_g_* (denoted by *γ_g_* ∈ *R*^|*g*|^) and the linear map *P_g_* that maps *γ_g_* to a p-dimensional vector *β_g_* ∈ *R^p^* in such a way that the elements with indexes in *g* are equal to those of *γ_g_* and all the other elements are zero. For example, if *g* = {2,3} and *γ_g_* = (−1,1), then *β_g_* = (0, −1, 1, 0,… 0) and *P_g_* is a *p×p* diagonal matrix with one on the 2nd and 3rd diagonal component with all the other components being zero. Then the desired penalty is

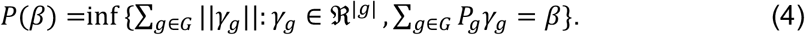

In other words, the regression coefficient *β* is decomposed as a sum of latent components *β_g_* = *P_g_γ_g_* and it is the norm of these components that is penalized (note ║*γ_g_*║ = ║*β_g_*║). In this way, overlaps between the groups are allowed. When there is no overlap, penalty (4) reduces to the classical group lasso penalty.^2^ Although the latter penalty may be straightforwardly defined for overlapping groups, it tends to select the complement of a union of groups -- if two groups share a gene but one group is not selected, then the coefficient for the shared gene must be zero and the other group is only partially selected. The penalty (4), on the other hand, promotes the opposite and this property is desired for pathway selection.

Estimation of the generalized linear model under the penalty (4) can be formulated as the following optimization problem

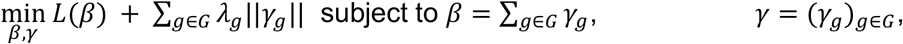

which, with an appropriate matrix *A* such that *β = Aγ*, can be equivalently written as an unconstrained optimization problem

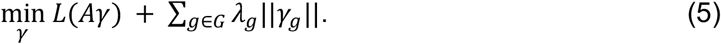

It can be shown that the second term ∑_*g*∈*G*_ *λ_g_*║*γ_g_*║ is a norm of the aggregated latent vector *γ* = (*γ_g_*)_*g*∈*G*_ and hence problem (5) has the same structure as problems (1) and (2) of the **Results** section. We then can apply the proximal gradient descent (PGD) algorithm of the **Results** section to solve (5) efficiently. For iteration *k* + 1, PGD updates *γ_g_* by the formula (3). Since there is no overlap between *γ_g_s* in *γ*, this update has a closed form:

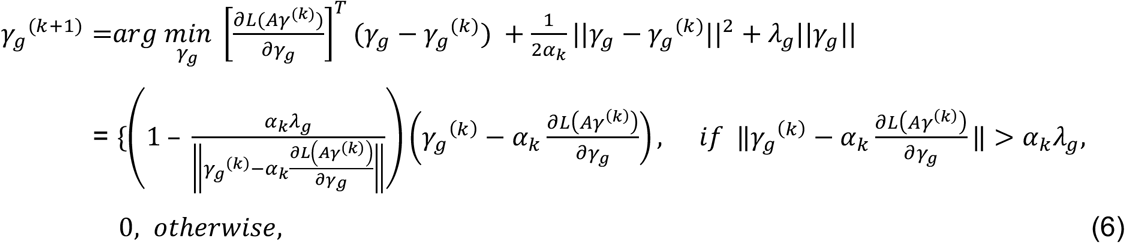

for all *g* ∈ *G*; *α_k_* is the step size between 0 and *2/M*. Most importantly, PGD allows a distributed update of latent groups since they are independent of each other, hence can be significantly accelerated by modern parallel computing devices such as a GPU or distributed computing environments such as clouds.

### Comparison with other software packages

The software package grpreg for the R statistical computing environment fits linear, logistic, and Cox regression models with non-overlapping group lasso penalties, hence solves problems (1) and (2). The package grpregOverlap extends grpreg to handle the latent group lasso penalty (4) to allow overlaps between groups, hence solves problem (5). The key differences between these packages and ParProx are three-fold: (a) solution algorithm, (b) memory management, and (c) standardization of variables. As for the algorithm, grpreg/grpregOverlap employs a (block) coordinate descent (BCD) method instead of proximal gradient of ParProx. BCD is a simple algorithm that updates a (latent) variable group at a time with the other groups held fixed - each group update has a closed form. Hence the complexity of each update is low. While inherently sequential, BCD is very efficient when the data size is moderate, as can be seen from the non-overlapping group-regularized analysis in the first case study. However, when there are overlaps in groups, BCD may expand the data size considerably, causing memory issues even if the original data size is modest. For instance, in the first case study, the somatic exome mutation data matrix is of size 9,707 x 55,961. With a latent group penalty in which there are 197,259 overlapping groups, the number of latent variables becomes 1,384,850. grpregOverlap creates a new effective data matrix of size 9,707 x 1,384,850 by duplicating the corresponding columns of the original data matrix in order to apply BCD, which requires more than 100 gigabytes of memory. On the other hand, ParProx evaluates the gradient of *L* by using the original data matrix and the linear map *A*, which is sparse and only has 0/1 entries. Hence, the additional memory requirement is small. For a detailed comparison of memory requirements in ParProx and grpreg/grpregOverlap, see **Table 1**. Recall that it is the independent nature of update (6) that allows the use of GPU acceleration and other parallel and distributed computing environments, which is not feasible for BCD. Even if the original data matrix does not fit into the memory of a single device, it can be distributed over multiple devices and coefficients of each group over multiple devices, and the coefficients can be updated simultaneously. Finally, grpreg/grpregOverlap standardizes variables by orthonormalizing the (latent) variables within the same group, while ParProx employs the common practice of standardizing each observed variable. Other minor differences between the two software implementations are as follows.

**Table 1.** Minimum memory requirement for each case study and variable grouping method. The number of variables may differ from the numbers in the main text due to inclusion of additional risk factors (covariates) from outside the respective omics data, e.g., age and gender.

|  | Group overlap | Sample size | No. variables | Latent dimension | Data size | grpreg/grp regOverlap | ParProx |
| --- | --- | --- | --- | --- | --- | --- | --- |
| Dataset 1 (somatic exome mutation) | No | 9707 | 55,963 |  | 4.346 GB | 4.346 GB | 4.346 GB |
|  | Yes | 9707 | 55,963 | 1,384,805 | 4.346 GB | 107.5 GB | 4.357 GB |
| Dataset 2 (pCR) | Yes | 469 | 12,309 | 127,083 | 46.18 MB | 476.8 MB | 47.20 MB |
| Dataset 3 (DNA methylation) | Yes | 428 | 289,508 | 370,473 | 991.3 MB | 1.268 GB | 994.2 MB |

#### Convergence criteria

In all of the examples, the PGD of ParProx was run until 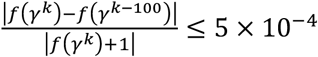 for cross-validation, and more stringently 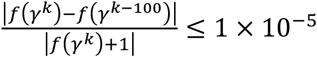 until for fitting the final model after the model selection. Here, *f*(*γ*) denotes the objective function of the optimization problem (5). For the BCD of grpreg/grpregOverlap, the default setting of the software was used, which stops the algorithm if 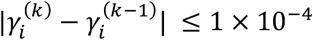.

#### Grid points for cross-validation

In ParProx, the regularization parameter *λ* was chosen among 31 equally log-spaced *λ* values between 10^-4^ and 10^-7^ in the first case study, among 31 equally log-spaced values between 10^-6^ and 10^-9^ in the second case study, and among 21 equally log-spaced values between 10^-6^ and 10^-8^ in the third case study. The grpreg and grpregOverlap packages automatically select 100 equally log-spaced values. The minimum is 0.05 if there are more variables than the number of samples, and 0.0001 otherwise. The maximum is chosen so that the model selects no variables with this *λ*.

#### Option for excluding variables from regularization

ParProx allows specification of variables excluded from regularization. This is a useful option in clinical omics data since certain variables represent known risk factors in a given disease context regardless of their statistical significance. grpreg R package allows this option, but grpregOverlap does not offer it as of version 2.2-0.

### TCGA pan-cancer somatic mutation data

TCGA pan-cancer somatic mutation data was downloaded from the Genomic Data Commons (MC3 Public MAF).^9^ In addition, we gathered 348,658 protein modification sites from PhosphoSitePlus^29^ and 45,607 domains, families and repeats for 19,076 genes from Pfam.^30^ Survival outcome data were downloaded from the TCGA Pan Cancer Clinical Data Resource,^8^ which contains curated clinical information for 10,793 patients. Among these, 367 non-primary skin cutaneous melanoma patients with metastatic tumors were excluded. Mapping somatic variants to protein units was described in our previous work.^16^ Protein information units (PIU) refer to the genomic regions encoding protein domains, or +/-5 amino acid-long windows around protein modification sites. Sequence regions between PIUs are defined as linker units (LU). The LUs not only include linker regions between domains but also cover unannotated, repeat or disordered regions. The regions outside the protein-coding sequences including untranslated regions, introns, and regulatory regions are collectively defined as noncoding units (NCU). NCUs are assigned to the closest gene in the genome. Aggregating somatic mutation mapped PIUs, LUs and NCUs from primary tumor samples categorized in 33 cancer types, we have 27,452 PIUs, 12,441 LUs and 16,068 NCUs, adding up to 55,961 units mapped by mutations from 9,707 individuals.

### Analysis of co-mutation frequency on protein interaction networks

The protein-protein interaction network has 133,146 unique pairs of interactions among 12,047 unique proteins. To estimate the significance of co-mutation frequency (the number of subjects having simultaneous mutation on both interacting proteins) of each pair of interacting proteins, we randomly sampled 133,146 pairs of interaction from the pool of proteins 1,000 times and calculated the co-mutation frequency for randomly sampled pairs in each iteration. The p-value of pair with co-mutation frequency F is defined as the number of pairs with co-mutation frequency higher than F divided by the total number of interactions (133,146), averaged across the 1000 iterations.

### Breast cancer neoadjuvant chemotherapy complete response microarray data sets

We downloaded gene expression microarray data sets with sample annotation information from the Gene Expression Omnibus database, based on the information from Prat *et al*.^13^: Hatzis *et al*. from GSE25006,^11^ Miyake *et al*. from GSE32646,^12^ and Horak *et al*. from GSE 41998.^10^ Each data set was normalized by equalizing the median and median absolute deviation of expression values across the samples. For regression analysis, we applied logarithmic transform (base 2) and substrate the mean from expression values in each gene. For univariate differential expression analysis, we performed two-sample *t*-test and computed *q*-values^31^ from the raw p-values to account for multiple testing.

### Interaction networks and gene pathways for variable group information

For variable group information in the TCGA somatic mutation data analysis with network penalty, we used protein-protein interaction network data from iRefIndex^32^ and BioPlex.^33^ For the group information used in the breast cancer data, we used a composite database of pathway databases called Consensus Pathway DataBase (CPDB)^18^ and Gene Ontology.^17^

### Liver cancer DNA methylation data

We downloaded the Illumina human methylation 450 array data set from Broad GDAC Firehose, corresponding to the liver hepatocellular carcinoma study (N=428, with duplicates from 377 patients).^34^ 52 patients had two replicates and we used each biopsy as an independent tumor sample for this illustration. We have selected 369,194 probes that are annotated to belong to one of the following regions of genes for the regression analysis: TSS1500, TSS200, 5’ UTR, 1^st^ exon, gene body, and 3’ UTR. Individual sequence regions were considered to be variable groups (e.g. A1BG_TSS1500, A1BG_TSS200, A1BG_1^st^ exon, A1BG_5’UTR, A1BG_body, A1BG_3’UTR are different variable groups for penalization). This resulted in a total of 90,099 groups, reflecting on average four methylation probes per group. In genomic regions with dense population of genes, the adjacent groups sometimes shared the same methylation probes, creating overlapping groups.

## Acknowledgements

This work was supported in part by grants from the National Research Foundation of Korea and Ministry of Science and ICT of Republic of Korea (No. 2019R1A2C1007126 to J.W.), Singapore Ministry of Education (MOE2018-T2-2-058 to H.C.), and National Medical Research Council of Singapore (NMRC-CG-M009 to H.C.).

## Author contributions

SK and JW implemented ParProx in Julia language. GL and HC prepared data sets and interpreted results in consultation with SK and JW. All authors contributed to manuscript writing.

## Software Availability

ParProx is freely available through GitHub repository at https://github.com/kose-y/ParProx.jl under the MIT license.

