## Supplementary Information for "Computationally scalable regression modeling for ultrahigh-dimensional omics data with ParProx"

### ParProx

This package aims to perform penalized logistic regression and Cox proportional hazard regression using the proximal gradient method. It supports $\ell_1$ penalty (lasso) and the group lasso penalty. The groups may overlap, as postulated by the latent group lasso (Obozinski,Jacob, and Vert, 2011 (https://arxiv.org/abs/1110.0413)). This package also supports computation on a single Nvidia GPU via the package CUDA.jl (https://juliagpu.gitlab.io/CUDA.jl/).

### Installation

#### Julia

Download Julia from https://julialang.org/downloads/ (https://julialang.org/downloads/) according to your OS and hardware, and follow platform-specific instruction here (https://julialang.org/downloads/platform/).

You can also run Julia from the Jupyter Notebook after installing the package IJulia (https://github.com/JuliaLang/IJulia.jl):

```
using Pkg
pkg"add IJulia"
```

#### The Package

The `ParProx` package is installed by using the command:

```
using Pkg
pkg"add https://github.com/kose-y/ParProx.jl"
```

To install the other packages used in this tutoral, use the command:

```
pkg"add Printf Statistics CSV Mmap CodecZlib ROCAnalysis DataFrames"
```

For optional CUDA support, the following is also needed:

```
pkg"add CUDA Adapt"
```

It requires a CUDA-capable GPU with compute capability 5.0 (Maxwell) or higher, and an accompanying Nvidia driver with support for CUDA 10.1 or newer. CUDA is automatically downloaded before the first use, unless configured otherwise through an environment variable. See this page (https://juliagpu.gitlab.io/CUDA.jl/installation/overview/) for further information.

The following shows the version of Julia and system information.

In [59]:
```
versioninfo()
```

```
Julia Version 1.5.2
Commit 539f3ce943 (2020-09-23 23:17 UTC)
Platform Info:
  OS: Linux (x86_64-pc-linux-gnu)
  CPU: Intel(R) Xeon(R) CPU E5-2680 v2 @ 2.80GHz
  WORD_SIZE: 64
  LIBM: libopenlibm
  LLVM: libLLVM-9.0.1 (ORCJIT, ivybridge)
```

### Example 1: Cox regression with nonoverlapping groups

In this example, we perform the Cox regression on somatic mutation count data of Glioblastoma patients. We use count of mutations in protein units as the covariates, and the group lasso penalty is applied, where the variables are grouped by the gene where the unit belongs to.

#### Loading packages

In [60]:
```
using ParProx, Printf, Statistics # load the packages
using CSV, DataFrames, CodecZlib, Mmap # packages for data reading. GZip is used to read the gzipped t
ext file.
```

#### Reading data files

The file "somatic_sum_table_GBM.tsv.gz" contains the information the mutations and the count of mutations. Columns 1-9 contains the information of the mutations, and the tenth column and later contains the mutation count for each subject.

In [3]:
```
somatic = DataFrame!(CSV.File(transcode(GzipDecompressor, Mmap.mmap(
            ParProx.datadir("somatic_sum_table_GBM.tsv.gz")))));
```

For regular tab- or comma-delimited files, `DataFrame!(CSV.File("test.txt"))` should be enough.

We print first six rows and five columns of the dataframe:

In [4]:
```
first(somatic[!, 1:5], 6)
```

Out[4]: 6 rows × 5 columns

| | uniprot_accession | start_position | end_position | center_position | unit_name |
|---|---|---|---|---|---|
| | String | String | String | String | String |
| 1 | Q9H2S6 | NA | NA | NA | NCU |
| 2 | O60762 | 28 | 199 | 113 | Glycos_transf_2 |
| 3 | Q8IZE3 | 26 | 245 | 135 | Pkinase |
| 4 | Q8IZE3 | NA | NA | NA | LU |
| 5 | Q8IZE3 | NA | NA | NA | NCU |
| 6 | Q9NSG2 | 176 | 728 | 452 | DUF4487 |

and next four columns are:

In [5]:
```
first(somatic[!, 6:9], 6)
```

Out[5]: 6 rows × 4 columns

| | gene_name | gene_id | unit_label | sum |
|---|---|---|---|---|
| | String? | String | String | Int64 |
| 1 | TNMD | ENSG00000000005 | NCU | 1 |
| 2 | DPM1 | ENSG00000000419 | PIU | 1 |
| 3 | SCYL3 | ENSG00000000457 | PIU | 1 |
| 4 | SCYL3 | ENSG00000000457 | LU | 2 |
| 5 | SCYL3 | ENSG00000000457 | NCU | 1 |
| 6 | C1orf112 | ENSG00000000460 | PIU | 1 |

The actual count data starts at column 10.

In [6]:
```
first(somatic[!, 10:15], 6)
```

Out[6]: 6 rows × 6 columns

| | TCGA-02-0003 | TCGA-02-0033 | TCGA-02-0047 | TCGA-02-0055 | TCGA-02-2466 | TCGA-02-2470 |
|---|---|---|---|---|---|---|
| | Int64 | Int64 | Int64 | Int64 | Int64 | Int64 |
| 1 | 0 | 0 | 0 | 0 | 0 | 0 |
| 2 | 0 | 0 | 0 | 0 | 0 | 0 |
| 3 | 0 | 0 | 0 | 0 | 0 | 0 |
| 4 | 0 | 0 | 0 | 0 | 0 | 0 |
| 5 | 0 | 0 | 0 | 0 | 0 | 0 |
| 6 | 0 | 0 | 0 | 0 | 0 | 0 |

The file "primary_TCGA_CDR_GBM.tsv" contains the clinical information of the subjects.

In [7]:
```
# read data
cdr = CSV.read("primary_TCGA_CDR_GBM.tsv", DataFrame);
```

```
In [8]: first(cdr, 6)
```

Out[8]: 6 rows × 34 columns (omitted printing of 29 columns)

| | V1 | bcr_patient_barcode | type | age_at_initial_pathologic_diagnosis | gender |
|---|---|---|---|---|---|
| | Int64 | String | String | Int64 | String |
| 1 | 2647 | TCGA-02-0003 | GBM | 50 | MALE |
| 2 | 2664 | TCGA-02-0033 | GBM | 54 | MALE |
| 3 | 2671 | TCGA-02-0047 | GBM | 78 | MALE |
| 4 | 2676 | TCGA-02-0055 | GBM | 62 | FEMALE |
| 5 | 2734 | TCGA-02-2466 | GBM | 61 | MALE |
| 6 | 2735 | TCGA-02-2470 | GBM | 57 | MALE |

We take the survival outcome and survival time from the clinical data:

```
In [9]: survival_event = convert(Vector, cdr[!, :OS])
        survival_time = convert(Vector, cdr[!, Symbol("OS.time")]);
```

We will also add age at inital pathologic diagnosis and gender as unpenalized covariates. We will code males to 1 and females to 0.

```
In [10]: # age at initial pathological diagnosis
         age_at_diagnosis = cdr[!, :age_at_initial_pathologic_diagnosis]

         # gender: 1: male, 0: not male
         gender = map(x -> x == "MALE", cdr[!, :gender])

         # we will adjust for these two covariates by adding them as nonpenalized variables.
         clinical = hcat(age_at_diagnosis, gender);
```

We take the predictors in a single matrix:

```
In [11]: somatic_predictors = hcat(convert(Array{Float64}, transpose(convert(Array{Float64}, somatic[:, 10:end
         ]))), clinical)
```

```
Out[11]: 390×25309 Array{Float64,2}:
 0.0  0.0  0.0  0.0  0.0  0.0  0.0  …  0.0  0.0  0.0  0.0  0.0  50.0  1.0
 0.0  0.0  0.0  0.0  0.0  0.0  0.0     0.0  0.0  0.0  0.0  0.0  54.0  1.0
 0.0  0.0  0.0  0.0  0.0  0.0  0.0     0.0  0.0  0.0  0.0  0.0  78.0  1.0
 0.0  0.0  0.0  0.0  0.0  0.0  0.0     0.0  0.0  0.0  0.0  0.0  62.0  0.0
 0.0  0.0  0.0  0.0  0.0  0.0  0.0     0.0  0.0  0.0  0.0  0.0  61.0  1.0
 0.0  0.0  0.0  0.0  0.0  0.0  0.0  …  0.0  0.0  0.0  0.0  0.0  57.0  1.0
 0.0  0.0  0.0  0.0  0.0  0.0  0.0     0.0  0.0  0.0  0.0  0.0  43.0  1.0
 0.0  0.0  0.0  0.0  0.0  0.0  0.0     0.0  0.0  0.0  0.0  0.0  53.0  1.0
 0.0  0.0  0.0  0.0  0.0  0.0  0.0     0.0  0.0  0.0  0.0  0.0  64.0  1.0
 0.0  0.0  0.0  0.0  0.0  0.0  0.0     0.0  0.0  0.0  0.0  0.0  81.0  0.0
 0.0  0.0  0.0  0.0  0.0  0.0  0.0  …  0.0  0.0  0.0  0.0  0.0  84.0  0.0
 0.0  0.0  0.0  0.0  0.0  0.0  0.0     0.0  0.0  0.0  0.0  0.0  67.0  1.0
 0.0  0.0  0.0  0.0  0.0  0.0  0.0     0.0  0.0  0.0  0.0  0.0  63.0  0.0
 ⋮                        ⋮         ⋱                   ⋮
 0.0  0.0  0.0  0.0  0.0  0.0  0.0     0.0  0.0  0.0  0.0  0.0  73.0  1.0
 0.0  0.0  0.0  0.0  0.0  0.0  0.0     0.0  0.0  0.0  0.0  0.0  54.0  1.0
 0.0  0.0  0.0  0.0  0.0  0.0  0.0  …  0.0  0.0  0.0  0.0  0.0  58.0  1.0
 0.0  0.0  0.0  0.0  0.0  0.0  0.0     0.0  0.0  0.0  0.0  0.0  44.0  0.0
 0.0  0.0  0.0  0.0  0.0  0.0  0.0     0.0  0.0  0.0  0.0  0.0  49.0  0.0
 0.0  0.0  0.0  0.0  0.0  0.0  0.0     0.0  0.0  0.0  0.0  0.0  64.0  1.0
 0.0  0.0  0.0  0.0  0.0  0.0  0.0     0.0  0.0  0.0  0.0  0.0  33.0  1.0
 0.0  0.0  0.0  0.0  0.0  0.0  0.0  …  0.0  0.0  0.0  0.0  0.0  50.0  0.0
 0.0  0.0  0.0  0.0  0.0  0.0  0.0     0.0  0.0  0.0  0.0  0.0  68.0  1.0
 0.0  0.0  0.0  0.0  0.0  0.0  0.0     0.0  0.0  0.0  0.0  0.0  72.0  0.0
 0.0  0.0  0.0  0.0  0.0  0.0  0.0     0.0  0.0  0.0  0.0  0.0  51.0  1.0
 0.0  0.0  0.0  0.0  0.0  0.0  0.0     0.0  0.0  0.0  0.0  0.0  55.0  1.0
```

Unpenalized variables must be placed at the last columns of the matrix.

we normalize the input variables:

In [12]:
```
normalize(x) = (x .- mean(x; dims=1))./ std(x; dims=1)
somatic_predictors = normalize(somatic_predictors)
```

Out[12]:
```
390×25309 Array{Float64,2}:
 -0.050637  -0.050637  -0.050637  …  -0.0879321  -0.748927    0.764111
 -0.050637  -0.050637  -0.050637     -0.0879321  -0.450504    0.764111
 -0.050637  -0.050637  -0.050637     -0.0879321   1.34003     0.764111
 -0.050637  -0.050637  -0.050637     -0.0879321   0.146342   -1.30536
 -0.050637  -0.050637  -0.050637     -0.0879321   0.0717363   0.764111
 -0.050637  -0.050637  -0.050637  …  -0.0879321  -0.226687    0.764111
 -0.050637  -0.050637  -0.050637     -0.0879321  -1.27117     0.764111
 -0.050637  -0.050637  -0.050637     -0.0879321  -0.52511     0.764111
 -0.050637  -0.050637  -0.050637     -0.0879321   0.295553    0.764111
 -0.050637  -0.050637  -0.050637     -0.0879321   1.56385    -1.30536
 -0.050637  -0.050637  -0.050637  …  -0.0879321   1.78767    -1.30536
 -0.050637  -0.050637  -0.050637     -0.0879321   0.519371    0.764111
 -0.050637  -0.050637  -0.050637     -0.0879321   0.220948   -1.30536
  ⋮                               ⋱                          
 -0.050637  -0.050637  -0.050637     -0.0879321   0.967005    0.764111
 -0.050637  -0.050637  -0.050637     -0.0879321  -0.450504    0.764111
 -0.050637  -0.050637  -0.050637  …  -0.0879321  -0.152081    0.764111
 -0.050637  -0.050637  -0.050637     -0.0879321  -1.19656    -1.30536
 -0.050637  -0.050637  -0.050637     -0.0879321  -0.823532   -1.30536
 -0.050637  -0.050637  -0.050637     -0.0879321   0.295553    0.764111
 -0.050637  -0.050637  -0.050637     -0.0879321  -2.01722     0.764111
 -0.050637  -0.050637  -0.050637  …  -0.0879321  -0.748927   -1.30536
 -0.050637  -0.050637  -0.050637     -0.0879321   0.593976    0.764111
 -0.050637  -0.050637  -0.050637     -0.0879321   0.892399   -1.30536
 -0.050637  -0.050637  -0.050637     -0.0879321  -0.674321    0.764111
 -0.050637  -0.050637  -0.050637     -0.0879321  -0.375898    0.764111
```

We are considering 390 subjects and 25309 variables (25307 somatic mutations, age at diagnosis, and gender).

We check if the order of clinical data and the mutation data matches.

In [13]:
```
all(cdr[!, :bcr_patient_barcode] .== names(somatic)[10:end])
```

Out[13]:
```
true
```

#### Sorting by nonincreasing order of observed time

The samples should be sorted in nonincreasing order of observed survival time.

In [14]:
```
sortorder = sortperm(survival_time; rev=true)
somatic_predictors = normalize(somatic_predictors[sortorder, :])
survival_event = survival_event[sortorder]
survival_time = survival_time[sortorder]
```

Out[14]:
```
390-element Array{Int64,1}:
 3881
 3667
 2883
 2791
 2246
 2126
 1818
 1788
 1481
 1458
 1448
 1426
 1417
    ⋮
   15
   13
   12
    6
    6
    6
    5
    4
    4
    3
    3
    0
```

The following code creates a vector of String to be used as the name of each variable. We will use the first eight columns of `somatic` data frame to make a variable name.

In [15]:
```
somatic_variable_names = map(x -> "$(x[1])\t$(x[2])\t$(x[3])\t" *
    "$(x[4])\t$(x[5])\t$(x[6])\t$(x[7])\t$(x[8])", eachrow(somatic))
somatic_variable_names = vcat(somatic_variable_names, map(x -> "$x\t\t\t\t\t\t\t",
        ["age_at_diagnosis", "gender"]));
```

#### Group definition

We figure out unique group names ( `somatic_groups` ), size of each group ( `somatic_group_sizes` ), and which group each variable belongs to ( `somatic_grpidx` ).

In [16]:
```
somatic_groups = unique(somatic[!, :gene_id])
somatic_group_sizes = map(somatic_groups) do x
    count(y -> y == x, somatic[!, :gene_id])
end
somatic_grpidx = map(somatic[!, :gene_id]) do x
    findfirst(y -> y .== x, somatic_groups)
end;
```

`somatic_grpidx` gives the group index each variable belongs. Note that Julia uses 1-based indexing.

In [17]:
```
somatic_grpidx
```

Out[17]:
```
25307-element Array{Int64,1}:
     1
     2
     3
     3
     3
     4
     5
     5
     5
     5
     5
     5
     5
     ⋮
 14247
 14248
 14249
 14250
 14251
 14252
 14253
 14253
 14253
 14253
 14253
 14254
```

#### Cross validation

For the cross validation, we create a `COXUpdate` object specifying the update rules: maximum number of iterations, steps between intermediate evaluation, relative tolerance for stopping, and whether to print the intermediate output.

For cross validation in Cox regression, "C-index" is used as the metric (the higher, the better). Stratified k-fold cross validation is implemented (k=5 here), i.e., we ensure that the proportion of event and censored subject as equal as possible across the folds.

In [18]:
```
using Random
Random.seed!(222);
```

We perform 5-fold corss validation using CPU only:

In [19]:
```
T = Float64
A = Array
U = ParProx.COXUpdate(; maxiter=10000, step=10, tol=5e-4, verbose=false)
lambdas = 10 .^ (range(-4, stop=-6, length=21)) # 21 values equally log-spaced in 10^-5 .. 10^-7
penalties = [GroupNormL2(l, somatic_grpidx) for l in lambdas]

@time score = ParProx.cross_validate(U, somatic_predictors,
    survival_event, survival_time, penalties, 5; T=Float64, A=Array);
```

```
 17.086662 seconds (8.32 M allocations: 2.010 GiB, 1.12% gc time)
```

The number of unpenalized variables are automatically detected from the penalty structure.

Simple lasso penalty could be used by replacing line 5 above with

```
penalties = [NormL1(l; unpen=2) for l in lambdas]
```

For lasso penalty, the user should provide the number of unpenalized variables at the last columns as a keyword argument.

The resulting c-index for each fold is listed as:

In [20]:
```
score
```

Out[20]:
```
21×5 Array{Float64,2}:
 0.630789  0.652882  0.68023   0.644217  0.584255
 0.630789  0.652882  0.68023   0.644217  0.584255
 0.630789  0.652882  0.68023   0.644217  0.584255
 0.630789  0.652882  0.68023   0.644217  0.584255
 0.630789  0.652882  0.68023   0.644217  0.584255
 0.630789  0.652882  0.68023   0.644217  0.584255
 0.630789  0.652882  0.68023   0.644217  0.584255
 0.629503  0.652882  0.68023   0.644217  0.584255
 0.629503  0.652882  0.68023   0.644217  0.584255
 0.627787  0.652882  0.68023   0.644217  0.584255
 0.625643  0.652882  0.681115  0.644682  0.584697
 0.622213  0.653322  0.680672  0.642824  0.588235
 0.616209  0.656841  0.678461  0.639573  0.591774
 0.607633  0.661241  0.678903  0.63725   0.5931
 0.596484  0.6674    0.678903  0.635857  0.597966
 0.587479  0.671359  0.671827  0.627497  0.598408
 0.58705   0.67004   0.6705    0.619136  0.59885
 0.580617  0.663     0.67448   0.60288   0.593543
 0.577187  0.653322  0.669173  0.601022  0.589562
 0.577187  0.652442  0.670057  0.599628  0.584255
 0.580189  0.644083  0.659     0.588946  0.58337
```

We select the penalty parameter that maximizes the mean c-index as following:

In [21]:
```
lambda_idx = argmax(mean(score; dims=2)[:])
lambda = lambdas[lambda_idx]
```

Out[21]:
```
0.0001
```

Now we fit the model with the selected value of lambda.

In [22]:
```
p = GroupNormL2(lambda, somatic_grpidx)
U = ParProx.COXUpdate(; maxiter=20000, step=20, tol=1e-6, verbose=false)
V = ParProx.COXVariables{Float64, Array}(somatic_predictors, survival_event, survival_time, p; eval_ob
j=true)
@time ParProx.fit!(U, V)
```

```
 55.770952 seconds (321.56 k allocations: 4.346 GiB, 0.78% gc time)
```

The tuned value selects the most sparse model possible.

We obtain the indexes of variables with nonzero coefficients:

In [23]:
```
nonzero_idxs = (1:length(V.β))[V.β .!= 0];
```

The selected variables, along with the coefficients can be printed as below:

In [24]:
```
for (variable_name, value) in zip(somatic_variable_names[nonzero_idxs], V.β[nonzero_idxs])
    println(variable_name, "\t", value)
end
```

```
age_at_diagnosis                                                                 0.12933261107254107
gender                                                          0.06219181385720149
```

#### (optional) Analysis using GPU

For using GPU, as stated before, the package CUDA.jl is required.

Cross validation on GPU is performed by the following code.

```
In [25]: using CUDA, Adapt
```

The following identifies the GPU we are running the code on.

```
In [26]: CUDA.device()

Out[26]: CuDevice(0): GeForce GTX 1080
```

It takes only a minor modification to run the cross validation on GPU.

```
In [27]: Random.seed!(222)
         T = Float64
         A = CuArray
         U = ParProx.COXUpdate(; maxiter=10000, step=10, tol=5e-4, verbose=false)
         lambdas = 10 .^ (range(-5, stop=-7, length=21)) # 21 values equally log-spaced in 10^-5 .. 10^-7
         penalties = [GroupNormL2{T,A}(l, somatic_grpidx) for l in lambdas]

         @time score = ParProx.cross_validate(U, somatic_predictors,
             survival_event, survival_time, penalties, 5; T=T, A=A);

          27.531135 seconds (39.08 M allocations: 5.213 GiB, 3.11% gc time)
```

It can be seen that analysis on GPU is 6-8 times faster except for the very first fit where the functions are compiled. It may be desirable to use `T=Float32` to reduce memory usage of the GPU.

```
In [28]: lambda_idx = argmax(mean(score; dims=2)[:])
         lambda = lambdas[lambda_idx]

Out[28]: 1.0e-5
```

```
In [29]: p = GroupNormL2{T,A}(lambda, somatic_grpidx)
         U = ParProx.COXUpdate(; maxiter=20000, step=20, tol=1e-6, verbose=false)
         V = ParProx.COXVariables{T,A}(adapt(A{T}, somatic_predictors),
             adapt(A{T}, survival_event),
             adapt(A{T}, survival_time), p; eval_obj=true)
         @time ParProx.fit!(U, V)

          11.971347 seconds (4.06 M allocations: 6.498 GiB, 2.23% gc time)
```

```
In [30]: nonzero_idxs = (1:length(V.β))[V.β .!= 0];

         ┌ Warning: Performing scalar operations on GPU arrays: This is very slow, consider disallowing these
         operations with `allowscalar(false)`
         └ @ GPUArrays /home/kose/.julia/packages/GPUArrays/WV76E/src/host/indexing.jl:43
```

```
In [31]: for (variable_name, value) in zip(somatic_variable_names[nonzero_idxs], V.β[nonzero_idxs])
             println(variable_name, "\t", value)
         end
```

```
Q6NUM9  NA      NA      NA      LU      RETSAT  ENSG00000042445 LU      -0.001632894180608977
Q5JXM2  115     342     228     Methyltransf_22 METTL24 ENSG00000053328 PIU     -0.001632894180608977
P27815  NA      NA      NA      LU      PDE4A   ENSG00000065989 LU      -0.003601501518672006
Q12888  NA      NA      NA      NCU     TP53BP1 ENSG00000067369 NCU     -0.003260300192141834 4
Q9NPF4  23      301     162     Peptidase_M22   OSGEP   ENSG00000092094 PIU     -0.000576944637489909
P25490  NA      NA      NA      LU      YY1     ENSG00000100811 LU      -0.003601501518672006
O00264  NA      NA      NA      LU      PGRMC1  ENSG00000101856 LU      -0.001632894180608977
Q9HC57  NA      NA      NA      LU      WFDC1   ENSG00000103175 LU      -0.003601501518672006
Q9NUM4  NA      NA      NA      NCU     TMEM106B        ENSG00000106460 NCU     -0.001632894180608977
Q9Y2J4  NA      NA      NA      LU      AMOTL2  ENSG00000114019 LU      -0.003260300192141834 4
Q9UI95  NA      NA      NA      LU      MAD2L2  ENSG00000116670 LU      -0.0003051353783785691
Q8TEH3  NA      NA      NA      LU      DENND1A ENSG00000119522 LU      -0.001632894180608977
Q8IX21  NA      NA      NA      LU      SLF2    ENSG00000119906 LU      -0.000576944637489909
Q9BUF7  NA      NA      NA      NCU     CRB3    ENSG00000130545 NCU     -0.000576944637489909
P36222  NA      NA      NA      NCU     CHI3L1  ENSG00000133048 NCU     -0.001632894180608977
P49638  NA      NA      NA      NCU     TTPA    ENSG00000137561 NCU     -0.001632894180608977
O75874  9       401     205     Iso_dh  IDH1    ENSG00000138413 PIU     -0.013317016909376768
O75874  130     140     135     py      IDH1    ENSG00000138413 PIU     -0.013317016909376768
Q92733  275     491     383     PRCC    PRCC    ENSG00000143294 PIU     -0.000576944637489909
O15492  NA      NA      NA      LU      RGS16   ENSG00000143333 LU      -0.001632894180608977
Q8IVH4  101     384     242     ArgK    MMAA    ENSG00000151611 PIU     -0.000576944637489909
Q96L08  32      91      61      Sushi   SUSD3   ENSG00000157303 PIU     -0.001632894180608977
Q9H0A9  1       153     77      Speriolin_N     SPATC1L ENSG00000160284 PIU     -0.003260300192141834
4
P50607  258     500     379     Tub     TUB     ENSG00000166402 PIU     -0.001632894180608977
Q96ER3  NA      NA      NA      LU      SAAL1   ENSG00000166788 LU      -0.001632894180608977
Q495B1  19      121     70      Ank_2   ANKDD1A ENSG00000166839 PIU     -1.15266492263703e-5
NA      NA      NA      NA      LU      FAM129C ENSG00000167483 LU      -0.0005395223579764626
P61018  10      171     90      Ras     RAB4B   ENSG00000167578 PIU     -1.15266492263703e-5
Q8WYH8  6       107     56      ING     ING5    ENSG00000168395 PIU     -0.0003051353783785691
Q9H5H4  NA      NA      NA      LU      ZNF768  ENSG00000169957 LU      -0.001632894180608977
Q9BZR9  NA      NA      NA      LU      TRIM8   ENSG00000171206 LU      -0.000576944637489909
O94811  52      207     129     p25-alpha       TPPP    ENSG00000171368 PIU     -0.001632894180608977
P14652  NA      NA      NA      LU      HOXB2   ENSG00000173917 LU      -0.001632894180608977
A6NIH7  88      247     167     GMP_PDE_delta   UNC119B ENSG00000175970 PIU     -0.003260300192141834
4
Q8N1N2  NA      NA      NA      LU      DYNAP   ENSG00000178690 LU      -1.15266492263703e-5
Q7Z5W3  NA      NA      NA      NCU     BCDIN3D ENSG00000186666 NCU     -0.001632894180608977
Q86Z23  111     235     173     C1q     C1QL4   ENSG00000186897 PIU     -0.001632894180608977
Q7Z444  43      201     122     Ras     ERAS    ENSG00000187682 PIU     -0.0003051353783785691
P42356  NA      NA      NA      LU      PI4KA   ENSG00000241973 LU      -1.15266492263703e-5
NA      NA      NA      NA      NCU     DNM1P47 ENSG00000259660 NCU     -0.003601501518672006
age_at_diagnosis                                                                0.11455939819834576
gender                                                          0.05527884841839899
```

### Example 2: Logistic regression with overlapping groups

In this example, we perform logistic regression with overlapping group lasso penalties. This package provides a specialized interface for overlapping groups. We use mRNA gene expression signature to classify breast cancer patients undergoing chemotherapy with anthracycline and neoadjuvant agents into two classes. Group of variables is defined by the pathway and GO terms. Genes not belonging to any of the pathways is included as a singleton group.

In latent group lasso model, variables in multiple groups are replicated each time, each replication corresponding to the contribution of a variable as a member of each group. We estimate the corresponding coefficient for each replication, and the sum of these replicated coefficients is the total coefficient for that variable. This is implemented as a lazy multiplication between the original data and a sparse matrix, reducing the memory usage.

For this analysis, we have four data files.

- `exprdata_set2_cut.txt.gz` contains the gene expression data of the subjects.
- `sample_set2.txt.gz` contains the clinical information of the subjects.
- `PATHWAYgroup.txt.gz` contains the information on the group structure: membership of each gene in a pathway.
- `gene_info.txt.gz` matches gene symbols to their functions.

For the preprocessing, we first read the gene expression and clinical data, then parse `PATHWAYgroup.txt.gz` to determine the group structure. In the process, we also attach the functions to each of the genes and pathways. Then, we construct variable names from the group structure and the functions.

#### Reading covariates

The following lines read the gene expression data. The first column of the file `exprdata_set2_cut.txt.gz` has the gene names, and the other columns represent the normalized gene expression level of each patient. We extract the gene names for later use.

In [32]:
```
brca_data = DataFrame!(CSV.File(transcode(GzipDecompressor, Mmap.mmap(
            ParProx.datadir("exprdata_set2_cut.txt.gz")))))
gene_names = brca_data[!, :Gene]
```

Out[32]:
```
1043-element Array{String,1}:
 "RFC2"
 "PAX8"
 "THRA"
 "EPHB3"
 "CFL1"
 "YY1"
 "ZPR1"
 "RHOA"
 "GUK1"
 "DSP"
 "RAD21"
 "SF3B2"
 "NDRG1"
 ⋮
 "ORAI3"
 "FAM174B"
 "ZNF335"
 "SEH1L"
 "LOC389906"
 "KIF18B"
 "RACGAP1"
 "CASP8AP2"
 "KLK5"
 "OR7E47P"
 "SCAF4"
 "SNHG17"
```

Now we load the covariates. Be carful when matching the subject information on the gene expression data and clinical data. We gather penalized variables and unpenalized variables separately.

In [33]:
```
# patient names in the gene expression file
patient_barcode = map(x -> string(x), names(brca_data)[2:end])

X = collect(transpose(convert(Matrix, brca_data[:, 2:end])))

subject_data = CSV.File(transcode(GzipDecompressor, Mmap.mmap(
            ParProx.datadir("sample_set2.txt.gz")))) |> DataFrame
normalize(x) = (x .- mean(x; dims=1))./ std(x; dims=1)
Age  = normalize(subject_data[!, :Age])
interceptcol = ones(size(X, 1))

X_unpen = [Age interceptcol]

y = subject_data[!, :PCR_outcome]

# match subject ids
subject_data = filter(x -> x[:GEO_accession] in patient_barcode, subject_data)
@assert all(subject_data[!, :GEO_accession] .== patient_barcode)
```

#### Building the groups

Now we read the group information. Each row of the file `PATHWAYgroup.txt.gz` lists a gene, a pathway id, function of the pathway and the gene symbol, indicating the membership of the gene in the pathway. We read the data file first:

In [34]:
```
group_info = DataFrame!(CSV.File(transcode(GzipDecompressor, Mmap.mmap(
            ParProx.datadir("PATHWAYgroup.txt.gz")))))
head(group_info)
```

Out[34]: 6 rows × 4 columns

| | ENSG | Pathwayid | Function | Genesym |
|---|---|---|---|---|
| | String | String | String | String |
| 1 | ENSG00000148584 | GO:0006397 | mRNA processing | A1CF |
| 2 | ENSG00000148584 | GO:0010467 | gene expression | A1CF |
| 3 | ENSG00000148584 | GO:0016554 | cytidine to uridine editing | A1CF |
| 4 | ENSG00000148584 | GO:0016556 | mRNA modification | A1CF |
| 5 | ENSG00000148584 | GO:0050821 | protein stabilization | A1CF |
| 6 | ENSG00000175899 | KEGG:04610 | Complement and coagulation cascades | A2M |

We create variables to store the group information.

- `gene_names` (defined above): Each element lists the gene symbols in the analysis. Location in this list matches the index of each gene.
- `pathwayids` : A unique list of pathway ids. Location in this list determines the index of each pathway.
- `variable_to_groups` : Each element lists the group each variable is in.
- `group_to_variables` : Each element lists the variable each group includes.
- `genesym_to_variableidx` : Matches gene symbol to variable index.
- `pathwayid_to_groupidx` : Matches pathway id to group index.

In [35]:
```
pathwayids = Vector{String}()
variable_to_groups = [Int[] for i in 1:length(gene_names)]
group_to_variables = Vector{Vector{Int}}()
genesym_to_variableidx = Dict{String, Int}()
pathwayid_to_groupidx = Dict{String, Int}();
```

These dictionaries map pathwayids and gene symbols to their functions.

In [36]:
```
pathwayid_to_pathwayfunction = Dict{String, String}()
genesym_to_genefunction = Dict{String, String}();
```

We build `genesym_to_variableidx` and `genesym_to_genefunction` :

In [37]:
```
for (i, v) in enumerate(gene_names)
    genesym_to_variableidx[v] = i
end

gene_info = CSV.File(transcode(GzipDecompressor, Mmap.mmap(
            ParProx.datadir("gene_info.txt.gz")))) |> DataFrame
for r in eachrow(gene_info)
    genesym_to_genefunction[r[2]] = r[4]
end
```

`pathwayids` , `variable_to_groups` , `group_to_variables` , `pathwayid_to_groupidx` , and `pathwayid_to_pathwayfunction` are all built by parsing `group_info` .

In [38]:
```
num_groups = 0
for r in eachrow(group_info)
    if !(r.Genesym in keys(genesym_to_variableidx)) # skip any gene not in our gene list
        continue
    end
    # get variable index from the gene symbol
    variable_idx = genesym_to_variableidx[r.Genesym]

    if !(r.Pathwayid in pathwayids) # if the pathway id is new ... #!haskey(pathwayid_to_genesym, r.Pa
thwayid)
        # add a new group
        num_groups += 1
        push!(pathwayids, r.Pathwayid)
        push!(group_to_variables, Int[])
        pathwayid_to_groupidx[r.Pathwayid] = num_groups
        group_idx = num_groups
    else
        group_idx = pathwayid_to_groupidx[r.Pathwayid]
    end

    # insert membership information
    push!(variable_to_groups[variable_idx], group_idx)
    push!(group_to_variables[group_idx], variable_idx)

    # retrieve pathway information
    if !haskey(pathwayid_to_pathwayfunction, r.Pathwayid)
        pathwayid_to_pathwayfunction[r.Pathwayid] = r.Function
    else
        # if redundant, just check correctness
        @assert pathwayid_to_pathwayfunction[r.Pathwayid] == r.Function
    end
end
```

We also enter non-pathway genes as singleton groups.

In [39]:
```
cnt = 0
for (i, v) in enumerate(gene_names)
    if length(variable_to_groups[i]) > 0 # gene is in some of the groups
        continue
    else # gene is not in any of the groups
        num_groups += 1
        cnt += 1
        # update group info
        push!(variable_to_groups[i], num_groups)
        push!(group_to_variables, [i])

        # update variable names
        pathwayid_placeholder =
            if haskey(genesym_to_genefunction, v)
                "Singleton of: $v"
            else # some genes do not appear in the gene function list
                "Singleton of: $v"
            end
        push!(pathwayids, pathwayid_placeholder)
    end
end
cnt
```

Out[39]: 90

90 singleton groups were added!

#### Constructing variable names

Here we create a vector of Strings for pathway ids and their functions.

First, we create a vector of variable names. We include age and intercept as nonpenalized variables.

In [40]:
```
variable_names = [gene_names; "Age\t\t"; "intercept\t\t"]
```

Out[40]:
```
1045-element Array{String,1}:
 "RFC2"
 "PAX8"
 "THRA"
 "EPHB3"
 "CFL1"
 "YY1"
 "ZPR1"
 "RHOA"
 "GUK1"
 "DSP"
 "RAD21"
 "SF3B2"
 "NDRG1"
 ⋮
 "ZNF335"
 "SEH1L"
 "LOC389906"
 "KIF18B"
 "RACGAP1"
 "CASP8AP2"
 "KLK5"
 "OR7E47P"
 "SCAF4"
 "SNHG17"
 "Age\t\t"
 "intercept\t\t"
```

We have 1045 variables for the analysis.

Now we concatenate pathway names and their functions.

In [41]:
```
pathways_with_functions = map(pathwayids) do x
    if haskey(pathwayid_to_pathwayfunction, x)
        "$x: $(pathwayid_to_pathwayfunction[x])"
    else
        x
    end
end
```

Out[41]:
```
5719-element Array{String,1}:
 "KEGG:00280: Valine, leucine and isoleucine degradation"
 "KEGG:04727: GABAergic synapse"
 "KEGG:01100: Metabolic pathways"
 "KEGG:00250: Alanine, aspartate and glutamate metabolism"
 "KEGG:00640: Propanoate metabolism"
 "KEGG:00410: beta-Alanine metabolism"
 "KEGG:00650: Butanoate metabolism"
 "GO:0001666: response to hypoxia"
 "GO:0007268: synaptic transmission"
 "GO:0007269: neurotransmitter secretion"
 "GO:0007620: copulation"
 "GO:0007626: locomotory behavior"
 "GO:0009449: gamma-aminobutyric acid biosynthetic process"
 ⋮
 "Singleton of: TEX13B"
 "Singleton of: IQCG"
 "Singleton of: LOC100131532"
 "Singleton of: C6orf25"
 "Singleton of: TMEM14B"
 "Singleton of: FAM64A"
 "Singleton of: GPR124"
 "Singleton of: FAM174B"
 "Singleton of: LOC389906"
 "Singleton of: OR7E47P"
 "Singleton of: SCAF4"
 "Singleton of: SNHG17"
```

We now create replicated variable names: each replicated variable corresponds to a membership of a gene to a group. The name should include pathway information and group information.

In [42]:
```
variable_names_replicated = String[]
cnt = 0
for i in 1:length(group_to_variables)
    for v in group_to_variables[i]
        pf = pathways_with_functions[i]
        g = gene_names[v]
        gt = genesym_to_genefunction[g]
        push!(variable_names_replicated, "$pf\t$g\t$gt")
    end
end
```

Finally, we append nonpenalized variables to the list.

In [43]:
```
variable_names_replicated = [variable_names_replicated; "Age\t\t"; "intercept\t\t"]
```

Out[43]:
```
21965-element Array{String,1}:
 "KEGG:00280: Valine, leucine and isoleucine degradation\tABAT\t4-aminobutyrate aminotransferase"
 "KEGG:00280: Valine, leucine and isoleucine degradation\tAUH\tAU RNA binding protein/enoyl-CoA hydra
tase"
 "KEGG:00280: Valine, leucine and isoleucine degradation\tHMGCL\t3-hydroxymethyl-3-methylglutaryl-CoA
lyase"
 "KEGG:00280: Valine, leucine and isoleucine degradation\tIVD\tisovaleryl-CoA dehydrogenase"
 "KEGG:00280: Valine, leucine and isoleucine degradation\tMCCC1\tmethylcrotonoyl-CoA carboxylase 1 (a
lpha)"
 "KEGG:00280: Valine, leucine and isoleucine degradation\tMCCC2\tmethylcrotonoyl-CoA carboxylase 2 (b
eta)"
 "KEGG:00280: Valine, leucine and isoleucine degradation\tMUT\tmethylmalonyl CoA mutase"
 "KEGG:04727: GABAergic synapse\tABAT\t4-aminobutyrate aminotransferase"
 "KEGG:04727: GABAergic synapse\tGABRA4\tgamma-aminobutyric acid (GABA) A receptor, alpha 4"
 "KEGG:04727: GABAergic synapse\tPRKCA\tprotein kinase C, alpha"
 "KEGG:04727: GABAergic synapse\tPRKX\tprotein kinase, X-linked"
 "KEGG:01100: Metabolic pathways\tABAT\t4-aminobutyrate aminotransferase"
 "KEGG:01100: Metabolic pathways\tACADL\tacyl-CoA dehydrogenase, long chain"
 ⋮
 "Singleton of: LOC100131532\tLOC100131532\tuncharacterized LOC100131532"
 "Singleton of: C6orf25\tC6orf25\tchromosome 6 open reading frame 25"
 "Singleton of: TMEM14B\tTMEM14B\ttransmembrane protein 14B"
 "Singleton of: FAM64A\tFAM64A\tfamily with sequence similarity 64, member A"
 "Singleton of: GPR124\tGPR124\tG protein-coupled receptor 124"
 "Singleton of: FAM174B\tFAM174B\tfamily with sequence similarity 174, member B"
 "Singleton of: LOC389906\tLOC389906\tzinc finger protein 839 pseudogene"
 "Singleton of: OR7E47P\tOR7E47P\tolfactory receptor, family 7, subfamily E, member 47 pseudogene"
 "Singleton of: SCAF4\tSCAF4\tSR-related CTD-associated factor 4"
 "Singleton of: SNHG17\tSNHG17\tsmall nucleolar RNA host gene 17 (non-protein coding)"
 "Age\t\t"
 "intercept\t\t"
```

We have 21965 variables on the replicated space.

#### Cross validation

Now we perform the 5-fold cross validation. The default criteria for cross validation is accuracy. It can be changed to area under the ROC curve (ROC) by adding a keyword parameter `criteria=auc` .

In [44]:
```
lambdas = 10 .^ (range(-6, stop=-9, length=31))
```

Out[44]:
```
31-element Array{Float64,1}:
 1.0e-6
 7.943282347242822e-7
 6.30957344480193e-7
 5.011872336272725e-7
 3.981071705534969e-7
 3.162277660168379e-7
 2.5118864315095823e-7
 1.9952623149688787e-7
 1.584893192461114e-7
 1.2589254117941662e-7
 1.0e-7
 7.943282347242822e-8
 6.30957344480193e-8
 ⋮
 1.2589254117941661e-8
 1.0e-8
 7.943282347242822e-9
 6.309573444801943e-9
 5.011872336272715e-9
 3.981071705534969e-9
 3.1622776601683795e-9
 2.511886431509582e-9
 1.9952623149688828e-9
 1.584893192461111e-9
 1.2589254117941663e-9
 1.0e-9
```

For overlapping groups, `group_to_variables` is used as an input, along with `lambdas`, rather than the pre-constructed penalties.

In [45]:
```
using Random
Random.seed!(222)
U = ParProx.LogisticUpdate(; maxiter=30000, step=20, tol=5e-4, verbose=false)
@time scores = ParProx.cross_validate(U, X, X_unpen, y, group_to_variables, lambdas, 5; T =Float64, cr
iteria=auc);
```

```
 14.409149 seconds (19.93 M allocations: 5.296 GiB, 3.57% gc time)
```

We select the lambda with the highest mean AUC.

In [46]:
```
lambda_idx = argmax(mean(scores; dims=2)[:])
lambda = lambdas[lambda_idx]
```

Out[46]:
```
1.2589254117941663e-9
```

Now fitting with the selected lambda:

In [47]:
```
U = ParProx.LogisticUpdate(; maxiter=30000, step=20, tol=1e-5, verbose=false)
V = ParProx.LogisticVariables{Float64}(X, X_unpen, y, lambda, group_to_variables)
@time ParProx.fit!(U, V)
```

```
 15.742283 seconds (1.75 M allocations: 10.678 GiB, 2.32% gc time)
```

In order to obtain the results in the original dimension, we need to obtain the "group matrix". Multiplying the group matrix by the *penalized* part of the coefficients gives the aggregate coefficient for each penalized variable. Non-penalized variables (in this case, the last two variables) should be treated separately.

In [48]:
```
_, grpmat, _ = ParProx.mapper_mat_idx(group_to_variables, length(gene_names));
```

In [49]:
```
size(grpmat)
```

Out[49]:
```
(1043, 21963)
```

In [50]:
```
β_orig = vcat(grpmat * collect(V.β[1:end-2]), collect(V.β)[end-1:end]);
```

The following prints the coefficients in the replicated dimensions:

In [51]:

```
for (v, β) in zip(variable_names_replicated[V.β .!= 0], V.β[V.β .!= 0])
    println("$v\t$β")
end
```

KEGG:00280: Valine, leucine and isoleucine degradation  ABAT    4-aminobutyrate aminotransferase
0.0008323544134017853
KEGG:00280: Valine, leucine and isoleucine degradation  AUH     AU RNA binding protein/enoyl-CoA hydr
atase   -0.0014593426888791468
KEGG:00280: Valine, leucine and isoleucine degradation  HMGCL   3-hydroxymethyl-3-methylglutaryl-CoA
lyase   -0.0010615725170478399
KEGG:00280: Valine, leucine and isoleucine degradation  IVD     isovaleryl-CoA dehydrogenase    -0.00
09348559171503029
KEGG:00280: Valine, leucine and isoleucine degradation  MCCC1   methylcrotonoyl-CoA carboxylase 1 (al
pha)    8.30660555675512e-5
KEGG:00280: Valine, leucine and isoleucine degradation  MCCC2   methylcrotonoyl-CoA carboxylase 2 (be
ta)     -0.000783494419844115
KEGG:00280: Valine, leucine and isoleucine degradation  MUT     methylmalonyl CoA mutase        0.002
754888016414 5425
KEGG:04727: GABAergic synapse   ABAT    4-aminobutyrate aminotransferase        0.0008232735329612729
KEGG:04727: GABAergic synapse   GABRA4  gamma-aminobutyric acid (GABA) A receptor, alpha 4      -0.00
14658463388531478
KEGG:04727: GABAergic synapse   PRKCA   protein kinase C, alpha -5.901736217620083e-6
KEGG:04727: GABAergic synapse   PRKX    protein kinase, X-linked        0.0003862539887885207
KEGG:01100: Metabolic pathways  ABAT    4-aminobutyrate aminotransferase        0.0008361539792630329
KEGG:01100: Metabolic pathways  ACADL   acyl-CoA dehydrogenase, long chain      0.000985316179371731
KEGG:01100: Metabolic pathways  ALDH1A2 aldehyde dehydrogenase 1 family, member A2      -0.0030292702
18180024
KEGG:01100: Metabolic pathways  ALDH5A1 aldehyde dehydrogenase 5 family, member A1      0.00233840571
78505286
KEGG:01100: Metabolic pathways  ALLC    allantoicase    -0.0009259344378902454
KEGG:01100: Metabolic pathways  ALOX12  arachidonate 12-lipoxygenase    -0.001960674381048051
KEGG:01100: Metabolic pathways  AMD1    adenosylmethionine decarboxylase 1      -0.002535890300186689
KEGG:01100: Metabolic pathways  ASS1    argininosuccinate synthase 1    0.0001442647137212186
KEGG:01100: Metabolic pathways  AUH     AU RNA binding protein/enoyl-CoA hydratase      -0.0014668253
656568745
KEGG:01100: Metabolic pathways  BDH1    3-hydroxybutyrate dehydrogenase, type 1 -0.000207475104397339
58
KEGG:01100: Metabolic pathways  CBS     cystathionine-beta-synthase     0.001522422413483285
KEGG:01100: Metabolic pathways  CPOX    coproporphyrinogen oxidase      0.0012789641964398664
KEGG:01100: Metabolic pathways  CTH     cystathionine gamma-lyase       0.0017546337885964076
KEGG:01100: Metabolic pathways  CTPS1   CTP synthase 1  -0.00042155251296389234
KEGG:01100: Metabolic pathways  CTPS2   CTP synthase 2  0.0017650016553290638
KEGG:01100: Metabolic pathways  CYP27B1 cytochrome P450, family 27, subfamily B, polypeptide 1  -0.00
047356623759338797
KEGG:01100: Metabolic pathways  DCT     dopachrome tautomerase  -0.002360766795992877
KEGG:01100: Metabolic pathways  DHCR24  24-dehydrocholesterol reductase -0.0017137397769336163
KEGG:01100: Metabolic pathways  DHFR    dihydrofolate reductase -0.0003076937765407043
KEGG:01100: Metabolic pathways  DTYMK   deoxythymidylate kinase (thymidylate kinase)    0.00349359546
24852592
KEGG:01100: Metabolic pathways  GAL3ST1 galactose-3-O-sulfotransferase 1        0.0018631853227812837
KEGG:01100: Metabolic pathways  GALK2   galactokinase 2 3.174981063618626e-5
KEGG:01100: Metabolic pathways  GALNT6  polypeptide N-acetylgalactosaminyltransferase 6 -0.0017830737
236985239
KEGG:01100: Metabolic pathways  GCDH    glutaryl-CoA dehydrogenase      -5.142324143013614e-6
KEGG:01100: Metabolic pathways  GCLM    glutamate-cysteine ligase, modifier subunit     -0.0037732217
757471516
KEGG:01100: Metabolic pathways  GCNT1   glucosaminyl (N-acetyl) transferase 1, core 2   0.00069857695
33269947
KEGG:01100: Metabolic pathways  GCNT2   glucosaminyl (N-acetyl) transferase 2, I-branching enzyme (I
blood group)    0.0024411127133358054
KEGG:01100: Metabolic pathways  GCNT3   glucosaminyl (N-acetyl) transferase 3, mucin type       -0.00
23204484305728522
KEGG:01100: Metabolic pathways  GCNT4   glucosaminyl (N-acetyl) transferase 4, core 2   0.00045352466
828224477
KEGG:01100: Metabolic pathways  GLUD2   glutamate dehydrogenase 2       -0.000660750767022681
KEGG:01100: Metabolic pathways  GMPS    guanine monphosphate synthase   0.0017809600837371215
KEGG:01100: Metabolic pathways  GUK1    guanylate kinase 1      0.0032840032282921345
KEGG:01100: Metabolic pathways  HMGCL   3-hydroxymethyl-3-methylglutaryl-CoA lyase      -0.0010678229
674893763
KEGG:01100: Metabolic pathways  HPRT1   hypoxanthine phosphoribosyltransferase 1        -0.0011894467
283326514
KEGG:01100: Metabolic pathways  HSD17B4 hydroxysteroid (17-beta) dehydrogenase 4        -0.0011670301
666150769
KEGG:01100: Metabolic pathways  HSD17B6 hydroxysteroid (17-beta) dehydrogenase 6        -0.0008712071
093238519
KEGG:01100: Metabolic pathways  IDO1    indoleamine 2,3-dioxygenase 1   0.0012307680167228244
KEGG:01100: Metabolic pathways  IMPDH1  IMP (inosine 5'-monophosphate) dehydrogenase 1  -0.0018789353
029630802
KEGG:01100: Metabolic pathways  IMPDH2  IMP (inosine 5'-monophosphate) dehydrogenase 2  -0.0003495009
6342178943
KEGG:01100: Metabolic pathways  INPP4B  inositol polyphosphate-4-phosphatase, type II, 105kDa   0.000
2893933596837731
KEGG:01100: Metabolic pathways  IVD     isovaleryl-CoA dehydrogenase    -0.0009384574293990815
KEGG:01100: Metabolic pathways  LDHC    lactate dehydrogenase C 0.0014884182403979084
KEGG:01100: Metabolic pathways  LIAS    lipoic acid synthetase  -0.0005559477430808897

KEGG:01100: Metabolic pathways  MCCC1   methylcrotonoyl-CoA carboxylase 1 (alpha)       8.57497893915
0512e-5
KEGG:01100: Metabolic pathways  MCCC2   methylcrotonoyl-CoA carboxylase 2 (beta)        -0.0007881466
527775406
KEGG:01100: Metabolic pathways  MGAM    maltase-glucoamylase (alpha-glucosidase)        -0.0013146570
407050662
KEGG:01100: Metabolic pathways  MTAP    methylthioadenosine phosphorylase       0.0026683610533921815
KEGG:01100: Metabolic pathways  MTHFD1  methylenetetrahydrofolate dehydrogenase (NADP+ dependent) 1,
methenyltetrahydrofolate cyclohydrolase, formyltetrahydrofolate synthetase      -0.000665594343038631
2
KEGG:01100: Metabolic pathways  MUT     methylmalonyl CoA mutase        0.0027674298852422047
KEGG:01100: Metabolic pathways  ODC1    ornithine decarboxylase 1       0.0008286228438253801
KEGG:01100: Metabolic pathways  OGDH    oxoglutarate (alpha-ketoglutarate) dehydrogenase (lipoamide)
-0.0002956255707684114
KEGG:01100: Metabolic pathways  PGLS    6-phosphogluconolactonase       0.0017539382257704043
KEGG:01100: Metabolic pathways  PHGDH   phosphoglycerate dehydrogenase  0.0002951364524401764
KEGG:01100: Metabolic pathways  PLA2G5  phospholipase A2, group V       0.0020854610553521975
KEGG:01100: Metabolic pathways  PLCB1   phospholipase C, beta 1 (phosphoinositide-specific)     0.000
15512507055808085
KEGG:01100: Metabolic pathways  PNPO    pyridoxamine 5'-phosphate oxidase       -0.001095584789131929
9
KEGG:01100: Metabolic pathways  POLA1   polymerase (DNA directed), alpha 1, catalytic subunit   0.001
2769114916353402
KEGG:01100: Metabolic pathways  POLA2   polymerase (DNA directed), alpha 2, accessory subunit   -0.00
1498436249890922
KEGG:01100: Metabolic pathways  POLD1   polymerase (DNA directed), delta 1, catalytic subunit   -0.00
044560935432407076
KEGG:01100: Metabolic pathways  POLD2   polymerase (DNA directed), delta 2, accessory subunit   0.000
7421477949456388
KEGG:01100: Metabolic pathways  POLD3   polymerase (DNA-directed), delta 3, accessory subunit   0.001
7466587412103055
KEGG:01100: Metabolic pathways  POLD4   polymerase (DNA-directed), delta 4, accessory subunit   -0.00
06409905849416635
KEGG:01100: Metabolic pathways  POLE    polymerase (DNA directed), epsilon, catalytic subunit   0.002
958138309852866
KEGG:01100: Metabolic pathways  POLE2   polymerase (DNA directed), epsilon 2, accessory subunit 0.001
3307107678158525
KEGG:01100: Metabolic pathways  POLE3   polymerase (DNA directed), epsilon 3, accessory subunit 0.000
8084517388967187
KEGG:01100: Metabolic pathways  POLR2D  polymerase (RNA) II (DNA directed) polypeptide D        0.001
8615916236063399
KEGG:01100: Metabolic pathways  PRIM1   primase, DNA, polypeptide 1 (49kDa)     0.0006376004836594553
KEGG:01100: Metabolic pathways  PRIM2   primase, DNA, polypeptide 2 (58kDa)     -0.000890332011945211
7
KEGG:01100: Metabolic pathways  PSAT1   phosphoserine aminotransferase 1        -0.000724130502531340
3
KEGG:01100: Metabolic pathways  PSPH    phosphoserine phosphatase       -0.0019408383378121971
KEGG:01100: Metabolic pathways  QDPR    quinoid dihydropteridine reductase      -0.002865829961782042
8
KEGG:01100: Metabolic pathways  RRM1    ribonucleotide reductase M1     0.0019181942335011165
KEGG:01100: Metabolic pathways  RRM2    ribonucleotide reductase M2     -0.0005663338491549913
KEGG:01100: Metabolic pathways  SHMT2   serine hydroxymethyltransferase 2 (mitochondrial)       -0.00
11001374615877283
KEGG:01100: Metabolic pathways  SORD    sorbitol dehydrogenase  -0.002027239084487823
KEGG:01100: Metabolic pathways  SPAM1   sperm adhesion molecule 1 (PH-20 hyaluronidase, zona pellucid
a binding)      -0.0019057089696524509
KEGG:01100: Metabolic pathways  SPHK1   sphingosine kinase 1    0.001817463641216558
KEGG:01100: Metabolic pathways  SPR     sepiapterin reductase (7,8-dihydrobiopterin:NADP+ oxidoreduct
ase)    -0.0012378419013538608
KEGG:01100: Metabolic pathways  TAT     tyrosine aminotransferase       0.0010569571988997722
KEGG:01100: Metabolic pathways  TH      tyrosine hydroxylase    -0.00034287296760169455
KEGG:01100: Metabolic pathways  TYMS    thymidylate synthetase  0.0015685929022796513
KEGG:01100: Metabolic pathways  UGCG    UDP-glucose ceramide glucosyltransferase        0.00036935096
319734475
KEGG:01100: Metabolic pathways  UGDH    UDP-glucose 6-dehydrogenase     0.0005025506410051844
KEGG:01100: Metabolic pathways  UGP2    UDP-glucose pyrophosphorylase 2 0.0007901636826903181
KEGG:01100: Metabolic pathways  UGT8    UDP glycosyltransferase 8       0.002404617422909584
KEGG:00250: Alanine, aspartate and glutamate metabolism ABAT    4-aminobutyrate aminotransferase
0.0008330016098156056
KEGG:00250: Alanine, aspartate and glutamate metabolism ALDH5A1 aldehyde dehydrogenase 5 family, memb
er A1   0.002328028914348744
KEGG:00250: Alanine, aspartate and glutamate metabolism ASS1    argininosuccinate synthase 1    0.000
14221332941440813
KEGG:00250: Alanine, aspartate and glutamate metabolism GLUD2   glutamate dehydrogenase 2       -0.00
06588423570298238
KEGG:00640: Propanoate metabolism       ABAT    4-aminobutyrate aminotransferase        0.00083838480
57490674
KEGG:00640: Propanoate metabolism       LDHC    lactate dehydrogenase C 0.0014929699399719597
KEGG:00640: Propanoate metabolism       MUT     methylmalonyl CoA mutase        0.0027741889752544036
KEGG:00410: beta-Alanine metabolism     ABAT    4-aminobutyrate aminotransferase        0.00082173624
43510854

KEGG:00650: Butanoate metabolism        ABAT    4-aminobutyrate aminotransferase        0.00083427856
86320111
KEGG:00650: Butanoate metabolism        ALDH5A1 aldehyde dehydrogenase 5 family, member A1      0.002
3314907318213746
KEGG:00650: Butanoate metabolism        BDH1    3-hydroxybutyrate dehydrogenase, type 1 -0.0002073436
3851731552
KEGG:00650: Butanoate metabolism        HMGCL   3-hydroxymethyl-3-methylglutaryl-CoA lyase      -0.00
1064392202790947
GO:0001666: response to hypoxia ABAT    4-aminobutyrate aminotransferase        0.0008334434534379153
GO:0001666: response to hypoxia ADM     adrenomedullin  0.002238812559897347
GO:0001666: response to hypoxia AGER    advanced glycosylation end product-specific receptor    -0.00
01742645423739692
GO:0001666: response to hypoxia BNIP3   BCL2/adenovirus E1B 19kDa interacting protein 3 0.00291415084
3654701
GO:0001666: response to hypoxia CAV1    caveolin 1, caveolae protein, 22kDa     -0.000535471952848709
9
GO:0001666: response to hypoxia CCL2    chemokine (C-C motif) ligand 2  0.0008172213365582933
GO:0001666: response to hypoxia CD24    CD24 molecule   0.0010725532147139423
GO:0001666: response to hypoxia CHRNB2  cholinergic receptor, nicotinic, beta 2 (neuronal)      -0.00
07788010520266243
GO:0001666: response to hypoxia CRYAB   crystallin, alpha B     0.0009855156589479086
GO:0001666: response to hypoxia CST3    cystatin C      -7.084825260829509e-5
GO:0001666: response to hypoxia CX3CL1  chemokine (C-X3-C motif) ligand 1       0.002118492984099296
GO:0001666: response to hypoxia CXCL12  chemokine (C-X-C motif) ligand 12       -0.001209203588890177
8
GO:0001666: response to hypoxia CXCR4   chemokine (C-X-C motif) receptor 4      0.0007954308332594612
GO:0001666: response to hypoxia DRD2    dopamine receptor D2    -0.00023411324962894844
GO:0001666: response to hypoxia ENG     endoglin        0.0008271176427567826
GO:0001666: response to hypoxia HIF1A   hypoxia inducible factor 1, alpha subunit (basic helix-loop-h
elix transcription factor)      -0.0006464543407707821
GO:0001666: response to hypoxia ITPR1   inositol 1,4,5-trisphosphate receptor, type 1   -0.0007812013
655777396
GO:0001666: response to hypoxia KCNMA1  potassium large conductance calcium-activated channel, subfam
ily M, alpha member 1   -0.0006595603547464593
GO:0001666: response to hypoxia LEP     leptin  0.0031927134861560623
GO:0001666: response to hypoxia MMP2    matrix metallopeptidase 2 (gelatinase A, 72kDa gelatinase, 72
kDa type IV collagenase)        -0.0011323873096091154
GO:0001666: response to hypoxia PML     promyelocytic leukemia  -0.0006798120904419597
GO:0001666: response to hypoxia RAF1    Raf-1 proto-oncogene, serine/threonine kinase   0.00149944125
90703223
GO:0001666: response to hypoxia SLC11A2 solute carrier family 11 (proton-coupled divalent metal ion t
ransporter), member 2   -0.0006653504084603696
GO:0001666: response to hypoxia TFRC    transferrin receptor    0.0010849330467224139
GO:0001666: response to hypoxia TGFB1   transforming growth factor, beta 1      -7.324269878783433e-5
GO:0001666: response to hypoxia TGFB2   transforming growth factor, beta 2      -0.001059988161415871
1
GO:0001666: response to hypoxia TGFB3   transforming growth factor, beta 3      -0.001825167790215568
7
GO:0001666: response to hypoxia TH      tyrosine hydroxylase    -0.00034212341489976434
GO:0001666: response to hypoxia THBS1   thrombospondin 1        -0.001036550292497587
GO:0001666: response to hypoxia TXN2    thioredoxin 2   0.0013835636435942488
GO:0001666: response to hypoxia VEGFA   vascular endothelial growth factor A    0.0005957623131915809
GO:0007268: synaptic transmission       ABAT    4-aminobutyrate aminotransferase        0.00083441285
82778238
GO:0007268: synaptic transmission       ALDH5A1 aldehyde dehydrogenase 5 family, member A1      0.002
325736785758144
GO:0007268: synaptic transmission       ASIC2   acid-sensing (proton-gated) ion channel 2       0.002
2770056248329206
GO:0007268: synaptic transmission       BCHE    butyrylcholinesterase   -5.891714260375113e-6
GO:0007268: synaptic transmission       CACNB3  calcium channel, voltage-dependent, beta 3 subunit
0.00021248498616726832
GO:0007268: synaptic transmission       CHRNB2  cholinergic receptor, nicotinic, beta 2 (neuronal)
-0.0007771671025138414
GO:0007268: synaptic transmission       CREB1   cAMP responsive element binding protein 1       0.000
6560337387386202
GO:0007268: synaptic transmission       CTNNB1  catenin (cadherin-associated protein), beta 1, 88kDa
-0.00011771406287013872
GO:0007268: synaptic transmission       DRD4    dopamine receptor D4    -0.001942610174312302
GO:0007268: synaptic transmission       GABRA4  gamma-aminobutyric acid (GABA) A receptor, alpha 4
-0.0014819946432346228
GO:0007268: synaptic transmission       GRIK5   glutamate receptor, ionotropic, kainate 5       -0.00
023303218514897286
GO:0007268: synaptic transmission       HTR2C   5-hydroxytryptamine (serotonin) receptor 2C, G protei
n-coupled       -0.0005243373591696025
GO:0007268: synaptic transmission       HTR6    5-hydroxytryptamine (serotonin) receptor 6, G protein
-coupled        -0.0010808645611208802
GO:0007268: synaptic transmission       KCNJ2   potassium inwardly-rectifying channel, subfamily J, m
ember 2 -0.000775720653809918
GO:0007268: synaptic transmission       KCNMA1  potassium large conductance calcium-activated channe
l, subfamily M, alpha member 1  -0.0006552123640800498
GO:0007268: synaptic transmission       KCNV2   potassium channel, subfamily V, member 2        0.000

92502378924598
GO:0007268: synaptic transmission LRP6 low density lipoprotein receptor-related protein 6 0.00014822359348205612
GO:0007268: synaptic transmission PICK1 protein interacting with PRKCA 1 -0.0009541127470417695
GO:0007268: synaptic transmission PLCB1 phospholipase C, beta 1 (phosphoinositide-specific) 0.00015258877278244228
GO:0007268: synaptic transmission PRKCA protein kinase C, alpha -6.141223100982468e-6
GO:0007268: synaptic transmission RAF1 Raf-1 proto-oncogene, serine/threonine kinase 0.001496415240446442
GO:0007268: synaptic transmission RPS6KA1 ribosomal protein S6 kinase, 90kDa, polypeptide 1 -0.0025108737549834137
GO:0007268: synaptic transmission SLC1A1 solute carrier family 1 (neuronal/epithelial high affinity glutamate transporter, system Xag), member 1 -0.0008991927780949922
GO:0007268: synaptic transmission SLC1A3 solute carrier family 1 (glial high affinity glutamate transporter), member 3 -0.0031174552126335183
GO:0007268: synaptic transmission SLC5A7 solute carrier family 5 (sodium/choline cotransporter), member 7 0.0020096060653920556
GO:0007268: synaptic transmission SYT1 synaptotagmin I -0.0014400204030124307
GO:0007269: neurotransmitter secretion ABAT 4-aminobutyrate aminotransferase 0.0008382109591487198
GO:0007269: neurotransmitter secretion ALDH5A1 aldehyde dehydrogenase 5 family, member A1 0.0023423281457344222
GO:0007269: neurotransmitter secretion SLC1A1 solute carrier family 1 (neuronal/epithelial high affinity glutamate transporter, system Xag), member 1 -0.0009061343508545292
GO:0007269: neurotransmitter secretion SLC1A3 solute carrier family 1 (glial high affinity glutamate transporter), member 3 -0.003136269992870771
GO:0007269: neurotransmitter secretion SLC5A7 solute carrier family 5 (sodium/choline cotransporter), member 7 0.0020197735643882743
GO:0007269: neurotransmitter secretion SYT1 synaptotagmin I -0.001451872580013471
GO:0007269: neurotransmitter secretion WNT7A wingless-type MMTV integration site family, member 7A 1.9218868493483478e-5
GO:0007620: copulation ABAT 4-aminobutyrate aminotransferase 0.000830649023831608
GO:0007620: copulation PI3 peptidase inhibitor 3, skin-derived -0.0010312990109310948
GO:0007626: locomotory behavior ABAT 4-aminobutyrate aminotransferase 0.0008188020442316773
GO:0007626: locomotory behavior AVP arginine vasopressin -0.000911917202172883
GO:0007626: locomotory behavior CHRNB2 cholinergic receptor, nicotinic, beta 2 (neuronal) -0.0007604539753433605
GO:0007626: locomotory behavior DRD2 dopamine receptor D2 -0.00023041388671371225
GO:0007626: locomotory behavior HPRT1 hypoxanthine phosphoribosyltransferase 1 -0.0011642187897423887
GO:0007626: locomotory behavior HTR2C 5-hydroxytryptamine (serotonin) receptor 2C, G protein-coupled -0.0005148148706419103
GO:0007626: locomotory behavior HTT huntingtin -0.0009518194507996049
GO:0007626: locomotory behavior PTEN phosphatase and tensin homolog 2.4547602115369354e-5
GO:0007626: locomotory behavior TH tyrosine hydroxylase -0.0003377675693174901
GO:0009449: gamma-aminobutyric acid biosynthetic process ABAT 4-aminobutyrate aminotransferase 0.0008414548467394141
GO:0009449: gamma-aminobutyric acid biosynthetic process SLC1A3 solute carrier family 1 (glial high affinity glutamate transporter), member 3 -0.00314633963443901
GO:0009450: gamma-aminobutyric acid catabolic process ABAT 4-aminobutyrate aminotransferase 0.0008384285830425878
GO:0009450: gamma-aminobutyric acid catabolic process ALDH5A1 aldehyde dehydrogenase 5 family, member A1 0.002345277617385604
GO:0010039: response to iron ion ABAT 4-aminobutyrate aminotransferase 0.0008341486169944331
GO:0010039: response to iron ion BCL2 B-cell CLL/lymphoma 2 -5.314774669704326e-6
GO:0010039: response to iron ion CCND1 cyclin D1 -0.002640764841484498
GO:0010039: response to iron ion CPOX coproporphyrinogen oxidase 0.0012771007140909795
GO:0010039: response to iron ion DRD2 dopamine receptor D2 -0.0002343433759645894
GO:0010039: response to iron ion SLC11A2 solute carrier family 11 (proton-coupled divalent metal ion transporter), member 2 -0.0006660535507818989
GO:0010039: response to iron ion TFRC transferrin receptor 0.0010863007139147046
GO:0031652: positive regulation of heat generation ABAT 4-aminobutyrate aminotransferase 0.0008217362443510854
GO:0032024: positive regulation of insulin secretion ABAT 4-aminobutyrate aminotransferase 0.0008012149650160562
GO:0032024: positive regulation of insulin secretion GJA1 gap junction protein, alpha 1, 43kDa -0.00014157107097421834
GO:0032024: positive regulation of insulin secretion ISL1 ISL LIM homeobox 1 8.229230395477169e-5
GO:0032024: positive regulation of insulin secretion JAK2 Janus kinase 2 -3.731858130527175e-5
GO:0032024: positive regulation of insulin secretion SOX4 SRY (sex determining region Y)-box 4 -4.3476389792572435e-5
GO:0032024: positive regulation of insulin secretion TCF7L2 transcription factor 7-like 2 (T-cell specific, HMG-box) 0.0005478985119265809
GO:0035094: response to nicotine ABAT 4-aminobutyrate aminotransferase 0.0008162557589217794
GO:0035094: response to nicotine AVP arginine vasopressin -0.0009092874589890048
GO:0035094: response to nicotine BCL2 B-cell CLL/lymphoma 2 -5.086636827417199e-6
GO:0035094: response to nicotine CHRNB2 cholinergic receptor, nicotinic, beta 2 (neuronal)

-0.0007583278132899442
GO:0035094: response to nicotine DRD2 dopamine receptor D2 -0.00023001114993561662
GO:0035094: response to nicotine GPX1 glutathione peroxidase 1 0.0003867305456564967
7
GO:0035094: response to nicotine HDAC2 histone deacetylase 2 -0.0011885414992245903
GO:0035094: response to nicotine HMOX1 heme oxygenase (decycling) 1 -0.000210711403920738
34
GO:0035094: response to nicotine PDX1 pancreatic and duodenal homeobox 1 0.00024693048
724199
GO:0035094: response to nicotine STAR steroidogenic acute regulatory protein 0.00086922330
19067949
GO:0042135: neurotransmitter catabolic process ABAT 4-aminobutyrate aminotransferase 0.000
8384285830425878
GO:0042135: neurotransmitter catabolic process ALDH5A1 aldehyde dehydrogenase 5 family, member A1
0.002345277617385604
GO:0042493: response to drug ABAT 4-aminobutyrate aminotransferase 0.0008321786943666631
GO:0042493: response to drug AOC1 amine oxidase, copper containing 1 -0.001420914896464388
5
GO:0042493: response to drug APOD apolipoprotein D 0.0026237076376514645
GO:0042493: response to drug ASS1 argininosuccinate synthase 1 0.00014194170651174364
GO:0042493: response to drug BAK1 BCL2-antagonist/killer 1 -0.0018729836222525886
GO:0042493: response to drug BAX BCL2-associated X protein -0.00042458025033647966
GO:0042493: response to drug BCHE butyrylcholinesterase -6.491329207779307e-6
GO:0042493: response to drug BCL2 B-cell CLL/lymphoma 2 -5.133173754918806e-6
GO:0042493: response to drug CAV1 caveolin 1, caveolae protein, 22kDa -0.000534839011063478
GO:0042493: response to drug CCNB1 cyclin B1 -0.0008812927404760949
GO:0042493: response to drug CCND1 cyclin D1 -0.002627638068207084
GO:0042493: response to drug CCNE1 cyclin E1 0.00037022802307575036
GO:0042493: response to drug CDH3 cadherin 3, type 1, P-cadherin (placental) -0.0012289660
774525374
GO:0042493: response to drug COL1A1 collagen, type I, alpha 1 -0.0005247261735010596
GO:0042493: response to drug CPT1A carnitine palmitoyltransferase 1A (liver) 0.00127106991
20922076
GO:0042493: response to drug CREB1 cAMP responsive element binding protein 1 0.00065678352
91725943
GO:0042493: response to drug CST3 cystatin C -6.962448601715963e-5
GO:0042493: response to drug CTNNB1 catenin (cadherin-associated protein), beta 1, 88kDa -0.00
011684991017535412
GO:0042493: response to drug CTPS1 CTP synthase 1 -0.0004206897806356908
GO:0042493: response to drug DRD2 dopamine receptor D2 -0.00023377312726287962
GO:0042493: response to drug FGF8 fibroblast growth factor 8 (androgen-induced) 0.00098204867
6332739
GO:0042493: response to drug GABRA4 gamma-aminobutyric acid (GABA) A receptor, alpha 4 -0.00
14818892879008391
GO:0042493: response to drug GATA3 GATA binding protein 3 -3.884231995012516e-5
GO:0042493: response to drug GATA4 GATA binding protein 4 -0.0010929060562918146
GO:0042493: response to drug GATA6 GATA binding protein 6 -2.8269350521338365e-5
GO:0042493: response to drug GCLM glutamate-cysteine ligase, modifier subunit -0.0037559422
30585455
GO:0042493: response to drug GGH gamma-glutamyl hydrolase (conjugase, folylpolygammaglutamyl h
ydrolase) -0.00031443847416488525
GO:0042493: response to drug HDAC2 histone deacetylase 2 -0.001205678237829658
GO:0042493: response to drug HMGB2 high mobility group box 2 0.000305739048590344
GO:0042493: response to drug HTR2C 5-hydroxytryptamine (serotonin) receptor 2C, G protein-couple
d -0.000523984514707995
GO:0042493: response to drug ICAM1 intercellular adhesion molecule 1 0.0007285199041821114
GO:0042493: response to drug IFNG interferon, gamma -4.8881451727045474e-5
GO:0042493: response to drug IGFBP2 insulin-like growth factor binding protein 2, 36kDa 0.000
1443812893048011
GO:0042493: response to drug IL4 interleukin 4 0.0002594464935783002
GO:0042493: response to drug INHBA inhibin, beta A -0.0013498319703095444
GO:0042493: response to drug ITGA3 integrin, alpha 3 (antigen CD49C, alpha 3 subunit of VLA-3 re
ceptor) 0.001397666933609066
GO:0042493: response to drug LOX lysyl oxidase -0.0005752529576941596
GO:0042493: response to drug LRP8 low density lipoprotein receptor-related protein 8, apolipopr
otein e receptor -0.0009304291800217338
GO:0042493: response to drug LYN LYN proto-oncogene, Src family tyrosine kinase -0.0014181651
652112257
GO:0042493: response to drug MAS1 MAS1 proto-oncogene, G protein-coupled receptor 0.00029183920
086953526
GO:0042493: response to drug MCM7 minichromosome maintenance complex component 7 -0.0016415981
644914998
GO:0042493: response to drug MDK midkine (neurite growth-promoting factor 2) 0.00166141231
98063545
GO:0042493: response to drug MGMT O-6-methylguanine-DNA methyltransferase 0.0004062990333310252
7
GO:0042493: response to drug MMP7 matrix metallopeptidase 7 (matrilysin, uterine) 0.00042493725
485866186
GO:0042493: response to drug MYC v-myc avian myelocytomatosis viral oncogene homolog -0.00
11347891996923505
GO:0042493: response to drug PDX1 pancreatic and duodenal homeobox 1 0.0002566008581639897

4
GO:0042493: response to drug    POR     P450 (cytochrome) oxidoreductase        0.0004564298553459019
GO:0042493: response to drug    PTCH1   patched 1       -6.604529345629732e-5
GO:0042493: response to drug    PTEN    phosphatase and tensin homolog  1.753387340373094e-5
GO:0042493: response to drug    PTN     pleiotrophin    0.0003004802981246495
GO:0042493: response to drug    RAD51   RAD51 recombinase       -0.0018583682484856418
GO:0042493: response to drug    RET     ret proto-oncogene      -0.0004905437144619259
GO:0042493: response to drug    SEMA3C  sema domain, immunoglobulin domain (Ig), short basic domain, secreted, (semaphorin) 3C       -0.0002209842248549008
GO:0042493: response to drug    SFRP1   secreted frizzled-related protein 1     0.0012784513667955754
GO:0042493: response to drug    SLC1A3  solute carrier family 1 (glial high affinity glutamate transporter), member 3        -0.0031160119825241386
GO:0042493: response to drug    SORD    sorbitol dehydrogenase  -0.0020159685579222656
GO:0042493: response to drug    SRD5A2  steroid-5-alpha-reductase, alpha polypeptide 2 (3-oxo-5 alpha-steroid delta 4-dehydrogenase alpha 2) -0.000641781192456762
GO:0042493: response to drug    STAR    steroidogenic acute regulatory protein  0.000892818398124577
GO:0042493: response to drug    TGFB1   transforming growth factor, beta 1      -7.280948934419745e-5
GO:0042493: response to drug    TGFB2   transforming growth factor, beta 2      -0.0010573617249044031
GO:0042493: response to drug    THBS1   thrombospondin 1        -0.0010341382726313892
GO:0042493: response to drug    THRA    thyroid hormone receptor, alpha 0.000762019914216386
GO:0042493: response to drug    TP73    tumor protein p73       0.0010261516026832198
GO:0042493: response to drug    TXN2    thioredoxin 2   0.0013811834239350366
GO:0042493: response to drug    TYMS    thymidylate synthetase  0.0015582612076874417
GO:0042493: response to drug    VEGFC   vascular endothelial growth factor C    -0.0033586098501147362
GO:0042493: response to drug    VLDLR   very low density lipoprotein receptor   0.0009510405671015782
GO:0042493: response to drug    XBP1    X-box binding protein 1 0.00026695250118882237
GO:0045471: response to ethanol ABAT    4-aminobutyrate aminotransferase        0.0008299947113849659
GO:0045471: response to ethanol AVP     arginine vasopressin    -0.0009337376715532955
GO:0045471: response to ethanol BAK1    BCL2-antagonist/killer 1        -0.0018691442589787509
GO:0045471: response to ethanol CCL2    chemokine (C-C motif) ligand 2  0.000813954396195279
GO:0045471: response to ethanol CCND1   cyclin D1       -0.0026223255119882023
GO:0045471: response to ethanol CCNE1   cyclin E1       0.00036987332070646903
GO:0045471: response to ethanol CHRNB2  cholinergic receptor, nicotinic, beta 2 (neuronal)      -0.0007754511338242427
GO:0045471: response to ethanol EGR1    early growth response 1 0.0010891728556308687
GO:0045471: response to ethanol GATA3   GATA binding protein 3  -3.929473943456721e-5
GO:0045471: response to ethanol GGH     gamma-glutamyl hydrolase (conjugase, folylpolygammaglutamyl hydrolase)       -0.00031353561000826134
GO:0045471: response to ethanol ICAM1   intercellular adhesion molecule 1       0.000727369329315799
GO:0045471: response to ethanol IL4     interleukin 4   0.00025901329791812745
GO:0045471: response to ethanol MGMT    O-6-methylguanine-DNA methyltransferase 0.0004056809053482333
GO:0045471: response to ethanol PTEN    phosphatase and tensin homolog  1.7586515911126337e-5
GO:0045471: response to ethanol RXRA    retinoid X receptor, alpha      0.001110996827210531
GO:0045471: response to ethanol STAR    steroidogenic acute regulatory protein  0.0008910161245458175
GO:0045471: response to ethanol TH      tyrosine hydroxylase    -0.0003412037158551921
GO:0045471: response to ethanol TYMS    thymidylate synthetase  0.0015551981636797304
GO:0045776: negative regulation of blood pressure       ABAT    4-aminobutyrate aminotransferase 0.0008395157508078793
GO:0045776: negative regulation of blood pressure       DRD2    dopamine receptor D2    -0.00023535337223040117
GO:0045776: negative regulation of blood pressure       VEGFC   vascular endothelial growth factor C -0.0033898641162820272
GO:0048148: behavioral response to cocaine      ABAT    4-aminobutyrate aminotransferase        0.0008314822091479581
GO:0048148: behavioral response to cocaine      DRD2    dopamine receptor D2    -0.000232865389467014
GO:0048148: behavioral response to cocaine      DRD4    dopamine receptor D4    -0.0019342023186998322
KEGG:02010: ABC transporters    ABCG1   ATP-binding cassette, sub-family G (WHITE), member 1    -0.0023606226924747856
GO:0006355: regulation of transcription, DNA-templated  ABCG1   ATP-binding cassette, sub-family G (WHITE), member 1 -0.002341665434571339
GO:0006355: regulation of transcription, DNA-templated  ACVRL1  activin A receptor type II-like 1 0.0019461451263920158
GO:0006355: regulation of transcription, DNA-templated  APEX1   APEX nuclease (multifunctional DNA repair enzyme) 1  0.0002001206630484574
GO:0006355: regulation of transcription, DNA-templated  ATF5    activating transcription factor 5 -0.002738417394466573
GO:0006355: regulation of transcription, DNA-templated  BACH1   BTB and CNC homology 1, basic leucine zipper transcription factor 1   0.0004858616370202321
GO:0006355: regulation of transcription, DNA-templated  CASP8AP2        caspase 8 associated protein 2       0.0012735027160679787
GO:0006355: regulation of transcription, DNA-templated  CEBPB   CCAAT/enhancer binding protein (C/EBP), beta        -0.0002745449759790139
GO:0006355: regulation of transcription, DNA-templated  CELSR2  cadherin, EGF LAG seven-pass G-type receptor 2       -0.0011360981432418866
GO:0006355: regulation of transcription, DNA-templated  CHAF1B  chromatin assembly factor 1, subunit B (p60) 0.006292511066352083
GO:0006355: regulation of transcription, DNA-templated  CHD3    chromodomain helicase DNA binding protein 3  0.00011917125375918135

GO:0006355: regulation of transcription, DNA-templated CLOCK clock circadian regulator 0.000
20035738705752545
GO:0006355: regulation of transcription, DNA-templated DAXX death-domain associated protein 0.000
8926960114535764
GO:0006355: regulation of transcription, DNA-templated DLX6 distal-less homeobox 6 -0.0001800747
707105529
GO:0006355: regulation of transcription, DNA-templated E2F1 E2F transcription factor 1 0.002
1001730920734312
GO:0006355: regulation of transcription, DNA-templated ENG endoglin 0.0008277045008275172
GO:0006355: regulation of transcription, DNA-templated ESR1 estrogen receptor 1 -0.0009490519
211270113
GO:0006355: regulation of transcription, DNA-templated EYA3 EYA transcriptional coactivator and p
hosphatase 3 0.0008375143974783718
GO:0006355: regulation of transcription, DNA-templated EZH2 enhancer of zeste 2 polycomb repressi
ve complex 2 subunit -0.00013690751833421626
GO:0006355: regulation of transcription, DNA-templated FOXC1 forkhead box C1 -2.2551636859671693e-
5
GO:0006355: regulation of transcription, DNA-templated FOXM1 forkhead box M1 0.0001971198857297299
8
GO:0006355: regulation of transcription, DNA-templated FZD7 frizzled class receptor 7 0.001
0715426643732194
GO:0006355: regulation of transcription, DNA-templated GATA4 GATA binding protein 4 -0.0010960469
90329257
GO:0006355: regulation of transcription, DNA-templated GLRX2 glutaredoxin 2 0.0011173084831466254
GO:0006355: regulation of transcription, DNA-templated HIF1A hypoxia inducible factor 1, alpha sub
unit (basic helix-loop-helix transcription factor) -0.0006473802200420288
GO:0006355: regulation of transcription, DNA-templated HINFP histone H4 transcription factor 0.000
9935293433366505
GO:0006355: regulation of transcription, DNA-templated HMGA1 high mobility group AT-hook 1 -0.00
031929834408582215
GO:0006355: regulation of transcription, DNA-templated HMGA2 high mobility group AT-hook 2 0.001
5108935737475357
GO:0006355: regulation of transcription, DNA-templated HMGB3 high mobility group box 3 0.000
46420608836882825
GO:0006355: regulation of transcription, DNA-templated HOXA3 homeobox A3 0.0009898676244079938
GO:0006355: regulation of transcription, DNA-templated HOXB1 homeobox B1 0.003690960254605468
GO:0006355: regulation of transcription, DNA-templated HOXB13 homeobox B13 0.0018279178412547708
GO:0006355: regulation of transcription, DNA-templated HOXB2 homeobox B2 0.0027778308018834626
GO:0006355: regulation of transcription, DNA-templated HOXD13 homeobox D13 -0.000593175132816979
9
GO:0006355: regulation of transcription, DNA-templated INSR insulin receptor -0.0013648946
786685415
GO:0006355: regulation of transcription, DNA-templated JMJD6 jumonji domain containing 6 0.003
595176943406386
GO:0006355: regulation of transcription, DNA-templated KANK1 KN motif and ankyrin repeat domains 1
0.002536265891080383
GO:0006355: regulation of transcription, DNA-templated KMT2D lysine (K)-specific methyltransferase
2D -0.0001995535186087161
GO:0006355: regulation of transcription, DNA-templated LHX6 LIM homeobox 6 0.0013455375779938114
GO:0006355: regulation of transcription, DNA-templated MEF2C myocyte enhancer factor 2C 0.000
9745956786313343
GO:0006355: regulation of transcription, DNA-templated MTERF1 mitochondrial transcription terminati
on factor 1 0.00028655484865196853
GO:0006355: regulation of transcription, DNA-templated NEUROD4 neuronal differentiation 4 0.001
5087356138340667
GO:0006355: regulation of transcription, DNA-templated NOTCH1 notch 1 0.0005192910047033064
GO:0006355: regulation of transcription, DNA-templated NRBF2 nuclear receptor binding factor 2
-0.0009522439220127656
GO:0006355: regulation of transcription, DNA-templated PADI4 peptidyl arginine deiminase, type IV
-0.0008276419108376789
GO:0006355: regulation of transcription, DNA-templated PITX2 paired-like homeodomain 2 0.002
178824277367717
GO:0006355: regulation of transcription, DNA-templated PML promyelocytic leukemia -0.0006808680
696586667
GO:0006355: regulation of transcription, DNA-templated POU1F1 POU class 1 homeobox 1 0.00013762929
217908887
GO:0006355: regulation of transcription, DNA-templated PRDM4 PR domain containing 4 0.00084140848
90226066
GO:0006355: regulation of transcription, DNA-templated PTTG1 pituitary tumor-transforming 1 -0.00
0362322343916256
GO:0006355: regulation of transcription, DNA-templated PTTG2 pituitary tumor-transforming 2 -0.00
17268107379317722
GO:0006355: regulation of transcription, DNA-templated PTTG3P pituitary tumor-transforming 3, pseud
ogene -4.0391214768816236e-5
GO:0006355: regulation of transcription, DNA-templated RBBP4 retinoblastoma binding protein 4
-0.0018629889162687448
GO:0006355: regulation of transcription, DNA-templated RFC1 replication factor C (activator 1) 1,
145kDa 0.0010605418258528654
GO:0006355: regulation of transcription, DNA-templated RREB1 ras responsive element binding protei
n 1 -0.00198407108758362
GO:0006355: regulation of transcription, DNA-templated SIX1 SIX homeobox 1 -0.00187227330062934

3
GO:0006355: regulation of transcription, DNA-templated SOX4 SRY (sex determining region Y)-box 4
-3.847491638180251e-5
GO:0006355: regulation of transcription, DNA-templated TBPL1 TBP-like 1 -0.000789896881277378
3
GO:0006355: regulation of transcription, DNA-templated TBX21 T-box 21 0.0003990177683590808
3
GO:0006355: regulation of transcription, DNA-templated TGFBR1 transforming growth factor, beta rece
ptor 1 0.00034029091337871563
GO:0006355: regulation of transcription, DNA-templated TP53 tumor protein p53 0.00117711530
16190811
GO:0006355: regulation of transcription, DNA-templated TULP3 tubby like protein 3 0.00095745576
78974636
GO:0006355: regulation of transcription, DNA-templated TXLNG taxilin gamma -0.000838522066702498
5
GO:0006355: regulation of transcription, DNA-templated WT1 Wilms tumor 1 -0.000507551555832777
9
GO:0006355: regulation of transcription, DNA-templated WWTR1 WW domain containing transcription re
gulator 1 0.0009013744323481756
GO:0006355: regulation of transcription, DNA-templated YBX1 Y box binding protein 1 -0.0009151955
467048587
GO:0006355: regulation of transcription, DNA-templated ZFP36L2 ZFP36 ring finger protein-like 2
-0.0003633828668500567
GO:0006355: regulation of transcription, DNA-templated ZNF205 zinc finger protein 205 0.00059682523
10036016
GO:0006355: regulation of transcription, DNA-templated ZNF207 zinc finger protein 207 -0.0003898340
106473997
GO:0006355: regulation of transcription, DNA-templated ZNF266 zinc finger protein 266 0.00041304641
85346677
GO:0006355: regulation of transcription, DNA-templated ZNF335 zinc finger protein 335 -0.0003652254
1044257413
GO:0006355: regulation of transcription, DNA-templated ZNF442 zinc finger protein 442 0.00105958638
9757607
GO:0008203: cholesterol metabolic process ABCG1 ATP-binding cassette, sub-family G (WHITE), m
ember 1 -0.002345487810176392
GO:0008203: cholesterol metabolic process CYP7B1 cytochrome P450, family 7, subfamily B, polyp
eptide 1 0.00016361504466810508
GO:0008203: cholesterol metabolic process IL4 interleukin 4 0.00025965180694784943
GO:0008203: cholesterol metabolic process LEP leptin 0.0032021505124831787
GO:0008203: cholesterol metabolic process LRP5 low density lipoprotein receptor-related prot
ein 5 2.981304340339902e-5
GO:0008203: cholesterol metabolic process RXRA retinoid X receptor, alpha 0.00111843405
90287424
GO:0008203: cholesterol metabolic process SOAT1 sterol O-acyltransferase 1 -0.0004083545
7833197046
GO:0008203: cholesterol metabolic process STAR steroidogenic acute regulatory protein 0.000
897649972770958
GO:0008203: cholesterol metabolic process VLDLR very low density lipoprotein receptor 0.000
9564620301798084
GO:0009720: detection of hormone stimulus ABCG1 ATP-binding cassette, sub-family G (WHITE), m
ember 1 -0.0023606226924747856
GO:0010033: response to organic substance ABCG1 ATP-binding cassette, sub-family G (WHITE), m
ember 1 -0.0023330672278115442
GO:0010033: response to organic substance CREB1 cAMP responsive element binding protein 1
0.0006548028848150214
GO:0010033: response to organic substance CRIP1 cysteine-rich protein 1 (intestinal) 0.001
720436410130986
GO:0010033: response to organic substance GLRX2 glutaredoxin 2 0.001113492904762377
GO:0010033: response to organic substance MAPT microtubule-associated protein tau 0.001
5495420225335123
GO:0010033: response to organic substance SERPINF2 serpin peptidase inhibitor, clade F
(alpha-2 antiplasmin, pigment epithelium derived factor), member 2 0.0006573269261983929
GO:0010033: response to organic substance WNT5A wingless-type MMTV integration site family, m
ember 5A -0.0006650579785260218
GO:0010745: negative regulation of macrophage derived foam cell differentiation ABCG1 ATP-binding c
assette, sub-family G (WHITE), member 1 -0.0023466386573099125
GO:0010745: negative regulation of macrophage derived foam cell differentiation NR1H3 nuclear recep
tor subfamily 1, group H, member 3 0.0008032537504780686
GO:0010872: regulation of cholesterol esterification ABCG1 ATP-binding cassette, sub-family G (W
HITE), member 1 -0.0023606226924747856
GO:0010875: positive regulation of cholesterol efflux ABCG1 ATP-binding cassette, sub-family G (W
HITE), member 1 -0.002326271039840745
GO:0010875: positive regulation of cholesterol efflux NR1H3 nuclear receptor subfamily 1, group
H, member 3 0.000798170453867906
GO:0010875: positive regulation of cholesterol efflux PTCH1 patched 1 -6.733786411850906e-5
GO:0010875: positive regulation of cholesterol efflux SIRT1 sirtuin 1 -3.108596316558766e-6
GO:0010887: negative regulation of cholesterol storage ABCG1 ATP-binding cassette, sub-family G (W
HITE), member 1 -0.0023466386573099125
GO:0010887: negative regulation of cholesterol storage NR1H3 nuclear receptor subfamily 1, group
H, member 3 0.0008032537504780686
GO:0032367: intracellular cholesterol transport ABCG1 ATP-binding cassette, sub-family G (WHITE), m

ember 1 -0.0023414183670599836
GO:0032367: intracellular cholesterol transport STAR    steroidogenic acute regulatory protein  0.000
8955133587820498
GO:0032367: intracellular cholesterol transport VPS4A   vacuolar protein sorting 4 homolog A (S. cere
visiae) -0.00068578347082942
GO:0033344: cholesterol efflux  ABCG1   ATP-binding cassette, sub-family G (WHITE), member 1    -0.00
23351541238514502
GO:0033344: cholesterol efflux  CAV1    caveolin 1, caveolae protein, 22kDa     -0.000535137360978536
6
GO:0033344: cholesterol efflux  SOAT1   sterol O-acyltransferase 1      -0.00040815997631377283
GO:0033700: phospholipid efflux ABCG1   ATP-binding cassette, sub-family G (WHITE), member 1    -0.00
23606226924747856
GO:0033993: response to lipid   ABCG1   ATP-binding cassette, sub-family G (WHITE), member 1    -0.00
2348053988443847
GO:0033993: response to lipid   GATA2   GATA binding protein 2  -0.0004546861240076834
GO:0033993: response to lipid   PCNA    proliferating cell nuclear antigen      0.0012333088339832425
GO:0034374: low-density lipoprotein particle remodeling ABCG1   ATP-binding cassette, sub-family G (W
HITE), member 1 -0.0023477349653539255
GO:0034374: low-density lipoprotein particle remodeling AGT     angiotensinogen (serpin peptidase inh
ibitor, clade A, member 8)      -0.0011404141840943832
GO:0034374: low-density lipoprotein particle remodeling AGTR1   angiotensin II receptor, type 1 0.000
3465689823358322
GO:0034375: high-density lipoprotein particle remodeling        ABCG1   ATP-binding cassette, sub-fam
ily G (WHITE), member 1 -0.0023606226924747856
GO:0034436: glycoprotein transport      ABCG1   ATP-binding cassette, sub-family G (WHITE), member 1
-0.0023604555192209145
GO:0034436: glycoprotein transport      GUK1    guanylate kinase 1      0.0033018436125547713
GO:0034436: glycoprotein transport      VLDLR   very low density lipoprotein receptor   0.00096462165
33888693
GO:0042157: lipoprotein metabolic process       ABCG1   ATP-binding cassette, sub-family G (WHITE), m
ember 1 -0.0023606226924747856
GO:0042632: cholesterol homeostasis     ABCG1   ATP-binding cassette, sub-family G (WHITE), member 1
-0.002333236756042257
GO:0042632: cholesterol homeostasis     AKR1C1  aldo-keto reductase family 1, member C1 -0.0014868902
932814732
GO:0042632: cholesterol homeostasis     CAV1    caveolin 1, caveolae protein, 22kDa     -0.0005350171
271438002
GO:0042632: cholesterol homeostasis     CD24    CD24 molecule   0.0010673039896267873
GO:0042632: cholesterol homeostasis     HPN     hepsin  0.0031532344758925988
GO:0042632: cholesterol homeostasis     LRP5    low density lipoprotein receptor-related protein 5
3.0402724128309627e-5
GO:0042632: cholesterol homeostasis     NR1H3   nuclear receptor subfamily 1, group H, member 3 0.000
8000871231898231
GO:0042632: cholesterol homeostasis     SIRT1   sirtuin 1       -2.7065479982397913e-6
GO:0042632: cholesterol homeostasis     SOAT1   sterol O-acyltransferase 1      -0.000408650022457234
46
GO:0042632: cholesterol homeostasis     XBP1    X-box binding protein 1 0.00026859745969275557
GO:0042987: amyloid precursor protein catabolic process ABCG1   ATP-binding cassette, sub-family G (W
HITE), member 1 -0.002356926099937954
GO:0042987: amyloid precursor protein catabolic process DHCR24  24-dehydrocholesterol reductase -0.00
17218261808735022
GO:0043691: reverse cholesterol transport       ABCG1   ATP-binding cassette, sub-family G (WHITE), m
ember 1 -0.0023606226924747856
GO:0044281: small molecule metabolic process    ABCG1   ATP-binding cassette, sub-family G (WHITE), m
ember 1 -0.0023373006184539994
GO:0044281: small molecule metabolic process    ACADL   acyl-CoA dehydrogenase, long chain      0.000
9806184984740072
GO:0044281: small molecule metabolic process    ACOT8   acyl-CoA thioesterase 8 0.0011354676229892832
GO:0044281: small molecule metabolic process    AGT     angiotensinogen (serpin peptidase inhibitor,
clade A, member 8)      -0.0011316057979557496
GO:0044281: small molecule metabolic process    AKT1    v-akt murine thymoma viral oncogene homolog 1
0.0007299030112991716
GO:0044281: small molecule metabolic process    ALOX12  arachidonate 12-lipoxygenase    -0.0019519243
432385431
GO:0044281: small molecule metabolic process    AMD1    adenosylmethionine decarboxylase 1      -0.00
25271199410123514
GO:0044281: small molecule metabolic process    ASS1    argininosuccinate synthase 1    0.00014210037
478324937
GO:0044281: small molecule metabolic process    AUH     AU RNA binding protein/enoyl-CoA hydratase
-0.001460609543166981
GO:0044281: small molecule metabolic process    AZIN1   antizyme inhibitor 1    -0.000127118829897413
5
GO:0044281: small molecule metabolic process    BDH1    3-hydroxybutyrate dehydrogenase, type 1 -0.00
020721964152094292
GO:0044281: small molecule metabolic process    CAV1    caveolin 1, caveolae protein, 22kDa     -0.00
05352462335167767
GO:0044281: small molecule metabolic process    CBS     cystathionine-beta-synthase     0.00151552901
05719084
GO:0044281: small molecule metabolic process    CHST15  carbohydrate (N-acetylgalactosamine 4-sulfate
6-O) sulfotransferase 15        -0.0008251421276839758
GO:0044281: small molecule metabolic process    CPOX    coproporphyrinogen oxidase      0.00127175671

78886773
GO:0044281: small molecule metabolic process CPT1A carnitine palmitoyltransferase 1A (liver) 0.0012720760621700972
GO:0044281: small molecule metabolic process CROT carnitine O-octanoyltransferase -0.0004380009443825063
GO:0044281: small molecule metabolic process CTH cystathionine gamma-lyase 0.0017469343321310446
GO:0044281: small molecule metabolic process CTPS1 CTP synthase 1 -0.0004210446504633483
GO:0044281: small molecule metabolic process CTPS2 CTP synthase 2 0.001759035235166584
GO:0044281: small molecule metabolic process CYP1B1 cytochrome P450, family 1, subfamily B, polypeptide 1 0.00039755471279155895
GO:0044281: small molecule metabolic process CYP27B1 cytochrome P450, family 27, subfamily B, polypeptide 1 -0.0004704327577764955
GO:0044281: small molecule metabolic process CYP4B1 cytochrome P450, family 4, subfamily B, polypeptide 1 -0.0013549255428817466
GO:0044281: small molecule metabolic process CYP7B1 cytochrome P450, family 7, subfamily B, polypeptide 1 0.00016319722932243985
GO:0044281: small molecule metabolic process DCT dopachrome tautomerase -0.002351662469383146
5
GO:0044281: small molecule metabolic process DHCR24 24-dehydrocholesterol reductase -0.0017061457477303887
GO:0044281: small molecule metabolic process DHFR dihydrofolate reductase -0.00030787753765544417
GO:0044281: small molecule metabolic process DTYMK deoxythymidylate kinase (thymidylate kinase) 0.0034793847935863345
GO:0044281: small molecule metabolic process ELOVL2 ELOVL fatty acid elongase 2 0.0005244086502352604
GO:0044281: small molecule metabolic process ELOVL5 ELOVL fatty acid elongase 5 0.000206789833912712
GO:0044281: small molecule metabolic process GCDH glutaryl-CoA dehydrogenase -4.876941985395083e-6
GO:0044281: small molecule metabolic process GCHFR GTP cyclohydrolase I feedback regulator -0.0008418507162816584
GO:0044281: small molecule metabolic process GCLM glutamate-cysteine ligase, modifier subunit -0.003759066733413574
GO:0044281: small molecule metabolic process GMPS guanine monphosphate synthase 0.001772553998425755
GO:0044281: small molecule metabolic process GPD1L glycerol-3-phosphate dehydrogenase 1-like 0.0005669395518063695
GO:0044281: small molecule metabolic process GPX1 glutathione peroxidase 1 0.00038770796335673157
GO:0044281: small molecule metabolic process GSTM1 glutathione S-transferase mu 1 -0.0014730010201691572
GO:0044281: small molecule metabolic process GSTM2 glutathione S-transferase mu 2 (muscle) -0.001912820285543965
GO:0044281: small molecule metabolic process GSTM3 glutathione S-transferase mu 3 (brain) -0.0012880251258405968
GO:0044281: small molecule metabolic process GSTP1 glutathione S-transferase pi 1 0.00023126604079261746
GO:0044281: small molecule metabolic process GUK1 guanylate kinase 1 0.0032717822596017567
GO:0044281: small molecule metabolic process HMGCL 3-hydroxymethyl-3-methylglutaryl-CoA lyase -0.0010627053518895476
GO:0044281: small molecule metabolic process HMOX1 heme oxygenase (decycling) 1 -0.00021731471238639437
GO:0044281: small molecule metabolic process HPRT1 hypoxanthine phosphoribosyltransferase 1 -0.0011850174365291715
GO:0044281: small molecule metabolic process HS2ST1 heparan sulfate 2-O-sulfotransferase 1 -0.000809822166168696
GO:0044281: small molecule metabolic process HSD17B4 hydroxysteroid (17-beta) dehydrogenase 4 -0.0011616703408892084
GO:0044281: small molecule metabolic process IDO1 indoleamine 2,3-dioxygenase 1 0.001224705151066539
GO:0044281: small molecule metabolic process IMPDH1 IMP (inosine 5'-monophosphate) dehydrogenase 1 -0.0018705519376337988
GO:0044281: small molecule metabolic process IMPDH2 IMP (inosine 5'-monophosphate) dehydrogenase 2 -0.0003471367382160138
GO:0044281: small molecule metabolic process INPP4B inositol polyphosphate-4-phosphatase, type II, 105kDa 0.0002900299912188722
GO:0044281: small molecule metabolic process ITPR1 inositol 1,4,5-trisphosphate receptor, type 1 -0.0007795693559122963
GO:0044281: small molecule metabolic process IVD isovaleryl-CoA dehydrogenase -0.000935309863094636
GO:0044281: small molecule metabolic process KPNB1 karyopherin (importin) beta 1 0.0007714731747169504
GO:0044281: small molecule metabolic process MCCC1 methylcrotonoyl-CoA carboxylase 1 (alpha) 8.37635285703379e-5
GO:0044281: small molecule metabolic process MCCC2 methylcrotonoyl-CoA carboxylase 2 (beta) -0.0007843290744811766
GO:0044281: small molecule metabolic process MED1 mediator complex subunit 1 0.0011280056167164041
GO:0044281: small molecule metabolic process MGAM maltase-glucoamylase (alpha-glucosidase)

-0.0013076289398003891
GO:0044281: small molecule metabolic process    MTAP    methylthioadenosine phosphorylase       0.002
6559830175411385
GO:0044281: small molecule metabolic process    MTHFD1  methylenetetrahydrofolate dehydrogenase (NADP
+ dependent) 1, methenyltetrahydrofolate cyclohydrolase, formyltetrahydrofolate synthetase      -0.00
06638600248168457
GO:0044281: small molecule metabolic process    MTMR2   myotubularin related protein 2  -0.0004608399
9738915407
GO:0044281: small molecule metabolic process    MUT     methylmalonyl CoA mutase        0.00275690303
28873346
GO:0044281: small molecule metabolic process    NUDT1   nudix (nucleoside diphosphate linked moiety
X)-type motif 1 0.0018356933277766565
GO:0044281: small molecule metabolic process    NUP153  nucleoporin 153kDa      0.0008242990270838395
GO:0044281: small molecule metabolic process    ODC1    ornithine decarboxylase 1       0.00082453386
45328437
GO:0044281: small molecule metabolic process    OGDH    oxoglutarate (alpha-ketoglutarate) dehydrogen
ase (lipoamide) -0.00029449443265991074
GO:0044281: small molecule metabolic process    PDK1    pyruvate dehydrogenase kinase, isozyme 1
0.0014229685868522122
GO:0044281: small molecule metabolic process    PDK2    pyruvate dehydrogenase kinase, isozyme 2
0.0014785758412337595
GO:0044281: small molecule metabolic process    PDK3    pyruvate dehydrogenase kinase, isozyme 3
0.0007714443325769016
GO:0044281: small molecule metabolic process    PEX11A  peroxisomal biogenesis factor 11 alpha  0.000
7169087210258612
GO:0044281: small molecule metabolic process    PGLS    6-phosphogluconolactonase       0.00174889045
9729801
GO:0044281: small molecule metabolic process    PHGDH   phosphoglycerate dehydrogenase  0.00029190524
00168678
GO:0044281: small molecule metabolic process    PIK3CD  phosphatidylinositol-4,5-bisphosphate 3-kinas
e, catalytic subunit delta      -0.0007801866571449217
GO:0044281: small molecule metabolic process    PLA2G5  phospholipase A2, group V       0.00207747489
26489024
GO:0044281: small molecule metabolic process    PLCB1   phospholipase C, beta 1 (phosphoinositide-spe
cific)  0.00015338744229837382
GO:0044281: small molecule metabolic process    PNPO    pyridoxamine 5'-phosphate oxidase       -0.00
1090369235513957
GO:0044281: small molecule metabolic process    POLD1   polymerase (DNA directed), delta 1, catalytic
subunit -0.0004447284406650489
GO:0044281: small molecule metabolic process    PRKCA   protein kinase C, alpha -5.866899738273962e-6
GO:0044281: small molecule metabolic process    PSAT1   phosphoserine aminotransferase 1        -0.00
07224615812606296
GO:0044281: small molecule metabolic process    PSMA5   proteasome (prosome, macropain) subunit, alph
a type, 5       0.0003316121703645838
GO:0044281: small molecule metabolic process    PSMD11  proteasome (prosome, macropain) 26S subunit,
non-ATPase, 11  -0.001094473474880452
GO:0044281: small molecule metabolic process    PSMD13  proteasome (prosome, macropain) 26S subunit,
non-ATPase, 13  -5.98873717603125e-5
GO:0044281: small molecule metabolic process    PSPH    phosphoserine phosphatase       -0.0019339171
391770393
GO:0044281: small molecule metabolic process    PTEN    phosphatase and tensin homolog  1.74955099864
62638e-5
GO:0044281: small molecule metabolic process    QDPR    quinoid dihydropteridine reductase      -0.00
28537157178306163
GO:0044281: small molecule metabolic process    RRM1    ribonucleotide reductase M1     0.00190797770
13236193
GO:0044281: small molecule metabolic process    RRM2    ribonucleotide reductase M2     -0.0005665510
292744961
GO:0044281: small molecule metabolic process    RXRA    retinoid X receptor, alpha      0.00111426772
89570058
GO:0044281: small molecule metabolic process    SEH1L   SEH1-like (S. cerevisiae)       -0.0007321582
541877544
GO:0044281: small molecule metabolic process    SLC25A37        solute carrier family 25 (mitochondri
al iron transporter), member 37 -0.00027879478646128375
GO:0044281: small molecule metabolic process    SLC2A5  solute carrier family 2 (facilitated glucose/
fructose transporter), member 5 0.00023113879380596704
GO:0044281: small molecule metabolic process    SLC35D1 solute carrier family 35 (UDP-GlcA/UDP-GalNAc
transporter), member D1 -6.698629543688241e-5
GO:0044281: small molecule metabolic process    SLC44A4 solute carrier family 44, member 4      0.002
852317752395857
GO:0044281: small molecule metabolic process    SLC6A8  solute carrier family 6 (neurotransmitter tra
nsporter), member 8     -0.0010431893460143566
GO:0044281: small molecule metabolic process    SMARCD3 SWI/SNF related, matrix associated, actin dep
endent regulator of chromatin, subfamily d, member 3    0.00034516995628864613
GO:0044281: small molecule metabolic process    SORD    sorbitol dehydrogenase  -0.002017714520828913
3
GO:0044281: small molecule metabolic process    SPHK1   sphingosine kinase 1    0.0018094047069558865
GO:0044281: small molecule metabolic process    SPR     sepiapterin reductase (7,8-dihydrobiopterin:N
ADP+ oxidoreductase)    -0.0012311909799859445
GO:0044281: small molecule metabolic process    SRD5A2  steroid-5-alpha-reductase, alpha polypeptide
2 (3-oxo-5 alpha-steroid delta 4-dehydrogenase alpha 2) -0.0006423794498907138

GO:0044281: small molecule metabolic process    STAR    steroidogenic acute regulatory protein  0.000
8936042515129592
GO:0044281: small molecule metabolic process    SULT4A1 sulfotransferase family 4A, member 1    -0.00
11680790625527699
GO:0044281: small molecule metabolic process    TAT     tyrosine aminotransferase       0.00105200948
38986239
GO:0044281: small molecule metabolic process    TH      tyrosine hydroxylase    -0.00034198743623431O
5
GO:0044281: small molecule metabolic process    TNFRSF21        tumor necrosis factor receptor superf
amily, member 21        0.00034818714596347814
GO:0044281: small molecule metabolic process    TPR     translocated promoter region, nuclear basket
protein -0.0006965563073315005
GO:0044281: small molecule metabolic process    TYMS    thymidylate synthetase  0.0015596642566728517
GO:0044281: small molecule metabolic process    UGCG    UDP-glucose ceramide glucosyltransferase
0.00036898840251944497
GO:0044281: small molecule metabolic process    UGDH    UDP-glucose 6-dehydrogenase     0.00050179489
44982196
GO:0044281: small molecule metabolic process    UGP2    UDP-glucose pyrophosphorylase 2 0.00078446674
07606123
GO:0045542: positive regulation of cholesterol biosynthetic process     ABCG1   ATP-binding cassette,
sub-family G (WHITE), member 1  -0.0023467586157089023
GO:0045542: positive regulation of cholesterol biosynthetic process     POR     P450 (cytochrome) oxi
doreductase     0.00045894675342146325
GO:0055085: transmembrane transport     ABCG1   ATP-binding cassette, sub-family G (WHITE), member 1
-0.002337396313274773
GO:0055085: transmembrane transport     ASIC2   acid-sensing (proton-gated) ion channel 2       0.002
27841985917225
GO:0055085: transmembrane transport     ATP1B3  ATPase, Na+/K+ transporting, beta 3 polypeptide -0.00
14972326308670327
GO:0055085: transmembrane transport     AVP     arginine vasopressin    -0.0009360706646363331
GO:0055085: transmembrane transport     BCL2    B-cell CLL/lymphoma 2   -4.8237896521702126e-6
GO:0055085: transmembrane transport     CLCN2   chloride channel, voltage-sensitive 2   0.00105915612
87887508
GO:0055085: transmembrane transport     CP      ceruloplasmin (ferroxidase)     0.0013070700736619442
GO:0055085: transmembrane transport     GABRA4  gamma-aminobutyric acid (GABA) A receptor, alpha 4
-0.0014832472302783266
GO:0055085: transmembrane transport     HMOX1   heme oxygenase (decycling) 1    -0.000217209734321947
13
GO:0055085: transmembrane transport     MRS2    MRS2 magnesium transporter      0.0006567646168481231
GO:0055085: transmembrane transport     NUP153  nucleoporin 153kDa      0.0008235870970852995
GO:0055085: transmembrane transport     RAF1    Raf-1 proto-oncogene, serine/threonine kinase   0.001
4977626876629012
GO:0055085: transmembrane transport     SEH1L   SEH1-like (S. cerevisiae)       -0.000732851748420286
5
GO:0055085: transmembrane transport     SLC11A2 solute carrier family 11 (proton-coupled divalent met
al ion transporter), member 2   -0.0006650349956668878
GO:0055085: transmembrane transport     SLC15A1 solute carrier family 15 (oligopeptide transporter),
member 1        -0.002758565564227252
GO:0055085: transmembrane transport     SLC15A2 solute carrier family 15 (oligopeptide transporter),
member 2        -0.00012308663479612314
GO:0055085: transmembrane transport     SLC17A3 solute carrier family 17 (organic anion transporter),
member 3        -0.0008661666739755724
GO:0055085: transmembrane transport     SLC1A1  solute carrier family 1 (neuronal/epithelial high aff
inity glutamate transporter, system Xag), member 1      -0.0009006843346179448
GO:0055085: transmembrane transport     SLC1A3  solute carrier family 1 (glial high affinity glutamat
e transporter), member 3        -0.0031193849803142844
GO:0055085: transmembrane transport     SLC1A5  solute carrier family 1 (neutral amino acid transport
er), member 5   0.0015350964550970097
GO:0055085: transmembrane transport     SLC22A18        solute carrier family 22, member 18     0.000
7342816565890956
GO:0055085: transmembrane transport     SLC22A4 solute carrier family 22 (organic cation/zwitterion t
ransporter), member 4   -0.0004797831934512718
GO:0055085: transmembrane transport     SLC22A5 solute carrier family 22 (organic cation/carnitine tr
ansporter), member 5    -0.001852304629286961
GO:0055085: transmembrane transport     SLC25A44        solute carrier family 25, member 44     0.000
3453198541108317
GO:0055085: transmembrane transport     SLC2A5  solute carrier family 2 (facilitated glucose/fructose
transporter), member 5  0.00023101582432796122
GO:0055085: transmembrane transport     SLC31A1 solute carrier family 31 (copper transporter), member
1       -2.6521411238811124e-5
GO:0055085: transmembrane transport     SLC35D1 solute carrier family 35 (UDP-GlcA/UDP-GalNAc transpo
rter), member D1        -6.695539968320712e-5
GO:0055085: transmembrane transport     SLC39A14        solute carrier family 39 (zinc transporter),
member 14       0.0001967108181535064
GO:0055085: transmembrane transport     SLC39A6 solute carrier family 39 (zinc transporter), member 6
0.0003055208990129552
GO:0055085: transmembrane transport     SLC44A4 solute carrier family 44, member 4      0.00285394415
53379397
GO:0055085: transmembrane transport     SLC4A7  solute carrier family 4, sodium bicarbonate cotranspo
rter, member 7  0.0008821552977918824
GO:0055085: transmembrane transport     SLC5A7  solute carrier family 5 (sodium/choline cotransporte

r), member 7    0.002010652153209131
GO:0055085: transmembrane transport     SLC7A5  solute carrier family 7 (amino acid transporter light chain, L system), member 5      -0.0025734284231520616
GO:0055085: transmembrane transport     SLC7A7  solute carrier family 7 (amino acid transporter light chain, y+L system), member 7    -0.0005985999534865756
GO:0055085: transmembrane transport     SLC7A8  solute carrier family 7 (amino acid transporter light chain, L system), member 8      -0.0004690251307187532
GO:0055085: transmembrane transport     STEAP3  STEAP family member 3, metalloreductase 0.002607597853962525
GO:0055085: transmembrane transport     TFRC    transferrin receptor    0.0010839266451801008
GO:0055085: transmembrane transport     TPR     translocated promoter region, nuclear basket protein    -0.0006968639987634029
GO:0055085: transmembrane transport     TTYH1   tweety family member 1  0.0008688345226562565
GO:0055091: phospholipid homeostasis    ABCG1   ATP-binding cassette, sub-family G (WHITE), member 1    -0.0023606226924747856
GO:0055099: response to high density lipoprotein particle       ABCG1   ATP-binding cassette, sub-family G (WHITE), member 1 -0.0023606226924747856
GO:1901998: toxin transport     ABCG1   ATP-binding cassette, sub-family G (WHITE), member 1    -0.002346795806007113
GO:1901998: toxin transport     BNIP3   BCL2/adenovirus E1B 19kDa interacting protein 3 0.002923295895721561
GO:1901998: toxin transport     DNM1    dynamin 1       -0.0013148182842815043
GO:1901998: toxin transport     LRP6    low density lipoprotein receptor-related protein 6      0.00015035712009481189
GO:1901998: toxin transport     NRP1    neuropilin 1    -0.0006458480958389628
GO:1901998: toxin transport     SLC17A3 solute carrier family 17 (organic anion transporter), member 3       -0.0008704568727983816
GO:1901998: toxin transport     SLC7A8  solute carrier family 7 (amino acid transporter light chain, L system), member 8     -0.0004731488263616598
GO:0000447: endonucleolytic cleavage in ITS1 to separate SSU-rRNA from 5.8S rRNA and LSU-rRNA from tricistronic rRNA transcript (SSU-rRNA, 5.8S rRNA, LSU-rRNA)      ABT1    activator of basal transcription 1   0.0025808828118788615
GO:0000472: endonucleolytic cleavage to generate mature 5'-end of SSU-rRNA from (SSU-rRNA, 5.8S rRNA, LSU-rRNA)       ABT1    activator of basal transcription 1      0.0025808828118788615
GO:0000480: endonucleolytic cleavage in 5'-ETS of tricistronic rRNA transcript (SSU-rRNA, 5.8S rRNA, LSU-rRNA)       ABT1    activator of basal transcription 1      0.0025808828118788615
GO:0006357: regulation of transcription from RNA polymerase II promoter ABT1    activator of basal transcription 1   0.0025485994256678884
GO:0006357: regulation of transcription from RNA polymerase II promoter ATF5    activating transcription factor 5    -0.0027426922287843605
GO:0006357: regulation of transcription from RNA polymerase II promoter BATF    basic leucine zipper transcription factor, ATF-like  0.0020423455617131865
GO:0006357: regulation of transcription from RNA polymerase II promoter BBS7    Bardet-Biedl syndrome 7       -0.0009800454774115735
GO:0006357: regulation of transcription from RNA polymerase II promoter CHD3    chromodomain helicase DNA binding protein 3   0.00011870809770782354
GO:0006357: regulation of transcription from RNA polymerase II promoter CITED1  Cbp/p300-interacting transactivator, with Glu/Asp-rich carboxy-terminal domain, 1    0.0028266656620635072
GO:0006357: regulation of transcription from RNA polymerase II promoter CLOCK   clock circadian regulator    0.000199564512483896
GO:0006357: regulation of transcription from RNA polymerase II promoter DEK     DEK proto-oncogene      0.002496676092612851
GO:0006357: regulation of transcription from RNA polymerase II promoter ECM1    extracellular matrix protein 1       -0.0015634305285417939
GO:0006357: regulation of transcription from RNA polymerase II promoter ELF5    E74-like factor 5 (ets domain transcription factor)  0.0011776085549585963
GO:0006357: regulation of transcription from RNA polymerase II promoter FOXA2   forkhead box A2 -1.631171154918581e-5
GO:0006357: regulation of transcription from RNA polymerase II promoter FOXC2   forkhead box C2 (MFH-1, mesenchyme forkhead 1)       0.0017528806946937776
GO:0006357: regulation of transcription from RNA polymerase II promoter FOXE3   forkhead box E3 0.0010535980170645683
GO:0006357: regulation of transcription from RNA polymerase II promoter FOXO3   forkhead box O3 0.0012039046713783101
GO:0006357: regulation of transcription from RNA polymerase II promoter GRHL2   grainyhead-like 2 (Drosophila)       0.0010091329198541844
GO:0006357: regulation of transcription from RNA polymerase II promoter HMGB1   high mobility group box 1    -0.0007775343098847414
GO:0006357: regulation of transcription from RNA polymerase II promoter HMGB2   high mobility group box 2    0.00031056276397000327
GO:0006357: regulation of transcription from RNA polymerase II promoter INHBA   inhibin, beta A -0.0013592777854372045
GO:0006357: regulation of transcription from RNA polymerase II promoter PITX2   paired-like homeodomain 2    0.0021825337567691187
GO:0006357: regulation of transcription from RNA polymerase II promoter PKN1    protein kinase N1    -0.001908390609875552
GO:0006357: regulation of transcription from RNA polymerase II promoter PURA    purine-rich element binding protein A        -0.00017148223536069093
GO:0006357: regulation of transcription from RNA polymerase II promoter RAD21   RAD21 homolog (S. pombe)     -0.00010205743301588322
GO:0006357: regulation of transcription from RNA polymerase II promoter SMARCD3 SWI/SNF related, matr

ix associated, actin dependent regulator of chromatin, subfamily d, member 3    0.0003465312712153093
GO:0006357: regulation of transcription from RNA polymerase II promoter SOX10   SRY (sex determining region Y)-box 10       0.00019354372544261626
GO:0006357: regulation of transcription from RNA polymerase II promoter STAT5A  signal transducer and activator of transcription 5A   0.0016005449419091411
GO:0006357: regulation of transcription from RNA polymerase II promoter TBX3    T-box 3 0.0012283722603860599
GO:0006357: regulation of transcription from RNA polymerase II promoter TCF15   transcription factor 15 (basic helix-loop-helix)    -0.00266139228011423
GO:0006357: regulation of transcription from RNA polymerase II promoter TCF7L2  transcription factor 7-like 2 (T-cell specific, HMG-box)    0.0005761274762007429
GO:0006357: regulation of transcription from RNA polymerase II promoter TFAP2C  transcription factor AP-2 gamma (activating enhancer binding protein 2 gamma)       0.0010656573993014762
GO:0006357: regulation of transcription from RNA polymerase II promoter TFCP2L1 transcription factor CP2-like 1     0.00045058761855385225
GO:0006357: regulation of transcription from RNA polymerase II promoter TFDP1   transcription factor Dp-1   0.0014508550242735452
GO:0006357: regulation of transcription from RNA polymerase II promoter THRA    thyroid hormone receptor, alpha      0.0007651182613818635
GO:0006357: regulation of transcription from RNA polymerase II promoter VEGFA   vascular endothelial growth factor A        0.0005979345192633118
GO:0006357: regulation of transcription from RNA polymerase II promoter WDR77   WD repeat domain 77 0.0001343223180984341
GO:0006357: regulation of transcription from RNA polymerase II promoter WT1     Wilms tumor 1   -0.0005088512230075559
GO:0006357: regulation of transcription from RNA polymerase II promoter YY1     YY1 transcription factor     0.0013448430086207975
GO:0006366: transcription from RNA polymerase II promoter       ABT1    activator of basal transcription 1   0.0025472722006984585
GO:0006366: transcription from RNA polymerase II promoter       ALX1    ALX homeobox 1  0.0022972543529411636
GO:0006366: transcription from RNA polymerase II promoter       ATF5    activating transcription factor 5    -0.0027416416805262334
GO:0006366: transcription from RNA polymerase II promoter       BACH1   BTB and CNC homology 1, basic leucine zipper transcription factor 1   0.00048689341453654145
GO:0006366: transcription from RNA polymerase II promoter       BATF    basic leucine zipper transcription factor, ATF-like  0.002042081310521299
GO:0006366: transcription from RNA polymerase II promoter       CDC40   cell division cycle 40  0.004175257673659743
GO:0006366: transcription from RNA polymerase II promoter       CEBPB   CCAAT/enhancer binding protein (C/EBP), beta -0.00027425600777316895
GO:0006366: transcription from RNA polymerase II promoter       CHD5    chromodomain helicase DNA binding protein 5  -0.000520211073697748
GO:0006366: transcription from RNA polymerase II promoter       CLOCK   clock circadian regulator 0.00019972569283432376
GO:0006366: transcription from RNA polymerase II promoter       CREB1   cAMP responsive element binding protein 1    0.0006608582004217471
GO:0006366: transcription from RNA polymerase II promoter       DEK     DEK proto-oncogene      0.0024953548997541427
GO:0006366: transcription from RNA polymerase II promoter       DLX5    distal-less homeobox 5  -0.0032907753226698367
GO:0006366: transcription from RNA polymerase II promoter       DMRT1   doublesex and mab-3 related transcription factor 1   0.001508744824397996
GO:0006366: transcription from RNA polymerase II promoter       EGR1    early growth response 1 0.0010957056680258271
GO:0006366: transcription from RNA polymerase II promoter       EGR2    early growth response 2 0.0014438711865747285
GO:0006366: transcription from RNA polymerase II promoter       ELF5    E74-like factor 5 (ets domain transcription factor)   0.0011765086377914667
GO:0006366: transcription from RNA polymerase II promoter       ESR1    estrogen receptor 1     -0.0009513485164322882
GO:0006366: transcription from RNA polymerase II promoter       ETV4    ets variant 4   0.00019594803348935586
GO:0006366: transcription from RNA polymerase II promoter       FOXA1   forkhead box A1 2.645648113322068e-5
GO:0006366: transcription from RNA polymerase II promoter       FOXA2   forkhead box A2 -1.639489731283133e-5
GO:0006366: transcription from RNA polymerase II promoter       FOXC1   forkhead box C1 -2.1213414545364027e-5
GO:0006366: transcription from RNA polymerase II promoter       FOXC2   forkhead box C2 (MFH-1, mesenchyme forkhead 1)       0.0017522536239867731
GO:0006366: transcription from RNA polymerase II promoter       FOXE3   forkhead box E3 0.0010532326424197403
GO:0006366: transcription from RNA polymerase II promoter       FOXH1   forkhead box H1 -0.0013755641386214127
GO:0006366: transcription from RNA polymerase II promoter       FOXM1   forkhead box M1 0.00019856383697957944
GO:0006366: transcription from RNA polymerase II promoter       GATA2   GATA binding protein 2  -0.00045336436088381273
GO:0006366: transcription from RNA polymerase II promoter       GATA3   GATA binding protein 3  -4.008793872914455e-5

GO:0006366: transcription from RNA polymerase II promoter	GATA4	GATA binding protein 4	-0.00
1097749460076959
GO:0006366: transcription from RNA polymerase II promoter	GATA6	GATA binding protein 6	-2.76
54404874889762e-5
GO:0006366: transcription from RNA polymerase II promoter	GLI2	GLI family zinc finger 2
0.0018581536683981124
GO:0006366: transcription from RNA polymerase II promoter	GLI3	GLI family zinc finger 3
-0.0021581246674813683
GO:0006366: transcription from RNA polymerase II promoter	GRHL2	grainyhead-like 2 (Drosophil
a)	0.001008849373771277
GO:0006366: transcription from RNA polymerase II promoter	HEYL	hes-related family bHLH trans
cription factor with YRPW motif-like	0.0006793749302363296
GO:0006366: transcription from RNA polymerase II promoter	HMGA1	high mobility group AT-hook 1
-0.0003188397277203899
GO:0006366: transcription from RNA polymerase II promoter	HMGA2	high mobility group AT-hook 2
0.0015122341438013314
GO:0006366: transcription from RNA polymerase II promoter	HOXA10	homeobox A10	-0.0029890960
40837077
GO:0006366: transcription from RNA polymerase II promoter	HOXA5	homeobox A5	0.00106936443
14674717
GO:0006366: transcription from RNA polymerase II promoter	HOXD13	homeobox D13	-0.0005931588
345513613
GO:0006366: transcription from RNA polymerase II promoter	IRF7	interferon regulatory factor
 7	-0.00130961299992098
GO:0006366: transcription from RNA polymerase II promoter	ISL1	ISL LIM homeobox 1	7.844
580151951685e-5
GO:0006366: transcription from RNA polymerase II promoter	KLF11	Kruppel-like factor 11	0.000
6196512224498407
GO:0006366: transcription from RNA polymerase II promoter	KLF15	Kruppel-like factor 15	0.000
4664762545163968
GO:0006366: transcription from RNA polymerase II promoter	KMT2A	lysine (K)-specific methyltra
nsferase 2A	0.0006623352953525668
GO:0006366: transcription from RNA polymerase II promoter	LEF1	lymphoid enhancer-binding fac
tor 1	-0.00010032359727424007
GO:0006366: transcription from RNA polymerase II promoter	LHX1	LIM homeobox 1	-0.0007608294
814884723
GO:0006366: transcription from RNA polymerase II promoter	LMO4	LIM domain only 4	0.002
117183041789784
GO:0006366: transcription from RNA polymerase II promoter	MEF2C	myocyte enhancer factor 2C
0.000976217940555779
GO:0006366: transcription from RNA polymerase II promoter	MSX1	msh homeobox 1	-0.0027769676
04298558
GO:0006366: transcription from RNA polymerase II promoter	NFIB	nuclear factor I/B	0.002
92867480402316
GO:0006366: transcription from RNA polymerase II promoter	NFX1	nuclear transcription factor,
X-box binding 1 -6.898439822970423e-5
GO:0006366: transcription from RNA polymerase II promoter	PAX2	paired box 2	-0.0016186836
670777447
GO:0006366: transcription from RNA polymerase II promoter	PAX3	paired box 3	-0.0029521020
923864417
GO:0006366: transcription from RNA polymerase II promoter	PAX6	paired box 6	0.00194085395
7981715
GO:0006366: transcription from RNA polymerase II promoter	PAX8	paired box 8	0.00094970712
1402035
GO:0006366: transcription from RNA polymerase II promoter	PDX1	pancreatic and duodenal homeo
box 1	0.0002583528469544681
GO:0006366: transcription from RNA polymerase II promoter	PHOX2B	paired-like homeobox 2b 0.000
3908077470034439
GO:0006366: transcription from RNA polymerase II promoter	PITX2	paired-like homeodomain 2
0.0021816286096449774
GO:0006366: transcription from RNA polymerase II promoter	POLR2D	polymerase (RNA) II (DNA dire
cted) polypeptide D	0.0018597504149469595
GO:0006366: transcription from RNA polymerase II promoter	POU1F1	POU class 1 homeobox 1	0.000
13787380430972386
GO:0006366: transcription from RNA polymerase II promoter	PRDM4	PR domain containing 4	0.000
8418544216438414
GO:0006366: transcription from RNA polymerase II promoter	PTTG1	pituitary tumor-transforming
 1	-0.0003614587254092106
GO:0006366: transcription from RNA polymerase II promoter	RAD21	RAD21 homolog (S. pombe)
-0.00010210734744061763
GO:0006366: transcription from RNA polymerase II promoter	RREB1	ras responsive element bindin
g protein 1	-0.001985997174195349
GO:0006366: transcription from RNA polymerase II promoter	SIX1	SIX homeobox 1	-0.0018746941
995814437
GO:0006366: transcription from RNA polymerase II promoter	SIX3	SIX homeobox 3	0.00204341751
17996422
GO:0006366: transcription from RNA polymerase II promoter	SNRPB	small nuclear ribonucleoprote
in polypeptides B and B1	0.0006676086128169167
GO:0006366: transcription from RNA polymerase II promoter	SOX10	SRY (sex determining region
 Y)-box 10	0.00019295429910308522
GO:0006366: transcription from RNA polymerase II promoter	SOX11	SRY (sex determining region

Y)-box 11 -0.00020860374601907114
GO:0006366: transcription from RNA polymerase II promoter SOX4 SRY (sex determining region Y)-box 4 -3.747035500166587e-5
GO:0006366: transcription from RNA polymerase II promoter SOX9 SRY (sex determining region Y)-box 9 -0.000519702800589481
GO:0006366: transcription from RNA polymerase II promoter TBPL1 TBP-like 1 -0.0007895771694666626
GO:0006366: transcription from RNA polymerase II promoter TFAP2A transcription factor AP-2 alpha (activating enhancer binding protein 2 alpha) 0.00048071653699130744
GO:0006366: transcription from RNA polymerase II promoter TFAP2C transcription factor AP-2 gamma (activating enhancer binding protein 2 gamma) 0.0010647016641216046
GO:0006366: transcription from RNA polymerase II promoter THRA thyroid hormone receptor, alpha 0.0007648602333120475
GO:0006366: transcription from RNA polymerase II promoter TP73 tumor protein p73 0.0010304016491717314
GO:0006366: transcription from RNA polymerase II promoter TRPS1 trichorhinophalangeal syndrome I -0.00017915056168623752
GO:0006366: transcription from RNA polymerase II promoter WT1 Wilms tumor 1 -0.0005085229322826396
GO:0006366: transcription from RNA polymerase II promoter WWP2 WW domain containing E3 ubiquitin protein ligase 2 -0.0009593203466696433
GO:0006366: transcription from RNA polymerase II promoter XBP1 X-box binding protein 1 0.000266957992993091
GO:0006366: transcription from RNA polymerase II promoter YBX1 Y box binding protein 1 -0.0009148360015116451
GO:0006366: transcription from RNA polymerase II promoter ZNF148 zinc finger protein 148 0.0021093563376307187
GO:0021522: spinal cord motor neuron differentiation ABT1 activator of basal transcription 1 0.00254226626767 2149
GO:0021522: spinal cord motor neuron differentiation DICER1 dicer 1, ribonuclease type III -1.6282337556684913e-5
GO:0021522: spinal cord motor neuron differentiation ISL1 ISL LIM homeobox 1 7.842794313222352e-5
GO:0021522: spinal cord motor neuron differentiation LMO4 LIM domain only 4 0.0021133847233919356
GO:0021522: spinal cord motor neuron differentiation PTCH1 patched 1 -6.544972550372584e-5
GO:0021522: spinal cord motor neuron differentiation SHH sonic hedgehog 0.0006006347645539258
GO:0021522: spinal cord motor neuron differentiation SOX4 SRY (sex determining region Y)-box 4 -3.747943937512265e-5
GO:0034462: small-subunit processome assembly ABT1 activator of basal transcription 1 0.0025808828118788615
KEGG:03320: PPAR signaling pathway ACADL acyl-CoA dehydrogenase, long chain 0.0009646794784327428
KEGG:03320: PPAR signaling pathway CPT1A carnitine palmitoyltransferase 1A (liver) 0.0012641780921739988
KEGG:03320: PPAR signaling pathway NR1H3 nuclear receptor subfamily 1, group H, member 3 0.0007937988522384975
KEGG:03320: PPAR signaling pathway RXRA retinoid X receptor, alpha 0.0011022238142090106
KEGG:00071: Fatty acid degradation ACADL acyl-CoA dehydrogenase, long chain 0.0009596892229527937
KEGG:00071: Fatty acid degradation CPT1A carnitine palmitoyltransferase 1A (liver) 0.0012597313822292199
KEGG:00071: Fatty acid degradation GCDH glutaryl-CoA dehydrogenase -3.881428098773422e-6
GO:0001659: temperature homeostasis ACADL acyl-CoA dehydrogenase, long chain 0.0009707249903913636
GO:0001659: temperature homeostasis DRD2 dopamine receptor D2 -0.0002321098541831773
GO:0001659: temperature homeostasis FOXO1 forkhead box O1 0.0017839815989336965
GO:0001659: temperature homeostasis GPX1 glutathione peroxidase 1 0.0003892603849823641
GO:0006635: fatty acid beta-oxidation ACADL acyl-CoA dehydrogenase, long chain 0.0009899911006884669
GO:0006635: fatty acid beta-oxidation CPT1A carnitine palmitoyltransferase 1A (liver) 0.0012798922242036421
GO:0006635: fatty acid beta-oxidation CROT carnitine O-octanoyltransferase -0.0004431177503181291
GO:0006635: fatty acid beta-oxidation HSD17B4 hydroxysteroid (17-beta) dehydrogenase 4 -0.001171959150548374
GO:0006635: fatty acid beta-oxidation LEP leptin 0.0032193828718324272
GO:0006635: fatty acid beta-oxidation MUT methylmalonyl CoA mutase 0.002777952976479921
GO:0019254: carnitine metabolic process, CoA-linked ACADL acyl-CoA dehydrogenase, long chain 0.0009725879762096732
GO:0033539: fatty acid beta-oxidation using acyl-CoA dehydrogenase ACADL acyl-CoA dehydrogenase, long chain 0.0009573338085733319
GO:0033539: fatty acid beta-oxidation using acyl-CoA dehydrogenase GCDH glutaryl-CoA dehydrogenase -3.6161329727610987e-6
GO:0033539: fatty acid beta-oxidation using acyl-CoA dehydrogenase IVD isovaleryl-CoA dehydrogenase -0.0009192251735 6215
GO:0042413: carnitine catabolic process ACADL acyl-CoA dehydrogenase, long chain 0.0009725879762096732
GO:0042758: long-chain fatty acid catabolic process ACADL acyl-CoA dehydrogenase, long chain 0.0009725879762096732
GO:0044242: cellular lipid catabolic process ACADL acyl-CoA dehydrogenase, long chain 0.000

9700934142145932
GO:0044242: cellular lipid catabolic process    SIRT2   sirtuin 2       -0.0008421256606398701
GO:0044255: cellular lipid metabolic process    ACADL   acyl-CoA dehydrogenase, long chain      0.000
9737906064588631
GO:0044255: cellular lipid metabolic process    ACOT8   acyl-CoA thioesterase 8 0.001131112167355671
GO:0044255: cellular lipid metabolic process    AGT     angiotensinogen (serpin peptidase inhibitor,
 clade A, member 8)     -0.0011210587670683874
GO:0044255: cellular lipid metabolic process    BDH1    3-hydroxybutyrate dehydrogenase, type 1 -0.00
02068315791200919
GO:0044255: cellular lipid metabolic process    CPT1A   carnitine palmitoyltransferase 1A (liver)
0.0012664374856221055
GO:0044255: cellular lipid metabolic process    CROT    carnitine O-octanoyltransferase -0.0004339150
4498231236
GO:0044255: cellular lipid metabolic process    ELOVL2  ELOVL fatty acid elongase 2     0.00052016469
31292288
GO:0044255: cellular lipid metabolic process    ELOVL5  ELOVL fatty acid elongase 5     0.00020743620
68424031
GO:0044255: cellular lipid metabolic process    GPD1L   glycerol-3-phosphate dehydrogenase 1-like
0.0005636324959056468
GO:0044255: cellular lipid metabolic process    HMGCL   3-hydroxymethyl-3-methylglutaryl-CoA lyase
-0.0010552294126392034
GO:0044255: cellular lipid metabolic process    HSD17B4 hydroxysteroid (17-beta) dehydrogenase 4
-0.0011538294707715091
GO:0044255: cellular lipid metabolic process    MED1    mediator complex subunit 1      0.00111908854
81381849
GO:0044255: cellular lipid metabolic process    MUT     methylmalonyl CoA mutase        0.00274152268
5246229
GO:0044255: cellular lipid metabolic process    PEX11A  peroxisomal biogenesis factor 11 alpha  0.000
7132308480431776
GO:0044255: cellular lipid metabolic process    RXRA    retinoid X receptor, alpha      0.00110766672
5026051
GO:0044255: cellular lipid metabolic process    SMARCD3 SWI/SNF related, matrix associated, actin dep
endent regulator of chromatin, subfamily d, member 3    0.0003428031927468975
GO:0044255: cellular lipid metabolic process    TNFRSF21        tumor necrosis factor receptor superf
amily, member 21        0.0003450018055870307
GO:0045717: negative regulation of fatty acid biosynthetic process      ACADL   acyl-CoA dehydrogenas
e, long chain   0.0009725879762096732
GO:0046322: negative regulation of fatty acid oxidation ACADL   acyl-CoA dehydrogenase, long chain
0.0009725879762096732
GO:0051289: protein homotetramerization ACADL   acyl-CoA dehydrogenase, long chain      0.00097903809
0400719
GO:0051289: protein homotetramerization ALDH5A1 aldehyde dehydrogenase 5 family, member A1      0.002
3240892942921546
GO:0051289: protein homotetramerization CTH     cystathionine gamma-lyase       0.001744065915903084
GO:0051289: protein homotetramerization EVL     Enah/Vasp-like  0.001936933421172402
GO:0051289: protein homotetramerization GOLGA2  golgin A2       0.00016771156185921486
GO:0051289: protein homotetramerization HPRT1   hypoxanthine phosphoribosyltransferase 1        -0.00
11848873314536897
GO:0051289: protein homotetramerization IMPDH2  IMP (inosine 5'-monophosphate) dehydrogenase 2  -0.00
034555137705332794
GO:0051289: protein homotetramerization KCNJ2   potassium inwardly-rectifying channel, subfamily J, m
ember 2 -0.000775272205327048
GO:0051289: protein homotetramerization RXRA    retinoid X receptor, alpha      0.0011134638179271623
GO:0051289: protein homotetramerization SHMT2   serine hydroxymethyltransferase 2 (mitochondrial)
-0.0010981723116027264
GO:0055088: lipid homeostasis   ACADL   acyl-CoA dehydrogenase, long chain      0.0009540101374639423
GO:0055088: lipid homeostasis   GCDH    glutaryl-CoA dehydrogenase      -3.4228792044652572e-6
GO:0055088: lipid homeostasis   IVD     isovaleryl-CoA dehydrogenase    -0.0009170692873098373
GO:0055088: lipid homeostasis   NR1H3   nuclear receptor subfamily 1, group H, member 3 0.00078771538
67270804
GO:0055114: oxidation-reduction process ACADL   acyl-CoA dehydrogenase, long chain      0.00098322535
73415454
GO:0055114: oxidation-reduction process AKR1C1  aldo-keto reductase family 1, member C1 -0.0014911606
652435315
GO:0055114: oxidation-reduction process ALDH1A2 aldehyde dehydrogenase 1 family, member A2      -0.00
3023800666767997
GO:0055114: oxidation-reduction process AOC1    amine oxidase, copper containing 1      -0.0014253945
956716426
GO:0055114: oxidation-reduction process APEX1   APEX nuclease (multifunctional DNA repair enzyme) 1
0.0002005883040891176
GO:0055114: oxidation-reduction process BDH1    3-hydroxybutyrate dehydrogenase, type 1 -0.0002074045
4285367665
GO:0055114: oxidation-reduction process CBS     cystathionine-beta-synthase     0.0015192448254002203
GO:0055114: oxidation-reduction process CP      ceruloplasmin (ferroxidase)     0.0013095237798756388
GO:0055114: oxidation-reduction process CPOX    coproporphyrinogen oxidase      0.001275685117244933
GO:0055114: oxidation-reduction process CYB5R4  cytochrome b5 reductase 4       -2.8434697809665187e-
5
GO:0055114: oxidation-reduction process CYP1B1  cytochrome P450, family 1, subfamily B, polypeptide 1
0.0003982236045467949
GO:0055114: oxidation-reduction process CYP27B1 cytochrome P450, family 27, subfamily B, polypeptide
 1      -0.00047215586872921685

GO:0055114: oxidation-reduction process CYP4B1  cytochrome P450, family 4, subfamily B, polypeptide 1
-0.0013589948123098427
GO:0055114: oxidation-reduction process CYP7B1  cytochrome P450, family 7, subfamily B, polypeptide 1
0.00016352255250777286
GO:0055114: oxidation-reduction process DCT     dopachrome tautomerase  -0.002356737353633095
GO:0055114: oxidation-reduction process DHCR24  24-dehydrocholesterol reductase -0.001710259766274021
9
GO:0055114: oxidation-reduction process DHFR    dihydrofolate reductase -0.00030786505875509155
GO:0055114: oxidation-reduction process GLRX2   glutaredoxin 2  0.0011177438867716778
GO:0055114: oxidation-reduction process GLUD2   glutamate dehydrogenase 2       -0.000659897177300448
6
GO:0055114: oxidation-reduction process GPD1L   glycerol-3-phosphate dehydrogenase 1-like       0.000
5683640146230379
GO:0055114: oxidation-reduction process HSD17B4 hydroxysteroid (17-beta) dehydrogenase 4        -0.00
11645341890049062
GO:0055114: oxidation-reduction process HSD17B6 hydroxysteroid (17-beta) dehydrogenase 6        -0.00
08687761628787207
GO:0055114: oxidation-reduction process HSDL2   hydroxysteroid dehydrogenase like 2     0.00137043369
03226139
GO:0055114: oxidation-reduction process IDO1    indoleamine 2,3-dioxygenase 1   0.0012279669183532962
GO:0055114: oxidation-reduction process IMPDH1  IMP (inosine 5'-monophosphate) dehydrogenase 1  -0.00
18751854036111269
GO:0055114: oxidation-reduction process IMPDH2  IMP (inosine 5'-monophosphate) dehydrogenase 2  -0.00
034838345354934333
GO:0055114: oxidation-reduction process JMJD6   jumonji domain containing 6     0.003596320288057919
GO:0055114: oxidation-reduction process LOX     lysyl oxidase   -0.0005774280777408365
GO:0055114: oxidation-reduction process MTHFD1  methylenetetrahydrofolate dehydrogenase (NADP+ depend
ent) 1, methenyltetrahydrofolate cyclohydrolase, formyltetrahydrofolate synthetase      -0.0006649171
409238842
GO:0055114: oxidation-reduction process PAX2    paired box 2    -0.0016168312553755798
GO:0055114: oxidation-reduction process PHGDH   phosphoglycerate dehydrogenase  0.0002934951681328271
GO:0055114: oxidation-reduction process PNPO    pyridoxamine 5'-phosphate oxidase       -0.0010931856
105589374
GO:0055114: oxidation-reduction process POR     P450 (cytochrome) oxidoreductase        0.00045800762
38634595
GO:0055114: oxidation-reduction process PRDX4   peroxiredoxin 4 0.0015269658420196228
GO:0055114: oxidation-reduction process QDPR    quinoid dihydropteridine reductase      -0.0028603630
968169467
GO:0055114: oxidation-reduction process RRM1    ribonucleotide reductase M1     0.001913561896799083
GO:0055114: oxidation-reduction process RRM2    ribonucleotide reductase M2     -0.000566635548573676
3
GO:0055114: oxidation-reduction process SORD    sorbitol dehydrogenase  -0.0020229309345302115
GO:0055114: oxidation-reduction process SPR     sepiapterin reductase (7,8-dihydrobiopterin:NADP+ oxi
doreductase)    -0.0012347649997144635
GO:0055114: oxidation-reduction process SRD5A2  steroid-5-alpha-reductase, alpha polypeptide 2 (3-oxo
-5 alpha-steroid delta 4-dehydrogenase alpha 2) -0.0006443451086806787
GO:0055114: oxidation-reduction process STEAP3  STEAP family member 3, metalloreductase 0.00261307963
5988976
GO:0055114: oxidation-reduction process STEAP4  STEAP family member 4   -0.0027720278485580843
GO:0055114: oxidation-reduction process TH      tyrosine hydroxylase    -0.00034249651780864814
GO:0055114: oxidation-reduction process TXN2    thioredoxin 2   0.0013852052524674077
GO:0055114: oxidation-reduction process UGDH    UDP-glucose 6-dehydrogenase     0.0005023683750838108
GO:0090181: regulation of cholesterol metabolic process ACADL   acyl-CoA dehydrogenase, long chain
0.0009725879762096732
KEGG:00120: Primary bile acid biosynthesis      ACOT8   acyl-CoA thioesterase 8 0.0011250415158007095
KEGG:00120: Primary bile acid biosynthesis      CYP7B1  cytochrome P450, family 7, subfamily B, polyp
eptide 1        0.00016078807902458235
KEGG:00120: Primary bile acid biosynthesis      HSD17B4 hydroxysteroid (17-beta) dehydrogenase 4
-0.0011418915082748657
KEGG:04146: Peroxisome  ACOT8   acyl-CoA thioesterase 8 0.0011267202985504377
KEGG:04146: Peroxisome  CROT    carnitine O-octanoyltransferase -0.0004311288076447868
KEGG:04146: Peroxisome  HMGCL   3-hydroxymethyl-3-methylglutaryl-CoA lyase      -0.001049136670314674
4
KEGG:04146: Peroxisome  HSD17B4 hydroxysteroid (17-beta) dehydrogenase 4        -0.001147517066830305
9
KEGG:04146: Peroxisome  PEX11A  peroxisomal biogenesis factor 11 alpha  0.0007093964743551378
GO:0006637: acyl-CoA metabolic process  ACOT8   acyl-CoA thioesterase 8 0.0011296365421344246
GO:0006637: acyl-CoA metabolic process  HMGCL   3-hydroxymethyl-3-methylglutaryl-CoA lyase      -0.00
10515053781074923
GO:0006699: bile acid biosynthetic process      ACOT8   acyl-CoA thioesterase 8 0.0011264948182661992
GO:0006699: bile acid biosynthetic process      CYP7B1  cytochrome P450, family 7, subfamily B, polyp
eptide 1        0.00016106635738449642
GO:0006699: bile acid biosynthetic process      HSD17B4 hydroxysteroid (17-beta) dehydrogenase 4
-0.001145062651649703
GO:0006699: bile acid biosynthetic process      STAR    steroidogenic acute regulatory protein  0.000
8797005036514014
GO:0008206: bile acid metabolic process ACOT8   acyl-CoA thioesterase 8 0.0011400378260621648
GO:0008206: bile acid metabolic process AKR1C1  aldo-keto reductase family 1, member C1 -0.0014955608
77114569
GO:0008206: bile acid metabolic process CYP7B1  cytochrome P450, family 7, subfamily B, polypeptide 1
0.00016378248230008337

GO:0008206: bile acid metabolic process HSD17B4 hydroxysteroid (17-beta) dehydrogenase 4 -0.00
11683509725662015
GO:0008206: bile acid metabolic process LEP leptin 0.003209837568461649
GO:0008206: bile acid metabolic process RXRA retinoid X receptor, alpha 0.001120879310317521
GO:0016032: viral process ACOT8 acyl-CoA thioesterase 8 0.0011332087985093959
GO:0016032: viral process BAX BCL2-associated X protein -0.0004239889194544545
GO:0016032: viral process BICD1 bicaudal D homolog 1 (Drosophila) 0.0024310978956263865
GO:0016032: viral process BNIP3 BCL2/adenovirus E1B 19kDa interacting protein 3 0.00289938193
3048572
GO:0016032: viral process BNIP3L BCL2/adenovirus E1B 19kDa interacting protein 3-like -5.76
37289103369734e-5
GO:0016032: viral process BRD4 bromodomain containing 4 0.0005528164370892126
GO:0016032: viral process BUB1 BUB1 mitotic checkpoint serine/threonine kinase 0.00131793572
73544674
GO:0016032: viral process CALCOCO2 calcium binding and coiled-coil domain 2 0.002
4467567733152218
GO:0016032: viral process CCDC86 coiled-coil domain containing 86 0.0007146750595481012
GO:0016032: viral process CREB1 cAMP responsive element binding protein 1 0.00065389482
45720486
GO:0016032: viral process CUL7 cullin 7 -0.00011804496107927607
GO:0016032: viral process CX3CR1 chemokine (C-X3-C motif) receptor 1 -0.000853631577973925
9
GO:0016032: viral process CXCR4 chemokine (C-X-C motif) receptor 4 0.0007903798646155335
GO:0016032: viral process DAXX death-domain associated protein 0.0008866141034356592
GO:0016032: viral process E4F1 E4F transcription factor 1 0.001477511507960714
GO:0016032: viral process EIF4G1 eukaryotic translation initiation factor 4 gamma, 1 0.000
7391653691435422
GO:0016032: viral process GFI1 growth factor independent 1 transcription repressor 0.001
658638564957058
GO:0016032: viral process GRB2 growth factor receptor-bound protein 2 0.0004279285947750677
GO:0016032: viral process HMGA1 high mobility group AT-hook 1 -0.0003182512018519802
GO:0016032: viral process IL6ST interleukin 6 signal transducer 0.00186879017975036
GO:0016032: viral process IPO5 importin 5 0.0016757694134188303
GO:0016032: viral process KPNB1 karyopherin (importin) beta 1 0.0007693123028523805
GO:0016032: viral process KRT18 keratin 18 -0.001012147259849316
GO:0016032: viral process LYN LYN proto-oncogene, Src family tyrosine kinase -0.0014147386
365213094
GO:0016032: viral process MAP3K5 mitogen-activated protein kinase kinase kinase 5 0.000
3407132397526862
GO:0016032: viral process MFGE8 milk fat globule-EGF factor 8 protein 0.0017956025031863698
GO:0016032: viral process MSH6 mutS homolog 6 0.0010715194755071408
GO:0016032: viral process NFX1 nuclear transcription factor, X-box binding 1 -7.1670514024
0045e-5
GO:0016032: viral process NUP153 nucleoporin 153kDa 0.0008197281779380463
GO:0016032: viral process PML promyelocytic leukemia -0.0006785609772968494
GO:0016032: viral process POLA1 polymerase (DNA directed), alpha 1, catalytic subunit 0.001
2637540074285634
GO:0016032: viral process POLR2D polymerase (RNA) II (DNA directed) polypeptide D 0.001
8466217380317508
GO:0016032: viral process PSMA5 proteasome (prosome, macropain) subunit, alpha type, 5 0.000
33006252451522997
GO:0016032: viral process PSMD11 proteasome (prosome, macropain) 26S subunit, non-ATPase, 11
-0.0010919739958158616
GO:0016032: viral process PSMD13 proteasome (prosome, macropain) 26S subunit, non-ATPase, 13
-6.0484852038221646e-5
GO:0016032: viral process RB1 retinoblastoma 1 -0.0014873904032028232
GO:0016032: viral process RHOA ras homolog family member A 0.0006670271923979144
GO:0016032: viral process SEH1L SEH1-like (S. cerevisiae) -0.0007307140972634647
GO:0016032: viral process SF3B2 splicing factor 3b, subunit 2, 145kDa 0.0006803728928588609
GO:0016032: viral process SIRT1 sirtuin 1 -2.557371602793684e-6
GO:0016032: viral process TFRC transferrin receptor 0.001080488028984342
GO:0016032: viral process TP53 tumor protein p53 0.00116871038602404
GO:0016032: viral process TP73 tumor protein p73 0.0010235047059473923
GO:0016032: viral process TPR translocated promoter region, nuclear basket protein -0.00
0694545133445699
GO:0016032: viral process UNG uracil-DNA glycosylase 0.0005621207451607175
GO:0016032: viral process VPS4A vacuolar protein sorting 4 homolog A (S. cerevisiae) -0.00
06818751482759127
GO:0016559: peroxisome fission ACOT8 acyl-CoA thioesterase 8 0.001124290631692677
GO:0016559: peroxisome fission PEX11A peroxisomal biogenesis factor 11 alpha 0.0007019294818520601
GO:0033540: fatty acid beta-oxidation using acyl-CoA oxidase ACOT8 acyl-CoA thioesterase 8 0.001
1263428857322378
GO:0033540: fatty acid beta-oxidation using acyl-CoA oxidase CROT carnitine O-octanoyltransfera
se -0.0004311244172930995
GO:0033540: fatty acid beta-oxidation using acyl-CoA oxidase HSD17B4 hydroxysteroid (17-beta) dehy
drogenase 4 -0.0011472315828132782
GO:0033559: unsaturated fatty acid metabolic process ACOT8 acyl-CoA thioesterase 8 0.00112413389
60080746
GO:0033559: unsaturated fatty acid metabolic process ELOVL2 ELOVL fatty acid elongase 2 0.000
5126286214051827
GO:0033559: unsaturated fatty acid metabolic process ELOVL5 ELOVL fatty acid elongase 5 0.000

2080830990032366
GO:0033559: unsaturated fatty acid metabolic process HSD17B4 hydroxysteroid (17-beta) dehydrogenas
e 4 -0.00114184805293816l
GO:0036109: alpha-linolenic acid metabolic process ACOT8 acyl-CoA thioesterase 8 0.00112413389
60080746
GO:0036109: alpha-linolenic acid metabolic process ELOVL2 ELOVL fatty acid elongase 2 0.000
5126286214051827
GO:0036109: alpha-linolenic acid metabolic process ELOVL5 ELOVL fatty acid elongase 5 0.000
2080830990032366
GO:0036109: alpha-linolenic acid metabolic process HSD17B4 hydroxysteroid (17-beta) dehydrogenas
e 4 -0.00114184805293816l
GO:0043649: dicarboxylic acid catabolic process ACOT8 acyl-CoA thioesterase 8 0.0011250785466930266
GO:0007275: multicellular organismal development ACRV1 acrosomal vesicle protein 1 -0.00
0602632480003769
GO:0007275: multicellular organismal development AES amino-terminal enhancer of split
0.0013608250886879669
GO:0007275: multicellular organismal development ALX1 ALX homeobox 1 0.002304854683493938
GO:0007275: multicellular organismal development CENPE centromere protein E, 312kDa 0.002
1641843690307526
GO:0007275: multicellular organismal development DSPP dentin sialophosphoprotein 0.000
8332002663454993
GO:0007275: multicellular organismal development EGFL6 EGF-like-domain, multiple 6 0.000
13791280033759865
GO:0007275: multicellular organismal development EYA3 EYA transcriptional coactivator and p
hosphatase 3 0.0008414052963303427
GO:0007275: multicellular organismal development GCNT2 glucosaminyl (N-acetyl) transferase
2, I-branching enzyme (I blood group) 0.0024467691116296084
GO:0007275: multicellular organismal development GNRHR gonadotropin-releasing hormone recept
or 0.0007448181806739657
GO:0007275: multicellular organismal development HELLS helicase, lymphoid-specific 0.003
4841415762245946
GO:0007275: multicellular organismal development HMGA2 high mobility group AT-hook 2 0.001
5176454559921598
GO:0007275: multicellular organismal development HMGB3 high mobility group box 3 0.000
46591417626646643
GO:0007275: multicellular organismal development HOXA10 homeobox A10 -0.002999580579960397
4
GO:0007275: multicellular organismal development HOXB1 homeobox B1 0.0037064002506680916
GO:0007275: multicellular organismal development HOXB2 homeobox B2 0.0027884294923930476
GO:0007275: multicellular organismal development HOXD13 homeobox D13 -0.000594838497653423
GO:0007275: multicellular organismal development ID2 inhibitor of DNA binding 2, dominant
negative helix-loop-helix protein 5.999482185643325e-5
GO:0007275: multicellular organismal development PAEP progestagen-associated endometrial pr
otein 0.0010706068741512256
GO:0007275: multicellular organismal development RREB1 ras responsive element binding protei
n 1 -0.0019922654238996515
GO:0007275: multicellular organismal development SPDEF SAM pointed domain containing ETS tra
nscription factor 0.0031544458464539717
GO:0007275: multicellular organismal development SUCO SUN domain containing ossification fa
ctor 0.002261786124292338
GO:0007275: multicellular organismal development TBX21 T-box 21 0.0004003452354227904
GO:0007275: multicellular organismal development TP53 tumor protein p53 0.00118414751
65336951
GO:0001525: angiogenesis ACVRL1 activin A receptor type II-like 1 0.0019380999107114756
GO:0001525: angiogenesis APOD apolipoprotein D 0.0026198779648346974
GO:0001525: angiogenesis ARHGAP24 Rho GTPase activating protein 24 -0.0009316971
671739714
GO:0001525: angiogenesis CAV1 caveolin 1, caveolae protein, 22kDa -0.000534296501562445
3
GO:0001525: angiogenesis CCL2 chemokine (C-C motif) ligand 2 0.0008139262642777207
GO:0001525: angiogenesis CCL8 chemokine (C-C motif) ligand 8 -0.0005893403229684579
GO:0001525: angiogenesis CIB1 calcium and integrin binding 1 (calmyrin) 5.56054062538
9192e-5
GO:0001525: angiogenesis CYP1B1 cytochrome P450, family 1, subfamily B, polypeptide 1 0.000
39674262040591463
GO:0001525: angiogenesis DICER1 dicer 1, ribonuclease type III -1.6371415982969357e-5
GO:0001525: angiogenesis ECM1 extracellular matrix protein 1 -0.0015511685488916948
GO:0001525: angiogenesis EFNA1 ephrin-A1 -0.0005195861981931491
GO:0001525: angiogenesis EPHB1 EPH receptor B1 0.0019584066018005605
GO:0001525: angiogenesis EPHB2 EPH receptor B2 -0.0007209679109037152
GO:0001525: angiogenesis EPHB3 EPH receptor B3 0.0008808647439006741
GO:0001525: angiogenesis FAP fibroblast activation protein, alpha -0.000484703068957529
96
GO:0001525: angiogenesis FGFR2 fibroblast growth factor receptor 2 0.0007605775669832327
GO:0001525: angiogenesis HIF1A hypoxia inducible factor 1, alpha subunit (basic helix-loop-h
elix transcription factor) -0.0006448825921510365
GO:0001525: angiogenesis HMOX1 heme oxygenase (decycling) 1 -0.00021674681670637937
GO:0001525: angiogenesis HOXA3 homeobox A3 0.0009858971129960937
GO:0001525: angiogenesis HOXB13 homeobox B13 0.0018204492376021164
GO:0001525: angiogenesis JAG1 jagged 1 0.0017523636559871694
GO:0001525: angiogenesis MED1 mediator complex subunit 1 0.0011248810657117853

GO:0001525: angiogenesis MFGE8 milk fat globule-EGF factor 8 protein 0.0017984068326029315
GO:0001525: angiogenesis MMP2 matrix metallopeptidase 2 (gelatinase A, 72kDa gelatinase, 72 kDa type IV collagenase) -0.0011267150998895675
GO:0001525: angiogenesis NRP1 neuropilin 1 -0.0006407896487169694
GO:0001525: angiogenesis PRKCA protein kinase C, alpha -5.918547948125366e-6
GO:0001525: angiogenesis PRKX protein kinase, X-linked 0.00039253274308216586
GO:0001525: angiogenesis PTEN phosphatase and tensin homolog 1.7995857133031807e-5
GO:0001525: angiogenesis RBPJ recombination signal binding protein for immunoglobulin kappa J region 0.0009790331924208256
GO:0001525: angiogenesis S100A7 S100 calcium binding protein A7 0.0015846662048894951
GO:0001525: angiogenesis SERPINE1 serpin peptidase inhibitor, clade E (nexin, plasminogen activator inhibitor type 1), member 1 0.0001126352820872885
GO:0001525: angiogenesis SIRT1 sirtuin 1 -2.1779228749138805e-6
GO:0001525: angiogenesis SOX18 SRY (sex determining region Y)-box 18 0.001285129056028295
GO:0001525: angiogenesis TGFB2 transforming growth factor, beta 2 -0.0010555947352951399
GO:0001525: angiogenesis TGFBR1 transforming growth factor, beta receptor 1 0.0003384609672932746
GO:0001525: angiogenesis VASH1 vasohibin 1 0.0005968959576848901
GO:0001525: angiogenesis VEGFA vascular endothelial growth factor A 0.0005921704392206564
GO:0001525: angiogenesis VEGFC vascular endothelial growth factor C -0.003353691530253524
GO:0001525: angiogenesis WNT7A wingless-type MMTV integration site family, member 7A 1.965404811143531e-5
GO:0001525: angiogenesis XBP1 X-box binding protein 1 0.0002670899867725779
GO:0001701: in utero embryonic development ACVRL1 activin A receptor type II-like 1 0.001942799063721505
GO:0001701: in utero embryonic development ADAR adenosine deaminase, RNA-specific -0.0001365954719026238
GO:0001701: in utero embryonic development ANGPT1 angiopoietin 1 0.0009002207230357271
GO:0001701: in utero embryonic development AXIN1 axin 1 -0.0007326864437604173
GO:0001701: in utero embryonic development BCL2L11 BCL2-like 11 (apoptosis facilitator) -0.0007743436337115881
GO:0001701: in utero embryonic development CCNB1 cyclin B1 -0.000881726117418746
GO:0001701: in utero embryonic development CTNNB1 catenin (cadherin-associated protein), beta 1, 88kDa -0.0001168451481805875
GO:0001701: in utero embryonic development EPN1 epsin 1 -0.0005464726383597676
GO:0001701: in utero embryonic development FGFR2 fibroblast growth factor receptor 2 0.0007627787719822483
GO:0001701: in utero embryonic development FOXA2 forkhead box A2 -1.671896804208024e-5
GO:0001701: in utero embryonic development FOXC1 forkhead box C1 -2.2151633814181584e-5
GO:0001701: in utero embryonic development GATA3 GATA binding protein 3 -3.9114628763146046e-5
GO:0001701: in utero embryonic development GATA4 GATA binding protein 4 -0.0010941244136378355
GO:0001701: in utero embryonic development GATA6 GATA binding protein 6 -2.812049431485987e-5
GO:0001701: in utero embryonic development GDF3 growth differentiation factor 3 -0.0018829331093555765
GO:0001701: in utero embryonic development GJA1 gap junction protein, alpha 1, 43kDa -0.0001614499533835121
GO:0001701: in utero embryonic development GLI2 GLI family zinc finger 2 0.00185233581312177
GO:0001701: in utero embryonic development GLI3 GLI family zinc finger 3 -0.002150375404943683
GO:0001701: in utero embryonic development GRHL2 grainyhead-like 2 (Drosophila) 0.001006214636632591
GO:0001701: in utero embryonic development HINFP histone H4 transcription factor 0.0009920066048793145
GO:0001701: in utero embryonic development IHH indian hedgehog -0.0020393182980781824
GO:0001701: in utero embryonic development ITGB1 integrin, beta 1 (fibronectin receptor, beta polypeptide, antigen CD29 includes MDF2, MSK12) 0.0024249919024742876
GO:0001701: in utero embryonic development JAG2 jagged 2 -5.707070766657243e-6
GO:0001701: in utero embryonic development KLF2 Kruppel-like factor 2 -0.0012315113992547775
GO:0001701: in utero embryonic development KMT2D lysine (K)-specific methyltransferase 2D -0.00019928881893223975
GO:0001701: in utero embryonic development MSH2 mutS homolog 2 0.001391454110996456
GO:0001701: in utero embryonic development MSX1 msh homeobox 1 -0.002767249173470441
GO:0001701: in utero embryonic development MYH10 myosin, heavy chain 10, non-muscle -0.0003286039508551057
GO:0001701: in utero embryonic development NOTCH1 notch 1 0.0005185223076809104
GO:0001701: in utero embryonic development PCNT pericentrin 0.0017862910354671168
GO:0001701: in utero embryonic development PDGFRB platelet-derived growth factor receptor, beta polypeptide -0.00036093766342838756
GO:0001701: in utero embryonic development PITX2 paired-like homeodomain 2 0.0021747140578855214
GO:0001701: in utero embryonic development POLE polymerase (DNA directed), epsilon, catalytic subunit 0.002946621927784630З
GO:0001701: in utero embryonic development PTCH1 patched 1 -6.590551620196477e-5
GO:0001701: in utero embryonic development RNASEH2B ribonuclease H2, subunit B -4.3543304634598356e-5
GO:0001701: in utero embryonic development RPA1 replication protein A1, 70kDa 0.00043890123

8480494
GO:0001701: in utero embryonic development RXRA retinoid X receptor, alpha 0.00111439159
42975636
GO:0001701: in utero embryonic development SLIT2 slit homolog 2 (Drosophila) -0.0016191656
339764272
GO:0001701: in utero embryonic development SMO smoothened, frizzled class receptor 0.002
1379492638268164
GO:0001701: in utero embryonic development SOX10 SRY (sex determining region Y)-box 10 0.000
19024190012834546
GO:0001701: in utero embryonic development SOX18 SRY (sex determining region Y)-box 18 0.001
2877585846430403
GO:0001701: in utero embryonic development TBX3 T-box 3 0.0012255975362851358
GO:0001701: in utero embryonic development TGFB3 transforming growth factor, beta 3 -0.00
18226037846556836
GO:0001701: in utero embryonic development TGFBR1 transforming growth factor, beta receptor 1
0.0003396651374221338
GO:0001701: in utero embryonic development TP53 tumor protein p53 0.001174888864625216
GO:0001701: in utero embryonic development TWIST1 twist family bHLH transcription factor 1
-0.0013406405390829822
GO:0001701: in utero embryonic development VEGFA vascular endothelial growth factor A 0.000
5945729999305591
GO:0001701: in utero embryonic development WDR19 WD repeat domain 19 -0.000721730544462502
2
GO:0001701: in utero embryonic development YBX1 Y box binding protein 1 -0.000913341286919911
8
GO:0001701: in utero embryonic development ZBTB18 zinc finger and BTB domain containing 18
0.001271302402325237
GO:0001701: in utero embryonic development ZNF335 zinc finger protein 335 -0.000364760986451832
16
GO:0001936: regulation of endothelial cell proliferation ACVRL1 activin A receptor type II-li
ke 1 0.0019639660590229095
GO:0001936: regulation of endothelial cell proliferation ALDH1A2 aldehyde dehydrogenase 1 fami
ly, member A2 -0.0030504783420228164
GO:0001937: negative regulation of endothelial cell proliferation ACVRL1 activin A receptor ty
pe II-like 1 0.0019250874938716658
GO:0001937: negative regulation of endothelial cell proliferation AGER advanced glycosylatio
n end product-specific receptor -0.00017320352237057754
GO:0001937: negative regulation of endothelial cell proliferation CAV1 caveolin 1, caveolae
protein, 22kDa -0.0005315075031537392
GO:0001937: negative regulation of endothelial cell proliferation ENG endoglin 0.000
816090469351905
GO:0001937: negative regulation of endothelial cell proliferation GJA1 gap junction protein,
alpha 1, 43kDa -0.00015587520759072218
GO:0001937: negative regulation of endothelial cell proliferation SULF1 sulfatase 1 -0.00
07928164889860239
GO:0001937: negative regulation of endothelial cell proliferation TGFBR1 transforming growth f
actor, beta receptor 1 0.0003354764853332718
GO:0001937: negative regulation of endothelial cell proliferation THBS1 thrombospondin 1
-0.001025212600209794
GO:0001937: negative regulation of endothelial cell proliferation VASH1 vasohibin 1 0.000
5934720064626207
GO:0001938: positive regulation of endothelial cell proliferation ACVRL1 activin A receptor ty
pe II-like 1 0.001923187106381941
GO:0001938: positive regulation of endothelial cell proliferation AKT1 v-akt murine thymoma
viral oncogene homolog 1 0.0007213552958856502
GO:0001938: positive regulation of endothelial cell proliferation BMP4 bone morphogenetic pr
otein 4 -0.00031903632616140775
GO:0001938: positive regulation of endothelial cell proliferation CAV1 caveolin 1, caveolae
protein, 22kDa -0.0005312057885718259
GO:0001938: positive regulation of endothelial cell proliferation CCL2 chemokine (C-C motif)
ligand 2 0.000807347083386153
GO:0001938: positive regulation of endothelial cell proliferation CXCL12 chemokine (C-X-C moti
f) ligand 12 -0.0011941913984007723
GO:0001938: positive regulation of endothelial cell proliferation ECM1 extracellular matrix
protein 1 -0.0015363472780654827
GO:0001938: positive regulation of endothelial cell proliferation FGFR3 fibroblast growth fac
tor receptor 3 0.00021856251848388451
GO:0001938: positive regulation of endothelial cell proliferation HIF1A hypoxia inducible fac
tor 1, alpha subunit (basic helix-loop-helix transcription factor) -0.000640315571894329
GO:0001938: positive regulation of endothelial cell proliferation HMGB2 high mobility group b
ox 2 0.0002979610535235676
GO:0001938: positive regulation of endothelial cell proliferation NRP1 neuropilin 1 -0.00
06346855167324196
GO:0001938: positive regulation of endothelial cell proliferation PRKCA protein kinase C, alp
ha -5.930897761783508e-6
GO:0001938: positive regulation of endothelial cell proliferation PROX1 prospero homeobox 1
0.001107536286675277
GO:0001938: positive regulation of endothelial cell proliferation TGFBR1 transforming growth f
actor, beta receptor 1 0.0003349533583460526
GO:0001938: positive regulation of endothelial cell proliferation THBS4 thrombospondin 4
-0.000441521132324233

GO:0001938: positive regulation of endothelial cell proliferation	VEGFA	vascular endothelial growth factor A	0.0005856396246766731
GO:0001938: positive regulation of endothelial cell proliferation	WNT5A	wingless-type MMTV integration site family, member 5A	-0.0006593814258707266
GO:0001946: lymphangiogenesis	ACVRL1	activin A receptor type II-like 1	0.00194612909198142
GO:0001946: lymphangiogenesis	FOXC2	forkhead box C2 (MFH-1, mesenchyme forkhead 1)	0.0017512462823182423
GO:0001946: lymphangiogenesis	PROX1	prospero homeobox 1	0.0011227685110076972
GO:0001946: lymphangiogenesis	SOX18	SRY (sex determining region Y)-box 18	0.0012909123757203743
GO:0001955: blood vessel maturation	ACVRL1	activin A receptor type II-like 1	0.001952714881760225
GO:0001955: blood vessel maturation	MMP2	matrix metallopeptidase 2 (gelatinase A, 72kDa gelatinase, 72kDa type IV collagenase)	-0.0011387836895902068
GO:0001974: blood vessel remodeling	ACVRL1	activin A receptor type II-like 1	0.0019370946521689786
GO:0001974: blood vessel remodeling	AGT	angiotensinogen (serpin peptidase inhibitor, clade A, member 8)	-0.0011264545369124863
GO:0001974: blood vessel remodeling	BAK1	BCL2-antagonist/killer 1	-0.0018697820976758988
GO:0001974: blood vessel remodeling	BAX	BCL2-associated X protein	-0.00042436271413715535
GO:0001974: blood vessel remodeling	FGF8	fibroblast growth factor 8 (androgen-induced)	0.0009795387522410447
GO:0001974: blood vessel remodeling	FOXC1	forkhead box C1 -2.3998618519730685e-5
GO:0001974: blood vessel remodeling	FOXC2	forkhead box C2 (MFH-1, mesenchyme forkhead 1)	0.0017423736182592222
GO:0001974: blood vessel remodeling	HOXA3	homeobox A3	0.0009854992287546631
GO:0001974: blood vessel remodeling	IGF1	insulin-like growth factor 1 (somatomedin C)	0.0001297890536110023
GO:0001974: blood vessel remodeling	JAG1	jagged 1	0.001752178132189106
GO:0001974: blood vessel remodeling	MEF2C	myocyte enhancer factor 2C	0.0009691194573440719
GO:0001974: blood vessel remodeling	RBPJ	recombination signal binding protein for immunoglobulin kappa J region	0.0009788711086604442
GO:0001974: blood vessel remodeling	SEMA3C	sema domain, immunoglobulin domain (Ig), short basic domain, secreted, (semaphorin) 3C	-0.00021923608706525435
GO:0001974: blood vessel remodeling	TGFB2	transforming growth factor, beta 2	-0.0010548733572331348
GO:0002043: blood vessel endothelial cell proliferation involved in sprouting angiogenesis	ACVRL1	activin A receptor type II-like 1	0.0019410890620574757
GO:0002043: blood vessel endothelial cell proliferation involved in sprouting angiogenesis	BMP4	bone morphogenetic protein 4	-0.0003218816800786749
GO:0006275: regulation of DNA replication	ACVRL1	activin A receptor type II-like 1	0.0019557258365855866
GO:0006468: protein phosphorylation	ACVRL1	activin A receptor type II-like 1	0.0019485157138485385
GO:0006468: protein phosphorylation	AKT1	v-akt murine thymoma viral oncogene homolog 1	0.0007327183796848716
GO:0006468: protein phosphorylation	AKT2	v-akt murine thymoma viral oncogene homolog 2	-0.000627221375344193
GO:0006468: protein phosphorylation	AURKA	aurora kinase A 0.0010107899453714244
GO:0006468: protein phosphorylation	AURKB	aurora kinase B 0.0003518632567351575
GO:0006468: protein phosphorylation	AURKC	aurora kinase C 0.0008881894274280078
GO:0006468: protein phosphorylation	BIRC5	baculoviral IAP repeat containing 5	-0.00026068578214866975
GO:0006468: protein phosphorylation	BRD4	bromodomain containing 4	0.0005545687792096687
GO:0006468: protein phosphorylation	BUB1	BUB1 mitotic checkpoint serine/threonine kinase 0.0013285965783585126
GO:0006468: protein phosphorylation	BUB1B	BUB1 mitotic checkpoint serine/threonine kinase B	-0.0012838557908560738
GO:0006468: protein phosphorylation	CAMK1D	calcium/calmodulin-dependent protein kinase ID	-0.002594530103074179
GO:0006468: protein phosphorylation	CCL2	chemokine (C-C motif) ligand 2	0.000818640106632378
GO:0006468: protein phosphorylation	CCL8	chemokine (C-C motif) ligand 8	-0.000591441696383413
GO:0006468: protein phosphorylation	CCND1	cyclin D1	-0.0026407986233100127
GO:0006468: protein phosphorylation	CCNE1	cyclin E1	0.00037201059294231096
GO:0006468: protein phosphorylation	CDK8	cyclin-dependent kinase 8	0.002618452964160652
GO:0006468: protein phosphorylation	CDKL5	cyclin-dependent kinase-like 5	0.0022902028514099303
GO:0006468: protein phosphorylation	CFL1	cofilin 1 (non-muscle)	-0.0011782671739032856
GO:0006468: protein phosphorylation	CREB1	cAMP responsive element binding protein 1	0.0006606053514895301
GO:0006468: protein phosphorylation	CSNK2B	casein kinase 2, beta polypeptide	0.0015328764946920695
GO:0006468: protein phosphorylation	GSK3B	glycogen synthase kinase 3 beta 0.0015531071795888414
GO:0006468: protein phosphorylation	GUCY2C	guanylate cyclase 2C (heat stable enterotoxin receptor)	-0.0016372526591578642
GO:0006468: protein phosphorylation	HIPK2	homeodomain interacting protein kinase 2	0.0007724939082316297
GO:0006468: protein phosphorylation	IGFBP3	insulin-like growth factor binding protein 3	0.0008382644580709871
GO:0006468: protein phosphorylation	IRAK1	interleukin-1 receptor-associated kinase 1	-0.0016130541618468864

GO:0006468: protein phosphorylation     JAK2    Janus kinase 2  -3.137260031916692e-5
GO:0006468: protein phosphorylation     LYN     LYN proto-oncogene, Src family tyrosine kinase  -0.00
14239427431591296
GO:0006468: protein phosphorylation     MAP3K5  mitogen-activated protein kinase kinase kinase 5
0.00034350186035423595
GO:0006468: protein phosphorylation     MMD     monocyte to macrophage differentiation-associated
0.0021537390797454724
GO:0006468: protein phosphorylation     MOK     MOK protein kinase      0.001793913507996557
GO:0006468: protein phosphorylation     NEK2    NIMA-related kinase 2   4.958374578080435e-5
GO:0006468: protein phosphorylation     PAK1    p21 protein (Cdc42/Rac)-activated kinase 1      -0.00
2193587250243135
GO:0006468: protein phosphorylation     PBK     PDZ binding kinase      0.0008433623468465556
GO:0006468: protein phosphorylation     PDK1    pyruvate dehydrogenase kinase, isozyme 1        0.001
4293432237704438
GO:0006468: protein phosphorylation     PDK2    pyruvate dehydrogenase kinase, isozyme 2        0.001
4834715708303674
GO:0006468: protein phosphorylation     PICK1   protein interacting with PRKCA 1        -0.0009574612
23837719
GO:0006468: protein phosphorylation     PIK3CD  phosphatidylinositol-4,5-bisphosphate 3-kinase, catal
ytic subunit delta      -0.0007828086515596967
GO:0006468: protein phosphorylation     PKN1    protein kinase N1       -0.0019077501569560065
GO:0006468: protein phosphorylation     PLK1    polo-like kinase 1      0.0010206881598149636
GO:0006468: protein phosphorylation     PLK3    polo-like kinase 3      0.0025700428188143378
GO:0006468: protein phosphorylation     PLK4    polo-like kinase 4      0.0029937321456941063
GO:0006468: protein phosphorylation     PRKCA   protein kinase C, alpha -5.876725573190922e-6
GO:0006468: protein phosphorylation     PRKCD   protein kinase C, delta -0.0011315960280285284
GO:0006468: protein phosphorylation     PRKCZ   protein kinase C, zeta  -0.001604605622205434
GO:0006468: protein phosphorylation     RAF1    Raf-1 proto-oncogene, serine/threonine kinase   0.001
5024938699221525
GO:0006468: protein phosphorylation     RET     ret proto-oncogene      -0.0004922159180050821
GO:0006468: protein phosphorylation     RPS6KA1 ribosomal protein S6 kinase, 90kDa, polypeptide 1
-0.002520030028593816
GO:0006468: protein phosphorylation     STK24   serine/threonine kinase 24      0.002701339451967088
GO:0006468: protein phosphorylation     TEX14   testis expressed 14     0.0017302251876234879
GO:0006468: protein phosphorylation     TGFB1   transforming growth factor, beta 1      -7.3280217973
23938e-5
GO:0006468: protein phosphorylation     TGFB2   transforming growth factor, beta 2      -0.0010623869
165459127
GO:0006468: protein phosphorylation     TGFBR1  transforming growth factor, beta receptor 1     0.000
3409874916133344
GO:0007162: negative regulation of cell adhesion        ACVRL1  activin A receptor type II-like 1
0.0019268955101691455
GO:0007162: negative regulation of cell adhesion        AGER    advanced glycosylation end product-sp
ecific receptor -0.00017354573912856195
GO:0007162: negative regulation of cell adhesion        ANGPT1  angiopoietin 1  0.0008935206257666981
GO:0007162: negative regulation of cell adhesion        RASA1   RAS p21 protein activator (GTPase act
ivating protein) 1      -0.00034402750509101936
GO:0007165: signal transduction ACVRL1  activin A receptor type II-like 1       0.0019434882906068775
GO:0007165: signal transduction ADM     adrenomedullin  0.002237431057860733
GO:0007165: signal transduction AKT1    v-akt murine thymoma viral oncogene homolog 1   0.00073040738
70056507
GO:0007165: signal transduction AKT2    v-akt murine thymoma viral oncogene homolog 2   -0.0006257732
791294791
GO:0007165: signal transduction AMHR2   anti-Mullerian hormone receptor, type II        -0.0002246880
3078219975
GO:0007165: signal transduction ANK3    ankyrin 3, node of Ranvier (ankyrin G)  0.0007000095618207143
GO:0007165: signal transduction AR      androgen receptor       0.002637732403507614
GO:0007165: signal transduction AVP     arginine vasopressin    -0.0009371013629622088
GO:0007165: signal transduction BCL11A  B-cell CLL/lymphoma 11A (zinc finger protein)   -0.0009769692
989772172
GO:0007165: signal transduction C3      complement component 3  0.002024181314913599
GO:0007165: signal transduction CASP8AP2        caspase 8 associated protein 2  0.0012716188181623074
GO:0007165: signal transduction CCL2    chemokine (C-C motif) ligand 2  0.0008164781588547721
GO:0007165: signal transduction CCL7    chemokine (C-C motif) ligand 7  -0.0024021925909729416
GO:0007165: signal transduction CCL8    chemokine (C-C motif) ligand 8  -0.0005902866570882142
GO:0007165: signal transduction CHRNB1  cholinergic receptor, nicotinic, beta 1 (muscle)        -6.97
4939505178637e-6
GO:0007165: signal transduction CHRNB2  cholinergic receptor, nicotinic, beta 2 (neuronal)      -0.00
07782428189131888
GO:0007165: signal transduction CLOCK   clock circadian regulator       0.0002001542150919359
GO:0007165: signal transduction CREB1   cAMP responsive element binding protein 1       0.00065806314
26769614
GO:0007165: signal transduction CSF3R   colony stimulating factor 3 receptor (granulocyte)      -8.33
9144135618882e-5
GO:0007165: signal transduction CSNK2B  casein kinase 2, beta polypeptide       0.001528290022274502
GO:0007165: signal transduction CXCL10  chemokine (C-X-C motif) ligand 10       6.252176489902882e-5
GO:0007165: signal transduction CXCL12  chemokine (C-X-C motif) ligand 12       -0.001208108420582850
3
GO:0007165: signal transduction DEK     DEK proto-oncogene      0.002487075539820206
GO:0007165: signal transduction DOCK1   dedicator of cytokinesis 1      -0.000684075227750657
GO:0007165: signal transduction ECM1    extracellular matrix protein 1  -0.001556783514827781

GO:0007165: signal transduction EDA ectodysplasin A -0.0008142998396472177
GO:0007165: signal transduction EGFR epidermal growth factor receptor 0.0006855542326527749
GO:0007165: signal transduction ERBB4 v-erb-b2 avian erythroblastic leukemia viral oncogene homolog 4 -0.00018282737676140546
GO:0007165: signal transduction ESR1 estrogen receptor 1 -0.0009479087845693358
GO:0007165: signal transduction FAS Fas cell surface death receptor -3.3247989333608365e-5
GO:0007165: signal transduction FGF3 fibroblast growth factor 3 0.0015606321391585846
GO:0007165: signal transduction FGF7 fibroblast growth factor 7 0.0006335279489453929
GO:0007165: signal transduction GATA3 GATA binding protein 3 -3.902895825587418e-5
GO:0007165: signal transduction GDNF glial cell derived neurotrophic factor 0.00044469132721872227
GO:0007165: signal transduction GJA1 gap junction protein, alpha 1, 43kDa -0.00016145881297111375
GO:0007165: signal transduction GOLT1B golgi transport 1B -0.0018204313940101476
GO:0007165: signal transduction GREM1 gremlin 1, DAN family BMP antagonist -0.000827025232494863 1
GO:0007165: signal transduction HIF1A hypoxia inducible factor 1, alpha subunit (basic helix-loop-helix transcription factor) -0.0006463380068602735
GO:0007165: signal transduction IGBP1 immunoglobulin (CD79A) binding protein 1 0.002848761021368232
GO:0007165: signal transduction IGF1 insulin-like growth factor 1 (somatomedin C) 0.00013073943590575084
GO:0007165: signal transduction IGF1R insulin-like growth factor 1 receptor 0.0010718343173586856
GO:0007165: signal transduction IGFBP1 insulin-like growth factor binding protein 1 0.0003878467873103586
GO:0007165: signal transduction IGFBP2 insulin-like growth factor binding protein 2, 36kDa 0.00014475319462091486
GO:0007165: signal transduction IGFBP4 insulin-like growth factor binding protein 4 -0.0012781286022579324
GO:0007165: signal transduction IGFBP6 insulin-like growth factor binding protein 6 -0.0020276698443934186
GO:0007165: signal transduction INHA inhibin, alpha 0.0002116812875126061
GO:0007165: signal transduction INPP4B inositol polyphosphate-4-phosphatase, type II, 105kDa 0.00028982401353553803
GO:0007165: signal transduction IRAK1 interleukin-1 receptor-associated kinase 1 -0.001608217676736459
GO:0007165: signal transduction ITPR1 inositol 1,4,5-trisphosphate receptor, type 1 -0.0007801932860876997
GO:0007165: signal transduction JAK2 Janus kinase 2 -3.184119929580906e-5
GO:0007165: signal transduction KIT v-kit Hardy-Zuckerman 4 feline sarcoma viral oncogene homolog 0.0002746279766027155
GO:0007165: signal transduction LGALS9 lectin, galactoside-binding, soluble, 9 -0.00011686347685723615
GO:0007165: signal transduction LRP8 low density lipoprotein receptor-related protein 8, apolipoprotein e receptor -0.0009315977289665427
GO:0007165: signal transduction LYN LYN proto-oncogene, Src family tyrosine kinase -0.0014201196210848548
GO:0007165: signal transduction MDK midkine (neurite growth-promoting factor 2) 0.0016645696032585083
GO:0007165: signal transduction MOK MOK protein kinase 0.0017891484430256099
GO:0007165: signal transduction NCK2 NCK adaptor protein 2 -0.00023207792304874017
GO:0007165: signal transduction NDRG2 NDRG family member 2 0.002172917084249931
GO:0007165: signal transduction NMU neuromedin U 0.00185005461017852
GO:0007165: signal transduction NRP1 neuropilin 1 -0.0006429554165685961
GO:0007165: signal transduction OR5I1 olfactory receptor, family 5, subfamily I, member 1 0.0018819441389188885
GO:0007165: signal transduction PDGFRB platelet-derived growth factor receptor, beta polypeptide -0.00036099984777883046
GO:0007165: signal transduction PEX11A peroxisomal biogenesis factor 11 alpha 0.0007172058370507033
GO:0007165: signal transduction PGR progesterone receptor 0.00035181374841912174
GO:0007165: signal transduction PIK3CD phosphatidylinositol-4,5-bisphosphate 3-kinase, catalytic subunit delta -0.0007806355534794369
GO:0007165: signal transduction PKN1 protein kinase N1 -0.0019025145368926836
GO:0007165: signal transduction PLCB1 phospholipase C, beta 1 (phosphoinositide-specific) 0.00015369129227155208
GO:0007165: signal transduction PRDM4 PR domain containing 4 0.0008401522472600571
GO:0007165: signal transduction PRKCA protein kinase C, alpha -5.85871572846618e-6
GO:0007165: signal transduction PRKCD protein kinase C, delta -0.0011280425279524247
GO:0007165: signal transduction PRKCZ protein kinase C, zeta -0.001600397382618745
GO:0007165: signal transduction PTK7 protein tyrosine kinase 7 -0.00022523708569734612
GO:0007165: signal transduction RAF1 Raf-1 proto-oncogene, serine/threonine kinase 0.0014984721965360787
GO:0007165: signal transduction RASA1 RAS p21 protein activator (GTPase activating protein) 1 -0.0003489944204112377 6
GO:0007165: signal transduction RASSF8 Ras association (RalGDS/AF-6) domain family (N-terminal) member 8 0.003320192670512 8
GO:0007165: signal transduction RET ret proto-oncogene -0.00049115622013241
GO:0007165: signal transduction RPS6KA1 ribosomal protein S6 kinase, 90kDa, polypeptide 1 -0.0025137535542865577
GO:0007165: signal transduction SNX17 sorting nexin 17 0.0016412085197914947
GO:0007165: signal transduction SOX9 SRY (sex determining region Y)-box 9 -0.000518432093799608

GO:0007165: signal transduction SPHK1 sphingosine kinase 1 0.001810464388038678
GO:0007165: signal transduction STK24 serine/threonine kinase 24 0.0026941011093451223
GO:0007165: signal transduction STMN1 stathmin 1 0.0005589882128292235
GO:0007165: signal transduction STX2 syntaxin 2 0.0003079835404112421
GO:0007165: signal transduction TGFBR1 transforming growth factor, beta receptor 1 0.00033977308512114475
GO:0007165: signal transduction TNFRSF10C tumor necrosis factor receptor superfamily, member 10c, decoy without an intracellular domain 0.00031715867109832004
GO:0007165: signal transduction TNFRSF11A tumor necrosis factor receptor superfamily, member 11a, NFKB activator 0.002775153287852285
GO:0007165: signal transduction TRAIP TRAF interacting protein 0.0010222919425565639
GO:0007165: signal transduction VDR vitamin D (1,25- dihydroxyvitamin D3) receptor 0.0005634964089186078
GO:0007165: signal transduction VEGFC vascular endothelial growth factor C -0.003363283081559841
GO:0007165: signal transduction VLDLR very low density lipoprotein receptor 0.0009526634738594306
GO:0007165: signal transduction ZPR1 ZPR1 zinc finger -0.00020945048144596618
GO:0007179: transforming growth factor beta receptor signaling pathway ACVRL1 activin A receptor type II-like 1 0.0019477170205270313
GO:0007179: transforming growth factor beta receptor signaling pathway AMHR2 anti-Mullerian hormone receptor, type II -0.000224928737072981
GO:0007179: transforming growth factor beta receptor signaling pathway CCL2 chemokine (C-C motif) ligand 2 0.0008183175303271569
GO:0007179: transforming growth factor beta receptor signaling pathway CCNC cyclin C 0.0005688221525635011
GO:0007179: transforming growth factor beta receptor signaling pathway CDK8 cyclin-dependent kinase 8 0.0026173712241212847
GO:0007179: transforming growth factor beta receptor signaling pathway CDKN2B cyclin-dependent kinase inhibitor 2B (p15, inhibits CDK4) -0.001839271938372105
GO:0007179: transforming growth factor beta receptor signaling pathway CITED1 Cbp/p300-interacting transactivator, with Glu/Asp-rich carboxy-terminal domain, 1 0.0028244334171750657
GO:0007179: transforming growth factor beta receptor signaling pathway CREB1 cAMP responsive element binding protein 1 0.0006603604642300682
GO:0007179: transforming growth factor beta receptor signaling pathway E2F4 E2F transcription factor 4, p107/p130-binding -0.0023502401060925245
GO:0007179: transforming growth factor beta receptor signaling pathway ENG endoglin 0.000828789595926268
GO:0007179: transforming growth factor beta receptor signaling pathway FOXH1 forkhead box H1 -0.0013747248438405282
GO:0007179: transforming growth factor beta receptor signaling pathway GCNT2 glucosaminyl (N-acetyl) transferase 2, I-branching enzyme (I blood group) 0.0024371068357180556
GO:0007179: transforming growth factor beta receptor signaling pathway HIPK2 homeodomain interacting protein kinase 2 0.0007721425017938067
GO:0007179: transforming growth factor beta receptor signaling pathway MYC v-myc avian myelocytomatosis viral oncogene homolog -0.0011384836277658153
GO:0007179: transforming growth factor beta receptor signaling pathway PML promyelocytic leukemia -0.0006808554572287163
GO:0007179: transforming growth factor beta receptor signaling pathway PRKCZ protein kinase C, zeta -0.0016038789679217638
GO:0007179: transforming growth factor beta receptor signaling pathway RHOA ras homolog family member A 0.000671138134059341
GO:0007179: transforming growth factor beta receptor signaling pathway SERPINE1 serpin peptidase inhibitor, clade E (nexin, plasminogen activator inhibitor type 1), member 1 0.00011209206002378357
GO:0007179: transforming growth factor beta receptor signaling pathway SKI SKI proto-oncogene -0.0006716534248830595
GO:0007179: transforming growth factor beta receptor signaling pathway TFDP1 transcription factor Dp-1 0.0014487226922070134
GO:0007179: transforming growth factor beta receptor signaling pathway TGFB1 transforming growth factor, beta 1 -7.328683779457394e-5
GO:0007179: transforming growth factor beta receptor signaling pathway TGFB2 transforming growth factor, beta 2 -0.001061945933123548
GO:0007179: transforming growth factor beta receptor signaling pathway TGFB3 transforming growth factor, beta 3 -0.0018281370084749255
GO:0007179: transforming growth factor beta receptor signaling pathway TGFBR1 transforming growth factor, beta receptor 1 0.00034086779108912695
GO:0007179: transforming growth factor beta receptor signaling pathway TP53 tumor protein p53 0.0011789064079598338
GO:0007179: transforming growth factor beta receptor signaling pathway WWTR1 WW domain containing transcription regulator 1 0.0009030502462025904
GO:0008015: blood circulation ACVRL1 activin A receptor type II-like 1 0.0019546331310049602
GO:0008015: blood circulation ADM adrenomedullin 0.0022504405831868593
GO:0008015: blood circulation CXCL10 chemokine (C-X-C motif) ligand 10 6.365246412149694e-5
GO:0008015: blood circulation CXCL12 chemokine (C-X-C motif) ligand 12 -0.0012155952786464235
GO:0008015: blood circulation E2F4 E2F transcription factor 4, p107/p130-binding -0.0023586915419635014
GO:0008015: blood circulation HOXB2 homeobox B2 0.002787874089650523
GO:0008217: regulation of blood pressure ACVRL1 activin A receptor type II-like 1 0.001958271553016392
GO:0008217: regulation of blood pressure AGT angiotensinogen (serpin peptidase inhibitor, clade A, member 8) -0.0011463655435048643

GO:0008217: regulation of blood pressure        HMOX1   heme oxygenase (decycling) 1    -0.0002198384
5923779878
GO:0008217: regulation of blood pressure        LEP     leptin  0.003219784341984936
GO:0008217: regulation of blood pressure        LRP5    low density lipoprotein receptor-related prot
ein 5   2.941578442795664e-5
GO:0008285: negative regulation of cell proliferation   ACVRL1  activin A receptor type II-like 1
0.0019450423689848504
GO:0008285: negative regulation of cell proliferation   ADM     adrenomedullin  0.0022391940259310085
GO:0008285: negative regulation of cell proliferation   AGT     angiotensinogen (serpin peptidase inh
ibitor, clade A, member 8)      -0.0011340968087181545
GO:0008285: negative regulation of cell proliferation   AIMP2   aminoacyl tRNA synthetase complex-int
eracting multifunctional protein 2      0.0012097239612722768
GO:0008285: negative regulation of cell proliferation   ALDH1A2 aldehyde dehydrogenase 1 family, memb
er A2   -0.003020945674775801
GO:0008285: negative regulation of cell proliferation   APC     adenomatous polyposis coli      0.000
6440335004019952
GO:0008285: negative regulation of cell proliferation   AR      androgen receptor       0.00263976782
081629
GO:0008285: negative regulation of cell proliferation   ATF5    activating transcription factor 5
-0.002736752461099586
GO:0008285: negative regulation of cell proliferation   BAK1    BCL2-antagonist/killer 1        -0.00
18772370398121953
GO:0008285: negative regulation of cell proliferation   BCHE    butyrylcholinesterase   -6.9765294945
67089e-6
GO:0008285: negative regulation of cell proliferation   BMP4    bone morphogenetic protein 4    -0.00
03227789396435446
GO:0008285: negative regulation of cell proliferation   CD9     CD9 molecule    -0.002587709018700915
GO:0008285: negative regulation of cell proliferation   CDC6    cell division cycle 6   0.00135487999
59442628
GO:0008285: negative regulation of cell proliferation   CDKN2A  cyclin-dependent kinase inhibitor 2A
0.0017593414912815564
GO:0008285: negative regulation of cell proliferation   CDKN2B  cyclin-dependent kinase inhibitor 2B
 (p15, inhibits CDK4)   -0.0018366482265428053
GO:0008285: negative regulation of cell proliferation   CHD5    chromodomain helicase DNA binding pro
tein 5  -0.0005192094837780242
GO:0008285: negative regulation of cell proliferation   CIB1    calcium and integrin binding 1 (calmy
rin)    5.457377976086593e-5
GO:0008285: negative regulation of cell proliferation   CSNK2B  casein kinase 2, beta polypeptide
0.001529713561061444
GO:0008285: negative regulation of cell proliferation   CTNNB1  catenin (cadherin-associated protei
n), beta 1, 88kDa       -0.00011700726060175801
GO:0008285: negative regulation of cell proliferation   CYP1B1  cytochrome P450, family 1, subfamily
 B, polypeptide 1       0.0003978775054213934
GO:0008285: negative regulation of cell proliferation   CYP27B1 cytochrome P450, family 27, subfamily
B, polypeptide 1        -0.00047153962910640715
GO:0008285: negative regulation of cell proliferation   DHCR24  24-dehydrocholesterol reductase -0.00
17085568606880306
GO:0008285: negative regulation of cell proliferation   DRD2    dopamine receptor D2    -0.0002341446
7350260133
GO:0008285: negative regulation of cell proliferation   ERBB4   v-erb-b2 avian erythroblastic leukemi
a viral oncogene homolog 4      -0.00018313082969778535
GO:0008285: negative regulation of cell proliferation   FGFR3   fibroblast growth factor receptor 3
0.00022285984370167545
GO:0008285: negative regulation of cell proliferation   FLT3    fms-related tyrosine kinase 3   -0.00
06819895209086498
GO:0008285: negative regulation of cell proliferation   GATA3   GATA binding protein 3  -3.9013435158
00615e-5
GO:0008285: negative regulation of cell proliferation   GLI3    GLI family zinc finger 3        -0.00
2153050127690327
GO:0008285: negative regulation of cell proliferation   HMGA1   high mobility group AT-hook 1   -0.00
031887073191870735
GO:0008285: negative regulation of cell proliferation   IGF1    insulin-like growth factor 1 (somatom
edin C) 0.00013110964500248775
GO:0008285: negative regulation of cell proliferation   IGFBP3  insulin-like growth factor binding pr
otein 3 0.0008371353197730288
GO:0008285: negative regulation of cell proliferation   IGFBP6  insulin-like growth factor binding pr
otein 6 -0.002029294690595642
GO:0008285: negative regulation of cell proliferation   INHBA   inhibin, beta A -0.001354191065560574
8
GO:0008285: negative regulation of cell proliferation   IRF6    interferon regulatory factor 6  -0.00
16947300086202334
GO:0008285: negative regulation of cell proliferation   JAK2    Janus kinase 2  -3.1680093694854645e-
5
GO:0008285: negative regulation of cell proliferation   KLF11   Kruppel-like factor 11  0.00061723921
48230501
GO:0008285: negative regulation of cell proliferation   KMT2A   lysine (K)-specific methyltransferase
2A      0.0006610398783569432
GO:0008285: negative regulation of cell proliferation   LYN     LYN proto-oncogene, Src family tyrosi
ne kinase       -0.0014213375854111863
GO:0008285: negative regulation of cell proliferation   MSX1    msh homeobox 1  -0.002770886085654946
GO:0008285: negative regulation of cell proliferation   NCK2    NCK adaptor protein 2   -0.0002321366

8445202468
GO:0008285: negative regulation of cell proliferation   NDRG1   N-myc downstream regulated 1    -0.0009414853496860374
GO:0008285: negative regulation of cell proliferation   NF2     neurofibromin 2 (merlin)        -0.0012725663301033127
GO:0008285: negative regulation of cell proliferation   NOTCH1  notch 1 0.0005190883491611531
GO:0008285: negative regulation of cell proliferation   PDX1    pancreatic and duodenal homeobox 1      0.00025751202812593035
GO:0008285: negative regulation of cell proliferation   PHOX2B  paired-like homeobox 2b 0.00039063016146476787
GO:0008285: negative regulation of cell proliferation   PML     promyelocytic leukemia  -0.0006802973368918827
GO:0008285: negative regulation of cell proliferation   POU1F1  POU class 1 homeobox 1  0.00013753675609704238
GO:0008285: negative regulation of cell proliferation   PRKCA   protein kinase C, alpha -5.880755402195293e-6
GO:0008285: negative regulation of cell proliferation   PROX1   prospero homeobox 1     0.0011222581274418979
GO:0008285: negative regulation of cell proliferation   PTEN    phosphatase and tensin homolog  1.699164479750552e-5
GO:0008285: negative regulation of cell proliferation   PTPN2   protein tyrosine phosphatase, non-receptor type 2    -0.0003294657092011837
GO:0008285: negative regulation of cell proliferation   PTPRJ   protein tyrosine phosphatase, receptor type, J       -0.00033868018785245137
GO:0008285: negative regulation of cell proliferation   RAF1    Raf-1 proto-oncogene, serine/threonine kinase        0.0014997181483652174
GO:0008285: negative regulation of cell proliferation   RARG    retinoic acid receptor, gamma   -0.0023208919093867454
GO:0008285: negative regulation of cell proliferation   RBBP4   retinoblastoma binding protein 4        -0.0018615449201155013
GO:0008285: negative regulation of cell proliferation   RBPJ    recombination signal binding protein for immunoglobulin kappa J region      0.0009819875758064528
GO:0008285: negative regulation of cell proliferation   RXRA    retinoid X receptor, alpha      0.0011157885166322512
GO:0008285: negative regulation of cell proliferation   SERPINE2        serpin peptidase inhibitor, clade E (nexin, plasminogen activator inhibitor type 1), member 2        0.002504422102593758
GO:0008285: negative regulation of cell proliferation   SFRP1   secreted frizzled-related protein 1     0.0012828129405696837
GO:0008285: negative regulation of cell proliferation   SIRT2   sirtuin 2       -0.0008528284020049035
GO:0008285: negative regulation of cell proliferation   SKI     SKI proto-oncogene      -0.0006708552260220732
GO:0008285: negative regulation of cell proliferation   SLIT2   slit homolog 2 (Drosophila)     -0.0016212717145099955
GO:0008285: negative regulation of cell proliferation   SOX4    SRY (sex determining region Y)-box 4    -3.823810295875508e-5
GO:0008285: negative regulation of cell proliferation   TFAP2A  transcription factor AP-2 alpha (activating enhancer binding protein 2 alpha)        0.0004801644901072095
GO:0008285: negative regulation of cell proliferation   TGFB1   transforming growth factor, beta 1      -7.306558685080605e-5
GO:0008285: negative regulation of cell proliferation   TGFB2   transforming growth factor, beta 2      -0.001060124302051384
GO:0008285: negative regulation of cell proliferation   TGFB3   transforming growth factor, beta 3      -0.0018249530927945778
GO:0008285: negative regulation of cell proliferation   TP53    tumor protein p53       0.0011765917919464677
GO:0008285: negative regulation of cell proliferation   TP73    tumor protein p73       0.0010285762628292007
GO:0008285: negative regulation of cell proliferation   VDR     vitamin D (1,25- dihydroxyvitamin D3) receptor        0.0005638134434367345
GO:0008285: negative regulation of cell proliferation   VEGFC   vascular endothelial growth factor C    -0.0033660776143192538
GO:0008285: negative regulation of cell proliferation   WT1     Wilms tumor 1   -0.0005072784612433903
GO:0010596: negative regulation of endothelial cell migration   ACVRL1  activin A receptor type II-like 1    0.0019392544309793536
GO:0010596: negative regulation of endothelial cell migration   AGER    advanced glycosylation end product-specific receptor -0.00017405620776148918
GO:0010596: negative regulation of endothelial cell migration   SLIT2   slit homolog 2 (Drosophila)     -0.001615558291241135
GO:0010596: negative regulation of endothelial cell migration   THBS1   thrombospondin 1        -0.0010327682943718805
GO:0010596: negative regulation of endothelial cell migration   VASH1   vasohibin 1     0.0005975192973211145
GO:0010862: positive regulation of pathway-restricted SMAD protein phosphorylation      ACVRL1  activin A receptor type II-like 1    0.0019429630682778376
GO:0010862: positive regulation of pathway-restricted SMAD protein phosphorylation      BMP4    bone morphogenetic protein 4         -0.00032248209099153513
GO:0010862: positive regulation of pathway-restricted SMAD protein phosphorylation      BMP7    bone morphogenetic protein 7         0.0008550989828034137
GO:0010862: positive regulation of pathway-restricted SMAD protein phosphorylation      CSNK2B  casei

n kinase 2, beta polypeptide    0.0015282231022148666
GO:0010862: positive regulation of pathway-restricted SMAD protein phosphorylation      ENG     endoglin     0.0008261881775255032
GO:0010862: positive regulation of pathway-restricted SMAD protein phosphorylation      GDF3    growth differentiation factor 3      -0.0018830764182839468
GO:0010862: positive regulation of pathway-restricted SMAD protein phosphorylation      INHA    inhibin, alpha       0.00021133483828984223
GO:0010862: positive regulation of pathway-restricted SMAD protein phosphorylation      INHBA   inhibin, beta A      -0.0013528337548718006
GO:0010862: positive regulation of pathway-restricted SMAD protein phosphorylation      TGFB1   transforming growth factor, beta 1   -7.314932938012394e-5
GO:0010862: positive regulation of pathway-restricted SMAD protein phosphorylation      TGFB2   transforming growth factor, beta 2   -0.00105891742936502
GO:0010862: positive regulation of pathway-restricted SMAD protein phosphorylation      TGFB3   transforming growth factor, beta 3   -0.0018232911276634782
GO:0010862: positive regulation of pathway-restricted SMAD protein phosphorylation      TGFBR1  transforming growth factor, beta receptor 1  0.0003398629920313266
GO:0010862: positive regulation of pathway-restricted SMAD protein phosphorylation      TTK     TTK protein kinase   0.0010856640624797396
GO:0023014: signal transduction by protein phosphorylation      ACVRL1  activin A receptor type II-like 1    0.0019476480050772886
GO:0023014: signal transduction by protein phosphorylation      AMHR2   anti-Mullerian hormone receptor, type II     -0.00022496307320129467
GO:0023014: signal transduction by protein phosphorylation      INSR    insulin receptor        -0.0013661233431596816
GO:0023014: signal transduction by protein phosphorylation      LYN     LYN proto-oncogene, Src family tyrosine kinase       -0.0014238942193058263
GO:0023014: signal transduction by protein phosphorylation      STK24   serine/threonine kinase 24      0.002700100931704592
GO:0023014: signal transduction by protein phosphorylation      TGFB2   transforming growth factor, beta 2   -0.0010619596012031707
GO:0023014: signal transduction by protein phosphorylation      TGFBR1  transforming growth factor, beta receptor 1  0.0003406119658438517
GO:0030308: negative regulation of cell growth  ACVRL1  activin A receptor type II-like 1       0.001946593706219525
GO:0030308: negative regulation of cell growth  AGT     angiotensinogen (serpin peptidase inhibitor, clade A, member 8)     -0.0011366798436235894
GO:0030308: negative regulation of cell growth  BCL2    B-cell CLL/lymphoma 2   -5.235961386149368e-6
GO:0030308: negative regulation of cell growth  CDKN2A  cyclin-dependent kinase inhibitor 2A    0.0017620138049313189
GO:0030308: negative regulation of cell growth  CRYAB   crystallin, alpha B     0.0009872064460255266
GO:0030308: negative regulation of cell growth  CYP27B1 cytochrome P450, family 27, subfamily B, polypeptide 1       -0.0004725078552287108
GO:0030308: negative regulation of cell growth  GJA1    gap junction protein, alpha 1, 43kDa    -0.0016379374024781912
GO:0030308: negative regulation of cell growth  GREM1   gremlin 1, DAN family BMP antagonist    -0.0008299974383682905
GO:0030308: negative regulation of cell growth  INHBA   inhibin, beta A -0.001357020545078564
GO:0030308: negative regulation of cell growth  MSX1    msh homeobox 1  -0.002774196213296862
GO:0030308: negative regulation of cell growth  NF2     neurofibromin 2 (merlin)        -0.0012725211664658478
GO:0030308: negative regulation of cell growth  PML     promyelocytic leukemia  -0.0006799242559832311
GO:0030308: negative regulation of cell growth  PRDM4   PR domain containing 4  0.0008408507580063661
GO:0030308: negative regulation of cell growth  PSRC1   proline/serine-rich coiled-coil 1       0.0006490186907055039
GO:0030308: negative regulation of cell growth  PTPRJ   protein tyrosine phosphatase, receptor type, J      -0.0003385804136027844
GO:0030308: negative regulation of cell growth  SERPINE2        serpin peptidase inhibitor, clade E (nexin, plasminogen activator inhibitor type 1), member 2      0.002506776530894684
GO:0030308: negative regulation of cell growth  SFRP1   secreted frizzled-related protein 1     0.0012865837019925064
GO:0030308: negative regulation of cell growth  SIRT1   sirtuin 1       -1.0999163846878799e-6
GO:0030308: negative regulation of cell growth  SLIT2   slit homolog 2 (Drosophila)     -0.0016231315668071005
GO:0030308: negative regulation of cell growth  TGFB1   transforming growth factor, beta 1      -7.352850406007194e-5
GO:0030308: negative regulation of cell growth  TGFB2   transforming growth factor, beta 2      -0.0010613630616584167
GO:0030308: negative regulation of cell growth  TP53    tumor protein p53       0.001178636425246807
GO:0030308: negative regulation of cell growth  WT1     Wilms tumor 1   -0.000508029502215423
GO:0030336: negative regulation of cell migration       ACVRL1  activin A receptor type II-like 1 0.001941793158108352
GO:0030336: negative regulation of cell migration       ARPIN   actin-related protein 2/3 complex inhibitor  -0.00014142792688151743
GO:0030336: negative regulation of cell migration       BCL2    B-cell CLL/lymphoma 2   -5.09805562685636e-6
GO:0030336: negative regulation of cell migration       CX3CR1  chemokine (C-X3-C motif) receptor 1 -0.0008561084499457646
GO:0030336: negative regulation of cell migration       CYP1B1  cytochrome P450, family 1, subfamily B, polypeptide 1       0.00039733772998888823

GO:0030336: negative regulation of cell migration	DACH1	dachshund family transcription factor 1	0.0020864773603336663
GO:0030336: negative regulation of cell migration	DAG1	dystroglycan 1 (dystrophin-associated glycoprotein 1) 0.0003612098521768637
GO:0030336: negative regulation of cell migration	DRD2	dopamine receptor D2	-0.0002338500151894021
GO:0030336: negative regulation of cell migration	ENG	endoglin	0.0008254162355259508
GO:0030336: negative regulation of cell migration	KANK1	KN motif and ankyrin repeat domains 1 0.0025304489406819818
GO:0030336: negative regulation of cell migration	NF2	neurofibromin 2 (merlin)	-0.001270521802846068
GO:0030336: negative regulation of cell migration	PTEN	phosphatase and tensin homolog	1.7255852294450517e-5
GO:0030336: negative regulation of cell migration	PTPRJ	protein tyrosine phosphatase, receptor type, J	-0.0003382301403196741
GO:0030336: negative regulation of cell migration	SERPINE1	serpin peptidase inhibitor, clade E (nexin, plasminogen activator inhibitor type 1), member 1	0.00011234506285016845
GO:0030336: negative regulation of cell migration	SFRP1	secreted frizzled-related protein 1 0.0012798458679897996
GO:0030336: negative regulation of cell migration	SHH	sonic hedgehog	0.0005990921539456026
GO:0030336: negative regulation of cell migration	SLIT2	slit homolog 2 (Drosophila)	-0.0016182399277552251
GO:0030336: negative regulation of cell migration	STK24	serine/threonine kinase 24	0.0026916715576177784
GO:0030336: negative regulation of cell migration	SULF1	sulfatase 1	-0.0008044807316197394
GO:0030336: negative regulation of cell migration	WNT4	wingless-type MMTV integration site family, member 4 -0.00025158545905788987
GO:0030509: BMP signaling pathway	ACVRL1	activin A receptor type II-like 1	0.0019435026586675321
GO:0030509: BMP signaling pathway	BMP4	bone morphogenetic protein 4	-0.00032207533292147485
GO:0030509: BMP signaling pathway	BMP7	bone morphogenetic protein 7	0.0008562001551039508
GO:0030509: BMP signaling pathway	DLX5	distal-less homeobox 5	-0.0032850254520641357
GO:0030509: BMP signaling pathway	EGR1	early growth response 1 0.0010933469015226146
GO:0030509: BMP signaling pathway	ENG	endoglin	0.0008263497207037059
GO:0030509: BMP signaling pathway	FGF8	fibroblast growth factor 8 (androgen-induced)	0.000983422510341201
GO:0030509: BMP signaling pathway	GATA4	GATA binding protein 4	-0.0010940148851977716
GO:0030509: BMP signaling pathway	GDF3	growth differentiation factor 3 -0.001883718475608416
GO:0030509: BMP signaling pathway	LEF1	lymphoid enhancer-binding factor 1	-9.96163711067701e-5
GO:0030509: BMP signaling pathway	SKI	SKI proto-oncogene	-0.0006709456968314366
GO:0030513: positive regulation of BMP signaling pathway	ACVRL1	activin A receptor type II-like 1	0.0019424986883671079
GO:0030513: positive regulation of BMP signaling pathway	BMP4	bone morphogenetic protein 4 -0.00032236645373537483
GO:0030513: positive regulation of BMP signaling pathway	ENG	endoglin	0.0008258856499784682
GO:0030513: positive regulation of BMP signaling pathway	GATA4	GATA binding protein 4	-0.0010939836837037238
GO:0030513: positive regulation of BMP signaling pathway	GATA6	GATA binding protein 6	-2.8058408187496483e-5
GO:0030513: positive regulation of BMP signaling pathway	HES1	hes family bHLH transcription factor 1	-0.0009102898038528625
GO:0030513: positive regulation of BMP signaling pathway	MSX1	msh homeobox 1	-0.0027669619221695565
GO:0030513: positive regulation of BMP signaling pathway	NOTCH1	notch 1 0.0005184455403721949
GO:0030513: positive regulation of BMP signaling pathway	RBPJ	recombination signal binding protein for immunoglobulin kappa J region	0.000980730626113904
GO:0030513: positive regulation of BMP signaling pathway	SOX11	SRY (sex determining region Y)-box 11	-0.00020882416755528332
GO:0030513: positive regulation of BMP signaling pathway	SULF1	sulfatase 1	-0.0008052128386622086
GO:0032332: positive regulation of chondrocyte differentiation	ACVRL1	activin A receptor type II-like 1	0.00195365183251654
GO:0032332: positive regulation of chondrocyte differentiation	GLI3	GLI family zinc finger 3 -0.00216507680922713
GO:0032332: positive regulation of chondrocyte differentiation	IHH	indian hedgehog -0.002052110766871769
GO:0032332: positive regulation of chondrocyte differentiation	POR	P450 (cytochrome) oxidoreductase	0.0004601507885058436
GO:0032332: positive regulation of chondrocyte differentiation	SOX9	SRY (sex determining region Y)-box 9	-0.0005210372975415022
GO:0032924: activin receptor signaling pathway	ACVRL1	activin A receptor type II-like 1	0.0019500099733605397
GO:0032924: activin receptor signaling pathway	INHBA	inhibin, beta A -0.0013605193067040258
GO:0032924: activin receptor signaling pathway	TGFBR1	transforming growth factor, beta receptor 1 0.00034160273668654595
GO:0035313: wound healing, spreading of epidermal cells ACVRL1	activin A receptor type II-like 1 0.001939685751628529

GO:0035313: wound healing, spreading of epidermal cells ARHGAP24 Rho GTPase activating protein 24 -0.000932441174060325
GO:0035313: wound healing, spreading of epidermal cells COL5A1 collagen, type V, alpha 1 -4.841118603584268e-5
GO:0043535: regulation of blood vessel endothelial cell migration ACVRL1 activin A receptor type II-like 1 0.0019411176309128094
GO:0043535: regulation of blood vessel endothelial cell migration EFNA1 ephrin-A1 -0.0005207594575803089
GO:0043537: negative regulation of blood vessel endothelial cell migration ACVRL1 activin A receptor type II-like 1 0.0019350997676852314
GO:0043537: negative regulation of blood vessel endothelial cell migration CSNK2B casein kinase 2, beta polypeptide 0.001520208378070712
GO:0043537: negative regulation of blood vessel endothelial cell migration HMGB1 high mobility group box 1 -0.0007741255427552253
GO:0043537: negative regulation of blood vessel endothelial cell migration TGFB1 transforming growth factor, beta 1 -7.219485695222219e-5
GO:0043537: negative regulation of blood vessel endothelial cell migration THBS1 thrombospondin 1 -0.0010305616462237103
GO:0043537: negative regulation of blood vessel endothelial cell migration VASH1 vasohibin 1 0.000596413089658595
GO:0045602: negative regulation of endothelial cell differentiation ACVRL1 activin A receptor type II-like 1 0.0019557258365855866
GO:0045603: positive regulation of endothelial cell differentiation ACVRL1 activin A receptor type II-like 1 0.0019386631618379942
GO:0045603: positive regulation of endothelial cell differentiation ALOX12 arachidonate 12-lipoxygenase -0.0019479874804598785
GO:0045603: positive regulation of endothelial cell differentiation BMP4 bone morphogenetic protein 4 -0.0003214854620348941
GO:0045603: positive regulation of endothelial cell differentiation CTNNB1 catenin (cadherin-associated protein), beta 1, 88kDa -0.00011743315777938882
GO:0045603: positive regulation of endothelial cell differentiation NOTCH1 notch 1 0.000516657458786623
GO:0045766: positive regulation of angiogenesis ACVRL1 activin A receptor type II-like 1 0.0019484576661752933
GO:0045766: positive regulation of angiogenesis ADM adrenomedullin 0.002243134031581997
GO:0045766: positive regulation of angiogenesis ALOX12 arachidonate 12-lipoxygenase -0.0019585587277376458
GO:0045766: positive regulation of angiogenesis C3 complement component 3 0.002029742039891386
GO:0045766: positive regulation of angiogenesis CX3CL1 chemokine (C-X3-C motif) ligand 1 0.0021230040437213837
GO:0045766: positive regulation of angiogenesis CX3CR1 chemokine (C-X3-C motif) receptor 1 -0.000858762972197624
GO:0045766: positive regulation of angiogenesis CYP1B1 cytochrome P450, family 1, subfamily B, polypeptide 1 0.00039841652308391946
GO:0045766: positive regulation of angiogenesis ECM1 extracellular matrix protein 1 -0.0015619311254693367
GO:0045766: positive regulation of angiogenesis GATA2 GATA binding protein 2 -0.0004529823385431144
GO:0045766: positive regulation of angiogenesis GATA4 GATA binding protein 4 -0.001097592728648809
GO:0045766: positive regulation of angiogenesis GATA6 GATA binding protein 6 -2.7957635891901486e-5
GO:0045766: positive regulation of angiogenesis GREM1 gremlin 1, DAN family BMP antagonist -0.0008303460376319977
GO:0045766: positive regulation of angiogenesis HIF1A hypoxia inducible factor 1, alpha subunit (basic helix-loop-helix transcription factor) -0.0006478370845735985
GO:0045766: positive regulation of angiogenesis HIPK2 homeodomain interacting protein kinase 2 0.0007724192858138796
GO:0045766: positive regulation of angiogenesis HMOX1 heme oxygenase (decycling) 1 -0.0002182584415909265
GO:0045766: positive regulation of angiogenesis ISL1 ISL LIM homeobox 1 7.865100337365996e-5
GO:0045766: positive regulation of angiogenesis NR2E1 nuclear receptor subfamily 2, group E, member 1 -0.002981664247892107
GO:0045766: positive regulation of angiogenesis PRKCA protein kinase C, alpha -5.8659134872811035e-6
GO:0045766: positive regulation of angiogenesis SERPINE1 serpin peptidase inhibitor, clade E (nexin, plasminogen activator inhibitor type 1), member 1 0.00011210625265471011
GO:0045766: positive regulation of angiogenesis SPHK1 sphingosine kinase 1 0.0018153777778733007
GO:0045766: positive regulation of angiogenesis THBS1 thrombospondin 1 -0.0010384229136312746
GO:0045766: positive regulation of angiogenesis TWIST1 twist family bHLH transcription factor 1 -0.0013453177836251868
GO:0045766: positive regulation of angiogenesis VEGFA vascular endothelial growth factor A 0.0005970545210815666
GO:0045766: positive regulation of angiogenesis VEGFC vascular endothelial growth factor C -0.003372177995694796
GO:0045766: positive regulation of angiogenesis WNT5A wingless-type MMTV integration site family, member 5A -0.0006692282805074454
GO:0045893: positive regulation of transcription, DNA-templated ACVRL1 activin A receptor type II-like 1 0.0019424699523251574
GO:0045893: positive regulation of transcription, DNA-templated AGT angiotensinogen (serpin peptidase inhibitor, clade A, member 8) -0.0011319510858498066

GO:0045893: positive regulation of transcription, DNA-templated ALX1 ALX homeobox 1 0.0022904363409177466
GO:0045893: positive regulation of transcription, DNA-templated AR androgen receptor 0.002636593340557388
GO:0045893: positive regulation of transcription, DNA-templated ATAD2 ATPase family, AAA domain containing 2 -4.789011084695563e-5
GO:0045893: positive regulation of transcription, DNA-templated ATF5 activating transcription factor 5 -0.0027330533475965753
GO:0045893: positive regulation of transcription, DNA-templated AXIN1 axin 1 -0.0007324973990418599
GO:0045893: positive regulation of transcription, DNA-templated BLM Bloom syndrome, RecQ helicase-like 0.0005237542326881846
GO:0045893: positive regulation of transcription, DNA-templated BMP4 bone morphogenetic protein 4 -0.000322339146843713
GO:0045893: positive regulation of transcription, DNA-templated BMP7 bone morphogenetic protein 7 0.0008551480720348394
GO:0045893: positive regulation of transcription, DNA-templated BRCA2 breast cancer 2, early onset -1.019444672387354le-5
GO:0045893: positive regulation of transcription, DNA-templated CCNE1 cyclin E1 0.00037051512478497605
GO:0045893: positive regulation of transcription, DNA-templated CDKN2A cyclin-dependent kinase inhibitor 2A 0.0017565407423127842
GO:0045893: positive regulation of transcription, DNA-templated CITED1 Cbp/p300-interacting transactivator, with Glu/Asp-rich carboxy-terminal domain, 1 0.0028158909298131732
GO:0045893: positive regulation of transcription, DNA-templated CLOCK clock circadian regulator 0.000200143886382209
GO:0045893: positive regulation of transcription, DNA-templated COL1A1 collagen, type I, alpha 1 -0.0005254580146218142
GO:0045893: positive regulation of transcription, DNA-templated CREB1 cAMP responsive element binding protein 1 0.0006576257344525607
GO:0045893: positive regulation of transcription, DNA-templated CTNNB1 catenin (cadherin-associated protein), beta 1, 88kDa -0.00011693183698617324
GO:0045893: positive regulation of transcription, DNA-templated DLX5 distal-less homeobox 5 -0.0032809466100890573
GO:0045893: positive regulation of transcription, DNA-templated DNAJC2 DnaJ (Hsp40) homolog, subfamily C, member 2 -0.0009830605135970505
GO:0045893: positive regulation of transcription, DNA-templated E2F1 E2F transcription factor 1 0.0020960157520390453
GO:0045893: positive regulation of transcription, DNA-templated E2F3 E2F transcription factor 3 0.00216029774663673
GO:0045893: positive regulation of transcription, DNA-templated EGR1 early growth response 1 0.001092277571488665
GO:0045893: positive regulation of transcription, DNA-templated EGR2 early growth response 2 0.0014373729383497186
GO:0045893: positive regulation of transcription, DNA-templated ERBB4 v-erb-b2 avian erythroblastic leukemia viral oncogene homolog 4 -0.00018253052870426654
GO:0045893: positive regulation of transcription, DNA-templated ESR1 estrogen receptor 1 -0.0009472384920453687
GO:0045893: positive regulation of transcription, DNA-templated ETS2 v-ets avian erythroblastosis virus E26 oncogene homolog 2 -0.0010147518385565995
GO:0045893: positive regulation of transcription, DNA-templated FGF7 fibroblast growth factor 7 0.0006332387776418405
GO:0045893: positive regulation of transcription, DNA-templated FOXA2 forkhead box A2 -1.674222153304094e-5
GO:0045893: positive regulation of transcription, DNA-templated FOXC1 forkhead box C1 -2.2422478273078076e-5
GO:0045893: positive regulation of transcription, DNA-templated FOXC2 forkhead box C2 (MFH-1, mesenchyme forkhead 1) 0.0017469206154163352
GO:0045893: positive regulation of transcription, DNA-templated FOXH1 forkhead box H1 -0.0013699111435599694
GO:0045893: positive regulation of transcription, DNA-templated FOXM1 forkhead box M1 0.00019684151868649364
GO:0045893: positive regulation of transcription, DNA-templated FOXO1 forkhead box O1 0.0017988886680514536
GO:0045893: positive regulation of transcription, DNA-templated FOXO3 forkhead box O3 0.0011990804052633508
GO:0045893: positive regulation of transcription, DNA-templated FZD7 frizzled class receptor 7 0.0010694453897585469
GO:0045893: positive regulation of transcription, DNA-templated GATA3 GATA binding protein 3 -3.879720197696264e-5
GO:0045893: positive regulation of transcription, DNA-templated GATA4 GATA binding protein 4 -0.0010938863353998412
GO:0045893: positive regulation of transcription, DNA-templated GLI1 GLI family zinc finger 1 -0.0012959183367459665
GO:0045893: positive regulation of transcription, DNA-templated GLI2 GLI family zinc finger 2 0.0018520766205114227
GO:0045893: positive regulation of transcription, DNA-templated GLI3 GLI family zinc finger 3 -0.002149767998237585
GO:0045893: positive regulation of transcription, DNA-templated HDAC2 histone deacetylase 2 -0.001206595709543229
GO:0045893: positive regulation of transcription, DNA-templated HIF1A hypoxia inducible factor 1, a

lpha subunit (basic helix-loop-helix transcription factor) -0.0006460986295605767
GO:0045893: positive regulation of transcription, DNA-templated HINFP histone H4 transcription fact
or 0.0009917191211945696
GO:0045893: positive regulation of transcription, DNA-templated HIPK2 homeodomain interacting prote
in kinase 2 0.0007703154598295992
GO:0045893: positive regulation of transcription, DNA-templated HMGA1 high mobility group AT-hook 1
-0.00031855269002779304
GO:0045893: positive regulation of transcription, DNA-templated HMGA2 high mobility group AT-hook 2
0.0015078669541719338
GO:0045893: positive regulation of transcription, DNA-templated HMGB2 high mobility group box 2
0.00030640789679783946
GO:0045893: positive regulation of transcription, DNA-templated ID2 inhibitor of DNA binding 2, d
ominant negative helix-loop-helix protein 6.014733886170636e-5
GO:0045893: positive regulation of transcription, DNA-templated IFNA2 interferon, alpha 2 -0.00
17160038228046428
GO:0045893: positive regulation of transcription, DNA-templated IGF1 insulin-like growth factor 1
(somatomedin C) 0.00013067562121328783
GO:0045893: positive regulation of transcription, DNA-templated IL4 interleukin 4 0.00025950438
234369504
GO:0045893: positive regulation of transcription, DNA-templated INHBA inhibin, beta A -0.0013515850
421893514
GO:0045893: positive regulation of transcription, DNA-templated INSR insulin receptor -0.00
13621044597654384
GO:0045893: positive regulation of transcription, DNA-templated IRAK1 interleukin-1 receptor-associ
ated kinase 1 -0.0016074433700917167
GO:0045893: positive regulation of transcription, DNA-templated IRF6 interferon regulatory factor
6 -0.0016923647209914838
GO:0045893: positive regulation of transcription, DNA-templated IRF7 interferon regulatory factor
7 -0.0013057533261672223
GO:0045893: positive regulation of transcription, DNA-templated KLF2 Kruppel-like factor 2 -0.00
1231290713583946
GO:0045893: positive regulation of transcription, DNA-templated KMT2A lysine (K)-specific methyltra
nsferase 2A 0.0006600827015158213
GO:0045893: positive regulation of transcription, DNA-templated LEF1 lymphoid enhancer-binding fac
tor 1 -0.00010027652822402436
GO:0045893: positive regulation of transcription, DNA-templated LGR4 leucine-rich repeat containin
g G protein-coupled receptor 4 0.00030823856512951985
GO:0045893: positive regulation of transcription, DNA-templated LHX1 LIM homeobox 1 -0.0007581026
733613869
GO:0045893: positive regulation of transcription, DNA-templated LRP5 low density lipoprotein recep
tor-related protein 5 3.030856743159438e-5
GO:0045893: positive regulation of transcription, DNA-templated LRP6 low density lipoprotein recep
tor-related protein 6 0.0001493542632949018
GO:0045893: positive regulation of transcription, DNA-templated MDK midkine (neurite growth-promo
ting factor 2) 0.0016634429200248321
GO:0045893: positive regulation of transcription, DNA-templated MED1 mediator complex subunit 1
0.0011281820256609692
GO:0045893: positive regulation of transcription, DNA-templated MEF2C myocyte enhancer factor 2C
0.0009725002200719217
GO:0045893: positive regulation of transcription, DNA-templated MYC v-myc avian myelocytomatosis
viral oncogene homolog -0.0011357788935995243
GO:0045893: positive regulation of transcription, DNA-templated NOTCH1 notch 1 0.0005183413877689858
GO:0045893: positive regulation of transcription, DNA-templated NPAT nuclear protein, ataxia-telan
giectasia locus -0.0007091645380148388
GO:0045893: positive regulation of transcription, DNA-templated NR1H3 nuclear receptor subfamily 1,
group H, member 3 0.000800427455734805
GO:0045893: positive regulation of transcription, DNA-templated NRIP1 nuclear receptor interacting
protein 1 0.001070494863550287
GO:0045893: positive regulation of transcription, DNA-templated PAX2 paired box 2 -0.0016127127
247679831
GO:0045893: positive regulation of transcription, DNA-templated PAX3 paired box 3 -0.0029431297
390478755
GO:0045893: positive regulation of transcription, DNA-templated PAX6 paired box 6 0.00193418783
61513528
GO:0045893: positive regulation of transcription, DNA-templated PAX8 paired box 8 0.00094766014
32854529
GO:0045893: positive regulation of transcription, DNA-templated PLCB1 phospholipase C, beta 1 (phos
phoinositide-specific) 0.00015354664083042096
GO:0045893: positive regulation of transcription, DNA-templated POU1F1 POU class 1 homeobox 1 0.000
13727846243815676
GO:0045893: positive regulation of transcription, DNA-templated PROX1 prospero homeobox 1 0.001
1205378911210572
GO:0045893: positive regulation of transcription, DNA-templated PSRC1 proline/serine-rich coiled-co
il 1 0.0006466591884429119
GO:0045893: positive regulation of transcription, DNA-templated PTCH1 patched 1 -6.6001761657
07603e-5
GO:0045893: positive regulation of transcription, DNA-templated RB1 retinoblastoma 1 -0.00
1492829789323435
GO:0045893: positive regulation of transcription, DNA-templated RET ret proto-oncogene -0.00
04908191583948981
GO:0045893: positive regulation of transcription, DNA-templated RNF187 ring finger protein 187 -0.00

22244566023502032
GO:0045893: positive regulation of transcription, DNA-templated RREB1 ras responsive element bindin
g protein 1 -0.0019804178693297887
GO:0045893: positive regulation of transcription, DNA-templated SALL1 spalt-like transcription fact
or 1 -0.0024350576906513027
GO:0045893: positive regulation of transcription, DNA-templated SFRP1 secreted frizzled-related pro
tein 1 0.0012801395176055667
GO:0045893: positive regulation of transcription, DNA-templated SHH sonic hedgehog 0.00059932514
29798954
GO:0045893: positive regulation of transcription, DNA-templated SIX1 SIX homeobox 1 -0.0018685348
88584419
GO:0045893: positive regulation of transcription, DNA-templated SMARCD3 SWI/SNF related, matrix assoc
iated, actin dependent regulator of chromatin, subfamily d, member 3 0.0003452148327330156
GO:0045893: positive regulation of transcription, DNA-templated SOX11 SRY (sex determining region
Y)-box 11 -0.00020913766128075338
GO:0045893: positive regulation of transcription, DNA-templated SOX18 SRY (sex determining region
Y)-box 18 0.0012876607654467276
GO:0045893: positive regulation of transcription, DNA-templated SOX4 SRY (sex determining region
Y)-box 4 -3.847273016829249e-5
GO:0045893: positive regulation of transcription, DNA-templated SOX9 SRY (sex determining region
Y)-box 9 -0.0005182236558763609
GO:0045893: positive regulation of transcription, DNA-templated TBX21 T-box 21 0.00039829661
307276207
GO:0045893: positive regulation of transcription, DNA-templated TBX3 T-box 3 0.0012256285385466956
GO:0045893: positive regulation of transcription, DNA-templated TFAP2A transcription factor AP-2 alp
ha (activating enhancer binding protein 2 alpha) 0.00047961282795407875
GO:0045893: positive regulation of transcription, DNA-templated TGFB1 transforming growth factor, b
eta 1 -7.288968852722864e-5
GO:0045893: positive regulation of transcription, DNA-templated TGFB3 transforming growth factor, b
eta 3 -0.0018220658739433056
GO:0045893: positive regulation of transcription, DNA-templated TGFBR1 transforming growth factor, b
eta receptor 1 0.0003395534066790808
GO:0045893: positive regulation of transcription, DNA-templated TP53 tumor protein p53 0.001
174553693524912
GO:0045893: positive regulation of transcription, DNA-templated TP73 tumor protein p73 0.001
0271237257381716
GO:0045893: positive regulation of transcription, DNA-templated TRIM16 tripartite motif containing 1
6 -0.0014876241924654263
GO:0045893: positive regulation of transcription, DNA-templated WNT1 wingless-type MMTV integratio
n site family, member 1 0.0007863796247952719
GO:0045893: positive regulation of transcription, DNA-templated WNT4 wingless-type MMTV integratio
n site family, member 4 -0.00025161322209507034
GO:0045893: positive regulation of transcription, DNA-templated WNT5A wingless-type MMTV integratio
n site family, member 5A -0.0006668628482081897
GO:0045893: positive regulation of transcription, DNA-templated WNT7A wingless-type MMTV integratio
n site family, member 7A 1.9536767214064923e-5
GO:0045893: positive regulation of transcription, DNA-templated WT1 Wilms tumor 1 -0.0005064029
700403651
GO:0045944: positive regulation of transcription from RNA polymerase II promoter ACVRL1 activ
in A receptor type II-like 1 0.0019443040642050957
GO:0045944: positive regulation of transcription from RNA polymerase II promoter ADIRF adipo
genesis regulatory factor -0.0001664788414672762
GO:0045944: positive regulation of transcription from RNA polymerase II promoter AKT1 v-akt
murine thymoma viral oncogene homolog 1 0.0007307987633961781
GO:0045944: positive regulation of transcription from RNA polymerase II promoter AKT2 v-akt
murine thymoma viral oncogene homolog 2 -0.0006260107247967155
GO:0045944: positive regulation of transcription from RNA polymerase II promoter ALX1 ALX h
omeobox 1 0.0022925475500006395
GO:0045944: positive regulation of transcription from RNA polymerase II promoter AR andro
gen receptor 0.002638822738909427
GO:0045944: positive regulation of transcription from RNA polymerase II promoter ASCL1 achae
te-scute family bHLH transcription factor 1 -0.0014328599434172357
GO:0045944: positive regulation of transcription from RNA polymerase II promoter ATAD2 ATPas
e family, AAA domain containing 2 -4.7625349177237565e-5
GO:0045944: positive regulation of transcription from RNA polymerase II promoter ATAD2B ATPas
e family, AAA domain containing 2B -0.0006257229810351446
GO:0045944: positive regulation of transcription from RNA polymerase II promoter ATF5 activ
ating transcription factor 5 -0.0027356679899733227
GO:0045944: positive regulation of transcription from RNA polymerase II promoter BACH1 BTB a
nd CNC homology 1, basic leucine zipper transcription factor 1 0.0004854033082600652
GO:0045944: positive regulation of transcription from RNA polymerase II promoter BATF basic
leucine zipper transcription factor, ATF-like 0.00203955795620932
GO:0045944: positive regulation of transcription from RNA polymerase II promoter BCL11A B-cel
l CLL/lymphoma 11A (zinc finger protein) -0.000977247548991509
GO:0045944: positive regulation of transcription from RNA polymerase II promoter BMP4 bone
morphogenetic protein 4 -0.0003226663399600548З
GO:0045944: positive regulation of transcription from RNA polymerase II promoter BMP7 bone
morphogenetic protein 7 0.0008558708714268643
GO:0045944: positive regulation of transcription from RNA polymerase II promoter BRD4 bromo
domain containing 4 0.0005538715296433381
GO:0045944: positive regulation of transcription from RNA polymerase II promoter CCNC cycli

n C	0.0005675530115576567
GO:0045944: positive regulation of transcription from RNA polymerase II promoter	CDK5RAP3	CDK5 regulatory subunit associated protein 3	0.0011383878037459396
GO:0045944: positive regulation of transcription from RNA polymerase II promoter	CDK8	cyclin-dependent kinase 8	0.0026132659090249357
GO:0045944: positive regulation of transcription from RNA polymerase II promoter	CDKN2A	cyclin-dependent kinase inhibitor 2A	0.0017585356679328025
GO:0045944: positive regulation of transcription from RNA polymerase II promoter	CDKN2B	cyclin-dependent kinase inhibitor 2B (p15, inhibits CDK4)	-0.001835902323059661
GO:0045944: positive regulation of transcription from RNA polymerase II promoter	CEBPB	CCAAT/enhancer binding protein (C/EBP), beta	-0.0002740273814654646
GO:0045944: positive regulation of transcription from RNA polymerase II promoter	CKAP2	cytoskeleton associated protein 2	4.914027723800857e-5
GO:0045944: positive regulation of transcription from RNA polymerase II promoter	CLOCK	clock circadian regulator	0.00020012973136357572
GO:0045944: positive regulation of transcription from RNA polymerase II promoter	CREB1	cAMP responsive element binding protein 1	0.0006584918070843077
GO:0045944: positive regulation of transcription from RNA polymerase II promoter	CTNNB1	catenin (cadherin-associated protein), beta 1, 88kDa	-0.0001169835294219796
GO:0045944: positive regulation of transcription from RNA polymerase II promoter	CXCL10	chemokine (C-X-C motif) ligand 10	6.257547612097429e-5
GO:0045944: positive regulation of transcription from RNA polymerase II promoter	DDX17	DEAD (Asp-Glu-Ala-Asp) box helicase 17	0.001762259909627512
GO:0045944: positive regulation of transcription from RNA polymerase II promoter	DMRT1	doublesex and mab-3 related transcription factor 1	0.0015048980570308553
GO:0045944: positive regulation of transcription from RNA polymerase II promoter	DRD2	dopamine receptor D2	-0.0002340815848517177
GO:0045944: positive regulation of transcription from RNA polymerase II promoter	E2F1	E2F transcription factor 1	0.002098146047396107
GO:0045944: positive regulation of transcription from RNA polymerase II promoter	E2F4	E2F transcription factor 4, p107/p130-binding	-0.0023461456649955417
GO:0045944: positive regulation of transcription from RNA polymerase II promoter	E2F8	E2F transcription factor 8	0.0017113278291275697
GO:0045944: positive regulation of transcription from RNA polymerase II promoter	EGFR	epidermal growth factor receptor	0.0006859607328492662
GO:0045944: positive regulation of transcription from RNA polymerase II promoter	EGR1	early growth response 1	0.0010933264118176435
GO:0045944: positive regulation of transcription from RNA polymerase II promoter	EGR2	early growth response 2	0.0014391647228194574
GO:0045944: positive regulation of transcription from RNA polymerase II promoter	ELF5	E74-like factor 5 (ets domain transcription factor)	0.0011713153360883152
GO:0045944: positive regulation of transcription from RNA polymerase II promoter	EN1	engrailed homeobox 1	-1.5794318398248944e-5
GO:0045944: positive regulation of transcription from RNA polymerase II promoter	ENG	endoglin	0.0008267683638286395
GO:0045944: positive regulation of transcription from RNA polymerase II promoter	ESR1	estrogen receptor 1	-0.0009483419545981599
GO:0045944: positive regulation of transcription from RNA polymerase II promoter	ETS2	v-ets avian erythroblastosis virus E26 oncogene homolog 2	-0.0010158235714558567
GO:0045944: positive regulation of transcription from RNA polymerase II promoter	ETV4	ets variant 4	0.0001949228002014306
GO:0045944: positive regulation of transcription from RNA polymerase II promoter	EYA1	EYA transcriptional coactivator and phosphatase 1	5.3112802003076535e-5
GO:0045944: positive regulation of transcription from RNA polymerase II promoter	FGFR2	fibroblast growth factor receptor 2	0.0007634000106427033
GO:0045944: positive regulation of transcription from RNA polymerase II promoter	FOXA1	forkhead box A1	2.774794035939197e-5
GO:0045944: positive regulation of transcription from RNA polymerase II promoter	FOXA2	forkhead box A2	-1.6667603280648776e-5
GO:0045944: positive regulation of transcription from RNA polymerase II promoter	FOXC1	forkhead box C1	-2.223171321108798e-5
GO:0045944: positive regulation of transcription from RNA polymerase II promoter	FOXC2	forkhead box C2 (MFH-1, mesenchyme forkhead 1)	0.0017485811525600034
GO:0045944: positive regulation of transcription from RNA polymerase II promoter	FOXH1	forkhead box H1	-0.0013714971674223674
GO:0045944: positive regulation of transcription from RNA polymerase II promoter	FOXM1	forkhead box M1	0.0001972126308271496
GO:0045944: positive regulation of transcription from RNA polymerase II promoter	FOXO1	forkhead box O1	0.0018006087849650712
GO:0045944: positive regulation of transcription from RNA polymerase II promoter	FOXO3	forkhead box O3	0.001200338976378505
GO:0045944: positive regulation of transcription from RNA polymerase II promoter	GATA2	GATA binding protein 2	-0.00045165603093946826
GO:0045944: positive regulation of transcription from RNA polymerase II promoter	GATA3	GATA binding protein 3	-3.900704642718199e-5
GO:0045944: positive regulation of transcription from RNA polymerase II promoter	GATA4	GATA binding protein 4	-0.0010950113941441482
GO:0045944: positive regulation of transcription from RNA polymerase II promoter	GATA6	GATA binding protein 6	-2.817318758581481e-5
GO:0045944: positive regulation of transcription from RNA polymerase II promoter	GDNF	glial cell derived neurotrophic factor	0.00044485219823784327

GO:0045944: positive regulation of transcription from RNA polymerase II promoter GLI1 GLI family zinc finger 1 -0.0012970825571942209
GO:0045944: positive regulation of transcription from RNA polymerase II promoter GLI2 GLI family zinc finger 2 0.0018539204116941608
GO:0045944: positive regulation of transcription from RNA polymerase II promoter GLI3 GLI family zinc finger 3 -0.002152105633066085
GO:0045944: positive regulation of transcription from RNA polymerase II promoter GREM1 gremlin 1, DAN family BMP antagonist -0.0008275668341616591
GO:0045944: positive regulation of transcription from RNA polymerase II promoter GRHL2 grainyhead-like 2 (Drosophila) 0.001006963233202589
GO:0045944: positive regulation of transcription from RNA polymerase II promoter GSK3B glycogen synthase kinase 3 beta 0.0015494892993057788
GO:0045944: positive regulation of transcription from RNA polymerase II promoter HDAC2 histone deacetylase 2 -0.0012074866134420418
GO:0045944: positive regulation of transcription from RNA polymerase II promoter HES1 hes family bHLH transcription factor 1 -0.0009111680624738231
GO:0045944: positive regulation of transcription from RNA polymerase II promoter HEY2 hes-related family bHLH transcription factor with YRPW motif 2 0.0026995028122499577
GO:0045944: positive regulation of transcription from RNA polymerase II promoter HEYL hes-related family bHLH transcription factor with YRPW motif-like 0.0006777681632462492
GO:0045944: positive regulation of transcription from RNA polymerase II promoter HIF1A hypoxia inducible factor 1, alpha subunit (basic helix-loop-helix transcription factor) -0.0006466180484311898
GO:0045944: positive regulation of transcription from RNA polymerase II promoter HIPK2 homeodomain interacting protein kinase 2 0.0007709706169694359
GO:0045944: positive regulation of transcription from RNA polymerase II promoter HMGA1 high mobility group AT-hook 1 -0.00031875032539067813
GO:0045944: positive regulation of transcription from RNA polymerase II promoter HMGA2 high mobility group AT-hook 2 0.0015092633026494988
GO:0045944: positive regulation of transcription from RNA polymerase II promoter HMGB1 high mobility group box 1 -0.0007765246540718914
GO:0045944: positive regulation of transcription from RNA polymerase II promoter HMGB2 high mobility group box 2 0.00030719307885096166
GO:0045944: positive regulation of transcription from RNA polymerase II promoter HOXA10 homeobox A10 -0.0029825409453013807
GO:0045944: positive regulation of transcription from RNA polymerase II promoter HOXA5 homeobox A5 0.001066454837108327
GO:0045944: positive regulation of transcription from RNA polymerase II promoter HOXB1 homeobox B1 0.003687280807666003
GO:0045944: positive regulation of transcription from RNA polymerase II promoter HOXD13 homeobox D13 -0.0005925971480680946
GO:0045944: positive regulation of transcription from RNA polymerase II promoter IFNG interferon, gamma -4.8911649086303884e-5
GO:0045944: positive regulation of transcription from RNA polymerase II promoter IGBP1 immunoglobulin (CD79A) binding protein 1 0.002849935930368596
GO:0045944: positive regulation of transcription from RNA polymerase II promoter IGF1 insulin-like growth factor 1 (somatomedin C) 0.00013093438469579307
GO:0045944: positive regulation of transcription from RNA polymerase II promoter IHH indian hedgehog -0.002041046971787435
GO:0045944: positive regulation of transcription from RNA polymerase II promoter IL4 interleukin 4 0.0002595770430440888
GO:0045944: positive regulation of transcription from RNA polymerase II promoter INHBA inhibin, beta A -0.0013533643395976804
GO:0045944: positive regulation of transcription from RNA polymerase II promoter IRF7 interferon regulatory factor 7 -0.001306922319073659
GO:0045944: positive regulation of transcription from RNA polymerase II promoter ISL1 ISL LIM homeobox 1 7.899059658237332e-5
GO:0045944: positive regulation of transcription from RNA polymerase II promoter ITGA6 integrin, alpha 6 0.001669402506389009
GO:0045944: positive regulation of transcription from RNA polymerase II promoter JAG1 jagged 1 0.0017569637542202388
GO:0045944: positive regulation of transcription from RNA polymerase II promoter JAK2 Janus kinase 2 -3.1764008224896414e-5
GO:0045944: positive regulation of transcription from RNA polymerase II promoter KLF15 Kruppel-like factor 15 0.00046526701612469186
GO:0045944: positive regulation of transcription from RNA polymerase II promoter KLF2 Kruppel-like factor 2 -0.0012323207467792867
GO:0045944: positive regulation of transcription from RNA polymerase II promoter KMT2A lysine (K)-specific methyltransferase 2A 0.0006607590505708988
GO:0045944: positive regulation of transcription from RNA polymerase II promoter KMT2D lysine (K)-specific methyltransferase 2D -0.00019942560142910442
GO:0045944: positive regulation of transcription from RNA polymerase II promoter LEF1 lymphoid enhancer-binding factor 1 -0.00010029885240289659
GO:0045944: positive regulation of transcription from RNA polymerase II promoter LMO4 LIM domain only 4 0.0021118349984596855
GO:0045944: positive regulation of transcription from RNA polymerase II promoter LRP5 low density lipoprotein receptor-related protein 5 3.0262688070737596e-5
GO:0045944: positive regulation of transcription from RNA polymerase II promoter LRP6 low density lipoprotein receptor-related protein 6 0.00014955188373028675
GO:0045944: positive regulation of transcription from RNA polymerase II promoter MED1 mediator complex subunit 1 0.001129496435543127

GO:0045944: positive regulation of transcription from RNA polymerase II promoter MEF2C myocyte enhancer factor 2C 0.0009735755401232784
GO:0045944: positive regulation of transcription from RNA polymerase II promoter MLLT10 myeloid/lymphoid or mixed-lineage leukemia (trithorax homolog, Drosophila); translocated to, 10 -0.0001632855831282176З
GO:0045944: positive regulation of transcription from RNA polymerase II promoter MSX1 msh homeobox 1 -0.0027696324611834644
GO:0045944: positive regulation of transcription from RNA polymerase II promoter MYC v-myc avian myelocytomatosis viral oncogene homolog -0.0011367180778710075
GO:0045944: positive regulation of transcription from RNA polymerase II promoter NCK1 NCK adaptor protein 1 -0.0007844735590932211
GO:0045944: positive regulation of transcription from RNA polymerase II promoter NCK2 NCK adaptor protein 2 -0.0002321168743430567 6
GO:0045944: positive regulation of transcription from RNA polymerase II promoter NFIB nuclear factor I/B 0.002919954450386313
GO:0045944: positive regulation of transcription from RNA polymerase II promoter NOTCH1 notch 1 0.0005188996920113451
GO:0045944: positive regulation of transcription from RNA polymerase II promoter NR1H3 nuclear receptor subfamily 1, group H, member 3 0.0008010554039037999
GO:0045944: positive regulation of transcription from RNA polymerase II promoter NR2E1 nuclear receptor subfamily 2, group E, member 1 -0.0029756095966291317
GO:0045944: positive regulation of transcription from RNA polymerase II promoter NRIP1 nuclear receptor interacting protein 1 0.001071349028194392
GO:0045944: positive regulation of transcription from RNA polymerase II promoter PAX2 paired box 2 -0.0016144289836253674
GO:0045944: positive regulation of transcription from RNA polymerase II promoter PAX3 paired box 3 -0.0029458861572175302
GO:0045944: positive regulation of transcription from RNA polymerase II promoter PAX6 paired box 6 0.0019361357343503501
GO:0045944: positive regulation of transcription from RNA polymerase II promoter PAX8 paired box 8 0.0009483815195790173
GO:0045944: positive regulation of transcription from RNA polymerase II promoter PDX1 pancreatic and duodenal homeobox 1 0.00025733616349433606
GO:0045944: positive regulation of transcription from RNA polymerase II promoter PGR progesterone receptor 0.0003519403727872666
GO:0045944: positive regulation of transcription from RNA polymerase II promoter PHOX2B paired-like homeobox 2b 0.0003905472967702659
GO:0045944: positive regulation of transcription from RNA polymerase II promoter PITX2 paired-like homeodomain 2 0.0021765648777488655
GO:0045944: positive regulation of transcription from RNA polymerase II promoter POU1F1 POU class 1 homeobox 1 0.00013744987738267359
GO:0045944: positive regulation of transcription from RNA polymerase II promoter PRKDC protein kinase, DNA-activated, catalytic polypeptide -0.001856019476409261
GO:0045944: positive regulation of transcription from RNA polymerase II promoter PROX1 prospero homeobox 1 0.0011217509861606585
GO:0045944: positive regulation of transcription from RNA polymerase II promoter RAD21 RAD21 homolog (S. pombe) -0.00010233687057396984
GO:0045944: positive regulation of transcription from RNA polymerase II promoter RAF1 Raf-1 proto-oncogene, serine/threonine kinase 0.001499129217626
GO:0045944: positive regulation of transcription from RNA polymerase II promoter RARG retinoic acid receptor, gamma -0.002319992912529788
GO:0045944: positive regulation of transcription from RNA polymerase II promoter RB1 retinoblastoma 1 -0.0014943260016971119
GO:0045944: positive regulation of transcription from RNA polymerase II promoter RBPJ recombination signal binding protein for immunoglobulin kappa J region 0.0009816834092332136
GO:0045944: positive regulation of transcription from RNA polymerase II promoter RPS6KA1 ribosomal protein S6 kinase, 90kDa, polypeptide 1 -0.0025147744791885517
GO:0045944: positive regulation of transcription from RNA polymerase II promoter RXRA retinoid X receptor, alpha 0.0011153419614766482
GO:0045944: positive regulation of transcription from RNA polymerase II promoter SALL1 spalt-like transcription factor 1 -0.0024379079972816297
GO:0045944: positive regulation of transcription from RNA polymerase II promoter SERPINE1 serpin peptidase inhibitor, clade E (nexin, plasminogen activator inhibitor type 1), member 1 0.000112352903574655
GO:0045944: positive regulation of transcription from RNA polymerase II promoter SERPINF2 serpin peptidase inhibitor, clade F (alpha-2 antiplasmin, pigment epithelium derived factor), member 2 0.0006588716807296493
GO:0045944: positive regulation of transcription from RNA polymerase II promoter SHH sonic hedgehog 0.0006000688860940622
GO:0045944: positive regulation of transcription from RNA polymerase II promoter SIRT1 sirtuin 1 -1.6533830443268866e-6
GO:0045944: positive regulation of transcription from RNA polymerase II promoter SIRT2 sirtuin 2 -0.0008524578540420826
GO:0045944: positive regulation of transcription from RNA polymerase II promoter SIX1 SIX homeobox 1 -0.0018703894772010047
GO:0045944: positive regulation of transcription from RNA polymerase II promoter SIX3 SIX homeobox 3 0.0020377068984229955
GO:0045944: positive regulation of transcription from RNA polymerase II promoter SKI SKI proto-oncogene -0.0006706217304763651
GO:0045944: positive regulation of transcription from RNA polymerase II promoter SMO smoothened, frizzled class receptor 0.002139710458429633

GO:0045944: positive regulation of transcription from RNA polymerase II promoter SOX10 SRY
(sex determining region Y)-box 10 0.00019050168791481697
GO:0045944: positive regulation of transcription from RNA polymerase II promoter SOX11 SRY
(sex determining region Y)-box 11 -0.00020912628944135043
GO:0045944: positive regulation of transcription from RNA polymerase II promoter SOX18 SRY
(sex determining region Y)-box 18 0.0012887650551480564
GO:0045944: positive regulation of transcription from RNA polymerase II promoter SOX4 SRY
(sex determining region Y)-box 4 -3.8302354977654686e-5
GO:0045944: positive regulation of transcription from RNA polymerase II promoter SOX9 SRY
(sex determining region Y)-box 9 -0.0005187298703103373
GO:0045944: positive regulation of transcription from RNA polymerase II promoter SPDEF SAM p
ointed domain containing ETS transcription factor 0.0031376050999923424
GO:0045944: positive regulation of transcription from RNA polymerase II promoter STAT5A signa
l transducer and activator of transcription 5A 0.0015966587800545346
GO:0045944: positive regulation of transcription from RNA polymerase II promoter TCF7L2 trans
cription factor 7-like 2 (T-cell specific, HMG-box) 0.0005738109595772474
GO:0045944: positive regulation of transcription from RNA polymerase II promoter TFAP2A trans
cription factor AP-2 alpha (activating enhancer binding protein 2 alpha) 0.0004799991172967250
6
GO:0045944: positive regulation of transcription from RNA polymerase II promoter TFAP2C trans
cription factor AP-2 gamma (activating enhancer binding protein 2 gamma) 0.0010603330294379238
GO:0045944: positive regulation of transcription from RNA polymerase II promoter TFDP1 trans
cription factor Dp-1 0.0014450484802161858
GO:0045944: positive regulation of transcription from RNA polymerase II promoter TGFB1 trans
forming growth factor, beta 1 -7.302009650972842e-5
GO:0045944: positive regulation of transcription from RNA polymerase II promoter TGFB3 trans
forming growth factor, beta 3 -0.00182411692809807
GO:0045944: positive regulation of transcription from RNA polymerase II promoter THRA thyro
id hormone receptor, alpha 0.0007633944560357848
GO:0045944: positive regulation of transcription from RNA polymerase II promoter THRB thyro
id hormone receptor, beta 0.0019444257709734128
GO:0045944: positive regulation of transcription from RNA polymerase II promoter TOP2A topoi
somerase (DNA) II alpha 170kDa -8.25118984213841e-5
GO:0045944: positive regulation of transcription from RNA polymerase II promoter TP53 tumor
protein p53 0.0011759598638691273
GO:0045944: positive regulation of transcription from RNA polymerase II promoter TP73 tumor
protein p73 0.0010281381911382771
GO:0045944: positive regulation of transcription from RNA polymerase II promoter TRPS1 trich
orhinophalangeal syndrome I -0.0001777800864428266
GO:0045944: positive regulation of transcription from RNA polymerase II promoter TWIST1 twist
family bHLH transcription factor 1 -0.001341832189189522
GO:0045944: positive regulation of transcription from RNA polymerase II promoter VDR vitam
in D (1,25- dihydroxyvitamin D3) receptor 0.0005636709231557238
GO:0045944: positive regulation of transcription from RNA polymerase II promoter VEGFA vascu
lar endothelial growth factor A 0.0005951064225557961
GO:0045944: positive regulation of transcription from RNA polymerase II promoter WNT1 wingl
ess-type MMTV integration site family, member 1 0.0007871140874052553
GO:0045944: positive regulation of transcription from RNA polymerase II promoter WNT5A wingl
ess-type MMTV integration site family, member 5A -0.0006675786331229681
GO:0045944: positive regulation of transcription from RNA polymerase II promoter WNT7A wingl
ess-type MMTV integration site family, member 7A 1.9509947385214043e-5
GO:0045944: positive regulation of transcription from RNA polymerase II promoter WT1 Wilms
tumor 1 -0.0005070002401581366
GO:0045944: positive regulation of transcription from RNA polymerase II promoter WWP2 WW do
main containing E3 ubiquitin protein ligase 2 -0.0009573922296456003
GO:0045944: positive regulation of transcription from RNA polymerase II promoter WWTR1 WW do
main containing transcription regulator 1 0.0009006582974867813
GO:0045944: positive regulation of transcription from RNA polymerase II promoter XBP1 X-box
binding protein 1 0.0002673236707801496
GO:0045944: positive regulation of transcription from RNA polymerase II promoter YAP1 Yes-a
ssociated protein 1 -0.0001643789816582154
GO:0045944: positive regulation of transcription from RNA polymerase II promoter YBX1 Y box
binding protein 1 -0.0009140689035934039
GO:0045944: positive regulation of transcription from RNA polymerase II promoter ZNF148 zinc
finger protein 148 0.002104489054872158
GO:0051895: negative regulation of focal adhesion assembly ACVRL1 activin A receptor type II-li
ke 1 0.0019530649656913576
GO:0051895: negative regulation of focal adhesion assembly APOD apolipoprotein D 0.002
6421423650420162
GO:0051895: negative regulation of focal adhesion assembly PTEN phosphatase and tensin homolo
g 1.6114585803782153e-5
GO:0051895: negative regulation of focal adhesion assembly THBS1 thrombospondin 1 -0.00
10406856223369905
GO:0060836: lymphatic endothelial cell differentiation ACVRL1 activin A receptor type II-like 1
0.001944657952279954 8
GO:0060836: lymphatic endothelial cell differentiation PROX1 prospero homeobox 1 0.00112179365
13517406
GO:0060836: lymphatic endothelial cell differentiation SOX18 SRY (sex determining region Y)-box 18
0.001289813463985067
GO:0060840: artery development ACVRL1 activin A receptor type II-like 1 0.0019561802069634356
GO:0060840: artery development GLI3 GLI family zinc finger 3 -0.002168766150195870 4

GO:0060840: artery development  SHH     sonic hedgehog  0.0006054598683210002
GO:0060841: venous blood vessel development     ACVRL1  activin A receptor type II-like 1       0.001
9557258365855866
GO:0061154: endothelial tube morphogenesis      ACVRL1  activin A receptor type II-like 1       0.001
9468136600092449
GO:0061154: endothelial tube morphogenesis      CSNK2B  casein kinase 2, beta polypeptide       0.001
5314226415575785
GO:0061154: endothelial tube morphogenesis      CTNNB1  catenin (cadherin-associated protein), beta
1, 88kDa       -0.00011698370345540074
GO:0061298: retina vasculature development in camera-type eye   ACVRL1  activin A receptor type II-li
ke 1    0.0019461897695409877
GO:0061298: retina vasculature development in camera-type eye   HIF1A   hypoxia inducible factor 1, a
lpha subunit (basic helix-loop-helix transcription factor)      -0.0006476998418521947
GO:0061298: retina vasculature development in camera-type eye   PDGFRB  platelet-derived growth facto
r receptor, beta polypeptide    -0.0003609549333895543
GO:0061298: retina vasculature development in camera-type eye   ROM1    retinal outer segment membran
e protein 1     0.002463912437196278
GO:0071560: cellular response to transforming growth factor beta stimulus       ACVRL1  activin A rec
eptor type II-like 1    0.001926261397488942
GO:0071560: cellular response to transforming growth factor beta stimulus       CAV1    caveolin 1, c
aveolae protein, 22kDa  -0.0005316917174526753
GO:0071560: cellular response to transforming growth factor beta stimulus       COL1A1  collagen, typ
e I, alpha 1    -0.0005185334556705554
GO:0071560: cellular response to transforming growth factor beta stimulus       CX3CR1  chemokine (C-
X3-C motif) receptor 1  -0.0008498669779676426
GO:0071560: cellular response to transforming growth factor beta stimulus       HDAC2   histone deace
tylase 2        -0.0011978096555682778
GO:0071560: cellular response to transforming growth factor beta stimulus       MEF2C   myocyte enhan
cer factor 2C   0.0009628080732897106
GO:0071560: cellular response to transforming growth factor beta stimulus       SFRP1   secreted friz
zled-related protein 1  0.0012641309059244078
GO:0071560: cellular response to transforming growth factor beta stimulus       SOX9    SRY (sex dete
rmining region Y)-box 9 -0.0005131145359244434
GO:0071560: cellular response to transforming growth factor beta stimulus       STAR    steroidogenic
acute regulatory protein        0.0008850365768218511
GO:0071560: cellular response to transforming growth factor beta stimulus       TGFB1   transforming
growth factor, beta 1  -7.196941732189385e-5
GO:0071560: cellular response to transforming growth factor beta stimulus       TGFBR1  transforming
growth factor, beta receptor 1 0.0003358247940326843
GO:0071560: cellular response to transforming growth factor beta stimulus       WNT4    wingless-type
MMTV integration site family, member 4  -0.00024749006372700263
GO:0071560: cellular response to transforming growth factor beta stimulus       WNT5A   wingless-type
MMTV integration site family, member 5A -0.0006607161693427856
GO:0071560: cellular response to transforming growth factor beta stimulus       WNT7A   wingless-type
MMTV integration site family, member 7A 2.0016579181722838e-5
GO:0071773: cellular response to BMP stimulus   ACVRL1  activin A receptor type II-like 1       0.001
9469161414038603
GO:0071773: cellular response to BMP stimulus   BMP4    bone morphogenetic protein 4    -0.0003231866
0085560853
GO:0071773: cellular response to BMP stimulus   BMP7    bone morphogenetic protein 7    0.00085672490
5735478
GO:0071773: cellular response to BMP stimulus   DLX5    distal-less homeobox 5  -0.003287992159580729
3
GO:0071773: cellular response to BMP stimulus   GATA3   GATA binding protein 3  -3.99441458484947e-5
GO:0071773: cellular response to BMP stimulus   GATA6   GATA binding protein 6  -2.773494593162437e-5
GO:0071773: cellular response to BMP stimulus   HEYL    hes-related family bHLH transcription factor
with YRPW motif-like   0.0006787436791520908
GO:0071773: cellular response to BMP stimulus   PHOX2B  paired-like homeobox 2b 0.0003906186163447779
GO:0071773: cellular response to BMP stimulus   SFRP1   secreted frizzled-related protein 1     0.001
285631768528616
GO:2000279: negative regulation of DNA biosynthetic process     ACVRL1  activin A receptor type II-li
ke 1    0.0019437890735744716
GO:2000279: negative regulation of DNA biosynthetic process     DACH1   dachshund family transcriptio
n factor 1      0.0020895788088585035
GO:2000279: negative regulation of DNA biosynthetic process     DNAJC2  DnaJ (Hsp40) homolog, subfami
ly C, member 2  -0.0009839168760175266
GO:2000279: negative regulation of DNA biosynthetic process     GJA1    gap junction protein, alpha
1, 43kDa       -0.00016054343281828392
GO:0006508: proteolysis ADAM2   ADAM metallopeptidase domain 2  0.0008093986193826227
GO:0006508: proteolysis AZU1    azurocidin 1    0.0001974591915609093
GO:0006508: proteolysis C2      complement component 2  -0.0003032388728206672
GO:0006508: proteolysis CHMP1A  charged multivesicular body protein 1A  -0.0007779082330697745
GO:0006508: proteolysis CPA4    carboxypeptidase A4     -0.0006964846600073002
GO:0006508: proteolysis CTSK    cathepsin K     -0.0003612139197387843
GO:0006508: proteolysis CUL7    cullin 7        -0.00011845549940195489
GO:0006508: proteolysis ESPL1   extra spindle pole bodies homolog 1 (S. cerevisiae)     0.00019570583
77817672
GO:0006508: proteolysis FAP     fibroblast activation protein, alpha    -0.00048527449866830755
GO:0006508: proteolysis GGH     gamma-glutamyl hydrolase (conjugase, folylpolygammaglutamyl hydrolas
e)      -0.00031457450643210375
GO:0006508: proteolysis HPN     hepsin  0.003151719539580326

GO:0006508: proteolysis KLK5    kallikrein-related peptidase 5  0.0017293365197943115
GO:0006508: proteolysis KLK7    kallikrein-related peptidase 7  0.0016212867507967707
GO:0006508: proteolysis LRP8    low density lipoprotein receptor-related protein 8, apolipoprotein e
 receptor       -0.0009299303650135817
GO:0006508: proteolysis METAP1  methionyl aminopeptidase 1      -0.0010139925529321567
GO:0006508: proteolysis MMP2    matrix metallopeptidase 2 (gelatinase A, 72kDa gelatinase, 72kDa type
IV collagenase) -0.0011276573466487817
GO:0006508: proteolysis MMP20   matrix metallopeptidase 20      -0.0024265001319835316
GO:0006508: proteolysis MMP24   matrix metallopeptidase 24 (membrane-inserted)  0.0014861614325918419
GO:0006508: proteolysis MMP7    matrix metallopeptidase 7 (matrilysin, uterine) 0.0004245508828506052
GO:0006508: proteolysis NAALADL1        N-acetylated alpha-linked acidic dipeptidase-like 1     -0.00
10165593282607669
GO:0006508: proteolysis NDEL1   nudE neurodevelopment protein 1-like 1  -0.0006025021567582332
GO:0006508: proteolysis PCSK2   proprotein convertase subtilisin/kexin type 2   0.0006040018243175648
GO:0006508: proteolysis PGC     progastricsin (pepsinogen C)    0.0013209284413645671
GO:0006508: proteolysis RELN    reelin  0.0015327924562527383
GO:0006508: proteolysis SFRP1   secreted frizzled-related protein 1     0.0012765291137464023
GO:0007155: cell adhesion       ADAM2   ADAM metallopeptidase domain 2  0.0008104134556117934
GO:0007155: cell adhesion       APC     adenomatous polyposis coli      0.0006434239199560318
GO:0007155: cell adhesion       CCL2    chemokine (C-C motif) ligand 2  0.0008159160900474181
GO:0007155: cell adhesion       CD22    CD22 molecule   0.0008367696149558021
GO:0007155: cell adhesion       CD9     CD9 molecule    -0.0025837983727500687
GO:0007155: cell adhesion       CDH2    cadherin 2, type 1, N-cadherin (neuronal)       -0.0006440520
366830424
GO:0007155: cell adhesion       CDH3    cadherin 3, type 1, P-cadherin (placental)      -0.0012296514
424777675
GO:0007155: cell adhesion       CIB1    calcium and integrin binding 1 (calmyrin)       5.48341576874
97897e-5
GO:0007155: cell adhesion       COL5A1  collagen, type V, alpha 1       -4.90033960257594e-5
GO:0007155: cell adhesion       CSF3R   colony stimulating factor 3 receptor (granulocyte)      -8.31
2030252776043e-5
GO:0007155: cell adhesion       CTNNB1  catenin (cadherin-associated protein), beta 1, 88kDa    -0.00
011686020462413972
GO:0007155: cell adhesion       CX3CL1  chemokine (C-X3-C motif) ligand 1       0.0021146751004839093
GO:0007155: cell adhesion       CX3CR1  chemokine (C-X3-C motif) receptor 1     -0.000856317316241553
6
GO:0007155: cell adhesion       CXCL12  chemokine (C-X-C motif) ligand 12       -0.001207260672067927
GO:0007155: cell adhesion       CYP1B1  cytochrome P450, family 1, subfamily B, polypeptide 1   0.000
39745795367300123
GO:0007155: cell adhesion       DSC2    desmocollin 2   0.00046989499160073105
GO:0007155: cell adhesion       DSG2    desmoglein 2    1.680558590932093e-5
GO:0007155: cell adhesion       EFS     embryonal Fyn-associated substrate      0.0039049070512974632
GO:0007155: cell adhesion       ENG     endoglin        0.0008255748753880444
GO:0007155: cell adhesion       FAP     fibroblast activation protein, alpha    -0.000486835305309508
7
GO:0007155: cell adhesion       GRHL2   grainyhead-like 2 (Drosophila)  0.0010059665278473026
GO:0007155: cell adhesion       HES1    hes family bHLH transcription factor 1  -0.000910111658682948
GO:0007155: cell adhesion       ICAM1   intercellular adhesion molecule 1       0.0007288206912410561
GO:0007155: cell adhesion       ITGB4   integrin, beta 4        0.0005795781480714008
GO:0007155: cell adhesion       LAMB2   laminin, beta 2 (laminin S)     -0.0013077434196658699
GO:0007155: cell adhesion       MFGE8   milk fat globule-EGF factor 8 protein   0.0018029424111920078
GO:0007155: cell adhesion       MYH10   myosin, heavy chain 10, non-muscle      -0.000328695499810490
8
GO:0007155: cell adhesion       PRKCA   protein kinase C, alpha -5.862354995876375e-6
GO:0007155: cell adhesion       PRKX    protein kinase, X-linked        0.00039422521787568283
GO:0007155: cell adhesion       PTK7    protein tyrosine kinase 7       -0.00022510246975984075
GO:0007155: cell adhesion       RELN    reelin  0.0015356399567435894
GO:0007155: cell adhesion       ROBO1   roundabout, axon guidance receptor, homolog 1 (Drosophila)
-0.002245569196689939
GO:0007155: cell adhesion       ROM1    retinal outer segment membrane protein 1        0.00245861614
3535573
GO:0007155: cell adhesion       SELPLG  selectin P ligand       0.00028658804116511174
GO:0007155: cell adhesion       SPAM1   sperm adhesion molecule 1 (PH-20 hyaluronidase, zona pellucid
a binding)      -0.0018981817539297056
GO:0007155: cell adhesion       THBS1   thrombospondin 1        -0.0010349187003224824
GO:0007155: cell adhesion       TNC     tenascin C      0.0007335921996569873
GO:0007338: single fertilization        ADAM2   ADAM metallopeptidase domain 2  0.0008170994606222098
GO:0007338: single fertilization        CD9     CD9 molecule    -0.0026086310294060446
GO:0007338: single fertilization        HOXA10  homeobox A10    -0.0030081021179170003
GO:0007338: single fertilization        MFGE8   milk fat globule-EGF factor 8 protein   0.00182231378
48137063
GO:0007338: single fertilization        SPAM1   sperm adhesion molecule 1 (PH-20 hyaluronidase, zona
 pellucida binding)     -0.0019147568464803436
GO:0007339: binding of sperm to zona pellucida  ADAM2   ADAM metallopeptidase domain 2  0.00080521936
31786543
GO:0007339: binding of sperm to zona pellucida  CRISP1  cysteine-rich secretory protein 1       9.152
250557947086e-5
GO:0007339: binding of sperm to zona pellucida  SPAM1   sperm adhesion molecule 1 (PH-20 hyaluronidas
e, zona pellucida binding)      -0.00188338477310234
GO:0007342: fusion of sperm to egg plasma membrane      ADAM2   ADAM metallopeptidase domain 2  0.000
8106749459156312

GO:0007342: fusion of sperm to egg plasma membrane CD9 CD9 molecule -0.002584966462837272
GO:0007342: fusion of sperm to egg plasma membrane CRISP1 cysteine-rich secretory protein 1
9.054040874410055e-5
GO:0007342: fusion of sperm to egg plasma membrane ROPN1B rhophilin associated tail protein 1B
0.0005975338370428852
GO:0007342: fusion of sperm to egg plasma membrane SERPINA5 serpin peptidase inhibitor, c
lade A (alpha-1 antiproteinase, antitrypsin), member 5 -0.0007106164040647417
GO:0007342: fusion of sperm to egg plasma membrane SPAM1 sperm adhesion molecule 1 (PH-20 hyal
uronidase, zona pellucida binding) -0.0018989810548254603
GO:0007342: fusion of sperm to egg plasma membrane TPST2 tyrosylprotein sulfotransferase 2
0.0007132110022684666
GO:0008542: visual learning ADAM2 ADAM metallopeptidase domain 2 0.000809818972871966
GO:0008542: visual learning CHRNB2 cholinergic receptor, nicotinic, beta 2 (neuronal) -0.00
0776983568262972
GO:0008542: visual learning DRD2 dopamine receptor D2 -0.00023370452311025205
GO:0008542: visual learning HIF1A hypoxia inducible factor 1, alpha subunit (basic helix-loop-h
elix transcription factor) -0.0006456804283043314
GO:0008542: visual learning HTT huntingtin -0.0009699089190292239
GO:0008542: visual learning ITGB1 integrin, beta 1 (fibronectin receptor, beta polypeptide, ant
igen CD29 includes MDF2, MSK12) 0.0024216526595014867
GO:0008542: visual learning KIT v-kit Hardy-Zuckerman 4 feline sarcoma viral oncogene homolog
0.0002734956918947965
GO:0008542: visual learning RGS14 regulator of G-protein signaling 14 -0.001580169586523755
3
GO:0030534: adult behavior ADAM2 ADAM metallopeptidase domain 2 0.0007804693401182167
GO:0030534: adult behavior BBS4 Bardet-Biedl syndrome 4 -0.0004923309987886044
GO:0030534: adult behavior PTEN phosphatase and tensin homolog 3.0920466977540914e-5
GO:0032504: multicellular organism reproduction ADAM2 ADAM metallopeptidase domain 2 0.00081496505
17966671
GO:0032504: multicellular organism reproduction CD9 CD9 molecule -0.002599742537263737
GO:0032504: multicellular organism reproduction SPAM1 sperm adhesion molecule 1 (PH-20 hyaluronidas
e, zona pellucida binding) -0.0019088849472196896
KEGG:05162: Measles ADAR adenosine deaminase, RNA-specific -0.0001367128882662882
KEGG:05162: Measles AKT1 v-akt murine thymoma viral oncogene homolog 1 0.0007287490684174982
KEGG:05162: Measles AKT2 v-akt murine thymoma viral oncogene homolog 2 -0.000624665508244975
KEGG:05162: Measles CCND1 cyclin D1 -0.0026253025827962886
KEGG:05162: Measles CCNE1 cyclin E1 0.0003695413706937553
KEGG:05162: Measles CCNE2 cyclin E2 0.0011036413978699482
KEGG:05162: Measles CLEC4M C-type lectin domain family 4, member M -0.0013869551595511627
KEGG:05162: Measles CSNK2B casein kinase 2, beta polypeptide 0.0015246119643528599
KEGG:05162: Measles FAS Fas cell surface death receptor -3.3219924775461695e-5
KEGG:05162: Measles GSK3B glycogen synthase kinase 3 beta 0.0015454716412320682
KEGG:05162: Measles HSPA2 heat shock 70kDa protein 2 -0.00010593951113440394
KEGG:05162: Measles IFNA2 interferon, alpha 2 -0.00171334098322788
KEGG:05162: Measles IFNG interferon, gamma -4.891538804258324e-5
KEGG:05162: Measles IL12B interleukin 12B 0.0012745687353103083
KEGG:05162: Measles IL4 interleukin 4 0.00025943445894402933
KEGG:05162: Measles IRAK1 interleukin-1 receptor-associated kinase 1 -0.001604958793645529
1
KEGG:05162: Measles IRF7 interferon regulatory factor 7 -0.001303843663897984
KEGG:05162: Measles JAK2 Janus kinase 2 -3.222539521968384e-5
KEGG:05162: Measles PIK3CD phosphatidylinositol-4,5-bisphosphate 3-kinase, catalytic subunit del
ta -0.0007789979071440775
KEGG:05162: Measles RCHY1 ring finger and CHY zinc finger domain containing 1, E3 ubiquitin pro
tein ligase -0.0010955495125526656
KEGG:05162: Measles STAT5A signal transducer and activator of transcription 5A 0.00159278950
10371643
KEGG:05162: Measles TNFAIP3 tumor necrosis factor, alpha-induced protein 3 0.0011138320160672033
KEGG:05162: Measles TNFRSF10C tumor necrosis factor receptor superfamily, member 10c, decoy
without an intracellular domain 0.00031694623567988454
KEGG:05162: Measles TP53 tumor protein p53 0.0011722475970647732
KEGG:05162: Measles TP73 tumor protein p73 0.0010255305142626693
KEGG:05164: Influenza A ADAR adenosine deaminase, RNA-specific -0.00013830670650294934
KEGG:05164: Influenza A AKT1 v-akt murine thymoma viral oncogene homolog 1 0.0007154224829440939
KEGG:05164: Influenza A AKT2 v-akt murine thymoma viral oncogene homolog 2 -0.000617530756299901
8
KEGG:05164: Influenza A CCL2 chemokine (C-C motif) ligand 2 0.000800229940272966
KEGG:05164: Influenza A CXCL10 chemokine (C-X-C motif) ligand 10 5.592860593354246e-5
KEGG:05164: Influenza A DNAJB1 DnaJ (Hsp40) homolog, subfamily B, member 1 -0.000233764990300689
5
KEGG:05164: Influenza A FAS Fas cell surface death receptor -3.269710544663396e-5
KEGG:05164: Influenza A GSK3B glycogen synthase kinase 3 beta 0.0015211235210667042
KEGG:05164: Influenza A HSPA2 heat shock 70kDa protein 2 -9.445156633811744e-5
KEGG:05164: Influenza A ICAM1 intercellular adhesion molecule 1 0.0007222394142386343
KEGG:05164: Influenza A IFNA2 interferon, alpha 2 -0.0016872938024025945
KEGG:05164: Influenza A IFNG interferon, gamma -4.922826925518964e-5
KEGG:05164: Influenza A IL12B interleukin 12B 0.0012597054317034246
KEGG:05164: Influenza A IRF7 interferon regulatory factor 7 -0.0012855425305344482
KEGG:05164: Influenza A JAK2 Janus kinase 2 -3.703844849291684e-5
KEGG:05164: Influenza A PIK3CD phosphatidylinositol-4,5-bisphosphate 3-kinase, catalytic subunit del
ta -0.000766608869124104

KEGG:05164: Influenza A PML     promyelocytic leukemia  -0.0006740708732245696
KEGG:05164: Influenza A PRKCA   protein kinase C, alpha -6.329865720102731e-6
KEGG:05164: Influenza A RAF1    Raf-1 proto-oncogene, serine/threonine kinase   0.0014722450759271897
KEGG:05164: Influenza A TNFRSF10C       tumor necrosis factor receptor superfamily, member 10c, decoy without an intracellular domain 0.00031677110527670953
KEGG:04623: Cytosolic DNA-sensing pathway       ADAR    adenosine deaminase, RNA-specific       -0.0001375756361813 3967
KEGG:04623: Cytosolic DNA-sensing pathway       CXCL10  chemokine (C-X-C motif) ligand 10       5.91756771637628e-5
KEGG:04623: Cytosolic DNA-sensing pathway       IFNA2   interferon, alpha 2     -0.0017025244418005948
KEGG:04623: Cytosolic DNA-sensing pathway       IRF7    interferon regulatory factor 7  -0.001296299782524989
GO:0001649: osteoblast differentiation  ADAR    adenosine deaminase, RNA-specific       -0.00013647345641657494
GO:0001649: osteoblast differentiation  AKT1    v-akt murine thymoma viral oncogene homolog 1   0.0007310432064968361
GO:0001649: osteoblast differentiation  ASF1A   anti-silencing function 1A histone chaperone    0.001050646417952133
GO:0001649: osteoblast differentiation  BMP4    bone morphogenetic protein 4    -0.0003227616967355373 4
GO:0001649: osteoblast differentiation  COL1A1  collagen, type I, alpha 1       -0.0005270021809212881
GO:0001649: osteoblast differentiation  DLX5    distal-less homeobox 5  -0.003284447664753778
GO:0001649: osteoblast differentiation  FBL     fibrillarin     -0.0002730574108555112
GO:0001649: osteoblast differentiation  GJA1    gap junction protein, alpha 1, 43kDa    -0.00016236491303947556
GO:0001649: osteoblast differentiation  GLI1    GLI family zinc finger 1        -0.0012973024908317552
GO:0001649: osteoblast differentiation  GLI2    GLI family zinc finger 2        0.0018543058393342203
GO:0001649: osteoblast differentiation  HSD17B4 hydroxysteroid (17-beta) dehydrogenase 4        -0.0011633178153449323
GO:0001649: osteoblast differentiation  IGFBP3  insulin-like growth factor binding protein 3    0.0008368225771456333
GO:0001649: osteoblast differentiation  IHH     indian hedgehog -0.0020416422971991904
GO:0001649: osteoblast differentiation  LEF1    lymphoid enhancer-binding factor 1      -0.0001002677334069404
GO:0001649: osteoblast differentiation  LGR4    leucine-rich repeat containing G protein-coupled receptor 4  0.0003086131415210507
GO:0001649: osteoblast differentiation  MEF2C   myocyte enhancer factor 2C      0.000973933940438167
GO:0001649: osteoblast differentiation  SFRP1   secreted frizzled-related protein 1     0.0012832922784907618
GO:0001649: osteoblast differentiation  SMO     smoothened, frizzled class receptor     0.0021402102557966196
GO:0001649: osteoblast differentiation  TNC     tenascin C      0.0007346243974645873
GO:0001649: osteoblast differentiation  TWIST1  twist family bHLH transcription factor 1        -0.0013423248367231636
GO:0001649: osteoblast differentiation  WWTR1   WW domain containing transcription regulator 1  0.000901384285867 7846
GO:0002244: hematopoietic progenitor cell differentiation       ADAR    adenosine deaminase, RNA-specific    -0.00013602220375612058
GO:0002244: hematopoietic progenitor cell differentiation       BMP4    bone morphogenetic protein 4 -0.00032074253230548387
GO:0002244: hematopoietic progenitor cell differentiation       INHBA   inhibin, beta A -0.0013437987085223018
GO:0002244: hematopoietic progenitor cell differentiation       PLEK    pleckstrin      0.0009786768375722964
GO:0002244: hematopoietic progenitor cell differentiation       SFRP1   secreted frizzled-related protein 1  0.0012736910229649457
GO:0002244: hematopoietic progenitor cell differentiation       TGFB1   transforming growth factor, beta 1   -7.285023274196377e-5
GO:0002244: hematopoietic progenitor cell differentiation       TOP2A   topoisomerase (DNA) II alpha 170kDa -8.170358880283573e-5
GO:0002566: somatic diversification of immune receptors via somatic mutation    ADAR    adenosine deaminase, RNA-specific    -0.00013990644784975787
GO:0006382: adenosine to inosine editing        ADAR    adenosine deaminase, RNA-specific       -0.0001399064478497 5787
GO:0006397: mRNA processing     ADAR    adenosine deaminase, RNA-specific       -0.0001368073612369 9487
GO:0006397: mRNA processing     JMJD6   jumonji domain containing 6     0.0036057524520759334
GO:0006397: mRNA processing     SF3B2   splicing factor 3b, subunit 2, 145kDa   0.0006868879876014877
GO:0006397: mRNA processing     ZPR1    ZPR1 zinc finger        -0.000211143475087956
GO:0006606: protein import into nucleus ADAR    adenosine deaminase, RNA-specific       -0.00013936023645948146
GO:0006606: protein import into nucleus HTT     huntingtin      -0.0009425434904552673
GO:0006606: protein import into nucleus KPNB1   karyopherin (importin) beta 1   0.0007528784128011074
GO:0006611: protein export from nucleus ADAR    adenosine deaminase, RNA-specific       -0.00013675380177956807
GO:0006611: protein export from nucleus EGR2    early growth response 2 0.001433452999880 5965
GO:0006611: protein export from nucleus GSK3B   glycogen synthase kinase 3 beta 0.0015447044502712784
GO:0006611: protein export from nucleus TGFB1   transforming growth factor, beta 1      -7.2450906819

10457e-5
GO:0009615: response to virus   ADAR    adenosine deaminase, RNA-specific       -0.00013723254133641113
GO:0009615: response to virus   CCL8    chemokine (C-C motif) ligand 8  -0.000585060030621215
GO:0009615: response to virus   CFL1    cofilin 1 (non-muscle)  -0.001156690027099323
GO:0009615: response to virus   CXCL12  chemokine (C-X-C motif) ligand 12       -0.00119025062443047
GO:0009615: response to virus   CXCR4   chemokine (C-X-C motif) receptor 4      0.0007819109697478861
GO:0009615: response to virus   GATA3   GATA binding protein 3  -3.6762809584126454e-5
GO:0009615: response to virus   HMGA1   high mobility group AT-hook 1   -0.00031544907302057996
GO:0009615: response to virus   HMGA2   high mobility group AT-hook 2   0.0014888663935048638
GO:0009615: response to virus   IFNG    interferon, gamma       -4.881796501282638e-5
GO:0009615: response to virus   IRF7    interferon regulatory factor 7  -0.0012902643344611482
GO:0009615: response to virus   MEF2C   myocyte enhancer factor 2C      0.00095719994647468
GO:0009615: response to virus   ODC1    ornithine decarboxylase 1       0.0008129549933178987
GO:0009615: response to virus   STMN1   stathmin 1      0.0005468524571078884
GO:0009615: response to virus   TBX21   T-box 21        0.00039470139805897615
GO:0010467: gene expression     ADAR    adenosine deaminase, RNA-specific       -0.00013715040135377294
GO:0010467: gene expression     AIMP2   aminoacyl tRNA synthetase complex-interacting multifunctional protein 2       0.0012074138188740522
GO:0010467: gene expression     AKT1    v-akt murine thymoma viral oncogene homolog 1   0.000727950094718941
GO:0010467: gene expression     AR      androgen receptor       0.0026330799688653655
GO:0010467: gene expression     CCNC    cyclin C        0.000564711047212912
GO:0010467: gene expression     CDC40   cell division cycle 40  0.00415355630825537
GO:0010467: gene expression     CDK8    cyclin-dependent kinase 8       0.0026058946731098613
GO:0010467: gene expression     CDKN2B  cyclin-dependent kinase inhibitor 2B (p15, inhibits CDK4) -0.001830242611780264
GO:0010467: gene expression     DCP1A   decapping mRNA 1A       -0.0011889563362459156
GO:0010467: gene expression     DICER1  dicer 1, ribonuclease type III  -1.6280070019297506e-5
GO:0010467: gene expression     E2F4    E2F transcription factor 4, p107/p130-binding   -0.002339110389203165
GO:0010467: gene expression     EDA     ectodysplasin A -0.0008111067719026453
GO:0010467: gene expression     EIF4G1  eukaryotic translation initiation factor 4 gamma, 1     0.0007399557932371727
GO:0010467: gene expression     ESR1    estrogen receptor 1     -0.0009437023007935861
GO:0010467: gene expression     EZH2    enhancer of zeste 2 polycomb repressive complex 2 subunit -0.00013822702540941705
GO:0010467: gene expression     HDAC2   histone deacetylase 2   -0.001205966651079079
GO:0010467: gene expression     IGF2BP2 insulin-like growth factor 2 mRNA binding protein 2     -0.0009345954717638301
GO:0010467: gene expression     IGF2BP3 insulin-like growth factor 2 mRNA binding protein 3     -0.0003840562437325937
GO:0010467: gene expression     LARS2   leucyl-tRNA synthetase 2, mitochondrial 0.0004659636778055991
GO:0010467: gene expression     MED1    mediator complex subunit 1      0.0011243407578242998
GO:0010467: gene expression     MTERF1  mitochondrial transcription termination factor 1        0.0002856379806586485
GO:0010467: gene expression     MYC     v-myc avian myelocytomatosis viral oncogene homolog     -0.001134678885788122
GO:0010467: gene expression     NOTCH1  notch 1 0.000516714335666352
GO:0010467: gene expression     NR1H3   nuclear receptor subfamily 1, group H, member 3 0.0007994962327485641
GO:0010467: gene expression     NR2E1   nuclear receptor subfamily 2, group E, member 1 -0.002967461502036658
GO:0010467: gene expression     NRBF2   nuclear receptor binding factor 2       -0.0009485175135698687
GO:0010467: gene expression     NUP153  nucleoporin 153kDa      0.0008206700334699571
GO:0010467: gene expression     PGR     progesterone receptor   0.0003525480759175039
GO:0010467: gene expression     POLR2D  polymerase (RNA) II (DNA directed) polypeptide D        0.001848778582361848
GO:0010467: gene expression     PRKCA   protein kinase C, alpha -6.002875484826545e-6
GO:0010467: gene expression     PRKCD   protein kinase C, delta -0.0011237942402427213
GO:0010467: gene expression     PSMA5   proteasome (prosome, macropain) subunit, alpha type, 5  0.0003303653219311149
GO:0010467: gene expression     PSMD11  proteasome (prosome, macropain) 26S subunit, non-ATPase, 11 -0.0010933626387221022
GO:0010467: gene expression     PSMD13  proteasome (prosome, macropain) 26S subunit, non-ATPase, 13 -6.052002956505354e-5
GO:0010467: gene expression     PTEN    phosphatase and tensin homolog  1.87232133069115e-5
GO:0010467: gene expression     RARG    retinoic acid receptor, gamma   -0.0023127320494180426
GO:0010467: gene expression     RBBP4   retinoblastoma binding protein 4        -0.0018583257610416829
GO:0010467: gene expression     RBPJ    recombination signal binding protein for immunoglobulin kappa J region        0.0009794693242463483
GO:0010467: gene expression     RXRA    retinoid X receptor, alpha      0.0011121463300903574
GO:0010467: gene expression     SEH1L   SEH1-like (S. cerevisiae)       -0.0007317701174542198
GO:0010467: gene expression     SERPINE1        serpin peptidase inhibitor, clade E (nexin, plasminogen activator inhibitor type 1), member 1        0.00011305298860409827
GO:0010467: gene expression     SF3B2   splicing factor 3b, subunit 2, 145kDa   0.0006811560016530254
GO:0010467: gene expression     SIRT1   sirtuin 1       -2.5139683459379486e-6
GO:0010467: gene expression     SKI     SKI proto-oncogene      -0.000668966627975757

GO:0010467: gene expression     SMC1A   structural maintenance of chromosomes 1A       -0.0010173157563004764
GO:0010467: gene expression     SNRPB   small nuclear ribonucleoprotein polypeptides B and B1   0.0006619905665319625
GO:0010467: gene expression     TFDP1   transcription factor Dp-1       0.0014375611998349554
GO:0010467: gene expression     THRA    thyroid hormone receptor, alpha 0.0007613593629614998
GO:0010467: gene expression     THRB    thyroid hormone receptor, beta  0.0019392091276832675
GO:0010467: gene expression     TP53    tumor protein p53       0.0011701994104631075
GO:0010467: gene expression     TPR     translocated promoter region, nuclear basket protein    -0.0006954732631280886
GO:0010467: gene expression     VDR     vitamin D (1,25- dihydroxyvitamin D3) receptor  0.0005629512267216586
GO:0010467: gene expression     WDR77   WD repeat domain 77     0.0001314474914996118
GO:0010467: gene expression     WWTR1   WW domain containing transcription regulator 1  0.000895308704260595
GO:0010467: gene expression     YAP1    Yes-associated protein 1        -0.0001648417837384787
GO:0010467: gene expression     YBX1    Y box binding protein 1 -0.0009133835980316579
GO:0016553: base conversion or substitution editing     ADAR    adenosine deaminase, RNA-specific -0.00013990644784975787
GO:0016556: mRNA modification   ADAR    adenosine deaminase, RNA-specific       -0.00013990644784975787
GO:0019221: cytokine-mediated signaling pathway ADAR    adenosine deaminase, RNA-specific       -0.00013724565779600143
GO:0019221: cytokine-mediated signaling pathway CCL2    chemokine (C-C motif) ligand 2  0.0008103532508516595
GO:0019221: cytokine-mediated signaling pathway CSF3R   colony stimulating factor 3 receptor (granulocyte)   -7.92633545069137e-5
GO:0019221: cytokine-mediated signaling pathway CX3CL1  chemokine (C-X3-C motif) ligand 1       0.002098623960739224
GO:0019221: cytokine-mediated signaling pathway EGR1    early growth response 1 0.0010857561953761558
GO:0019221: cytokine-mediated signaling pathway EIF4G1  eukaryotic translation initiation factor 4 gamma, 1  0.0007371722881954919
GO:0019221: cytokine-mediated signaling pathway FLT3    fms-related tyrosine kinase 3   -0.0006749201034360919
GO:0019221: cytokine-mediated signaling pathway ICAM1   intercellular adhesion molecule 1       0.0007265161158796249
GO:0019221: cytokine-mediated signaling pathway IFNA2   interferon, alpha 2     -0.001705710759405729
GO:0019221: cytokine-mediated signaling pathway IFNG    interferon, gamma       -4.9015339485445666e-5
GO:0019221: cytokine-mediated signaling pathway IL12B   interleukin 12B 0.0012703394824661383
GO:0019221: cytokine-mediated signaling pathway IL20RA  interleukin 20 receptor, alpha  0.00017454558999419149
GO:0019221: cytokine-mediated signaling pathway IL6ST   interleukin 6 signal transducer 0.0018646748087262468
GO:0019221: cytokine-mediated signaling pathway IRF6    interferon regulatory factor 6  -0.0016815704969173608
GO:0019221: cytokine-mediated signaling pathway IRF7    interferon regulatory factor 7  -0.0012985258191372687
GO:0019221: cytokine-mediated signaling pathway JAK2    Janus kinase 2  -3.3756220379334146e-5
GO:0019221: cytokine-mediated signaling pathway KIT     v-kit Hardy-Zuckerman 4 feline sarcoma viral oncogene homolog       0.00026899525560018187
GO:0019221: cytokine-mediated signaling pathway KPNA3   karyopherin alpha 3 (importin alpha 4)  -0.0008139575058525815
GO:0019221: cytokine-mediated signaling pathway KPNB1   karyopherin (importin) beta 1   0.0007677265884137691
GO:0019221: cytokine-mediated signaling pathway LRP8    low density lipoprotein receptor-related protein 8, apolipoprotein e receptor       -0.000926951950682721
GO:0019221: cytokine-mediated signaling pathway NEDD4   neural precursor cell expressed, developmentally down-regulated 4, E3 ubiquitin protein ligase       0.0022499782277308183
GO:0019221: cytokine-mediated signaling pathway NUP153  nucleoporin 153kDa      0.0008163710379416999
GO:0019221: cytokine-mediated signaling pathway PML     promyelocytic leukemia  -0.0006776433942854828
GO:0019221: cytokine-mediated signaling pathway PRKCD   protein kinase C, delta -0.0011187482464835316
GO:0019221: cytokine-mediated signaling pathway PTPN1   protein tyrosine phosphatase, non-receptor type 1    -0.0010340136506469672
GO:0019221: cytokine-mediated signaling pathway PTPN2   protein tyrosine phosphatase, non-receptor type 2    -0.0003280394691560234
GO:0019221: cytokine-mediated signaling pathway SEH1L   SEH1-like (S. cerevisiae)       -0.0007295901577404032
GO:0019221: cytokine-mediated signaling pathway TPR     translocated promoter region, nuclear basket protein        -0.00069302240210558
GO:0030218: erythrocyte differentiation ADAR    adenosine deaminase, RNA-specific       -0.00013700878779790182
GO:0030218: erythrocyte differentiation BMP4    bone morphogenetic protein 4    -0.00031723226721966445
GO:0030218: erythrocyte differentiation GATA3   GATA binding protein 3  -3.632909010990944e-5
GO:0030218: erythrocyte differentiation HIPK2   homeodomain interacting protein kinase 2        0.0007597017919034308
GO:0030218: erythrocyte differentiation INHA    inhibin, alpha  0.0002109105270446142
GO:0030218: erythrocyte differentiation INHBA   inhibin, beta A -0.001321821689531759

GO:0030218: erythrocyte differentiation JAK2    Janus kinase 2  -3.458261394232065e-5
GO:0030218: erythrocyte differentiation KIT     v-kit Hardy-Zuckerman 4 feline sarcoma viral oncogene homolog 0.00026441402624056585
GO:0030218: erythrocyte differentiation LYN     LYN proto-oncogene, Src family tyrosine kinase  -0.0013975350218323362
GO:0030218: erythrocyte differentiation PTPN2   protein tyrosine phosphatase, non-receptor type 2 -0.0003247990752857284
GO:0030218: erythrocyte differentiation THRA    thyroid hormone receptor, alpha 0.0007511726389885195
GO:0031054: pre-miRNA processing        ADAR    adenosine deaminase, RNA-specific       -0.0001365135916260654
GO:0031054: pre-miRNA processing        DICER1  dicer 1, ribonuclease type III  -3.720827120463265e-5
GO:0035280: miRNA loading onto RISC involved in gene silencing by miRNA ADAR    adenosine deaminase, RNA-specific   -0.0001365135916260654
GO:0035280: miRNA loading onto RISC involved in gene silencing by miRNA DICER1  dicer 1, ribonuclease type III        -3.720827120463265e-5
GO:0035455: response to interferon-alpha        ADAR    adenosine deaminase, RNA-specific       -0.0013990644784975787
GO:0043066: negative regulation of apoptotic process    ADAR    adenosine deaminase, RNA-specific -0.00013656307425511367
GO:0043066: negative regulation of apoptotic process    AKT1    v-akt murine thymoma viral oncogene homolog 1        0.000731883291528895
GO:0043066: negative regulation of apoptotic process    ALOX12  arachidonate 12-lipoxygenase    -0.0019565985768180473
GO:0043066: negative regulation of apoptotic process    ANGPT1  angiopoietin 1  0.0009019132492647955
GO:0043066: negative regulation of apoptotic process    ASCL1   achaete-scute family bHLH transcription factor 1     -0.0014349478609823748
GO:0043066: negative regulation of apoptotic process    ASIC2   acid-sensing (proton-gated) ion channel 2    0.0022830086044381883
GO:0043066: negative regulation of apoptotic process    ATF5    activating transcription factor 5 -0.0027390757032819033
GO:0043066: negative regulation of apoptotic process    AURKA   aurora kinase A 0.0010097033297251838
GO:0043066: negative regulation of apoptotic process    AVP     arginine vasopressin    -0.0009390430798288736
GO:0043066: negative regulation of apoptotic process    AZU1    azurocidin 1    0.0001982389521986351
GO:0043066: negative regulation of apoptotic process    BARD1   BRCA1 associated RING domain 1  0.001746074914910911
GO:0043066: negative regulation of apoptotic process    BCL2    B-cell CLL/lymphoma 2   -4.964459510846809e-6
GO:0043066: negative regulation of apoptotic process    BFAR    bifunctional apoptosis regulator -0.0003708538608130184
GO:0043066: negative regulation of apoptotic process    BIRC5   baculoviral IAP repeat containing 5 -0.0002604841018292551
GO:0043066: negative regulation of apoptotic process    BMP4    bone morphogenetic protein 4    -0.0003230735012952702
GO:0043066: negative regulation of apoptotic process    BNIP3   BCL2/adenovirus E1B 19kDa interacting protein 3       0.002917156946463022
GO:0043066: negative regulation of apoptotic process    BNIP3L  BCL2/adenovirus E1B 19kDa interacting protein 3-like  -5.759533705013394e-5
GO:0043066: negative regulation of apoptotic process    CAMK1D  calcium/calmodulin-dependent protein kinase ID      -0.0025917899344116116
GO:0043066: negative regulation of apoptotic process    CDK1    cyclin-dependent kinase 1       0.0003119884456876906
GO:0043066: negative regulation of apoptotic process    CFL1    cofilin 1 (non-muscle)  -0.0011770057379542303
GO:0043066: negative regulation of apoptotic process    CIB1    calcium and integrin binding 1 (calmyrin)    5.4286365686468264e-5
GO:0043066: negative regulation of apoptotic process    CRYAA   crystallin, alpha A     -0.0006763605736025916
GO:0043066: negative regulation of apoptotic process    CRYAB   crystallin, alpha B     0.0009862795861388942
GO:0043066: negative regulation of apoptotic process    DHCR24  24-dehydrocholesterol reductase -0.0017100981450029037
GO:0043066: negative regulation of apoptotic process    EGFR    epidermal growth factor receptor 0.000687225415849133
GO:0043066: negative regulation of apoptotic process    EGR1    early growth response 1 0.0010946950640368608
GO:0043066: negative regulation of apoptotic process    EGR2    early growth response 2 0.001441658660244234
GO:0043066: negative regulation of apoptotic process    ERBB4   v-erb-b2 avian erythroblastic leukemia viral oncogene homolog 4      -0.00018360934112878757
GO:0043066: negative regulation of apoptotic process    FAS     Fas cell surface death receptor -3.334421057655442e-5
GO:0043066: negative regulation of apoptotic process    FOXE3   forkhead box E3 0.0010523252216038269
GO:0043066: negative regulation of apoptotic process    FOXO1   forkhead box O1 0.0018028519283239256
GO:0043066: negative regulation of apoptotic process    GATA6   GATA binding protein 6  -2.8040580375928132e-5
GO:0043066: negative regulation of apoptotic process    GDNF    glial cell derived neurotrophic factor       0.0004452756652995561
GO:0043066: negative regulation of apoptotic process    GLI3    GLI family zinc finger 3        -0.0021552163228767017
GO:0043066: negative regulation of apoptotic process    GREM1   gremlin 1, DAN family BMP antagonist

-0.000829158487145094
GO:0043066: negative regulation of apoptotic process    GSK3B   glycogen synthase kinase 3 beta 0.001
5515058164287826
GO:0043066: negative regulation of apoptotic process    GSTP1   glutathione S-transferase pi 1  0.000
23224553744555224
GO:0043066: negative regulation of apoptotic process    HDAC2   histone deacetylase 2   -0.0012085838
81178543
GO:0043066: negative regulation of apoptotic process    HMGA2   high mobility group AT-hook 2   0.001
5110520694431187
GO:0043066: negative regulation of apoptotic process    HPN     hepsin  0.0031625224127230685
GO:0043066: negative regulation of apoptotic process    IGF1    insulin-like growth factor 1 (somatom
edin C) 0.0001313927588091418
GO:0043066: negative regulation of apoptotic process    IGF1R   insulin-like growth factor 1 receptor
0.0010731641395217626
GO:0043066: negative regulation of apoptotic process    IHH     indian hedgehog -0.002043848834352691
GO:0043066: negative regulation of apoptotic process    IL4     interleukin 4   0.0002596022801489126
GO:0043066: negative regulation of apoptotic process    IL6ST   interleukin 6 signal transducer 0.001
8780714790041751
GO:0043066: negative regulation of apoptotic process    IL7     interleukin 7   0.0008728000895207234
GO:0043066: negative regulation of apoptotic process    IRAK1   interleukin-1 receptor-associated kin
ase 1   -0.001611254775255239
GO:0043066: negative regulation of apoptotic process    KIF14   kinesin family member 14        0.000
4867689475413461
GO:0043066: negative regulation of apoptotic process    KRT18   keratin 18      -0.001019567446735863
2
GO:0043066: negative regulation of apoptotic process    LEF1    lymphoid enhancer-binding factor 1
-0.00010025273476476138
GO:0043066: negative regulation of apoptotic process    LEP     leptin  0.003196334262655334
GO:0043066: negative regulation of apoptotic process    MAD2L1  MAD2 mitotic arrest deficient-like 1
 (yeast)        0.0003193345155635912
GO:0043066: negative regulation of apoptotic process    MED1    mediator complex subunit 1      0.001
1312947550757774
GO:0043066: negative regulation of apoptotic process    MSX1    msh homeobox 1  -0.002773633304305703
6
GO:0043066: negative regulation of apoptotic process    MYC     v-myc avian myelocytomatosis viral on
cogene homolog  -0.0011379328297025277
GO:0043066: negative regulation of apoptotic process    NR2E1   nuclear receptor subfamily 2, group
 E, member 1    -0.002979045549123312
GO:0043066: negative regulation of apoptotic process    PAX2    paired box 2    -0.001616701654041186
3
GO:0043066: negative regulation of apoptotic process    PCNT    pericentrin     0.001789667874352318
GO:0043066: negative regulation of apoptotic process    PDGFRB  platelet-derived growth factor recept
or, beta polypeptide    -0.0003621973767381215
GO:0043066: negative regulation of apoptotic process    PHB2    prohibitin 2    -0.000737110001924458
7
GO:0043066: negative regulation of apoptotic process    PLK1    polo-like kinase 1      0.00101960923
10914523
GO:0043066: negative regulation of apoptotic process    PLK3    polo-like kinase 3      0.00256763538
44121015
GO:0043066: negative regulation of apoptotic process    PRKCZ   protein kinase C, zeta  -0.0016030255
100953586
GO:0043066: negative regulation of apoptotic process    PRKDC   protein kinase, DNA-activated, cataly
tic polypeptide -0.0018581700433447819
GO:0043066: negative regulation of apoptotic process    PRLR    prolactin receptor      0.00218635787
5551232
GO:0043066: negative regulation of apoptotic process    PSMA5   proteasome (prosome, macropain) subun
it, alpha type, 5       0.00033248937418795107
GO:0043066: negative regulation of apoptotic process    PSMD11  proteasome (prosome, macropain) 26S s
ubunit, non-ATPase, 11  -0.0010961632989426242
GO:0043066: negative regulation of apoptotic process    PSMD13  proteasome (prosome, macropain) 26S s
ubunit, non-ATPase, 13  -5.951709231238094e-5
GO:0043066: negative regulation of apoptotic process    PTEN    phosphatase and tensin homolog  1.672
7331316222756e-5
GO:0043066: negative regulation of apoptotic process    RAF1    Raf-1 proto-oncogene, serine/threonin
e kinase        0.0015010131652367297
GO:0043066: negative regulation of apoptotic process    RARG    retinoic acid receptor, gamma   -0.00
23228513270552547
GO:0043066: negative regulation of apoptotic process    RPS6KA1 ribosomal protein S6 kinase, 90kDa, p
olypeptide 1    -0.0025176883886868016
GO:0043066: negative regulation of apoptotic process    SFRP1   secreted frizzled-related protein 1
0.0012846742657332071
GO:0043066: negative regulation of apoptotic process    SHH     sonic hedgehog  0.000601123478103724
GO:0043066: negative regulation of apoptotic process    SIRT1   sirtuin 1       -1.4205056609003693e-
6
GO:0043066: negative regulation of apoptotic process    SMO     smoothened, frizzled class receptor
0.0021424862395646744
GO:0043066: negative regulation of apoptotic process    SOX10   SRY (sex determining region Y)-box 10
0.00019152662733506722
GO:0043066: negative regulation of apoptotic process    SOX9    SRY (sex determining region Y)-box 9
-0.000519435686652471
GO:0043066: negative regulation of apoptotic process    SPHK1   sphingosine kinase 1    0.00181361813

50504928
GO:0043066: negative regulation of apoptotic process    TBX3    T-box 3 0.001227679395935684
GO:0043066: negative regulation of apoptotic process    TFAP2A  transcription factor AP-2 alpha (activating enhancer binding protein 2 alpha)       0.00048048132567914245
GO:0043066: negative regulation of apoptotic process    THBS1   thrombospondin 1        -0.0010374064289076868
GO:0043066: negative regulation of apoptotic process    TMBIM6  transmembrane BAX inhibitor motif containing 6       -0.00014791609261999048
GO:0043066: negative regulation of apoptotic process    TP53    tumor protein p53       0.0011779338336137713
GO:0043066: negative regulation of apoptotic process    TWIST1  twist family bHLH transcription factor 1     -0.0013438239194713578
GO:0043066: negative regulation of apoptotic process    VEGFA   vascular endothelial growth factor A 0.0005962379960733929
GO:0043066: negative regulation of apoptotic process    WNT5A   wingless-type MMTV integration site family, member 5A        -0.0006685224561495124
GO:0043066: negative regulation of apoptotic process    WNT7A   wingless-type MMTV integration site family, member 7A        1.9431653555381154e-5
GO:0043066: negative regulation of apoptotic process    WT1     Wilms tumor 1   -0.0005078466928052352
GO:0043066: negative regulation of apoptotic process    XBP1    X-box binding protein 1 0.0002674536241916083
GO:0044387: negative regulation of protein kinase activity by regulation of protein phosphorylation ADAR    adenosine deaminase, RNA-specific       -0.00014172340387966562
GO:0044387: negative regulation of protein kinase activity by regulation of protein phosphorylation CDK5RAP3        CDK5 regulatory subunit associated protein 3    0.0011148389451219039
GO:0045070: positive regulation of viral genome replication     ADAR    adenosine deaminase, RNA-specific    -0.00012748806385473504
GO:0045070: positive regulation of viral genome replication     TOP2A   topoisomerase (DNA) II alpha 170kDa -0.0001035351432502088
GO:0045071: negative regulation of viral genome replication     ADAR    adenosine deaminase, RNA-specific    -0.00013782742167199437
GO:0045071: negative regulation of viral genome replication     PROX1   prospero homeobox 1     0.0011028137011656733
GO:0045087: innate immune response      ADAR    adenosine deaminase, RNA-specific       -0.0001369097389079575
GO:0045087: innate immune response      AGER    advanced glycosylation end product-specific receptor -0.00017408994298191992
GO:0045087: innate immune response      AKT1    v-akt murine thymoma viral oncogene homolog 1   0.0007291312940672268
GO:0045087: innate immune response      ANGPT1  angiopoietin 1  0.000899208547749044
GO:0045087: innate immune response      BCL2    B-cell CLL/lymphoma 2   -4.63977917433393e-6
GO:0045087: innate immune response      C1QA    complement component 1, q subcomponent, A chain 0.0005413367481662143
GO:0045087: innate immune response      C2      complement component 2  -0.0003033928982477654
GO:0045087: innate immune response      C3      complement component 3  0.002020819095584042
GO:0045087: innate immune response      C8B     complement component 8, beta polypeptide        0.00023004215933241588
GO:0045087: innate immune response      C8G     complement component 8, gamma polypeptide       -0.0011927590870437687
GO:0045087: innate immune response      CDC42   cell division cycle 42  0.0012380849751142213
GO:0045087: innate immune response      CFL1    cofilin 1 (non-muscle)  -0.0011725829120191697
GO:0045087: innate immune response      CLEC4M  C-type lectin domain family 4, member M -0.001387617981078733
GO:0045087: innate immune response      CREB1   cAMP responsive element binding protein 1       0.0006563420940543394
GO:0045087: innate immune response      CTNNB1  catenin (cadherin-associated protein), beta 1, 88kDa -0.00011721946459964424
GO:0045087: innate immune response      CTSK    cathepsin K     -0.0003610400192848525
GO:0045087: innate immune response      DOCK1   dedicator of cytokinesis 1      -0.0006824929634463713
GO:0045087: innate immune response      EGFR    epidermal growth factor receptor        0.0006835099630099177
GO:0045087: innate immune response      ERBB4   v-erb-b2 avian erythroblastic leukemia viral oncogene homolog 4       -0.00018118034906298736
GO:0045087: innate immune response      FGF3    fibroblast growth factor 3      0.0015582032755426124
GO:0045087: innate immune response      FGF5    fibroblast growth factor 5      -0.001092025283465081
GO:0045087: innate immune response      FGF7    fibroblast growth factor 7      0.0006327941327195193
GO:0045087: innate immune response      FGF8    fibroblast growth factor 8 (androgen-induced)   0.00098167543278885
GO:0045087: innate immune response      FGFR2   fibroblast growth factor receptor 2     0.0007614734031902966
GO:0045087: innate immune response      FGFR3   fibroblast growth factor receptor 3     0.00022228057360437894
GO:0045087: innate immune response      FOXO1   forkhead box O1 0.001797152413089744
GO:0045087: innate immune response      FOXO3   forkhead box O3 0.0011976005448450797
GO:0045087: innate immune response      FRS2    fibroblast growth factor receptor substrate 2   0.0009277356986840541
GO:0045087: innate immune response      GATA3   GATA binding protein 3  -3.750096269679546e-5
GO:0045087: innate immune response      GRB2    growth factor receptor-bound protein 2  0.00042900931604482535

GO:0045087: innate immune response GSK3B glycogen synthase kinase 3 beta 0.0015463276968181818
GO:0045087: innate immune response HMGB1 high mobility group box 1 -0.00077585209691955
89
GO:0045087: innate immune response IFNA2 interferon, alpha 2 -0.001714251316911292
GO:0045087: innate immune response IRAK1 interleukin-1 receptor-associated kinase 1 -0.00
16061162290777725
GO:0045087: innate immune response IRF7 interferon regulatory factor 7 -0.001304373328927446
2
GO:0045087: innate immune response ITPR1 inositol 1,4,5-trisphosphate receptor, type 1 -0.00
07779355211256258
GO:0045087: innate immune response JAK2 Janus kinase 2 -3.250870257995503e-5
GO:0045087: innate immune response KIT v-kit Hardy-Zuckerman 4 feline sarcoma viral oncogene
homolog 0.000272815337166814 4
GO:0045087: innate immune response LGR4 leucine-rich repeat containing G protein-coupled rece
ptor 4 0.000307816695487188
GO:0045087: innate immune response LYN LYN proto-oncogene, Src family tyrosine kinase -0.00
14182182717488387
GO:0045087: innate immune response MAP3K5 mitogen-activated protein kinase kinase kinase 5
0.00034169417563195193
GO:0045087: innate immune response MEF2C myocyte enhancer factor 2C 0.0009711998910311991
GO:0045087: innate immune response NCK1 NCK adaptor protein 1 -0.0007843203023177386
GO:0045087: innate immune response PADI4 peptidyl arginine deiminase, type IV -0.0008246749
297307712
GO:0045087: innate immune response PAK1 p21 protein (Cdc42/Rac)-activated kinase 1 -0.00
21837200619320833
GO:0045087: innate immune response PAK3 p21 protein (Cdc42/Rac)-activated kinase 3 -0.00
12570743350347783
GO:0045087: innate immune response PDGFRB platelet-derived growth factor receptor, beta polypep
tide -0.00035921679789667065
GO:0045087: innate immune response PIK3CD phosphatidylinositol-4,5-bisphosphate 3-kinase, catal
ytic subunit delta -0.0007793959194915437
GO:0045087: innate immune response PML promyelocytic leukemia -0.0006796648975348978
GO:0045087: innate immune response PRKCA protein kinase C, alpha -6.014805301749246e-6
GO:0045087: innate immune response PRKCD protein kinase C, delta -0.0011256273762663808
GO:0045087: innate immune response PRKDC protein kinase, DNA-activated, catalytic polypeptide
-0.0018530690548876861
GO:0045087: innate immune response PSMA5 proteasome (prosome, macropain) subunit, alpha type,
 5 0.0003309004949413005
GO:0045087: innate immune response PSMD11 proteasome (prosome, macropain) 26S subunit, non-ATPa
se, 11 -0.0010939357184463607
GO:0045087: innate immune response PSMD13 proteasome (prosome, macropain) 26S subunit, non-ATPa
se, 13 -6.0146189128108326e-5
GO:0045087: innate immune response PTEN phosphatase and tensin homolog 1.798925006716671e-5
GO:0045087: innate immune response RAF1 Raf-1 proto-oncogene, serine/threonine kinase 0.001
496055178920178
GO:0045087: innate immune response RASA1 RAS p21 protein activator (GTPase activating protein)
1 -0.00034784005484039734
GO:0045087: innate immune response RPS6KA1 ribosomal protein S6 kinase, 90kDa, polypeptide 1
-0.0025100788340712012
GO:0045087: innate immune response S100A7 S100 calcium binding protein A7 0.0015866887479776808
GO:0045087: innate immune response S100B S100 calcium binding protein B 0.0051741189838140356
GO:0045087: innate immune response SIRT2 sirtuin 2 -0.0008504675302753777
GO:0045087: innate immune response SPTBN1 spectrin, beta, non-erythrocytic 1 0.00160441979
51686752
GO:0045087: innate immune response TNFAIP3 tumor necrosis factor, alpha-induced protein 3 0.001
1142374994470715
GO:0051607: defense response to virus ADAR adenosine deaminase, RNA-specific -0.0001373714
8453157137
GO:0051607: defense response to virus AZU1 azurocidin 1 0.000196616828794953
GO:0051607: defense response to virus BCL2 B-cell CLL/lymphoma 2 -4.800103538771492e-6
GO:0051607: defense response to virus BNIP3 BCL2/adenovirus E1B 19kDa interacting protein 3 0.002
8890941672137897
GO:0051607: defense response to virus BNIP3L BCL2/adenovirus E1B 19kDa interacting protein 3-like
-5.756032665825834e-5
GO:0051607: defense response to virus CXADR coxsackie virus and adenovirus receptor 0.00043004343
585807453
GO:0051607: defense response to virus CXCL10 chemokine (C-X-C motif) ligand 10 5.96967268539
8532e-5
GO:0051607: defense response to virus DICER1 dicer 1, ribonuclease type III -1.6544603795708154e-
5
GO:0051607: defense response to virus IFNA2 interferon, alpha 2 -0.0017045525350927982
GO:0051607: defense response to virus IFNG interferon, gamma -4.90508257323948e-5
GO:0051607: defense response to virus IL12B interleukin 12B 0.0012697272945275389
GO:0051607: defense response to virus PML promyelocytic leukemia -0.0006775239138134028
GO:0060216: definitive hemopoiesis ADAR adenosine deaminase, RNA-specific -0.0001386256
6561192042
GO:0060216: definitive hemopoiesis GATA2 GATA binding protein 2 -0.00042300767220347906
GO:0060216: definitive hemopoiesis KMT2A lysine (K)-specific methyltransferase 2A 0.000
6264229801728786
GO:0060216: definitive hemopoiesis ZFP36L2 ZFP36 ring finger protein-like 2 -0.0003555471
445140842

GO:0060337: type I interferon signaling pathway ADAR    adenosine deaminase, RNA-specific       -0.0001373960988728699
GO:0060337: type I interferon signaling pathway EGR1    early growth response 1 0.001090203709091139
GO:0060337: type I interferon signaling pathway IFNA2   interferon, alpha 2     -0.001712599673081713
GO:0060337: type I interferon signaling pathway IRF6    interferon regulatory factor 6  -0.0016882263485073642
GO:0060337: type I interferon signaling pathway IRF7    interferon regulatory factor 7  -0.0013029546957386532
GO:0060337: type I interferon signaling pathway PTPN1   protein tyrosine phosphatase, non-receptor type 1    -0.0010385328019515672
GO:0060339: negative regulation of type I interferon-mediated signaling pathway ADAR    adenosine deaminase, RNA-specific    -0.0001440556406552134
GO:0060339: negative regulation of type I interferon-mediated signaling pathway PTPN2   protein tyrosine phosphatase, non-receptor type 2    -0.0002988267418296442
GO:0061484: hematopoietic stem cell homeostasis ADAR    adenosine deaminase, RNA-specific       -0.0001399064478497578 7
GO:1900369: negative regulation of RNA interference     ADAR    adenosine deaminase, RNA-specific -0.0001399064478497578 7
GO:0006351: transcription, DNA-templated        ADIRF   adipogenesis regulatory factor  -0.0001675617209196798
GO:0006351: transcription, DNA-templated        AES     amino-terminal enhancer of split        0.0013577569053053603
GO:0006351: transcription, DNA-templated        APEX1   APEX nuclease (multifunctional DNA repair enzyme) 1  0.00020166743038709678
GO:0006351: transcription, DNA-templated        AR      androgen receptor       0.002646246582789125
GO:0006351: transcription, DNA-templated        ASCL1   achaete-scute family bHLH transcription factor 1     -0.0014379690025452749
GO:0006351: transcription, DNA-templated        ASF1A   anti-silencing function 1A histone chaperone 0.0010541688723845269
GO:0006351: transcription, DNA-templated        ATAD2   ATPase family, AAA domain containing 2  -4.68007668326296le-5
GO:0006351: transcription, DNA-templated        BCL11A  B-cell CLL/lymphoma 11A (zinc finger protein) -0.0009789705885269245
GO:0006351: transcription, DNA-templated        BIRC5   baculoviral IAP repeat containing 5     -0.0002607198766295170 5
GO:0006351: transcription, DNA-templated        BRD4    bromodomain containing 4        0.000554872295429083
GO:0006351: transcription, DNA-templated        CASP8AP2        caspase 8 associated protein 2  0.0012777215111830394
GO:0006351: transcription, DNA-templated        CCNC    cyclin C        0.0005696399025622202
GO:0006351: transcription, DNA-templated        CCND1   cyclin D1       -0.0026437378838172427
GO:0006351: transcription, DNA-templated        CDK8    cyclin-dependent kinase 8       0.002620616238663111
GO:0006351: transcription, DNA-templated        CDKN2A  cyclin-dependent kinase inhibitor 2A    0.0017650103453912
GO:0006351: transcription, DNA-templated        CDKN2B  cyclin-dependent kinase inhibitor 2B (p15, inhibits CDK4)    -0.0018417462822618968
GO:0006351: transcription, DNA-templated        CHAF1B  chromatin assembly factor 1, subunit B (p60) 0.006306137171951331
GO:0006351: transcription, DNA-templated        CHD3    chromodomain helicase DNA binding protein 3 0.00011887975520542764
GO:0006351: transcription, DNA-templated        CHMP1A  charged multivesicular body protein 1A  -0.0007825243721964978
GO:0006351: transcription, DNA-templated        CITED1  Cbp/p300-interacting transactivator, with Glu/Asp-rich carboxy-terminal domain, 1   0.002828390608548773
GO:0006351: transcription, DNA-templated        CTNNB1  catenin (cadherin-associated protein), beta 1, 88kDa       -0.00011719238664682533
GO:0006351: transcription, DNA-templated        DAXX    death-domain associated protein 0.0008952233162214325
GO:0006351: transcription, DNA-templated        DDX17   DEAD (Asp-Glu-Ala-Asp) box helicase 17  0.0017684564306945576
GO:0006351: transcription, DNA-templated        DNAJC2  DnaJ (Hsp40) homolog, subfamily C, member 2 -0.000986900211967874
GO:0006351: transcription, DNA-templated        E2F1    E2F transcription factor 1      0.002105069127653711
GO:0006351: transcription, DNA-templated        E2F4    E2F transcription factor 4, p107/p130-binding -0.0023533151198579854
GO:0006351: transcription, DNA-templated        E2F6    E2F transcription factor 6      0.0007202667271313004
GO:0006351: transcription, DNA-templated        E2F8    E2F transcription factor 8      0.0017162204414668022
GO:0006351: transcription, DNA-templated        E4F1    E4F transcription factor 1      0.0014855914032716618
GO:0006351: transcription, DNA-templated        ERBB4   v-erb-b2 avian erythroblastic leukemia viral oncogene homolog 4     -0.00018432545847019125
GO:0006351: transcription, DNA-templated        ESR1    estrogen receptor 1     -0.0009517868341633654
GO:0006351: transcription, DNA-templated        ETS2    v-ets avian erythroblastosis virus E26 oncogene homolog 2    -0.0010194748802169614
GO:0006351: transcription, DNA-templated        EYA1    EYA transcriptional coactivator and phosphatase 1    5.359143556317774e-5

GO:0006351: transcription, DNA-templated	EYA3	EYA transcriptional coactivator and phosphatase 3	0.0008393394739330924
GO:0006351: transcription, DNA-templated	EZH2	enhancer of zeste 2 polycomb repressive complex 2 subunit	-0.00013623578626891617
GO:0006351: transcription, DNA-templated	FOXM1	forkhead box M1 0.00019821275763450393
GO:0006351: transcription, DNA-templated	FOXO1	forkhead box O1 0.0018062678046832804
GO:0006351: transcription, DNA-templated	FOXO3	forkhead box O3 0.0012044867691304321
GO:0006351: transcription, DNA-templated	GATA2	GATA binding protein 2	-0.0004534173767054064
GO:0006351: transcription, DNA-templated	GFI1	growth factor independent 1 transcription repressor	0.0016694925462020502
GO:0006351: transcription, DNA-templated	GLI1	GLI family zinc finger 1	-0.0013007656246102237
GO:0006351: transcription, DNA-templated	HDAC2	histone deacetylase 2	-0.0012105156916769363
GO:0006351: transcription, DNA-templated	HELLS	helicase, lymphoid-specific	0.003475868617673022
GO:0006351: transcription, DNA-templated	HES1	hes family bHLH transcription factor 1	-0.0009141658046763733
GO:0006351: transcription, DNA-templated	HEXIM1	hexamethylene bis-acetamide inducible 1 0.00015327641706974976
GO:0006351: transcription, DNA-templated	HEY2	hes-related family bHLH transcription factor with YRPW motif 2	0.0027093792983095472
GO:0006351: transcription, DNA-templated	HINFP	histone H4 transcription factor 0.000995932531679305
GO:0006351: transcription, DNA-templated	HIPK2	homeodomain interacting protein kinase 2	0.0007731044956616214
GO:0006351: transcription, DNA-templated	HOXA3	homeobox A3	0.000991802762943586
GO:0006351: transcription, DNA-templated	HOXB1	homeobox B1	0.0036978126034359054
GO:0006351: transcription, DNA-templated	HOXB13	homeobox B13	0.0018316255175505562
GO:0006351: transcription, DNA-templated	HOXB2	homeobox B2	0.0027820391586367075
GO:0006351: transcription, DNA-templated	ID2	inhibitor of DNA binding 2, dominant negative helix-loop-helix protein	6.002927244346837e-5
GO:0006351: transcription, DNA-templated	IRF6	interferon regulatory factor 6	-0.001699513123758845
GO:0006351: transcription, DNA-templated	JMJD6	jumonji domain containing 6	0.0036023216885157773
GO:0006351: transcription, DNA-templated	KANK1	KN motif and ankyrin repeat domains 1	0.0025423350294964246
GO:0006351: transcription, DNA-templated	KLF2	Kruppel-like factor 2	-0.001235482982606846
GO:0006351: transcription, DNA-templated	KMT2B	lysine (K)-specific methyltransferase 2B	0.00046328569974643855
GO:0006351: transcription, DNA-templated	KMT2D	lysine (K)-specific methyltransferase 2D	-0.00020011455246932134
GO:0006351: transcription, DNA-templated	LHX6	LIM homeobox 6	0.0013492924793433738
GO:0006351: transcription, DNA-templated	MLLT10	myeloid/lymphoid or mixed-lineage leukemia (trithorax homolog, Drosophila); translocated to, 10	-0.0001638780188007765
GO:0006351: transcription, DNA-templated	MLXIP	MLX interacting protein 0.0027297553060307406
GO:0006351: transcription, DNA-templated	MYC	v-myc avian myelocytomatosis viral oncogene homolog	-0.0011399598187555918
GO:0006351: transcription, DNA-templated	NEUROD4	neuronal differentiation 4	0.001511503763484542
GO:0006351: transcription, DNA-templated	NOC2L	nucleolar complex associated 2 homolog (S. cerevisiae)	-0.0001514458693923486
GO:0006351: transcription, DNA-templated	NPAT	nuclear protein, ataxia-telangiectasia locus	-0.0007116665604177845
GO:0006351: transcription, DNA-templated	NRIP1	nuclear receptor interacting protein 1	0.0010742903363907153
GO:0006351: transcription, DNA-templated	PADI4	peptidyl arginine deiminase, type IV	-0.0008298530598771132
GO:0006351: transcription, DNA-templated	PAX2	paired box 2	-0.0016200068037620424
GO:0006351: transcription, DNA-templated	PAX8	paired box 8	0.0009506601049226415
GO:0006351: transcription, DNA-templated	PER2	period circadian clock 2	0.0012424599014061403
GO:0006351: transcription, DNA-templated	PHB2	prohibitin 2	-0.0007384633744968913
GO:0006351: transcription, DNA-templated	PKN1	protein kinase N1	-0.0019094981210912743
GO:0006351: transcription, DNA-templated	PML	promyelocytic leukemia	-0.0006815997443380531
GO:0006351: transcription, DNA-templated	PPP1R13L	protein phosphatase 1, regulatory subunit 13 like	-0.0014789465380928755
GO:0006351: transcription, DNA-templated	PROX1	prospero homeobox 1	0.0011257513484109628
GO:0006351: transcription, DNA-templated	PURA	purine-rich element binding protein A	-0.0017128735392898028
GO:0006351: transcription, DNA-templated	RB1	retinoblastoma 1	-0.0014992189923968755
GO:0006351: transcription, DNA-templated	RBBP4	retinoblastoma binding protein 4	-0.0018653430843826367
GO:0006351: transcription, DNA-templated	RFC1	replication factor C (activator 1) 1, 145kDa	0.0010631081672241947
GO:0006351: transcription, DNA-templated	SALL1	spalt-like transcription factor 1	-0.00

2447497162819614
GO:0006351: transcription, DNA-templated SERPINE1 serpin peptidase inhibitor, clade E (nexin, plasminogen activator inhibitor type 1), member 1 0.00011211078648559636
GO:0006351: transcription, DNA-templated SIRT1 sirtuin 1 -1.1689012488519055e-6
GO:0006351: transcription, DNA-templated SIRT2 sirtuin 2 -0.0008553464471506225
GO:0006351: transcription, DNA-templated SKI SKI proto-oncogene -0.000672465064142849 9
GO:0006351: transcription, DNA-templated SMARCD3 SWI/SNF related, matrix associated, actin dependent regulator of chromatin, subfamily d, member 3 0.00034685432064183335
GO:0006351: transcription, DNA-templated SPDEF SAM pointed domain containing ETS transcription factor 0.003146651632048523
GO:0006351: transcription, DNA-templated STAT5A signal transducer and activator of transcription 5A 0.001601556676844244
GO:0006351: transcription, DNA-templated TBX21 T-box 21 0.00039949044251085696
GO:0006351: transcription, DNA-templated TBX3 T-box 3 0.0012296117248930631
GO:0006351: transcription, DNA-templated TCF15 transcription factor 15 (basic helix-loop-helix) -0.002663303608079843
GO:0006351: transcription, DNA-templated TCF7L2 transcription factor 7-like 2 (T-cell specific, HMG-box) 0.000576051457913259
GO:0006351: transcription, DNA-templated TFCP2L1 transcription factor CP2-like 1 0.0004506590126016773
GO:0006351: transcription, DNA-templated TFDP1 transcription factor Dp-1 0.0014509939541695938
GO:0006351: transcription, DNA-templated THRB thyroid hormone receptor, beta 0.0019501091583035255
GO:0006351: transcription, DNA-templated TWIST1 twist family bHLH transcription factor 1 -0.0013467142954703033
GO:0006351: transcription, DNA-templated TXLNG taxilin gamma -0.0008396131646636599
GO:0006351: transcription, DNA-templated VPS72 vacuolar protein sorting 72 homolog (S. cerevisiae) 0.0005604743224801479
GO:0006351: transcription, DNA-templated WWTR1 WW domain containing transcription regulator 1 0.0009044947774101575
GO:0006351: transcription, DNA-templated YY1 YY1 transcription factor 0.0013454257525090118
GO:0006351: transcription, DNA-templated ZBTB18 zinc finger and BTB domain containing 18 0.0012760552569729337
GO:0006351: transcription, DNA-templated ZNF205 zinc finger protein 205 0.0005971545050382172
GO:0006351: transcription, DNA-templated ZNF24 zinc finger protein 24 0.001295675804133213
GO:0006351: transcription, DNA-templated ZNF266 zinc finger protein 266 0.0004135283287241028
GO:0006351: transcription, DNA-templated ZNF335 zinc finger protein 335 -0.000365896095415836
GO:0006351: transcription, DNA-templated ZNF442 zinc finger protein 442 0.0010621231350483454
GO:0030154: cell differentiation ADIRF adipogenesis regulatory factor -0.00016681887194860251
GO:0030154: cell differentiation AKT1 v-akt murine thymoma viral oncogene homolog 1 0.000730676154289761
GO:0030154: cell differentiation ARHGAP24 Rho GTPase activating protein 24 -0.0009343006482775348
GO:0030154: cell differentiation AXIN1 axin 1 -0.0007333463540473516
GO:0030154: cell differentiation CPLX2 complexin 2 0.002031753329072328
GO:0030154: cell differentiation EDA ectodysplasin A -0.0008146890664469235
GO:0030154: cell differentiation EDAR ectodysplasin A receptor 0.0005936239813974121
GO:0030154: cell differentiation EGFL6 EGF-like-domain, multiple 6 0.0001366426792905654
GO:0030154: cell differentiation ELF5 E74-like factor 5 (ets domain transcription factor) 0.0011718001958474474
GO:0030154: cell differentiation ETS2 v-ets avian erythroblastosis virus E26 oncogene homolog 2 -0.001015585100172126
GO:0030154: cell differentiation ETV4 ets variant 4 0.00019503508883322835
GO:0030154: cell differentiation FOXA1 forkhead box A1 2.7263894893049837e-5
GO:0030154: cell differentiation FOXA2 forkhead box A2 -1.6605610178417844e-5
GO:0030154: cell differentiation FOXC1 forkhead box C1 -2.185149261822637e-5
GO:0030154: cell differentiation FOXC2 forkhead box C2 (MFH-1, mesenchyme forkhead 1) 0.0017480670298103472
GO:0030154: cell differentiation FOXE3 forkhead box E3 0.0010507447838200954
GO:0030154: cell differentiation FOXH1 forkhead box H1 -0.0013714256138066298
GO:0030154: cell differentiation FOXM1 forkhead box M1 0.00019748677887642297
GO:0030154: cell differentiation FOXO1 forkhead box O1 0.0018001303307412367
GO:0030154: cell differentiation FOXO3 forkhead box O3 0.0012001196054617594
GO:0030154: cell differentiation GLRX2 glutaredoxin 2 0.0011160844527823416
GO:0030154: cell differentiation INHA inhibin, alpha 0.00021147559013901275
GO:0030154: cell differentiation INHBA inhibin, beta A -0.0013534974399642865
GO:0030154: cell differentiation JAG2 jagged 2 -5.620467313112927e-6
GO:0030154: cell differentiation JAK2 Janus kinase 2 -3.152112797203175e-5
GO:0030154: cell differentiation KIF2A kinesin heavy chain member 2A -0.0007843984089759799
GO:0030154: cell differentiation MDK midkine (neurite growth-promoting factor 2) 0.0016657587701324876
GO:0030154: cell differentiation NDRG2 NDRG family member 2 0.0021741396950714755
GO:0030154: cell differentiation NELL1 NEL-like 1 (chicken) 0.0015329098517756217
GO:0030154: cell differentiation PURA purine-rich element binding protein A -0.0001705929206954963
GO:0030154: cell differentiation SLC7A5 solute carrier family 7 (amino acid transporter light

chain, L system), member 5	-0.002574075147499151
GO:0030154: cell differentiation	SPDEF	SAM pointed domain containing ETS transcription facto
r	0.003136377589828832
GO:0030154: cell differentiation	SRD5A2	steroid-5-alpha-reductase, alpha polypeptide 2 (3-oxo
-5 alpha-steroid delta 4-dehydrogenase alpha 2) -0.0006432281117309362
GO:0030154: cell differentiation	TP53	tumor protein p53	0.0011759806987857292
GO:0030154: cell differentiation	YY1	YY1 transcription factor	0.0013402815459465903
GO:0045600: positive regulation of fat cell differentiation	ADIRF	adipogenesis regulatory facto
r	-0.00016418273076696812
GO:0045600: positive regulation of fat cell differentiation	AKT1	v-akt murine thymoma viral on
cogene homolog 1	0.0007232830270269223
GO:0045600: positive regulation of fat cell differentiation	CEBPB	CCAAT/enhancer binding protei
n (C/EBP), beta -0.0002710629310775028
GO:0045600: positive regulation of fat cell differentiation	CREB1	cAMP responsive element bindi
ng protein 1	0.0006504646253352424
GO:0045600: positive regulation of fat cell differentiation	HTR2C	5-hydroxytryptamine (serotoni
n) receptor 2C, G protein-coupled	-0.0005203800227167744
GO:0045600: positive regulation of fat cell differentiation	ID2	inhibitor of DNA binding 2, d
ominant negative helix-loop-helix protein	6.0352946227578056e-5
GO:0045600: positive regulation of fat cell differentiation	LRP5	low density lipoprotein recep
tor-related protein 5	3.132876831481811e-5
GO:0045600: positive regulation of fat cell differentiation	SAV1	salvador family WW domain con
taining protein 1	-0.0023207872964762953
GO:0045600: positive regulation of fat cell differentiation	SFRP1	secreted frizzled-related pro
tein 1	0.0012656508995616943
GO:0045600: positive regulation of fat cell differentiation	XBP1	X-box binding protein 1 0.000
26508061877325773
GO:0071478: cellular response to radiation	ADIRF	adipogenesis regulatory factor	-0.0001435353
193695775
GO:0072719: cellular response to cisplatin	ADIRF	adipogenesis regulatory factor	-0.0001633631
8119645004
GO:0072719: cellular response to cisplatin	RAD51	RAD51 recombinase	-0.001848593006405684
GO:0072719: cellular response to cisplatin	SLC31A1 solute carrier family 31 (copper transporte
r), member 1	-2.758633427670951e-5
GO:2001023: regulation of response to drug	ADIRF	adipogenesis regulatory factor	-0.0001648556
9311122276
GO:2001023: regulation of response to drug	GABRA4	gamma-aminobutyric acid (GABA) A receptor, al
pha 4	-0.0014745183714208066
GO:0001570: vasculogenesis	ADM	adrenomedullin	0.0022434078885771925
GO:0001570: vasculogenesis	CAV1	caveolin 1, caveolae protein, 22kDa	-0.000536389671852577
2
GO:0001570: vasculogenesis	CITED1	Cbp/p300-interacting transactivator, with Glu/Asp-rich carbox
y-terminal domain, 1	0.00282634847978138
GO:0001570: vasculogenesis	CUL7	cullin 7	-0.0001196282552138675
GO:0001570: vasculogenesis	ENG	endoglin	0.0008293706626310097
GO:0001570: vasculogenesis	FOXM1	forkhead box M1 0.00019753488647465217
GO:0001570: vasculogenesis	HEY2	hes-related family bHLH transcription factor with YRPW motif
2	0.0027069690513191087
GO:0001570: vasculogenesis	PITX2	paired-like homeodomain 2	0.0021820487507270485
GO:0001570: vasculogenesis	PTPRJ	protein tyrosine phosphatase, receptor type, J	-0.0003391328
524076697
GO:0001570: vasculogenesis	RASA1	RAS p21 protein activator (GTPase activating protein) 1 -0.00
03499982278031426
GO:0001570: vasculogenesis	SHH	sonic hedgehog	0.0006020355406372963
GO:0001570: vasculogenesis	SMO	smoothened, frizzled class receptor	0.002144905640206822
GO:0001570: vasculogenesis	SOX18	SRY (sex determining region Y)-box 18	0.0012914910099911382
GO:0001570: vasculogenesis	VEGFA	vascular endothelial growth factor A	0.0005968811974446955
GO:0001570: vasculogenesis	WT1	Wilms tumor 1	-0.0005086154486246667
GO:0001570: vasculogenesis	YAP1	Yes-associated protein 1	-0.00016470097204287582
GO:0001843: neural tube closure ADM	adrenomedullin	0.0022446428940379594
GO:0001843: neural tube closure ALX1	ALX homeobox 1	0.0022988626457801633
GO:0001843: neural tube closure BBS4	Bardet-Biedl syndrome 4 -0.0005105909852871569
GO:0001843: neural tube closure BMP4	bone morphogenetic protein 4	-0.00032355394167502964
GO:0001843: neural tube closure FZD3	frizzled class receptor 3	0.0003804666054673862
GO:0001843: neural tube closure GRHL2	grainyhead-like 2 (Drosophila)	0.001009523584328426
GO:0001843: neural tube closure IFT122	intraflagellar transport 122 homolog (Chlamydomonas)	-0.00
16152546694120262
GO:0001843: neural tube closure LIAS	lipoic acid synthetase	-0.0005556733559023034
GO:0001843: neural tube closure LMO4	LIM domain only 4	0.0021179655193401233
GO:0001843: neural tube closure LRP6	low density lipoprotein receptor-related protein 6	0.000
14996070998739782
GO:0001843: neural tube closure MTHFD1	methylenetetrahydrofolate dehydrogenase (NADP+ dependent) 1,
methenyltetrahydrofolate cyclohydrolase, formyltetrahydrofolate synthetase	-0.000665730350697600
2
GO:0001843: neural tube closure PAX2	paired box 2	-0.0016195239316690046
GO:0001843: neural tube closure PAX3	paired box 3	-0.0029541363399052762
GO:0001843: neural tube closure PHACTR4 phosphatase and actin regulator 4	-0.001796271394250020
8
GO:0001843: neural tube closure PTCH1	patched 1	-6.577755257534153e-5
GO:0001843: neural tube closure RARG	retinoic acid receptor, gamma	-0.0023266427241055336
GO:0001843: neural tube closure SALL1	spalt-like transcription factor 1	-0.002446828641577958

6
GO:0001843: neural tube closure SKI SKI proto-oncogene -0.0006723450428151536
GO:0001843: neural tube closure TULP3 tubby like protein 3 0.0009591934635432486
GO:0001843: neural tube closure TWIST1 twist family bHLH transcription factor 1 -0.0013463727
26287963
GO:0002026: regulation of the force of heart contraction ADM adrenomedullin 0.00222994670
94114254
GO:0002026: regulation of the force of heart contraction IFNG interferon, gamma -4.91
7862048942021e-5
GO:0002026: regulation of the force of heart contraction PRKCA protein kinase C, alpha -6.13
0061387185221e-6
GO:0002031: G-protein coupled receptor internalization ADM adrenomedullin 0.002241989129939734
GO:0002031: G-protein coupled receptor internalization DRD2 dopamine receptor D2 -0.0002343534
6128204245
GO:0006171: cAMP biosynthetic process ADM adrenomedullin 0.002257456012228351
GO:0006701: progesterone biosynthetic process ADM adrenomedullin 0.002247342140914187
GO:0006701: progesterone biosynthetic process STAR steroidogenic acute regulatory protein 0.000
8989063754188108
GO:0007204: positive regulation of cytosolic calcium ion concentration ADM adrenomedullin 0.002
23819578503534
GO:0007204: positive regulation of cytosolic calcium ion concentration AGTR1 angiotensin II recept
or, type 1 0.0003469501432738616
GO:0007204: positive regulation of cytosolic calcium ion concentration AVP arginine vasopressin
-0.0009377923473621333
GO:0007204: positive regulation of cytosolic calcium ion concentration CD24 CD24 molecule 0.001
072304631472206
GO:0007204: positive regulation of cytosolic calcium ion concentration CXCL13 chemokine (C-X-C moti
f) ligand 13 0.002993858781088234З
GO:0007204: positive regulation of cytosolic calcium ion concentration CXCR4 chemokine (C-X-C moti
f) receptor 4 0.0007952823600648638
GO:0007204: positive regulation of cytosolic calcium ion concentration ESR1 estrogen receptor 1
-0.0009489711330357443
GO:0007204: positive regulation of cytosolic calcium ion concentration GATA2 GATA binding protein
2 -0.00045219039679756696
GO:0007204: positive regulation of cytosolic calcium ion concentration GJA1 gap junction protein,
alpha 1, 43kDa -0.00016247681117484476
GO:0007204: positive regulation of cytosolic calcium ion concentration HMGB1 high mobility group b
ox 1 -0.0007759442251216442
GO:0007204: positive regulation of cytosolic calcium ion concentration JAK2 Janus kinase 2 -3.13
9059991722578e-5
GO:0007267: cell-cell signaling ADM adrenomedullin 0.0022421583587430224
GO:0007267: cell-cell signaling AGT angiotensinogen (serpin peptidase inhibitor, clade A, member
8) -0.0011363402136848192
GO:0007267: cell-cell signaling AR androgen receptor 0.002643287950150788
GO:0007267: cell-cell signaling AREG amphiregulin 0.002462287461729004
GO:0007267: cell-cell signaling AVP arginine vasopressin -0.0009395884492577825
GO:0007267: cell-cell signaling C1QA complement component 1, q subcomponent, A chain 0.00054351884
46500535
GO:0007267: cell-cell signaling CCL7 chemokine (C-C motif) ligand 7 -0.0024068730976171135
GO:0007267: cell-cell signaling CCL8 chemokine (C-C motif) ligand 8 -0.0005913692449657669
GO:0007267: cell-cell signaling CXCL10 chemokine (C-X-C motif) ligand 10 6.27737423453633e-5
GO:0007267: cell-cell signaling CXCL13 chemokine (C-X-C motif) ligand 13 0.0029995297624913412
GO:0007267: cell-cell signaling EFNA1 ephrin-A1 -0.0005215606381449765
GO:0007267: cell-cell signaling FGF3 fibroblast growth factor 3 0.0015642588656650518
GO:0007267: cell-cell signaling FGF5 fibroblast growth factor 5 -0.001096136656745229
GO:0007267: cell-cell signaling FGFR2 fibroblast growth factor receptor 2 0.000764917678410162
GO:0007267: cell-cell signaling GATA4 GATA binding protein 4 -0.0010970487169320359
GO:0007267: cell-cell signaling GJA1 gap junction protein, alpha 1, 43kDa -0.00016270092097271l
26
GO:0007267: cell-cell signaling GRB2 growth factor receptor-bound protein 2 0.0004305622912941412
6
GO:0007267: cell-cell signaling GREM1 gremlin 1, DAN family BMP antagonist -0.000829799277102856
6
GO:0007267: cell-cell signaling IFNA2 interferon, alpha 2 -0.001720762184470687
GO:0007267: cell-cell signaling IHH indian hedgehog -0.0020450571847204346
GO:0007267: cell-cell signaling IL7 interleukin 7 0.0008733124452919575
GO:0007267: cell-cell signaling INHA inhibin, alpha 0.00021174228440315048
GO:0007267: cell-cell signaling INHBA inhibin, beta A -0.0013570390310194817
GO:0007267: cell-cell signaling LHX1 LIM homeobox 1 -0.0007604609081624374
GO:0007267: cell-cell signaling NRP1 neuropilin 1 -0.000644826869648602
GO:0007267: cell-cell signaling PGR progesterone receptor 0.00035247299751778565
GO:0007267: cell-cell signaling SHH sonic hedgehog 0.0006015980003569587
GO:0007267: cell-cell signaling SRD5A2 steroid-5-alpha-reductase, alpha polypeptide 2 (3-oxo-5 alpha
-steroid delta 4-dehydrogenase alpha 2) -0.0006447623364778302
GO:0007267: cell-cell signaling TFAP2C transcription factor AP-2 gamma (activating enhancer binding
protein 2 gamma) 0.00106335307409465
GO:0007267: cell-cell signaling TGFB2 transforming growth factor, beta 2 -0.001061919689039696
5
GO:0007267: cell-cell signaling TNFRSF11A tumor necrosis factor receptor superfamily, member 11
a, NFKB activator 0.0027819203611569674
GO:0007267: cell-cell signaling WNT1 wingless-type MMTV integration site family, member 1 0.000

78846941802616
GO:0007507: heart development   ADM     adrenomedullin  0.0022403976541209695
GO:0007507: heart development   CRIP1   cysteine-rich protein 1 (intestinal)    0.0017234078132703317
GO:0007507: heart development   CXADR   coxsackie virus and adenovirus receptor 0.0004359879102501653
GO:0007507: heart development   ERBB4   v-erb-b2 avian erythroblastic leukemia viral oncogene homolog 4       -0.0001836264204598173
GO:0007507: heart development   FOXC1   forkhead box C1 -2.1815688535006127e-5
GO:0007507: heart development   FOXC2   forkhead box C2 (MFH-1, mesenchyme forkhead 1)  0.0017500887252007928
GO:0007507: heart development   GATA2   GATA binding protein 2  -0.0004523452001406851
GO:0007507: heart development   GATA3   GATA binding protein 3  -3.9466253800904126e-5
GO:0007507: heart development   GJA1    gap junction protein, alpha 1, 43kDa    -0.0001624874359230264
GO:0007507: heart development   GLI2    GLI family zinc finger 2        0.0018556407034636836
GO:0007507: heart development   GLI3    GLI family zinc finger 3        -0.0021545456124907633
GO:0007507: heart development   HEXIM1  hexamethylene bis-acetamide inducible 1 0.0001527033173223713
GO:0007507: heart development   ITGA3   integrin, alpha 3 (antigen CD49C, alpha 3 subunit of VLA-3 receptor) 0.0014011512107650484
GO:0007507: heart development   JMJD6   jumonji domain containing 6     0.0035947509921381436
GO:0007507: heart development   MEF2C   myocyte enhancer factor 2C      0.0009746357047119457
GO:0007507: heart development   MTHFD1  methylenetetrahydrofolate dehydrogenase (NADP+ dependent) 1, methenyltetrahydrofolate cyclohydrolase, formyltetrahydrofolate synthetase     -0.0006645436796725466
GO:0007507: heart development   NOTCH1  notch 1 0.0005195080009200783
GO:0007507: heart development   PAX3    paired box 3    -0.0029484446967786584
GO:0007507: heart development   PCNA    proliferating cell nuclear antigen      0.001228039162480632
GO:0007507: heart development   PRKDC   protein kinase, DNA-activated, catalytic polypeptide    -0.00185744841481684
GO:0007507: heart development   PTEN    phosphatase and tensin homolog  1.677183192616889e-5
GO:0007507: heart development   PTPRJ   protein tyrosine phosphatase, receptor type, J  -0.0003387748230284296
GO:0007507: heart development   RAF1    Raf-1 proto-oncogene, serine/threonine kinase   0.0015005038199883132
GO:0007507: heart development   SALL1   spalt-like transcription factor 1       -0.0024407948070585936
GO:0007507: heart development   SHH     sonic hedgehog  0.0006008317048758139
GO:0007507: heart development   SOX4    SRY (sex determining region Y)-box 4    -3.797852553967947e-5
GO:0007507: heart development   STRA6   stimulated by retinoic acid 6   -0.00168244616709004
GO:0007507: heart development   TGFB2   transforming growth factor, beta 2      -0.0010607370183366232
GO:0007507: heart development   TGFBR1  transforming growth factor, beta receptor 1     0.0003404305818267974
GO:0007507: heart development   TH      tyrosine hydroxylase    -0.00034235394491181665
GO:0007507: heart development   TRPS1   trichorhinophalangeal syndrome I        -0.0001783128055240674
GO:0007507: heart development   VLDLR   very low density lipoprotein receptor   0.0009544257474276615
GO:0007507: heart development   WT1     Wilms tumor 1   -0.0005075979069531801
GO:0007565: female pregnancy    ADM     adrenomedullin  0.0022416185676936698
GO:0007565: female pregnancy    AGT     angiotensinogen (serpin peptidase inhibitor, clade A, member 8)     -0.0011362942712785191
GO:0007565: female pregnancy    BCL2    B-cell CLL/lymphoma 2   -4.9559769533562945e-6
GO:0007565: female pregnancy    EPN1    epsin 1 -0.0005480359649646314
GO:0007565: female pregnancy    IDO1    indoleamine 2,3-dioxygenase 1   0.0012283431506251445
GO:0007565: female pregnancy    IGFBP2  insulin-like growth factor binding protein 2, 36kDa     0.00014538508354895372
GO:0007565: female pregnancy    IL4     interleukin 4   0.0002594868369637104
GO:0007565: female pregnancy    LEP     leptin  0.003197561759565936
GO:0007565: female pregnancy    LGALS9  lectin, galactoside-binding, soluble, 9 -0.00011664718794397956
GO:0007565: female pregnancy    MAPT    microtubule-associated protein tau      0.0015532975118281052
GO:0007565: female pregnancy    PSG4    pregnancy specific beta-1-glycoprotein 4        -0.0010328661238374207
GO:0007565: female pregnancy    STAT5A  signal transducer and activator of transcription 5A     0.0015989561575371672
GO:0007565: female pregnancy    TFCP2L1 transcription factor CP2-like 1 0.000449947406450043
GO:0007565: female pregnancy    TGFB1   transforming growth factor, beta 1      -7.329792419287217e-5
GO:0007565: female pregnancy    TGFB3   transforming growth factor, beta 3      -0.001827661027896623
GO:0007568: aging       ADM     adrenomedullin  0.002243579445825092
GO:0007568: aging       AGT     angiotensinogen (serpin peptidase inhibitor, clade A, member 8) -0.0011377192141996976
GO:0007568: aging       AKT1    v-akt murine thymoma viral oncogene homolog 1   0.0007328830645346385
GO:0007568: aging       ALOX12  arachidonate 12-lipoxygenase    -0.0019589806508740013
GO:0007568: aging       APOD    apolipoprotein D        0.0026352863126499508
GO:0007568: aging       ASS1    argininosuccinate synthase 1    0.0001439403730751679З
GO:0007568: aging       BAK1    BCL2-antagonist/killer 1        -0.001880879756494619
GO:0007568: aging       CAST    calpastatin     -0.003041868365457816
GO:0007568: aging       CCL2    chemokine (C-C motif) ligand 2  0.0008189559148773358
GO:0007568: aging       CDKN2B  cyclin-dependent kinase inhibitor 2B (p15, inhibits CDK4)       -0.0018402570942091761
GO:0007568: aging       CRYAB   crystallin, alpha B     0.0009879638542821013
GO:0007568: aging       CX3CL1  chemokine (C-X3-C motif) ligand 1       0.002123769647916821

GO:0007568: aging       GLRX2   glutaredoxin 2  0.001118792896039328
GO:0007568: aging       GPX1    glutathione peroxidase 1        0.000387701507454065
GO:0007568: aging       GRB2    growth factor receptor-bound protein 2  0.00043072339321570776
GO:0007568: aging       IGFBP2  insulin-like growth factor binding protein 2, 36kDa     0.00014553747
74046118
GO:0007568: aging       IRAK1   interleukin-1 receptor-associated kinase 1      -0.001613150040502562
GO:0007568: aging       KCNE2   potassium voltage-gated channel, Isk-related family, member 2   -0.00
05766565756590681
GO:0007568: aging       KRT33B  keratin 33B     0.0009447727301874563
GO:0007568: aging       MMP7    matrix metallopeptidase 7 (matrilysin, uterine) 0.0004263936853100033
GO:0007568: aging       PAX2    paired box 2    -0.0016188877523914803
GO:0007568: aging       PTEN    phosphatase and tensin homolog  1.6239143543625582e-5
GO:0007568: aging       RXRA    retinoid X receptor, alpha      0.0011178851661669657
GO:0007568: aging       TFRC    transferrin receptor    0.0010869358788891505
GO:0007568: aging       TGFB1   transforming growth factor, beta 1      -7.344736873760857e-5
GO:0007568: aging       TGFB3   transforming growth factor, beta 3      -0.00182961432299378
GO:0007568: aging       TYMS    thymidylate synthetase  0.0015670155233409742
GO:0008209: androgen metabolic process  ADM     adrenomedullin  0.0022357531165483374
GO:0008209: androgen metabolic process  ESR1    estrogen receptor 1     -0.0009473512583995904
GO:0008209: androgen metabolic process  HSD17B4 hydroxysteroid (17-beta) dehydrogenase 4        -0.00
11616394572610347
GO:0008209: androgen metabolic process  SHH     sonic hedgehog  0.0005990696765356112
GO:0008209: androgen metabolic process  SRD5A2  steroid-5-alpha-reductase, alpha polypeptide 2 (3-oxo
-5 alpha-steroid delta 4-dehydrogenase alpha 2) -0.0006422839767221325
GO:0008284: positive regulation of cell proliferation   ADM     adrenomedullin  0.0022410950993080577
GO:0008284: positive regulation of cell proliferation   ALDH1A2 aldehyde dehydrogenase 1 family, memb
er A2   -0.0030233719864110917
GO:0008284: positive regulation of cell proliferation   ALOX12  arachidonate 12-lipoxygenase    -0.00
1956588532168183
GO:0008284: positive regulation of cell proliferation   AR      androgen receptor       0.00264123026
72169136
GO:0008284: positive regulation of cell proliferation   AREG    amphiregulin    0.0024605282476191067
GO:0008284: positive regulation of cell proliferation   AVP     arginine vasopressin    -0.0009391515
786076601
GO:0008284: positive regulation of cell proliferation   BIRC5   baculoviral IAP repeat containing 5
-0.0002602362256852859
GO:0008284: positive regulation of cell proliferation   CDC20   cell division cycle 20  -0.0001883575
1784495341
GO:0008284: positive regulation of cell proliferation   CDC7    cell division cycle 7   -0.0006803484
2796161
GO:0008284: positive regulation of cell proliferation   CIB1    calcium and integrin binding 1 (calmy
rin)    5.403418133617768e-5
GO:0008284: positive regulation of cell proliferation   CST3    cystatin C      -7.09271232806805e-5
GO:0008284: positive regulation of cell proliferation   CXCL10  chemokine (C-X-C motif) ligand 10
6.311418221515931e-5
GO:0008284: positive regulation of cell proliferation   E2F3    E2F transcription factor 3      0.002
1651822739979297
GO:0008284: positive regulation of cell proliferation   EGFR    epidermal growth factor receptor
0.0006873945507094657
GO:0008284: positive regulation of cell proliferation   ERBB4   v-erb-b2 avian erythroblastic leukemi
a viral oncogene homolog 4      -0.00018389225564362325
GO:0008284: positive regulation of cell proliferation   FGF3    fibroblast growth factor 3      0.001
563306565160581
GO:0008284: positive regulation of cell proliferation   FGF5    fibroblast growth factor 5      -0.00
10955850081850228
GO:0008284: positive regulation of cell proliferation   FGF7    fibroblast growth factor 7      0.000
6345221184297735
GO:0008284: positive regulation of cell proliferation   FGF8    fibroblast growth factor 8 (androgen-
induced)        0.0009857303561165117
GO:0008284: positive regulation of cell proliferation   FGFR2   fibroblast growth factor receptor 2
0.0007645573325409214
GO:0008284: positive regulation of cell proliferation   FGFR3   fibroblast growth factor receptor 3
0.00022307374408331033
GO:0008284: positive regulation of cell proliferation   FLT3    fms-related tyrosine kinase 3   -0.00
06828990140541774
GO:0008284: positive regulation of cell proliferation   FOXM1   forkhead box M1 0.0001979683675731128
4
GO:0008284: positive regulation of cell proliferation   FZR1    fizzy/cell division cycle 20 related
 1 (Drosophila) 0.0009496567670515858
GO:0008284: positive regulation of cell proliferation   GCNT2   glucosaminyl (N-acetyl) transferase
 2, I-branching enzyme (I blood group)  0.0024357318828497183
GO:0008284: positive regulation of cell proliferation   GDNF    glial cell derived neurotrophic facto
r       0.00044520541170774364
GO:0008284: positive regulation of cell proliferation   GLI1    GLI family zinc finger 1        -0.00
12985550423469132
GO:0008284: positive regulation of cell proliferation   GREM1   gremlin 1, DAN family BMP antagonist
-0.0008293204466775923
GO:0008284: positive regulation of cell proliferation   HDAC2   histone deacetylase 2   -0.0012082371
845297463
GO:0008284: positive regulation of cell proliferation   HES1    hes family bHLH transcription factor
 1      -0.0009123981761961904

GO:0008284: positive regulation of cell proliferation   HIPK2   homeodomain interacting protein kinas
e 2     0.0007717085127869681
GO:0008284: positive regulation of cell proliferation   HOXA3   homeobox A3     0.0009900277491969958
GO:0008284: positive regulation of cell proliferation   IFNG    interferon, gamma       -4.8874569978
34808e-5
GO:0008284: positive regulation of cell proliferation   IGF1    insulin-like growth factor 1 (somatom
edin C) 0.0001313531632458009
GO:0008284: positive regulation of cell proliferation   IGF1R   insulin-like growth factor 1 receptor
0.0010728707701114512
GO:0008284: positive regulation of cell proliferation   IL6ST   interleukin 6 signal transducer 0.001
8777446167258076
GO:0008284: positive regulation of cell proliferation   IL7     interleukin 7   0.0008729160679010597
GO:0008284: positive regulation of cell proliferation   INSR    insulin receptor        -0.0013654856
869018532
GO:0008284: positive regulation of cell proliferation   ISL1    ISL LIM homeobox 1      7.86717480732
209e-5
GO:0008284: positive regulation of cell proliferation   ITGB1   integrin, beta 1 (fibronectin recepto
r, beta polypeptide, antigen CD29 includes MDF2, MSK12) 0.0024307867666587537
GO:0008284: positive regulation of cell proliferation   JAK2    Janus kinase 2  -3.1338239867384334e-
5
GO:0008284: positive regulation of cell proliferation   KIF14   kinesin family member 14        0.000
4869357289194409
GO:0008284: positive regulation of cell proliferation   KIT     v-kit Hardy-Zuckerman 4 feline sarcom
a viral oncogene homolog        0.00027607878718469327
GO:0008284: positive regulation of cell proliferation   KMT2D   lysine (K)-specific methyltransferase
2D      -0.0001997727000891164
GO:0008284: positive regulation of cell proliferation   LEF1    lymphoid enhancer-binding factor 1
-0.00010029784314462719
GO:0008284: positive regulation of cell proliferation   LEP     leptin  0.0031963665156191177
GO:0008284: positive regulation of cell proliferation   LRP5    low density lipoprotein receptor-rela
ted protein 5   3.0206857642998332e-5
GO:0008284: positive regulation of cell proliferation   LYN     LYN proto-oncogene, Src family tyrosi
ne kinase       -0.0014224121197098166
GO:0008284: positive regulation of cell proliferation   MAS1    MAS1 proto-oncogene, G protein-couple
d receptor      0.00029116216256672183
GO:0008284: positive regulation of cell proliferation   MFGE8   milk fat globule-EGF factor 8 protein
0.0018078374057839495
GO:0008284: positive regulation of cell proliferation   MYC     v-myc avian myelocytomatosis viral on
cogene homolog  -0.0011376324669968025
GO:0008284: positive regulation of cell proliferation   NOTCH1  notch 1 0.0005197075762901456
GO:0008284: positive regulation of cell proliferation   ODC1    ornithine decarboxylase 1       0.000
8268257255228956
GO:0008284: positive regulation of cell proliferation   PAX3    paired box 3    -0.002949347521846710
5
GO:0008284: positive regulation of cell proliferation   PDGFRB  platelet-derived growth factor recept
or, beta polypeptide    -0.0003624394164056994
GO:0008284: positive regulation of cell proliferation   PDX1    pancreatic and duodenal homeobox 1
0.0002579178362100724
GO:0008284: positive regulation of cell proliferation   POU1F1  POU class 1 homeobox 1  0.00013769257
418485355
GO:0008284: positive regulation of cell proliferation   PRC1    protein regulator of cytokinesis 1
0.0012311469731749358
GO:0008284: positive regulation of cell proliferation   PRKCZ   protein kinase C, zeta  -0.0016029215
175104086
GO:0008284: positive regulation of cell proliferation   PROX1   prospero homeobox 1     0.00112341641
77144777
GO:0008284: positive regulation of cell proliferation   PTEN    phosphatase and tensin homolog  1.658
078482338241e-5
GO:0008284: positive regulation of cell proliferation   PTN     pleiotrophin    0.0003027089957155453
4
GO:0008284: positive regulation of cell proliferation   PURA    purine-rich element binding protein A
-0.00017096419545517539
GO:0008284: positive regulation of cell proliferation   RARG    retinoic acid receptor, gamma   -0.00
2322808048203535
GO:0008284: positive regulation of cell proliferation   RNF187  ring finger protein 187 -0.0022298669
757898812
GO:0008284: positive regulation of cell proliferation   RPA1    replication protein A1, 70kDa   0.000
44064174226367904
GO:0008284: positive regulation of cell proliferation   S100B   S100 calcium binding protein B  0.005
193531837173987
GO:0008284: positive regulation of cell proliferation   SFRP1   secreted frizzled-related protein 1
0.0012851051968391038
GO:0008284: positive regulation of cell proliferation   SHH     sonic hedgehog  0.0006011691842453295
GO:0008284: positive regulation of cell proliferation   SHMT2   serine hydroxymethyltransferase 2 (mi
tochondrial)    -0.0010984777529084396
GO:0008284: positive regulation of cell proliferation   SIRT1   sirtuin 1       -1.3705262127162033e-
6
GO:0008284: positive regulation of cell proliferation   SOX11   SRY (sex determining region Y)-box 11
-0.00020882667022459914
GO:0008284: positive regulation of cell proliferation   SOX4    SRY (sex determining region Y)-box 4
-3.78243907630414e-5

GO:0008284: positive regulation of cell proliferation   SOX9    SRY (sex determining region Y)-box 9
-0.0005192837332730816
GO:0008284: positive regulation of cell proliferation   STAMBP  STAM binding protein    -0.0001705258
241929094
GO:0008284: positive regulation of cell proliferation   TBX3    T-box 3 0.0012274221775040348
GO:0008284: positive regulation of cell proliferation   TGFB1   transforming growth factor, beta 1
-7.329146074140709e-5
GO:0008284: positive regulation of cell proliferation   TGFB2   transforming growth factor, beta 2
-0.0010612035139734976
GO:0008284: positive regulation of cell proliferation   TGFBR1  transforming growth factor, beta rece
ptor 1  0.0003406303055982047
GO:0008284: positive regulation of cell proliferation   TIPIN   TIMELESS interacting protein    -0.00
05408522058055696
GO:0008284: positive regulation of cell proliferation   TNC     tenascin C      0.0007354561470014528
GO:0008284: positive regulation of cell proliferation   TNFRSF11A       tumor necrosis factor recepto
r superfamily, member 11a, NFKB activator       0.002780359294954262
GO:0008284: positive regulation of cell proliferation   TTK     TTK protein kinase      0.00108792083
10257387
GO:0008284: positive regulation of cell proliferation   VEGFA   vascular endothelial growth factor A
0.0005964425569342986
GO:0008284: positive regulation of cell proliferation   VEGFC   vascular endothelial growth factor C
-0.003368988555231368
GO:0008284: positive regulation of cell proliferation   WDR77   WD repeat domain 77     0.00013368823
95141298
GO:0008284: positive regulation of cell proliferation   WNT1    wingless-type MMTV integration site f
amily, member 1 0.0007880109313703516
GO:0008284: positive regulation of cell proliferation   WNT10B  wingless-type MMTV integration site f
amily, member 10B       -0.0013909401968136062
GO:0008284: positive regulation of cell proliferation   WWTR1   WW domain containing transcription re
gulator 1       0.0009025750375793526
GO:0008284: positive regulation of cell proliferation   YAP1    Yes-associated protein 1        -0.00
01642635831546427З
GO:0009409: response to cold    ADM     adrenomedullin  0.0022397904488824032
GO:0009409: response to cold    AGT     angiotensinogen (serpin peptidase inhibitor, clade A, member
 8)     -0.0011351167577243072
GO:0009409: response to cold    CXCL10  chemokine (C-X-C motif) ligand 10       6.310089253870686e-5
GO:0009409: response to cold    HSPA2   heat shock 70kDa protein 2      -0.00010820053490843656
GO:0009409: response to cold    IMPDH1  IMP (inosine 5'-monophosphate) dehydrogenase 1  -0.0018739381
381500842
GO:0009409: response to cold    THRA    thyroid hormone receptor, alpha 0.0007637331060292847
GO:0009611: response to wounding        ADM     adrenomedullin  0.0022363247259476046
GO:0009611: response to wounding        AGER    advanced glycosylation end product-specific receptor
-0.0001742117748962843
GO:0009611: response to wounding        CCL2    chemokine (C-C motif) ligand 2  0.0008155802961288844
GO:0009611: response to wounding        CX3CR1  chemokine (C-X3-C motif) receptor 1     -0.0008561588
55469583
GO:0009611: response to wounding        FGF7    fibroblast growth factor 7      0.000633525478579317
GO:0009611: response to wounding        GAP43   growth associated protein 43    -0.000129617682620766
96
GO:0009611: response to wounding        HOXB13  homeobox B13    0.0018246993757456676
GO:0009611: response to wounding        ITGB4   integrin, beta 4        0.0005794066538673223
GO:0009611: response to wounding        MDK     midkine (neurite growth-promoting factor 2)     0.001
662821949830099
GO:0009611: response to wounding        PAX6    paired box 6    0.0019339674064337217
GO:0009611: response to wounding        PDX1    pancreatic and duodenal homeobox 1      0.00025689463
20263224
GO:0009611: response to wounding        SLC1A3  solute carrier family 1 (glial high affinity glutamat
e transporter), member 3        -0.003119516609261705
GO:0009611: response to wounding        TGFB1   transforming growth factor, beta 1      -7.2615426280
97549e-5
GO:0009611: response to wounding        TGFB2   transforming growth factor, beta 2      -0.0010585455
482918817
GO:0009611: response to wounding        TNC     tenascin C      0.0007340740888897685
GO:0009611: response to wounding        VASH1   vasohibin 1     0.0005983295308603423
GO:0009611: response to wounding        WNT1    wingless-type MMTV integration site family, member 1
0.0007865486151669959
GO:0010460: positive regulation of heart rate   ADM     adrenomedullin  0.0022598983240230894
GO:0010460: positive regulation of heart rate   HEY2    hes-related family bHLH transcription factor
 with YRPW motif 2      0.002731516827600816
GO:0010460: positive regulation of heart rate   SLC1A1  solute carrier family 1 (neuronal/epithelial
 high affinity glutamate transporter, system Xag), member 1     -0.0009115214102514174
GO:0019731: antibacterial humoral response      ADM     adrenomedullin  0.0022574560122283 51
GO:0019933: cAMP-mediated signaling     ADM     adrenomedullin  0.00223539206214254З
GO:0019933: cAMP-mediated signaling     EPHA5   EPH receptor A5 0.000598134029290315
GO:0019933: cAMP-mediated signaling     SOX9    SRY (sex determining region Y)-box 9    -0.0005187964
341177323
GO:0030819: positive regulation of cAMP biosynthetic process    ADM     adrenomedullin  0.00224866908
88087212
GO:0030819: positive regulation of cAMP biosynthetic process    AVP     arginine vasopressin    -0.00
09429485082957513
GO:0031100: organ regeneration  ADM     adrenomedullin  0.002245904287921161

GO:0031100: organ regeneration  BAK1    BCL2-antagonist/killer 1        -0.001882565407390322
GO:0031100: organ regeneration  CAST    calpastatin     -0.003045237234049009
GO:0031100: organ regeneration  CAV1    caveolin 1, caveolae protein, 22kDa     -0.000536641460892182
1
GO:0031100: organ regeneration  CCL2    chemokine (C-C motif) ligand 2  0.0008201448042826113
GO:0031100: organ regeneration  CCND1   cyclin D1       -0.002646093423570791
GO:0031100: organ regeneration  CCNE1   cyclin E1       0.00037361656676417873
GO:0031100: organ regeneration  CXCL12  chemokine (C-X-C motif) ligand 12       -0.001213591761001855
7
GO:0031100: organ regeneration  LEF1    lymphoid enhancer-binding factor 1      -0.000100321165752307
69
GO:0031100: organ regeneration  MED1    mediator complex subunit 1      0.0011347722602584717
GO:0031100: organ regeneration  MKI67   marker of proliferation Ki-67   0.0015788570868711287
GO:0031100: organ regeneration  NOTCH1  notch 1 0.0005211581069085083
GO:0031100: organ regeneration  PDX1    pancreatic and duodenal homeobox 1      0.000258939959868928
7
GO:0031100: organ regeneration  TGFB1   transforming growth factor, beta 1      -7.379122373448564e-5
GO:0031100: organ regeneration  TYMS    thymidylate synthetase  0.0015697154640689755
GO:0031100: organ regeneration  WNT1    wingless-type MMTV integration site family, member 1    0.000
7895807975310407
GO:0031102: neuron projection regeneration      ADM     adrenomedullin  0.00225745601222835
GO:0031623: receptor internalization    ADM     adrenomedullin  0.002246176257821061
GO:0031623: receptor internalization    GRB2    growth factor receptor-bound protein 2  0.00043168060
687490844
GO:0031623: receptor internalization    NEDD4   neural precursor cell expressed, developmentally down
-regulated 4, E3 ubiquitin protein ligase       0.0022725957418444046
GO:0032496: response to lipopolysaccharide      ADM     adrenomedullin  0.0022475306611521566
GO:0032496: response to lipopolysaccharide      CEBPB   CCAAT/enhancer binding protein (C/EBP), beta
-0.0002749827394197019
GO:0032496: response to lipopolysaccharide      CITED1  Cbp/p300-interacting transactivator, with Gl
u/Asp-rich carboxy-terminal domain, 1   0.002831618417272932
GO:0032496: response to lipopolysaccharide      CXCL13  chemokine (C-X-C motif) ligand 13       0.003
0073685792690887
GO:0032496: response to lipopolysaccharide      CYP27B1 cytochrome P450, family 27, subfamily B, poly
peptide 1       -0.0004743644212478374
GO:0032496: response to lipopolysaccharide      HDAC2   histone deacetylase 2   -0.001210945989045251
7
GO:0032496: response to lipopolysaccharide      HMGB2   high mobility group box 2       0.00031142181
42957026
GO:0032496: response to lipopolysaccharide      IDO1    indoleamine 2,3-dioxygenase 1   0.00123211170
53351704
GO:0032496: response to lipopolysaccharide      IRAK1   interleukin-1 receptor-associated kinase 1
-0.0016163172384537738
GO:0032496: response to lipopolysaccharide      JAK2    Janus kinase 2  -3.072034762998232e-5
GO:0032496: response to lipopolysaccharide      LGALS9  lectin, galactoside-binding, soluble, 9 -0.00
011650656355459592
GO:0032496: response to lipopolysaccharide      LIAS    lipoic acid synthetase  -0.000556379619369132
4
GO:0032496: response to lipopolysaccharide      NOTCH1  notch 1 0.0005213228839302022
GO:0032496: response to lipopolysaccharide      S100A14 S100 calcium binding protein A14        -0.00
2171255778808952
GO:0032496: response to lipopolysaccharide      S100A7  S100 calcium binding protein A7 0.00159819453
12622562
GO:0032496: response to lipopolysaccharide      TFAP2A  transcription factor AP-2 alpha (activating e
nhancer binding protein 2 alpha)        0.0004815641135286411
GO:0032496: response to lipopolysaccharide      TH      tyrosine hydroxylase    -0.000343046910413423
3
GO:0032496: response to lipopolysaccharide      TNFRSF11A       tumor necrosis factor receptor superf
amily, member 11a, NFKB activator       0.002789346206407167
GO:0032868: response to insulin ADM     adrenomedullin  0.0022571304207396298
GO:0032868: response to insulin AGRP    agouti related protein homolog (mouse)  -0.002593156783284863
7
GO:0032868: response to insulin CITED1  Cbp/p300-interacting transactivator, with Glu/Asp-rich carbox
y-terminal domain, 1    0.002845316550106531
GO:0032868: response to insulin EGR2    early growth response 2 0.0014564520686788352
GO:0032868: response to insulin GCNT1   glucosaminyl (N-acetyl) transferase 1, core 2   0.00070523077
00962994
GO:0032868: response to insulin GGH     gamma-glutamyl hydrolase (conjugase, folylpolygammaglutamyl h
ydrolase)       -0.00031518036116047845
GO:0032868: response to insulin LEP     leptin  0.0032240763975226903
GO:0032868: response to insulin LYN     LYN proto-oncogene, Src family tyrosine kinase  -0.0014327910
398884238
GO:0032868: response to insulin SIRT1   sirtuin 1       1.6888794085940226e-9
GO:0042475: odontogenesis of dentin-containing tooth    ADM     adrenomedullin  0.002237900733244428
GO:0042475: odontogenesis of dentin-containing tooth    BAX     BCL2-associated X protein       -0.00
0425344349383711
GO:0042475: odontogenesis of dentin-containing tooth    BCL2L11 BCL2-like 11 (apoptosis facilitator)
-0.0007749680304709944
GO:0042475: odontogenesis of dentin-containing tooth    BMP4    bone morphogenetic protein 4    -0.00
03225660506252650 4
GO:0042475: odontogenesis of dentin-containing tooth    BMP7    bone morphogenetic protein 7    0.000

8558030048578339
GO:0042475: odontogenesis of dentin-containing tooth    CTNNB1  catenin (cadherin-associated protei
n), beta 1, 88kDa       -0.00011709937218164791
GO:0042475: odontogenesis of dentin-containing tooth    EDA     ectodysplasin A -0.000814413405726349
7
GO:0042475: odontogenesis of dentin-containing tooth    EDAR    ectodysplasin A receptor        0.000
5940033102101944
GO:0042475: odontogenesis of dentin-containing tooth    FOXC1   forkhead box C1 -2.258048567095939e-5
GO:0042475: odontogenesis of dentin-containing tooth    GLI2    GLI family zinc finger 2        0.001
853532447217153
GO:0042475: odontogenesis of dentin-containing tooth    GLI3    GLI family zinc finger 3        -0.00
2151267354800685
GO:0042475: odontogenesis of dentin-containing tooth    HDAC2   histone deacetylase 2   -0.0012076448
724121483
GO:0042475: odontogenesis of dentin-containing tooth    JAG2    jagged 2        -5.740313696177119e-6
GO:0042475: odontogenesis of dentin-containing tooth    LEF1    lymphoid enhancer-binding factor 1
-0.00010029132528500789
GO:0042475: odontogenesis of dentin-containing tooth    LRP6    low density lipoprotein receptor-rela
ted protein 6   0.0001493770263094102
GO:0042475: odontogenesis of dentin-containing tooth    MSX1    msh homeobox 1  -0.002768754521346488
GO:0042475: odontogenesis of dentin-containing tooth    NF2     neurofibromin 2 (merlin)        -0.00
12721457975192092
GO:0042475: odontogenesis of dentin-containing tooth    PITX2   paired-like homeodomain 2       0.002
176081270647936
GO:0042475: odontogenesis of dentin-containing tooth    SHH     sonic hedgehog  0.0005998055592181083
GO:0042475: odontogenesis of dentin-containing tooth    SMO     smoothened, frizzled class receptor
0.002139208736228798
GO:0042475: odontogenesis of dentin-containing tooth    SOSTDC1 sclerostin domain containing 1  -0.00
15248043678562425
GO:0042594: response to starvation      ADM     adrenomedullin  0.002249074129288369
GO:0042594: response to starvation      HMGCL   3-hydroxymethyl-3-methylglutaryl-CoA lyase      -0.00
10697342762814776
GO:0043065: positive regulation of apoptotic process    ADM     adrenomedullin  0.0022498471391765914
GO:0043065: positive regulation of apoptotic process    ALDH1A2 aldehyde dehydrogenase 1 family, memb
er A2   -0.0030351067118519893
GO:0043065: positive regulation of apoptotic process    APC     adenomatous polyposis coli      0.000
6454770127754742
GO:0043065: positive regulation of apoptotic process    ARL6IP5 ADP-ribosylation factor-like 6 intera
cting protein 5 0.00016989518960989095
GO:0043065: positive regulation of apoptotic process    BAK1    BCL2-antagonist/killer 1        -0.00
18861589047304052
GO:0043065: positive regulation of apoptotic process    BARD1   BRCA1 associated RING domain 1  0.001
7548024382458183
GO:0043065: positive regulation of apoptotic process    BAX     BCL2-associated X protein       -0.00
042692508516908206
GO:0043065: positive regulation of apoptotic process    BCL2L11 BCL2-like 11 (apoptosis facilitator)
-0.0007779063600343187
GO:0043065: positive regulation of apoptotic process    BMP4    bone morphogenetic protein 4    -0.00
032443722104737464
GO:0043065: positive regulation of apoptotic process    BMP7    bone morphogenetic protein 7    0.000
8594945586820189
GO:0043065: positive regulation of apoptotic process    BNIP3   BCL2/adenovirus E1B 19kDa interacting
protein 3       0.0029296395185289745
GO:0043065: positive regulation of apoptotic process    BNIP3L  BCL2/adenovirus E1B 19kDa interacting
protein 3-like  -5.7595195156419006e-5
GO:0043065: positive regulation of apoptotic process    CAMK1D  calcium/calmodulin-dependent protein
 kinase ID      -0.002602852291366281
GO:0043065: positive regulation of apoptotic process    CDKN2A  cyclin-dependent kinase inhibitor 2A
0.0017700596267689462
GO:0043065: positive regulation of apoptotic process    CTNNB1  catenin (cadherin-associated protei
n), beta 1, 88kDa       -0.00011699420714832125
GO:0043065: positive regulation of apoptotic process    CYP1B1  cytochrome P450, family 1, subfamily
 B, polypeptide 1       0.0003993754843956998
GO:0043065: positive regulation of apoptotic process    DIABLO  diablo, IAP-binding mitochondrial pro
tein    -0.0005983876797996916
GO:0043065: positive regulation of apoptotic process    ECT2    epithelial cell transforming 2  0.001
0689481078891747
GO:0043065: positive regulation of apoptotic process    EEF1A2  eukaryotic translation elongation fac
tor 1 alpha 2   -0.0018661239554174096
GO:0043065: positive regulation of apoptotic process    FAS     Fas cell surface death receptor -3.34
9400984284594e-5
GO:0043065: positive regulation of apoptotic process    FOXO1   forkhead box O1 0.001809917847530766
GO:0043065: positive regulation of apoptotic process    HMGA2   high mobility group AT-hook 2   0.001
5165756674686438
GO:0043065: positive regulation of apoptotic process    HMGB1   high mobility group box 1       -0.00
07791148104959677
GO:0043065: positive regulation of apoptotic process    HOXA5   homeobox A5     0.00107329093703349
GO:0043065: positive regulation of apoptotic process    IGFBP3  insulin-like growth factor binding pr
otein 3 0.0008397833620326271
GO:0043065: positive regulation of apoptotic process    ITGA6   integrin, alpha 6       0.00168251559
5332933

GO:0043065: positive regulation of apoptotic process    ITGB1   integrin, beta 1 (fibronectin receptor, beta polypeptide, antigen CD29 includes MDF2, MSK12) 0.002442181192397091
GO:0043065: positive regulation of apoptotic process    KCNMA1  potassium large conductance calcium-activated channel, subfamily M, alpha member 1   -0.0006647765072681668
GO:0043065: positive regulation of apoptotic process    KLF11   Kruppel-like factor 11  0.0006228236785852949
GO:0043065: positive regulation of apoptotic process    MAP3K5  mitogen-activated protein kinase kinase kinase 5     0.00034480602709511117
GO:0043065: positive regulation of apoptotic process    MELK    maternal embryonic leucine zipper kinase     0.002082269148067369
GO:0043065: positive regulation of apoptotic process    NOTCH1  notch 1 0.0005220117098999359
GO:0043065: positive regulation of apoptotic process    PRKDC   protein kinase, DNA-activated, catalytic polypeptide -0.0018647529353817882
GO:0043065: positive regulation of apoptotic process    RARG    retinoic acid receptor, gamma   -0.002331950121265366
GO:0043065: positive regulation of apoptotic process    RXRA    retinoid X receptor, alpha      0.0011209440490071176
GO:0043065: positive regulation of apoptotic process    S100B   S100 calcium binding protein B  0.005216725688016132
GO:0043065: positive regulation of apoptotic process    SAV1    salvador family WW domain containing protein 1      -0.0023482111932885863
GO:0043065: positive regulation of apoptotic process    SFRP1   secreted frizzled-related protein 1 0.0012937941625074981
GO:0043065: positive regulation of apoptotic process    SHQ1    SHQ1, H/ACA ribonucleoprotein assembly factor        -0.0014430001259238606
GO:0043065: positive regulation of apoptotic process    SIRT1   sirtuin 1       -6.269227723450117e-7
GO:0043065: positive regulation of apoptotic process    SLIT2   slit homolog 2 (Drosophila)     -0.001630296568624057
GO:0043065: positive regulation of apoptotic process    SOX4    SRY (sex determining region Y)-box 4 -3.684691026260639e-5
GO:0043065: positive regulation of apoptotic process    SPDEF   SAM pointed domain containing ETS transcription factor       0.003151609414133948
GO:0043065: positive regulation of apoptotic process    STEAP3  STEAP family member 3, metalloreductase      0.002622569480875091
GO:0043065: positive regulation of apoptotic process    TGFB1   transforming growth factor, beta 1 -7.391976541094114e-5
GO:0043065: positive regulation of apoptotic process    TGFB3   transforming growth factor, beta 3 -0.001836027210878173
GO:0043065: positive regulation of apoptotic process    TGM2    transglutaminase 2      -0.00018249272792951118
GO:0043065: positive regulation of apoptotic process    TOP2A   topoisomerase (DNA) II alpha 170kDa -8.027966129823012e-5
GO:0043065: positive regulation of apoptotic process    TP53    tumor protein p53       0.001184408141686088
GO:0043065: positive regulation of apoptotic process    WNT10B  wingless-type MMTV integration site family, member 10B       -0.0013974797399692578
GO:0043065: positive regulation of apoptotic process    WT1     Wilms tumor 1   -0.0005105571750694807
GO:0043116: negative regulation of vascular permeability        ADM     adrenomedullin  0.002240060658406547
GO:0043116: negative regulation of vascular permeability        ANGPT1  angiopoietin 1  0.0009015850685944626
GO:0043116: negative regulation of vascular permeability        PTPRJ   protein tyrosine phosphatase, receptor type, J        -0.0003389540755447038
GO:0043116: negative regulation of vascular permeability        SLIT2   slit homolog 2 (Drosophila) -0.001621789123813906
GO:0045906: negative regulation of vasoconstriction     ADM     adrenomedullin  0.0022669326078214743
GO:0045906: negative regulation of vasoconstriction     CX3CL1  chemokine (C-X3-C motif) ligand 1 0.0021518927541464596
GO:0045906: negative regulation of vasoconstriction     LEP     leptin  0.0032405431344564323
GO:0045909: positive regulation of vasodilation ADM     adrenomedullin  0.0022443629968690025
GO:0045909: positive regulation of vasodilation AGT     angiotensinogen (serpin peptidase inhibitor, clade A, member 8)     -0.0011387460881769529
GO:0045909: positive regulation of vasodilation ALOX12  arachidonate 12-lipoxygenase    -0.0019597816912600807
GO:0045909: positive regulation of vasodilation GJA1    gap junction protein, alpha 1, 43kDa    -0.00016429640094642515
GO:0045909: positive regulation of vasodilation HMOX1   heme oxygenase (decycling) 1    -0.00021850438694969522
GO:0046879: hormone secretion   ADM     adrenomedullin  0.002257456012228351
GO:0048589: developmental growth        ADM     adrenomedullin  0.0022482873183828442
GO:0048589: developmental growth        ASPM    asp (abnormal spindle) homolog, microcephaly associated (Drosophila) 1.8428209697634427e-5
GO:0048589: developmental growth        GATA3   GATA binding protein 3  -4.186639102464043e-5
GO:0048589: developmental growth        GLI2    GLI family zinc finger 2        0.001862347567809895
GO:0048589: developmental growth        GLI3    GLI family zinc finger 3        -0.0021647469411024607
GO:0048589: developmental growth        SOX10   SRY (sex determining region Y)-box 10   0.00019614568829672116
GO:0048589: developmental growth        STRA6   stimulated by retinoic acid 6   -0.0016890001525057105

GO:0048589: developmental growth        TYMS    thymidylate synthetase  0.0015724753015463396
GO:0050829: defense response to Gram-negative bacterium ADM     adrenomedullin  0.0022295977012530515
GO:0050829: defense response to Gram-negative bacterium AZU1    azurocidin 1    0.00019713893299273016
GO:0050829: defense response to Gram-negative bacterium HMGB2   high mobility group box 2       0.0003034666997796713
GO:0050829: defense response to Gram-negative bacterium IL12B   interleukin 12B 0.0012732256978753317
GO:0050829: defense response to Gram-negative bacterium MMP7    matrix metallopeptidase 7 (matrilysin, uterine)     0.0004242195853160617
GO:0050829: defense response to Gram-negative bacterium S100A7  S100 calcium binding protein A7 0.0015833155525608472
GO:0050829: defense response to Gram-negative bacterium SERPINE1        serpin peptidase inhibitor, clade E (nexin, plasminogen activator inhibitor type 1), member 1        0.00011281464216740762
GO:0050830: defense response to Gram-positive bacterium ADM     adrenomedullin  0.002225749123551416
GO:0050830: defense response to Gram-positive bacterium HMGB2   high mobility group box 2       0.000302344660230977
GO:0050830: defense response to Gram-positive bacterium IL27RA  interleukin 27 receptor, alpha  -0.0004817948427053052
GO:0050830: defense response to Gram-positive bacterium MMP7    matrix metallopeptidase 7 (matrilysin, uterine)     0.00042389702871016083
GO:0050830: defense response to Gram-positive bacterium SEH1L   SEH1-like (S. cerevisiae)       -0.000729047521512466 2
GO:0051384: response to glucocorticoid  ADM     adrenomedullin  0.0022542552450695685
GO:0051384: response to glucocorticoid  AREG    amphiregulin    0.002475830499200363
GO:0051384: response to glucocorticoid  BCHE    butyrylcholinesterase   -8.878500389435954e-6
GO:0051384: response to glucocorticoid  BCL2    B-cell CLL/lymphoma 2   -4.979173179272789e-6
GO:0051384: response to glucocorticoid  BMP4    bone morphogenetic protein 4    -0.0003251113367670451
GO:0051384: response to glucocorticoid  C3      complement component 3  0.002040502706986942
GO:0051384: response to glucocorticoid  CAV1    caveolin 1, caveolae protein, 22kDa     -0.0005380439279837386
GO:0051384: response to glucocorticoid  CTSV    cathepsin V     0.0006611152623379376
GO:0051384: response to glucocorticoid  FAS     Fas cell surface death receptor -3.35856138899954e-5
GO:0051384: response to glucocorticoid  IGFBP2  insulin-like growth factor binding protein 2, 36kDa     0.0001467815782874009
GO:0051384: response to glucocorticoid  MDK     midkine (neurite growth-promoting factor 2)     0.001683973947861747
GO:0051384: response to glucocorticoid  PDX1    pancreatic and duodenal homeobox 1      0.00026055980019655675
GO:0051384: response to glucocorticoid  RXRA    retinoid X receptor, alpha      0.0011231159560783006
GO:0051384: response to glucocorticoid  S100B   S100 calcium binding protein B  0.005228628183890329
GO:0051384: response to glucocorticoid  TAT     tyrosine aminotransferase       0.0010617844211096921
GO:0051384: response to glucocorticoid  TYMS    thymidylate synthetase  0.001577414410666 4016
GO:0060670: branching involved in labyrinthine layer morphogenesis      ADM     adrenomedullin  0.0022358403364742176
GO:0060670: branching involved in labyrinthine layer morphogenesis      FGFR2   fibroblast growth factor receptor 2  0.0007620538491925126
GO:0060670: branching involved in labyrinthine layer morphogenesis      GRB2    growth factor receptor-bound protein 2       0.00042949293077006694
GO:0060712: spongiotrophoblast layer development        ADM     adrenomedullin  0.0022633438948317305
GO:0060712: spongiotrophoblast layer development        CITED1  Cbp/p300-interacting transactivator, with Glu/Asp-rich carboxy-terminal domain, 1   0.002854266054516458
GO:0097084: vascular smooth muscle cell development     ADM     adrenomedullin  0.0022597339534806216
GO:0097084: vascular smooth muscle cell development     HES1    hes family bHLH transcription factor 1      -0.0009206495187076083
GO:0097084: vascular smooth muscle cell development     HEY2    hes-related family bHLH transcription factor with YRPW motif 2        0.0027312608271300976
GO:2001214: positive regulation of vasculogenesis       ADM     adrenomedullin  0.002257456012228351
GO:0000122: negative regulation of transcription from RNA polymerase II promoter        AES     amino-terminal enhancer of split     0.0013551182258712238
GO:0000122: negative regulation of transcription from RNA polymerase II promoter        ALX1    ALX homeobox 1       0.0022943594153471167
GO:0000122: negative regulation of transcription from RNA polymerase II promoter        ASCL1   achaete-scute family bHLH transcription factor 1     -0.0014341810788094525
GO:0000122: negative regulation of transcription from RNA polymerase II promoter        AURKB   aurora kinase B      0.0003510242537475111
GO:0000122: negative regulation of transcription from RNA polymerase II promoter        BACH1   BTB and CNC homology 1, basic leucine zipper transcription factor 1  0.0004858791321776827
GO:0000122: negative regulation of transcription from RNA polymerase II promoter        BCL11A  B-cell CLL/lymphoma 11A (zinc finger protein)        -0.0009778037521593697
GO:0000122: negative regulation of transcription from RNA polymerase II promoter        BMP4    bone morphogenetic protein 4        -0.0003229166554189383
GO:0000122: negative regulation of transcription from RNA polymerase II promoter        CAV1    caveolin 1, caveolae protein, 22kDa  -0.0005358515151663144
GO:0000122: negative regulation of transcription from RNA polymerase II promoter        CCND1   cyclin D1    -0.002636169611349596 2
GO:0000122: negative regulation of transcription from RNA polymerase II promoter        CTNNB1  catenin (cadherin-associated protein), beta 1, 88kDa -0.0001170720241604335
GO:0000122: negative regulation of transcription from RNA polymerase II promoter        DACH1   dachshund family transcription factor 1      0.0020906120418930363
GO:0000122: negative regulation of transcription from RNA polymerase II promoter        DAXX    death

-domain associated protein	0.0008926760602729517
GO:0000122: negative regulation of transcription from RNA polymerase II promoter	DICER1	dicer 1, ribonuclease type III	-1.6085242419160052e-5
GO:0000122: negative regulation of transcription from RNA polymerase II promoter	DKK1	dickkopf WNT signaling pathway inhibitor 1	-0.0001808628076458805
GO:0000122: negative regulation of transcription from RNA polymerase II promoter	DMRT1	doublesex and mab-3 related transcription factor 1	0.001506282695641828
GO:0000122: negative regulation of transcription from RNA polymerase II promoter	E2F1	E2F transcription factor 1	0.0020999519103675314
GO:0000122: negative regulation of transcription from RNA polymerase II promoter	E2F6	E2F transcription factor 6	0.000718393620700157
GO:0000122: negative regulation of transcription from RNA polymerase II promoter	E2F8	E2F transcription factor 8	0.0017125554910919414
GO:0000122: negative regulation of transcription from RNA polymerase II promoter	E4F1	E4F transcription factor 1	0.0014829497580189303
GO:0000122: negative regulation of transcription from RNA polymerase II promoter	EFNA1	ephrin-A1	-0.0005211819739195807
GO:0000122: negative regulation of transcription from RNA polymerase II promoter	EGR1	early growth response 1	0.0010942465833379375
GO:0000122: negative regulation of transcription from RNA polymerase II promoter	EN1	engrailed homeobox 1	-1.5604905726076528e-5
GO:0000122: negative regulation of transcription from RNA polymerase II promoter	ENG	endoglin	0.0008276634707962915
GO:0000122: negative regulation of transcription from RNA polymerase II promoter	ETS2	v-ets avian erythroblastosis virus E26 oncogene homolog 2	-0.0010168219306386026
GO:0000122: negative regulation of transcription from RNA polymerase II promoter	EZH2	enhancer of zeste 2 polycomb repressive complex 2 subunit	-0.0001365219345557046
GO:0000122: negative regulation of transcription from RNA polymerase II promoter	FGFR2	fibroblast growth factor receptor 2	0.0007640876482155909
GO:0000122: negative regulation of transcription from RNA polymerase II promoter	FGFR3	fibroblast growth factor receptor 3	0.00022303140380648304
GO:0000122: negative regulation of transcription from RNA polymerase II promoter	FOXC2	forkhead box C2 (MFH-1, mesenchyme forkhead 1)	0.0017500454485836895
GO:0000122: negative regulation of transcription from RNA polymerase II promoter	FOXM1	forkhead box M1	0.0001973847451437482
GO:0000122: negative regulation of transcription from RNA polymerase II promoter	FOXO1	forkhead box O1	0.0018021228107439747
GO:0000122: negative regulation of transcription from RNA polymerase II promoter	FOXO3	forkhead box O3	0.0012014358019426352
GO:0000122: negative regulation of transcription from RNA polymerase II promoter	GATA2	GATA binding protein 2	-0.0004520628084075147
GO:0000122: negative regulation of transcription from RNA polymerase II promoter	GATA3	GATA binding protein 3	-3.899696096053636e-5
GO:0000122: negative regulation of transcription from RNA polymerase II promoter	GATA6	GATA binding protein 6	-2.8164220902766755e-5
GO:0000122: negative regulation of transcription from RNA polymerase II promoter	GFI1	growth factor independent 1 transcription repressor	0.0016660985531133793
GO:0000122: negative regulation of transcription from RNA polymerase II promoter	GLI2	GLI family zinc finger 2	0.0018555577160233197
GO:0000122: negative regulation of transcription from RNA polymerase II promoter	GLI3	GLI family zinc finger 3	-0.002154025316445987
GO:0000122: negative regulation of transcription from RNA polymerase II promoter	HDAC2	histone deacetylase 2	-0.0012083777929821988
GO:0000122: negative regulation of transcription from RNA polymerase II promoter	HES1	hes family bHLH transcription factor 1	-0.0009119510641972796
GO:0000122: negative regulation of transcription from RNA polymerase II promoter	HEXIM1	hexamethylene bis-acetamide inducible 1	0.0001529605964295342
GO:0000122: negative regulation of transcription from RNA polymerase II promoter	HEY2	hes-related family bHLH transcription factor with YRPW motif 2	0.002702102519300041
GO:0000122: negative regulation of transcription from RNA polymerase II promoter	HINFP	histone H4 transcription factor	0.0009935279983226887
GO:0000122: negative regulation of transcription from RNA polymerase II promoter	HIPK2	homeodomain interacting protein kinase 2	0.000771549302232574
GO:0000122: negative regulation of transcription from RNA polymerase II promoter	HMGA2	high mobility group AT-hook 2	0.001510506126961629
GO:0000122: negative regulation of transcription from RNA polymerase II promoter	HMGB1	high mobility group box 1	-0.000777058587311547
GO:0000122: negative regulation of transcription from RNA polymerase II promoter	ID2	inhibitor of DNA binding 2, dominant negative helix-loop-helix protein	6.010272536522299e-5
GO:0000122: negative regulation of transcription from RNA polymerase II promoter	IFNG	interferon, gamma	-4.8930795400852984e-5
GO:0000122: negative regulation of transcription from RNA polymerase II promoter	IGBP1	immunoglobulin (CD79A) binding protein 1	0.0028521663821031665
GO:0000122: negative regulation of transcription from RNA polymerase II promoter	IRF7	interferon regulatory factor 7	-0.0013078823059719273
GO:0000122: negative regulation of transcription from RNA polymerase II promoter	ISL1	ISL LIM homeobox 1	7.891835877104974e-5
GO:0000122: negative regulation of transcription from RNA polymerase II promoter	KLF11	Kruppel-like factor 11	0.0006176055842431094
GO:0000122: negative regulation of transcription from RNA polymerase II promoter	LEF1	lymphoid enhancer-binding factor 1	-0.00010024651916415745

GO:0000122: negative regulation of transcription from RNA polymerase II promoter LEP lepti
n 0.0031947153487910497
GO:0000122: negative regulation of transcription from RNA polymerase II promoter LRP8 low d
ensity lipoprotein receptor-related protein 8, apolipoprotein e receptor -0.000932766265972705
1
GO:0000122: negative regulation of transcription from RNA polymerase II promoter MED1 media
tor complex subunit 1 0.0011306282915268353
GO:0000122: negative regulation of transcription from RNA polymerase II promoter MEF2C myocy
te enhancer factor 2C 0.0009745291680395772
GO:0000122: negative regulation of transcription from RNA polymerase II promoter MSX1 msh h
omeobox 1 -0.002772207992396805
GO:0000122: negative regulation of transcription from RNA polymerase II promoter MYC v-myc
avian myelocytomatosis viral oncogene homolog -0.0011376644613760585
GO:0000122: negative regulation of transcription from RNA polymerase II promoter NFIB nucle
ar factor I/B 0.0029227627622900347
GO:0000122: negative regulation of transcription from RNA polymerase II promoter NFX1 nucle
ar transcription factor, X-box binding 1 -6.97792408087923e-5
GO:0000122: negative regulation of transcription from RNA polymerase II promoter NOC2L nucle
olar complex associated 2 homolog (S. cerevisiae) -0.00015122390490234476
GO:0000122: negative regulation of transcription from RNA polymerase II promoter NOTCH1 notch
1 0.000519308328221836
GO:0000122: negative regulation of transcription from RNA polymerase II promoter NR1H3 nucle
ar receptor subfamily 1, group H, member 3 0.0008015691707731485
GO:0000122: negative regulation of transcription from RNA polymerase II promoter NR2E1 nucle
ar receptor subfamily 2, group E, member 1 -0.0029779697158289634
GO:0000122: negative regulation of transcription from RNA polymerase II promoter NRIP1 nucle
ar receptor interacting protein 1 0.0010722096159011534
GO:0000122: negative regulation of transcription from RNA polymerase II promoter ORC2 origi
n recognition complex, subunit 2 -0.0006128708749429622
GO:0000122: negative regulation of transcription from RNA polymerase II promoter PAX3 paire
d box 3 -0.0029482489937430593
GO:0000122: negative regulation of transcription from RNA polymerase II promoter PAX6 paire
d box 6 0.0019377480187608097
GO:0000122: negative regulation of transcription from RNA polymerase II promoter PDX1 pancr
eatic and duodenal homeobox 1 0.00025767106892919694
GO:0000122: negative regulation of transcription from RNA polymerase II promoter PER2 perio
d circadian clock 2 0.0012395282488476763
GO:0000122: negative regulation of transcription from RNA polymerase II promoter PHF14 PHD f
inger protein 14 0.0032282891775884657
GO:0000122: negative regulation of transcription from RNA polymerase II promoter PITX2 paire
d-like homeodomain 2 0.002178479479224185
GO:0000122: negative regulation of transcription from RNA polymerase II promoter PLK1 polo-
like kinase 1 0.0010189579927959973
GO:0000122: negative regulation of transcription from RNA polymerase II promoter PLK3 polo-
like kinase 3 0.0025667371334023837
GO:0000122: negative regulation of transcription from RNA polymerase II promoter POU1F1 POU c
lass 1 homeobox 1 0.00013762064483145763
GO:0000122: negative regulation of transcription from RNA polymerase II promoter PPP1R13L
protein phosphatase 1, regulatory subunit 13 like -0.001475378327993628
GO:0000122: negative regulation of transcription from RNA polymerase II promoter PROX1 prosp
ero homeobox 1 0.0011228040046543475
GO:0000122: negative regulation of transcription from RNA polymerase II promoter PTCH1 patch
ed 1 -6.590360048554093e-5
GO:0000122: negative regulation of transcription from RNA polymerase II promoter RARG retin
oic acid receptor, gamma -0.0023218968713357028
GO:0000122: negative regulation of transcription from RNA polymerase II promoter RBPJ recom
bination signal binding protein for immunoglobulin kappa J region 0.0009823683903396026
GO:0000122: negative regulation of transcription from RNA polymerase II promoter RREB1 ras r
esponsive element binding protein 1 -0.0019836430271720495
GO:0000122: negative regulation of transcription from RNA polymerase II promoter RXRA retin
oid X receptor, alpha 0.0011162949316676943
GO:0000122: negative regulation of transcription from RNA polymerase II promoter S100A1 S100
calcium binding protein A1 0.001587087287117167
GO:0000122: negative regulation of transcription from RNA polymerase II promoter SALL1 spalt
-like transcription factor 1 -0.0024404452992737844
GO:0000122: negative regulation of transcription from RNA polymerase II promoter SHH sonic
hedgehog 0.0006007415421319317
GO:0000122: negative regulation of transcription from RNA polymerase II promoter SIRT1 sirtu
in 1 -1.5243308058533812e-6
GO:0000122: negative regulation of transcription from RNA polymerase II promoter SIRT2 sirtu
in 2 -0.0008532116563844861
GO:0000122: negative regulation of transcription from RNA polymerase II promoter SIX1 SIX h
omeobox 1 -0.0018719885232860096
GO:0000122: negative regulation of transcription from RNA polymerase II promoter SKI SKI p
roto-oncogene -0.0006711169904449967
GO:0000122: negative regulation of transcription from RNA polymerase II promoter SMO smoot
hened, frizzled class receptor 0.0021415373065663676
GO:0000122: negative regulation of transcription from RNA polymerase II promoter SOX11 SRY
(sex determining region Y)-box 11 -0.000209235570876596
GO:0000122: negative regulation of transcription from RNA polymerase II promoter SOX18 SRY
(sex determining region Y)-box 18 0.0012897248055904992

GO:0000122: negative regulation of transcription from RNA polymerase II promoter SOX9 SRY (sex determining region Y)-box 9 -0.0005192689492928105
GO:0000122: negative regulation of transcription from RNA polymerase II promoter SPDEF SAM pointed domain containing ETS transcription factor 0.003140107582151736
GO:0000122: negative regulation of transcription from RNA polymerase II promoter TBX3 T-box 3 0.0012274167587069326
GO:0000122: negative regulation of transcription from RNA polymerase II promoter TCF7L2 transcription factor 7-like 2 (T-cell specific, HMG-box) 0.0005743624922343588
GO:0000122: negative regulation of transcription from RNA polymerase II promoter TFAP2A transcription factor AP-2 alpha (activating enhancer binding protein 2 alpha) 0.0004803589178179184
7
GO:0000122: negative regulation of transcription from RNA polymerase II promoter TFAP2C transcription factor AP-2 gamma (activating enhancer binding protein 2 gamma) 0.001061660309989715
GO:0000122: negative regulation of transcription from RNA polymerase II promoter TFCP2L1 transcription factor CP2-like 1 0.0004495484372106232
GO:0000122: negative regulation of transcription from RNA polymerase II promoter TGFB1 transforming growth factor, beta 1 -7.309557659658694e-5
GO:0000122: negative regulation of transcription from RNA polymerase II promoter THRB thyroid hormone receptor, beta 0.0019459537768192004
GO:0000122: negative regulation of transcription from RNA polymerase II promoter TP53 tumor protein p53 0.0011771957338800399
GO:0000122: negative regulation of transcription from RNA polymerase II promoter TP73 tumor protein p73 0.0010290644601768277
GO:0000122: negative regulation of transcription from RNA polymerase II promoter TPR translocated promoter region, nuclear basket protein -0.0006977416686542844
GO:0000122: negative regulation of transcription from RNA polymerase II promoter TRPS1 trichorhinophalangeal syndrome I -0.0001781453355674055
GO:0000122: negative regulation of transcription from RNA polymerase II promoter TWIST1 twist family bHLH transcription factor 1 -0.0013431266966506346
GO:0000122: negative regulation of transcription from RNA polymerase II promoter VDR vitamin D (1,25- dihydroxyvitamin D3) receptor 0.0005640252179659059
GO:0000122: negative regulation of transcription from RNA polymerase II promoter VEGFA vascular endothelial growth factor A 0.0005957531111647733
GO:0000122: negative regulation of transcription from RNA polymerase II promoter VLDLR very low density lipoprotein receptor 0.0009540857561877541
GO:0000122: negative regulation of transcription from RNA polymerase II promoter VPS72 vacuolar protein sorting 72 homolog (S. cerevisiae) 0.0005591059954160044
GO:0000122: negative regulation of transcription from RNA polymerase II promoter WFS1 Wolfram syndrome 1 (wolframin) 0.0005773100865995265
GO:0000122: negative regulation of transcription from RNA polymerase II promoter WNT10B wingless-type MMTV integration site family, member 10B -0.0013901551851193844
GO:0000122: negative regulation of transcription from RNA polymerase II promoter WT1 Wilms tumor 1 -0.0005075491450663937
GO:0000122: negative regulation of transcription from RNA polymerase II promoter WWP2 WW domain containing E3 ubiquitin protein ligase 2 -0.0009581368781906829
GO:0000122: negative regulation of transcription from RNA polymerase II promoter WWTR1 WW domain containing transcription regulator 1 0.0009016010907040479
GO:0000122: negative regulation of transcription from RNA polymerase II promoter XBP1 X-box binding protein 1 0.0002675704031140029
GO:0000122: negative regulation of transcription from RNA polymerase II promoter YBX1 Y box binding protein 1 -0.0009146851717373777
GO:0000122: negative regulation of transcription from RNA polymerase II promoter YY1 YY1 transcription factor 0.0013417771859454127
GO:0000122: negative regulation of transcription from RNA polymerase II promoter ZBTB18 zinc finger and BTB domain containing 18 0.001273315113422191
GO:0000122: negative regulation of transcription from RNA polymerase II promoter ZNF148 zinc finger protein 148 0.002106336098087779
GO:0000122: negative regulation of transcription from RNA polymerase II promoter ZNF205 zinc finger protein 205 0.0005964780761849497
GO:0009887: organ morphogenesis AES amino-terminal enhancer of split 0.0013551700108189616
GO:0009887: organ morphogenesis CCL2 chemokine (C-C motif) ligand 2 0.0008169353018639715
GO:0009887: organ morphogenesis E2F4 E2F transcription factor 4, p107/p130-binding -0.0023475824
803697344
GO:0009887: organ morphogenesis EVL Enah/Vasp-like 0.0019399869451577245
GO:0009887: organ morphogenesis FGFR2 fibroblast growth factor receptor 2 0.0007636188960153762
GO:0009887: organ morphogenesis GATA2 GATA binding protein 2 -0.0004513740507206311
GO:0009887: organ morphogenesis GATA3 GATA binding protein 3 -3.812156262349606e-5
GO:0009887: organ morphogenesis GSK3B glycogen synthase kinase 3 beta 0.001550460969730984
GO:0009887: organ morphogenesis IL7 interleukin 7 0.00087157097582893
GO:0009887: organ morphogenesis LHX1 LIM homeobox 1 -0.000759297118267382
GO:0009887: organ morphogenesis NRP1 neuropilin 1 -0.0006436307598317074
GO:0009887: organ morphogenesis PAX3 paired box 3 -0.002947608275728009
GO:0009887: organ morphogenesis PAX6 paired box 6 0.0019368754649154137
GO:0009887: organ morphogenesis PDX1 pancreatic and duodenal homeobox 1 0.0002573621353780432
7
GO:0009887: organ morphogenesis PTCH1 patched 1 -6.62651310595012e-5
GO:0009887: organ morphogenesis STX2 syntaxin 2 0.00030844837225686944
GO:0009887: organ morphogenesis TBX3 T-box 3 0.0012278099074311086
GO:0009887: organ morphogenesis TH tyrosine hydroxylase -0.00034241678024209463
GO:0009887: organ morphogenesis THRB thyroid hormone receptor, beta 0.0019457902119361344
GO:0009887: organ morphogenesis TRPS1 trichorhinophalangeal syndrome I -0.000177554815867657

GO:0009887: organ morphogenesis VEGFC vascular endothelial growth factor C -0.00336647139016756
GO:0010629: negative regulation of gene expression AES amino-terminal enhancer of split 0.0013519140010937415
GO:0010629: negative regulation of gene expression BAK1 BCL2-antagonist/killer 1 -0.0018730431728721149
GO:0010629: negative regulation of gene expression BBS4 Bardet-Biedl syndrome 4 -0.0005090853651926164
GO:0010629: negative regulation of gene expression C1QTNF3 C1q and tumor necrosis factor related protein 3 0.00013892932752127408
GO:0010629: negative regulation of gene expression CCNB1 cyclin B1 -0.0008809332885227912
GO:0010629: negative regulation of gene expression CDC42 cell division cycle 42 0.001238289788968595
GO:0010629: negative regulation of gene expression CRYAB crystallin, alpha B 0.0009830222652376518
GO:0010629: negative regulation of gene expression ESR1 estrogen receptor 1 -0.0009468039846844982
GO:0010629: negative regulation of gene expression GJA1 gap junction protein, alpha 1, 43kDa -0.00016121412881582505
GO:0010629: negative regulation of gene expression HEY2 hes-related family bHLH transcription factor with YRPW motif 2 0.0026944214350133095
GO:0010629: negative regulation of gene expression HINFP histone H4 transcription factor 0.0009911689595940656
GO:0010629: negative regulation of gene expression ID2 inhibitor of DNA binding 2, dominant negative helix-loop-helix protein 6.011590669622907e-5
GO:0010629: negative regulation of gene expression IFNA2 interferon, alpha 2 -0.001714628700838684
GO:0010629: negative regulation of gene expression IFNG interferon, gamma -4.883233209385633e-5
GO:0010629: negative regulation of gene expression LGALS9 lectin, galactoside-binding, soluble, 9 -0.00011678216342476746
GO:0010629: negative regulation of gene expression MEF2C myocyte enhancer factor 2C 0.0009717040659472457
GO:0010629: negative regulation of gene expression PGR progesterone receptor 0.00035122284772455564
GO:0010629: negative regulation of gene expression RNASEH2B ribonuclease H2, subunit B -4.3467780817131196e-5
GO:0010629: negative regulation of gene expression SFRP1 secreted frizzled-related protein 1 0.0012794387325746081
GO:0010629: negative regulation of gene expression SIRT1 sirtuin 1 -1.7852213432194947e-6
GO:0010629: negative regulation of gene expression SLIT2 slit homolog 2 (Drosophila) -0.0016175920743850963
GO:0010629: negative regulation of gene expression SMO smoothened, frizzled class receptor 0.0021359331492284035
GO:0010629: negative regulation of gene expression SOX11 SRY (sex determining region Y)-box 11 -0.00020867327667661625
GO:0010629: negative regulation of gene expression STC2 stanniocalcin 2 -0.0013951345470435844
GO:0010629: negative regulation of gene expression TGFB1 transforming growth factor, beta 1 -7.294610457040557e-5
GO:0010629: negative regulation of gene expression WNT4 wingless-type MMTV integration site family, member 4 -0.0002515425559214698
GO:0010629: negative regulation of gene expression WWP2 WW domain containing E3 ubiquitin protein ligase 2 -0.0009557896463014889
GO:0010629: negative regulation of gene expression YY1 YY1 transcription factor 0.001337878804402301
GO:0010629: negative regulation of gene expression ZNF148 zinc finger protein 148 0.0021006780488864877
GO:0016055: Wnt signaling pathway AES amino-terminal enhancer of split 0.0013527395166038428
GO:0016055: Wnt signaling pathway CCNE1 cyclin E1 0.00037081171272804154
GO:0016055: Wnt signaling pathway CD24 CD24 molecule 0.001070781637080905
GO:0016055: Wnt signaling pathway CELSR2 cadherin, EGF LAG seven-pass G-type receptor 2 -0.0011340597826993825
GO:0016055: Wnt signaling pathway CPE carboxypeptidase E -0.0025422313126101325
GO:0016055: Wnt signaling pathway CSNK2B casein kinase 2, beta polypeptide 0.0015273538504854429
GO:0016055: Wnt signaling pathway CTNNB1 catenin (cadherin-associated protein), beta 1, 88kDa -0.00011677085235793604
GO:0016055: Wnt signaling pathway CTNNBIP1 catenin, beta interacting protein 1 0.0005851476176684202
GO:0016055: Wnt signaling pathway DRD2 dopamine receptor D2 -0.00023390629743860674
GO:0016055: Wnt signaling pathway LEF1 lymphoid enhancer-binding factor 1 -0.0001003597289257529
GO:0016055: Wnt signaling pathway LRP5 low density lipoprotein receptor-related protein 5 3.0445528481599608e-5
GO:0016055: Wnt signaling pathway LRP6 low density lipoprotein receptor-related protein 6 0.00014952042784598294
GO:0016055: Wnt signaling pathway NDRG2 NDRG family member 2 0.0021715200009711005

GO:0016055: Wnt signaling pathway       PITX2   paired-like homeodomain 2       0.0021740623016134673
GO:0016055: Wnt signaling pathway       SOSTDC1 sclerostin domain containing 1  -0.001522919432176506
1
GO:0016055: Wnt signaling pathway       WNT1    wingless-type MMTV integration site family, member 1
0.0007862487173985867
GO:0016055: Wnt signaling pathway       WNT10B  wingless-type MMTV integration site family, member 10
B       -0.0013872120620972378
GO:0016055: Wnt signaling pathway       WNT5A   wingless-type MMTV integration site family, member 5A
-0.0006668920517706575
GO:0032091: negative regulation of protein binding      AES     amino-terminal enhancer of split
0.0013463307260487643
GO:0032091: negative regulation of protein binding      AURKA   aurora kinase A 0.0009991882076638278
GO:0032091: negative regulation of protein binding      AURKB   aurora kinase B 0.0003460323517961098
4
GO:0032091: negative regulation of protein binding      BAX     BCL2-associated X protein       -0.00
04228922884857201
GO:0032091: negative regulation of protein binding      CAV1    caveolin 1, caveolae protein, 22kDa
-0.000532948143787745
GO:0032091: negative regulation of protein binding      CTNNBIP1        catenin, beta interacting pro
tein 1  0.0005811182393533249
GO:0032091: negative regulation of protein binding      GOLGA2  golgin A2       0.0001670497670462498
5
GO:0032091: negative regulation of protein binding      GSK3B   glycogen synthase kinase 3 beta 0.001
5382435686406638
GO:0032091: negative regulation of protein binding      NES     nestin  0.0013608046463423498
GO:0032091: negative regulation of protein binding      PRKCD   protein kinase C, delta -0.0011190671
462786525
GO:0032091: negative regulation of protein binding      TEX14   testis expressed 14     0.00171536115
52606788
GO:0032091: negative regulation of protein binding      TMBIM6  transmembrane BAX inhibitor motif con
taining 6       -0.0001469758710778853
GO:0045892: negative regulation of transcription, DNA-templated AES     amino-terminal enhancer of sp
lit     0.001351784730574072
GO:0045892: negative regulation of transcription, DNA-templated ASCL1   achaete-scute family bHLH tra
nscription factor 1     -0.0014295509173068216
GO:0045892: negative regulation of transcription, DNA-templated ATF5    activating transcription fact
or 5    -0.0027301283776263673
GO:0045892: negative regulation of transcription, DNA-templated BASP1   brain abundant, membrane atta
ched signal protein 1   0.0004775931535688066
GO:0045892: negative regulation of transcription, DNA-templated BIRC5   baculoviral IAP repeat contai
ning 5  -0.00026031859065165546
GO:0045892: negative regulation of transcription, DNA-templated BMP4    bone morphogenetic protein 4
-0.0003219902464734653
GO:0045892: negative regulation of transcription, DNA-templated BMP7    bone morphogenetic protein 7
0.0008543523023319433
GO:0045892: negative regulation of transcription, DNA-templated CDKN2A  cyclin-dependent kinase inhib
itor 2A 0.0017543369588370777
GO:0045892: negative regulation of transcription, DNA-templated CEBPB   CCAAT/enhancer binding protei
n (C/EBP), beta -0.0002735178826884575
GO:0045892: negative regulation of transcription, DNA-templated CHMP1A  charged multivesicular body p
rotein 1A       -0.0007783748268909475
GO:0045892: negative regulation of transcription, DNA-templated CITED1  Cbp/p300-interacting transact
ivator, with Glu/Asp-rich carboxy-terminal domain, 1    0.002812628287906496
GO:0045892: negative regulation of transcription, DNA-templated CLOCK   clock circadian regulator
0.0002001716583007587
GO:0045892: negative regulation of transcription, DNA-templated CTNNB1  catenin (cadherin-associated
 protein), beta 1, 88kDa        -0.00011686077626024047
GO:0045892: negative regulation of transcription, DNA-templated DACH1   dachshund family transcriptio
n factor 1      0.002085597153487456
GO:0045892: negative regulation of transcription, DNA-templated DAXX    death-domain associated prote
in      0.0008895562116497326
GO:0045892: negative regulation of transcription, DNA-templated E2F1    E2F transcription factor 1
0.0020936531362438795
GO:0045892: negative regulation of transcription, DNA-templated EZH2    enhancer of zeste 2 polycomb
 repressive complex 2 subunit   -0.0001367235849565081
GO:0045892: negative regulation of transcription, DNA-templated FOXM1   forkhead box M1 0.00019648954
15783686
GO:0045892: negative regulation of transcription, DNA-templated FOXO1   forkhead box O1 0.00179695934
86101643
GO:0045892: negative regulation of transcription, DNA-templated GATA3   GATA binding protein 3  -3.86
4292371758242e-5
GO:0045892: negative regulation of transcription, DNA-templated GATA6   GATA binding protein 6  -2.83
353326952951e-5
GO:0045892: negative regulation of transcription, DNA-templated GFI1    growth factor independent 1 t
ranscription repressor  0.00166184971780774З
GO:0045892: negative regulation of transcription, DNA-templated GLI3    GLI family zinc finger 3
-0.0021471929500517495
GO:0045892: negative regulation of transcription, DNA-templated GREM1   gremlin 1, DAN family BMP ant
agonist -0.0008252087915197293
GO:0045892: negative regulation of transcription, DNA-templated HDAC2   histone deacetylase 2   -0.00
12055610861570981

GO:0045892: negative regulation of transcription, DNA-templated HES1    hes family bHLH transcription factor 1        -0.0009092210165938033
GO:0045892: negative regulation of transcription, DNA-templated HEXIM1  hexamethylene bis-acetamide inducible 1      0.00015247737052206335
GO:0045892: negative regulation of transcription, DNA-templated HEY2    hes-related family bHLH transcription factor with YRPW motif 2       0.002693177601517478
GO:0045892: negative regulation of transcription, DNA-templated HEYL    hes-related family bHLH transcription factor with YRPW motif-like    0.0006763193724865664
GO:0045892: negative regulation of transcription, DNA-templated HINFP   histone H4 transcription factor      0.0009906205915676393
GO:0045892: negative regulation of transcription, DNA-templated HMGA1   high mobility group AT-hook 1 -0.00031828935388999443
GO:0045892: negative regulation of transcription, DNA-templated HMGA2   high mobility group AT-hook 2 0.001506300927691984
GO:0045892: negative regulation of transcription, DNA-templated HMGB2   high mobility group box 2 0.0003055442002599929
GO:0045892: negative regulation of transcription, DNA-templated ID2     inhibitor of DNA binding 2, dominant negative helix-loop-helix protein       6.017573893963402e-5
GO:0045892: negative regulation of transcription, DNA-templated IFNA2   interferon, alpha 2     -0.0017141527027006909
GO:0045892: negative regulation of transcription, DNA-templated IL4     interleukin 4   0.0002594261121362925
GO:0045892: negative regulation of transcription, DNA-templated LANCL2  LanC lantibiotic synthetase component C-like 2 (bacterial)   0.0011831124860059638
GO:0045892: negative regulation of transcription, DNA-templated LEF1    lymphoid enhancer-binding factor 1   -0.00010028864894957699
GO:0045892: negative regulation of transcription, DNA-templated LGR4    leucine-rich repeat containing G protein-coupled receptor 4  0.00030794274591630844
GO:0045892: negative regulation of transcription, DNA-templated LHX1    LIM homeobox 1  -0.000757199167502307
GO:0045892: negative regulation of transcription, DNA-templated NOTCH1  notch 1 0.0005177525193474509
GO:0045892: negative regulation of transcription, DNA-templated PAX2    paired box 2    -0.001610811635700988
GO:0045892: negative regulation of transcription, DNA-templated PER2    period circadian clock 2 0.0012358609273226847
GO:0045892: negative regulation of transcription, DNA-templated PHB2    prohibitin 2    -0.000734865088509974
GO:0045892: negative regulation of transcription, DNA-templated PLCB1   phospholipase C, beta 1 (phosphoinositide-specific)  0.00015317901283391243
GO:0045892: negative regulation of transcription, DNA-templated PML     promyelocytic leukemia  -0.0006791444323532432
GO:0045892: negative regulation of transcription, DNA-templated PROX1   prospero homeobox 1     0.0011191792502334788
GO:0045892: negative regulation of transcription, DNA-templated PURA    purine-rich element binding protein A        -0.00016995048969449845
GO:0045892: negative regulation of transcription, DNA-templated RB1     retinoblastoma 1        -0.0014911611868604357
GO:0045892: negative regulation of transcription, DNA-templated RBPJ    recombination signal binding protein for immunoglobulin kappa J region      0.0009799788864599795
GO:0045892: negative regulation of transcription, DNA-templated SALL1   spalt-like transcription factor 1    -0.0024318228424323636
GO:0045892: negative regulation of transcription, DNA-templated SFRP1   secreted frizzled-related protein 1  0.001278059449732701
GO:0045892: negative regulation of transcription, DNA-templated SIRT1   sirtuin 1       -1.9291066704488234e-6
GO:0045892: negative regulation of transcription, DNA-templated SIRT2   sirtuin 2       -0.0008505944997300519
GO:0045892: negative regulation of transcription, DNA-templated SIX3    SIX homeobox 3  0.0020329987152231705
GO:0045892: negative regulation of transcription, DNA-templated SOX10   SRY (sex determining region Y)-box 10      0.00018912903029736042
GO:0045892: negative regulation of transcription, DNA-templated SOX18   SRY (sex determining region Y)-box 18      0.0012864361700746579
GO:0045892: negative regulation of transcription, DNA-templated SOX9    SRY (sex determining region Y)-box 9       -0.0005176144539026877
GO:0045892: negative regulation of transcription, DNA-templated TBX3    T-box 3 0.0012245786504059767
GO:0045892: negative regulation of transcription, DNA-templated TCF7L2  transcription factor 7-like 2 (T-cell specific, HMG-box)      0.0005723457182564316
GO:0045892: negative regulation of transcription, DNA-templated TFAP2A  transcription factor AP-2 alpha (activating enhancer binding protein 2 alpha)        0.000479173327917902
GO:0045892: negative regulation of transcription, DNA-templated TGFB1   transforming growth factor, beta 1   -7.275729444127839e-5
GO:0045892: negative regulation of transcription, DNA-templated THRA    thyroid hormone receptor, alpha      0.0007619017160781874
GO:0045892: negative regulation of transcription, DNA-templated TP53    tumor protein p53       0.0011729599534958288
GO:0045892: negative regulation of transcription, DNA-templated TWIST1  twist family bHLH transcription factor 1     -0.001338717112133745
GO:0045892: negative regulation of transcription, DNA-templated VDR     vitamin D (1,25- dihydroxyvitamin D3) receptor       0.0005628198711417726
GO:0045892: negative regulation of transcription, DNA-templated WNT4    wingless-type MMTV integratio

n site family, member 4 -0.00025106657156398254
GO:0045892: negative regulation of transcription, DNA-templated WNT5A   wingless-type MMTV integratio
n site family, member 5A        -0.0006660780260055667
GO:0045892: negative regulation of transcription, DNA-templated WT1     Wilms tumor 1   -0.0005057158
283539179
GO:0045892: negative regulation of transcription, DNA-templated WWP2    WW domain containing E3 ubiqu
itin protein ligase 2   -0.000955562380329093
GO:0045892: negative regulation of transcription, DNA-templated ZBTB18  zinc finger and BTB domain co
ntaining 18     0.0012698825729290388
GO:0045892: negative regulation of transcription, DNA-templated ZNF148  zinc finger protein 148 0.002
100034896123665
GO:0045892: negative regulation of transcription, DNA-templated ZNF24   zinc finger protein 24  0.001
2894295712332702
GO:0060761: negative regulation of response to cytokine stimulus        AES     amino-terminal enhanc
er of split     0.0013463150483353048
GO:0070555: response to interleukin-1   AES     amino-terminal enhancer of split        0.00136291437
06802714
GO:0070555: response to interleukin-1   CITED1  Cbp/p300-interacting transactivator, with Glu/Asp-ric
h carboxy-terminal domain, 1    0.002841623051359837
GO:0070555: response to interleukin-1   IGBP1   immunoglobulin (CD79A) binding protein 1        0.002
8698158549162107
GO:0070555: response to interleukin-1   IRAK1   interleukin-1 receptor-associated kinase 1      -0.00
1622894614721151
GO:0070555: response to interleukin-1   LGALS9  lectin, galactoside-binding, soluble, 9 -0.0001166480
4654304971
GO:0070555: response to interleukin-1   PRKCA   protein kinase C, alpha -6.143989571451896e-6
GO:0070555: response to interleukin-1   TNFRSF11A       tumor necrosis factor receptor superfamily, m
ember 11a, NFKB activator       0.002797670307330731
GO:0090090: negative regulation of canonical Wnt signaling pathway      AES     amino-terminal enhanc
er of split     0.0013446295997899037
GO:0090090: negative regulation of canonical Wnt signaling pathway      APC     adenomatous polyposis
coli    0.0006405561210755343
GO:0090090: negative regulation of canonical Wnt signaling pathway      AXIN1   axin 1  -0.0007259334
725522164
GO:0090090: negative regulation of canonical Wnt signaling pathway      CAV1    caveolin 1, caveolae
 protein, 22kDa -0.000532305282418228
GO:0090090: negative regulation of canonical Wnt signaling pathway      CDH2    cadherin 2, type 1, N
-cadherin (neuronal)    -0.0006365518356656023
GO:0090090: negative regulation of canonical Wnt signaling pathway      DKK1    dickkopf WNT signalin
g pathway inhibitor 1   -0.0001796534172234697
GO:0090090: negative regulation of canonical Wnt signaling pathway      EGR1    early growth response
1       0.0010843701458420837
GO:0090090: negative regulation of canonical Wnt signaling pathway      FOXO1   forkhead box O1 0.001
7858973853553523
GO:0090090: negative regulation of canonical Wnt signaling pathway      FOXO3   forkhead box O3 0.001
1896221452105987
GO:0090090: negative regulation of canonical Wnt signaling pathway      GLI1    GLI family zinc finge
r 1     -0.0012874867377596435
GO:0090090: negative regulation of canonical Wnt signaling pathway      GLI3    GLI family zinc finge
r 3     -0.002132751476636087
GO:0090090: negative regulation of canonical Wnt signaling pathway      GREM1   gremlin 1, DAN family
BMP antagonist  -0.0008181393414796465
GO:0090090: negative regulation of canonical Wnt signaling pathway      GSK3B   glycogen synthase kin
ase 3 beta      0.001536306766493082
GO:0090090: negative regulation of canonical Wnt signaling pathway      ISL1    ISL LIM homeobox 1
7.974583911044468e-5
GO:0090090: negative regulation of canonical Wnt signaling pathway      LEF1    lymphoid enhancer-bin
ding factor 1   -0.00010044236999972472
GO:0090090: negative regulation of canonical Wnt signaling pathway      LRP6    low density lipoprote
in receptor-related protein 6   0.00014822228708482923
GO:0090090: negative regulation of canonical Wnt signaling pathway      NOTCH1  notch 1 0.00051450565
65424627
GO:0090090: negative regulation of canonical Wnt signaling pathway      PPP2R3A protein phosphatase
 2, regulatory subunit B'', alpha       -3.401723456631783e-5
GO:0090090: negative regulation of canonical Wnt signaling pathway      PSMA5   proteasome (prosome,
 macropain) subunit, alpha type, 5      0.0003290731665883212
GO:0090090: negative regulation of canonical Wnt signaling pathway      PSMD11  proteasome (prosome,
 macropain) 26S subunit, non-ATPase, 11 -0.0010879595894439222
GO:0090090: negative regulation of canonical Wnt signaling pathway      PSMD13  proteasome (prosome,
 macropain) 26S subunit, non-ATPase, 13 -6.047868398404913e-5
GO:0090090: negative regulation of canonical Wnt signaling pathway      SFRP1   secreted frizzled-rel
ated protein 1  0.0012664764144225064
GO:0090090: negative regulation of canonical Wnt signaling pathway      SHH     sonic hedgehog  0.000
5936221018870155
GO:0090090: negative regulation of canonical Wnt signaling pathway      SOSTDC1 sclerostin domain con
taining 1       -0.0015163835930333239
GO:0090090: negative regulation of canonical Wnt signaling pathway      SOX10   SRY (sex determining
 region Y)-box 10       0.0001853021654686114
GO:0090090: negative regulation of canonical Wnt signaling pathway      SOX9    SRY (sex determining
 region Y)-box 9        -0.000513964178902587
GO:0090090: negative regulation of canonical Wnt signaling pathway      TCF7L2  transcription factor

7-like 2 (T-cell specific, HMG-box) 0.0005681147050651933
GO:0090090: negative regulation of canonical Wnt signaling pathway WNT4 wingless-type MMTV integration site family, member 4 -0.0002480656864292685
GO:0090090: negative regulation of canonical Wnt signaling pathway WNT5A wingless-type MMTV integration site family, member 5A -0.0006616863540872481
GO:0090090: negative regulation of canonical Wnt signaling pathway WWTR1 WW domain containing transcription regulator 1 0.0008909372223944106
GO:2000210: positive regulation of anoikis AES amino-terminal enhancer of split 0.0013463150483353048
GO:0051180: vitamin transport AFM afamin -0.001693919942297489
GO:0001933: negative regulation of protein phosphorylation AGER advanced glycosylation end product-specific receptor -0.0001735806048704551
GO:0001933: negative regulation of protein phosphorylation ANGPT1 angiopoietin 1 0.0008942613539592709
GO:0001933: negative regulation of protein phosphorylation CCNB1 cyclin B1 -0.0008789843383837509
GO:0001933: negative regulation of protein phosphorylation CDK5RAP3 CDK5 regulatory subunit associated protein 3 0.0011304767666894013
GO:0001933: negative regulation of protein phosphorylation CIB1 calcium and integrin binding 1 (calmyrin) 5.769625484162538e-5
GO:0001933: negative regulation of protein phosphorylation IGFBP3 insulin-like growth factor binding protein 3 0.0008324809561600393
GO:0001933: negative regulation of protein phosphorylation LRP6 low density lipoprotein receptor-related protein 6 0.000147590793305975
GO:0001933: negative regulation of protein phosphorylation LYN LYN proto-oncogene, Src family tyrosine kinase -0.0014095478897937037
GO:0001933: negative regulation of protein phosphorylation MAS1 MAS1 proto-oncogene, G protein-coupled receptor 0.000293665967793492
GO:0001933: negative regulation of protein phosphorylation PAX6 paired box 6 0.0019193292556878086
GO:0001933: negative regulation of protein phosphorylation PRKDC protein kinase, DNA-activated, catalytic polypeptide -0.0018424092921994436
GO:0001933: negative regulation of protein phosphorylation PTEN phosphatase and tensin homolog 1.984896351515504e-5
GO:0001933: negative regulation of protein phosphorylation SLIT2 slit homolog 2 (Drosophila) -0.0016053719849884746
GO:0001933: negative regulation of protein phosphorylation TGFB1 transforming growth factor, beta 1 -7.162701065306365e-5
GO:0001933: negative regulation of protein phosphorylation WWTR1 WW domain containing transcription regulator 1 0.0008893523319973953
GO:0006954: inflammatory response AGER advanced glycosylation end product-specific receptor -0.0001741605193325246
GO:0006954: inflammatory response AKT1 v-akt murine thymoma viral oncogene homolog 1 0.0007293226205558226
GO:0006954: inflammatory response AZU1 azurocidin 1 0.00019770024192330277
GO:0006954: inflammatory response C3 complement component 3 0.002021493529410979
GO:0006954: inflammatory response CAMK1D calcium/calmodulin-dependent protein kinase ID -0.0025835848378670543
GO:0006954: inflammatory response CCL2 chemokine (C-C motif) ligand 2 0.0008151701822056868
GO:0006954: inflammatory response CCL7 chemokine (C-C motif) ligand 7 -0.0024000209296410925
GO:0006954: inflammatory response CCL8 chemokine (C-C motif) ligand 8 -0.0005901406027813145
GO:0006954: inflammatory response CEBPB CCAAT/enhancer binding protein (C/EBP), beta -0.0027383613982190493
GO:0006954: inflammatory response CXCL10 chemokine (C-X-C motif) ligand 10 6.185463254846113e-5
GO:0006954: inflammatory response CXCL13 chemokine (C-X-C motif) ligand 13 0.0029883681309419165
GO:0006954: inflammatory response CXCR4 chemokine (C-X-C motif) receptor 4 0.0007930653805844545
GO:0006954: inflammatory response ECM1 extracellular matrix protein 1 -0.0015538446464899176
GO:0006954: inflammatory response HMGB1 high mobility group box 1 -0.0007758328482250262
GO:0006954: inflammatory response IFNA2 interferon, alpha 2 -0.0017147744852922715
GO:0006954: inflammatory response IGFBP4 insulin-like growth factor binding protein 4 -0.0012759153273434408
GO:0006954: inflammatory response KIT v-kit Hardy-Zuckerman 4 feline sarcoma viral oncogene homolog 0.00027329672935830077
GO:0006954: inflammatory response LGALS9 lectin, galactoside-binding, soluble, 9 -0.00011702440917983025
GO:0006954: inflammatory response LIAS lipoic acid synthetase -0.0005533963804851617
GO:0006954: inflammatory response NFX1 nuclear transcription factor, X-box binding 1 -7.071783108679627e-5
GO:0006954: inflammatory response ORM1 orosomucoid 1 -0.0004722964838708868
GO:0006954: inflammatory response PIK3CD phosphatidylinositol-4,5-bisphosphate 3-kinase, catalytic subunit delta -0.0007796206153916808
GO:0006954: inflammatory response PRKCZ protein kinase C, zeta -0.0015985844588706389
GO:0006954: inflammatory response RXRA retinoid X receptor, alpha 0.0011135967399531442
GO:0006954: inflammatory response SPHK1 sphingosine kinase 1 0.0018079082150883495

GO:0006954: inflammatory response TGFB1 transforming growth factor, beta 1 -7.265089659943401e-5
GO:0006954: inflammatory response THBS1 thrombospondin 1 -0.0010339970738101817
GO:0006954: inflammatory response TNFAIP3 tumor necrosis factor, alpha-induced protein 3 0.0011147675776687195
GO:0006954: inflammatory response TP73 tumor protein p73 0.0010263950253054854
GO:0007166: cell surface receptor signaling pathway AGER advanced glycosylation end product-specific receptor -0.0001747321234888183
GO:0007166: cell surface receptor signaling pathway AGT angiotensinogen (serpin peptidase inhibitor, clade A, member 8) -0.0011409484028379244
GO:0007166: cell surface receptor signaling pathway CCL2 chemokine (C-C motif) ligand 2 0.0008207016857744447
GO:0007166: cell surface receptor signaling pathway CD9 CD9 molecule -0.0025985720497186987
GO:0007166: cell surface receptor signaling pathway CXCL10 chemokine (C-X-C motif) ligand 10 6.367702381689452e-5
GO:0007166: cell surface receptor signaling pathway CXCL13 chemokine (C-X-C motif) ligand 13 0.0030085060487865323
GO:0007166: cell surface receptor signaling pathway EGFR epidermal growth factor receptor 0.0006904557236067012
GO:0007166: cell surface receptor signaling pathway EVL Enah/Vasp-like 0.0019458134711758195
GO:0007166: cell surface receptor signaling pathway FAS Fas cell surface death receptor -3.351252345089459e-5
GO:0007166: cell surface receptor signaling pathway IFNA2 interferon, alpha 2 -0.0017255790808465062
GO:0007166: cell surface receptor signaling pathway IFNG interferon, gamma -4.894359067742936e-5
GO:0007166: cell surface receptor signaling pathway IL27RA interleukin 27 receptor, alpha -0.00048582361139973783
GO:0007166: cell surface receptor signaling pathway INHA inhibin, alpha 0.00021179489211757058
GO:0007166: cell surface receptor signaling pathway INHBA inhibin, beta A -0.0013625139485714598
GO:0007166: cell surface receptor signaling pathway JMJD6 jumonji domain containing 6 0.0036071729847609803
GO:0007166: cell surface receptor signaling pathway PRLR prolactin receptor 0.0021934715255802865
GO:0007166: cell surface receptor signaling pathway TACSTD2 tumor-associated calcium signal transducer 2 -0.002270967381742542
GO:0007166: cell surface receptor signaling pathway TSPAN6 tetraspanin 6 0.001374710145542575
GO:0007259: JAK-STAT cascade AGER advanced glycosylation end product-specific receptor -0.00017234174170604297
GO:0007259: JAK-STAT cascade CCL2 chemokine (C-C motif) ligand 2 0.0007961782341408852
GO:0007259: JAK-STAT cascade FGFR3 fibroblast growth factor receptor 3 0.00021486648949821377
GO:0007259: JAK-STAT cascade JAK2 Janus kinase 2 -3.807257164588723e-5
GO:0007259: JAK-STAT cascade STAMBP STAM binding protein -0.0001682348269519361
GO:0007259: JAK-STAT cascade STAT5A signal transducer and activator of transcription 5A 0.0015624547873845597
GO:0007420: brain development AGER advanced glycosylation end product-specific receptor -0.0001744685042829096
GO:0007420: brain development APOD apolipoprotein D 0.0026341666368316255
GO:0007420: brain development BAK1 BCL2-antagonist/killer 1 -0.001880152323493059
GO:0007420: brain development BBS7 Bardet-Biedl syndrome 7 -0.0009791393949996827
GO:0007420: brain development BCL2L11 BCL2-like 11 (apoptosis facilitator) -0.000776140215065747
GO:0007420: brain development BRCA2 breast cancer 2, early onset -1.0011110254309207e-5
GO:0007420: brain development CAST calpastatin -0.003040365683747042
GO:0007420: brain development CD9 CD9 molecule -0.002591700643047095
GO:0007420: brain development CDK5RAP3 CDK5 regulatory subunit associated protein 3 0.0011403608568741014
GO:0007420: brain development CITED1 Cbp/p300-interacting transactivator, with Glu/Asp-rich carboxy-terminal domain, 1 0.0028247663778074544
GO:0007420: brain development CST3 cystatin C -7.082053828187304e-5
GO:0007420: brain development EGR2 early growth response 2 0.0014429781073361958
GO:0007420: brain development FOXC1 forkhead box C1 -2.190496884133556e-5
GO:0007420: brain development GRHL2 grainyhead-like 2 (Drosophila) 0.0010086452618165902
GO:0007420: brain development IGF1R insulin-like growth factor 1 receptor 0.001073753351488752
GO:0007420: brain development MAP1S microtubule-associated protein 1S 0.0005311389918535142
GO:0007420: brain development MAPT microtubule-associated protein tau 0.0015541305624082937
GO:0007420: brain development MED1 mediator complex subunit 1 0.0011322471481942738
GO:0007420: brain development NES nestin 0.001377504396871962
GO:0007420: brain development PHGDH phosphoglycerate dehydrogenase 0.00029394755769203716
GO:0007420: brain development PRKDC protein kinase, DNA-activated, catalytic polypeptide -0.0018594000602884392
GO:0007420: brain development PROX1 prospero homeobox 1 0.001124256151056211
GO:0007420: brain development PTCH1 patched 1 -6.575980054019745e-5
GO:0007420: brain development RELN reelin 0.0015412381043283842
GO:0007420: brain development SHROOM2 shroom family member 2 0.0011148750556103648
GO:0007420: brain development SIX3 SIX homeobox 3 0.0020422795242461176
GO:0007420: brain development SPHK1 sphingosine kinase 1 0.0018148802383100922
GO:0007420: brain development STAR steroidogenic acute regulatory protein 0.0008967393806491165

GO:0007420: brain development   STMN1   stathmin 1      0.0005610502424353884
GO:0007420: brain development   THRA    thyroid hormone receptor, alpha 0.0007648297005168199
GO:0007420: brain development   TULP3   tubby like protein 3    0.0009584191021333225
GO:0007420: brain development   ZNF335  zinc finger protein 335 -0.000365528325018728
GO:0009100: glycoprotein metabolic process      AGER    advanced glycosylation end product-specific r
eceptor -0.00014834649160886415
GO:0009750: response to fructose        AGER    advanced glycosylation end product-specific receptor
-0.00014834649160886415
GO:0010508: positive regulation of autophagy    AGER    advanced glycosylation end product-specific r
eceptor -0.0001748070748723673
GO:0010508: positive regulation of autophagy    BNIP3   BCL2/adenovirus E1B 19kDa interacting protein
3       0.002929798361141282
GO:0010508: positive regulation of autophagy    FOXO1   forkhead box O1 0.0018100350428352063
GO:0010508: positive regulation of autophagy    XBP1    X-box binding protein 1 0.0002673288898503275
GO:0010718: positive regulation of epithelial to mesenchymal transition AGER    advanced glycosylatio
n end product-specific receptor -0.00017412770780643642
GO:0010718: positive regulation of epithelial to mesenchymal transition ALX1    ALX homeobox 1  0.002
2896753805271277
GO:0010718: positive regulation of epithelial to mesenchymal transition COL1A1  collagen, type I, alp
ha 1    -0.0005260914185390754
GO:0010718: positive regulation of epithelial to mesenchymal transition CTNNB1  catenin (cadherin-ass
ociated protein), beta 1, 88kDa -0.00011654993116169157
GO:0010718: positive regulation of epithelial to mesenchymal transition EZH2    enhancer of zeste 2 p
olycomb repressive complex 2 subunit    -0.0001356722093780021
GO:0010718: positive regulation of epithelial to mesenchymal transition GCNT2   glucosaminyl (N-acety
l) transferase 2, I-branching enzyme (I blood group)    0.0024294293566783534
GO:0010718: positive regulation of epithelial to mesenchymal transition HDAC2   histone deacetylase 2
-0.0012052943575696273
GO:0010718: positive regulation of epithelial to mesenchymal transition LEF1    lymphoid enhancer-bin
ding factor 1   -0.0001004325859400352
GO:0010718: positive regulation of epithelial to mesenchymal transition NOTCH1  notch 1 0.00051854533
7136905
GO:0010718: positive regulation of epithelial to mesenchymal transition TGFB1   transforming growth f
actor, beta 1   -7.313794013420128e-5
GO:0010718: positive regulation of epithelial to mesenchymal transition TGFB2   transforming growth f
actor, beta 2   -0.0010581970171750206
GO:0010718: positive regulation of epithelial to mesenchymal transition TGFB3   transforming growth f
actor, beta 3   -0.0018222936478021797
GO:0010718: positive regulation of epithelial to mesenchymal transition TWIST1  twist family bHLH tra
nscription factor 1     -0.001340089402570324
GO:0010718: positive regulation of epithelial to mesenchymal transition WWTR1   WW domain containing
transcription regulator 1      0.0009000166833594047
GO:0010763: positive regulation of fibroblast migration AGER    advanced glycosylation end product-sp
ecific receptor -0.00017177467981926522
GO:0010763: positive regulation of fibroblast migration AKT1    v-akt murine thymoma viral oncogene h
omolog 1        0.0007084604504139823
GO:0010763: positive regulation of fibroblast migration TGFB1   transforming growth factor, beta 1
-6.900388222676307e-5
GO:0010763: positive regulation of fibroblast migration THBS1   thrombospondin 1        -0.0010069743
726467085
GO:0014823: response to activity        AGER    advanced glycosylation end product-specific receptor
-0.00017291152542450692
GO:0014823: response to activity        CCL2    chemokine (C-C motif) ligand 2  0.000802419071465806
GO:0014823: response to activity        MAS1    MAS1 proto-oncogene, G protein-coupled receptor 0.000
29584200617529925
GO:0014823: response to activity        PRKDC   protein kinase, DNA-activated, catalytic polypeptide
-0.0018321034931001409
GO:0014823: response to activity        PTN     pleiotrophin    0.00029075271927588573
GO:0014823: response to activity        STAR    steroidogenic acute regulatory protein  0.00087860704
66180656
GO:0014823: response to activity        TH      tyrosine hydroxylase    -0.00033916097826923106
GO:0014911: positive regulation of smooth muscle cell migration AGER    advanced glycosylation end pr
oduct-specific receptor -0.00016892625099548163
GO:0014911: positive regulation of smooth muscle cell migration BCL2    B-cell CLL/lymphoma 2   -6.25
35302081430594e-6
GO:0014911: positive regulation of smooth muscle cell migration IGF1    insulin-like growth factor 1
(somatomedin C)        0.00011656591892340848
GO:0014911: positive regulation of smooth muscle cell migration NRP1    neuropilin 1    -0.0006070586
606722714
GO:0014911: positive regulation of smooth muscle cell migration PDGFRB  platelet-derived growth facto
r receptor, beta polypeptide    -0.0003310873939760629
GO:0030324: lung development    AGER    advanced glycosylation end product-specific receptor    -0.00
017443942649393202
GO:0030324: lung development    ALDH1A2 aldehyde dehydrogenase 1 family, member A2      -0.0030238104
633179795
GO:0030324: lung development    DICER1  dicer 1, ribonuclease type III  -1.598508287682783e-5
GO:0030324: lung development    FGFR2   fibroblast growth factor receptor 2     0.00076443047210283
GO:0030324: lung development    GLI1    GLI family zinc finger 1        -0.0012985894525954192
GO:0030324: lung development    GLI2    GLI family zinc finger 2        0.0018565882658731474
GO:0030324: lung development    GLI3    GLI family zinc finger 3        -0.002154876813410871
GO:0030324: lung development    HES1    hes family bHLH transcription factor 1  -0.000912387133289415

GO:0030324: lung development    ITGA3   integrin, alpha 3 (antigen CD49C, alpha 3 subunit of VLA-3 receptor) 0.0014022278178806888
GO:0030324: lung development    JMJD6   jumonji domain containing 6     0.003596253880307908
GO:0030324: lung development    LOX     lysyl oxidase   -0.0005774279246892739
GO:0030324: lung development    NOTCH1  notch 1 0.0005194326424966054
GO:0030324: lung development    PROX1   prospero homeobox 1     0.0011233759559207202
GO:0030324: lung development    PTN     pleiotrophin    0.0003025318227062162
GO:0030324: lung development    SHH     sonic hedgehog  0.0006010797438065888
GO:0030324: lung development    STRA6   stimulated by retinoic acid 6   -0.001683118619888543
GO:0030324: lung development    TMBIM6  transmembrane BAX inhibitor motif containing 6  -0.00014760836380781412
GO:0030324: lung development    VEGFA   vascular endothelial growth factor A    0.0005959007297280883
GO:0030324: lung development    WNT5A   wingless-type MMTV integration site family, member 5A   -0.0006684151812011535
GO:0031175: neuron projection development       AGER    advanced glycosylation end product-specific receptor -0.00017444118644246632
GO:0031175: neuron projection development       AREG    amphiregulin    0.0024601188297535635
GO:0031175: neuron projection development       GDNF    glial cell derived neurotrophic factor  0.00044549446832151346
GO:0031175: neuron projection development       HMGB1   high mobility group box 1       -0.0007779385934992564
GO:0031175: neuron projection development       LYN     LYN proto-oncogene, Src family tyrosine kinase       -0.0014223060883479437
GO:0031175: neuron projection development       NEDD4   neural precursor cell expressed, developmentally down-regulated 4, E3 ubiquitin protein ligase       0.0022661973448634093
GO:0031175: neuron projection development       PHGDH   phosphoglycerate dehydrogenase  0.00029135468481888597
GO:0031175: neuron projection development       RB1     retinoblastoma 1        -0.0014955249723514231
GO:0031175: neuron projection development       STMN1   stathmin 1      0.0005586829782879251
GO:0031175: neuron projection development       STMN2   stathmin 2      -0.0019494785260562404
GO:0031175: neuron projection development       STMN3   stathmin-like 3 0.00259343418697851
GO:0031175: neuron projection development       STMN4   stathmin-like 4 0.0003485622681119767
GO:0032966: negative regulation of collagen biosynthetic process        AGER    advanced glycosylation end product-specific receptor -0.00014834649160886415
GO:0033189: response to vitamin A       AGER    advanced glycosylation end product-specific receptor -0.00017482761798466026
GO:0033189: response to vitamin A       ALDH1A2 aldehyde dehydrogenase 1 family, member A2      -0.0030371512431999165
GO:0033189: response to vitamin A       GATA4   GATA binding protein 4  -0.0011019367534209122
GO:0033189: response to vitamin A       PITX2   paired-like homeodomain 2       0.0021901196203252256
GO:0033189: response to vitamin A       RXRA    retinoid X receptor, alpha      0.0011218503108525505
GO:0033189: response to vitamin A       TYMS    thymidylate synthetase  0.0015738413411606836
GO:0033595: response to genistein       AGER    advanced glycosylation end product-specific receptor -0.00014834649160886415
GO:0033689: negative regulation of osteoblast proliferation     AGER    advanced glycosylation end product-specific receptor -0.0001737606478537304
GO:0033689: negative regulation of osteoblast proliferation     BCL2    B-cell CLL/lymphoma 2   -5.531878170583663e-6
GO:0033689: negative regulation of osteoblast proliferation     GREM1   gremlin 1, DAN family BMP antagonist -0.0008254190077890399
GO:0033689: negative regulation of osteoblast proliferation     NELL1   NEL-like 1 (chicken)    0.001528246646504543
GO:0033689: negative regulation of osteoblast proliferation     SFRP1   secreted frizzled-related protein 1  0.0012792945079847498
GO:0035690: cellular response to drug   AGER    advanced glycosylation end product-specific receptor -0.000174049047621811
GO:0035690: cellular response to drug   CCL2    chemokine (C-C motif) ligand 2  0.0008138434299684032
GO:0035690: cellular response to drug   EGR1    early growth response 1 0.0010900244760895233
GO:0035690: cellular response to drug   KCNE2   potassium voltage-gated channel, Isk-related family, member 2       -0.0005740932946616276
GO:0035690: cellular response to drug   MEF2C   myocyte enhancer factor 2C      0.0009698909105021104
GO:0035690: cellular response to drug   MYC     v-myc avian myelocytomatosis viral oncogene homolog -0.00113443138895615
GO:0035690: cellular response to drug   QDPR    quinoid dihydropteridine reductase      -0.00284750467888784
GO:0035690: cellular response to drug   TFRC    transferrin receptor    0.001081740301690953
GO:0035690: cellular response to drug   TH      tyrosine hydroxylase    -0.00034158854759942244
GO:0035690: cellular response to drug   TP53    tumor protein p53       0.001170761336715618
GO:0043507: positive regulation of JUN kinase activity  AGER    advanced glycosylation end product-specific receptor -0.00017483046779262425
GO:0043507: positive regulation of JUN kinase activity  MAP3K5  mitogen-activated protein kinase kinase kinase 5     0.00034513897915360806
GO:0043507: positive regulation of JUN kinase activity  PAK1    p21 protein (Cdc42/Rac)-activated kinase 1   -0.0022036882015784498
GO:0043507: positive regulation of JUN kinase activity  TNFRSF11A       tumor necrosis factor receptor superfamily, member 11a, NFKB activator       0.0027958343078884375
GO:0043525: positive regulation of neuron apoptotic process     AGER    advanced glycosylation end product-specific receptor -0.00017316224415798743
GO:0043525: positive regulation of neuron apoptotic process     ASCL1   achaete-scute family bHLH transcription factor 1     -0.0014125259486000286

GO:0043525: positive regulation of neuron apoptotic process     BAX     BCL2-associated X protein
-0.00042247701907483393
GO:0043525: positive regulation of neuron apoptotic process     BCL2L11 BCL2-like 11 (apoptosis facil
itator) -0.0007689878285913827
GO:0043525: positive regulation of neuron apoptotic process     CDC34   cell division cycle 34  -0.00
016003391003193722
GO:0043525: positive regulation of neuron apoptotic process     CDC42   cell division cycle 42  0.001
2258436174132363
GO:0043525: positive regulation of neuron apoptotic process     EGR1    early growth response 1 0.001
0810191041238106
GO:0043525: positive regulation of neuron apoptotic process     FGFR3   fibroblast growth factor rece
ptor 3  0.00021964963809514143
GO:0043525: positive regulation of neuron apoptotic process     FOXO3   forkhead box O3 0.00118482002
39213142
GO:0043525: positive regulation of neuron apoptotic process     GRIK5   glutamate receptor, ionotropi
c, kainate 5    -0.00023169200191268337
GO:0043525: positive regulation of neuron apoptotic process     PAK3    p21 protein (Cdc42/Rac)-activ
ated kinase 3   -0.0012475001292670772
GO:0043525: positive regulation of neuron apoptotic process     TFAP2A  transcription factor AP-2 alp
ha (activating enhancer binding protein 2 alpha)        0.0004763184137753916
GO:0043525: positive regulation of neuron apoptotic process     TGFB2   transforming growth factor, b
eta 2   -0.00104532887082841
GO:0043525: positive regulation of neuron apoptotic process     TP53    tumor protein p53       0.001
1571306736293352
GO:0048146: positive regulation of fibroblast proliferation     AGER    advanced glycosylation end pr
oduct-specific receptor -0.00017375890780748623
GO:0048146: positive regulation of fibroblast proliferation     AGT     angiotensinogen (serpin pepti
dase inhibitor, clade A, member 8)      -0.0011236518161791584
GO:0048146: positive regulation of fibroblast proliferation     CCNB1   cyclin B1       -0.0008795881
028420063
GO:0048146: positive regulation of fibroblast proliferation     E2F1    E2F transcription factor 1
0.002085148869595043
GO:0048146: positive regulation of fibroblast proliferation     EGFR    epidermal growth factor recep
tor     0.0006800807786013381
GO:0048146: positive regulation of fibroblast proliferation     ESR1    estrogen receptor 1     -0.00
09415233693240736
GO:0048146: positive regulation of fibroblast proliferation     IGF1    insulin-like growth factor 1
 (somatomedin C)        0.00012893953284377714
GO:0048146: positive regulation of fibroblast proliferation     MYC     v-myc avian myelocytomatosis
 viral oncogene homolog -0.0011312139237075827
GO:0048146: positive regulation of fibroblast proliferation     PML     promyelocytic leukemia  -0.00
06776744080157952
GO:0048146: positive regulation of fibroblast proliferation     PRKDC   protein kinase, DNA-activate
d, catalytic polypeptide        -0.0018460968198891938
GO:0048146: positive regulation of fibroblast proliferation     RNASEH2B        ribonuclease H2, subu
nit B   -4.26625034167724e-5
GO:0048146: positive regulation of fibroblast proliferation     SPHK1   sphingosine kinase 1    0.001
8002259895043603
GO:0048146: positive regulation of fibroblast proliferation     TGFB1   transforming growth factor, b
eta 1   -7.21400493849189e-5
GO:0048146: positive regulation of fibroblast proliferation     WNT1    wingless-type MMTV integratio
n site family, member 1 0.0007827611742660043
GO:0048146: positive regulation of fibroblast proliferation     WNT5A   wingless-type MMTV integratio
n site family, member 5A        -0.0006631535226664452
GO:0048661: positive regulation of smooth muscle cell proliferation     AGER    advanced glycosylatio
n end product-specific receptor -0.0001736169086333461
GO:0048661: positive regulation of smooth muscle cell proliferation     ALOX12  arachidonate 12-lipox
ygenase -0.001940539874844371
GO:0048661: positive regulation of smooth muscle cell proliferation     BMP4    bone morphogenetic pr
otein 4 -0.0003204751497244805
GO:0048661: positive regulation of smooth muscle cell proliferation     CX3CL1  chemokine (C-X3-C mot
if) ligand 1    0.002100798079705573
GO:0048661: positive regulation of smooth muscle cell proliferation     FGFR2   fibroblast growth fac
tor receptor 2  0.00075776608380576
GO:0048661: positive regulation of smooth muscle cell proliferation     HMOX1   heme oxygenase (decyc
ling) 1 -0.00021587041829301996
GO:0048661: positive regulation of smooth muscle cell proliferation     ID2     inhibitor of DNA bind
ing 2, dominant negative helix-loop-helix protein       6.0341930686014506e-5
GO:0048661: positive regulation of smooth muscle cell proliferation     IGF1    insulin-like growth f
actor 1 (somatomedin C) 0.00012889561004382225
GO:0048661: positive regulation of smooth muscle cell proliferation     IRAK1   interleukin-1 recepto
r-associated kinase 1   -0.001597864610406035
GO:0048661: positive regulation of smooth muscle cell proliferation     ORC1    origin recognition co
mplex, subunit 1        -0.0007351319920685377
GO:0048661: positive regulation of smooth muscle cell proliferation     PDGFRB  platelet-derived grow
th factor receptor, beta polypeptide    -0.00035670154597099634
GO:0048661: positive regulation of smooth muscle cell proliferation     SERPINF2        serpin peptid
ase inhibitor, clade F (alpha-2 antiplasmin, pigment epithelium derived factor), member 2       0.000
6549047756364493
GO:0048661: positive regulation of smooth muscle cell proliferation     SKP2    S-phase kinase-associ
ated protein 2, E3 ubiquitin protein ligase     -0.0005360494011624165

GO:0050729: positive regulation of inflammatory response        AGER    advanced glycosylation end pr
oduct-specific receptor -0.0001736865309078786
GO:0050729: positive regulation of inflammatory response        AGT     angiotensinogen (serpin pepti
dase inhibitor, clade A, member 8)      -0.0011252073986677228
GO:0050729: positive regulation of inflammatory response        AGTR1   angiotensin II receptor, type
1       0.0003474740820156128
GO:0050729: positive regulation of inflammatory response        CX3CL1  chemokine (C-X3-C motif) liga
nd 1    0.0021041179405726627
GO:0050729: positive regulation of inflammatory response        IL12B   interleukin 12B 0.00127146959
73251736
GO:0050729: positive regulation of inflammatory response        JAK2    Janus kinase 2  -3.2577995143
77629e-5
GO:0050729: positive regulation of inflammatory response        PRKCA   protein kinase C, alpha -5.92
56636190428846e-6
GO:0050729: positive regulation of inflammatory response        SERPINE1        serpin peptidase inhi
bitor, clade E (nexin, plasminogen activator inhibitor type 1), member 1        0.0001126160737699245
7
GO:0050729: positive regulation of inflammatory response        STAT5A  signal transducer and activat
or of transcription 5A  0.0015882386383049955
GO:0050729: positive regulation of inflammatory response        WNT5A   wingless-type MMTV integratio
n site family, member 5A        -0.0006636463894865768
GO:0050930: induction of positive chemotaxis    AGER    advanced glycosylation end product-specific r
eceptor -0.00017447057463708084
GO:0050930: induction of positive chemotaxis    AZU1    azurocidin 1    0.00019835981167533648
GO:0050930: induction of positive chemotaxis    CXCL12  chemokine (C-X-C motif) ligand 12       -0.00
12109384003218438
GO:0050930: induction of positive chemotaxis    PRKCA   protein kinase C, alpha -5.8755268757426515e-
6
GO:0050930: induction of positive chemotaxis    VEGFA   vascular endothelial growth factor A    0.000
5965945992619954
GO:0050930: induction of positive chemotaxis    VEGFC   vascular endothelial growth factor C    -0.00
33707730129195765
GO:0051092: positive regulation of NF-kappaB transcription factor activity      AGER    advanced glyc
osylation end product-specific receptor -0.00017432335669368248
GO:0051092: positive regulation of NF-kappaB transcription factor activity      AGT     angiotensinog
en (serpin peptidase inhibitor, clade A, member 8)      -0.0011337690988374248
GO:0051092: positive regulation of NF-kappaB transcription factor activity      AR      androgen rece
ptor    0.0026406868806194955
GO:0051092: positive regulation of NF-kappaB transcription factor activity      CIB1    calcium and i
ntegrin binding 1 (calmyrin)    5.504244238658219e-5
GO:0051092: positive regulation of NF-kappaB transcription factor activity      CLOCK   clock circadi
an regulator    0.00020029395010825264
GO:0051092: positive regulation of NF-kappaB transcription factor activity      CTH     cystathionine
gamma-lyase     0.0017494893888894
GO:0051092: positive regulation of NF-kappaB transcription factor activity      EDA     ectodysplasin
A       -0.0008150419386035184
GO:0051092: positive regulation of NF-kappaB transcription factor activity      GREM1   gremlin 1, DA
N family BMP antagonist -0.0008278792419378842
GO:0051092: positive regulation of NF-kappaB transcription factor activity      ICAM1   intercellular
adhesion molecule 1     0.0007295545166121158
GO:0051092: positive regulation of NF-kappaB transcription factor activity      IRAK1   interleukin-1
receptor-associated kinase 1    -0.0016102921022455629
GO:0051092: positive regulation of NF-kappaB transcription factor activity      LGALS9  lectin, galac
toside-binding, soluble, 9      -0.00011687984728779951
GO:0051092: positive regulation of NF-kappaB transcription factor activity      PRKCZ   protein kinas
e C, zeta       -0.0016019594428588926
GO:0051092: positive regulation of NF-kappaB transcription factor activity      SPHK1   sphingosine k
inase 1 0.001811821086001211
GO:0051092: positive regulation of NF-kappaB transcription factor activity      TGFB1   transforming
growth factor, beta 1  -7.288180145474266e-5
GO:0051092: positive regulation of NF-kappaB transcription factor activity      TNFRSF11A       tumor
necrosis factor receptor superfamily, member 11a, NFKB activator        0.0027777048444944798
GO:0051092: positive regulation of NF-kappaB transcription factor activity      WNT5A   wingless-type
MMTV integration site family, member 5A -0.0006677078796261199
GO:0051101: regulation of DNA binding   AGER    advanced glycosylation end product-specific receptor
-0.0001728263481397249
GO:0051101: regulation of DNA binding   CSNK2B  casein kinase 2, beta polypeptide       0.00150495023
58291903
GO:0051101: regulation of DNA binding   HJURP   Holliday junction recognition protein   -7.6928380050
05403e-5
GO:0051101: regulation of DNA binding   TGFB1   transforming growth factor, beta 1      -7.1443911215
37735e-5
GO:0051595: response to methylglyoxal   AGER    advanced glycosylation end product-specific receptor
-0.00014834649160886415
GO:0055074: calcium ion homeostasis     AGER    advanced glycosylation end product-specific receptor
-0.0001738194344385704
GO:0055074: calcium ion homeostasis     CAV1    caveolin 1, caveolae protein, 22kDa     -0.0005338583
631218383
GO:0055074: calcium ion homeostasis     CYP27B1 cytochrome P450, family 27, subfamily B, polypeptide
1      -0.0004686574635031665
GO:0055074: calcium ion homeostasis     S100A14 S100 calcium binding protein A14        -0.0021505132

543200903
GO:0055074: calcium ion homeostasis WFS1 Wolfram syndrome 1 (wolframin) 0.0005753734761084318
GO:0055093: response to hyperoxia AGER advanced glycosylation end product-specific receptor -0.0001744937618179712
GO:0055093: response to hyperoxia BNIP3 BCL2/adenovirus E1B 19kDa interacting protein 3 0.002921075434424916
GO:0055093: response to hyperoxia COL1A1 collagen, type I, alpha 1 -0.0005294384627999855
GO:0055093: response to hyperoxia HDAC2 histone deacetylase 2 -0.001208643089586805
GO:0055093: response to hyperoxia PDGFRB platelet-derived growth factor receptor, beta polypeptide -0.00036376425484009546
GO:0060100: positive regulation of phagocytosis, engulfment AGER advanced glycosylation end product-specific receptor -0.00017437269042957547
GO:0060100: positive regulation of phagocytosis, engulfment GATA2 GATA binding protein 2 -0.00045139068839606143
GO:0060100: positive regulation of phagocytosis, engulfment STAP1 signal transducing adaptor family member 1 0.00244564877336297
GO:0060290: transdifferentiation AGER advanced glycosylation end product-specific receptor -0.00017186886896242517
GO:0060290: transdifferentiation GATA4 GATA binding protein 4 -0.0010650683605545053
GO:0060290: transdifferentiation PDX1 pancreatic and duodenal homeobox 1 0.00024676608728110697
GO:0070301: cellular response to hydrogen peroxide AGER advanced glycosylation end product-specific receptor -0.00017398449921313602
GO:0070301: cellular response to hydrogen peroxide BNIP3 BCL2/adenovirus E1B 19kDa interacting protein 3 0.002901768597232071
GO:0070301: cellular response to hydrogen peroxide CST3 cystatin C -6.895543295744764e-5
GO:0070301: cellular response to hydrogen peroxide CYP1B1 cytochrome P450, family 1, subfamily B, polypeptide 1 0.0003967770016374729
GO:0070301: cellular response to hydrogen peroxide ECT2 epithelial cell transforming 2 0.0010582300901999435
GO:0070301: cellular response to hydrogen peroxide EZH2 enhancer of zeste 2 polycomb repressive complex 2 subunit -0.00013681638465076716
GO:0070301: cellular response to hydrogen peroxide HDAC2 histone deacetylase 2 -0.001203910021992933
GO:0070301: cellular response to hydrogen peroxide KLF2 Kruppel-like factor 2 -0.0012287365066860787
GO:0070301: cellular response to hydrogen peroxide MAP3K5 mitogen-activated protein kinase kinase kinase 5 0.0003410789784105757
GO:0070301: cellular response to hydrogen peroxide PAX2 paired box 2 -0.001607723892580603
GO:0070301: cellular response to hydrogen peroxide SIRT1 sirtuin 1 -2.3537799167040125e-6
GO:0070301: cellular response to hydrogen peroxide TNFAIP3 tumor necrosis factor, alpha-induced protein 3 0.0011126139106054412
GO:0071333: cellular response to glucose stimulus AGER advanced glycosylation end product-specific receptor -0.00017279936772989385
GO:0071333: cellular response to glucose stimulus GATA4 GATA binding protein 4 -0.0010778050662791766
GO:0071333: cellular response to glucose stimulus GRIK5 glutamate receptor, ionotropic, kainate 5 -0.000231424604226445
GO:0071333: cellular response to glucose stimulus ICAM1 intercellular adhesion molecule 1 0.000722410998062139
GO:0071333: cellular response to glucose stimulus PAX2 paired box 2 -0.0015880765090634655
GO:0071333: cellular response to glucose stimulus PDK3 pyruvate dehydrogenase kinase, isozyme 3 0.0007545067307062336
GO:0071333: cellular response to glucose stimulus SOX4 SRY (sex determining region Y)-box 4 -4.09899900218994e-5
GO:0071333: cellular response to glucose stimulus STAR steroidogenic acute regulatory protein 0.000879486952714341
GO:0071333: cellular response to glucose stimulus TH tyrosine hydroxylase -0.00033880877546808055
GO:0071333: cellular response to glucose stimulus XBP1 X-box binding protein 1 0.00026540600671913294
GO:0071398: cellular response to fatty acid AGER advanced glycosylation end product-specific receptor -0.0001740364875257397
GO:0071398: cellular response to fatty acid CCNB1 cyclin B1 -0.0008811124135056447
GO:0071398: cellular response to fatty acid CPT1A carnitine palmitoyltransferase 1A (liver) 0.0012703301162785693
GO:0071398: cellular response to fatty acid E2F1 E2F transcription factor 1 0.002091937316160375
GO:0071398: cellular response to fatty acid PDK3 pyruvate dehydrogenase kinase, isozyme 3 0.0007692072096432322
GO:0071407: cellular response to organic cyclic compound AGER advanced glycosylation end product-specific receptor -0.00017355911864847432
GO:0071407: cellular response to organic cyclic compound AKT1 v-akt murine thymoma viral oncogene homolog 1 0.0007233967173948213
GO:0071407: cellular response to organic cyclic compound AKT2 v-akt murine thymoma viral oncogene homolog 2 -0.0006222397836267628
GO:0071407: cellular response to organic cyclic compound AXIN1 axin 1 -0.0007244654666458465

GO:0071407: cellular response to organic cyclic compound        CCL2    chemokine (C-C motif) ligand
 2      0.0008079755974389148
GO:0071407: cellular response to organic cyclic compound        CCNB1   cyclin B1       -0.0008807212
614018302
GO:0071407: cellular response to organic cyclic compound        CYP1B1  cytochrome P450, family 1, su
bfamily B, polypeptide 1        0.0003948647056333417
GO:0071407: cellular response to organic cyclic compound        GLI2    GLI family zinc finger 2
0.0018375816077707459
GO:0071407: cellular response to organic cyclic compound        MGMT    O-6-methylguanine-DNA methylt
ransferase      0.0004081079628331256
GO:0071407: cellular response to organic cyclic compound        PAK1    p21 protein (Cdc42/Rac)-activ
ated kinase 1   -0.0021684321580586097
GO:0071407: cellular response to organic cyclic compound        PAK3    p21 protein (Cdc42/Rac)-activ
ated kinase 3   -0.0012506631005173505
GO:0071407: cellular response to organic cyclic compound        TGFB1   transforming growth factor, b
eta 1   -7.087309899284884e-5
GO:0072657: protein localization to membrane    AGER    advanced glycosylation end product-specific r
eceptor -0.00017484498155257284
GO:0072657: protein localization to membrane    CPE     carboxypeptidase E      -0.002559902518296103
GO:0072714: response to selenite ion    AGER    advanced glycosylation end product-specific receptor
 -0.00014834649160886415
GO:1901018: positive regulation of potassium ion transmembrane transporter activity     AGER    advan
ced glycosylation end product-specific receptor -0.00017284645634956522
GO:1901018: positive regulation of potassium ion transmembrane transporter activity     ATP1B3  ATPas
e, Na+/K+ transporting, beta 3 polypeptide      -0.0014828321350051443
GO:2000353: positive regulation of endothelial cell apoptotic process   AGER    advanced glycosylatio
n end product-specific receptor -0.00017100918134362427
GO:2000353: positive regulation of endothelial cell apoptotic process   BMP4    bone morphogenetic pr
otein 4 -0.00031126678406137045
GO:2000353: positive regulation of endothelial cell apoptotic process   THBS1   thrombospondin 1
 -0.0009981485010827531
GO:2000353: positive regulation of endothelial cell apoptotic process   XBP1    X-box binding protein
1       0.0002619672047020384
GO:2000379: positive regulation of reactive oxygen species metabolic process    AGER    advanced glyc
osylation end product-specific receptor -0.00017445715956479607
GO:2000379: positive regulation of reactive oxygen species metabolic process    AGT     angiotensinog
en (serpin peptidase inhibitor, clade A, member 8)      -0.001139848434904905
GO:2000379: positive regulation of reactive oxygen species metabolic process    AGTR1   angiotensin I
I receptor, type 1      0.0003468076530572347
GO:2000379: positive regulation of reactive oxygen species metabolic process    GRB2    growth factor
receptor-bound protein 2        0.00043099356827873116
GO:2000379: positive regulation of reactive oxygen species metabolic process    LEP     leptin  0.003
2042520517062507
GO:2000379: positive regulation of reactive oxygen species metabolic process    PDGFRB  platelet-deri
ved growth factor receptor, beta polypeptide    -0.00036446269083052963
GO:2000379: positive regulation of reactive oxygen species metabolic process    THBS1   thrombospondi
n 1     -0.0010398131143783795
GO:2000379: positive regulation of reactive oxygen species metabolic process    TP53    tumor protein
p53     0.0011816822614750906
GO:2000676: positive regulation of type B pancreatic cell apoptotic process     AGER    advanced glyc
osylation end product-specific receptor -0.00014834649160886415
GO:0010628: positive regulation of gene expression      AGR2    anterior gradient 2     0.00163102585
40125503
GO:0010628: positive regulation of gene expression      ALDH1A2 aldehyde dehydrogenase 1 family, memb
er A2   -0.0030214035597184883
GO:0010628: positive regulation of gene expression      ALOX12  arachidonate 12-lipoxygenase    -0.00
19549918846050425
GO:0010628: positive regulation of gene expression      ANK3    ankyrin 3, node of Ranvier (ankyrin
 G)     0.0007006326936388541
GO:0010628: positive regulation of gene expression      AR      androgen receptor       0.00264130702
73786825
GO:0010628: positive regulation of gene expression      AVP     arginine vasopressin    -0.0009377555
621954948
GO:0010628: positive regulation of gene expression      AZU1    azurocidin 1    0.0001982533945105536
3
GO:0010628: positive regulation of gene expression      CAV1    caveolin 1, caveolae protein, 22kDa
 -0.0005358314464250328
GO:0010628: positive regulation of gene expression      CDC42   cell division cycle 42  0.00124141809
09225774
GO:0010628: positive regulation of gene expression      CDH3    cadherin 3, type 1, P-cadherin (place
ntal)   -0.0012325089245543426
GO:0010628: positive regulation of gene expression      CITED1  Cbp/p300-interacting transactivator,
 with Glu/Asp-rich carboxy-terminal domain, 1   0.002820644403418464
GO:0010628: positive regulation of gene expression      DICER1  dicer 1, ribonuclease type III  -1.59
39035530301243e-5
GO:0010628: positive regulation of gene expression      E2F1    E2F transcription factor 1      0.002
0989260280933333
GO:0010628: positive regulation of gene expression      FGF8    fibroblast growth factor 8 (androgen-
induced)        0.000984587459185798
GO:0010628: positive regulation of gene expression      GJA1    gap junction protein, alpha 1, 43kDa
 -0.00016104021761663878

GO:0010628: positive regulation of gene expression HINFP histone H4 transcription factor 0.000
9927224392016238
GO:0010628: positive regulation of gene expression HMGA2 high mobility group AT-hook 2 0.001
510370069653839
GO:0010628: positive regulation of gene expression HPN hepsin 0.0031614623473557807
GO:0010628: positive regulation of gene expression ID2 inhibitor of DNA binding 2, dominant
negative helix-loop-helix protein 6.018731043526236e-5
GO:0010628: positive regulation of gene expression IFNG interferon, gamma -4.9042182046
04563e-5
GO:0010628: positive regulation of gene expression INHBA inhibin, beta A -0.001353712520955763
GO:0010628: positive regulation of gene expression ITGA3 integrin, alpha 3 (antigen CD49C, alp
ha 3 subunit of VLA-3 receptor) 0.001401477696653707
GO:0010628: positive regulation of gene expression LEF1 lymphoid enhancer-binding factor 1
-0.00010009965043653972
GO:0010628: positive regulation of gene expression MED1 mediator complex subunit 1 0.001
1297923713243696
GO:0010628: positive regulation of gene expression MEF2C myocyte enhancer factor 2C 0.000
9740408352369696
GO:0010628: positive regulation of gene expression PAX6 paired box 6 0.0019366682096019544
GO:0010628: positive regulation of gene expression PIK3CD phosphatidylinositol-4,5-bisphosphate
3-kinase, catalytic subunit delta -0.0007814128934088676
GO:0010628: positive regulation of gene expression SMO smoothened, frizzled class receptor
0.0021407068134982185
GO:0010628: positive regulation of gene expression SOX11 SRY (sex determining region Y)-box 11
-0.00021004944823715404
GO:0010628: positive regulation of gene expression STAP1 signal transducing adaptor family mem
ber 1 0.0024454884414603626
GO:0010628: positive regulation of gene expression STAR steroidogenic acute regulatory protei
n 0.0008950555723609031
GO:0010628: positive regulation of gene expression TFAP2A transcription factor AP-2 alpha (acti
vating enhancer binding protein 2 alpha) 0.00048047176610611475
GO:0010628: positive regulation of gene expression TGFB1 transforming growth factor, beta 1
-7.277166689020003e-5
GO:0010628: positive regulation of gene expression TGFB2 transforming growth factor, beta 2
-0.0010601601271692355
GO:0010628: positive regulation of gene expression TGFBR1 transforming growth factor, beta rece
ptor 1 0.00034000856531030057
GO:0010628: positive regulation of gene expression TNC tenascin C 0.0007351709405452431
GO:0010628: positive regulation of gene expression TP53 tumor protein p53 0.00117625292
07385668
GO:0010628: positive regulation of gene expression TWIST1 twist family bHLH transcription facto
r 1 -0.0013424544095350145
GO:0010628: positive regulation of gene expression VDR vitamin D (1,25- dihydroxyvitamin D3)
receptor 0.0005641977550405355
GO:0010628: positive regulation of gene expression VEGFA vascular endothelial growth factor A
0.00059481865849399
GO:0010628: positive regulation of gene expression ZPR1 ZPR1 zinc finger -0.0002100850
547252754
GO:0010811: positive regulation of cell-substrate adhesion AGR2 anterior gradient 2 0.001
6160594067224795
GO:0010811: positive regulation of cell-substrate adhesion EGFL6 EGF-like-domain, multiple 6
0.00013387785099819088
GO:0010811: positive regulation of cell-substrate adhesion ITGA3 integrin, alpha 3 (antigen CD
49C, alpha 3 subunit of VLA-3 receptor) 0.0013851329370988385
GO:0010811: positive regulation of cell-substrate adhesion JAK2 Janus kinase 2 -3.5414436133
21719e-5
GO:0010811: positive regulation of cell-substrate adhesion PTN pleiotrophin 0.00029182850
01311842
GO:0010811: positive regulation of cell-substrate adhesion THBS1 thrombospondin 1 -0.00
10220888368528103
GO:0045742: positive regulation of epidermal growth factor receptor signaling pathway AGR2 anter
ior gradient 2 0.0016287095961735926
GO:0045742: positive regulation of epidermal growth factor receptor signaling pathway AGT angio
tensinogen (serpin peptidase inhibitor, clade A, member 8) -0.001138511708397122
GO:0048546: digestive tract morphogenesis AGR2 anterior gradient 2 0.0016204944302982797
GO:0048546: digestive tract morphogenesis BBS7 Bardet-Biedl syndrome 7 -0.000971161426819931
1
GO:0048546: digestive tract morphogenesis BCL2 B-cell CLL/lymphoma 2 -5.614563355390727e-6
GO:0048546: digestive tract morphogenesis EGFR epidermal growth factor receptor 0.000
6825754042497429
GO:0048546: digestive tract morphogenesis EPHB3 EPH receptor B3 0.000881300789621169
GO:0048546: digestive tract morphogenesis FGFR3 fibroblast growth factor receptor 3 0.000
220799334068332
GO:0048546: digestive tract morphogenesis GLI1 GLI family zinc finger 1 -0.0012917663
02057122
GO:0048546: digestive tract morphogenesis HIF1A hypoxia inducible factor 1, alpha subunit (ba
sic helix-loop-helix transcription factor) -0.000643431109054301
GO:0048546: digestive tract morphogenesis SFRP1 secreted frizzled-related protein 1 0.001
2754926175704388
GO:0048546: digestive tract morphogenesis SOX10 SRY (sex determining region Y)-box 10 0.000
18939330145281624

GO:0048546: digestive tract morphogenesis	STRA6	stimulated by retinoic acid 6	-0.0016729989147031131
GO:0048546: digestive tract morphogenesis	STX2	syntaxin 2	0.0003062489153723763
GO:0048546: digestive tract morphogenesis	TP73	tumor protein p73	0.001022991935448292
GO:0048639: positive regulation of developmental growth AGR2	anterior gradient 2	0.001637915018435805
GO:0048639: positive regulation of developmental growth C3	complement component 3	0.0020417287878284718
GO:0048639: positive regulation of developmental growth INSR	insulin receptor	-0.0013755111282643966
GO:0048639: positive regulation of developmental growth LEP	leptin	0.0032210207512611143
GO:0048639: positive regulation of developmental growth PLCB1	phospholipase C, beta 1 (phosphoinositide-specific)	0.00015698768375104583
GO:0048639: positive regulation of developmental growth PRKDC	protein kinase, DNA-activated, catalytic polypeptide -0.0018694495197106365
GO:0060480: lung goblet cell differentiation	AGR2	anterior gradient 2	0.0016381287197342284
GO:0060480: lung goblet cell differentiation	HOXA5	homeobox A5	0.0010735335675550143
GO:0060480: lung goblet cell differentiation	SPDEF	SAM pointed domain containing ETS transcription factor	0.003155937456267705
GO:0060548: negative regulation of cell death	AGR2	anterior gradient 2	0.0015923902205138094
GO:0060548: negative regulation of cell death	BMP4	bone morphogenetic protein 4	-0.00031256338030417554
GO:0060548: negative regulation of cell death	BMP7	bone morphogenetic protein 7	0.0008343654848161874
GO:0060548: negative regulation of cell death	CST3	cystatin C	-5.6923105634038216e-5
GO:0060548: negative regulation of cell death	MGMT	O-6-methylguanine-DNA methyltransferase 0.0004086293854645374
GO:0060548: negative regulation of cell death	NCK1	NCK adaptor protein 1	-0.0007719424766182613
GO:0060548: negative regulation of cell death	SOX11	SRY (sex determining region Y)-box 11	-0.0021052977470417688
GO:0060548: negative regulation of cell death	SOX4	SRY (sex determining region Y)-box 4	-4.6357151762039666e-5
GO:0070254: mucus secretion	AGR2	anterior gradient 2	0.001616559967177164
GO:0070254: mucus secretion	VAMP8	vesicle-associated membrane protein 8	-5.7398246831976084e-5
GO:0090004: positive regulation of establishment of protein localization to plasma membrane	AGR2	anterior gradient 2	0.0016314391645785957
GO:0090004: positive regulation of establishment of protein localization to plasma membrane	AKT1	v-akt murine thymoma viral oncogene homolog 1	0.0007330372891685904
GO:0090004: positive regulation of establishment of protein localization to plasma membrane	CIB1	calcium and integrin binding 1 (calmyrin)	5.371600698006697e-5
GO:0090004: positive regulation of establishment of protein localization to plasma membrane	ITGA3	integrin, alpha 3 (antigen CD49C, alpha 3 subunit of VLA-3 receptor)	0.0014033841205646697
GO:0090004: positive regulation of establishment of protein localization to plasma membrane	ITGB1	integrin, beta 1 (fibronectin receptor, beta polypeptide, antigen CD29 includes MDF2, MSK12)	0.002434774046461785
GO:1903896: positive regulation of IRE1-mediated unfolded protein response	AGR2	anterior gradient 2	0.0016261012102319812
GO:1903896: positive regulation of IRE1-mediated unfolded protein response	BAK1	BCL2-antagonist/killer 1	-0.0018674303700608119
GO:1903896: positive regulation of IRE1-mediated unfolded protein response	BAX	BCL2-associated X protein	-0.0004245322910266186
GO:1903896: positive regulation of IRE1-mediated unfolded protein response	BCL2L11 BCL2-like 11 (apoptosis facilitator)	-0.0007730341545850334
GO:1903896: positive regulation of IRE1-mediated unfolded protein response	PTPN1	protein tyrosine phosphatase, non-receptor type 1	-0.0010358127098714785
GO:1903899: positive regulation of PERK-mediated unfolded protein response	AGR2	anterior gradient 2	0.0016154099039124756
GO:1903899: positive regulation of PERK-mediated unfolded protein response	PTPN2	protein tyrosine phosphatase, non-receptor type 2	-0.00032766327631981345
KEGG:04920: Adipocytokine signaling pathway	AGRP	agouti related protein homolog (mouse)	-0.00258458293485761
KEGG:04920: Adipocytokine signaling pathway	AKT1	v-akt murine thymoma viral oncogene homolog 1	0.0007355932430650508
KEGG:04920: Adipocytokine signaling pathway	AKT2	v-akt murine thymoma viral oncogene homolog 2	-0.0006290229838738278
KEGG:04920: Adipocytokine signaling pathway	CPT1A	carnitine palmitoyltransferase 1A (liver)	0.0012785223405717785
KEGG:04920: Adipocytokine signaling pathway	JAK2	Janus kinase 2	-3.11322706299306e-5
KEGG:04920: Adipocytokine signaling pathway	LEP	leptin	0.0032118289187881366
KEGG:04920: Adipocytokine signaling pathway	RXRA	retinoid X receptor, alpha	0.0011217176846465627
GO:0007218: neuropeptide signaling pathway	AGRP	agouti related protein homolog (mouse)	-0.002598079954431895
GO:0007218: neuropeptide signaling pathway	CPE	carboxypeptidase E	-0.0025727351448356565
GO:0007218: neuropeptide signaling pathway	NMU	neuromedin U	0.001873342195725202
GO:0007623: circadian rhythm	AGRP	agouti related protein homolog (mouse)	-0.0025814852074817173
GO:0007623: circadian rhythm	ATF5	activating transcription factor 5	-0.002746917064340821

5
GO:0007623: circadian rhythm    AVP     arginine vasopressin    -0.0009420770898513263
GO:0007623: circadian rhythm    CLOCK   clock circadian regulator       0.0001999754857051074
GO:0007623: circadian rhythm    CPT1A   carnitine palmitoyltransferase 1A (liver)       0.00127703879
22196994
GO:0007623: circadian rhythm    CREB1   cAMP responsive element binding protein 1       0.00066239449
54809945
GO:0007623: circadian rhythm    DRD4    dopamine receptor D4    -0.0019527946500199067
GO:0007623: circadian rhythm    EGR1    early growth response 1 0.0010978600316305883
GO:0007623: circadian rhythm    FAS     Fas cell surface death receptor -3.360411498469055e-5
GO:0007623: circadian rhythm    GSK3B   glycogen synthase kinase 3 beta 0.0015561253915316107
GO:0007623: circadian rhythm    LEP     leptin  0.003207028097573989
GO:0007623: circadian rhythm    NRIP1   nuclear receptor interacting protein 1  0.0010756727878886592
GO:0007623: circadian rhythm    PER2    period circadian clock 2        0.0012439650648792557
GO:0007623: circadian rhythm    PROX1   prospero homeobox 1     0.001126948966941315
GO:0007623: circadian rhythm    SERPINE1        serpin peptidase inhibitor, clade E (nexin, plasminog
en activator inhibitor type 1), member 1        0.0001119474631762424
GO:0007623: circadian rhythm    TYMS    thymidylate synthetase  0.0015696074884617001
GO:0007631: feeding behavior    AGRP    agouti related protein homolog (mouse)  -0.002576312040367615
GO:0007631: feeding behavior    DRD2    dopamine receptor D2    -0.00023435807408985762
GO:0007631: feeding behavior    HTR2C   5-hydroxytryptamine (serotonin) receptor 2C, G protein-couple
d       -0.000526196174293973
GO:0007631: feeding behavior    STRA6   stimulated by retinoic acid 6   -0.0016843392582535418
GO:0008343: adult feeding behavior      AGRP    agouti related protein homolog (mouse)  -0.0026068701
655804676
GO:0008343: adult feeding behavior      LEP     leptin  0.0032441430554579808
GO:0009648: photoperiodism      AGRP    agouti related protein homolog (mouse)  -0.002586827397754356
2
GO:0009648: photoperiodism      CLOCK   clock circadian regulator       0.0001995736604176251
GO:0009648: photoperiodism      DRD4    dopamine receptor D4    -0.0019562601533895537
GO:0009648: photoperiodism      NMU     neuromedin U    0.0018635970788099955
GO:0009755: hormone-mediated signaling pathway  AGRP    agouti related protein homolog (mouse)  -0.00
25752002813014107
GO:0009755: hormone-mediated signaling pathway  JAK2    Janus kinase 2  -3.209043029876428e-5
GO:0009755: hormone-mediated signaling pathway  THRA    thyroid hormone receptor, alpha 0.00076507448
11826131
GO:0042755: eating behavior     AGRP    agouti related protein homolog (mouse)  -0.002594858428084233
2
GO:0042755: eating behavior     CPT1A   carnitine palmitoyltransferase 1A (liver)       0.00128159287
13301437
GO:0042755: eating behavior     LEP     leptin  0.0032263055093409328
GO:0042755: eating behavior     NMU     neuromedin U    0.0018697811514943195
GO:0042755: eating behavior     TCF15   transcription factor 15 (basic helix-loop-helix)        -0.00
26763268812153925
GO:0042755: eating behavior     TH      tyrosine hydroxylase    -0.000344178260329221
GO:0060135: maternal process involved in female pregnancy       AGRP    agouti related protein homolo
g (mouse)       -0.002571362204035076
GO:0060135: maternal process involved in female pregnancy       CCL2    chemokine (C-C motif) ligand
 2      0.0008167085215575093
GO:0060135: maternal process involved in female pregnancy       DSG2    desmoglein 2    1.66297648020
01456e-5
GO:0060135: maternal process involved in female pregnancy       IHH     indian hedgehog -0.0020411997
218876187
GO:0060135: maternal process involved in female pregnancy       ITGA3   integrin, alpha 3 (antigen CD
49C, alpha 3 subunit of VLA-3 receptor) 0.0014006125361972629
GO:0060135: maternal process involved in female pregnancy       LGALS9  lectin, galactoside-binding,
 soluble, 9     -0.00011694111963838872
GO:0060135: maternal process involved in female pregnancy       MMP7    matrix metallopeptidase 7 (ma
trilysin, uterine)      0.00042535684990099333
GO:0060259: regulation of feeding behavior      AGRP    agouti related protein homolog (mouse)  -0.00
25994695453300225
GO:2000253: positive regulation of feeding behavior     AGRP    agouti related protein homolog (mous
e)      -0.0025994695453300225
KEGG:04614: Renin-angiotensin system    AGT     angiotensinogen (serpin peptidase inhibitor, clade A,
member 8)       -0.0011169922119025708
KEGG:04614: Renin-angiotensin system    AGTR1   angiotensin II receptor, type 1 0.0003452163775896154
7
KEGG:04614: Renin-angiotensin system    MAS1    MAS1 proto-oncogene, G protein-coupled receptor 0.000
2909766497993822
KEGG:04924: Renin secretion     AGT     angiotensinogen (serpin peptidase inhibitor, clade A, member
 8)     -0.0011139206084095315
KEGG:04924: Renin secretion     AGTR1   angiotensin II receptor, type 1 0.00034539412973386146
KEGG:04924: Renin secretion     CREB1   cAMP responsive element binding protein 1       0.00064678059
74784756
KEGG:04924: Renin secretion     ITPR1   inositol 1,4,5-trisphosphate receptor, type 1   -0.0007686695
942128237
KEGG:04924: Renin secretion     KCNJ2   potassium inwardly-rectifying channel, subfamily J, member 2
-0.000766749771381275
KEGG:04924: Renin secretion     KCNMA1  potassium large conductance calcium-activated channel, subfam
ily M, alpha member 1   -0.0006468055333396439
KEGG:04924: Renin secretion     PLCB1   phospholipase C, beta 1 (phosphoinositide-specific)     0.000

14994329191748548
KEGG:04924: Renin secretion PRKX protein kinase, X-linked 0.00038800348559060347
GO:0001543: ovarian follicle rupture AGT angiotensinogen (serpin peptidase inhibitor, clade A, member 8) -0.001133536416752873
GO:0001543: ovarian follicle rupture NRIP1 nuclear receptor interacting protein 1 0.0010690839378154195
GO:0001558: regulation of cell growth AGT angiotensinogen (serpin peptidase inhibitor, clade A, member 8) -0.0011239591957926947
GO:0001558: regulation of cell growth AGTR1 angiotensin II receptor, type 1 0.00034646295979797034
GO:0001558: regulation of cell growth FOXM1 forkhead box M1 0.00019628552953102386
GO:0001558: regulation of cell growth IGFBP2 insulin-like growth factor binding protein 2, 36kDa 0.00014265375258051428
GO:0001558: regulation of cell growth IGFBP3 insulin-like growth factor binding protein 3 0.0008327229314907618
GO:0001558: regulation of cell growth IGFBP4 insulin-like growth factor binding protein 4 -0.0012706687737660475
GO:0001558: regulation of cell growth IGFBP6 insulin-like growth factor binding protein 6 -0.0020158103178430824
GO:0001558: regulation of cell growth KIF14 kinesin family member 14 0.000484010442894907
GO:0001568: blood vessel development AGT angiotensinogen (serpin peptidase inhibitor, clade A, member 8) -0.0011386435333327553
GO:0001568: blood vessel development ALDH1A2 aldehyde dehydrogenase 1 family, member A2 -0.0030282294877035385
GO:0001568: blood vessel development COL1A1 collagen, type I, alpha 1 -0.0005294449881268177
GO:0001568: blood vessel development COL5A1 collagen, type V, alpha 1 -5.199023414011665e-5
GO:0001568: blood vessel development FOXO1 forkhead box O1 0.0018057365213942007
GO:0001568: blood vessel development LOX lysyl oxidase -0.0005785892056026903
GO:0001568: blood vessel development MEF2C myocyte enhancer factor 2C 0.0009770104438555638
GO:0001568: blood vessel development PAX6 paired box 6 0.0019421263098318656
GO:0001568: blood vessel development SPHK1 sphingosine kinase 1 0.0018168231948020308
GO:0001568: blood vessel development STRA6 stimulated by retinoic acid 6 -0.0016860521972014942
GO:0001568: blood vessel development TBX3 T-box 3 0.001228910923017606
GO:0001568: blood vessel development TCF7L2 transcription factor 7-like 2 (T-cell specific, HMG-box) 0.0005761762234375021
GO:0001658: branching involved in ureteric bud morphogenesis AGT angiotensinogen (serpin peptidase inhibitor, clade A, member 8) -0.001132421595568316
GO:0001658: branching involved in ureteric bud morphogenesis BCL2 B-cell CLL/lymphoma 2 -4.783028522101539e-6
GO:0001658: branching involved in ureteric bud morphogenesis BMP4 bone morphogenetic protein 4 -0.00032231007745216536
GO:0001658: branching involved in ureteric bud morphogenesis CITED1 Cbp/p300-interacting transactivator, with Glu/Asp-rich carboxy-terminal domain, 1 0.0028166717397520528
GO:0001658: branching involved in ureteric bud morphogenesis CTNNB1 catenin (cadherin-associated protein), beta 1, 88kDa -0.00011696429615207499
GO:0001658: branching involved in ureteric bud morphogenesis CTNNBIP1 catenin, beta interacting protein 1 0.0005853909251136402
GO:0001658: branching involved in ureteric bud morphogenesis EYA1 EYA transcriptional coactivator and phosphatase 1 5.310144828986267e-5
GO:0001658: branching involved in ureteric bud morphogenesis FGF8 fibroblast growth factor 8 (androgen-induced) 0.0009833334907729438
GO:0001658: branching involved in ureteric bud morphogenesis GDNF glial cell derived neurotrophic factor 0.0004445285581730752
GO:0001658: branching involved in ureteric bud morphogenesis GLI3 GLI family zinc finger 3 -0.002150071239082692
GO:0001658: branching involved in ureteric bud morphogenesis GREM1 gremlin 1, DAN family BMP antagonist -0.0008268716043978847
GO:0001658: branching involved in ureteric bud morphogenesis LHX1 LIM homeobox 1 -0.0007582861704903152
GO:0001658: branching involved in ureteric bud morphogenesis MYC v-myc avian myelocytomatosis viral oncogene homolog -0.0011360947770484152
GO:0001658: branching involved in ureteric bud morphogenesis PAX2 paired box 2 -0.0016129605463831643
GO:0001658: branching involved in ureteric bud morphogenesis PAX8 paired box 8 0.000947632249931198
GO:0001658: branching involved in ureteric bud morphogenesis PTCH1 patched 1 -6.596917879542931e-5
GO:0001658: branching involved in ureteric bud morphogenesis SALL1 spalt-like transcription factor 1 -0.0024359193832538747
GO:0001658: branching involved in ureteric bud morphogenesis SHH sonic hedgehog 0.000599594129670786
GO:0001658: branching involved in ureteric bud morphogenesis SIX1 SIX homeobox 1 -0.0018688091832661294
GO:0001658: branching involved in ureteric bud morphogenesis SOX9 SRY (sex determining region Y)-box 9 -0.0005185016850486457
GO:0001658: branching involved in ureteric bud morphogenesis WNT1 wingless-type MMTV integration site family, member 1 0.0007864630843607674
GO:0001658: branching involved in ureteric bud morphogenesis WNT4 wingless-type MMTV integration site family, member 4 -0.0002517234722073026

GO:0001658: branching involved in ureteric bud morphogenesis    WT1     Wilms tumor 1   -0.000506633399667247
GO:0001819: positive regulation of cytokine production  AGT     angiotensinogen (serpin peptidase inhibitor, clade A, member 8)      -0.0011514506577944982
GO:0001819: positive regulation of cytokine production  FLOT1   flotillin 1     -0.000770043146373940 5
GO:0001819: positive regulation of cytokine production  LEP     leptin  0.003230302030180371
GO:0001822: kidney development  AGT     angiotensinogen (serpin peptidase inhibitor, clade A, member 8)     -0.0011322885087761373
GO:0001822: kidney development  AGTR1   angiotensin II receptor, type 1 0.0003476434851386544
GO:0001822: kidney development  ALDH1A2 aldehyde dehydrogenase 1 family, member A2      -0.003017440452369914
GO:0001822: kidney development  APC     adenomatous polyposis coli      0.0006435382371252014
GO:0001822: kidney development  ARL3    ADP-ribosylation factor-like 3  -0.0011802692847468813
GO:0001822: kidney development  ASS1    argininosuccinate synthase 1    0.0001423666715683511
GO:0001822: kidney development  BAX     BCL2-associated X protein       -0.00042495573157465386
GO:0001822: kidney development  BCL2L11 BCL2-like 11 (apoptosis facilitator)    -0.000774409498806054 3
GO:0001822: kidney development  BMP4    bone morphogenetic protein 4    -0.00032241516797292745
GO:0001822: kidney development  GATA3   GATA binding protein 3  -3.896905088019714e-5
GO:0001822: kidney development  GLI2    GLI family zinc finger 2        0.0018523887542204842
GO:0001822: kidney development  JMJD6   jumonji domain containing 6     0.003588983878481649
GO:0001822: kidney development  LEF1    lymphoid enhancer-binding factor 1      -0.00010030675272366698
GO:0001822: kidney development  LHX1    LIM homeobox 1  -0.0007582482598129826
GO:0001822: kidney development  ODC1    ornithine decarboxylase 1       0.0008248122290811511
GO:0001822: kidney development  PAX8    paired box 8    0.0009477606784070618
GO:0001822: kidney development  PROX1   prospero homeobox 1     0.001120769548385456
GO:0001822: kidney development  SALL1   spalt-like transcription factor 1       -0.002435606551412037 4
GO:0001822: kidney development  SIX1    SIX homeobox 1  -0.0018688725541587952
GO:0001822: kidney development  SOX11   SRY (sex determining region Y)-box 11   -0.000209040638474912 6
GO:0001822: kidney development  STRA6   stimulated by retinoic acid 6   -0.0016794296292797035
GO:0001822: kidney development  SULF1   sulfatase 1     -0.0008049564743280624
GO:0001822: kidney development  TFAP2A  transcription factor AP-2 alpha (activating enhancer binding protein 2 alpha)       0.0004796553809574217
GO:0001822: kidney development  TGFBR1  transforming growth factor, beta receptor 1     0.00033964589359753415
GO:0001822: kidney development  THRA    thyroid hormone receptor, alpha 0.0007627917589666244
GO:0001822: kidney development  TP73    tumor protein p73       0.0010272788843812657
GO:0001822: kidney development  VEGFA   vascular endothelial growth factor A    0.0005944993942241002
GO:0001822: kidney development  WFS1    Wolfram syndrome 1 (wolframin)  0.000576583569760183
GO:0001822: kidney development  WNT4    wingless-type MMTV integration site family, member 4    -0.00025176071467857426
GO:0001822: kidney development  WT1     Wilms tumor 1   -0.000506511294641922
GO:0001998: angiotensin mediated vasoconstriction involved in regulation of systemic arterial blood pressure AGT     angiotensinogen (serpin peptidase inhibitor, clade A, member 8) -0.001141324461898953 3
GO:0001999: renal response to blood flow involved in circulatory renin-angiotensin regulation of systemic arterial blood pressure    AGT     angiotensinogen (serpin peptidase inhibitor, clade A, member 8)     -0.0011413244618989533
GO:0002003: angiotensin maturation      AGT     angiotensinogen (serpin peptidase inhibitor, clade A, member 8)       -0.0011413244618989533
GO:0002016: regulation of blood volume by renin-angiotensin     AGT     angiotensinogen (serpin peptidase inhibitor, clade A, member 8)     -0.0011413244618989533
GO:0002018: renin-angiotensin regulation of aldosterone production      AGT     angiotensinogen (serpin peptidase inhibitor, clade A, member 8)      -0.0011273061613599608
GO:0002018: renin-angiotensin regulation of aldosterone production      AGTR1   angiotensin II receptor, type 1      0.00034518432154975775
GO:0002019: regulation of renal output by angiotensin   AGT     angiotensinogen (serpin peptidase inhibitor, clade A, member 8)      -0.0011413244618989533
GO:0002027: regulation of heart rate    AGT     angiotensinogen (serpin peptidase inhibitor, clade A, member 8)       -0.0011190207888007175
GO:0002027: regulation of heart rate    DRD2    dopamine receptor D2    -0.0002322212772190437
GO:0002027: regulation of heart rate    GPD1L   glycerol-3-phosphate dehydrogenase 1-like       0.0005611155218810467
GO:0002034: regulation of blood vessel size by renin-angiotensin        AGT     angiotensinogen (serpin peptidase inhibitor, clade A, member 8)      -0.0011197730280089002
GO:0002034: regulation of blood vessel size by renin-angiotensin        AGTR1   angiotensin II receptor, type 1      0.00034590288283904464
GO:0002034: regulation of blood vessel size by renin-angiotensin        SERPINF2        serpin peptidase inhibitor, clade F (alpha-2 antiplasmin, pigment epithelium derived factor), member 2       0.0006525904449321849
GO:0003014: renal system process        AGT     angiotensinogen (serpin peptidase inhibitor, clade A, member 8)       -0.001124249487657679
GO:0003014: renal system process        BCL2    B-cell CLL/lymphoma 2   -4.809622110100739e-6
GO:0003014: renal system process        BMP4    bone morphogenetic protein 4    -0.00032065295276132507
GO:0003014: renal system process        FAS     Fas cell surface death receptor -3.312797888114488e-5
GO:0003014: renal system process        PKN1    protein kinase N1       -0.0018923139906094526

GO:0003051: angiotensin-mediated drinking behavior	AGT	angiotensinogen (serpin peptidase inhibitor, clade A, member 8)	-0.0011413244618989533
GO:0003331: positive regulation of extracellular matrix constituent secretion	AGT	angiotensinogen (serpin peptidase inhibitor, clade A, member 8)	-0.0011413244618989533
GO:0006883: cellular sodium ion homeostasis	AGT	angiotensinogen (serpin peptidase inhibitor, clade A, member 8)	-0.0011378125879293243
GO:0006883: cellular sodium ion homeostasis	ATP1B3	ATPase, Na+/K+ transporting, beta 3 polypeptide	-0.0015001052654050246
GO:0007160: cell-matrix adhesion	AGT	angiotensinogen (serpin peptidase inhibitor, clade A, member 8)	-0.0011379943618682917
GO:0007160: cell-matrix adhesion	BCL2L11	BCL2-like 11 (apoptosis facilitator)	-0.0007757746956300358
GO:0007160: cell-matrix adhesion	CD63	CD63 molecule	-0.0010646265330701344
GO:0007160: cell-matrix adhesion	CTNNB1	catenin (cadherin-associated protein), beta 1, 88kDa	-0.00011670494850022754
GO:0007160: cell-matrix adhesion	EDA	ectodysplasin A	-0.0008172001685629274
GO:0007160: cell-matrix adhesion	ITGA3	integrin, alpha 3 (antigen CD49C, alpha 3 subunit of VLA-3 receptor)	0.0014023025785692758
GO:0007160: cell-matrix adhesion	ITGA6	integrin, alpha 6	0.001675856995409526
GO:0007160: cell-matrix adhesion	ITGB1	integrin, beta 1 (fibronectin receptor, beta polypeptide, antigen CD29 includes MDF2, MSK12)	0.002433633248060729
GO:0007160: cell-matrix adhesion	ITGB4	integrin, beta 4	0.0005828388908821713
GO:0007160: cell-matrix adhesion	THBS3	thrombospondin 3	0.001620301316347829
GO:0007186: G-protein coupled receptor signaling pathway	AGT	angiotensinogen (serpin peptidase inhibitor, clade A, member 8)	-0.001135681886770785
GO:0007186: G-protein coupled receptor signaling pathway	AGTR1	angiotensin II receptor, type 1	0.0003472775024194787
GO:0007186: G-protein coupled receptor signaling pathway	AKT1	v-akt murine thymoma viral oncogene homolog 1	0.0007317522629380932
GO:0007186: G-protein coupled receptor signaling pathway	AREG	amphiregulin	0.0024600912856273306
GO:0007186: G-protein coupled receptor signaling pathway	C3	complement component 3	0.002027451858856666
GO:0007186: G-protein coupled receptor signaling pathway	CCL2	chemokine (C-C motif) ligand 2	0.0008179630959293952
GO:0007186: G-protein coupled receptor signaling pathway	CCL7	chemokine (C-C motif) ligand 7	-0.0024049453512226576
GO:0007186: G-protein coupled receptor signaling pathway	CCL8	chemokine (C-C motif) ligand 8	-0.0005905841499930062
GO:0007186: G-protein coupled receptor signaling pathway	CELSR2	cadherin, EGF LAG seven-pass G-type receptor 2	-0.0011365478355988014
GO:0007186: G-protein coupled receptor signaling pathway	CX3CL1	chemokine (C-X3-C motif) ligand 1	0.0021205962195314804
GO:0007186: G-protein coupled receptor signaling pathway	CX3CR1	chemokine (C-X3-C motif) receptor 1	-0.0008580729258881207
GO:0007186: G-protein coupled receptor signaling pathway	CXCL10	chemokine (C-X-C motif) ligand 10	6.31941933739969e-5
GO:0007186: G-protein coupled receptor signaling pathway	CXCL12	chemokine (C-X-C motif) ligand 12	-0.0012103273851138332
GO:0007186: G-protein coupled receptor signaling pathway	CXCL13	chemokine (C-X-C motif) ligand 13	0.0029976064731152066
GO:0007186: G-protein coupled receptor signaling pathway	CXCR4	chemokine (C-X-C motif) receptor 4	0.0007961901815883596
GO:0007186: G-protein coupled receptor signaling pathway	FRS2	fibroblast growth factor receptor substrate 2	0.0009311209785255437
GO:0007186: G-protein coupled receptor signaling pathway	FZD3	frizzled class receptor 3	0.00037958442708152366
GO:0007186: G-protein coupled receptor signaling pathway	FZD7	frizzled class receptor 7	0.001072572538989316
GO:0007186: G-protein coupled receptor signaling pathway	GNRHR	gonadotropin-releasing hormone receptor	0.0007414722958120305
GO:0007186: G-protein coupled receptor signaling pathway	GPR87	G protein-coupled receptor 87	-2.497251587351049e-5
GO:0007186: G-protein coupled receptor signaling pathway	GPSM2	G-protein signaling modulator 2	0.0017753013506732127
GO:0007186: G-protein coupled receptor signaling pathway	INSR	insulin receptor	-0.0013653012317385407
GO:0007186: G-protein coupled receptor signaling pathway	JAK2	Janus kinase 2	-3.129415782048627e-5
GO:0007186: G-protein coupled receptor signaling pathway	LGR4	leucine-rich repeat containing G protein-coupled receptor 4	0.0003088865309290892
GO:0007186: G-protein coupled receptor signaling pathway	MAS1	MAS1 proto-oncogene, G protein-coupled receptor	0.00029110400707595406
GO:0007186: G-protein coupled receptor signaling pathway	NMU	neuromedin U	0.001853474823473163
GO:0007186: G-protein coupled receptor signaling pathway	OR5I1	olfactory receptor, family 5, subfamily I, member 1	0.0018843550241663888
GO:0007186: G-protein coupled receptor signaling pathway	PDGFRB	platelet-derived growth factor receptor, beta polypeptide	-0.0003624778230943918
GO:0007186: G-protein coupled receptor signaling pathway	SMO	smoothened, frizzled class receptor	0.002142193010941400З

GO:0007186: G-protein coupled receptor signaling pathway TULP3 tubby like protein 3 0.000
9579090710494531
GO:0007199: G-protein coupled receptor signaling pathway coupled to cGMP nucleotide second messenger
AGT angiotensinogen (serpin peptidase inhibitor, clade A, member 8) -0.0011413244618989533
GO:0007200: phospholipase C-activating G-protein coupled receptor signaling pathway AGT angio
tensinogen (serpin peptidase inhibitor, clade A, member 8) -0.0011148533931377089
GO:0007200: phospholipase C-activating G-protein coupled receptor signaling pathway AGTR1 angio
tensin II receptor, type 1 0.0003456715919302292
GO:0007200: phospholipase C-activating G-protein coupled receptor signaling pathway ESR1 estro
gen receptor 1 -0.0009352668420446902
GO:0007200: phospholipase C-activating G-protein coupled receptor signaling pathway HTR2C 5-hyd
roxytryptamine (serotonin) receptor 2C, G protein-coupled -0.0005181719878916906
GO:0007200: phospholipase C-activating G-protein coupled receptor signaling pathway TGM2 trans
glutaminase 2 -0.00017696815049185778
GO:0007202: activation of phospholipase C activity AGT angiotensinogen (serpin peptidase inh
ibitor, clade A, member 8) -0.0011236894533061515
GO:0007202: activation of phospholipase C activity CREB1 cAMP responsive element binding prote
in 1 0.0006526255386096972
GO:0007202: activation of phospholipase C activity EGFR epidermal growth factor receptor
0.0006800670113925343
GO:0007202: activation of phospholipase C activity ITPR1 inositol 1,4,5-trisphosphate recepto
r, type 1 -0.0007744885115874016
GO:0007202: activation of phospholipase C activity PRKCA protein kinase C, alpha -5.8315058233
73601e-6
GO:0007202: activation of phospholipase C activity PRKCD protein kinase C, delta -0.0011195788
348135891
GO:0007250: activation of NF-kappaB-inducing kinase activity AGT angiotensinogen (serpin pepti
dase inhibitor, clade A, member 8) -0.0011342443589731634
GO:0007250: activation of NF-kappaB-inducing kinase activity IRAK1 interleukin-1 receptor-associ
ated kinase 1 -0.00160908965092749
GO:0007250: activation of NF-kappaB-inducing kinase activity MAS1 MAS1 proto-oncogene, G protei
n-coupled receptor 0.000290753142077939
GO:0007263: nitric oxide mediated signal transduction AGT angiotensinogen (serpin peptidase inh
ibitor, clade A, member 8) -0.0011272561800915579
GO:0007263: nitric oxide mediated signal transduction PDX1 pancreatic and duodenal homeobox 1
0.0002558342245161704
GO:0007588: excretion AGT angiotensinogen (serpin peptidase inhibitor, clade A, member 8) -0.00
11194164532164915
GO:0007588: excretion HMOX1 heme oxygenase (decycling) 1 -0.00021522995210703462
GO:0007588: excretion SLC22A18 solute carrier family 22, member 18 0.0007274064541036471
GO:0008065: establishment of blood-nerve barrier AGT angiotensinogen (serpin peptidase inh
ibitor, clade A, member 8) -0.0011364185548147984
GO:0008065: establishment of blood-nerve barrier GSTM3 glutathione S-transferase mu 3 (brai
n) -0.0012904510104553937
GO:0009651: response to salt stress AGT angiotensinogen (serpin peptidase inhibitor, clade A,
member 8) -0.0011250371577881835
GO:0009651: response to salt stress BAX BCL2-associated X protein -0.000422612395972620
63
GO:0009651: response to salt stress TH tyrosine hydroxylase -0.00034049874484744767
GO:0009651: response to salt stress TP53 tumor protein p53 0.0011676634794120424
GO:0010535: positive regulation of activation of JAK2 kinase activity AGT angiotensinogen (serp
in peptidase inhibitor, clade A, member 8) -0.001134097097218038
GO:0010535: positive regulation of activation of JAK2 kinase activity IL12B interleukin 12B 0.001
276451706293287
GO:0010595: positive regulation of endothelial cell migration AGT angiotensinogen (serpin pepti
dase inhibitor, clade A, member 8) -0.0011220269800965624
GO:0010595: positive regulation of endothelial cell migration ALOX12 arachidonate 12-lipoxygenase
-0.0019394077472052154
GO:0010595: positive regulation of endothelial cell migration ANGPT1 angiopoietin 1 0.00089504716
83722528
GO:0010595: positive regulation of endothelial cell migration BMP4 bone morphogenetic protein 4
-0.0003203270222288026
GO:0010595: positive regulation of endothelial cell migration FOXC2 forkhead box C2 (MFH-1, mesen
chyme forkhead 1) 0.001736344833359962
GO:0010595: positive regulation of endothelial cell migration GATA3 GATA binding protein 3 -3.76
6319822270048e-5
GO:0010595: positive regulation of endothelial cell migration NRP1 neuropilin 1 -0.0006378237
944011305
GO:0010595: positive regulation of endothelial cell migration PRKCA protein kinase C, alpha -5.93
64492213859905e-6
GO:0010595: positive regulation of endothelial cell migration PROX1 prospero homeobox 1 0.001
1126813719566882
GO:0010595: positive regulation of endothelial cell migration THBS1 thrombospondin 1 -0.00
10283456165712283
GO:0010595: positive regulation of endothelial cell migration VEGFA vascular endothelial growth f
actor A 0.0005889216945167238
GO:0010595: positive regulation of endothelial cell migration WNT5A wingless-type MMTV integratio
n site family, member 5A -0.0006622910120876477
GO:0010595: positive regulation of endothelial cell migration WNT7A wingless-type MMTV integratio
n site family, member 7A 1.984883657709688e-5
GO:0010613: positive regulation of cardiac muscle hypertrophy AGT angiotensinogen (serpin pepti

dase inhibitor, clade A, member 8)	-0.0011277061382266415
GO:0010613: positive regulation of cardiac muscle hypertrophy	IGF1	insulin-like growth factor 1 (somatomedin C)	0.0001299906142537372
GO:0010613: positive regulation of cardiac muscle hypertrophy	IL6ST	interleukin 6 signal transducer	0.0018700966561512522
GO:0010613: positive regulation of cardiac muscle hypertrophy	PRKCA	protein kinase C, alpha -5.952841332274639e-6
GO:0010744: positive regulation of macrophage derived foam cell differentiation AGT	angiotensinogen (serpin peptidase inhibitor, clade A, member 8)	-0.0011273061613599608
GO:0010744: positive regulation of macrophage derived foam cell differentiation AGTR1	angiotensin II receptor, type 1	0.00034518432154975775
GO:0010873: positive regulation of cholesterol esterification	AGT	angiotensinogen (serpin peptidase inhibitor, clade A, member 8)	-0.0011273061613599608
GO:0010873: positive regulation of cholesterol esterification	AGTR1	angiotensin II receptor, type 1	0.00034518432154975775
GO:0010951: negative regulation of endopeptidase activity	AGT	angiotensinogen (serpin peptidase inhibitor, clade A, member 8)	-0.0011427867962670623
GO:0010951: negative regulation of endopeptidase activity	AKT1	v-akt murine thymoma viral oncogene homolog 1	0.0007357118999286487
GO:0010951: negative regulation of endopeptidase activity	C3	complement component 3	0.002037330251145308
GO:0010951: negative regulation of endopeptidase activity	CAST	calpastatin	-0.0030523211956222976
GO:0010951: negative regulation of endopeptidase activity	CST3	cystatin C	-7.225017747036872e-5
GO:0010951: negative regulation of endopeptidase activity	PI3	peptidase inhibitor 3, skin-derived	-0.0010467915172388356
GO:0010951: negative regulation of endopeptidase activity	PTTG1	pituitary tumor-transforming 1	-0.00036217313646242266
GO:0010951: negative regulation of endopeptidase activity	SERPINA5	serpin peptidase inhibitor, clade A (alpha-1 antiproteinase, antitrypsin), member 5	-0.0007139125929340433
GO:0010951: negative regulation of endopeptidase activity	SERPINB5	serpin peptidase inhibitor, clade B (ovalbumin), member 5	-0.004556683309996608
GO:0010951: negative regulation of endopeptidase activity	SERPINE1	serpin peptidase inhibitor, clade E (nexin, plasminogen activator inhibitor type 1), member 1	0.0001118708448074291
GO:0010951: negative regulation of endopeptidase activity	SERPINE2	serpin peptidase inhibitor, clade E (nexin, plasminogen activator inhibitor type 1), member 2	0.002517990175649599
GO:0010951: negative regulation of endopeptidase activity	SERPINF2	serpin peptidase inhibitor, clade F (alpha-2 antiplasmin, pigment epithelium derived factor), member 2	0.0006623869558177233
GO:0010951: negative regulation of endopeptidase activity	SERPINH1	serpin peptidase inhibitor, clade H (heat shock protein 47), member 1, (collagen binding protein 1)	-0.0010955084418645016
GO:0014061: regulation of norepinephrine secretion	AGT	angiotensinogen (serpin peptidase inhibitor, clade A, member 8)	-0.0011413244618989533
GO:0014068: positive regulation of phosphatidylinositol 3-kinase signaling	AGT	angiotensinogen (serpin peptidase inhibitor, clade A, member 8)	-0.0011254763501075068
GO:0014068: positive regulation of phosphatidylinositol 3-kinase signaling	ANGPT1	angiopoietin 1	0.0008966105857940089
GO:0014068: positive regulation of phosphatidylinositol 3-kinase signaling	FLT3	fms-related tyrosine kinase 3	-0.0006772422850948269
GO:0014068: positive regulation of phosphatidylinositol 3-kinase signaling	IGF1	insulin-like growth factor 1 (somatomedin C)	0.00012950036429621651
GO:0014068: positive regulation of phosphatidylinositol 3-kinase signaling	JAK2	Janus kinase 2	-3.262564335094834e-5
GO:0014068: positive regulation of phosphatidylinositol 3-kinase signaling	KIT	v-kit Hardy-Zuckerman 4 feline sarcoma viral oncogene homolog	0.0002716325925501927
GO:0014068: positive regulation of phosphatidylinositol 3-kinase signaling	NEDD4	neural precursor cell expressed, developmentally down-regulated 4, E3 ubiquitin protein ligase	0.002252824626825657
GO:0014068: positive regulation of phosphatidylinositol 3-kinase signaling	PDGFRB	platelet-derived growth factor receptor, beta polypeptide	-0.0003578912793950715
GO:0014068: positive regulation of phosphatidylinositol 3-kinase signaling	RELN	reelin	0.0015284506475392245
GO:0014068: positive regulation of phosphatidylinositol 3-kinase signaling	SOX9	SRY (sex determining region Y)-box 9 -0.0005160082523193126
GO:0014068: positive regulation of phosphatidylinositol 3-kinase signaling	TGFB2	transforming growth factor, beta 2	-0.0010534663100429144
GO:0014824: artery smooth muscle contraction	AGT	angiotensinogen (serpin peptidase inhibitor, clade A, member 8)	-0.0011413244618989533
GO:0014873: response to muscle activity involved in regulation of muscle adaptation	AGT	angiotensinogen (serpin peptidase inhibitor, clade A, member 8)	-0.0011413244618989533
GO:0016525: negative regulation of angiogenesis AGT	angiotensinogen (serpin peptidase inhibitor, clade A, member 8)	-0.0011142345282872882
GO:0016525: negative regulation of angiogenesis CCL2	chemokine (C-C motif) ligand 2	0.0008062675702981245
GO:0016525: negative regulation of angiogenesis CX3CR1	chemokine (C-X3-C motif) receptor 1	-0.0008474072551148822
GO:0016525: negative regulation of angiogenesis CXCL10	chemokine (C-X-C motif) ligand 10	6.037374343293444e-5
GO:0016525: negative regulation of angiogenesis FOXC1	forkhead box C1 -2.3138581521752672e-5

GO:0016525: negative regulation of angiogenesis HOXA5 homeobox A5 0.0010503760111728367
GO:0016525: negative regulation of angiogenesis PML promyelocytic leukemia -0.000673340743943491
6
GO:0016525: negative regulation of angiogenesis SULF1 sulfatase 1 -0.0007903565640150291
GO:0016525: negative regulation of angiogenesis THBS1 thrombospondin 1 -0.001022742126216288
8
GO:0016525: negative regulation of angiogenesis THBS4 thrombospondin 4 -0.000441214516278042
GO:0016525: negative regulation of angiogenesis VASH1 vasohibin 1 0.0005917174396845665
GO:0019229: regulation of vasoconstriction AGT angiotensinogen (serpin peptidase inhibitor, clade A, member 8) -0.0011377222488670726
GO:0019229: regulation of vasoconstriction AGTR1 angiotensin II receptor, type 1 0.00034766174
511833127
GO:0019229: regulation of vasoconstriction ASIC2 acid-sensing (proton-gated) ion channel 2
0.002285878334650619
GO:0019229: regulation of vasoconstriction PER2 period circadian clock 2 0.00124167361
4763525
GO:0030198: extracellular matrix organization AGT angiotensinogen (serpin peptidase inhibitor, clade A, member 8) -0.0011353962802214252
GO:0030198: extracellular matrix organization BMP4 bone morphogenetic protein 4 -0.0003230041
7784592294
GO:0030198: extracellular matrix organization BMP7 bone morphogenetic protein 7 0.00085644808
38143902
GO:0030198: extracellular matrix organization COL11A1 collagen, type XI, alpha 1 -0.0004423323
769293622
GO:0030198: extracellular matrix organization COL1A1 collagen, type I, alpha 1 -0.0005275772
33248182
GO:0030198: extracellular matrix organization COL5A1 collagen, type V, alpha 1 -5.0651252001
06433e-5
GO:0030198: extracellular matrix organization COL5A2 collagen, type V, alpha 2 -0.0003465949
792463337
GO:0030198: extracellular matrix organization CTSK cathepsin K -0.0003640321303025609
GO:0030198: extracellular matrix organization CTSV cathepsin V 0.0006550577223109958
GO:0030198: extracellular matrix organization DAG1 dystroglycan 1 (dystrophin-associated glycopr
otein 1) 0.0003620444556816926
GO:0030198: extracellular matrix organization DSPP dentin sialophosphoprotein 0.00083063623
25807083
GO:0030198: extracellular matrix organization EGFL6 EGF-like-domain, multiple 6 0.00013687380
078647796
GO:0030198: extracellular matrix organization FBN2 fibrillin 2 -0.0007423944454956597
GO:0030198: extracellular matrix organization ICAM1 intercellular adhesion molecule 1 0.000
7296354396173889
GO:0030198: extracellular matrix organization ITGA3 integrin, alpha 3 (antigen CD49C, alpha 3 sub
unit of VLA-3 receptor) 0.001401145283838552
GO:0030198: extracellular matrix organization ITGA6 integrin, alpha 6 0.0016721877762541249
GO:0030198: extracellular matrix organization ITGB1 integrin, beta 1 (fibronectin receptor, beta polypeptide, antigen CD29 includes MDF2, MSK12) 0.002430008993952343
GO:0030198: extracellular matrix organization ITGB4 integrin, beta 4 0.0005814874489974715
GO:0030198: extracellular matrix organization KLK7 kallikrein-related peptidase 7 0.00162800037
5796593
GO:0030198: extracellular matrix organization LAMB2 laminin, beta 2 (laminin S) -0.0013108738
780589942
GO:0030198: extracellular matrix organization LOX lysyl oxidase -0.0005772349323089475
GO:0030198: extracellular matrix organization MMP2 matrix metallopeptidase 2 (gelatinase A, 72kD
a gelatinase, 72kDa type IV collagenase) -0.0011332886696424152
GO:0030198: extracellular matrix organization MMP20 matrix metallopeptidase 20 -0.0024346693
98792978
GO:0030198: extracellular matrix organization MMP7 matrix metallopeptidase 7 (matrilysin, uterin
e) 0.00042591564956760925
GO:0030198: extracellular matrix organization NR2E1 nuclear receptor subfamily 2, group E, member
1 -0.0029780332048168334
GO:0030198: extracellular matrix organization PRDX4 peroxiredoxin 4 0.0015264475327929476
GO:0030198: extracellular matrix organization PRKCA protein kinase C, alpha -5.839797129212113e-6
GO:0030198: extracellular matrix organization SERPINB5 serpin peptidase inhibitor, clade B (ovalbumin), member 5 -0.004536539408318589
GO:0030198: extracellular matrix organization SERPINE1 serpin peptidase inhibitor, clade E (nexin, plasminogen activator inhibitor type 1), member 1 0.0001121322413896657
GO:0030198: extracellular matrix organization SERPINH1 serpin peptidase inhibitor, clade H (heat shock protein 47), member 1, (collagen binding protein 1) -0.0010902762951661936
GO:0030198: extracellular matrix organization SOX9 SRY (sex determining region Y)-box 9 -0.00
05191319035850543
GO:0030198: extracellular matrix organization TGFB1 transforming growth factor, beta 1 -7.32
6144857069033e-5
GO:0030198: extracellular matrix organization TGFB2 transforming growth factor, beta 2 -0.00
10608832620807036
GO:0030198: extracellular matrix organization TGFB3 transforming growth factor, beta 3 -0.00
18265300195037515
GO:0030198: extracellular matrix organization THBS1 thrombospondin 1 -0.001037232948127688
3
GO:0030198: extracellular matrix organization TNC tenascin C 0.0007352193829639208
GO:0030432: peristalsis AGT angiotensinogen (serpin peptidase inhibitor, clade A, member 8) -0.00
1115861005758054

GO:0030432: peristalsis DRD2    dopamine receptor D2    -0.00023187057536279692
GO:0030432: peristalsis GDNF    glial cell derived neurotrophic factor  0.0004402006314804402
GO:0032270: positive regulation of cellular protein metabolic process   AGT     angiotensinogen (serp
in peptidase inhibitor, clade A, member 8)      -0.0011252869951244597
GO:0032270: positive regulation of cellular protein metabolic process   AGTR1   angiotensin II recept
or, type 1      0.0003465033700135899
GO:0032270: positive regulation of cellular protein metabolic process   AKT1    v-akt murine thymoma
 viral oncogene homolog 1       0.0007258870202631819
GO:0032270: positive regulation of cellular protein metabolic process   INHBA   inhibin, beta A -0.00
1343336194750891
GO:0032270: positive regulation of cellular protein metabolic process   NR1H3   nuclear receptor subf
amily 1, group H, member 3      0.0007965623366097778
GO:0032270: positive regulation of cellular protein metabolic process   TGFB1   transforming growth f
actor, beta 1   -7.252275201797462e-5
GO:0032930: positive regulation of superoxide anion generation  AGT     angiotensinogen (serpin pepti
dase inhibitor, clade A, member 8)      -0.001123035459903848
GO:0032930: positive regulation of superoxide anion generation  GSTP1   glutathione S-transferase pi
 1      0.00023020869859850975
GO:0032930: positive regulation of superoxide anion generation  PRKCD   protein kinase C, delta -0.00
11189176103720965
GO:0032930: positive regulation of superoxide anion generation  TGFB1   transforming growth factor, b
eta 1   -7.25348456252524e-5
GO:0033138: positive regulation of peptidyl-serine phosphorylation      AGT     angiotensinogen (serp
in peptidase inhibitor, clade A, member 8)      -0.0011282006662961489
GO:0033138: positive regulation of peptidyl-serine phosphorylation      AKT1    v-akt murine thymoma
 viral oncogene homolog 1       0.0007282841687977955
GO:0033138: positive regulation of peptidyl-serine phosphorylation      AKT2    v-akt murine thymoma
 viral oncogene homolog 2       -0.0006244734032888663
GO:0033138: positive regulation of peptidyl-serine phosphorylation      ANGPT1  angiopoietin 1  0.000
8983012903031152
GO:0033138: positive regulation of peptidyl-serine phosphorylation      AVP     arginine vasopressin
-0.0009339310067446521
GO:0033138: positive regulation of peptidyl-serine phosphorylation      AXIN1   axin 1  -0.0007304631
354543683
GO:0033138: positive regulation of peptidyl-serine phosphorylation      BCL2    B-cell CLL/lymphoma 2
-4.701486539407933e-6
GO:0033138: positive regulation of peptidyl-serine phosphorylation      CAV1    caveolin 1, caveolae
 protein, 22kDa -0.0005343319103674757
GO:0033138: positive regulation of peptidyl-serine phosphorylation      CDC42   cell division cycle 4
2       0.0012366886409960764
GO:0033138: positive regulation of peptidyl-serine phosphorylation      GSK3B   glycogen synthase kin
ase 3 beta      0.0015446040341675613
GO:0033138: positive regulation of peptidyl-serine phosphorylation      PAK1    p21 protein (Cdc42/Ra
c)-activated kinase 1   -0.0021812362555912596
GO:0033138: positive regulation of peptidyl-serine phosphorylation      PFN2    profilin 2      0.001
9989673366821257
GO:0033138: positive regulation of peptidyl-serine phosphorylation      RAF1    Raf-1 proto-oncogene,
serine/threonine kinase 0.001494459029090121
GO:0033138: positive regulation of peptidyl-serine phosphorylation      TGFB1   transforming growth f
actor, beta 1   -7.240759311739797e-5
GO:0033138: positive regulation of peptidyl-serine phosphorylation      VEGFA   vascular endothelial
 growth factor A        0.0005920034529647973
GO:0033138: positive regulation of peptidyl-serine phosphorylation      WNT5A   wingless-type MMTV in
tegration site family, member 5A        -0.0006650898199579927
GO:0033864: positive regulation of NAD(P)H oxidase activity     AGT     angiotensinogen (serpin pepti
dase inhibitor, clade A, member 8)      -0.0011273061613599608
GO:0033864: positive regulation of NAD(P)H oxidase activity     AGTR1   angiotensin II receptor, type
1       0.00034518432154975775
GO:0034104: negative regulation of tissue remodeling    AGT     angiotensinogen (serpin peptidase inh
ibitor, clade A, member 8)      -0.0011413244618989533
GO:0035411: catenin import into nucleus AGT     angiotensinogen (serpin peptidase inhibitor, clade A,
member 8)       -0.0011413244618989533
GO:0035813: regulation of renal sodium excretion        AGT     angiotensinogen (serpin peptidase inh
ibitor, clade A, member 8)      -0.001126686660165578
GO:0035813: regulation of renal sodium excretion        AGTR1   angiotensin II receptor, type 1 0.000
34571201260529845
GO:0035813: regulation of renal sodium excretion        AVP     arginine vasopressin    -0.0009321890
790038756
GO:0035815: positive regulation of renal sodium excretion       AGT     angiotensinogen (serpin pepti
dase inhibitor, clade A, member 8)      -0.001126036425668974
GO:0035815: positive regulation of renal sodium excretion       DRD2    dopamine receptor D2    -0.00
023290838230142068
GO:0040018: positive regulation of multicellular organism growth        AGT     angiotensinogen (serp
in peptidase inhibitor, clade A, member 8)      -0.0011253416474882984
GO:0040018: positive regulation of multicellular organism growth        BBS4    Bardet-Biedl syndrome
4       -0.000507321817866552
GO:0040018: positive regulation of multicellular organism growth        BCL2    B-cell CLL/lymphoma 2
-4.803184511103192e-6
GO:0040018: positive regulation of multicellular organism growth        CREB1   cAMP responsive eleme
nt binding protein 1    0.0006538172107678811
GO:0040018: positive regulation of multicellular organism growth        DRD2    dopamine receptor D2

-0.00023319793886676708
GO:0040018: positive regulation of multicellular organism growth        POU1F1  POU class 1 homeobox 1      0.0001365720790053925
GO:0040018: positive regulation of multicellular organism growth        SMO     smoothened, frizzled class receptor 0.002128862632710565
GO:0040018: positive regulation of multicellular organism growth        STAT5A  signal transducer and activator of transcription 5A   0.0015892017426617903
GO:0042127: regulation of cell proliferation    AGT     angiotensinogen (serpin peptidase inhibitor, clade A, member 8)     -0.001134182620963468
GO:0042127: regulation of cell proliferation    AGTR1   angiotensin II receptor, type 1 0.00034719940364801255
GO:0042127: regulation of cell proliferation    CHEK1   checkpoint kinase 1     0.0009009769811606011
GO:0042127: regulation of cell proliferation    CIB1    calcium and integrin binding 1 (calmyrin) 5.411435171859002e-5
GO:0042127: regulation of cell proliferation    CXCL10  chemokine (C-X-C motif) ligand 10       6.302211511482034e-5
GO:0042127: regulation of cell proliferation    CXCL13  chemokine (C-X-C motif) ligand 13       0.0029945756728374947
GO:0042127: regulation of cell proliferation    E2F4    E2F transcription factor 4, p107/p130-binding -0.0023463311704491623
GO:0042127: regulation of cell proliferation    ENG     endoglin        0.0008269777790662984
GO:0042127: regulation of cell proliferation    EZH2    enhancer of zeste 2 polycomb repressive complex 2 subunit    -0.00013584486316601662
GO:0042127: regulation of cell proliferation    FOXM1   forkhead box M1 0.0001978327308672379
GO:0042127: regulation of cell proliferation    GUCY2C  guanylate cyclase 2C (heat stable enterotoxin receptor)       -0.0016341243245748657
GO:0042127: regulation of cell proliferation    HOXD13  homeobox D13    -0.0005924310398957535
GO:0042127: regulation of cell proliferation    INHA    inhibin, alpha  0.00021149484023045943
GO:0042127: regulation of cell proliferation    JAG1    jagged 1        0.0017568436223542353
GO:0042127: regulation of cell proliferation    JAG2    jagged 2        -5.5988298097584525e-6
GO:0042127: regulation of cell proliferation    JAK2    Janus kinase 2  -3.144520313963049e-5
GO:0042127: regulation of cell proliferation    JUP     junction plakoglobin    -0.001479485583929459
GO:0042127: regulation of cell proliferation    KIT     v-kit Hardy-Zuckerman 4 feline sarcoma viral oncogene homolog       0.0002755503498915295
GO:0042127: regulation of cell proliferation    MMP7    matrix metallopeptidase 7 (matrilysin, uterine)      0.0004257983501875644
GO:0042127: regulation of cell proliferation    RPA3    replication protein A3, 14kDa   0.003138171179304926
GO:0042127: regulation of cell proliferation    SHH     sonic hedgehog  0.0006002971515527101
GO:0042127: regulation of cell proliferation    SIRT1   sirtuin 1       -1.518382478995567e-6
GO:0042127: regulation of cell proliferation    SIX3    SIX homeobox 3  0.0020385109905892216
GO:0042127: regulation of cell proliferation    SOX9    SRY (sex determining region Y)-box 9    -0.0005185173871042266
GO:0042127: regulation of cell proliferation    TFAP2C  transcription factor AP-2 gamma (activating enhancer binding protein 2 gamma)        0.0010611623802875964
GO:0042127: regulation of cell proliferation    TGFB3   transforming growth factor, beta 3      -0.0018249526898884544
GO:0042311: vasodilation        AGT     angiotensinogen (serpin peptidase inhibitor, clade A, member 8)     -0.0011265381176853437
GO:0042311: vasodilation        GPX1    glutathione peroxidase 1        0.00038570470654197825
GO:0043410: positive regulation of MAPK cascade AGT     angiotensinogen (serpin peptidase inhibitor, clade A, member 8)     -0.0011327409337338347
GO:0043410: positive regulation of MAPK cascade CDH2    cadherin 2, type 1, N-cadherin (neuronal) -0.0006448969576984283
GO:0043410: positive regulation of MAPK cascade CTNNB1  catenin (cadherin-associated protein), beta 1, 88kDa       -0.0001167548191732344
GO:0043410: positive regulation of MAPK cascade FGFR2   fibroblast growth factor receptor 2     0.000762806861838605
GO:0043410: positive regulation of MAPK cascade FGFR3   fibroblast growth factor receptor 3     0.00022251376683993427
GO:0043410: positive regulation of MAPK cascade FLT3    fms-related tyrosine kinase 3   -0.0006811615422099966
GO:0043410: positive regulation of MAPK cascade HMGB1   high mobility group box 1       -0.0007756731930923223
GO:0043410: positive regulation of MAPK cascade IGF1    insulin-like growth factor 1 (somatomedin C) 0.0001309116690577526
GO:0043410: positive regulation of MAPK cascade IGF1R   insulin-like growth factor 1 receptor   0.0010710913076569985
GO:0043410: positive regulation of MAPK cascade IGFBP3  insulin-like growth factor binding protein 3 0.0008360134930686311
GO:0043410: positive regulation of MAPK cascade IGFBP4  insulin-like growth factor binding protein 4 -0.0012776354032527765
GO:0043410: positive regulation of MAPK cascade INSR    insulin receptor        -0.001362366543641768
GO:0043410: positive regulation of MAPK cascade KIT     v-kit Hardy-Zuckerman 4 feline sarcoma viral oncogene homolog       0.00027491895017796975
GO:0043410: positive regulation of MAPK cascade LEP     leptin  0.0031889836225433912
GO:0043524: negative regulation of neuron apoptotic process     AGT     angiotensinogen (serpin peptidase inhibitor, clade A, member 8)      -0.001133237625270088
GO:0043524: negative regulation of neuron apoptotic process     ANGPT1  angiopoietin 1  0.0009006358254645089
GO:0043524: negative regulation of neuron apoptotic process     BAX     BCL2-associated X protein

-0.0004251634550346602
GO:0043524: negative regulation of neuron apoptotic process BCL2 B-cell CLL/lymphoma 2 -4.953281541930707e-6
GO:0043524: negative regulation of neuron apoptotic process CCL2 chemokine (C-C motif) ligand 2 0.0008165370033201197
GO:0043524: negative regulation of neuron apoptotic process CEBPB CCAAT/enhancer binding protein (C/EBP), beta -0.0002740039177823732
GO:0043524: negative regulation of neuron apoptotic process CITED1 Cbp/p300-interacting transactivator, with Glu/Asp-rich carboxy-terminal domain, 1 0.0028181002469272443
GO:0043524: negative regulation of neuron apoptotic process FGF8 fibroblast growth factor 8 (androgen-induced) 0.000983943107358738
GO:0043524: negative regulation of neuron apoptotic process GCLM glutamate-cysteine ligase, modifier subunit -0.003761309374100509
GO:0043524: negative regulation of neuron apoptotic process GDNF glial cell derived neurotrophic factor 0.0004447190543489691
GO:0043524: negative regulation of neuron apoptotic process HIPK2 homeodomain interacting protein kinase 2 0.0007707519811739049
GO:0043524: negative regulation of neuron apoptotic process HMOX1 heme oxygenase (decycling) 1 -0.0002176040508603668
GO:0043524: negative regulation of neuron apoptotic process HTT huntingtin -0.0009715676832402412
GO:0043524: negative regulation of neuron apoptotic process ISL1 ISL LIM homeobox 1 7.89267365097406e-5
GO:0043524: negative regulation of neuron apoptotic process JAK2 Janus kinase 2 -3.1710086875264245e-5
GO:0043524: negative regulation of neuron apoptotic process KIF14 kinesin family member 14 0.00048599383726066026
GO:0043524: negative regulation of neuron apoptotic process MDK midkine (neurite growth-promoting factor 2) 0.0016652841063607656
GO:0043524: negative regulation of neuron apoptotic process MEF2C myocyte enhancer factor 2C 0.0009733500044987471
GO:0043524: negative regulation of neuron apoptotic process MSH2 mutS homolog 2 0.0013922790619047267
GO:0043524: negative regulation of neuron apoptotic process NES nestin 0.0013735926438886211
GO:0043524: negative regulation of neuron apoptotic process NRP1 neuropilin 1 -0.0006432072180678691
GO:0043524: negative regulation of neuron apoptotic process RASA1 RAS p21 protein activator (GTPase activating protein) 1 -0.0003490480491911304
GO:0043524: negative regulation of neuron apoptotic process SIX1 SIX homeobox 1 -0.001869884356182418
GO:0043524: negative regulation of neuron apoptotic process STAMBP STAM binding protein -0.0017040826720829682
GO:0043524: negative regulation of neuron apoptotic process STAR steroidogenic acute regulatory protein 0.0008945065528124638
GO:0043524: negative regulation of neuron apoptotic process TGFB3 transforming growth factor, beta 3 -0.001823656656031718
GO:0043524: negative regulation of neuron apoptotic process TP73 tumor protein p73 0.0010278837325965122
GO:0043524: negative regulation of neuron apoptotic process WFS1 Wolfram syndrome 1 (wolframin) 0.0005767752251105922
GO:0044267: cellular protein metabolic process AGT angiotensinogen (serpin peptidase inhibitor, clade A, member 8) -0.0011298646866433663
GO:0044267: cellular protein metabolic process BCHE butyrylcholinesterase -6.363623417656086e-6
GO:0044267: cellular protein metabolic process BLM Bloom syndrome, RecQ helicase-like 0.0005233192652441642
GO:0044267: cellular protein metabolic process CCL2 chemokine (C-C motif) ligand 2 0.0008150974570626101
GO:0044267: cellular protein metabolic process CDKN2A cyclin-dependent kinase inhibitor 2A 0.0017540359393327823
GO:0044267: cellular protein metabolic process CPE carboxypeptidase E -0.0025393514208383347
GO:0044267: cellular protein metabolic process CUL7 cullin 7 -0.00011849200300347666
GO:0044267: cellular protein metabolic process EIF4G1 eukaryotic translation initiation factor 4 gamma, 1 0.0007410971874029052
GO:0044267: cellular protein metabolic process GALNT6 polypeptide N-acetylgalactosaminyltransferase 6 -0.0017719174850972363
GO:0044267: cellular protein metabolic process GCNT1 glucosaminyl (N-acetyl) transferase 1, core 2 0.0006921481954517552
GO:0044267: cellular protein metabolic process GCNT3 glucosaminyl (N-acetyl) transferase 3, mucin type -0.0023071710945766865
GO:0044267: cellular protein metabolic process GCNT4 glucosaminyl (N-acetyl) transferase 4, core 2 0.0004512824233740165
GO:0044267: cellular protein metabolic process IGF1 insulin-like growth factor 1 (somatomedin C) 0.00012999108687664686
GO:0044267: cellular protein metabolic process IGFBP1 insulin-like growth factor binding protein 1 0.00038737283713590826
GO:0044267: cellular protein metabolic process IGFBP2 insulin-like growth factor binding protein 2, 36kDa 0.0001441149318450228
GO:0044267: cellular protein metabolic process IGFBP3 insulin-like growth factor binding protein 3 0.0008356939504056863
GO:0044267: cellular protein metabolic process IGFBP4 insulin-like growth factor binding protein 4

-0.0012758392785670852
GO:0044267: cellular protein metabolic process  IGFBP6  insulin-like growth factor binding protein 6
-0.0020241788444113123
GO:0044267: cellular protein metabolic process  MMP2    matrix metallopeptidase 2 (gelatinase A, 72kDa gelatinase, 72kDa type IV collagenase)        -0.001128373983199132
GO:0044267: cellular protein metabolic process  MUC1    mucin 1, cell surface associated        0.0012824971077652503
GO:0044267: cellular protein metabolic process  MUC7    mucin 7, secreted       -0.003154680300308304
GO:0044267: cellular protein metabolic process  NUP153  nucleoporin 153kDa      0.0008231568379220467
GO:0044267: cellular protein metabolic process  PCSK2   proprotein convertase subtilisin/kexin type 2
0.0006041548261440691
GO:0044267: cellular protein metabolic process  PGM3    phosphoglucomutase 3    0.00198326723008385
GO:0044267: cellular protein metabolic process  PML     promyelocytic leukemia  -0.0006790064186019185
GO:0044267: cellular protein metabolic process  RAD21   RAD21 homolog (S. pombe)        -0.00010208722922996495
GO:0044267: cellular protein metabolic process  RPA1    replication protein A1, 70kDa   0.00043766112917028585
GO:0044267: cellular protein metabolic process  SEH1L   SEH1-like (S. cerevisiae)       -0.0007311997696591838
GO:0044267: cellular protein metabolic process  SMC1A   structural maintenance of chromosomes 1A
-0.0010182806749436464
GO:0044267: cellular protein metabolic process  SMC5    structural maintenance of chromosomes 5 -0.0007013261910983885
GO:0044267: cellular protein metabolic process  SMC6    structural maintenance of chromosomes 6 -0.0017478535236534372
GO:0044267: cellular protein metabolic process  SPHK1   sphingosine kinase 1    0.001807209072782821
GO:0044267: cellular protein metabolic process  THBS1   thrombospondin 1        -0.0010338199462024342
GO:0044267: cellular protein metabolic process  TPR     translocated promoter region, nuclear basket protein        -0.000695650520048142
GO:0044267: cellular protein metabolic process  WFS1    Wolfram syndrome 1 (wolframin)  0.0005760037162688469
GO:0044267: cellular protein metabolic process  XBP1    X-box binding protein 1 0.00026666778204140884
GO:0045429: positive regulation of nitric oxide biosynthetic process    AGT     angiotensinogen (serpin peptidase inhibitor, clade A, member 8)      -0.0011140948995178083
GO:0045429: positive regulation of nitric oxide biosynthetic process    AKT1    v-akt murine thymoma viral oncogene homolog 1       0.0007201755139836965
GO:0045429: positive regulation of nitric oxide biosynthetic process    AKT2    v-akt murine thymoma viral oncogene homolog 2       -0.000618964829837747
GO:0045429: positive regulation of nitric oxide biosynthetic process    ASS1    argininosuccinate synthase 1 0.00013797008423045557
GO:0045429: positive regulation of nitric oxide biosynthetic process    EGFR    epidermal growth factor receptor     0.0006742966701912761
GO:0045429: positive regulation of nitric oxide biosynthetic process    ESR1    estrogen receptor 1
-0.0009346559442157986
GO:0045429: positive regulation of nitric oxide biosynthetic process    ICAM1   intercellular adhesion molecule 1    0.0007229834608800636
GO:0045429: positive regulation of nitric oxide biosynthetic process    IFNG    interferon, gamma
-4.871539041186913e-5
GO:0045429: positive regulation of nitric oxide biosynthetic process    INSR    insulin receptor
-0.0013447000675839106
GO:0045429: positive regulation of nitric oxide biosynthetic process    JAK2    Janus kinase 2  -3.36323847778102e-5
GO:0045723: positive regulation of fatty acid biosynthetic process      AGT     angiotensinogen (serpin peptidase inhibitor, clade A, member 8)      -0.0011295254163965216
GO:0045723: positive regulation of fatty acid biosynthetic process      NR1H3   nuclear receptor subfamily 1, group H, member 3      0.0007982188691035323
GO:0046622: positive regulation of organ growth AGT     angiotensinogen (serpin peptidase inhibitor, clade A, member 8)     -0.0011348383041610308
GO:0046622: positive regulation of organ growth FGF8    fibroblast growth factor 8 (androgen-induced)
0.0009848521483020664
GO:0046622: positive regulation of organ growth IL7     interleukin 7   0.000872138404154457
GO:0046622: positive regulation of organ growth SMO     smoothened, frizzled class receptor     0.0021405929497700935
GO:0046622: positive regulation of organ growth YAP1    Yes-associated protein 1        -0.00016420445902064833
GO:0048143: astrocyte activation        AGT     angiotensinogen (serpin peptidase inhibitor, clade A, member 8)       -0.0011464001484450567
GO:0048143: astrocyte activation        SMO     smoothened, frizzled class receptor     0.002155616885929775
GO:0048144: fibroblast proliferation    AGT     angiotensinogen (serpin peptidase inhibitor, clade A, member 8)       -0.0011413244618989533
GO:0048169: regulation of long-term neuronal synaptic plasticity        AGT     angiotensinogen (serpin peptidase inhibitor, clade A, member 8)      -0.0011247664212435055
GO:0048169: regulation of long-term neuronal synaptic plasticity        DRD2    dopamine receptor D2
-0.00023294888617112795
GO:0048169: regulation of long-term neuronal synaptic plasticity        EGR1    early growth response 1       0.0010866701092259235
GO:0048659: smooth muscle cell proliferation    AGT     angiotensinogen (serpin peptidase inhibitor,

clade A, member 8)	-0.0011413244618989533
GO:0050663: cytokine secretion	AGT	angiotensinogen (serpin peptidase inhibitor, clade A, member 8)	-0.0011379983167102846
GO:0050663: cytokine secretion	LYN	LYN proto-oncogene, Src family tyrosine kinase	-0.0014235085791756512
GO:0050731: positive regulation of peptidyl-tyrosine phosphorylation	AGT	angiotensinogen (serpin peptidase inhibitor, clade A, member 8)	-0.0011283689576810593
GO:0050731: positive regulation of peptidyl-tyrosine phosphorylation	ANGPT1	angiopoietin 1	0.0008983233655464457
GO:0050731: positive regulation of peptidyl-tyrosine phosphorylation	EFNA1	ephrin-A1	-0.0005197146846847352
GO:0050731: positive regulation of peptidyl-tyrosine phosphorylation	FGF7	fibroblast growth factor 7	0.0006320722858868137
GO:0050731: positive regulation of peptidyl-tyrosine phosphorylation	ICAM1	intercellular adhesion molecule 1	0.0007278150678050746
GO:0050731: positive regulation of peptidyl-tyrosine phosphorylation	IGF1	insulin-like growth factor 1 (somatomedin C)	0.00013018510362595802
GO:0050731: positive regulation of peptidyl-tyrosine phosphorylation	JAK2	Janus kinase 2	-3.246110063992825e-5
GO:0050731: positive regulation of peptidyl-tyrosine phosphorylation	LRP8	low density lipoprotein receptor-related protein 8, apolipoprotein e receptor	-0.0009298787235284282
GO:0050731: positive regulation of peptidyl-tyrosine phosphorylation	NRP1	neuropilin 1	-0.0006410515846548176
GO:0050731: positive regulation of peptidyl-tyrosine phosphorylation	RELN	reelin	0.0015320091530040389
GO:0050731: positive regulation of peptidyl-tyrosine phosphorylation	TGFB1	transforming growth factor, beta 1	-7.247029046501877e-5
GO:0050731: positive regulation of peptidyl-tyrosine phosphorylation	THBS4	thrombospondin 4	-0.00044362526692384285
GO:0050731: positive regulation of peptidyl-tyrosine phosphorylation	TP53	tumor protein p53	0.0011712953670444948
GO:0050731: positive regulation of peptidyl-tyrosine phosphorylation	VEGFA	vascular endothelial growth factor A	0.0005921379831008294
GO:0050731: positive regulation of peptidyl-tyrosine phosphorylation	VEGFC	vascular endothelial growth factor C	-0.0033543365913086005
GO:0051145: smooth muscle cell differentiation	AGT	angiotensinogen (serpin peptidase inhibitor, clade A, member 8)	-0.001138583949785588
GO:0051145: smooth muscle cell differentiation	CTNNB1	catenin (cadherin-associated protein), beta 1, 88kDa	-0.00011669252822893965
GO:0051145: smooth muscle cell differentiation	GATA6	GATA binding protein 6	-2.744803819977668e-5
GO:0051145: smooth muscle cell differentiation	HEY2	hes-related family bHLH transcription factor with YRPW motif 2	0.0027079462603562827
GO:0051145: smooth muscle cell differentiation	MEF2C	myocyte enhancer factor 2C	0.0009766005189961668
GO:0051145: smooth muscle cell differentiation	WNT4	wingless-type MMTV integration site family, member 4	-0.0002541690884966172
GO:0051387: negative regulation of neurotrophin TRK receptor signaling pathway	AGT	angiotensinogen (serpin peptidase inhibitor, clade A, member 8)	-0.0011413244618989533
GO:0051403: stress-activated MAPK cascade	AGT	angiotensinogen (serpin peptidase inhibitor, clade A, member 8)	-0.0011381000621126767
GO:0051403: stress-activated MAPK cascade	CREB1	cAMP responsive element binding protein 1	0.0006612426418666483
GO:0051403: stress-activated MAPK cascade	CRYAB	crystallin, alpha B	0.0009881431402928255
GO:0051403: stress-activated MAPK cascade	IRAK1	interleukin-1 receptor-associated kinase 1	-0.00161383780582968
GO:0051403: stress-activated MAPK cascade	MAP3K19	mitogen-activated protein kinase kinase kinase 19	-0.0016542761039534373
GO:0051403: stress-activated MAPK cascade	MEF2C	myocyte enhancer factor 2C	0.0009767451355542315
GO:0051403: stress-activated MAPK cascade	RPS6KA1	ribosomal protein S6 kinase, 90kDa, polypeptide 1	-0.002520954957733001
GO:0051924: regulation of calcium ion transport	AGT	angiotensinogen (serpin peptidase inhibitor, clade A, member 8)	-0.0011194082351256936
GO:0051924: regulation of calcium ion transport	BCL2	B-cell CLL/lymphoma 2	-5.881241212492345e-6
GO:0051924: regulation of calcium ion transport	GJA1	gap junction protein, alpha 1, 43kDa	-0.0001585097141952876
GO:0051969: regulation of transmission of nerve impulse	AGT	angiotensinogen (serpin peptidase inhibitor, clade A, member 8)	-0.0011413244618989533
GO:0061049: cell growth involved in cardiac muscle cell development	AGT	angiotensinogen (serpin peptidase inhibitor, clade A, member 8)	-0.0011356652797173757
GO:0061049: cell growth involved in cardiac muscle cell development	GATA4	GATA binding protein 4	-0.0010955738267244865
GO:0061098: positive regulation of protein tyrosine kinase activity	AGT	angiotensinogen (serpin peptidase inhibitor, clade A, member 8)	-0.0011336556975768703
GO:0061098: positive regulation of protein tyrosine kinase activity	CD24	CD24 molecule	0.001071639702462 6808
GO:0061098: positive regulation of protein tyrosine kinase activity	GREM1	gremlin 1, DAN family BMP antagonist	-0.0008277527067870902
GO:0061098: positive regulation of protein tyrosine kinase activity	LRP8	low density lipoprotein receptor-related protein 8, apolipoprotein e receptor	-0.0009308798552369754
GO:0061098: positive regulation of protein tyrosine kinase activity	PTPN1	protein tyrosine phos

phatase, non-receptor type 1    -0.001041824103006975
GO:0061098: positive regulation of protein tyrosine kinase activity     RELN    reelin  0.00153698460
24346169
GO:0070371: ERK1 and ERK2 cascade       AGT     angiotensinogen (serpin peptidase inhibitor, clade A,
member 8)       -0.0011245650318756973
GO:0070371: ERK1 and ERK2 cascade       AVP     arginine vasopressin    -0.0009309082047241995
GO:0070371: ERK1 and ERK2 cascade       IGF1    insulin-like growth factor 1 (somatomedin C)    0.000
12932958021281922
GO:0070371: ERK1 and ERK2 cascade       MED1    mediator complex subunit 1      0.0011212181192247915
GO:0070371: ERK1 and ERK2 cascade       SOX9    SRY (sex determining region Y)-box 9    -0.0005150462
600783925
GO:0070471: uterine smooth muscle contraction   AGT     angiotensinogen (serpin peptidase inhibitor,
clade A, member 8)     -0.0011413244618989533
GO:0071260: cellular response to mechanical stimulus    AGT     angiotensinogen (serpin peptidase inh
ibitor, clade A, member 8)      -0.001130849001973462
GO:0071260: cellular response to mechanical stimulus    AKT1    v-akt murine thymoma viral oncogene h
omolog 1        0.0007290802436596668
GO:0071260: cellular response to mechanical stimulus    BAK1    BCL2-antagonist/killer 1        -0.00
1872236108850714
GO:0071260: cellular response to mechanical stimulus    BMP4    bone morphogenetic protein 4    -0.00
0322026651887928
GO:0071260: cellular response to mechanical stimulus    BNIP3   BCL2/adenovirus E1B 19kDa interacting
protein 3       0.0029070189298056165
GO:0071260: cellular response to mechanical stimulus    CASP8AP2        caspase 8 associated protein
2      0.0012695237955783517
GO:0071260: cellular response to mechanical stimulus    CAV1    caveolin 1, caveolae protein, 22kDa
-0.0005345261633046916
GO:0071260: cellular response to mechanical stimulus    CHEK1   checkpoint kinase 1     0.00089834208
85444516
GO:0071260: cellular response to mechanical stimulus    COL1A1  collagen, type I, alpha 1       -0.00
05252066452635591
GO:0071260: cellular response to mechanical stimulus    EGR1    early growth response 1 0.00109093848
85159701
GO:0071260: cellular response to mechanical stimulus    FAS     Fas cell surface death receptor -3.31
5298479412207e-5
GO:0071260: cellular response to mechanical stimulus    GJA1    gap junction protein, alpha 1, 43kDa
-0.00016123313902589005
GO:0071260: cellular response to mechanical stimulus    KCNJ2   potassium inwardly-rectifying channe
l, subfamily J, member 2        -0.0007752898729378255
GO:0071260: cellular response to mechanical stimulus    MMP7    matrix metallopeptidase 7 (matrilysi
n, uterine)     0.0004249327201853574
GO:0071260: cellular response to mechanical stimulus    SOX9    SRY (sex determining region Y)-box 9
-0.0005172766424310745
GO:0090190: positive regulation of branching involved in ureteric bud morphogenesis     AGT     angio
tensinogen (serpin peptidase inhibitor, clade A, member 8)      -0.001135879394108056
GO:0090190: positive regulation of branching involved in ureteric bud morphogenesis     GDNF    glial
cell derived neurotrophic factor        0.0004451213142344934
GO:0090190: positive regulation of branching involved in ureteric bud morphogenesis     GREM1   greml
in 1, DAN family BMP antagonist -0.0008294600210623037
GO:0090190: positive regulation of branching involved in ureteric bud morphogenesis     LGR4    leuci
ne-rich repeat containing G protein-coupled receptor 4  0.000308807182139114
GO:0090190: positive regulation of branching involved in ureteric bud morphogenesis     LHX1    LIM h
omeobox 1       -0.0007599456780626497
GO:0090190: positive regulation of branching involved in ureteric bud morphogenesis     PAX2    paire
d box 2 -0.001616570649938755
GO:0090190: positive regulation of branching involved in ureteric bud morphogenesis     PAX8    paire
d box 8 0.0009488374160705639
GO:0090190: positive regulation of branching involved in ureteric bud morphogenesis     SALL1   spalt
-like transcription factor 1    -0.0024421208757663335
GO:0090190: positive regulation of branching involved in ureteric bud morphogenesis     SIX1    SIX h
omeobox 1       -0.001872528142639172
GO:0090190: positive regulation of branching involved in ureteric bud morphogenesis     SMO     smoot
hened, frizzled class receptor  0.002142175255551561
GO:0090190: positive regulation of branching involved in ureteric bud morphogenesis     SOX9    SRY
(sex determining region Y)-box 9       -0.0005194276940881324
GO:0090190: positive regulation of branching involved in ureteric bud morphogenesis     TGFB1   trans
forming growth factor, beta 1   -7.324219900892433e-5
GO:0090190: positive regulation of branching involved in ureteric bud morphogenesis     VEGFA   vascu
lar endothelial growth factor A 0.0005962983015804187
GO:1900020: positive regulation of protein kinase C activity    AGT     angiotensinogen (serpin pepti
dase inhibitor, clade A, member 8)      -0.0011306837922656996
GO:1900020: positive regulation of protein kinase C activity    WNT5A   wingless-type MMTV integratio
n site family, member 5A        -0.0006657721693316111
GO:1901201: regulation of extracellular matrix assembly AGT     angiotensinogen (serpin peptidase inh
ibitor, clade A, member 8)      -0.0011289600098693805
GO:1901201: regulation of extracellular matrix assembly NOTCH1  notch 1 0.0005170083157855374
GO:1902632: positive regulation of membrane hyperpolarization   AGT     angiotensinogen (serpin pepti
dase inhibitor, clade A, member 8)      -0.0011413244618989533
GO:1903598: positive regulation of gap junction assembly        AGT     angiotensinogen (serpin pepti
dase inhibitor, clade A, member 8)      -0.001127421096015043
GO:1903598: positive regulation of gap junction assembly        CAV1    caveolin 1, caveolae protein,

22kDa   -0.000533000302787689
GO:1903779: regulation of cardiac conduction    AGT     angiotensinogen (serpin peptidase inhibitor, clade A, member 8)     -0.0011413244618989533
GO:2000650: negative regulation of sodium ion transmembrane transporter activity        AGT     angiotensinogen (serpin peptidase inhibitor, clade A, member 8)      -0.0011440163609418427
GO:2000650: negative regulation of sodium ion transmembrane transporter activity        NEDD4   neural precursor cell expressed, developmentally down-regulated 4, E3 ubiquitin protein ligase       0.0022756847164011945
GO:2001238: positive regulation of extrinsic apoptotic signaling pathway        AGT     angiotensinogen (serpin peptidase inhibitor, clade A, member 8)      -0.0011188591065510772
GO:2001238: positive regulation of extrinsic apoptotic signaling pathway        CAV1    caveolin 1, caveolae protein, 22kDa  -0.0005313378843129163
GO:2001238: positive regulation of extrinsic apoptotic signaling pathway        PML     promyelocytic leukemia        -0.0006749960214677707
KEGG:04080: Neuroactive ligand-receptor interaction     AGTR1   angiotensin II receptor, type 1 0.0003505979712411405
KEGG:04080: Neuroactive ligand-receptor interaction     CHRNB1  cholinergic receptor, nicotinic, beta 1 (muscle)      -7.444182280315222e-6
KEGG:04080: Neuroactive ligand-receptor interaction     CHRNB2  cholinergic receptor, nicotinic, beta 2 (neuronal)    -0.0007775952737220377
KEGG:04080: Neuroactive ligand-receptor interaction     CSH1    chorionic somatomammotropin hormone 1 (placental lactogen)    -0.0002574152848051912
KEGG:04080: Neuroactive ligand-receptor interaction     DRD2    dopamine receptor D2    -0.00023368620618888992
KEGG:04080: Neuroactive ligand-receptor interaction     DRD4    dopamine receptor D4    -0.0019435849857920122
KEGG:04080: Neuroactive ligand-receptor interaction     GABRA4  gamma-aminobutyric acid (GABA) A receptor, alpha 4   -0.00148262966614 3303
KEGG:04080: Neuroactive ligand-receptor interaction     GNRHR   gonadotropin-releasing hormone receptor      0.0007407863776446282
KEGG:04080: Neuroactive ligand-receptor interaction     GRIK5   glutamate receptor, ionotropic, kainate 5    -0.0002329090563643156
KEGG:04080: Neuroactive ligand-receptor interaction     HTR2C   5-hydroxytryptamine (serotonin) receptor 2C, G protein-coupled       -0.0005246970516044352
KEGG:04080: Neuroactive ligand-receptor interaction     HTR6    5-hydroxytryptamine (serotonin) receptor 6, G protein-coupled        -0.001081363304210822
KEGG:04080: Neuroactive ligand-receptor interaction     LEP     leptin  0.0031865997634206895
KEGG:04080: Neuroactive ligand-receptor interaction     MAS1    MAS1 proto-oncogene, G protein-coupled receptor      0.0002931804741008749
KEGG:04080: Neuroactive ligand-receptor interaction     PRLR    prolactin receptor      0.0021826627020479807
KEGG:04080: Neuroactive ligand-receptor interaction     THRA    thyroid hormone receptor, alpha 0.000763370738236480 9
KEGG:04080: Neuroactive ligand-receptor interaction     THRB    thyroid hormone receptor, beta  0.0019430701777553032
KEGG:04270: Vascular smooth muscle contraction  AGTR1   angiotensin II receptor, type 1 0.00034777176244726307
KEGG:04270: Vascular smooth muscle contraction  ITPR1   inositol 1,4,5-trisphosphate receptor, type 1 -0.0007726285830895636
KEGG:04270: Vascular smooth muscle contraction  KCNMA1  potassium large conductance calcium-activated channel, subfamily M, alpha member 1    -0.0006502154596969317
KEGG:04270: Vascular smooth muscle contraction  PLA2G5  phospholipase A2, group V       0.002063440001743987
KEGG:04270: Vascular smooth muscle contraction  PLCB1   phospholipase C, beta 1 (phosphoinositide-specific)  0.00015076845374553335
KEGG:04270: Vascular smooth muscle contraction  PRKCA   protein kinase C, alpha -5.907479652625517e-6
KEGG:04270: Vascular smooth muscle contraction  PRKCD   protein kinase C, delta -0.0011175627256523005
KEGG:04270: Vascular smooth muscle contraction  PRKX    protein kinase, X-linked        0.0003893818631832804
KEGG:04270: Vascular smooth muscle contraction  RAF1    Raf-1 proto-oncogene, serine/threonine kinase 0.0014867845994448316
KEGG:04270: Vascular smooth muscle contraction  RHOA    ras homolog family member A     0.0006644111183773567
KEGG:05200: Pathways in cancer  AGTR1   angiotensin II receptor, type 1 0.0003475353240859817
KEGG:05200: Pathways in cancer  AKT1    v-akt murine thymoma viral oncogene homolog 1   0.0007292479628088175
KEGG:05200: Pathways in cancer  AKT2    v-akt murine thymoma viral oncogene homolog 2   -0.0006249550526819739
KEGG:05200: Pathways in cancer  APC     adenomatous polyposis coli      0.0006430765155879746
KEGG:05200: Pathways in cancer  AR      androgen receptor       0.0026342428208375097
KEGG:05200: Pathways in cancer  AXIN1   axin 1  -0.0007317074748467241
KEGG:05200: Pathways in cancer  BAX     BCL2-associated X protein       -0.0004245212723179917
KEGG:05200: Pathways in cancer  BCL2    B-cell CLL/lymphoma 2   -5.116964969240721e-6
KEGG:05200: Pathways in cancer  BIRC5   baculoviral IAP repeat containing 5     -0.0002601839706338840 7
KEGG:05200: Pathways in cancer  BMP4    bone morphogenetic protein 4    -0.0003220751179902788 5
KEGG:05200: Pathways in cancer  BRCA2   breast cancer 2, early onset    -1.0177525361501894e-5
KEGG:05200: Pathways in cancer  CCNA1   cyclin A1       -0.0017566511546665898
KEGG:05200: Pathways in cancer  CCND1   cyclin D1       -0.002627716223086892
KEGG:05200: Pathways in cancer  CCNE1   cyclin E1       0.0003702560262258885 5

KEGG:05200: Pathways in cancer  CCNE2   cyclin E2       0.0011049610661534755
KEGG:05200: Pathways in cancer  CDC42   cell division cycle 42  0.0012381436869618634
KEGG:05200: Pathways in cancer  CDKN2A  cyclin-dependent kinase inhibitor 2A    0.001754834093439353
KEGG:05200: Pathways in cancer  CDKN2B  cyclin-dependent kinase inhibitor 2B (p15, inhibits CDK4)
-0.0018324229590384968
KEGG:05200: Pathways in cancer  CKS2    CDC28 protein kinase regulatory subunit 2       -4.8289909527
43828e-5
KEGG:05200: Pathways in cancer  CSF3R   colony stimulating factor 3 receptor (granulocyte)      -8.28
5031067242025e-5
KEGG:05200: Pathways in cancer  CTNNB1  catenin (cadherin-associated protein), beta 1, 88kDa    -0.00
011680281936528776
KEGG:05200: Pathways in cancer  CXCL12  chemokine (C-X-C motif) ligand 12       -0.001206308219044905
KEGG:05200: Pathways in cancer  CXCR4   chemokine (C-X-C motif) receptor 4      0.0007932901940236521
KEGG:05200: Pathways in cancer  E2F1    E2F transcription factor 1      0.002094094751964618 5
KEGG:05200: Pathways in cancer  E2F3    E2F transcription factor 3      0.00215852160526122 2
KEGG:05200: Pathways in cancer  EGFR    epidermal growth factor receptor        0.0006843054887895298
KEGG:05200: Pathways in cancer  FAS     Fas cell surface death receptor -3.318796770827798e-5
KEGG:05200: Pathways in cancer  FGF3    fibroblast growth factor 3      0.0015583623166298707
KEGG:05200: Pathways in cancer  FGF5    fibroblast growth factor 5      -0.0010922603319737509
KEGG:05200: Pathways in cancer  FGF7    fibroblast growth factor 7      0.0006326358006618651
KEGG:05200: Pathways in cancer  FGF8    fibroblast growth factor 8 (androgen-induced)   0.00098205648
25246724
KEGG:05200: Pathways in cancer  FGFR2   fibroblast growth factor receptor 2     0.0007618709844360251
KEGG:05200: Pathways in cancer  FGFR3   fibroblast growth factor receptor 3     0.0002219821041618055
KEGG:05200: Pathways in cancer  FLT3    fms-related tyrosine kinase 3   -0.0006801293417308778
KEGG:05200: Pathways in cancer  FOXO1   forkhead box O1 0.0017972413502108642
KEGG:05200: Pathways in cancer  FZD3    frizzled class receptor 3       0.00037754096218346343
KEGG:05200: Pathways in cancer  FZD7    frizzled class receptor 7       0.0010684524267841435
KEGG:05200: Pathways in cancer  GLI1    GLI family zinc finger 1        -0.0012948875572445198
KEGG:05200: Pathways in cancer  GLI2    GLI family zinc finger 2        0.0018503085215558056
KEGG:05200: Pathways in cancer  GLI3    GLI family zinc finger 3        -0.002147785799533418 4
KEGG:05200: Pathways in cancer  GRB2    growth factor receptor-bound protein 2  0.0004289053722929083
KEGG:05200: Pathways in cancer  GSK3B   glycogen synthase kinase 3 beta 0.0015464803123016698
KEGG:05200: Pathways in cancer  GSTP1   glutathione S-transferase pi 1  0.00023115625791369747
KEGG:05200: Pathways in cancer  HDAC2   histone deacetylase 2   -0.0012055335931622562
KEGG:05200: Pathways in cancer  HIF1A   hypoxia inducible factor 1, alpha subunit (basic helix-loop-h
elix transcription factor)      -0.0006455051398532944
KEGG:05200: Pathways in cancer  IGF1    insulin-like growth factor 1 (somatomedin C)    0.00013034899
531345497
KEGG:05200: Pathways in cancer  IGF1R   insulin-like growth factor 1 receptor   0.0010705675369078608
KEGG:05200: Pathways in cancer  ITGA3   integrin, alpha 3 (antigen CD49C, alpha 3 subunit of VLA-3 re
ceptor) 0.0013975734579825929
KEGG:05200: Pathways in cancer  ITGA6   integrin, alpha 6       0.0016654431816611522
KEGG:05200: Pathways in cancer  ITGB1   integrin, beta 1 (fibronectin receptor, beta polypeptide, ant
igen CD29 includes MDF2, MSK12) 0.0024220572218738795
KEGG:05200: Pathways in cancer  JUP     junction plakoglobin    -0.001476930825441092
KEGG:05200: Pathways in cancer  KIT     v-kit Hardy-Zuckerman 4 feline sarcoma viral oncogene homolog
0.00027388354277701126
KEGG:05200: Pathways in cancer  LAMB2   laminin, beta 2 (laminin S)     -0.001306699058154702
KEGG:05200: Pathways in cancer  LEF1    lymphoid enhancer-binding factor 1      -0.000100325958478382
4
KEGG:05200: Pathways in cancer  MLH1    mutL homolog 1  0.0023325033266400016
KEGG:05200: Pathways in cancer  MMP2    matrix metallopeptidase 2 (gelatinase A, 72kDa gelatinase, 72
kDa type IV collagenase)        -0.001129030532031634
KEGG:05200: Pathways in cancer  MSH2    mutS homolog 2  0.0013896006625863
KEGG:05200: Pathways in cancer  MSH3    mutS homolog 3  -0.0005297582184598424
KEGG:05200: Pathways in cancer  MSH6    mutS homolog 6  0.001075835573800222
KEGG:05200: Pathways in cancer  MYC     v-myc avian myelocytomatosis viral oncogene homolog     -0.00
11346829922679583
KEGG:05200: Pathways in cancer  PAX8    paired box 8    0.0009469430853739513
KEGG:05200: Pathways in cancer  PDGFRB  platelet-derived growth factor receptor, beta polypeptide
-0.00036020480106274645
KEGG:05200: Pathways in cancer  PIK3CD  phosphatidylinositol-4,5-bisphosphate 3-kinase, catalytic sub
unit delta      -0.00077950179044433
KEGG:05200: Pathways in cancer  PLCB1   phospholipase C, beta 1 (phosphoinositide-specific)     0.000
15327982703546234
KEGG:05200: Pathways in cancer  PML     promyelocytic leukemia  -0.000679134335794394
KEGG:05200: Pathways in cancer  PRKCA   protein kinase C, alpha -5.867986599925351e-6
KEGG:05200: Pathways in cancer  PRKX    protein kinase, X-linked        0.0003937998854287269
KEGG:05200: Pathways in cancer  PTCH1   patched 1       -6.599530310629093e-5
KEGG:05200: Pathways in cancer  PTEN    phosphatase and tensin homolog  1.7445182939433086e-5
KEGG:05200: Pathways in cancer  RAD51   RAD51 recombinase       -0.0018582796598198698
KEGG:05200: Pathways in cancer  RAF1    Raf-1 proto-oncogene, serine/threonine kinase   0.00149632993
09306967
KEGG:05200: Pathways in cancer  RB1     retinoblastoma 1        -0.0014914358284641802
KEGG:05200: Pathways in cancer  RET     ret proto-oncogene      -0.0004905200765679358
KEGG:05200: Pathways in cancer  RHOA    ras homolog family member A     0.0006686522994744405
KEGG:05200: Pathways in cancer  RXRA    retinoid X receptor, alpha      0.0011132437576497677
KEGG:05200: Pathways in cancer  SHH     sonic hedgehog  0.0005986343010761835
KEGG:05200: Pathways in cancer  SKP2    S-phase kinase-associated protein 2, E3 ubiquitin protein lig
ase     -0.0005362216081415746

KEGG:05200: Pathways in cancer  SMO     smoothened, frizzled class receptor     0.0021356217119138377
KEGG:05200: Pathways in cancer  STAT5A  signal transducer and activator of transcription 5A     0.001
5937308840572014
KEGG:05200: Pathways in cancer  TCF7L2  transcription factor 7-like 2 (T-cell specific, HMG-box)
0.0005725618128288142
KEGG:05200: Pathways in cancer  TGFB1   transforming growth factor, beta 1      -7.283372914293484e-5
KEGG:05200: Pathways in cancer  TGFB2   transforming growth factor, beta 2      -0.001057378518212652
1
KEGG:05200: Pathways in cancer  TGFB3   transforming growth factor, beta 3      -0.001820325400538047
8
KEGG:05200: Pathways in cancer  TGFBR1  transforming growth factor, beta receptor 1     0.00033918499
231240317
KEGG:05200: Pathways in cancer  TP53    tumor protein p53       0.0011732916887733892
KEGG:05200: Pathways in cancer  TPR     translocated promoter region, nuclear basket protein    -0.00
06959176399215367
KEGG:05200: Pathways in cancer  VEGFA   vascular endothelial growth factor A    0.0005936513709879258
KEGG:05200: Pathways in cancer  VEGFC   vascular endothelial growth factor C    -0.003358503626733515
4
KEGG:05200: Pathways in cancer  WNT1    wingless-type MMTV integration site family, member 1    0.000
7856803371034068
KEGG:05200: Pathways in cancer  WNT10B  wingless-type MMTV integration site family, member 10B  -0.00
13859491656375256
KEGG:05200: Pathways in cancer  WNT4    wingless-type MMTV integration site family, member 4    -0.00
02512588301347477
KEGG:05200: Pathways in cancer  WNT5A   wingless-type MMTV integration site family, member 5A   -0.00
066626235155091
KEGG:05200: Pathways in cancer  WNT7A   wingless-type MMTV integration site family, member 7A   1.960
9304101290968e-5
KEGG:04022: cGMP-PKG signaling pathway  AGTR1   angiotensin II receptor, type 1 0.0003485926894849319
3
KEGG:04022: cGMP-PKG signaling pathway  AKT1    v-akt murine thymoma viral oncogene homolog 1   0.000
7225817719238702
KEGG:04022: cGMP-PKG signaling pathway  AKT2    v-akt murine thymoma viral oncogene homolog 2   -0.00
06210979171807566
KEGG:04022: cGMP-PKG signaling pathway  ATP1B3  ATPase, Na+/K+ transporting, beta 3 polypeptide -0.00
14853337829943082
KEGG:04022: cGMP-PKG signaling pathway  CREB1   cAMP responsive element binding protein 1       0.000
648961978164158
KEGG:04022: cGMP-PKG signaling pathway  GATA4   GATA binding protein 4  -0.0010834492017024245
KEGG:04022: cGMP-PKG signaling pathway  INSR    insulin receptor        -0.0013487715813365355
KEGG:04022: cGMP-PKG signaling pathway  ITPR1   inositol 1,4,5-trisphosphate receptor, type 1   -0.00
07704561243640077
KEGG:04022: cGMP-PKG signaling pathway  KCNMA1  potassium large conductance calcium-activated channe
l, subfamily M, alpha member 1  -0.000647997961960089
KEGG:04022: cGMP-PKG signaling pathway  MEF2C   myocyte enhancer factor 2C      0.0009623584081566344
KEGG:04022: cGMP-PKG signaling pathway  PIK3CD  phosphatidylinositol-4,5-bisphosphate 3-kinase, catal
ytic subunit delta      -0.000773199492227543
KEGG:04022: cGMP-PKG signaling pathway  PLCB1   phospholipase C, beta 1 (phosphoinositide-specific)
0.00015005615254737358
KEGG:04022: cGMP-PKG signaling pathway  RAF1    Raf-1 proto-oncogene, serine/threonine kinase   0.001
484508621691188
KEGG:04022: cGMP-PKG signaling pathway  RHOA    ras homolog family member A     0.0006635702649900566
KEGG:04261: Adrenergic signaling in cardiomyocytes      AGTR1   angiotensin II receptor, type 1 0.000
3485667211912762
KEGG:04261: Adrenergic signaling in cardiomyocytes      AKT1    v-akt murine thymoma viral oncogene h
omolog 1        0.0007080662896574123
KEGG:04261: Adrenergic signaling in cardiomyocytes      AKT2    v-akt murine thymoma viral oncogene h
omolog 2        -0.0006121077292545785
KEGG:04261: Adrenergic signaling in cardiomyocytes      ATP1B3  ATPase, Na+/K+ transporting, beta 3 p
olypeptide      -0.0014621576289393062
KEGG:04261: Adrenergic signaling in cardiomyocytes      BCL2    B-cell CLL/lymphoma 2   -5.5446879758
74964e-6
KEGG:04261: Adrenergic signaling in cardiomyocytes      CACNB3  calcium channel, voltage-dependent, b
eta 3 subunit   0.00020853217890244732
KEGG:04261: Adrenergic signaling in cardiomyocytes      CREB1   cAMP responsive element binding prote
in 1    0.0006326820546893797
KEGG:04261: Adrenergic signaling in cardiomyocytes      PIK3CD  phosphatidylinositol-4,5-bisphosphate
3-kinase, catalytic subunit delta       -0.0007594704790757685
KEGG:04261: Adrenergic signaling in cardiomyocytes      PLCB1   phospholipase C, beta 1 (phosphoinosi
tide-specific)  0.00014373582046099694
KEGG:04261: Adrenergic signaling in cardiomyocytes      PPP2R3A protein phosphatase 2, regulatory sub
unit B'', alpha -4.216054407415247e-5
KEGG:04261: Adrenergic signaling in cardiomyocytes      PRKCA   protein kinase C, alpha -6.1538647455
54917e-6
KEGG:04261: Adrenergic signaling in cardiomyocytes      PRKX    protein kinase, X-linked        0.000
37626611359626975
KEGG:04020: Calcium signaling pathway   AGTR1   angiotensin II receptor, type 1 0.0003466102480739934
4
KEGG:04020: Calcium signaling pathway   EGFR    epidermal growth factor receptor        0.00067196196
79738327
KEGG:04020: Calcium signaling pathway   ERBB4   v-erb-b2 avian erythroblastic leukemia viral oncogene

homolog 4	-0.00017706689218756385
KEGG:04020: Calcium signaling pathway	HTR2C	5-hydroxytryptamine (serotonin) receptor 2C, G protein-coupled	-0.0005172444913433181
KEGG:04020: Calcium signaling pathway	HTR6	5-hydroxytryptamine (serotonin) receptor 6, G protein-coupled	-0.0010658225355745852
KEGG:04020: Calcium signaling pathway	ITPR1	inositol 1,4,5-trisphosphate receptor, type 1	-0.00076669115042475
KEGG:04020: Calcium signaling pathway	ORAI2	ORAI calcium release-activated calcium modulator 2	0.0007224552284418564
KEGG:04020: Calcium signaling pathway	ORAI3	ORAI calcium release-activated calcium modulator 3	-0.0008751577028992491
KEGG:04020: Calcium signaling pathway	PDGFRB	platelet-derived growth factor receptor, beta polypeptide	-0.0003518927572837588
KEGG:04020: Calcium signaling pathway	PLCB1	phospholipase C, beta 1 (phosphoinositide-specific)	0.0001486891189611444
KEGG:04020: Calcium signaling pathway	PRKCA	protein kinase C, alpha -5.844379105646469e-6
KEGG:04020: Calcium signaling pathway	PRKX	protein kinase, X-linked	0.0003859988348092519
KEGG:04020: Calcium signaling pathway	SPHK1	sphingosine kinase 1	0.0017831484148631688
GO:0003081: regulation of systemic arterial blood pressure by renin-angiotensin AGTR1	angiotensin II receptor, type 1	0.00034087272072047303
GO:0007266: Rho protein signal transduction	AGTR1	angiotensin II receptor, type 1 0.00034745936097046185
GO:0007266: Rho protein signal transduction	CFL1	cofilin 1 (non-muscle)	-0.0011751921834503025
GO:0007266: Rho protein signal transduction	CNKSR1	connector enhancer of kinase suppressor of Ras 1	-0.0011061531138089678
GO:0007266: Rho protein signal transduction	PAK1	p21 protein (Cdc42/Rac)-activated kinase 1	-0.002187846403390788
GO:0007266: Rho protein signal transduction	PAK3	p21 protein (Cdc42/Rac)-activated kinase 3	-0.001258658592141223
GO:0007266: Rho protein signal transduction	PHACTR4 phosphatase and actin regulator 4	-0.00179095898007962
GO:0007266: Rho protein signal transduction	RHOA	ras homolog family member A	0.0006697706205168874
GO:0007266: Rho protein signal transduction	ROPN1B	rhophilin associated tail protein 1B	0.0005978015550404625
GO:0019722: calcium-mediated signaling	AGTR1	angiotensin II receptor, type 1 0.00034708652049723904
GO:0019722: calcium-mediated signaling	CXCR4	chemokine (C-X-C motif) receptor 4	0.0007919562854707339
GO:0019722: calcium-mediated signaling	SPHK1	sphingosine kinase 1	0.0018046616123086464
GO:0032430: positive regulation of phospholipase A2 activity	AGTR1	angiotensin II receptor, type 1	0.00034087272072047303
GO:0038166: angiotensin-activated signaling pathway	AGTR1	angiotensin II receptor, type 1 0.0003348582837193237
GO:0038166: angiotensin-activated signaling pathway	MAS1	MAS1 proto-oncogene, G protein-coupled receptor	0.0003017651563656741
GO:0042312: regulation of vasodilation	AGTR1	angiotensin II receptor, type 1 0.00034087272072047303
GO:0050727: regulation of inflammatory response AGTR1	angiotensin II receptor, type 1 0.00034965335038647933
GO:0050727: regulation of inflammatory response BRD4	bromodomain containing 4	0.0005486325697231976
GO:0050727: regulation of inflammatory response JAK2	Janus kinase 2	-3.679057871386259e-5
GO:0050727: regulation of inflammatory response LYN	LYN proto-oncogene, Src family tyrosine kinase	-0.0013931861534899259
GO:0050727: regulation of inflammatory response MAS1	MAS1 proto-oncogene, G protein-coupled receptor	0.0002963608384649916
GO:0050727: regulation of inflammatory response TNFAIP3 tumor necrosis factor, alpha-induced protein 3	0.0010936282909919455
GO:0051482: positive regulation of cytosolic calcium ion concentration involved in phospholipase C-activating G-protein coupled signaling pathway	AGTR1	angiotensin II receptor, type 1 0.000349560040539391
GO:0051482: positive regulation of cytosolic calcium ion concentration involved in phospholipase C-activating G-protein coupled signaling pathway	DRD2	dopamine receptor D2	-0.0002216842067900553
GO:0051482: positive regulation of cytosolic calcium ion concentration involved in phospholipase C-activating G-protein coupled signaling pathway	HTR2C	5-hydroxytryptamine (serotonin) receptor 2C, G protein-coupled	-0.0004873807284922389
GO:0060326: cell chemotaxis	AGTR1	angiotensin II receptor, type 1 0.0003469201288968819
GO:0060326: cell chemotaxis	AZU1	azurocidin 1	0.00019664738122811862
GO:0060326: cell chemotaxis	BIN2	bridging integrator 2	-0.001634800364921307
GO:0060326: cell chemotaxis	CXCL12	chemokine (C-X-C motif) ligand 12	-0.0012011471703642527
GO:0060326: cell chemotaxis	ENG	endoglin	0.0008205391086668363
GO:0060326: cell chemotaxis	EPHB1	EPH receptor B1 0.0019525787321272947
GO:0060326: cell chemotaxis	HMGB2	high mobility group box 2	0.00030318353180710725
GO:0060326: cell chemotaxis	KIT	v-kit Hardy-Zuckerman 4 feline sarcoma viral oncogene homolog	0.0002717553072549106З
GO:0060326: cell chemotaxis	LEF1	lymphoid enhancer-binding factor 1	-0.0001004153180837496З

GO:0060326: cell chemotaxis PDGFRB platelet-derived growth factor receptor, beta polypeptide -0.0003579372517893117
GO:0060326: cell chemotaxis PRKCD protein kinase C, delta -0.0011208776123495451
GO:0086097: phospholipase C-activating angiotensin-activated signaling pathway AGTR1 angiotensin II receptor, type 1 0.00034087272072047303
GO:0006418: tRNA aminoacylation for protein translation AIMP2 aminoacyl tRNA synthetase complex-interacting multifunctional protein 2 0.0011874080425433111
GO:0006418: tRNA aminoacylation for protein translation LARS2 leucyl-tRNA synthetase 2, mitochondrial 0.00044953839655748247
GO:0006915: apoptotic process AIMP2 aminoacyl tRNA synthetase complex-interacting multifunctional protein 2 0.0012090167990556286
GO:0006915: apoptotic process AKT1 v-akt murine thymoma viral oncogene homolog 1 0.0007302741753768608
GO:0006915: apoptotic process APC adenomatous polyposis coli 0.0006437735456922245
GO:0006915: apoptotic process AXIN1 axin 1 -0.0007327393983158618
GO:0006915: apoptotic process BAK1 BCL2-antagonist/killer 1 -0.0018753961632275344
GO:0006915: apoptotic process BAX BCL2-associated X protein -0.00042514058779966464
GO:0006915: apoptotic process BCL2 B-cell CLL/lymphoma 2 -4.9777653506392396e-6
GO:0006915: apoptotic process BCL2L11 BCL2-like 11 (apoptosis facilitator) -0.000774660858702622
GO:0006915: apoptotic process BFAR bifunctional apoptosis regulator -0.000369362576607406 6
GO:0006915: apoptotic process BNIP3 BCL2/adenovirus E1B 19kDa interacting protein 3 0.0029113762225990874
GO:0006915: apoptotic process BUB1 BUB1 mitotic checkpoint serine/threonine kinase 0.001324194682342641
GO:0006915: apoptotic process BUB1B BUB1 mitotic checkpoint serine/threonine kinase B -0.001281108354915063
GO:0006915: apoptotic process CDK1 cyclin-dependent kinase 1 0.00031063400550536944
GO:0006915: apoptotic process CIB1 calcium and integrin binding 1 (calmyrin) 5.498901725151033e-5
GO:0006915: apoptotic process CITED1 Cbp/p300-interacting transactivator, with Glu/Asp-rich carboxy-terminal domain, 1 0.00281694641768821
GO:0006915: apoptotic process CKAP2 cytoskeleton associated protein 2 4.881591577572257e-5
GO:0006915: apoptotic process CST3 cystatin C -6.975577470499725e-5
GO:0006915: apoptotic process CTNNB1 catenin (cadherin-associated protein), beta 1, 88kDa -0.0001170206239645121
GO:0006915: apoptotic process CXCR4 chemokine (C-X-C motif) receptor 4 0.000794235602377018
GO:0006915: apoptotic process DAXX death-domain associated protein 0.0008910378855435978
GO:0006915: apoptotic process DHCR24 24-dehydrocholesterol reductase -0.001706753077782305
GO:0006915: apoptotic process DIABLO diablo, IAP-binding mitochondrial protein -0.0005955950874973314
GO:0006915: apoptotic process DOCK1 dedicator of cytokinesis 1 -0.0006838636074784756
GO:0006915: apoptotic process DSG2 desmoglein 2 1.668456633629192e-5
GO:0006915: apoptotic process DSP desmoplakin -5.666357773000137e-5
GO:0006915: apoptotic process E2F1 E2F transcription factor 1 0.0020967558273057435
GO:0006915: apoptotic process EDAR ectodysplasin A receptor 0.0005937664990297261
GO:0006915: apoptotic process ESPL1 extra spindle pole bodies homolog 1 (S. cerevisiae) 0.00019668379559849655
GO:0006915: apoptotic process FAS Fas cell surface death receptor -3.326835424946714e-5
GO:0006915: apoptotic process FEM1B fem-1 homolog b (C. elegans) -0.004129437588366252
GO:0006915: apoptotic process FGF8 fibroblast growth factor 8 (androgen-induced) 0.0009834224895683026
GO:0006915: apoptotic process FGFR2 fibroblast growth factor receptor 2 0.0007628372960817822
GO:0006915: apoptotic process FOXO1 forkhead box O1 0.0017995590026224561
GO:0006915: apoptotic process GJA1 gap junction protein, alpha 1, 43kDa -0.00016112545416969718
GO:0006915: apoptotic process GLRX2 glutaredoxin 2 0.00111569509792518
GO:0006915: apoptotic process GREM1 gremlin 1, DAN family BMP antagonist -0.0008267059407351259
GO:0006915: apoptotic process HMGB1 high mobility group box 1 -0.0007762934518108861
GO:0006915: apoptotic process HMGB2 high mobility group box 2 0.00030641277187692316
GO:0006915: apoptotic process IFNA2 interferon, alpha 2 -0.001716644611279821
GO:0006915: apoptotic process IFNG interferon, gamma -4.8932342679812706e-5
GO:0006915: apoptotic process IGFBP3 insulin-like growth factor binding protein 3 0.0008366433195386398
GO:0006915: apoptotic process JAK2 Janus kinase 2 -3.197876945434212e-5
GO:0006915: apoptotic process KLF11 Kruppel-like factor 11 0.0006160522158830201
GO:0006915: apoptotic process KMT2A lysine (K)-specific methyltransferase 2A 0.0006603287159805295
GO:0006915: apoptotic process KPNB1 karyopherin (importin) beta 1 0.0007716663649921333
GO:0006915: apoptotic process MAP1S microtubule-associated protein 1S 0.0005305226458520358
GO:0006915: apoptotic process MAPT microtubule-associated protein tau 0.0015512119513917224
GO:0006915: apoptotic process MEF2C myocyte enhancer factor 2C 0.0009728470760646271
GO:0006915: apoptotic process MELK maternal embryonic leucine zipper kinase 0.002068095947441584
GO:0006915: apoptotic process NOC2L nucleolar complex associated 2 homolog (S. cerevisiae) -0.0015109743019154433
GO:0006915: apoptotic process PAEP progestagen-associated endometrial protein 0.0010649492454120443
GO:0006915: apoptotic process PAK1 p21 protein (Cdc42/Rac)-activated kinase 1 -0.002186863181408709

GO:0006915: apoptotic process   PAX3    paired box 3    -0.0029442138311813774
GO:0006915: apoptotic process   PLK3    polo-like kinase 3      0.002563436347137538
GO:0006915: apoptotic process   PML     promyelocytic leukemia  -0.0006799470690519059
GO:0006915: apoptotic process   PPP1R13L        protein phosphatase 1, regulatory subunit 13 like
-0.0014731505524553804
GO:0006915: apoptotic process   PRKCD   protein kinase C, delta -0.0011277401215461832
GO:0006915: apoptotic process   PSMA5   proteasome (prosome, macropain) subunit, alpha type, 5  0.000
3317158097982081
GO:0006915: apoptotic process   PSMD11  proteasome (prosome, macropain) 26S subunit, non-ATPase, 11
-0.0010947868970762168
GO:0006915: apoptotic process   PSMD13  proteasome (prosome, macropain) 26S subunit, non-ATPase, 13
-5.983052028420806e-5
GO:0006915: apoptotic process   PTEN    phosphatase and tensin homolog  1.7370802439597945e-5
GO:0006915: apoptotic process   RAD21   RAD21 homolog (S. pombe)        -0.00010240308952538399
GO:0006915: apoptotic process   RAF1    Raf-1 proto-oncogene, serine/threonine kinase   0.00149822185
51294762
GO:0006915: apoptotic process   S100A14 S100 calcium binding protein A14        -0.00215858573866971
3
GO:0006915: apoptotic process   SIX1    SIX homeobox 1  -0.0018692210724397717
GO:0006915: apoptotic process   STEAP3  STEAP family member 3, metalloreductase 0.0026082397026490167
GO:0006915: apoptotic process   STK24   serine/threonine kinase 24      0.002693624030796802
GO:0006915: apoptotic process   SULF1   sulfatase 1     -0.0008048270968838138
GO:0006915: apoptotic process   TFDP1   transcription factor Dp-1       0.001443707299958998
GO:0006915: apoptotic process   TGFBR1  transforming growth factor, beta receptor 1     0.00033965305
77626321
GO:0006915: apoptotic process   TNFAIP3 tumor necrosis factor, alpha-induced protein 3  0.00111607527
93497309
GO:0006915: apoptotic process   TNFRSF10C       tumor necrosis factor receptor superfamily, member 10
c, decoy without an intracellular domain        0.000317260355768524
GO:0006915: apoptotic process   TNFRSF21        tumor necrosis factor receptor superfamily, member 21
0.0003482531960852244
GO:0006915: apoptotic process   TP53    tumor protein p53       0.0011749295130844956
GO:0006915: apoptotic process   TPX2    TPX2, microtubule-associated    0.0013408903692152196
GO:0006915: apoptotic process   TRAIP   TRAF interacting protein        0.001022035919147697
GO:0031398: positive regulation of protein ubiquitination       AIMP2   aminoacyl tRNA synthetase com
plex-interacting multifunctional protein 2      0.0012010069598741677
GO:0031398: positive regulation of protein ubiquitination       ANGPT1  angiopoietin 1  0.00089136580
74095578
GO:0031398: positive regulation of protein ubiquitination       AXIN1   axin 1  -0.000721306845065099
9
GO:0031398: positive regulation of protein ubiquitination       CAV1    caveolin 1, caveolae protein,
22kDa   -0.0005316868040034787
GO:0031398: positive regulation of protein ubiquitination       CDK5RAP3        CDK5 regulatory subun
it associated protein 3 0.0011275586233072876
GO:0031398: positive regulation of protein ubiquitination       FGFR3   fibroblast growth factor rece
ptor 3  0.00021868572785721461
GO:0031398: positive regulation of protein ubiquitination       RCHY1   ring finger and CHY zinc fing
er domain containing 1, E3 ubiquitin protein ligase     -0.0010856415125516392
GO:0031398: positive regulation of protein ubiquitination       SPHK1   sphingosine kinase 1    0.001
7881180246662415
GO:0031398: positive regulation of protein ubiquitination       WFS1    Wolfram syndrome 1 (wolframi
n)      0.0005759734548774606
GO:0060510: Type II pneumocyte differentiation  AIMP2   aminoacyl tRNA synthetase complex-interacting
multifunctional protein 2       0.001213658882964944
GO:0060510: Type II pneumocyte differentiation  GATA6   GATA binding protein 6  -2.65581446105943e-5
GO:0060510: Type II pneumocyte differentiation  IGF1    insulin-like growth factor 1 (somatomedin C)
0.00013298208500483577
GO:0060510: Type II pneumocyte differentiation  NFIB    nuclear factor I/B      0.0029447623268373043
GO:1901216: positive regulation of neuron death AIMP2   aminoacyl tRNA synthetase complex-interacting
multifunctional protein 2       0.001196310183674719
GO:1901216: positive regulation of neuron death DAXX    death-domain associated protein 0.00087052662
13619483
GO:1901216: positive regulation of neuron death MAP3K5  mitogen-activated protein kinase kinase kinas
e 5     0.00033455092555201754
KEGG:00980: Metabolism of xenobiotics by cytochrome P450        AKR1C1  aldo-keto reductase family 1,
member C1       -0.00148423451174907
KEGG:00980: Metabolism of xenobiotics by cytochrome P450        CYP1B1  cytochrome P450, family 1, su
bfamily B, polypeptide 1        0.00039667067958173464
KEGG:00980: Metabolism of xenobiotics by cytochrome P450        GSTM1   glutathione S-transferase mu
 1      -0.0014679242835137236
KEGG:00980: Metabolism of xenobiotics by cytochrome P450        GSTM2   glutathione S-transferase mu
 2 (muscle)     -0.0019064539301062598
KEGG:00980: Metabolism of xenobiotics by cytochrome P450        GSTM3   glutathione S-transferase mu
 3 (brain)      -0.0012838890629065195
KEGG:00980: Metabolism of xenobiotics by cytochrome P450        GSTP1   glutathione S-transferase pi
 1      0.00022946726349868914
KEGG:00140: Steroid hormone biosynthesis        AKR1C1  aldo-keto reductase family 1, member C1 -0.00
14712069940209108
KEGG:00140: Steroid hormone biosynthesis        CYP1B1  cytochrome P450, family 1, subfamily B, polyp
eptide 1        0.0003925703910131021
KEGG:00140: Steroid hormone biosynthesis        CYP7B1  cytochrome P450, family 7, subfamily B, polyp

eptide 1        0.00016022857801800206
KEGG:00140: Steroid hormone biosynthesis        HSD17B6 hydroxysteroid (17-beta) dehydrogenase 6
-0.0008473314212231178
KEGG:00140: Steroid hormone biosynthesis        SRD5A2  steroid-5-alpha-reductase, alpha polypeptide
 2 (3-oxo-5 alpha-steroid delta 4-dehydrogenase alpha 2)        -0.0006299596753869804
GO:0001523: retinoid metabolic process  AKR1C1  aldo-keto reductase family 1, member C1 -0.0014878829
230930409
GO:0001523: retinoid metabolic process  LRP8    low density lipoprotein receptor-related protein 8, a
polipoprotein e receptor        -0.0009319875632310968
GO:0001523: retinoid metabolic process  STRA6   stimulated by retinoic acid 6   -0.001676578866016785
6
GO:0006805: xenobiotic metabolic process        AKR1C1  aldo-keto reductase family 1, member C1 -0.00
148942202367127
GO:0006805: xenobiotic metabolic process        CYP1B1  cytochrome P450, family 1, subfamily B, polyp
eptide 1        0.0003978334256150409
GO:0006805: xenobiotic metabolic process        CYP27B1 cytochrome P450, family 27, subfamily B, poly
peptide 1       -0.00047093615349892123
GO:0006805: xenobiotic metabolic process        CYP4B1  cytochrome P450, family 4, subfamily B, polyp
eptide 1        -0.0013561574475498969
GO:0006805: xenobiotic metabolic process        CYP7B1  cytochrome P450, family 7, subfamily B, polyp
eptide 1        0.00016330216790402918
GO:0006805: xenobiotic metabolic process        GCLM    glutamate-cysteine ligase, modifier subunit
-0.0037621196683362518
GO:0006805: xenobiotic metabolic process        GSTM1   glutathione S-transferase mu 1  -0.0014741932
94648619
GO:0006805: xenobiotic metabolic process        GSTM2   glutathione S-transferase mu 2 (muscle) -0.00
1914348873701864
GO:0006805: xenobiotic metabolic process        GSTM3   glutathione S-transferase mu 3 (brain)  -0.00
12889978412383453
GO:0006805: xenobiotic metabolic process        GSTP1   glutathione S-transferase pi 1  0.00023144083
87097843
GO:0006805: xenobiotic metabolic process        SLC35D1 solute carrier family 35 (UDP-GlcA/UDP-GalNAc
transporter), member D1 -6.696505371664462e-5
GO:0006805: xenobiotic metabolic process        SULT4A1 sulfotransferase family 4A, member 1    -0.00
11690724189763386
GO:0006805: xenobiotic metabolic process        UGDH    UDP-glucose 6-dehydrogenase     0.00050218317
91150162
GO:0006805: xenobiotic metabolic process        UGP2    UDP-glucose pyrophosphorylase 2 0.00078526807
16861065
GO:0007586: digestion   AKR1C1  aldo-keto reductase family 1, member C1 -0.0014971109160566137
GO:0007586: digestion   NMU     neuromedin U    0.001863059665924592
GO:0007586: digestion   PGC     progastricsin (pepsinogen C)    0.0013304868788324752
GO:0007586: digestion   SLC15A1 solute carrier family 15 (oligopeptide transporter), member 1   -0.00
277732506826375
GO:0007603: phototransduction, visible light    AKR1C1  aldo-keto reductase family 1, member C1 -0.00
14807883341618935
GO:0007603: phototransduction, visible light    LRP8    low density lipoprotein receptor-related prot
ein 8, apolipoprotein e receptor        -0.0009278360939371803
GO:0007603: phototransduction, visible light    METAP1  methionyl aminopeptidase 1      -0.0010119364
799306917
GO:0007603: phototransduction, visible light    PRKCA   protein kinase C, alpha -6.423218744170156e-6
GO:0007603: phototransduction, visible light    STRA6   stimulated by retinoic acid 6   -0.0016651468
689401101
GO:0015721: bile acid and bile salt transport   AKR1C1  aldo-keto reductase family 1, member C1 -0.00
14814815180658953
GO:0015721: bile acid and bile salt transport   RXRA    retinoid X receptor, alpha      0.00110894791
83730648
GO:0030299: intestinal cholesterol absorption   AKR1C1  aldo-keto reductase family 1, member C1 -0.00
14835116889152422
GO:0030855: epithelial cell differentiation     AKR1C1  aldo-keto reductase family 1, member C1 -0.00
14884134398433025
GO:0030855: epithelial cell differentiation     BMP7    bone morphogenetic protein 7    0.00085527282
25342672
GO:0030855: epithelial cell differentiation     CDK1    cyclin-dependent kinase 1       0.00030944164
835295746
GO:0030855: epithelial cell differentiation     CPT1A   carnitine palmitoyltransferase 1A (liver)
0.0012723765196986756
GO:0030855: epithelial cell differentiation     DLX5    distal-less homeobox 5  -0.003280923113797892
4
GO:0030855: epithelial cell differentiation     DLX6    distal-less homeobox 6  -0.000179923080002118
26
GO:0030855: epithelial cell differentiation     FGFR2   fibroblast growth factor receptor 2     0.000
7621441874261383
GO:0030855: epithelial cell differentiation     PCNA    proliferating cell nuclear antigen      0.001
2237219721258714
GO:0030855: epithelial cell differentiation     SIX1    SIX homeobox 1  -0.0018681328167706043
GO:0030855: epithelial cell differentiation     STX2    syntaxin 2      0.00030777425472991254
GO:0030855: epithelial cell differentiation     VEGFA   vascular endothelial growth factor A    0.000
5933402809700335
GO:0030855: epithelial cell differentiation     WT1     Wilms tumor 1   -0.0005061393445866927
GO:0042448: progesterone metabolic process      AKR1C1  aldo-keto reductase family 1, member C1 -0.00

14835116889152422
GO:0042574: retinal metabolic process   AKR1C1  aldo-keto reductase family 1, member C1 -0.0014965655797961742
GO:0042574: retinal metabolic process   ALDH1A2 aldehyde dehydrogenase 1 family, member A2      -0.0030348572373743066
GO:0042574: retinal metabolic process   CYP1B1  cytochrome P450, family 1, subfamily B, polypeptide 1 0.0003988872093612855
GO:0044597: daunorubicin metabolic process      AKR1C1  aldo-keto reductase family 1, member C1 -0.0014835116889152422
GO:0044598: doxorubicin metabolic process       AKR1C1  aldo-keto reductase family 1, member C1 -0.0014835116889152422
GO:0046683: response to organophosphorus        AKR1C1  aldo-keto reductase family 1, member C1 -0.001490446815926812
GO:0046683: response to organophosphorus        RFC3    replication factor C (activator 1) 3, 38kDa 0.0003143934576300519
GO:0046683: response to organophosphorus        TRIM16  tripartite motif containing 16  -0.0014908189289572127
GO:0046683: response to organophosphorus        TYMS    thymidylate synthetase  0.0015651544575019049
GO:0051260: protein homooligomerization AKR1C1  aldo-keto reductase family 1, member C1 -0.0014765187659678439
GO:0051260: protein homooligomerization AXIN1   axin 1  -0.0007229657882528062
GO:0051260: protein homooligomerization BAX     BCL2-associated X protein       -0.00042181360863916516
GO:0051260: protein homooligomerization CAV1    caveolin 1, caveolae protein, 22kDa     -0.0005317576552947028
GO:0051260: protein homooligomerization CEP57   centrosomal protein 57kDa       -0.0005350088650690891
GO:0051260: protein homooligomerization CPT1A   carnitine palmitoyltransferase 1A (liver)       0.0012631724568001816
GO:0051260: protein homooligomerization CRYAA   crystallin, alpha A     -0.000666426903659395
GO:0051260: protein homooligomerization CRYAB   crystallin, alpha B     0.0009705595613955785
GO:0051260: protein homooligomerization ECT2    epithelial cell transforming 2  0.0010495059974709273
GO:0051260: protein homooligomerization FAS     Fas cell surface death receptor -3.2841269541606685e-5
GO:0051260: protein homooligomerization FLOT1   flotillin 1     -0.0007567730086071729
GO:0051260: protein homooligomerization HMOX1   heme oxygenase (decycling) 1    -0.0002145807562788407 4
GO:0051260: protein homooligomerization KCNMA1  potassium large conductance calcium-activated channel, subfamily M, alpha member 1  -0.0006467593622725964
GO:0051260: protein homooligomerization KCNV2   potassium channel, subfamily V, member 2        0.0009177516646332518
GO:0051260: protein homooligomerization RAD51   RAD51 recombinase       -0.001842998967944961
GO:0051260: protein homooligomerization SLC1A1  solute carrier family 1 (neuronal/epithelial high affinity glutamate transporter, system Xag), member 1      -0.000891815451402955
GO:0051260: protein homooligomerization SYT1    synaptotagmin I -0.0014261697188205774
GO:0071395: cellular response to jasmonic acid stimulus AKR1C1  aldo-keto reductase family 1, member C1     -0.0014835116889152422
KEGG:04510: Focal adhesion      AKT1    v-akt murine thymoma viral oncogene homolog 1   0.0007311119635755517
KEGG:04510: Focal adhesion      AKT2    v-akt murine thymoma viral oncogene homolog 2   -0.0006259514437814399
KEGG:04510: Focal adhesion      BCL2    B-cell CLL/lymphoma 2   -5.0870349019443046e-6
KEGG:04510: Focal adhesion      CAV1    caveolin 1, caveolae protein, 22kDa     -0.0005354547470216653
KEGG:04510: Focal adhesion      CCND1   cyclin D1       -0.0026352127440208276
KEGG:04510: Focal adhesion      CDC42   cell division cycle 42  0.001240958670856986
KEGG:04510: Focal adhesion      CHAD    chondroadherin  -0.0021796141370551162
KEGG:04510: Focal adhesion      COL11A1 collagen, type XI, alpha 1      -0.00044194481889284687
KEGG:04510: Focal adhesion      COL1A1  collagen, type I, alpha 1       -0.0005271888644141815
KEGG:04510: Focal adhesion      COL5A1  collagen, type V, alpha 1       -5.055327799347168e-5
KEGG:04510: Focal adhesion      COL5A2  collagen, type V, alpha 2       -0.0003463078706705317
KEGG:04510: Focal adhesion      CTNNB1  catenin (cadherin-associated protein), beta 1, 88kDa    -0.0011676958739873385
KEGG:04510: Focal adhesion      DOCK1   dedicator of cytokinesis 1      -0.0006848151025178811
KEGG:04510: Focal adhesion      EGFR    epidermal growth factor receptor        0.0006866066273333796
KEGG:04510: Focal adhesion      GRB2    growth factor receptor-bound protein 2  0.00042977400146622716
KEGG:04510: Focal adhesion      GSK3B   glycogen synthase kinase 3 beta 0.0015498469809238434
KEGG:04510: Focal adhesion      IGF1    insulin-like growth factor 1 (somatomedin C)    0.00013115624351227092
KEGG:04510: Focal adhesion      IGF1R   insulin-like growth factor 1 receptor   0.0010719019030154217
KEGG:04510: Focal adhesion      ITGA3   integrin, alpha 3 (antigen CD49C, alpha 3 subunit of VLA-3 receptor) 0.0014001321876838254
KEGG:04510: Focal adhesion      ITGA6   integrin, alpha 6       0.0016709391283725886
KEGG:04510: Focal adhesion      ITGB1   integrin, beta 1 (fibronectin receptor, beta polypeptide, antigen CD29 includes MDF2, MSK12) 0.00242820302857357 8
KEGG:04510: Focal adhesion      ITGB4   integrin, beta 4        0.0005810363399926141
KEGG:04510: Focal adhesion      LAMB2   laminin, beta 2 (laminin S)     -0.001309954426513315
KEGG:04510: Focal adhesion      PAK1    p21 protein (Cdc42/Rac)-activated kinase 1      -0.0021888679209354223
KEGG:04510: Focal adhesion      PAK3    p21 protein (Cdc42/Rac)-activated kinase 3      -0.0012590159

858141344
KEGG:04510: Focal adhesion PDGFRB platelet-derived growth factor receptor, beta polypeptide -0.00036201551849594457
KEGG:04510: Focal adhesion PIK3CD phosphatidylinositol-4,5-bisphosphate 3-kinase, catalytic sub unit delta -0.0007812225179908653
KEGG:04510: Focal adhesion PRKCA protein kinase C, alpha -5.841197382942582e-6
KEGG:04510: Focal adhesion PTEN phosphatase and tensin homolog 1.6638750585002516e-5
KEGG:04510: Focal adhesion RAF1 Raf-1 proto-oncogene, serine/threonine kinase 0.00149950669 95844872
KEGG:04510: Focal adhesion RELN reelin 0.001538452145682455
KEGG:04510: Focal adhesion RHOA ras homolog family member A 0.0006700457466109883
KEGG:04510: Focal adhesion THBS1 thrombospondin 1 -0.0010365297244895536
KEGG:04510: Focal adhesion THBS3 thrombospondin 3 0.0016180337455121824
KEGG:04510: Focal adhesion THBS4 thrombospondin 4 -0.0004449362499914049
KEGG:04510: Focal adhesion TNC tenascin C 0.0007346498478172674
KEGG:04510: Focal adhesion VEGFA vascular endothelial growth factor A 0.0005957414552274813
KEGG:04510: Focal adhesion VEGFC vascular endothelial growth factor C -0.003365676955124797
KEGG:05166: HTLV-I infection AKT1 v-akt murine thymoma viral oncogene homolog 1 0.00073014872 65280057
KEGG:05166: HTLV-I infection AKT2 v-akt murine thymoma viral oncogene homolog 2 -0.0006254672 127444562
KEGG:05166: HTLV-I infection APC adenomatous polyposis coli 0.0006433803976048462
KEGG:05166: HTLV-I infection BAX BCL2-associated X protein -0.00042482099359576986
KEGG:05166: HTLV-I infection BUB1B BUB1 mitotic checkpoint serine/threonine kinase B -0.00 12803290688684194
KEGG:05166: HTLV-I infection BUB3 BUB3 mitotic checkpoint protein 0.0016214780334313293
KEGG:05166: HTLV-I infection CCND1 cyclin D1 -0.0026314383248254767
KEGG:05166: HTLV-I infection CDC20 cell division cycle 20 -0.00018846746314222135
KEGG:05166: HTLV-I infection CDC23 cell division cycle 23 -0.0015523879891386674
KEGG:05166: HTLV-I infection CDC27 cell division cycle 27 0.0005361343533335338
KEGG:05166: HTLV-I infection CDKN2A cyclin-dependent kinase inhibitor 2A 0.0017572163952829754
KEGG:05166: HTLV-I infection CDKN2B cyclin-dependent kinase inhibitor 2B (p15, inhibits CDK4) -0.0018343261379406247
KEGG:05166: HTLV-I infection CHEK1 checkpoint kinase 1 0.0008996601738664548
KEGG:05166: HTLV-I infection CREB1 cAMP responsive element binding protein 1 0.00065793162 82892348
KEGG:05166: HTLV-I infection CTNNB1 catenin (cadherin-associated protein), beta 1, 88kDa -0.00 011677765719283276
KEGG:05166: HTLV-I infection E2F1 E2F transcription factor 1 0.0020965077462631043
KEGG:05166: HTLV-I infection E2F3 E2F transcription factor 3 0.0021608927453620726
KEGG:05166: HTLV-I infection EGR1 early growth response 1 0.0010924201527298138
KEGG:05166: HTLV-I infection EGR2 early growth response 2 0.001437952547003632
KEGG:05166: HTLV-I infection ETS2 v-ets avian erythroblastosis virus E26 oncogene homolog 2 -0.0010148671813836451
KEGG:05166: HTLV-I infection FZD3 frizzled class receptor 3 0.00037831878734940306
KEGG:05166: HTLV-I infection FZD7 frizzled class receptor 7 0.001070010419455323
KEGG:05166: HTLV-I infection GSK3B glycogen synthase kinase 3 beta 0.0015481661412175241
KEGG:05166: HTLV-I infection ICAM1 intercellular adhesion molecule 1 0.0007288551674977716
KEGG:05166: HTLV-I infection MAD2L1 MAD2 mitotic arrest deficient-like 1 (yeast) 0.00031861014 213334544
KEGG:05166: HTLV-I infection MSX1 msh homeobox 1 -0.002767385544638794
KEGG:05166: HTLV-I infection MYC v-myc avian myelocytomatosis viral oncogene homolog -0.00 1135538305624405
KEGG:05166: HTLV-I infection NRP1 neuropilin 1 -0.0006427489251350104
KEGG:05166: HTLV-I infection PCNA proliferating cell nuclear antigen 0.001225529361077097
KEGG:05166: HTLV-I infection PDGFRB platelet-derived growth factor receptor, beta polypeptide -0.00036110174672602217
KEGG:05166: HTLV-I infection PIK3CD phosphatidylinositol-4,5-bisphosphate 3-kinase, catalytic sub unit delta -0.0007803532808301464
KEGG:05166: HTLV-I infection POLD1 polymerase (DNA directed), delta 1, catalytic subunit -0.00 04444761587691851
KEGG:05166: HTLV-I infection POLD2 polymerase (DNA directed), delta 2, accessory subunit 0.000 7395023232667374
KEGG:05166: HTLV-I infection POLD3 polymerase (DNA-directed), delta 3, accessory subunit 0.001 7395010973657735
KEGG:05166: HTLV-I infection POLD4 polymerase (DNA-directed), delta 4, accessory subunit -0.00 06380253171875899
KEGG:05166: HTLV-I infection POLE polymerase (DNA directed), epsilon, catalytic subunit 0.002 9465906056635707
KEGG:05166: HTLV-I infection POLE2 polymerase (DNA directed), epsilon 2, accessory subunit 0.001 3251314623131936
KEGG:05166: HTLV-I infection POLE3 polymerase (DNA directed), epsilon 3, accessory subunit 0.000 8044456814438908
KEGG:05166: HTLV-I infection PRKX protein kinase, X-linked 0.0003947014380226406
KEGG:05166: HTLV-I infection PTTG1 pituitary tumor-transforming 1 -0.000361399323128094
KEGG:05166: HTLV-I infection PTTG2 pituitary tumor-transforming 2 -0.0017237148085190964
KEGG:05166: HTLV-I infection RB1 retinoblastoma 1 -0.001493091369589595
KEGG:05166: HTLV-I infection STAT5A signal transducer and activator of transcription 5A 0.001 5953326831280432
KEGG:05166: HTLV-I infection TBPL1 TBP-like 1 -0.0007881631366525119
KEGG:05166: HTLV-I infection TGFB1 transforming growth factor, beta 1 -7.30389369201852e-5

```
KEGG:05166: HTLV-I infection    TGFB2   transforming growth factor, beta 2      -0.001058701643025118
8
KEGG:05166: HTLV-I infection    TGFB3   transforming growth factor, beta 3      -0.001822789925395239
4
KEGG:05166: HTLV-I infection    TGFBR1  transforming growth factor, beta receptor 1     0.00033971200
41990669
KEGG:05166: HTLV-I infection    TP53    tumor protein p53       0.0011750040561670679
KEGG:05166: HTLV-I infection    WNT1    wingless-type MMTV integration site family, member 1    0.000
786460923647678
KEGG:05166: HTLV-I infection    WNT10B  wingless-type MMTV integration site family, member 10B  -0.00
13876971599892582
KEGG:05166: HTLV-I infection    WNT4    wingless-type MMTV integration site family, member 4    -0.00
025196024583135567

Excessive output truncated after 524341 bytes.
```

KEGG:05166: HTLV-I infection    WNT5A   wingless-type MMTV integration site family, member 5A   -0.00
06671137872325665
KEGG:05166: HTLV-I infection    WNT7A   wingless-type MMTV integration site family, member 7A   1.957
9561150038017e-5
KEGG:05166: HTLV-I infection    XBP1    X-box binding protein 1 0.00026680014780548696
KEGG:05169: Epstein-Barr virus infection        AKT1    v-akt murine thymoma viral oncogene homolog 1
0.0007262566190756878
KEGG:05169: Epstein-Barr virus infection        AKT2    v-akt murine thymoma viral oncogene homolog 2
-0.0006235672077896338
KEGG:05169: Epstein-Barr virus infection        BCL2    B-cell CLL/lymphoma 2   -4.882937985058531e-6
KEGG:05169: Epstein-Barr virus infection        CCNA1   cyclin A1       -0.0017501487999748546
KEGG:05169: Epstein-Barr virus infection        CDK1    cyclin-dependent kinase 1       0.00030695291
3730044
KEGG:05169: Epstein-Barr virus infection        CSNK2B  casein kinase 2, beta polypeptide       0.001
5192769043707265
KEGG:05169: Epstein-Barr virus infection        GSK3B   glycogen synthase kinase 3 beta 0.00154119024
95898236
KEGG:05169: Epstein-Barr virus infection        HDAC2   histone deacetylase 2   -0.001203898890181675
KEGG:05169: Epstein-Barr virus infection        HSPA2   heat shock 70kDa protein 2      -0.0001034582
4148931563
KEGG:05169: Epstein-Barr virus infection        ICAM1   intercellular adhesion molecule 1       0.000
727362676052208
KEGG:05169: Epstein-Barr virus infection        IFNG    interferon, gamma       -4.900796379268886e-5
KEGG:05169: Epstein-Barr virus infection        IRAK1   interleukin-1 receptor-associated kinase 1
-0.0016002937427628528
KEGG:05169: Epstein-Barr virus infection        LYN     LYN proto-oncogene, Src family tyrosine kinas
e       -0.0014136847532094164
KEGG:05169: Epstein-Barr virus infection        MYC     v-myc avian myelocytomatosis viral oncogene h
omolog  -0.0011325117414121163
KEGG:05169: Epstein-Barr virus infection        NEDD4   neural precursor cell expressed, developmenta
lly down-regulated 4, E3 ubiquitin protein ligase       0.002253522858241044
KEGG:05169: Epstein-Barr virus infection        PIK3CD  phosphatidylinositol-4,5-bisphosphate 3-kinas
e, catalytic subunit delta      -0.0007767642901220772
KEGG:05169: Epstein-Barr virus infection        POLR2D  polymerase (RNA) II (DNA directed) polypeptid
e D     0.0018450093234024657
KEGG:05169: Epstein-Barr virus infection        PRKX    protein kinase, X-linked        0.00039030241
681122673
KEGG:05169: Epstein-Barr virus infection        PSMD11  proteasome (prosome, macropain) 26S subunit,
non-ATPase, 11  -0.001091548683016147
KEGG:05169: Epstein-Barr virus infection        PSMD13  proteasome (prosome, macropain) 26S subunit,
non-ATPase, 13  -6.068925883060731e-5
KEGG:05169: Epstein-Barr virus infection        RB1     retinoblastoma 1        -0.001486167860636004
KEGG:05169: Epstein-Barr virus infection        RBPJ    recombination signal binding protein for immu
noglobulin kappa J region       0.000977774103801858
KEGG:05169: Epstein-Barr virus infection        SKP2    S-phase kinase-associated protein 2, E3 ubiqu
itin protein ligase     -0.000537455985916273
KEGG:05169: Epstein-Barr virus infection        TBPL1   TBP-like 1      -0.0007871806823993504
KEGG:05169: Epstein-Barr virus infection        TNFAIP3 tumor necrosis factor, alpha-induced protein
3       0.0011106622138456054
KEGG:05169: Epstein-Barr virus infection        TP53    tumor protein p53       0.001167256464273668
KEGG:04630: Jak-STAT signaling pathway  AKT1    v-akt murine thymoma viral oncogene homolog 1   0.000
7322366576941181
KEGG:04630: Jak-STAT signaling pathway  AKT2    v-akt murine thymoma viral oncogene homolog 2   -0.00
06269139625152389
KEGG:04630: Jak-STAT signaling pathway  CCND1   cyclin D1       -0.002638355521223424
KEGG:04630: Jak-STAT signaling pathway  CSF3R   colony stimulating factor 3 receptor (granulocyte)
-8.433393498168513e-5
KEGG:04630: Jak-STAT signaling pathway  CSH1    chorionic somatomammotropin hormone 1 (placental lact
ogen)   -0.0002576642301649812
KEGG:04630: Jak-STAT signaling pathway  GRB2    growth factor receptor-bound protein 2  0.00043051485
57108661
KEGG:04630: Jak-STAT signaling pathway  IFNA2   interferon, alpha 2     -0.001720392587112037
KEGG:04630: Jak-STAT signaling pathway  IFNG    interferon, gamma       -4.899191761680743e-5
KEGG:04630: Jak-STAT signaling pathway  IL12B   interleukin 12B 0.001278848198633779
KEGG:04630: Jak-STAT signaling pathway  IL20RA  interleukin 20 receptor, alpha  0.0001745061911577260
4
KEGG:04630: Jak-STAT signaling pathway  IL4     interleukin 4   0.00025976679277823583
KEGG:04630: Jak-STAT signaling pathway  IL6ST   interleukin 6 signal transducer 0.0018793482781852787
KEGG:04630: Jak-STAT signaling pathway  IL7     interleukin 7   0.0008729639337432427
KEGG:04630: Jak-STAT signaling pathway  JAK2    Janus kinase 2  -3.159219747417667e-5
KEGG:04630: Jak-STAT signaling pathway  LEP     leptin  0.0031976777282567794
KEGG:04630: Jak-STAT signaling pathway  MYC     v-myc avian myelocytomatosis viral oncogene homolog
-0.001138868670510603
KEGG:04630: Jak-STAT signaling pathway  PIK3CD  phosphatidylinositol-4,5-bisphosphate 3-kinase, catal
ytic subunit delta      -0.0007822987779466497
KEGG:04630: Jak-STAT signaling pathway  PRLR    prolactin receptor      0.002187381210529882
KEGG:04630: Jak-STAT signaling pathway  STAT5A  signal transducer and activator of transcription 5A
0.0015991927405241142
KEGG:04071: Sphingolipid signaling pathway      AKT1    v-akt murine thymoma viral oncogene homolog 1
0.0007202122493181328
KEGG:04071: Sphingolipid signaling pathway      AKT2    v-akt murine thymoma viral oncogene homolog 2

```
-0.000619913285639517
KEGG:04071: Sphingolipid signaling pathway      BAX     BCL2-associated X protein       -0.0004216900
039775023
KEGG:04071: Sphingolipid signaling pathway      BCL2    B-cell CLL/lymphoma 2   -4.414007247247495e-6
```

The following prints the coefficients in the original dimensions:

In [52]:

```
for (v, β) in zip(variable_names[β_orig .!= 0], β_orig[β_orig .!= 0])
    println("$v\t$β")
end
```

| | |
|---|---|
| RFC2 | 0.017828370354349216 |
| PAX8 | 0.03684757874519627 |
| THRA | 0.02806710700734744 |
| EPHB3 | 0.019420774902517743 |
| CFL1 | -0.030460550648403955 |
| YY1 | 0.013402782876598905 |
| ZPR1 | -0.004170459413053395 |
| RHOA | 0.04322099220670631 |
| GUK1 | 0.05284161532839056 |
| DSP | -0.0008924333259685153 |
| RAD21 | -0.001631468584420956 8 |
| SF3B2 | 0.004117778492081893 |
| NDRG1 | -0.006543628992920165 5 |
| CD63 | -0.01588590376579995 |
| DNAJB1 | -0.002074023570791529 |
| XBP1 | 0.021161233616516405 |
| SPTBN1 | 0.03684346496032824 |
| HMGB1 | -0.030756596050312526 |
| GPX1 | 0.021234007378015077 |
| NUMA1 | 0.021341452238237472 |
| ARL6IP5 | 0.001092416179919603 9 |
| STMN1 | 0.007247922509085328 |
| ODC1 | 0.009846896972186728 |
| DAZAP2 | -0.0007855012627521644 |
| TMBIM6 | -0.00228187693048959 |
| GSTP1 | 0.008155197247241433 |
| ZNF207 | -0.00272055661792677 |
| DHCR24 | -0.035829513244511876 |
| DEK | 0.017525131494012178 |
| HIF1A | -0.04980124402515670 5 |
| SNX17 | 0.011478189129344098 |
| CD9 | -0.03889280343675502 |
| JUP | -0.038198768007394066 |
| TGM2 | -0.00172711866113033 |
| MMP2 | -0.02598193263059217 |
| THBS1 | -0.08446415723750327 |
| TUFM | -0.006526015058534712 |
| POLD2 | 0.017044076906068108 |
| CPE | -0.022985479474374404 |
| SLC7A5 | -0.04388439401899305 |
| AMD1 | -0.025402082120591248 |
| PCNA | 0.0417767591397736 |
| PSMD13 | -0.0024832832828435847 |
| RAF1 | 0.1433808502445361 |
| PSMA5 | 0.013051348177155975 |
| OGDH | -0.005067398489416659 |
| TOP2A | -0.0015644814822285936 |
| ETS2 | -0.012155698340348456 |
| CST3 | -0.002004327087036865 |
| ZFP36L2 | -0.004307263131398862 |
| CSNK2B | 0.03354336593236934 |
| PHGDH | 0.005731129900934959 |
| HSD17B4 | -0.021954771791886242 |
| SOX4 | -0.001843562616912388 |
| WDR77 | 0.0010212673455483972 |
| BUB3 | 0.020980692290832027 |
| ITGA3 | 0.053139657793133725 |
| RRM1 | 0.03261837436083391 |
| IGFBP4 | -0.01529495987980445 |
| APOD | 0.05281390344932129 |
| RPA1 | 0.016706070001905687 |
| CTNNB1 | -0.016692352559666623 |
| ZNHIT1 | -0.0016925861371024217 |
| MCM3 | 0.02142799502691732 |
| SORD | -0.03855195661545188 |
| ID2 | 0.003043658587362374 3 |
| IRAK1 | -0.08686643583075766 |
| SMC1A | -0.021331481725464577 |
| KRT18 | -0.010141434375582892 |
| PHB2 | -0.012382612300996647 |
| TNC | 0.010270550679614052 |
| ITGA6 | 0.06202647497027872 |
| SMC4 | 0.008291635589654026 |
| GJA1 | -0.009687495934931683 |
| EGR1 | 0.048905226314721474 |
| TPR | -0.028977825234667656 |
| TP53 | 0.1700158671284832 |
| MCM5 | 0.013891019627766395 |
| RPA2 | 0.008798613227269438 |
| DAXX | 0.022950864698423102 |
| AZIN1 | -0.0008413038683729087 |

| | |
|---|---|
| NCAPD2 | -0.006694478531962174 |
| ADAR | -0.004273578061227504 |
| ENG | 0.03035317517838372 |
| HDAC2 | -0.07936953927783136 |
| BNIP3 | 0.1025852155435736 |
| RRM2 | -0.00904653751058178 |
| IMPDH2 | -0.004533325675427472 |
| PSRC1 | 0.005150878919895948 |
| PRDX4 | 0.015233898271638825 |
| MCM6 | 0.013085227098779618 |
| CHMP1A | -0.011579771180144553 |
| CIB1 | 0.002494086200296393 |
| CCNC | 0.005668826446434938 |
| NASP | 0.016794814852552593 |
| EGFR | 0.05923885482145959 |
| LANCL1 | 0.00073398264946708 |
| EFNA1 | -0.012411014380812602 |
| SFRP1 | 0.1053762993216416l |
| SLC39A6 | 0.0011622219237774857 |
| BIRC5 | -0.006728392679664588 |
| NUP153 | 0.020401330786399764 |
| BRD4 | 0.009878615819780397 |
| IGBP1 | 0.03716659575857557 |
| MCM2 | 0.02964701901069167 |
| NOC2L | -0.0013018167154264428 |
| WWTR1 | 0.024221399551931373 |
| AIMP2 | 0.008424538648259098 |
| PKN1 | -0.0380894348100307 |
| PRKCZ | -0.08299626615623114 |
| MVP | -0.005039939366418271 |
| SLC6A8 | -0.010418167114822336 |
| PLK1 | 0.04441888968698376 |
| E2F4 | -0.04000230602549144 |
| VPS72 | 0.0027918177788449927 |
| PDGFRB | -0.030899955754132985 |
| TACSTD2 | -0.027265926405762794 |
| MTHFD1 | -0.01510667125644826 |
| COL1A1 | -0.02774187855127702 |
| UNG | 0.005040246685305159 |
| UBE2K | -0.002633814755276219 |
| HTT | -0.04333086573251333 |
| BASP1 | 0.004749137833453016 |
| IGSF3 | 0.0002846542519311496 |
| RXRA | 0.05780812177993816 |
| MYC | -0.07254841230400208 |
| CYP1B1 | 0.016791485175176302 |
| CTSK | -0.00537288792758819 |
| MLXIP | 0.010971441163870615 |
| MLH1 | 0.04920692668941197 |
| DHFR | -0.007368555092904128 |
| PRKCD | -0.06848826247053859 |
| VAMP8 | -0.0006123414941763355 |
| GSTM3 | -0.016771952909600263 |
| FOXM1 | 0.005470530397087533 |
| NFX1 | -0.0003601096625964127 |
| TYMS | 0.05804193843777028 |
| PDK2 | 0.022218697269260047 |
| SSBP1 | 0.003358191924518330З |
| ENSA | 0.011071869332661398 |
| IRF6 | -0.018578500771458274 |
| NRIP1 | 0.010689665804530204 |
| CTPS1 | -0.00503376916407351 |
| LYN | -0.13602000880799606 |
| SERPINE1 | 0.004549817149767904 |
| TOPBP1 | 0.008262583182653471 |
| ICAM1 | 0.04698813280501898 |
| ARL3 | -0.012944671028492987 |
| TNFAIP3 | 0.06196838444377323 |
| RASA1 | -0.010347310860967325 |
| PTPN1 | -0.03400858029035727 |
| IGFBP2 | 0.0021582946457342545 |
| FOXO1 | 0.08981646049044094 |
| LIG1 | 0.0193507999466856 |
| USP8 | -0.004295447026511489 |
| SLC7A8 | -0.007029917025023939 |
| HMGCL | -0.017920596453702684 |
| UBE2S | 0.00451754408104329 |
| SLC1A3 | -0.07846231096017797 |
| DBN1 | -0.007958079888046346 |
| WBP1L | -0.0006812879912496237 |
| STAMBP | -0.0013439330003021679 |

| | |
|---|---|
| HEXIM1 | 0.0009146807422728784 |
| AGT | -0.11777825302279481 |
| HPRT1 | -0.03429130520873595 |
| CDC20 | -0.0041580317140993426 |
| CDC23 | -0.029289849014534904 |
| E2F1 | 0.09452257827698175 |
| WFS1 | 0.016589595131934937 |
| MSH6 | 0.018275675693561977 |
| ADM | 0.09880822122504294 |
| SOX9 | -0.05534375159597975 |
| UBE2C | -0.00524690374511718 |
| MUT | 0.035950521811880666 |
| POLD4 | -0.015987318033880753 |
| STMN2 | -0.015563832488220849 |
| STAT5A | 0.09242498840939789 |
| RNASEH2A | 0.004135880019279893 |
| C2 | -0.003309436311102557 |
| CAV1 | -0.06306570615282246 |
| CHST15 | -0.005782691230612209 |
| CTNNBIP1 | 0.008138176628460057 |
| TGFB1 | -0.01286980654040744 |
| KIF2A | -0.009405835015435746 |
| CCDC86 | 0.0007146750595481012 |
| SLC11A2 | -0.025404068219129074 |
| RB1 | -0.08508075060728257 |
| KIAA0040 | -0.0022389706808229122 |
| SPAG5 | -0.0032009057004588662 |
| PVRL2 | -0.0006298298280750446 |
| RNF8 | 0.014322281504453256 |
| KATNB1 | -0.0011054373409471166 |
| FBN2 | -0.007380182720434389 |
| DOCK1 | -0.011621186923791521 |
| RFC5 | 0.04044914529319224 |
| MTMR2 | -0.008937113364945702 |
| CDK1 | 0.011067291839721308 |
| LGALS9 | -0.0045394352169789995 |
| ZNF24 | 0.0038769440123150203 |
| CDH3 | -0.028177404381542398 |
| DTYMK | 0.05614040994978363 |
| HS2ST1 | -0.006412566664718714 |
| NCK2 | -0.006297849220518809 |
| ZNF148 | 0.016871012238787716 |
| COL5A1 | -0.0011032697296370437 |
| UGDH | 0.006978188918291193 |
| ORC4 | -0.0003733583483527795 |
| EZH2 | -0.004516562690241726 |
| MAD2L1 | 0.005619561855304875 |
| CDC40 | 0.0378164276380198 |
| RPS6KA1 | -0.09063268000400478 |
| HES1 | -0.064127697024044 |
| PSMG1 | 0.00040652018292425306 |
| MMD | 0.010754803225344934 |
| KMT2B | 0.00504495307813712 |
| POLD1 | -0.014189772998580036 |
| ASF1A | 0.009452157844126936 |
| SUCO | 0.01123697897827811 |
| STC2 | -0.016718403399941687 |
| CDH2 | -0.01726046288243851 |
| SPR | -0.01968662444539362 |
| PLEK | 0.02437012161243763 |
| CEP57 | -0.004804746283049294 |
| MED1 | 0.06757529887822789 |
| GCDH | 7.034724478452115e-6 |
| SPG11 | -8.779000084344127e-5 |
| APC | 0.040377343009541176 |
| PTTG1 | -0.00578518542883349 |
| CUL7 | -0.001874500267223687 |
| AOC1 | -0.018470431644445617 |
| GGH | -0.0024980678538852963 |
| ADIRF | -0.001136796358779777 |
| CSF3R | -0.001001198050010839 |
| ALDH5A1 | 0.04921593880047606 |
| SKP2 | -0.011781180780861279 |
| IGF1R | 0.056589839552778735 |
| CPT1A | 0.02793053700663758 |
| FGFR2 | 0.09289228548238233 |
| PCNT | 0.02860162116186179 |
| POLR2D | 0.044608743610788586 |
| HMOX1 | -0.010639419074505253 |
| CXCL12 | -0.057742079037596795 |
| IVD | -0.007438199163742574 |

| | |
|---|---|
| BCL2 | -0.0008945318307243404 |
| CX3CL1 | 0.08062568872404574 |
| PI3 | -0.002078090528169930 4 |
| E2F3 | 0.04115371621305719 |
| RREB1 | -0.02182065647643502 |
| FZD7 | 0.0287529926189359 |
| ITPR1 | -0.041649108642733426 |
| KIAA0020 | 0.0009336617440814806 |
| BAK1 | -0.1122410424196227 7 |
| HMGB3 | 0.003201531061351816 |
| BUB1B | -0.020432339950024483 |
| DYNC2LI1 | 0.004614918362283697 |
| DLGAP5 | -0.007949800760510883 |
| MANBA | -0.01902581522427586 |
| SEMA3C | -0.0030735790631009406 |
| AKT2 | -0.0658944023468839 |
| IGF2BP3 | -0.0022821883184598103 |
| MAP3K5 | 0.009733993057483062 |
| IGFBP6 | -0.010108693926009854 |
| PIK3CD | -0.08446872456456452 |
| CXADR | 0.008125836938706728 |
| NR1H3 | 0.029204949713111036 |
| GCLM | -0.07189730152604519 |
| MAPT | 0.03556043947045428 |
| VASH1 | 0.005336994188221942 |
| KIF3B | -0.002474744514527213 4 |
| E2F6 | 0.0021573287632984505 |
| CDC6 | 0.021677609174670764 |
| SLC31A1 | -0.0002675017652726704 |
| CHAF1A | 0.0021731702471032423 |
| SYT1 | -0.0231985911200916 |
| LARS2 | 0.002685547772540581 |
| PURA | -0.0015289193774571034 |
| WWP2 | -0.018109591414804605 |
| RFC4 | 0.013094847932800606 |
| ZWINT | 0.005037357692006729 |
| CELSR2 | -0.015828809359290336 |
| PTEN | 0.002703874590459572 |
| PRKX | 0.02631090918211519 |
| TPST2 | 0.0021248479871390376 |
| AURKA | 0.026072099828434364 |
| RBMX2 | 0.0011525304047793952 |
| SMARCD3 | 0.005056510060143438 |
| LIG3 | 0.017937525776897875 |
| CDC45 | 0.03512874376469682 |
| RFC3 | 0.006289485493524521 |
| FOXO3 | 0.04304335440765909 |
| TFDP1 | 0.024597855220075616 |
| AKR1C1 | -0.028235242102568604 |
| NDC80 | -0.01299434899650587 |
| IMPDH1 | -0.022526243065154807 |
| CKS2 | -0.0002767823749429333 |
| CPOX | 0.015309998121761634 |
| RARG | -0.07205944464497151 |
| BACH1 | 0.0038557769403125748 |
| ACOT8 | 0.014671699445802378 |
| SMC2 | 0.0034757928847099304 |
| VDR | 0.015094705098909525 |
| MMP7 | 0.006718351487298258 |
| SKI | -0.025321698970961454 |
| RGS14 | -0.03319822659144861 |
| LOX | -0.005739266799934818 |
| PROM1 | -0.007435736621323742 |
| CREB1 | 0.05291850511267864 |
| COL11A1 | -0.009683886337841513 |
| TRIM16 | -0.013286678121464168 |
| BCAS1 | -0.00012129172027097935 |
| FGFR3 | 0.015061013692887142 |
| GOLGA2 | 0.002426158179985120 4 |
| EFS | 0.007825864377930104 |
| GSTM2 | -0.03453019093217699 |
| SLC2A5 | 0.002611578059423235 |
| POLA2 | -0.020954323183441054 |
| KIF11 | 0.012877878211679267 |
| GAP43 | -0.0010497450042150276 |
| CDC7 | -0.009474360240546228 |
| PHF14 | 0.012996775839253327 |
| CXCL10 | 0.0022540377037781707 |
| EEF1A2 | -0.011206457515607325 |
| KIAA0513 | -0.0026336085747406777 |
| STAR | 0.04875919120485766 6 |

| | |
|---|---|
| GSTM1 | -0.01622487243355555 |
| FKBP5 | 0.0009347797125010532 |
| ABCG1 | -0.06107660423264803 |
| CROT | -0.004733522036937515 |
| CD22 | 0.0033428343036181925 |
| SLC7A7 | -0.00835070353512725 |
| DKK1 | -0.00437664025968826 |
| EXO1 | -0.007169600632849006 |
| NEK2 | 0.0008735531837337527 |
| TFAP2A | 0.01474155768604491 |
| FOXA1 | 0.0007464072405028385 |
| KIF23 | -0.003814023656309869 |
| NCK1 | -0.023238340655216336 |
| CNKSR1 | -0.003309506835284202 |
| BICD1 | 0.029467848470741357 |
| PICK1 | -0.018798454080332136 |
| DSC2 | 0.0036990888717844854 |
| NUDT1 | 0.01658488722027079 |
| FEN1 | 0.03928002763295099 |
| CHAF1B | 0.06336904561093554 |
| THBS4 | -0.00793658615361211 |
| FAS | -0.0018303251038774492 |
| ZW10 | -0.009789089406096785 |
| ESPL1 | 0.0026349483067022576 |
| TTK | 0.008657784410876937 |
| MELK | 0.020732613609057232 |
| CCNF | 0.0017514255657294672 |
| RAD9A | 0.00781452458959319 |
| CDK8 | 0.026158579700205575 |
| POLA1 | 0.028022750740118882 |
| CP | 0.006502042373259052 |
| ORC2 | -0.0036782166622163805 |
| SERPINB5 | -0.03646908129424964 |
| IL6ST | 0.05064993589383436 |
| GCHFR | -0.00664382491807731 |
| PCSK2 | 0.004188397886511958 |
| MTERF1 | 0.0021837543354899405 |
| MGMT | 0.006087508732606019 |
| UGCG | 0.0032665560698968772 |
| PLK4 | 0.027056234063494847 |
| SOX11 | -0.010375705499242811 |
| INHBA | -0.07031194038416436 |
| HPN | 0.06343944613246817 |
| PTPN2 | -0.01185049915707216 |
| CARD8 | 0.0009737029049671201 |
| MTAP | 0.026794832993818404 |
| ORC5 | -0.0008066718761636689 |
| PLK3 | 0.056517760847479115 |
| SHROOM2 | 0.01657094105494471 |
| CLOCK | 0.005225377238792192 |
| SLC22A18 | 0.0036381804080160784 |
| ITGB4 | 0.013350845909100328 |
| NF2 | -0.03664467039870869 |
| PFN2 | 0.030182653760940353 |
| ATF5 | -0.0438895183388974 |
| RAD51 | -0.05959231669664289 |
| CCNE2 | 0.018853635423271606 |
| ORM1 | -0.0027846959939131564 |
| CENPE | 0.03231574820149773 |
| PSPH | -0.021255067676917733 |
| KIT | 0.023303054440520162 |
| AUH | -0.008742765931883346 |
| PRIM1 | 0.007640062297202151 |
| SLC22A5 | -0.02409020140972213 |
| SERPINF2 | 0.014325953337234935 |
| CRIP1 | 0.01716096384814977 |
| ORC1 | -0.0058874352807807015 |
| MOK | 0.005371912559628551 |
| PEX11A | 0.006381301901495141 |
| GLI3 | -0.14256324992470193 |
| KAL1 | -0.0011795132247510995 |
| GALK2 | 0.00012695444364892652 |
| ESR1 | -0.040574930847268 |
| AREG | 0.05169677369466026 |
| GPSM2 | 0.00711559391187069 |
| CXCL13 | 0.07837155826906521 |
| EGR2 | 0.03901184423000848 |
| PER2 | 0.03325074136646313 |
| PGC | 0.005304266152110366 |
| LRP8 | -0.020463024120600203 |
| TFAP2C | 0.020189519687844052 |

| | |
|---|---|
| CDC14A | 0.0023230844116615816 |
| IGFBP1 | 0.003091742143176988 |
| SLC15A2 | -0.0010016510545315749 |
| S100A1 | 0.007943911280445217 |
| DCT | -0.030649335406504687 |
| BARD1 | 0.020946437515230745 |
| AGTR1 | 0.012107886892131919 |
| INPP4B | 0.004323559735075351 |
| PDZK1 | -0.001581735302411606 |
| CHEK1 | 0.01974575498226775 |
| MLLT10 | -0.00032716360192899417 |
| DAG1 | 0.009834514052789567 |
| BCHE | 4.276323291334793e-5 |
| OCEL1 | -7.907603710843561e-5 |
| HOXB2 | 0.03343632574992888 |
| C6orf106 | 0.0014717327485321858 |
| DACH1 | 0.027188465761183736 |
| UGP2 | 0.010926257825547092 |
| GCNT1 | 0.008369281276925853 |
| GATA4 | -0.06217314871779478 |
| SLC44A4 | 0.025737067759456722 |
| TRAIP | 0.007132092222931834 |
| LRP6 | 0.005758161221066203 |
| ANGPT1 | 0.04651626867644292 |
| PRIM2 | -0.010671924399342714 |
| PAX6 | 0.10863461454752399 |
| GAL3ST1 | 0.013072702833115055 |
| CYP27B1 | -0.012502240478196407 |
| HSD17B6 | -0.005185492295727334 |
| BLM | 0.012447397255563744 |
| KLK7 | 0.008128490790703461 |
| FGF7 | 0.027689500012265018 |
| TCP11L1 | 0.002348702682587311 |
| ROM1 | 0.012298719107757091 |
| SIX1 | -0.09910802932249518 |
| CPA4 | -0.0013634362606130984 |
| JAK2 | -0.003768246769903389 |
| TULP3 | 0.01810592878314025 |
| GREB1 | -0.0006855056510944366 |
| RET | -0.01794275462279358 |
| MSH3 | -0.005832871205581579 |
| SLC22A4 | -0.005223516220411508 |
| CX3CR1 | -0.01877762630860521 |
| CCNA1 | -0.029898620261494976 |
| POLE2 | 0.022591169092778675 |
| S100A7 | 0.019112269245436565 |
| RELN | 0.06930054090657724 |
| IL27RA | -0.0038227634199274733 |
| MSX1 | -0.10305141919523 |
| BATF | 0.02850508666822337 |
| WNT5A | -0.07847281242458237 |
| GPR64 | 0.0018646637234725897 |
| NMU | 0.022327964446469235 |
| DICER1 | -0.0011654068895853803 |
| WT1 | -0.024530961590750838 |
| ACADL | 0.017504038742022124 |
| HMGA1 | -0.005375502601779681 |
| LRRC23 | 0.0008850087968634609 |
| CTH | 0.03683707201625856 |
| NELL1 | 0.01071026244982359 |
| GINS1 | 0.005782639831933588 |
| ISL1 | 0.004283152117975489 |
| PLA2G5 | 0.06866360274356276 |
| WNT10B | -0.05960610425415589 |
| EDA | -0.012953896218986422 |
| TNFRSF10C | 0.002220244284736504 |
| PAX2 | -0.10655206932697343 |
| LHX1 | -0.04795659968052898 |
| GUCY2C | -0.011412771717607458 |
| PRLR | 0.04369735823572912 |
| PDK3 | 0.009255883892137945 |
| KIF14 | 0.015300681464566706 |
| CPLX2 | 0.016297126542175237 |
| EYA3 | 0.00829170731718843 |
| ANK3 | 0.016523807369716512 |
| SLC1A1 | -0.012628381153536344 |
| ZNF205 | 0.0029722372034130277 |
| TBCCD1 | 0.002407925925447312 |
| NDRG2 | 0.019672720370709128 |
| KIAA0125 | 0.0004132393741945021 |
| HINFP | 0.014878738381224681 |

| | |
|---|---|
| PML | -0.04791240195835318 |
| MGAM | -0.011773852786785865 |
| ABAT | 0.01909954045691702 |
| KLHL24 | 0.00032070457449530636 |
| SMA4 | 0.0020387403396643564 |
| CDKL5 | 0.018288948441041876 |
| GFI1 | 0.031571796504911286 |
| DRD2 | -0.021851368214878313 |
| SIX3 | 0.06346645664152697 |
| CHRNB2 | -0.03847303987011225 |
| GLI1 | -0.03497752184708721 |
| FLT3 | -0.02025991321711388 |
| PDK1 | 0.02563803576820573 |
| ASIC2 | 0.052607582207338494 |
| IL7 | 0.015592135087047689 |
| CHRNB1 | -0.00019164268514129743 |
| KCNJ2 | -0.0175672057590206 |
| ACRV1 | -0.000602632480003769 |
| ERBB4 | -0.008486885619535424 |
| SCGB1D2 | 0.0004387758538985073 |
| STATH | -0.005395035560048601 |
| ALX1 | 0.02986215295181512 |
| AFM | -0.001693919942297489 |
| PAEP | 0.006354664338546461 |
| CHAD | -0.013078713959233526 |
| AMHR2 | -0.002368030217064603 |
| SALL1 | -0.0884056709242187 |
| TAT | 0.01686296407596211 |
| PRKCA | -0.0006614508433365902 |
| SRD5A2 | -0.015277224904423728 |
| TGFBR1 | 0.0258946932399959 |
| HTR6 | -0.009662179726478122 |
| C8B | 0.0025303041239801496 |
| PHOX2B | 0.011495585806956116 |
| PTK7 | -0.0035529964314248747 |
| ALDH1A2 | -0.12135910958184191 |
| LDHC | 0.016357644900862903 |
| CRISP1 | 0.00026445494265186885 |
| GLI2 | 0.10206865475980202 |
| TNFRSF11A | 0.07533420474772554 |
| SLC16A6 | 0.0020836993612578544 |
| CDKN2A | 0.1091711254000116 |
| THRB | 0.029168583684075385 |
| ASS1 | 0.004830149327111398 |
| LEP | 0.1998378331370313 |
| CITED1 | 0.11631010981891542 |
| AKT1 | 0.1325823850762814 |
| ZBTB18 | 0.0139589612891684 |
| AGRP | -0.031067674487839293 |
| ALOX12 | -0.07043025237679758 |
| SLC15A1 | -0.022132859662382626 |
| SLC17A3 | -0.013743047633786651 |
| TCF15 | -0.042577823785949695 |
| HTR2C | -0.011836730934854088 |
| TFRC | 0.022648351025412846 |
| STX2 | 0.0050443912429787244 |
| CYP7B1 | 0.00192962417579458 |
| HOXD13 | -0.008245711200176873 |
| PROX1 | 0.06596269150182502 |
| RAD17 | -0.0012069700278087014 |
| MSMB | -0.0013331384342858386 |
| NR2E1 | -0.09852641803918796 |
| CAST | -0.030527010441833227 |
| ATP5G3 | 0.0018423725159665723 |
| CDKN2B | -0.05876492166517276 |
| IL4 | 0.01882278098376838 |
| PITX2 | 0.10480238529079344 |
| MAGEB3 | -0.00691658100488872 |
| SHH | 0.08366559408643309 |
| MMP20 | -0.017020842905839707 |
| SLC4A7 | 0.010467453451719635 |
| FOXH1 | -0.02464150989437682 |
| ADAM2 | 0.00645805920949776 |
| PAX3 | -0.059068253533199576 |
| KIF25 | -0.009609396255554062 |
| SERPINH1 | -0.0075636364797095415 |
| PPP2R3A | -0.0006220842487928137 |
| RASSF8 | 0.0033201926705128 |
| RBPJ | 0.05165924746472584 |
| KRT33B | 0.0018854914919091375 |
| POU1F1 | 0.001968463608032277 |

| | |
|---|---|
| MUC1 | 0.015340453178736255 |
| AVP | -0.03623974017775202 |
| INSR | -0.07200559687285481 |
| NAALADL1 | -0.0010165593282607669 |
| IL12B | 0.09369902304859443 |
| GATA2 | -0.019232401633833163 |
| MEF2C | 0.09777138637302663 |
| CLEC4M | -0.02493680520433717 |
| RFC1 | 0.023320146009614223 |
| CDC14B | 0.0001570744684361441 |
| HMGA2 | 0.07233033972660002 |
| LAIR1 | 0.0012946246181076428 |
| CCL7 | -0.05525290613251585 |
| ZNF442 | 0.0021217095248059524 |
| NDEL1 | -0.014938650791860788 |
| ANP32E | 0.0008414262211640958 |
| DDX17 | 0.010624504900571072 |
| CHIT1 | 0.0017042028202982383 |
| PSG4 | -0.0010328661238374207 |
| TTN | 0.03969133562208978 |
| MAS1 | 0.005871102423210318 |
| DRD4 | -0.07577515006143414 |
| HOXB1 | 0.04817760522773429 |
| TH | -0.02298980468540805 |
| PGR | 0.0056064827069496 |
| CSH1 | -0.0010294405113189214 |
| UGT8 | 0.021797239011992684 |
| BRCA2 | -0.0004001008524303972 |
| FGF5 | -0.029454465531980573 |
| MMP24 | 0.010426711820916278 |
| TBPL1 | -0.009405012825298754 |
| PTN | 0.007133781784054912 |
| IRF7 | -0.054667593900032785 |
| FGF8 | 0.07140111278981238 |
| GABRA4 | -0.02364422891972487 |
| BAX | -0.047402195440266075 |
| PTTG3P | -0.00020876180747159026 |
| OR5I1 | 0.009398362467484083 |
| PTCH1 | -0.0035436404931479264 |
| BCL2L11 | -0.037703052688405965 |
| PDX1 | 0.010516918867804344 |
| WNT1 | 0.04369680334753988 |
| HOXA3 | 0.013754711326618273 |
| WNT4 | -0.016164899247737854 |
| EIF4G1 | 0.011039070602603434 |
| YBX1 | -0.010053564233932749 |
| CD24 | 0.02988863062484948 |
| MAGED2 | 0.0011459396930812245 |
| PRKDC | -0.07789656076747947 |
| NRD1 | -0.0010531918503088782 |
| CCND1 | -0.18493934739300477 |
| CDC42 | 0.11480020475125491 |
| FLOT1 | -0.020237890050965705 |
| PSMD11 | -0.04560858541280286 |
| ELOVL5 | 0.0026892349674367414 |
| MCM7 | -0.023035440617418333 |
| CHD3 | 0.001086365278281501 |
| HMGB2 | 0.010195853792630806 |
| SNRPB | 0.009292643196300646 |
| POLE3 | 0.008058861478672534 |
| ATP1B3 | -0.049105061357628356 |
| STK24 | 0.04315211111121438 |
| SLC1A5 | 0.013826124416096996 |
| POR | 0.009350598710947209 |
| KPNB1 | 0.016033673916676063 |
| MDK | 0.03179016915859888 |
| TXN2 | 0.020657983182013905 |
| JAG1 | 0.06489289364190193 |
| TSPAN6 | 0.005498471709099 |
| QDPR | -0.040191396953261585 |
| AGR2 | 0.01785423349179096 |
| CXCR4 | 0.01972775578729733 |
| LMO4 | 0.0232973500798145 |
| CRYAB | 0.021582002268535957 |
| NFIB | 0.056012093999129875 |
| ECM1 | -0.02181365703317276 |
| STXBP2 | -0.009931038665850007 |
| KIF2C | 0.007483289154081436 |
| FZR1 | 0.017831600102073526 |
| MSH2 | 0.04175805328364862 |
| SERPINA5 | -0.004965822186561536 |

| | |
|---|---|
| AURKB | 0.007573735252924805 |
| LRP5 | 0.00140006702662159 |
| RPA3 | 0.09428235665230203 |
| HSDL2 | 0.0027339165079795953 |
| CACNB3 | 0.0033579975839442143 |
| IGF1 | 0.01411703310451068 |
| YIPF4 | 0.0010880978793895207 |
| THBS3 | 0.016153803444977646 |
| EPHB2 | -0.01725448446227756 |
| BMP7 | 0.05857987441538792 |
| GATA3 | -0.003547293725863661 |
| PAK1 | -0.1202453059662451 |
| MCCC2 | -0.007785154522843759 |
| BUB1 | 0.023772293746745804 |
| KIFC1 | 0.008559426097992033 |
| S100B | 0.08900178023831674 |
| SLC35D1 | -0.0007554871546431539 |
| TGFB3 | -0.13674655463203483 |
| LLPH | -0.00021432664927591784 |
| JAG2 | -0.0001614591264209394 |
| PADI2 | 0.0051776286151684 |
| NPAT | -0.0035409649949677546 |
| VLDLR | 0.021889991829839632 |
| CDT1 | 0.013044432987764098 |
| SOX10 | 0.0037792197723009176 |
| HOXB13 | 0.016446173550618004 |
| SELPLG | 0.0022892769697621792 |
| SLIT2 | -0.07791903893529607 |
| TGFB2 | -0.1193780873760941 |
| PARP3 | 0.015434259944578655 |
| FBXL4 | 0.004265644090267209 |
| GSK3B | 0.13906893656117805 |
| VEGFC | -0.12818045716407228 |
| FAP | -0.007189592876954757 |
| ASCL1 | -0.06877170297567121 |
| GATA6 | -0.001112533276770192 |
| APEX1 | 0.0035320394450440934 |
| IDO1 | 0.026842061222851387 |
| PGM3 | 0.02388703108559418 |
| TPX2 | 0.009385015564002133 |
| CTSV | 0.012273549678511547 |
| AGER | -0.008157913176116412 |
| IGFBP3 | 0.016694774973335505 |
| CYP4B1 | -0.008157465159450511 |
| FOXA2 | -0.0008266461929169179 |
| INHA | 0.005488558464491148 |
| PTPRJ | -0.007016215211972421 |
| CRYAA | -0.0040075893217782895 |
| WNT7A | 0.0010529178437344014 |
| CYP2B7P | 0.00013132082203898819 |
| RECQL5 | 0.012637809951031557 |
| C8G | -0.010678030681020804 |
| BCL11A | -0.01459751416609687 |
| IFNG | -0.004231728952820177 |
| RBBP4 | -0.03534281334268238 |
| GLUD2 | -0.007176789123447404 |
| GPR125 | 0.0012335524835453975 |
| NRP1 | -0.042420062488698854 |
| VEGFA | 0.0666633083753416 |
| MSH4 | -0.0014911312037883824 |
| SPAM1 | -0.017120073185159724 |
| POLDIP3 | -0.0003304417171146348 |
| MFGE8 | 0.016281240676370125 |
| EPHB1 | 0.04531887315980476 |
| CALCOCO2 | 0.00987327046847262 |
| ACVRL1 | 0.08556964022928057 |
| LEF1 | -0.008738760148707024 |
| GCNT2 | 0.0365829825538686 |
| AURKC | 0.009679931011797385 |
| AR | 0.08189387341717855 |
| IFNA2 | -0.07366269960649005 |
| PADI4 | -0.00904727123774782 |
| BMP4 | -0.05194854849590268 |
| GNRHR | 0.0036862891801634624 |
| HSPA2 | -0.0021826014130113943 |
| ETV4 | 0.0017136601329301375 |
| FBL | -0.0018366177387389546 |
| BDH1 | -0.0018446615434380591 |
| PRG2 | -0.007673598049155526 |
| GINS4 | -0.011529877940197965 |
| KMT2D | -0.0025206311151364166 |

| | |
|---|---|
| IGLJ3 | 0.003248875669633096 |
| IGK | 0.0029472733191604097 |
| PLCB1 | 0.010705113862137874 |
| ITGB1 | 0.15327054269447868 |
| IPO5 | 0.010054180206643165 |
| TMEM123 | -0.0013691983958083512 |
| TOP2B | 0.0034448039717102003 |
| MKI67 | 0.014176230928602422 |
| TPGS2 | 0.001830430141403624 |
| KMT2A | 0.015485947628582028 |
| SLC39A14 | 0.0009186088282803245 |
| MCM4 | -0.012131811779754908 |
| RNF187 | -0.011075139468168136 |
| SERPINE2 | 0.06795055647096451 |
| MAP1B | -0.005304206896318236 |
| SEL1L3 | 0.001482943805671271 |
| YIPF6 | 0.0004924847107853044 |
| SULF1 | -0.017637187191465103 |
| FEM1B | -0.029090201626783614 |
| MYH10 | -0.00953057846447447 |
| ZCCHC24 | -0.0006336174544859331 |
| KIAA0232 | -0.0025749317470091723 |
| TENC1 | -0.0016540678942851385 |
| CEBPB | -0.009142993773767153 |
| GPD1L | 0.011155131643536167 |
| SPIDR | -0.01805494699474201 |
| CDC34 | -0.0021652700354158065 |
| ZNF609 | -0.0020932604936431992 |
| METAP1 | -0.00802137483893706 |
| SLC25A44 | 0.0003453198541108317 |
| JMJD6 | 0.07225247109460839 |
| C16orf45 | 0.001512985360780316 |
| RCHY1 | -0.012047180085251727 |
| BBS4 | -0.024491286185357834 |
| TCF7L2 | 0.022737387684181265 |
| FAM171A1 | -0.0008474461993496483 |
| OBSL1 | 0.0004996753392981059 |
| NCAPD3 | -0.0062919453240963125 |
| CBS | 0.02740814756860186 |
| CKAP5 | -0.00784421841174049 |
| POLD3 | 0.04009422205891675 |
| AXIN1 | -0.04008779072142446 |
| DNAJC16 | -0.00010774179655897079 |
| SMC5 | -0.00840435524672013 |
| NCAPH | 0.0005077986713116508 |
| KANK1 | 0.03570418486402972 |
| NEDD4 | 0.08611271769809645 |
| LOC100272216 | -0.0004901303998456613 |
| DNAJC2 | -0.00785480274112632 |
| HOXA10 | -0.03585348281798796 |
| ROBO1 | -0.03381507510547673 |
| TNNT1 | 0.0023616702764739797 |
| FOXC1 | -0.0010032433396977894 |
| YAP1 | -0.00521093327438735 |
| ATAD2B | -0.0012403210541986134 |
| SPDEF | 0.03149979170441675 |
| SOSTDC1 | -0.012182241980776625 |
| CLCN2 | 0.008450306755612699 |
| CCNE1 | 0.013357563921641034 |
| DNA2 | -0.0005924515358255857 |
| DLX5 | -0.06904625505431028 |
| ELOVL2 | 0.0060805461246128895 |
| HIPK2 | 0.0283944848486138 |
| HOXA5 | 0.02651939251917732 |
| TWIST1 | -0.07355870775008264 |
| CHD5 | -0.004060582690920923 |
| YPEL1 | 0.002844755706654635 |
| CCL8 | -0.014706434092778383 |
| LIAS | -0.0043664347027258495 |
| SHMT2 | -0.01310599052478396 |
| XIST | -0.0013837068271472972 |
| GMPS | 0.017799732080132098 |
| FOXC2 | 0.06632063111815426 |
| PTTG2 | -0.01371016425337387 |
| FGF3 | 0.04819322200362268 |
| AZU1 | 0.00530291009094901 |
| TNFRSF21 | 0.006502653335445799 |
| PAK3 | -0.04013955338735342 |
| EYA1 | 0.0017480075922933923 |
| ZNF266 | 0.0008265747472587705 |
| CCNB1 | -0.03598713934059483 |

| | |
|---|---|
| PP14571 | -0.0025847593046036026 |
| KCNMA1 | -0.021521345148707603 |
| GRIK5 | -0.0028890162313301598 |
| RNASEH2B | -0.0003486624368721747 |
| GRB2 | 0.028244434541645335 |
| DNM1 | -0.02218182942553517 |
| CEP152 | 0.016242950198095015 |
| STOML2 | 0.0051736598617511 |
| IHH | -0.0981336737412036 |
| EPHA5 | 0.006530393686639469 |
| RAD21L1 | -0.011228435521741088 |
| POLE | 0.05615385969439244 |
| RAB3GAP2 | 0.013865146208784956 |
| LAMB2 | -0.02355652281477592 |
| MUC7 | -0.015812317014986945 |
| CCL2 | 0.07777608367830818 |
| IFT122 | -0.03543911313873816 |
| LOC100293211 | 0.002250473974020906 |
| DISC1 | -0.03731677569455142 |
| KCNV2 | 0.003696900978180594 |
| ORAI2 | 0.002164600637607095 |
| ARPIN | -0.0005814888777068246 |
| STEAP1B | 0.002804840397185504 |
| SEC14L1P1 | 0.004847184538895704 |
| AES | 0.016231813272144718 |
| PERP | 0.014624321373756952 |
| NAA50 | 0.000696110791108242 1 |
| HN1 | -5.812779117379615e-5 |
| C3 | 0.0830268588437952 |
| UFC1 | 0.002081909240303291 |
| EVL | 0.023265625083061247 |
| CDC27 | 0.007420671443608753 |
| DSG2 | 0.00017329429395628 |
| KLHDC2 | -0.0003758990939784145 |
| VPS4A | -0.018885741648463586 |
| FAM127B | 0.002068826100125258 |
| PRC1 | 0.004887640416565872 |
| PPDPF | -0.004333499725805429 |
| FAM65A | -0.0033104341627444413 |
| RBM47 | 0.0008135753629212931 |
| NUSAP1 | -0.0031989576398166897 |
| BFAR | -0.0032646454557970607 |
| WDR11 | 0.0004084790987555989 |
| C21orf59 | 0.001276663114000037 |
| SLC25A37 | -0.001101948341530599 |
| SMAP1 | 0.004010691731589973 |
| GSDMD | 0.0023281873418337156 |
| GOLT1B | -0.009110027865084448 |
| STMN3 | 0.0207922491669522 |
| TMEM258 | 0.0036313222830984967 |
| LANCL2 | 0.003563536119480891 |
| C1QA | 0.005377251754758939 |
| TSKU | 0.00651410414586309 |
| CKAP2 | 0.0001859579560697081 |
| MKL2 | -0.0007206805073324864 |
| BCCIP | 0.005814671919895755 |
| SAV1 | -0.03272071156694 |
| LGR4 | 0.006571083295942736 |
| PRDM4 | 0.008370680347239904 |
| KIF4A | 0.015508808710484129 |
| PGLS | 0.010472817234260473 |
| FAM204A | -0.004478857609673523 |
| ABT1 | 0.0179616691415539 4 |
| NDE1 | -0.031881842308720244 |
| STEAP3 | 0.02882076318717924 |
| MCCC1 | 0.0007363760398943163 |
| TTC4 | 0.0003864077187552611 |
| CMC2 | -0.00024523649873989684 |
| GREM1 | -0.03628044658326054 |
| KLF11 | 0.0037117930552162098 |
| RNASEH1 | 0.0016941513093079123 |
| TRPS1 | -0.0028047543800425578 |
| DCP1A | -0.0059072595504989 |
| LPPR2 | -0.0013568981277233033 |
| PNPO | -0.011992318969413343 |
| MAP1S | 0.0042251920556763654 |
| E4F1 | 0.017773349853668404 |
| MRS2 | 0.001948042410551421 |
| C8orf4 | -0.0010783892034500855 |
| CEP55 | 0.00018269415358942 8 |
| KBTBD4 | 0.0021944441349361693 |

| | |
|---|---|
| RINT1 | -0.006919338654133502 |
| SMO | 0.1371278839293916 |
| NCAPG | 0.000693116427805255З |
| S100A14 | -0.015165150621937296 |
| NES | 0.015184277222533585 |
| GINS3 | -0.00027924086714144815 |
| HJURP | -0.00045926730263409 |
| CDK5RAP3 | 0.022624477001446944 |
| KIF20A | -0.001686633809426736 |
| SMC6 | -0.01398471781996997 |
| ATAD2 | -0.0001941253739088845 |
| FCGRT | 0.001765194609950125 |
| IGF2BP2 | -0.004613114653713914 |
| PPP1R13L | -0.017696251377432606 |
| FBXO5 | 0.007503733417388883 |
| SIRT1 | -0.0009599762338105395 |
| NOTCH1 | 0.06908067138798721 |
| INTS8 | -0.0045263903469869495 |
| RAB17 | 0.00235030694754029 |
| HSPB7 | 0.0015844318424696417 |
| KRT23 | -0.000734650426396615l |
| DNAJC12 | 0.0007439037392301431 |
| METRN | -0.005624192028500556 |
| CYB5R4 | -0.00021192099129545034 |
| CTPS2 | 0.014098873505983605 |
| SHQ1 | -0.004306154966906718 |
| IL20RA | 0.0008584941824028729 |
| PBK | 0.005020736063415776 |
| BIN2 | -0.006561411567025762 |
| SCUBE2 | 0.0025221661308145404 |
| HSPA14 | 0.0002787788487917095 |
| RBKS | 0.0008862409994960207 |
| PHACTR4 | -0.016079244020787184 |
| CEP63 | -0.00015742808684255963 |
| SPHK1 | 0.06878627113250543 |
| TIPIN | -0.0048284433759789025 |
| C1orf159 | -0.0013948275277618286 |
| DIABLO | -0.005344035959869676 |
| ATHL1 | -0.0009901807536505773 |
| KLF2 | -0.02328942670693097 |
| GRHL2 | 0.025092132700132403 |
| TTYH1 | 0.009566840787928561 |
| COA7 | 0.0005966881548589191 |
| TTC33 | -0.002621094875096663 |
| SULT4A1 | -0.005814538701167437 |
| EGFL6 | 0.000822197401509781 |
| C7orf63 | 0.0012350401404626558 |
| LAMP5 | -0.003579246121745118 |
| LRFN4 | 0.001393938640831942 |
| GCNT3 | -0.025506586914859186 |
| CEP72 | -0.016425670499335236 |
| BORA | 0.011964177974982481 |
| ROBO3 | 0.004799815360220849 |
| SOX18 | 0.0295000211128748l |
| KIF16B | 0.03523818751096763 |
| PIEZO2 | 0.0020898293231342603 |
| CEP83 | -0.0007677935602652956 |
| PTPLA | 0.002554874314263294 |
| TBX3 | 0.05010579689726048 |
| FZD3 | 0.009331988846620856 |
| BBS7 | -0.020445120694412327 |
| TFCP2L1 | 0.004859367748266085 |
| HEY2 | 0.13883585381515864 |
| VTCN1 | -0.011060391877614666 |
| GIMAP6 | -0.0007352876517308436 |
| MTL5 | -0.0025622556198957327 |
| ECT2 | 0.024320114192369513 |
| ARL15 | -0.004536728216233619 |
| NARF | 0.0014994215563336726 |
| FAM198B | 9.679876491163116e-5 |
| LHX6 | 0.009407866917839711 |
| SPAG4 | 0.0006137799805040558 |
| ASPM | 0.0002548203396439101 |
| GLRX2 | 0.016675863751174958 |
| GPR87 | -7.288109362732377e-5 |
| GALNT6 | -0.010690878808872865 |
| KIZ | 3.31175041513901e-5 |
| TXLNG | -0.005025118529839369 |
| E2F8 | 0.020587127716812 |
| AUNIP | -0.00044028577435419595 |
| ERO1LB | 0.0018389095331322214 |

```
MFI2	0.004359643072220506
EDAR	0.005933150194892732
GDF3	-0.039493719443255915
STAP1	0.03192302906622345
HELLS	0.02789855780890322
STEAP4	-0.01391388036506503
CAMK1D	-0.031182194982635195
ARPP21	-0.0008943828850588785
ALLC	-0.0027650432492677406
ROPN1B	0.004169631433242046
DMRT1	0.02853845282955854
EN1	-0.00021794323410886883
CCDC170	-0.0009410021876603598
SIRT2	-0.048816855988439746
FOXE3	0.01465172213949957
ELF5	0.011752955688516507
MCM10	-0.0041918103763885755
KIF24	-0.011283208386569682
HEYL	0.013492537971960218
TBX21	0.0038694007489718366
SLC5A7	0.02006523348792231
CCDC7	-0.001419948952359388
EDDM3B	0.0010086661202008459
TP73	0.0420453123927202
GCNT4	0.00538011446141364
C1orf112		0.001533231819017808
PSAT1	-0.005765858129629997
WDR19	-0.010047564609988216
C1QTNF3	0.0015438782131631129
ARHGAP24		-0.0074784006899104895
TEX13B	-0.0009161916424618885
TEX14	0.015492611545399734
KCNE2	-0.011773142899107383
EPN1	-0.004333698632650037
RPA4	-0.02490832447198217
MAP3K19	-0.008239112962222229
IQCG	0.0014698666492631838
LOC100131532	-0.005151893057862087
STMN4	0.0013959838112425872
KIF18A	0.012850376596106073
DLX6	-0.001800012326516208
KLF15	0.0036817542319530017
FRS2	0.02771376092040482
NEUROD4	0.015071128029595655
C6orf25	-0.0004983610788643351
GDNF	0.014056253615913419
TMEM14B	0.003590952595578687
BNIP3L	-0.0004637209207129531
KPNA3	-0.004073128755144697
GINS2	0.003650234479374486
SOAT1	-0.0035840099220547086
FAM64A	0.00022582903076745391
DSPP	0.004961435490698744
SPDL1	0.02250710639363425
STRA6	-0.06393712883327846
COL5A2	-0.00581222376250594
NRBF2	-0.006598199026985765
GPR124	-0.002425575121783928
KLHL22	0.003054254639106805
ORAI3	-0.0026187771148240353
FAM174B	-6.729041772863932e-5
ZNF335	-0.004675023254393562
SEH1L	-0.022554065088721768
LOC389906		0.005430567767702915
KIF18B	0.015702773273153524
RACGAP1	0.026271373792290355
CASP8AP2		0.012679791553399917
KLK5	0.00695160749560311
OR7E47P	0.0005763425767187261
SCAF4	-0.0003286804849717546
SNHG17	-0.0011774470329610934
Age			-0.002246735353315669
intercept			-0.06328011266889626
```

**(optional) analysis using GPU**

Once again, the analysis can be carried out using a GPU. At this time, data should be converted to `CuArray` s beforehand for overlapping groups.

In [53]:
```
Random.seed!(222)
T = Float64
A = CuArray
U = ParProx.LogisticUpdate(; maxiter=30000, step=20, tol=5e-4, verbose=false)
lambdas = 10 .^ (range(-6, stop=-9, length=31))

@time scores = ParProx.cross_validate(U,
    adapt(A{T}, X), adapt(A{T}, X_unpen), adapt(A{Int32}, y), group_to_variables, lambdas, 5;
    T=Float64, criteria=auc);
```

```
 18.327784 seconds (26.58 M allocations: 15.006 GiB, 3.71% gc time)
```

In [54]:
```
lambda_idx = argmax(mean(scores; dims=2)[:])
lambda = lambdas[lambda_idx]
```

Out[54]:
```
1.2589254117941663e-9
```

In [55]:
```
U = ParProx.LogisticUpdate(; maxiter=30000, step=20, tol=1e-5, verbose=false)
V = ParProx.LogisticVariables{Float64}(adapt(A{T}, X),
    adapt(A{T}, X_unpen), adapt(A{Int}, y), lambda, group_to_variables)
@time ParProx.fit!(U, V)
```

```
 23.080018 seconds (23.83 M allocations: 35.875 GiB, 5.21% gc time)
```

In [56]:
```
β_orig = vcat(grpmat * collect(V.β[1:end-2]), collect(V.β)[end-1:end]);
```

In [57]:

```
for (v, β) in zip(variable_names_replicated[V.β .!= 0], V.β[V.β .!= 0])
    println("$v\t$β")
end
```

KEGG:00280: Valine, leucine and isoleucine degradation ABAT 4-aminobutyrate aminotransferase 0.0008323544134017856
KEGG:00280: Valine, leucine and isoleucine degradation AUH AU RNA binding protein/enoyl-CoA hydratase -0.0014593426888791475
KEGG:00280: Valine, leucine and isoleucine degradation HMGCL 3-hydroxymethyl-3-methylglutaryl-CoA lyase -0.0010615725170478399
KEGG:00280: Valine, leucine and isoleucine degradation IVD isovaleryl-CoA dehydrogenase -0.0009348559171503037
KEGG:00280: Valine, leucine and isoleucine degradation MCCC1 methylcrotonoyl-CoA carboxylase 1 (alpha) 8.306605556755177e-5
KEGG:00280: Valine, leucine and isoleucine degradation MCCC2 methylcrotonoyl-CoA carboxylase 2 (beta) -0.0007834944198441148
KEGG:00280: Valine, leucine and isoleucine degradation MUT methylmalonyl CoA mutase 0.0027548880164145425
KEGG:04727: GABAergic synapse ABAT 4-aminobutyrate aminotransferase 0.0008232735329612731
KEGG:04727: GABAergic synapse GABRA4 gamma-aminobutyric acid (GABA) A receptor, alpha 4 -0.0014658463388531483
KEGG:04727: GABAergic synapse PRKCA protein kinase C, alpha -5.901736217620194e-6
KEGG:04727: GABAergic synapse PRKX protein kinase, X-linked 0.0003862539887885204
KEGG:01100: Metabolic pathways ABAT 4-aminobutyrate aminotransferase 0.0008361539792630331
KEGG:01100: Metabolic pathways ACADL acyl-CoA dehydrogenase, long chain 0.000985316179371731
KEGG:01100: Metabolic pathways ALDH1A2 aldehyde dehydrogenase 1 family, member A2 -0.0030292702181800245
KEGG:01100: Metabolic pathways ALDH5A1 aldehyde dehydrogenase 5 family, member A1 0.002338405717850529
KEGG:01100: Metabolic pathways ALLC allantoicase -0.0009259344378902447
KEGG:01100: Metabolic pathways ALOX12 arachidonate 12-lipoxygenase -0.001960674381048052
KEGG:01100: Metabolic pathways AMD1 adenosylmethionine decarboxylase 1 -0.0025358903001866883
KEGG:01100: Metabolic pathways ASS1 argininosuccinate synthase 1 0.0001442647137212183
KEGG:01100: Metabolic pathways AUH AU RNA binding protein/enoyl-CoA hydratase -0.0014668253656568752
KEGG:01100: Metabolic pathways BDH1 3-hydroxybutyrate dehydrogenase, type 1 -0.00020747510439733985
KEGG:01100: Metabolic pathways CBS cystathionine-beta-synthase 0.001522422413483285
KEGG:01100: Metabolic pathways CPOX coproporphyrinogen oxidase 0.0012789641964398662
KEGG:01100: Metabolic pathways CTH cystathionine gamma-lyase 0.0017546337885964072
KEGG:01100: Metabolic pathways CTPS1 CTP synthase 1 -0.0004215525129638922
KEGG:01100: Metabolic pathways CTPS2 CTP synthase 2 0.0017650016553290638
KEGG:01100: Metabolic pathways CYP27B1 cytochrome P450, family 27, subfamily B, polypeptide 1 -0.0047356623759338824
KEGG:01100: Metabolic pathways DCT dopachrome tautomerase -0.0023607667959928773
KEGG:01100: Metabolic pathways DHCR24 24-dehydrocholesterol reductase -0.001713739776933617
KEGG:01100: Metabolic pathways DHFR dihydrofolate reductase -0.0003076937765407045
KEGG:01100: Metabolic pathways DTYMK deoxythymidylate kinase (thymidylate kinase) 0.0034935954624852592
KEGG:01100: Metabolic pathways GAL3ST1 galactose-3-O-sulfotransferase 1 0.0018631853227812844
KEGG:01100: Metabolic pathways GALK2 galactokinase 2 3.174981063618625e-5
KEGG:01100: Metabolic pathways GALNT6 polypeptide N-acetylgalactosaminyltransferase 6 -0.0017830737236985237
KEGG:01100: Metabolic pathways GCDH glutaryl-CoA dehydrogenase -5.142324143013628e-6
KEGG:01100: Metabolic pathways GCLM glutamate-cysteine ligase, modifier subunit -0.0037732217757471516
KEGG:01100: Metabolic pathways GCNT1 glucosaminyl (N-acetyl) transferase 1, core 2 0.0006985769533269946
KEGG:01100: Metabolic pathways GCNT2 glucosaminyl (N-acetyl) transferase 2, I-branching enzyme (I blood group) 0.0024411127133358054
KEGG:01100: Metabolic pathways GCNT3 glucosaminyl (N-acetyl) transferase 3, mucin type -0.0023204484305728522
KEGG:01100: Metabolic pathways GCNT4 glucosaminyl (N-acetyl) transferase 4, core 2 0.0004535246682822453
KEGG:01100: Metabolic pathways GLUD2 glutamate dehydrogenase 2 -0.0006607507670226813
KEGG:01100: Metabolic pathways GMPS guanine monphosphate synthase 0.0017809600837371212
KEGG:01100: Metabolic pathways GUK1 guanylate kinase 1 0.003284003228292133
KEGG:01100: Metabolic pathways HMGCL 3-hydroxymethyl-3-methylglutaryl-CoA lyase -0.0010678229674893761
KEGG:01100: Metabolic pathways HPRT1 hypoxanthine phosphoribosyltransferase 1 -0.0011894467283326516
KEGG:01100: Metabolic pathways HSD17B4 hydroxysteroid (17-beta) dehydrogenase 4 -0.0011670301666150764
KEGG:01100: Metabolic pathways HSD17B6 hydroxysteroid (17-beta) dehydrogenase 6 -0.0008712071093238522
KEGG:01100: Metabolic pathways IDO1 indoleamine 2,3-dioxygenase 1 0.001230768016722825
KEGG:01100: Metabolic pathways IMPDH1 IMP (inosine 5'-monophosphate) dehydrogenase 1 -0.0018789353029630793
KEGG:01100: Metabolic pathways IMPDH2 IMP (inosine 5'-monophosphate) dehydrogenase 2 -0.0003495009634217896
KEGG:01100: Metabolic pathways INPP4B inositol polyphosphate-4-phosphatase, type II, 105kDa 0.000289393359683728
KEGG:01100: Metabolic pathways IVD isovaleryl-CoA dehydrogenase -0.0009384574293990822
KEGG:01100: Metabolic pathways LDHC lactate dehydrogenase C 0.001488418240397908

KEGG:01100: Metabolic pathways  LIAS    lipoic acid synthetase  -0.00055594774308089
KEGG:01100: Metabolic pathways  MCCC1   methylcrotonoyl-CoA carboxylase 1 (alpha)       8.57497893915
0576e-5
KEGG:01100: Metabolic pathways  MCCC2   methylcrotonoyl-CoA carboxylase 2 (beta)        -0.0007881466
527775403
KEGG:01100: Metabolic pathways  MGAM    maltase-glucoamylase (alpha-glucosidase)        -0.0013146570
407050655
KEGG:01100: Metabolic pathways  MTAP    methylthioadenosine phosphorylase       0.002668361053392182
KEGG:01100: Metabolic pathways  MTHFD1  methylenetetrahydrofolate dehydrogenase (NADP+ dependent) 1,
methenyltetrahydrofolate cyclohydrolase, formyltetrahydrofolate synthetase      -0.00066559434303863
5
KEGG:01100: Metabolic pathways  MUT     methylmalonyl CoA mutase        0.00276742988524220
KEGG:01100: Metabolic pathways  ODC1    ornithine decarboxylase 1       0.000828622843825379
KEGG:01100: Metabolic pathways  OGDH    oxoglutarate (alpha-ketoglutarate) dehydrogenase (lipoamide)
-0.000295625570768411
KEGG:01100: Metabolic pathways  PGLS    6-phosphogluconolactonase       0.001753938225770404
KEGG:01100: Metabolic pathways  PHGDH   phosphoglycerate dehydrogenase  0.000295136452440176
KEGG:01100: Metabolic pathways  PLA2G5  phospholipase A2, group V       0.00208546105535219
KEGG:01100: Metabolic pathways  PLCB1   phospholipase C, beta 1 (phosphoinositide-specific)     0.000
1551250705580811
KEGG:01100: Metabolic pathways  PNPO    pyridoxamine 5'-phosphate oxidase       -0.001095584789131929
9
KEGG:01100: Metabolic pathways  POLA1   polymerase (DNA directed), alpha 1, catalytic subunit   0.001
2769114916353394
KEGG:01100: Metabolic pathways  POLA2   polymerase (DNA directed), alpha 2, accessory subunit   -0.00
14984362498909225
KEGG:01100: Metabolic pathways  POLD1   polymerase (DNA directed), delta 1, catalytic subunit   -0.00
04456093543240708
KEGG:01100: Metabolic pathways  POLD2   polymerase (DNA directed), delta 2, accessory subunit   0.000
742147794945639
KEGG:01100: Metabolic pathways  POLD3   polymerase (DNA-directed), delta 3, accessory subunit   0.001
746658741210305
KEGG:01100: Metabolic pathways  POLD4   polymerase (DNA-directed), delta 4, accessory subunit   -0.00
06409905849416638
KEGG:01100: Metabolic pathways  POLE    polymerase (DNA directed), epsilon, catalytic subunit   0.002
9581383098528655
KEGG:01100: Metabolic pathways  POLE2   polymerase (DNA directed), epsilon 2, accessory subunit 0.001
3307107678158523
KEGG:01100: Metabolic pathways  POLE3   polymerase (DNA directed), epsilon 3, accessory subunit 0.000
8084517388967186
KEGG:01100: Metabolic pathways  POLR2D  polymerase (RNA) II (DNA directed) polypeptide D        0.001
8615916236063403
KEGG:01100: Metabolic pathways  PRIM1   primase, DNA, polypeptide 1 (49kDa)     0.000637600483659455
KEGG:01100: Metabolic pathways  PRIM2   primase, DNA, polypeptide 2 (58kDa)     -0.00089033201194521
5
KEGG:01100: Metabolic pathways  PSAT1   phosphoserine aminotransferase 1        -0.000724130502531340
9
KEGG:01100: Metabolic pathways  PSPH    phosphoserine phosphatase       -0.001940838337812198
KEGG:01100: Metabolic pathways  QDPR    quinoid dihydropteridine reductase      -0.002865829961782043
6
KEGG:01100: Metabolic pathways  RRM1    ribonucleotide reductase M1     0.0019181942335011152
KEGG:01100: Metabolic pathways  RRM2    ribonucleotide reductase M2     -0.0005663338491549913
KEGG:01100: Metabolic pathways  SHMT2   serine hydroxymethyltransferase 2 (mitochondrial)       -0.00
11001374615877285
KEGG:01100: Metabolic pathways  SORD    sorbitol dehydrogenase  -0.002027239084487823
KEGG:01100: Metabolic pathways  SPAM1   sperm adhesion molecule 1 (PH-20 hyaluronidase, zona pellucid
a binding)      -0.0019057089696524507
KEGG:01100: Metabolic pathways  SPHK1   sphingosine kinase 1    0.0018174636412165578
KEGG:01100: Metabolic pathways  SPR     sepiapterin reductase (7,8-dihydrobiopterin:NADP+ oxidoreduct
ase)    -0.0012378419013538608
KEGG:01100: Metabolic pathways  TAT     tyrosine aminotransferase       0.0010569571988997724
KEGG:01100: Metabolic pathways  TH      tyrosine hydroxylase    -0.0003428729676016945
KEGG:01100: Metabolic pathways  TYMS    thymidylate synthetase  0.0015685929022796515
KEGG:01100: Metabolic pathways  UGCG    UDP-glucose ceramide glucosyltransferase        0.00036935096
31973446
KEGG:01100: Metabolic pathways  UGDH    UDP-glucose 6-dehydrogenase     0.0005025506410051844
KEGG:01100: Metabolic pathways  UGP2    UDP-glucose pyrophosphorylase 2 0.0007901636826903183
KEGG:01100: Metabolic pathways  UGT8    UDP glycosyltransferase 8       0.0024046174229095845
KEGG:00250: Alanine, aspartate and glutamate metabolism ABAT    4-aminobutyrate aminotransferase
0.0008330016098156058
KEGG:00250: Alanine, aspartate and glutamate metabolism ALDH5A1 aldehyde dehydrogenase 5 family, memb
er A1   0.00232802891434874
KEGG:00250: Alanine, aspartate and glutamate metabolism ASS1    argininosuccinate synthase 1    0.000
142213329414408
KEGG:00250: Alanine, aspartate and glutamate metabolism GLUD2   glutamate dehydrogenase 2       -0.00
06588423570298241
KEGG:00640: Propanoate metabolism       ABAT    4-aminobutyrate aminotransferase        0.00083838480
57490676
KEGG:00640: Propanoate metabolism       LDHC    lactate dehydrogenase C 0.001492969939971959
KEGG:00640: Propanoate metabolism       MUT     methylmalonyl CoA mutase        0.0027741889752544036
KEGG:00410: beta-Alanine metabolism     ABAT    4-aminobutyrate aminotransferase        0.00082173624

43510856
KEGG:00650: Butanoate metabolism ABAT 4-aminobutyrate aminotransferase 0.00083427856
86320112
KEGG:00650: Butanoate metabolism ALDH5A1 aldehyde dehydrogenase 5 family, member A1 0.002
331490731821375
KEGG:00650: Butanoate metabolism BDH1 3-hydroxybutyrate dehydrogenase, type 1 -0.0002073436
3851731576
KEGG:00650: Butanoate metabolism HMGCL 3-hydroxymethyl-3-methylglutaryl-CoA lyase -0.00
1064392202790467
GO:0001666: response to hypoxia ABAT 4-aminobutyrate aminotransferase 0.0008334434534379155
GO:0001666: response to hypoxia ADM adrenomedullin 0.0022388125598973474
GO:0001666: response to hypoxia AGER advanced glycosylation end product-specific receptor -0.00
01742645423739626
GO:0001666: response to hypoxia BNIP3 BCL2/adenovirus E1B 19kDa interacting protein 3 0.00291415084
36547
GO:0001666: response to hypoxia CAV1 caveolin 1, caveolae protein, 22kDa -0.000535471952848710
1
GO:0001666: response to hypoxia CCL2 chemokine (C-C motif) ligand 2 0.0008172213365582937
GO:0001666: response to hypoxia CD24 CD24 molecule 0.0010725532147139426
GO:0001666: response to hypoxia CHRNB2 cholinergic receptor, nicotinic, beta 2 (neuronal) -0.00
0778801052026624
GO:0001666: response to hypoxia CRYAB crystallin, alpha B 0.0009855156589479084
GO:0001666: response to hypoxia CST3 cystatin C -7.084825260829531e-5
GO:0001666: response to hypoxia CX3CL1 chemokine (C-X3-C motif) ligand 1 0.002118492984099295
GO:0001666: response to hypoxia CXCL12 chemokine (C-X-C motif) ligand 12 -0.001209203588890178
GO:0001666: response to hypoxia CXCR4 chemokine (C-X-C motif) receptor 4 0.0007954308332594616
GO:0001666: response to hypoxia DRD2 dopamine receptor D2 -0.0002341132496289484
GO:0001666: response to hypoxia ENG endoglin 0.0008271176427567823
GO:0001666: response to hypoxia HIF1A hypoxia inducible factor 1, alpha subunit (basic helix-loop-h
elix transcription factor) -0.0006464543407707821
GO:0001666: response to hypoxia ITPR1 inositol 1,4,5-trisphosphate receptor, type 1 -0.0007812013
655777393
GO:0001666: response to hypoxia KCNMA1 potassium large conductance calcium-activated channel, subfam
ily M, alpha member 1 -0.0006595603547464599
GO:0001666: response to hypoxia LEP leptin 0.0031927134861560623
GO:0001666: response to hypoxia MMP2 matrix metallopeptidase 2 (gelatinase A, 72kDa gelatinase, 72
kDa type IV collagenase) -0.0011323873096091158
GO:0001666: response to hypoxia PML promyelocytic leukemia -0.0006798120904419596
GO:0001666: response to hypoxia RAF1 Raf-1 proto-oncogene, serine/threonine kinase 0.00149944125
9070323
GO:0001666: response to hypoxia SLC11A2 solute carrier family 11 (proton-coupled divalent metal ion t
ransporter), member 2 -0.0006653504084603694
GO:0001666: response to hypoxia TFRC transferrin receptor 0.0010849330467224143
GO:0001666: response to hypoxia TGFB1 transforming growth factor, beta 1 -7.324269878783433e-5
GO:0001666: response to hypoxia TGFB2 transforming growth factor, beta 2 -0.001059988161415871
GO:0001666: response to hypoxia TGFB3 transforming growth factor, beta 3 -0.00182516779021556
5
GO:0001666: response to hypoxia TH tyrosine hydroxylase -0.00034212341489976423
GO:0001666: response to hypoxia THBS1 thrombospondin 1 -0.001036550292497587
GO:0001666: response to hypoxia TXN2 thioredoxin 2 0.0013835636435942493
GO:0001666: response to hypoxia VEGFA vascular endothelial growth factor A 0.0005957623131915806
GO:0007268: synaptic transmission ABAT 4-aminobutyrate aminotransferase 0.00083441285
8277824
GO:0007268: synaptic transmission ALDH5A1 aldehyde dehydrogenase 5 family, member A1 0.002
325736785758145
GO:0007268: synaptic transmission ASIC2 acid-sensing (proton-gated) ion channel 2 0.002
277005624832919
GO:0007268: synaptic transmission BCHE butyrylcholinesterase -5.891714260374989e-6
GO:0007268: synaptic transmission CACNB3 calcium channel, voltage-dependent, beta 3 subunit
0.0002124849861672683
GO:0007268: synaptic transmission CHRNB2 cholinergic receptor, nicotinic, beta 2 (neuronal)
-0.0007771671025138416
GO:0007268: synaptic transmission CREB1 cAMP responsive element binding protein 1 0.000
6560337387386202
GO:0007268: synaptic transmission CTNNB1 catenin (cadherin-associated protein), beta 1, 88kDa
-0.00011771406287013875
GO:0007268: synaptic transmission DRD4 dopamine receptor D4 -0.001942610174312302
GO:0007268: synaptic transmission GABRA4 gamma-aminobutyric acid (GABA) A receptor, alpha 4
-0.0014819946432346233
GO:0007268: synaptic transmission GRIK5 glutamate receptor, ionotropic, kainate 5 -0.00
023303218514897277
GO:0007268: synaptic transmission HTR2C 5-hydroxytryptamine (serotonin) receptor 2C, G protei
n-coupled -0.0005243373591696027
GO:0007268: synaptic transmission HTR6 5-hydroxytryptamine (serotonin) receptor 6, G protein
-coupled -0.0010808645611208798
GO:0007268: synaptic transmission KCNJ2 potassium inwardly-rectifying channel, subfamily J, m
ember 2 -0.0007757206538099177
GO:0007268: synaptic transmission KCNMA1 potassium large conductance calcium-activated channe
l, subfamily M, alpha member 1 -0.0006552123640800504
GO:0007268: synaptic transmission KCNV2 potassium channel, subfamily V, member 2 0.000
925023789245980l

GO:0007268: synaptic transmission LRP6 low density lipoprotein receptor-related protein 6
0.00014822359348205615
GO:0007268: synaptic transmission PICK1 protein interacting with PRKCA 1 -0.0009541127
470417695
GO:0007268: synaptic transmission PLCB1 phospholipase C, beta 1 (phosphoinositide-specific)
0.00015258877278244255
GO:0007268: synaptic transmission PRKCA protein kinase C, alpha -6.141223100982578e-6
GO:0007268: synaptic transmission RAF1 Raf-1 proto-oncogene, serine/threonine kinase 0.001
4964152404464423
GO:0007268: synaptic transmission RPS6KA1 ribosomal protein S6 kinase, 90kDa, polypeptide 1
-0.0025108737549834137
GO:0007268: synaptic transmission SLC1A1 solute carrier family 1 (neuronal/epithelial high aff
inity glutamate transporter, system Xag), member 1 -0.0008991927780949918
GO:0007268: synaptic transmission SLC1A3 solute carrier family 1 (glial high affinity glutamat
e transporter), member 3 -0.0031174552126335174
GO:0007268: synaptic transmission SLC5A7 solute carrier family 5 (sodium/choline cotransporte
r), member 7 0.002009606065392056
GO:0007268: synaptic transmission SYT1 synaptotagmin I -0.0014400204030124307
GO:0007269: neurotransmitter secretion ABAT 4-aminobutyrate aminotransferase 0.00083821095
914872
GO:0007269: neurotransmitter secretion ALDH5A1 aldehyde dehydrogenase 5 family, member A1 0.002
3423281457344227
GO:0007269: neurotransmitter secretion SLC1A1 solute carrier family 1 (neuronal/epithelial high aff
inity glutamate transporter, system Xag), member 1 -0.0009061343508545289
GO:0007269: neurotransmitter secretion SLC1A3 solute carrier family 1 (glial high affinity glutamat
e transporter), member 3 -0.0031362699928707705
GO:0007269: neurotransmitter secretion SLC5A7 solute carrier family 5 (sodium/choline cotransporte
r), member 7 0.0020197735643882747
GO:0007269: neurotransmitter secretion SYT1 synaptotagmin I -0.0014518725800134712
GO:0007269: neurotransmitter secretion WNT7A wingless-type MMTV integration site family, member 7A
1.9218868493483488e-5
GO:0007620: copulation ABAT 4-aminobutyrate aminotransferase 0.0008306490238316081
GO:0007620: copulation PI3 peptidase inhibitor 3, skin-derived -0.0010312990109310953
GO:0007626: locomotory behavior ABAT 4-aminobutyrate aminotransferase 0.0008188020442316776
GO:0007626: locomotory behavior AVP arginine vasopressin -0.0009119172021728828
GO:0007626: locomotory behavior CHRNB2 cholinergic receptor, nicotinic, beta 2 (neuronal) -0.00
07604539753433607
GO:0007626: locomotory behavior DRD2 dopamine receptor D2 -0.00023041388671371223
GO:0007626: locomotory behavior HPRT1 hypoxanthine phosphoribosyltransferase 1 -0.0011642187
897423887
GO:0007626: locomotory behavior HTR2C 5-hydroxytryptamine (serotonin) receptor 2C, G protein-couple
d -0.0005148148706419105
GO:0007626: locomotory behavior HTT huntingtin -0.0009518194507996056
GO:0007626: locomotory behavior PTEN phosphatase and tensin homolog 2.4547602115369364e-5
GO:0007626: locomotory behavior TH tyrosine hydroxylase -0.00033776756931749
GO:0009449: gamma-aminobutyric acid biosynthetic process ABAT 4-aminobutyrate aminotransfer
ase 0.0008414548467394144
GO:0009449: gamma-aminobutyric acid biosynthetic process SLC1A3 solute carrier family 1 (glia
l high affinity glutamate transporter), member 3 -0.0031463396344390097
GO:0009450: gamma-aminobutyric acid catabolic process ABAT 4-aminobutyrate aminotransferase
0.0008384285830425881
GO:0009450: gamma-aminobutyric acid catabolic process ALDH5A1 aldehyde dehydrogenase 5 family, memb
er A1 0.0023452776173856047
GO:0010039: response to iron ion ABAT 4-aminobutyrate aminotransferase 0.00083414861
69944334
GO:0010039: response to iron ion BCL2 B-cell CLL/lymphoma 2 -5.3147746697044305e-6
GO:0010039: response to iron ion CCND1 cyclin D1 -0.002640764841484498
GO:0010039: response to iron ion CPOX coproporphyrinogen oxidase 0.0012771007140909793
GO:0010039: response to iron ion DRD2 dopamine receptor D2 -0.0002343433759645894
GO:0010039: response to iron ion SLC11A2 solute carrier family 11 (proton-coupled divalent met
al ion transporter), member 2 -0.0006660535507818987
GO:0010039: response to iron ion TFRC transferrin receptor 0.0010863007139147048
GO:0031652: positive regulation of heat generation ABAT 4-aminobutyrate aminotransferase
0.0008217362443510856
GO:0032024: positive regulation of insulin secretion ABAT 4-aminobutyrate aminotransferase
0.0008012149650160564
GO:0032024: positive regulation of insulin secretion GJA1 gap junction protein, alpha 1, 43kDa
-0.00014157107097421834
GO:0032024: positive regulation of insulin secretion ISL1 ISL LIM homeobox 1 8.22923039547
7173e-5
GO:0032024: positive regulation of insulin secretion JAK2 Janus kinase 2 -3.7318581305271725e-
5
GO:0032024: positive regulation of insulin secretion SOX4 SRY (sex determining region Y)-box 4
-4.347638979257247e-5
GO:0032024: positive regulation of insulin secretion TCF7L2 transcription factor 7-like 2 (T-cell
specific, HMG-box) 0.0005478985119265808
GO:0035094: response to nicotine ABAT 4-aminobutyrate aminotransferase 0.00081625575
89217797
GO:0035094: response to nicotine AVP arginine vasopressin -0.0009092874589890046
GO:0035094: response to nicotine BCL2 B-cell CLL/lymphoma 2 -5.086636827417259e-6
GO:0035094: response to nicotine CHRNB2 cholinergic receptor, nicotinic, beta 2 (neuronal)

-0.0007583278132899443
GO:0035094: response to nicotine	DRD2	dopamine receptor D2	-0.00023001114993561662
GO:0035094: response to nicotine	GPX1	glutathione peroxidase 1	0.0003867305456564968
GO:0035094: response to nicotine	HDAC2	histone deacetylase 2	-0.0011885414992245905
GO:0035094: response to nicotine	HMOX1	heme oxygenase (decycling) 1	-0.000210711403920738
4
GO:0035094: response to nicotine	PDX1	pancreatic and duodenal homeobox 1	0.00024693048
72419899
GO:0035094: response to nicotine	STAR	steroidogenic acute regulatory protein	0.00086922330
19067947
GO:0042135: neurotransmitter catabolic process	ABAT	4-aminobutyrate aminotransferase	0.000
8384285830425881
GO:0042135: neurotransmitter catabolic process	ALDH5A1 aldehyde dehydrogenase 5 family, member A1
0.0023452776173856047
GO:0042493: response to drug	ABAT	4-aminobutyrate aminotransferase	0.0008321786943666633
GO:0042493: response to drug	AOC1	amine oxidase, copper containing 1	-0.001420914896464388
7
GO:0042493: response to drug	APOD	apolipoprotein D	0.002623707637651465
GO:0042493: response to drug	ASS1	argininosuccinate synthase 1	0.00014194170651174356
GO:0042493: response to drug	BAK1	BCL2-antagonist/killer 1	-0.001872983622252589
GO:0042493: response to drug	BAX	BCL2-associated X protein	-0.00042458025033647977
GO:0042493: response to drug	BCHE	butyrylcholinesterase	-6.491329207779173e-6
GO:0042493: response to drug	BCL2	B-cell CLL/lymphoma 2	-5.133173754918898e-6
GO:0042493: response to drug	CAV1	caveolin 1, caveolae protein, 22kDa	-0.000534839011063478
GO:0042493: response to drug	CCNB1	cyclin B1	-0.0008812927404760955
GO:0042493: response to drug	CCND1	cyclin D1	-0.002627638068207084
GO:0042493: response to drug	CCNE1	cyclin E1	0.00037022802307575025
GO:0042493: response to drug	CDH3	cadherin 3, type 1, P-cadherin (placental)	-0.0012289660
774525374
GO:0042493: response to drug	COL1A1	collagen, type I, alpha 1	-0.0005247261735010598
GO:0042493: response to drug	CPT1A	carnitine palmitoyltransferase 1A (liver)	0.00127106991
20922074
GO:0042493: response to drug	CREB1	cAMP responsive element binding protein 1	0.00065678352
91725943
GO:0042493: response to drug	CST3	cystatin C	-6.962448601715986e-5
GO:0042493: response to drug	CTNNB1	catenin (cadherin-associated protein), beta 1, 88kDa	-0.00
011684991017535416
GO:0042493: response to drug	CTPS1	CTP synthase 1	-0.0004206897806356906
GO:0042493: response to drug	DRD2	dopamine receptor D2	-0.0002337731272628796
GO:0042493: response to drug	FGF8	fibroblast growth factor 8 (androgen-induced)	0.00098204867
63327395
GO:0042493: response to drug	GABRA4	gamma-aminobutyric acid (GABA) A receptor, alpha 4	-0.00
14818892879008396
GO:0042493: response to drug	GATA3	GATA binding protein 3	-3.884231995012493e-5
GO:0042493: response to drug	GATA4	GATA binding protein 4	-0.0010929060562918144
GO:0042493: response to drug	GATA6	GATA binding protein 6	-2.826935052133846e-5
GO:0042493: response to drug	GCLM	glutamate-cysteine ligase, modifier subunit	-0.0037559422
30585455
GO:0042493: response to drug	GGH	gamma-glutamyl hydrolase (conjugase, folylpolygammaglutamyl h
ydrolase)	-0.0003144384741648851 4
GO:0042493: response to drug	HDAC2	histone deacetylase 2	-0.0012056782378296585
GO:0042493: response to drug	HMGB2	high mobility group box 2	0.00030573904859034417
GO:0042493: response to drug	HTR2C	5-hydroxytryptamine (serotonin) receptor 2C, G protein-couple
d	-0.0005239845147079952
GO:0042493: response to drug	ICAM1	intercellular adhesion molecule 1	0.0007285199041821116
GO:0042493: response to drug	IFNG	interferon, gamma	-4.88814517270455e-5
GO:0042493: response to drug	IGFBP2	insulin-like growth factor binding protein 2, 36kDa	0.000
14438128930480118
GO:0042493: response to drug	IL4	interleukin 4	0.00025944649357830023
GO:0042493: response to drug	INHBA	inhibin, beta A -0.0013498319703095449
GO:0042493: response to drug	ITGA3	integrin, alpha 3 (antigen CD49C, alpha 3 subunit of VLA-3 re
ceptor) 0.0013976669336090667
GO:0042493: response to drug	LOX	lysyl oxidase	-0.0005752529576941598
GO:0042493: response to drug	LRP8	low density lipoprotein receptor-related protein 8, apolipopr
otein e receptor	-0.0009304291800217335
GO:0042493: response to drug	LYN	LYN proto-oncogene, Src family tyrosine kinase	-0.0014181651
65211226
GO:0042493: response to drug	MAS1	MAS1 proto-oncogene, G protein-coupled receptor 0.00029183920
08695353
GO:0042493: response to drug	MCM7	minichromosome maintenance complex component 7	-0.0016415981
644914996
GO:0042493: response to drug	MDK	midkine (neurite growth-promoting factor 2)	0.00166141231
98063554
GO:0042493: response to drug	MGMT	O-6-methylguanine-DNA methyltransferase 0.0004062990333310249
5
GO:0042493: response to drug	MMP7	matrix metallopeptidase 7 (matrilysin, uterine) 0.00042493725
4858662
GO:0042493: response to drug	MYC	v-myc avian myelocytomatosis viral oncogene homolog	-0.00
113478919969235
GO:0042493: response to drug	PDX1	pancreatic and duodenal homeobox 1	0.0002566008581639894
7

GO:0042493: response to drug POR P450 (cytochrome) oxidoreductase 0.0004564298553459014
GO:0042493: response to drug PTCH1 patched 1 -6.604529345629721e-5
GO:0042493: response to drug PTEN phosphatase and tensin homolog 1.753387340373098e-5
GO:0042493: response to drug PTN pleiotrophin 0.00030048029812464937
GO:0042493: response to drug RAD51 RAD51 recombinase -0.0018583682484856418
GO:0042493: response to drug RET ret proto-oncogene -0.0004905437144619261
GO:0042493: response to drug SEMA3C sema domain, immunoglobulin domain (Ig), short basic domain, secreted, (semaphorin) 3C -0.00022098422485490067
GO:0042493: response to drug SFRP1 secreted frizzled-related protein 1 0.0012784513667955754
GO:0042493: response to drug SLC1A3 solute carrier family 1 (glial high affinity glutamate transporter), member 3 -0.003116011982524138
GO:0042493: response to drug SORD sorbitol dehydrogenase -0.0020159685579222656
GO:0042493: response to drug SRD5A2 steroid-5-alpha-reductase, alpha polypeptide 2 (3-oxo-5 alpha-steroid delta 4-dehydrogenase alpha 2) -0.000641781192456762
GO:0042493: response to drug STAR steroidogenic acute regulatory protein 0.0008928183981245767
GO:0042493: response to drug TGFB1 transforming growth factor, beta 1 -7.28094893441974e-5
GO:0042493: response to drug TGFB2 transforming growth factor, beta 2 -0.001057361724904403
GO:0042493: response to drug THBS1 thrombospondin 1 -0.0010341382726313892
GO:0042493: response to drug THRA thyroid hormone receptor, alpha 0.0007620199142163863
GO:0042493: response to drug TP73 tumor protein p73 0.0010261516026832196
GO:0042493: response to drug TXN2 thioredoxin 2 0.0013811834239350373
GO:0042493: response to drug TYMS thymidylate synthetase 0.001558261207687442
GO:0042493: response to drug VEGFC vascular endothelial growth factor C -0.003358609850114736
7
GO:0042493: response to drug VLDLR very low density lipoprotein receptor 0.0009510405671015776
GO:0042493: response to drug XBP1 X-box binding protein 1 0.00026695250118882253
GO:0045471: response to ethanol ABAT 4-aminobutyrate aminotransferase 0.0008299947113849661
GO:0045471: response to ethanol AVP arginine vasopressin -0.0009337376715532953
GO:0045471: response to ethanol BAK1 BCL2-antagonist/killer 1 -0.0018691442589787515
GO:0045471: response to ethanol CCL2 chemokine (C-C motif) ligand 2 0.0008139543961952794
GO:0045471: response to ethanol CCND1 cyclin D1 -0.002622325511988202
GO:0045471: response to ethanol CCNE1 cyclin E1 0.0003698733207064688
GO:0045471: response to ethanol CHRNB2 cholinergic receptor, nicotinic, beta 2 (neuronal) -0.0007754511338242428
GO:0045471: response to ethanol EGR1 early growth response 1 0.0010891728556308685
GO:0045471: response to ethanol GATA3 GATA binding protein 3 -3.929473943456696e-5
GO:0045471: response to ethanol GGH gamma-glutamyl hydrolase (conjugase, folylpolygammaglutamyl hydrolase) -0.00031353561000826123
GO:0045471: response to ethanol ICAM1 intercellular adhesion molecule 1 0.0007273693293157991
GO:0045471: response to ethanol IL4 interleukin 4 0.00025901329791812755
GO:0045471: response to ethanol MGMT O-6-methylguanine-DNA methyltransferase 0.0004056809053482329
7
GO:0045471: response to ethanol PTEN phosphatase and tensin homolog 1.7586515911126354e-5
GO:0045471: response to ethanol RXRA retinoid X receptor, alpha 0.0011109968272105307
GO:0045471: response to ethanol STAR steroidogenic acute regulatory protein 0.0008910161245458171
GO:0045471: response to ethanol TH tyrosine hydroxylase -0.00034120371585519207
GO:0045471: response to ethanol TYMS thymidylate synthetase 0.0015551981636797308
GO:0045776: negative regulation of blood pressure ABAT 4-aminobutyrate aminotransferase 0.0008395157508078796
GO:0045776: negative regulation of blood pressure DRD2 dopamine receptor D2 -0.00023535337223040117
GO:0045776: negative regulation of blood pressure VEGFC vascular endothelial growth factor C -0.003389864116282028
GO:0048148: behavioral response to cocaine ABAT 4-aminobutyrate aminotransferase 0.0008314822091479583
GO:0048148: behavioral response to cocaine DRD2 dopamine receptor D2 -0.00023286538946701402
GO:0048148: behavioral response to cocaine DRD4 dopamine receptor D4 -0.001934202318699832
KEGG:02010: ABC transporters ABCG1 ATP-binding cassette, sub-family G (WHITE), member 1 -0.0023606226924747856
GO:0006355: regulation of transcription, DNA-templated ABCG1 ATP-binding cassette, sub-family G (WHITE), member 1 -0.002341665434571339
GO:0006355: regulation of transcription, DNA-templated ACVRL1 activin A receptor type II-like 1 0.0019461451263920165
GO:0006355: regulation of transcription, DNA-templated APEX1 APEX nuclease (multifunctional DNA repair enzyme) 1 0.00020012066304845745
GO:0006355: regulation of transcription, DNA-templated ATF5 activating transcription factor 5 -0.002738417394466573
GO:0006355: regulation of transcription, DNA-templated BACH1 BTB and CNC homology 1, basic leucine zipper transcription factor 1 0.0004858616370202321
GO:0006355: regulation of transcription, DNA-templated CASP8AP2 caspase 8 associated protein 2 0.0012735027160679787
GO:0006355: regulation of transcription, DNA-templated CEBPB CCAAT/enhancer binding protein (C/EBP), beta -0.0002745449759790138
GO:0006355: regulation of transcription, DNA-templated CELSR2 cadherin, EGF LAG seven-pass G-type receptor 2 -0.0011360981432418868
GO:0006355: regulation of transcription, DNA-templated CHAF1B chromatin assembly factor 1, subunit B (p60) 0.006292511066352086
GO:0006355: regulation of transcription, DNA-templated CHD3 chromodomain helicase DNA binding protein 3 0.00011917125375918149
GO:0006355: regulation of transcription, DNA-templated CLOCK clock circadian regulator 0.000

20035738705752543
GO:0006355: regulation of transcription, DNA-templated DAXX death-domain associated protein 0.000
8926960114535767
GO:0006355: regulation of transcription, DNA-templated DLX6 distal-less homeobox 6 -0.0001800747
707105529
GO:0006355: regulation of transcription, DNA-templated E2F1 E2F transcription factor 1 0.002
100173092073431
GO:0006355: regulation of transcription, DNA-templated ENG endoglin 0.0008277045008275168
GO:0006355: regulation of transcription, DNA-templated ESR1 estrogen receptor 1 -0.0009490519
211270114
GO:0006355: regulation of transcription, DNA-templated EYA3 EYA transcriptional coactivator and p
hosphatase 3 0.0008375143974783711
GO:0006355: regulation of transcription, DNA-templated EZH2 enhancer of zeste 2 polycomb repressi
ve complex 2 subunit -0.0001369075183342161
GO:0006355: regulation of transcription, DNA-templated FOXC1 forkhead box C1 -2.2551636859671754e-
5
GO:0006355: regulation of transcription, DNA-templated FOXM1 forkhead box M1 0.0001971198857297301
GO:0006355: regulation of transcription, DNA-templated FZD7 frizzled class receptor 7 0.001
0715426643732203
GO:0006355: regulation of transcription, DNA-templated GATA4 GATA binding protein 4 -0.0010960469
903292568
GO:0006355: regulation of transcription, DNA-templated GLRX2 glutaredoxin 2 0.0011173084831466258
GO:0006355: regulation of transcription, DNA-templated HIF1A hypoxia inducible factor 1, alpha sub
unit (basic helix-loop-helix transcription factor) -0.0006473802200420288
GO:0006355: regulation of transcription, DNA-templated HINFP histone H4 transcription factor 0.000
9935293433366503
GO:0006355: regulation of transcription, DNA-templated HMGA1 high mobility group AT-hook 1 -0.00
031929834408582215
GO:0006355: regulation of transcription, DNA-templated HMGA2 high mobility group AT-hook 2 0.001
5108935737475357
GO:0006355: regulation of transcription, DNA-templated HMGB3 high mobility group box 3 0.000
46420608836882825
GO:0006355: regulation of transcription, DNA-templated HOXA3 homeobox A3 0.0009898676244079934
GO:0006355: regulation of transcription, DNA-templated HOXB1 homeobox B1 0.0036909602546054683
GO:0006355: regulation of transcription, DNA-templated HOXB13 homeobox B13 0.0018279178412547701
GO:0006355: regulation of transcription, DNA-templated HOXB2 homeobox B2 0.002777830801883462
GO:0006355: regulation of transcription, DNA-templated HOXD13 homeobox D13 -0.000593175132816979
9
GO:0006355: regulation of transcription, DNA-templated INSR insulin receptor -0.0013648946
786685415
GO:0006355: regulation of transcription, DNA-templated JMJD6 jumonji domain containing 6 0.003
595176943406385
GO:0006355: regulation of transcription, DNA-templated KANK1 KN motif and ankyrin repeat domains 1
0.0025362658910803833
GO:0006355: regulation of transcription, DNA-templated KMT2D lysine (K)-specific methyltransferase
2D -0.00019955351860871605
GO:0006355: regulation of transcription, DNA-templated LHX6 LIM homeobox 6 0.0013455375779938107
GO:0006355: regulation of transcription, DNA-templated MEF2C myocyte enhancer factor 2C 0.000
9745956786313344
GO:0006355: regulation of transcription, DNA-templated MTERF1 mitochondrial transcription terminati
on factor 1 0.0002865548486519686
GO:0006355: regulation of transcription, DNA-templated NEUROD4 neuronal differentiation 4 0.001
5087356138340669
GO:0006355: regulation of transcription, DNA-templated NOTCH1 notch 1 0.000519291004703306
GO:0006355: regulation of transcription, DNA-templated NRBF2 nuclear receptor binding factor 2
-0.000952243922012765
GO:0006355: regulation of transcription, DNA-templated PADI4 peptidyl arginine deiminase, type IV
-0.000827641910837679
GO:0006355: regulation of transcription, DNA-templated PITX2 paired-like homeodomain 2 0.002
178824277367717
GO:0006355: regulation of transcription, DNA-templated PML promyelocytic leukemia -0.0006808680
696586666
GO:0006355: regulation of transcription, DNA-templated POU1F1 POU class 1 homeobox 1 0.00013762929
217908912
GO:0006355: regulation of transcription, DNA-templated PRDM4 PR domain containing 4 0.00084140848
90226066
GO:0006355: regulation of transcription, DNA-templated PTTG1 pituitary tumor-transforming 1 -0.00
036232234391625565
GO:0006355: regulation of transcription, DNA-templated PTTG2 pituitary tumor-transforming 2 -0.00
17268107379317717
GO:0006355: regulation of transcription, DNA-templated PTTG3P pituitary tumor-transforming 3, pseud
ogene -4.0391214768816385e-5
GO:0006355: regulation of transcription, DNA-templated RBBP4 retinoblastoma binding protein 4
-0.0018629889162687448
GO:0006355: regulation of transcription, DNA-templated RFC1 replication factor C (activator 1) 1,
145kDa 0.0010605418258528654
GO:0006355: regulation of transcription, DNA-templated RREB1 ras responsive element binding protei
n 1 -0.0019840710875836205
GO:0006355: regulation of transcription, DNA-templated SIX1 SIX homeobox 1 -0.001872273300629345
2
GO:0006355: regulation of transcription, DNA-templated SOX4 SRY (sex determining region Y)-box 4

-3.84749163818025e-5
GO:0006355: regulation of transcription, DNA-templated  TBPL1   TBP-like 1      -0.000789896881277378
4
GO:0006355: regulation of transcription, DNA-templated  TBX21   T-box 21        0.0003990177683590804
GO:0006355: regulation of transcription, DNA-templated  TGFBR1  transforming growth factor, beta rece
ptor 1  0.00034029091337871547
GO:0006355: regulation of transcription, DNA-templated  TP53    tumor protein p53       0.00117711530
1619081
GO:0006355: regulation of transcription, DNA-templated  TULP3   tubby like protein 3    0.00095745576
78974643
GO:0006355: regulation of transcription, DNA-templated  TXLNG   taxilin gamma   -0.000838522066702498
8
GO:0006355: regulation of transcription, DNA-templated  WT1     Wilms tumor 1   -0.000507551555832778
1
GO:0006355: regulation of transcription, DNA-templated  WWTR1   WW domain containing transcription re
gulator 1       0.0009013744323481764
GO:0006355: regulation of transcription, DNA-templated  YBX1    Y box binding protein 1 -0.0009151955
467048586
GO:0006355: regulation of transcription, DNA-templated  ZFP36L2 ZFP36 ring finger protein-like 2
-0.0003633828668500567
GO:0006355: regulation of transcription, DNA-templated  ZNF205  zinc finger protein 205 0.00059682523
10036018
GO:0006355: regulation of transcription, DNA-templated  ZNF207  zinc finger protein 207 -0.0003898340
1064739966
GO:0006355: regulation of transcription, DNA-templated  ZNF266  zinc finger protein 266 0.00041304641
85346677
GO:0006355: regulation of transcription, DNA-templated  ZNF335  zinc finger protein 335 -0.0003652254
1044257413
GO:0006355: regulation of transcription, DNA-templated  ZNF442  zinc finger protein 442 0.00105958638
9757607
GO:0008203: cholesterol metabolic process       ABCG1   ATP-binding cassette, sub-family G (WHITE), m
ember 1 -0.0023454878101763924
GO:0008203: cholesterol metabolic process       CYP7B1  cytochrome P450, family 7, subfamily B, polyp
eptide 1        0.00016361504466810514
GO:0008203: cholesterol metabolic process       IL4     interleukin 4   0.0002596518069478495
GO:0008203: cholesterol metabolic process       LEP     leptin  0.0032021505124831783
GO:0008203: cholesterol metabolic process       LRP5    low density lipoprotein receptor-related prot
ein 5   2.981304340339888e-5
GO:0008203: cholesterol metabolic process       RXRA    retinoid X receptor, alpha      0.00111843405
90287421
GO:0008203: cholesterol metabolic process       SOAT1   sterol O-acyltransferase 1      -0.0004083545
783319706
GO:0008203: cholesterol metabolic process       STAR    steroidogenic acute regulatory protein  0.000
8976499727709577
GO:0008203: cholesterol metabolic process       VLDLR   very low density lipoprotein receptor   0.000
9564620301798078
GO:0009720: detection of hormone stimulus       ABCG1   ATP-binding cassette, sub-family G (WHITE), m
ember 1 -0.0023606226924747856
GO:0010033: response to organic substance       ABCG1   ATP-binding cassette, sub-family G (WHITE), m
ember 1 -0.0023330672278115442
GO:0010033: response to organic substance       CREB1   cAMP responsive element binding protein 1
0.0006548028848150213
GO:0010033: response to organic substance       CRIP1   cysteine-rich protein 1 (intestinal)    0.001
7204364101309866
GO:0010033: response to organic substance       GLRX2   glutaredoxin 2  0.0011134929047623772
GO:0010033: response to organic substance       MAPT    microtubule-associated protein tau      0.001
5495420225335116
GO:0010033: response to organic substance       SERPINF2        serpin peptidase inhibitor, clade F
(alpha-2 antiplasmin, pigment epithelium derived factor), member 2      0.0006573269261983929
GO:0010033: response to organic substance       WNT5A   wingless-type MMTV integration site family, m
ember 5A        -0.0006650579785260216
GO:0010745: negative regulation of macrophage derived foam cell differentiation ABCG1   ATP-binding c
assette, sub-family G (WHITE), member 1 -0.002346638657309913
GO:0010745: negative regulation of macrophage derived foam cell differentiation NR1H3   nuclear recep
tor subfamily 1, group H, member 3      0.0008032537504780684
GO:0010872: regulation of cholesterol esterification    ABCG1   ATP-binding cassette, sub-family G (W
HITE), member 1 -0.0023606226924747856
GO:0010875: positive regulation of cholesterol efflux   ABCG1   ATP-binding cassette, sub-family G (W
HITE), member 1 -0.0023262710398407453
GO:0010875: positive regulation of cholesterol efflux   NR1H3   nuclear receptor subfamily 1, group
H, member 3     0.0007981704538679059
GO:0010875: positive regulation of cholesterol efflux   PTCH1   patched 1       -6.733786411850896e-5
GO:0010875: positive regulation of cholesterol efflux   SIRT1   sirtuin 1       -3.108596316558697e-6
GO:0010887: negative regulation of cholesterol storage  ABCG1   ATP-binding cassette, sub-family G (W
HITE), member 1 -0.002346638657309913
GO:0010887: negative regulation of cholesterol storage  NR1H3   nuclear receptor subfamily 1, group
H, member 3     0.0008032537504780684
GO:0032367: intracellular cholesterol transport ABCG1   ATP-binding cassette, sub-family G (WHITE), m
ember 1 -0.0023414183670599836
GO:0032367: intracellular cholesterol transport STAR    steroidogenic acute regulatory protein  0.000
8955133587820494

GO:0032367: intracellular cholesterol transport VPS4A   vacuolar protein sorting 4 homolog A (S. cerevisiae) -0.0006857834708294203
GO:0033344: cholesterol efflux  ABCG1   ATP-binding cassette, sub-family G (WHITE), member 1    -0.0023351541238514502
GO:0033344: cholesterol efflux  CAV1    caveolin 1, caveolae protein, 22kDa     -0.000535137360978536 8
GO:0033344: cholesterol efflux  SOAT1   sterol O-acyltransferase 1      -0.000408159976313773
GO:0033700: phospholipid efflux ABCG1   ATP-binding cassette, sub-family G (WHITE), member 1    -0.0023606226924747856
GO:0033993: response to lipid   ABCG1   ATP-binding cassette, sub-family G (WHITE), member 1    -0.002348053988443847
GO:0033993: response to lipid   GATA2   GATA binding protein 2  -0.0004546861240076834
GO:0033993: response to lipid   PCNA    proliferating cell nuclear antigen      0.0012333088339832432
GO:0034374: low-density lipoprotein particle remodeling ABCG1   ATP-binding cassette, sub-family G (WHITE), member 1 -0.0023477349653539255
GO:0034374: low-density lipoprotein particle remodeling AGT     angiotensinogen (serpin peptidase inhibitor, clade A, member 8)      -0.0011404141840943827
GO:0034374: low-density lipoprotein particle remodeling AGTR1   angiotensin II receptor, type 1 0.0003465689823358322
GO:0034375: high-density lipoprotein particle remodeling        ABCG1   ATP-binding cassette, sub-family G (WHITE), member 1 -0.0023606226924747856
GO:0034436: glycoprotein transport      ABCG1   ATP-binding cassette, sub-family G (WHITE), member 1 -0.002360455519220915
GO:0034436: glycoprotein transport      GUK1    guanylate kinase 1      0.0033018436125547704
GO:0034436: glycoprotein transport      VLDLR   very low density lipoprotein receptor   0.0009646216533888689
GO:0042157: lipoprotein metabolic process       ABCG1   ATP-binding cassette, sub-family G (WHITE), member 1 -0.0023606226924747856
GO:0042632: cholesterol homeostasis     ABCG1   ATP-binding cassette, sub-family G (WHITE), member 1 -0.002333236756042257
GO:0042632: cholesterol homeostasis     AKR1C1  aldo-keto reductase family 1, member C1 -0.001486890293281473
GO:0042632: cholesterol homeostasis     CAV1    caveolin 1, caveolae protein, 22kDa     -0.0005350171271438004
GO:0042632: cholesterol homeostasis     CD24    CD24 molecule   0.0010673039896267873
GO:0042632: cholesterol homeostasis     HPN     hepsin  0.0031532344758925983
GO:0042632: cholesterol homeostasis     LRP5    low density lipoprotein receptor-related protein 5 3.040272412830946e-5
GO:0042632: cholesterol homeostasis     NR1H3   nuclear receptor subfamily 1, group H, member 3 0.000800087123189823 1
GO:0042632: cholesterol homeostasis     SIRT1   sirtuin 1       -2.70654799823969e-6
GO:0042632: cholesterol homeostasis     SOAT1   sterol O-acyltransferase 1      -0.000408650022457234 5
GO:0042632: cholesterol homeostasis     XBP1    X-box binding protein 1 0.0002685974596927559
GO:0042987: amyloid precursor protein catabolic process ABCG1   ATP-binding cassette, sub-family G (WHITE), member 1 -0.002356926099937954
GO:0042987: amyloid precursor protein catabolic process DHCR24  24-dehydrocholesterol reductase -0.0017218261808735027
GO:0043691: reverse cholesterol transport       ABCG1   ATP-binding cassette, sub-family G (WHITE), member 1 -0.0023606226924747856
GO:0044281: small molecule metabolic process    ABCG1   ATP-binding cassette, sub-family G (WHITE), member 1 -0.002337300618454
GO:0044281: small molecule metabolic process    ACADL   acyl-CoA dehydrogenase, long chain      0.0009806184984740072
GO:0044281: small molecule metabolic process    ACOT8   acyl-CoA thioesterase 8 0.0011354676229892832
GO:0044281: small molecule metabolic process    AGT     angiotensinogen (serpin peptidase inhibitor, clade A, member 8)      -0.0011316057979557492
GO:0044281: small molecule metabolic process    AKT1    v-akt murine thymoma viral oncogene homolog 1 0.0007299030112991718
GO:0044281: small molecule metabolic process    ALOX12  arachidonate 12-lipoxygenase    -0.0019519243432385442
GO:0044281: small molecule metabolic process    AMD1    adenosylmethionine decarboxylase 1      -0.0025271199410123505
GO:0044281: small molecule metabolic process    ASS1    argininosuccinate synthase 1    0.00014210037478324923
GO:0044281: small molecule metabolic process    AUH     AU RNA binding protein/enoyl-CoA hydratase -0.001460609543166981 7
GO:0044281: small molecule metabolic process    AZIN1   antizyme inhibitor 1    -0.00012711882989741352
GO:0044281: small molecule metabolic process    BDH1    3-hydroxybutyrate dehydrogenase, type 1 -0.00020721964152094316
GO:0044281: small molecule metabolic process    CAV1    caveolin 1, caveolae protein, 22kDa     -0.0005352462335167768
GO:0044281: small molecule metabolic process    CBS     cystathionine-beta-synthase     0.0015155290105719084
GO:0044281: small molecule metabolic process    CHST15  carbohydrate (N-acetylgalactosamine 4-sulfate 6-O) sulfotransferase 15        -0.0008251421276839753
GO:0044281: small molecule metabolic process    CPOX    coproporphyrinogen oxidase      0.001271756717888677
GO:0044281: small molecule metabolic process    CPT1A   carnitine palmitoyltransferase 1A (liver) 0.001272076062170097 2

GO:0044281: small molecule metabolic process    CROT    carnitine O-octanoyltransferase -0.0004380009443825059
GO:0044281: small molecule metabolic process    CTH     cystathionine gamma-lyase       0.001746934332131044
GO:0044281: small molecule metabolic process    CTPS1   CTP synthase 1  -0.00042104465046334814
GO:0044281: small molecule metabolic process    CTPS2   CTP synthase 2  0.0017590352351665843
GO:0044281: small molecule metabolic process    CYP1B1  cytochrome P450, family 1, subfamily B, polypeptide 1        0.00039755471279155895
GO:0044281: small molecule metabolic process    CYP27B1 cytochrome P450, family 27, subfamily B, polypeptide 1       -0.00047043275777649575
GO:0044281: small molecule metabolic process    CYP4B1  cytochrome P450, family 4, subfamily B, polypeptide 1        -0.0013549255428817464
GO:0044281: small molecule metabolic process    CYP7B1  cytochrome P450, family 7, subfamily B, polypeptide 1        0.0001631972293224399
GO:0044281: small molecule metabolic process    DCT     dopachrome tautomerase  -0.002351662469383147
GO:0044281: small molecule metabolic process    DHCR24  24-dehydrocholesterol reductase -0.001706145747730389
GO:0044281: small molecule metabolic process    DHFR    dihydrofolate reductase -0.00030787753765544444
GO:0044281: small molecule metabolic process    DTYMK   deoxythymidylate kinase (thymidylate kinase)    0.003479384793586334
GO:0044281: small molecule metabolic process    ELOVL2  ELOVL fatty acid elongase 2     0.0005244086502352605
GO:0044281: small molecule metabolic process    ELOVL5  ELOVL fatty acid elongase 5     0.00020678983391271213
GO:0044281: small molecule metabolic process    GCDH    glutaryl-CoA dehydrogenase      -4.876941985395099e-6
GO:0044281: small molecule metabolic process    GCHFR   GTP cyclohydrolase I feedback regulator -0.0008418507162816586
GO:0044281: small molecule metabolic process    GCLM    glutamate-cysteine ligase, modifier subunit     -0.0037590667334135735
GO:0044281: small molecule metabolic process    GMPS    guanine monphosphate synthase   0.0017725539984257551
GO:0044281: small molecule metabolic process    GPD1L   glycerol-3-phosphate dehydrogenase 1-like       0.0005669395518063697
GO:0044281: small molecule metabolic process    GPX1    glutathione peroxidase 1        0.0003877079633567316
GO:0044281: small molecule metabolic process    GSTM1   glutathione S-transferase mu 1  -0.001473001020169157
GO:0044281: small molecule metabolic process    GSTM2   glutathione S-transferase mu 2 (muscle) -0.0019128202855439643
GO:0044281: small molecule metabolic process    GSTM3   glutathione S-transferase mu 3 (brain)  -0.0012880251258405966
GO:0044281: small molecule metabolic process    GSTP1   glutathione S-transferase pi 1  0.00023126604079261778
GO:0044281: small molecule metabolic process    GUK1    guanylate kinase 1      0.003271782259601756
GO:0044281: small molecule metabolic process    HMGCL   3-hydroxymethyl-3-methylglutaryl-CoA lyase      -0.0010627053518895476
GO:0044281: small molecule metabolic process    HMOX1   heme oxygenase (decycling) 1    -0.00021731471238639431
GO:0044281: small molecule metabolic process    HPRT1   hypoxanthine phosphoribosyltransferase 1        -0.0011850174365291718
GO:0044281: small molecule metabolic process    HS2ST1  heparan sulfate 2-O-sulfotransferase 1  -0.0008098221661686961
GO:0044281: small molecule metabolic process    HSD17B4 hydroxysteroid (17-beta) dehydrogenase 4        -0.001161670340889208
GO:0044281: small molecule metabolic process    IDO1    indoleamine 2,3-dioxygenase 1   0.0012247051510665397
GO:0044281: small molecule metabolic process    IMPDH1  IMP (inosine 5'-monophosphate) dehydrogenase 1       -0.001870551937633798
GO:0044281: small molecule metabolic process    IMPDH2  IMP (inosine 5'-monophosphate) dehydrogenase 2       -0.000347136738216014
GO:0044281: small molecule metabolic process    INPP4B  inositol polyphosphate-4-phosphatase, type II, 105kDa       0.0002900299912188719
GO:0044281: small molecule metabolic process    ITPR1   inositol 1,4,5-trisphosphate receptor, type 1   -0.000779569355912296
GO:0044281: small molecule metabolic process    IVD     isovaleryl-CoA dehydrogenase    -0.0009353098630946368
GO:0044281: small molecule metabolic process    KPNB1   karyopherin (importin) beta 1   0.0007714731747169501
GO:0044281: small molecule metabolic process    MCCC1   methylcrotonoyl-CoA carboxylase 1 (alpha)       8.376352857033848e-5
GO:0044281: small molecule metabolic process    MCCC2   methylcrotonoyl-CoA carboxylase 2 (beta)        -0.0007843290744811763
GO:0044281: small molecule metabolic process    MED1    mediator complex subunit 1      0.0011280056167164044
GO:0044281: small molecule metabolic process    MGAM    maltase-glucoamylase (alpha-glucosidase)        -0.0013076289398003885
GO:0044281: small molecule metabolic process    MTAP    methylthioadenosine phosphorylase       0.0026559830175411385
GO:0044281: small molecule metabolic process    MTHFD1  methylenetetrahydrofolate dehydrogenase (NADP

+ dependent) 1, methenyltetrahydrofolate cyclohydrolase, formyltetrahydrofolate synthetase -0.00
06638600248168461
GO:0044281: small molecule metabolic process MTMR2 myotubularin related protein 2 -0.0004608399
973891542
GO:0044281: small molecule metabolic process MUT methylmalonyl CoA mutase 0.00275690303
28873346
GO:0044281: small molecule metabolic process NUDT1 nudix (nucleoside diphosphate linked moiety
X)-type motif 1 0.001835693327776657
GO:0044281: small molecule metabolic process NUP153 nucleoporin 153kDa 0.0008242990270838397
GO:0044281: small molecule metabolic process ODC1 ornithine decarboxylase 1 0.00082453386
45328432
GO:0044281: small molecule metabolic process OGDH oxoglutarate (alpha-ketoglutarate) dehydrogen
ase (lipoamide) -0.0002944944326599106
GO:0044281: small molecule metabolic process PDK1 pyruvate dehydrogenase kinase, isozyme 1
0.0014229685868522122
GO:0044281: small molecule metabolic process PDK2 pyruvate dehydrogenase kinase, isozyme 2
0.0014785758412337597
GO:0044281: small molecule metabolic process PDK3 pyruvate dehydrogenase kinase, isozyme 3
0.0007714443325769016
GO:0044281: small molecule metabolic process PEX11A peroxisomal biogenesis factor 11 alpha 0.000
7169087210258609
GO:0044281: small molecule metabolic process PGLS 6-phosphogluconolactonase 0.00174889045
97298008
GO:0044281: small molecule metabolic process PHGDH phosphoglycerate dehydrogenase 0.00029190524
001686805
GO:0044281: small molecule metabolic process PIK3CD phosphatidylinositol-4,5-bisphosphate 3-kinas
e, catalytic subunit delta -0.0007801866571449219
GO:0044281: small molecule metabolic process PLA2G5 phospholipase A2, group V 0.00207747489
26489033
GO:0044281: small molecule metabolic process PLCB1 phospholipase C, beta 1 (phosphoinositide-spe
cific) 0.00015338744229837411
GO:0044281: small molecule metabolic process PNPO pyridoxamine 5'-phosphate oxidase -0.00
10903692355139567
GO:0044281: small molecule metabolic process POLD1 polymerase (DNA directed), delta 1, catalytic
subunit -0.00044472844066504897
GO:0044281: small molecule metabolic process PRKCA protein kinase C, alpha -5.8668997382740645e-
6
GO:0044281: small molecule metabolic process PSAT1 phosphoserine aminotransferase 1 -0.00
072246158126063
GO:0044281: small molecule metabolic process PSMA5 proteasome (prosome, macropain) subunit, alph
a type, 5 0.0003316121703645836
GO:0044281: small molecule metabolic process PSMD11 proteasome (prosome, macropain) 26S subunit,
non-ATPase, 11 -0.0010944734748804521
GO:0044281: small molecule metabolic process PSMD13 proteasome (prosome, macropain) 26S subunit,
non-ATPase, 13 -5.988737176031255e-5
GO:0044281: small molecule metabolic process PSPH phosphoserine phosphatase -0.0019339171
391770402
GO:0044281: small molecule metabolic process PTEN phosphatase and tensin homolog 1.74955099864
6266e-5
GO:0044281: small molecule metabolic process QDPR quinoid dihydropteridine reductase -0.00
28537157178306167
GO:0044281: small molecule metabolic process RRM1 ribonucleotide reductase M1 0.00190797770
13236183
GO:0044281: small molecule metabolic process RRM2 ribonucleotide reductase M2 -0.0005665510
292744959
GO:0044281: small molecule metabolic process RXRA retinoid X receptor, alpha 0.00111426772
89570054
GO:0044281: small molecule metabolic process SEH1L SEH1-like (S. cerevisiae) -0.0007321582
541877546
GO:0044281: small molecule metabolic process SLC25A37 solute carrier family 25 (mitochondri
al iron transporter), member 37 -0.0002787947864612839
GO:0044281: small molecule metabolic process SLC2A5 solute carrier family 2 (facilitated glucose/
fructose transporter), member 5 0.0002311387938059672
GO:0044281: small molecule metabolic process SLC35D1 solute carrier family 35 (UDP-GlcA/UDP-GalNAc
transporter), member D1 -6.698629543688238e-5
GO:0044281: small molecule metabolic process SLC44A4 solute carrier family 44, member 4 0.002
8523177523958588
GO:0044281: small molecule metabolic process SLC6A8 solute carrier family 6 (neurotransmitter tra
nsporter), member 8 -0.0010431893460143564
GO:0044281: small molecule metabolic process SMARCD3 SWI/SNF related, matrix associated, actin dep
endent regulator of chromatin, subfamily d, member 3 0.00034516995628864586
GO:0044281: small molecule metabolic process SORD sorbitol dehydrogenase -0.002017714520828913
3
GO:0044281: small molecule metabolic process SPHK1 sphingosine kinase 1 0.0018094047069558865
GO:0044281: small molecule metabolic process SPR sepiapterin reductase (7,8-dihydrobiopterin:N
ADP+ oxidoreductase) -0.0012311909799859445
GO:0044281: small molecule metabolic process SRD5A2 steroid-5-alpha-reductase, alpha polypeptide
2 (3-oxo-5 alpha-steroid delta 4-dehydrogenase alpha 2) -0.0006423794498907138
GO:0044281: small molecule metabolic process STAR steroidogenic acute regulatory protein 0.000
8936042515129589
GO:0044281: small molecule metabolic process SULT4A1 sulfotransferase family 4A, member 1 -0.00

11680790625527707
GO:0044281: small molecule metabolic process    TAT     tyrosine aminotransferase       0.00105200948
38986243
GO:0044281: small molecule metabolic process    TH      tyrosine hydroxylase    -0.00034198743623431
4
GO:0044281: small molecule metabolic process    TNFRSF21        tumor necrosis factor receptor superf
amily, member 21        0.00034818714596347803
GO:0044281: small molecule metabolic process    TPR     translocated promoter region, nuclear basket
protein -0.0006965563073315005
GO:0044281: small molecule metabolic process    TYMS    thymidylate synthetase  0.001559664256672852
GO:0044281: small molecule metabolic process    UGCG    UDP-glucose ceramide glucosyltransferase
0.00036898840251944475
GO:0044281: small molecule metabolic process    UGDH    UDP-glucose 6-dehydrogenase     0.00050179489
44982197
GO:0044281: small molecule metabolic process    UGP2    UDP-glucose pyrophosphorylase 2 0.00078446674
07606125
GO:0045542: positive regulation of cholesterol biosynthetic process     ABCG1   ATP-binding cassette,
sub-family G (WHITE), member 1  -0.0023467586157089023
GO:0045542: positive regulation of cholesterol biosynthetic process     POR     P450 (cytochrome) oxi
doreductase     0.00045894675342146276
GO:0055085: transmembrane transport     ABCG1   ATP-binding cassette, sub-family G (WHITE), member 1
-0.002337396313274773б
GO:0055085: transmembrane transport     ASIC2   acid-sensing (proton-gated) ion channel 2       0.002
2784198591722492
GO:0055085: transmembrane transport     ATP1B3  ATPase, Na+/K+ transporting, beta 3 polypeptide -0.00
1497232630867033
GO:0055085: transmembrane transport     AVP     arginine vasopressin    -0.0009360706646363328
GO:0055085: transmembrane transport     BCL2    B-cell CLL/lymphoma 2   -4.823789652170313e-6
GO:0055085: transmembrane transport     CLCN2   chloride channel, voltage-sensitive 2   0.00105915612
8788751
GO:0055085: transmembrane transport     CP      ceruloplasmin (ferroxidase)     0.0013070700736619446
GO:0055085: transmembrane transport     GABRA4  gamma-aminobutyric acid (GABA) A receptor, alpha 4
-0.001483247230278327
GO:0055085: transmembrane transport     HMOX1   heme oxygenase (decycling) 1    -0.00021720973432194
716
GO:0055085: transmembrane transport     MRS2    MRS2 magnesium transporter      0.0006567646168481227
GO:0055085: transmembrane transport     NUP153  nucleoporin 153kDa      0.0008235870970852998
GO:0055085: transmembrane transport     RAF1    Raf-1 proto-oncogene, serine/threonine kinase   0.001
497762687662902
GO:0055085: transmembrane transport     SEH1L   SEH1-like (S. cerevisiae)       -0.000732851748420286
5
GO:0055085: transmembrane transport     SLC11A2 solute carrier family 11 (proton-coupled divalent met
al ion transporter), member 2   -0.0006650349956668877
GO:0055085: transmembrane transport     SLC15A1 solute carrier family 15 (oligopeptide transporter),
member 1        -0.0027585655642272514
GO:0055085: transmembrane transport     SLC15A2 solute carrier family 15 (oligopeptide transporter),
member 2        -0.00012308663479612342
GO:0055085: transmembrane transport     SLC17A3 solute carrier family 17 (organic anion transporter),
member 3        -0.0008661666739755719
GO:0055085: transmembrane transport     SLC1A1  solute carrier family 1 (neuronal/epithelial high aff
inity glutamate transporter, system Xag), member 1      -0.0009006843346179444
GO:0055085: transmembrane transport     SLC1A3  solute carrier family 1 (glial high affinity glutamat
e transporter), member 3        -0.0031193849803142835
GO:0055085: transmembrane transport     SLC1A5  solute carrier family 1 (neutral amino acid transport
er), member 5   0.0015350964550970095
GO:0055085: transmembrane transport     SLC22A18        solute carrier family 22, member 18     0.000
7342816565890956
GO:0055085: transmembrane transport     SLC22A4 solute carrier family 22 (organic cation/zwitterion t
ransporter), member 4   -0.00047978319345127145
GO:0055085: transmembrane transport     SLC22A5 solute carrier family 22 (organic cation/carnitine tr
ansporter), member 5    -0.0018523046292869614
GO:0055085: transmembrane transport     SLC25A44        solute carrier family 25, member 44     0.000
3453198541108319
GO:0055085: transmembrane transport     SLC2A5  solute carrier family 2 (facilitated glucose/fructose
transporter), member 5  0.0002310158243279614
GO:0055085: transmembrane transport     SLC31A1 solute carrier family 31 (copper transporter), member
1       -2.652141123881126З e-5
GO:0055085: transmembrane transport     SLC35D1 solute carrier family 35 (UDP-GlcA/UDP-GalNAc transpo
rter), member D1        -6.69553996832071e-5
GO:0055085: transmembrane transport     SLC39A14        solute carrier family 39 (zinc transporter),
member 14       0.00019671081815350637
GO:0055085: transmembrane transport     SLC39A6 solute carrier family 39 (zinc transporter), member 6
0.00030552089901295527
GO:0055085: transmembrane transport     SLC44A4 solute carrier family 44, member 4      0.00285394415
53379406
GO:0055085: transmembrane transport     SLC4A7  solute carrier family 4, sodium bicarbonate cotranspo
rter, member 7  0.0008821552977918825
GO:0055085: transmembrane transport     SLC5A7  solute carrier family 5 (sodium/choline cotransporte
r), member 7    0.002010652153209131
GO:0055085: transmembrane transport     SLC7A5  solute carrier family 7 (amino acid transporter light
chain, L system), member 5      -0.002573428423152061

GO:0055085: transmembrane transport SLC7A7 solute carrier family 7 (amino acid transporter light chain, y+L system), member 7 -0.0005985999534865752
GO:0055085: transmembrane transport SLC7A8 solute carrier family 7 (amino acid transporter light chain, L system), member 8 -0.00046902513071875264
GO:0055085: transmembrane transport STEAP3 STEAP family member 3, metalloreductase 0.002607597853962525
GO:0055085: transmembrane transport TFRC transferrin receptor 0.0010839266451801012
GO:0055085: transmembrane transport TPR translocated promoter region, nuclear basket protein -0.0006968639987634029
GO:0055085: transmembrane transport TTYH1 tweety family member 1 0.0008688345226562565
GO:0055091: phospholipid homeostasis ABCG1 ATP-binding cassette, sub-family G (WHITE), member 1 -0.0023606226924747856
GO:0055099: response to high density lipoprotein particle ABCG1 ATP-binding cassette, sub-family G (WHITE), member 1 -0.0023606226924747856
GO:1901998: toxin transport ABCG1 ATP-binding cassette, sub-family G (WHITE), member 1 -0.002346795806007113
GO:1901998: toxin transport BNIP3 BCL2/adenovirus E1B 19kDa interacting protein 3 0.0029232958957215597
GO:1901998: toxin transport DNM1 dynamin 1 -0.0013148182842815034
GO:1901998: toxin transport LRP6 low density lipoprotein receptor-related protein 6 0.00015035712009481194
GO:1901998: toxin transport NRP1 neuropilin 1 -0.0006458480958389629
GO:1901998: toxin transport SLC17A3 solute carrier family 17 (organic anion transporter), member 3 -0.0008704568727983809
GO:1901998: toxin transport SLC7A8 solute carrier family 7 (amino acid transporter light chain, L system), member 8 -0.0004731488263616594
GO:0000447: endonucleolytic cleavage in ITS1 to separate SSU-rRNA from 5.8S rRNA and LSU-rRNA from tricistronic rRNA transcript (SSU-rRNA, 5.8S rRNA, LSU-rRNA) ABT1 activator of basal transcription 1 0.002580882811878861
GO:0000472: endonucleolytic cleavage to generate mature 5'-end of SSU-rRNA from (SSU-rRNA, 5.8S rRNA, LSU-rRNA) ABT1 activator of basal transcription 1 0.002580882811878861
GO:0000480: endonucleolytic cleavage in 5'-ETS of tricistronic rRNA transcript (SSU-rRNA, 5.8S rRNA, LSU-rRNA) ABT1 activator of basal transcription 1 0.002580882811878861
GO:0006357: regulation of transcription from RNA polymerase II promoter ABT1 activator of basal transcription 1 0.002548599425667888
GO:0006357: regulation of transcription from RNA polymerase II promoter ATF5 activating transcription factor 5 -0.0027426922287843605
GO:0006357: regulation of transcription from RNA polymerase II promoter BATF basic leucine zipper transcription factor, ATF-like 0.002042345561713186
GO:0006357: regulation of transcription from RNA polymerase II promoter BBS7 Bardet-Biedl syndrome 7 -0.0009800454774115737
GO:0006357: regulation of transcription from RNA polymerase II promoter CHD3 chromodomain helicase DNA binding protein 3 0.00011870809770782367
GO:0006357: regulation of transcription from RNA polymerase II promoter CITED1 Cbp/p300-interacting transactivator, with Glu/Asp-rich carboxy-terminal domain, 1 0.0028266656620635072
GO:0006357: regulation of transcription from RNA polymerase II promoter CLOCK clock circadian regulator 0.000199564512483896
GO:0006357: regulation of transcription from RNA polymerase II promoter DEK DEK proto-oncogene 0.002496676092612851
GO:0006357: regulation of transcription from RNA polymerase II promoter ECM1 extracellular matrix protein 1 -0.001563430528541794
GO:0006357: regulation of transcription from RNA polymerase II promoter ELF5 E74-like factor 5 (ets domain transcription factor) 0.0011776085549585966
GO:0006357: regulation of transcription from RNA polymerase II promoter FOXA2 forkhead box A2 -1.6311711549185648e-5
GO:0006357: regulation of transcription from RNA polymerase II promoter FOXC2 forkhead box C2 (MFH-1, mesenchyme forkhead 1) 0.0017528806946937778
GO:0006357: regulation of transcription from RNA polymerase II promoter FOXE3 forkhead box E3 0.0010535980170645683
GO:0006357: regulation of transcription from RNA polymerase II promoter FOXO3 forkhead box O3 0.00120390467137831
GO:0006357: regulation of transcription from RNA polymerase II promoter GRHL2 grainyhead-like 2 (Drosophila) 0.0010091329198541842
GO:0006357: regulation of transcription from RNA polymerase II promoter HMGB1 high mobility group box 1 -0.0007775343098847412
GO:0006357: regulation of transcription from RNA polymerase II promoter HMGB2 high mobility group box 2 0.00031056276397000343
GO:0006357: regulation of transcription from RNA polymerase II promoter INHBA inhibin, beta A -0.0013592777854372052
GO:0006357: regulation of transcription from RNA polymerase II promoter PITX2 paired-like homeodomain 2 0.002182533756769119
GO:0006357: regulation of transcription from RNA polymerase II promoter PKN1 protein kinase N1 -0.0019083906098755527
GO:0006357: regulation of transcription from RNA polymerase II promoter PURA purine-rich element binding protein A -0.00017148223536069093
GO:0006357: regulation of transcription from RNA polymerase II promoter RAD21 RAD21 homolog (S. pombe) -0.00010205743301588334
GO:0006357: regulation of transcription from RNA polymerase II promoter SMARCD3 SWI/SNF related, matrix associated, actin dependent regulator of chromatin, subfamily d, member 3 0.000346531271215309
GO:0006357: regulation of transcription from RNA polymerase II promoter SOX10 SRY (sex determining region Y)-box 10 0.00019354372544261634

GO:0006357: regulation of transcription from RNA polymerase II promoter STAT5A  signal transducer and activator of transcription 5A   0.0016005449419091407
GO:0006357: regulation of transcription from RNA polymerase II promoter TBX3    T-box 3 0.0012283722603860594
GO:0006357: regulation of transcription from RNA polymerase II promoter TCF15   transcription factor 15 (basic helix-loop-helix)    -0.0026613922801142303
GO:0006357: regulation of transcription from RNA polymerase II promoter TCF7L2  transcription factor 7-like 2 (T-cell specific, HMG-box)    0.0005761274762007429
GO:0006357: regulation of transcription from RNA polymerase II promoter TFAP2C  transcription factor AP-2 gamma (activating enhancer binding protein 2 gamma)       0.0010656573993014765
GO:0006357: regulation of transcription from RNA polymerase II promoter TFCP2L1 transcription factor CP2-like 1     0.00045058761855385263
GO:0006357: regulation of transcription from RNA polymerase II promoter TFDP1   transcription factor Dp-1   0.0014508550242735452
GO:0006357: regulation of transcription from RNA polymerase II promoter THRA    thyroid hormone receptor, alpha      0.0007651182613818636
GO:0006357: regulation of transcription from RNA polymerase II promoter VEGFA   vascular endothelial growth factor A        0.0005979345192633115
GO:0006357: regulation of transcription from RNA polymerase II promoter WDR77   WD repeat domain 77 0.00013432231809843394
GO:0006357: regulation of transcription from RNA polymerase II promoter WT1     Wilms tumor 1   -0.0005088512230075562
GO:0006357: regulation of transcription from RNA polymerase II promoter YY1     YY1 transcription factor     0.0013448430086207988
GO:0006366: transcription from RNA polymerase II promoter       ABT1    activator of basal transcription 1   0.0025472722006984585
GO:0006366: transcription from RNA polymerase II promoter       ALX1    ALX homeobox 1  0.002297254352941165
GO:0006366: transcription from RNA polymerase II promoter       ATF5    activating transcription factor 5    -0.0027416416805262334
GO:0006366: transcription from RNA polymerase II promoter       BACH1   BTB and CNC homology 1, basic leucine zipper transcription factor 1   0.0004868934145365414
GO:0006366: transcription from RNA polymerase II promoter       BATF    basic leucine zipper transcription factor, ATF-like  0.002042081310521299
GO:0006366: transcription from RNA polymerase II promoter       CDC40   cell division cycle 40  0.0041752576736597425
GO:0006366: transcription from RNA polymerase II promoter       CEBPB   CCAAT/enhancer binding protein (C/EBP), beta -0.0002742560077731689
GO:0006366: transcription from RNA polymerase II promoter       CHD5    chromodomain helicase DNA binding protein 5  -0.0005202110736977483
GO:0006366: transcription from RNA polymerase II promoter       CLOCK   clock circadian regulator 0.00019972569283432376
GO:0006366: transcription from RNA polymerase II promoter       CREB1   cAMP responsive element binding protein 1    0.0006608582004217471
GO:0006366: transcription from RNA polymerase II promoter       DEK     DEK proto-oncogene      0.0024953548997541427
GO:0006366: transcription from RNA polymerase II promoter       DLX5    distal-less homeobox 5  -0.003290775322669836
GO:0006366: transcription from RNA polymerase II promoter       DMRT1   doublesex and mab-3 related transcription factor 1   0.0015087448243979954
GO:0006366: transcription from RNA polymerase II promoter       EGR1    early growth response 1 0.0010957056680258271
GO:0006366: transcription from RNA polymerase II promoter       EGR2    early growth response 2 0.001443871186574729
GO:0006366: transcription from RNA polymerase II promoter       ELF5    E74-like factor 5 (ets domain transcription factor)   0.0011765086377914667
GO:0006366: transcription from RNA polymerase II promoter       ESR1    estrogen receptor 1     -0.0009513485164322883
GO:0006366: transcription from RNA polymerase II promoter       ETV4    ets variant 4   0.00019594803348935608
GO:0006366: transcription from RNA polymerase II promoter       FOXA1   forkhead box A1 2.645648113322075e-5
GO:0006366: transcription from RNA polymerase II promoter       FOXA2   forkhead box A2 -1.6394897312831178e-5
GO:0006366: transcription from RNA polymerase II promoter       FOXC1   forkhead box C1 -2.12134145453641e-5
GO:0006366: transcription from RNA polymerase II promoter       FOXC2   forkhead box C2 (MFH-1, mesenchyme forkhead 1)       0.0017522536239867733
GO:0006366: transcription from RNA polymerase II promoter       FOXE3   forkhead box E3 0.0010532326424197403
GO:0006366: transcription from RNA polymerase II promoter       FOXH1   forkhead box H1 -0.0013755641386214129
GO:0006366: transcription from RNA polymerase II promoter       FOXM1   forkhead box M1 0.0001985638369795796
GO:0006366: transcription from RNA polymerase II promoter       GATA2   GATA binding protein 2  -0.0004533643608838128
GO:0006366: transcription from RNA polymerase II promoter       GATA3   GATA binding protein 3  -4.008793872914433e-5
GO:0006366: transcription from RNA polymerase II promoter       GATA4   GATA binding protein 4  -0.0010977494600769588
GO:0006366: transcription from RNA polymerase II promoter       GATA6   GATA binding protein 6  -2.76

54404874889836e-5
GO:0006366: transcription from RNA polymerase II promoter GLI2 GLI family zinc finger 2 0.0018581536683981122
GO:0006366: transcription from RNA polymerase II promoter GLI3 GLI family zinc finger 3 -0.0021581246674813683
GO:0006366: transcription from RNA polymerase II promoter GRHL2 grainyhead-like 2 (Drosophila) 0.0010088493737712768
GO:0006366: transcription from RNA polymerase II promoter HEYL hes-related family bHLH transcription factor with YRPW motif-like 0.0006793749302363291
GO:0006366: transcription from RNA polymerase II promoter HMGA1 high mobility group AT-hook 1 -0.0003188397277203899
GO:0006366: transcription from RNA polymerase II promoter HMGA2 high mobility group AT-hook 2 0.0015122341438013312
GO:0006366: transcription from RNA polymerase II promoter HOXA10 homeobox A10 -0.0029890960408370778
GO:0006366: transcription from RNA polymerase II promoter HOXA5 homeobox A5 0.0010693644314674717
GO:0006366: transcription from RNA polymerase II promoter HOXD13 homeobox D13 -0.0005931588345513611
GO:0006366: transcription from RNA polymerase II promoter IRF7 interferon regulatory factor 7 -0.0013096129999209797
GO:0006366: transcription from RNA polymerase II promoter ISL1 ISL LIM homeobox 1 7.844580151951696e-5
GO:0006366: transcription from RNA polymerase II promoter KLF11 Kruppel-like factor 11 0.000619651222449841
GO:0006366: transcription from RNA polymerase II promoter KLF15 Kruppel-like factor 15 0.0004664762545163968
GO:0006366: transcription from RNA polymerase II promoter KMT2A lysine (K)-specific methyltransferase 2A 0.000662335295352567
GO:0006366: transcription from RNA polymerase II promoter LEF1 lymphoid enhancer-binding factor 1 -0.0001003235972742402
GO:0006366: transcription from RNA polymerase II promoter LHX1 LIM homeobox 1 -0.0007608294814884724
GO:0006366: transcription from RNA polymerase II promoter LMO4 LIM domain only 4 0.0021171830417897842
GO:0006366: transcription from RNA polymerase II promoter MEF2C myocyte enhancer factor 2C 0.0009762179405557791
GO:0006366: transcription from RNA polymerase II promoter MSX1 msh homeobox 1 -0.0027769676042985574
GO:0006366: transcription from RNA polymerase II promoter NFIB nuclear factor I/B 0.0029286748040231597
GO:0006366: transcription from RNA polymerase II promoter NFX1 nuclear transcription factor, X-box binding 1 -6.8984398229704e-5
GO:0006366: transcription from RNA polymerase II promoter PAX2 paired box 2 -0.0016186836670777447
GO:0006366: transcription from RNA polymerase II promoter PAX3 paired box 3 -0.0029521020923864425
GO:0006366: transcription from RNA polymerase II promoter PAX6 paired box 6 0.001940853957981715
GO:0006366: transcription from RNA polymerase II promoter PAX8 paired box 8 0.0009497071214020347
GO:0006366: transcription from RNA polymerase II promoter PDX1 pancreatic and duodenal homeobox 1 0.00025835284695446787
GO:0006366: transcription from RNA polymerase II promoter PHOX2B paired-like homeobox 2b 0.00039080774700344403
GO:0006366: transcription from RNA polymerase II promoter PITX2 paired-like homeodomain 2 0.00218162860964497 8
GO:0006366: transcription from RNA polymerase II promoter POLR2D polymerase (RNA) II (DNA directed) polypeptide D 0.00185975041494696
GO:0006366: transcription from RNA polymerase II promoter POU1F1 POU class 1 homeobox 1 0.0001378738043097241
GO:0006366: transcription from RNA polymerase II promoter PRDM4 PR domain containing 4 0.0008418544216438413
GO:0006366: transcription from RNA polymerase II promoter PTTG1 pituitary tumor-transforming 1 -0.0003614587254092103
GO:0006366: transcription from RNA polymerase II promoter RAD21 RAD21 homolog (S. pombe) -0.00010210734744061777
GO:0006366: transcription from RNA polymerase II promoter RREB1 ras responsive element binding protein 1 -0.001985997174195349
GO:0006366: transcription from RNA polymerase II promoter SIX1 SIX homeobox 1 -0.0018746941995814426
GO:0006366: transcription from RNA polymerase II promoter SIX3 SIX homeobox 3 0.002043417511799642
GO:0006366: transcription from RNA polymerase II promoter SNRPB small nuclear ribonucleoprotein polypeptides B and B1 0.0006676086128169167
GO:0006366: transcription from RNA polymerase II promoter SOX10 SRY (sex determining region Y)-box 10 0.0001929542991030853
GO:0006366: transcription from RNA polymerase II promoter SOX11 SRY (sex determining region Y)-box 11 -0.00020860374601907087
GO:0006366: transcription from RNA polymerase II promoter SOX4 SRY (sex determining region Y)-box 4 -3.747035500166586e-5

GO:0006366: transcription from RNA polymerase II promoter SOX9 SRY (sex determining region Y)-box 9 -0.0005197028005894814
GO:0006366: transcription from RNA polymerase II promoter TBPL1 TBP-like 1 -0.0007895771694666627
GO:0006366: transcription from RNA polymerase II promoter TFAP2A transcription factor AP-2 alpha (activating enhancer binding protein 2 alpha) 0.00048071653699130733
GO:0006366: transcription from RNA polymerase II promoter TFAP2C transcription factor AP-2 gamma (activating enhancer binding protein 2 gamma) 0.0010647016641216049
GO:0006366: transcription from RNA polymerase II promoter THRA thyroid hormone receptor, alpha 0.0007648602333120478
GO:0006366: transcription from RNA polymerase II promoter TP73 tumor protein p73 0.0010304016491717311
GO:0006366: transcription from RNA polymerase II promoter TRPS1 trichorhinophalangeal syndrome I -0.00017915056168623763
GO:0006366: transcription from RNA polymerase II promoter WT1 Wilms tumor 1 -0.0005085229322826398
GO:0006366: transcription from RNA polymerase II promoter WWP2 WW domain containing E3 ubiquitin protein ligase 2 -0.0009593203466696437
GO:0006366: transcription from RNA polymerase II promoter XBP1 X-box binding protein 1 0.0002669579929930911
GO:0006366: transcription from RNA polymerase II promoter YBX1 Y box binding protein 1 -0.000914836001511645
GO:0006366: transcription from RNA polymerase II promoter ZNF148 zinc finger protein 148 0.002109356337630719
GO:0021522: spinal cord motor neuron differentiation ABT1 activator of basal transcription 1 0.002542266267672149
GO:0021522: spinal cord motor neuron differentiation DICER1 dicer 1, ribonuclease type III -1.6282337556684818e-5
GO:0021522: spinal cord motor neuron differentiation ISL1 ISL LIM homeobox 1 7.84279431322236e-5
GO:0021522: spinal cord motor neuron differentiation LMO4 LIM domain only 4 0.002113384723391936
GO:0021522: spinal cord motor neuron differentiation PTCH1 patched 1 -6.544972550372576e-5
GO:0021522: spinal cord motor neuron differentiation SHH sonic hedgehog 0.000600634764553926
GO:0021522: spinal cord motor neuron differentiation SOX4 SRY (sex determining region Y)-box 4 -3.747943937512266e-5
GO:0034462: small-subunit processome assembly ABT1 activator of basal transcription 1 0.002580882811878861
KEGG:03320: PPAR signaling pathway ACADL acyl-CoA dehydrogenase, long chain 0.0009646794784327429
KEGG:03320: PPAR signaling pathway CPT1A carnitine palmitoyltransferase 1A (liver) 0.0012641780921739988
KEGG:03320: PPAR signaling pathway NR1H3 nuclear receptor subfamily 1, group H, member 3 0.0007937988522384975
KEGG:03320: PPAR signaling pathway RXRA retinoid X receptor, alpha 0.0011022238142090102
KEGG:00071: Fatty acid degradation ACADL acyl-CoA dehydrogenase, long chain 0.0009596892229527938
KEGG:00071: Fatty acid degradation CPT1A carnitine palmitoyltransferase 1A (liver) 0.0012597313822292199
KEGG:00071: Fatty acid degradation GCDH glutaryl-CoA dehydrogenase -3.88142809877343e-6
GO:0001659: temperature homeostasis ACADL acyl-CoA dehydrogenase, long chain 0.0009707249903913637
GO:0001659: temperature homeostasis DRD2 dopamine receptor D2 -0.0002321098541831773
GO:0001659: temperature homeostasis FOXO1 forkhead box O1 0.0017839815989336971
GO:0001659: temperature homeostasis GPX1 glutathione peroxidase 1 0.0003892603849823642
GO:0006635: fatty acid beta-oxidation ACADL acyl-CoA dehydrogenase, long chain 0.0009899911006884669
GO:0006635: fatty acid beta-oxidation CPT1A carnitine palmitoyltransferase 1A (liver) 0.0012798922242036421
GO:0006635: fatty acid beta-oxidation CROT carnitine O-octanoyltransferase -0.00044311775031812873
GO:0006635: fatty acid beta-oxidation HSD17B4 hydroxysteroid (17-beta) dehydrogenase 4 -0.0011719591505483737
GO:0006635: fatty acid beta-oxidation LEP leptin 0.003219382871832427
GO:0006635: fatty acid beta-oxidation MUT methylmalonyl CoA mutase 0.0027779529764799214
GO:0019254: carnitine metabolic process, CoA-linked ACADL acyl-CoA dehydrogenase, long chain 0.0009725879762096733
GO:0033539: fatty acid beta-oxidation using acyl-CoA dehydrogenase ACADL acyl-CoA dehydrogenase, long chain 0.0009573338085733191
GO:0033539: fatty acid beta-oxidation using acyl-CoA dehydrogenase GCDH glutaryl-CoA dehydrogenase -3.6161329727611156e-6
GO:0033539: fatty acid beta-oxidation using acyl-CoA dehydrogenase IVD isovaleryl-CoA dehydrogenase -0.0009192251735621507
GO:0042413: carnitine catabolic process ACADL acyl-CoA dehydrogenase, long chain 0.0009725879762096733
GO:0042758: long-chain fatty acid catabolic process ACADL acyl-CoA dehydrogenase, long chain 0.0009725879762096733
GO:0044242: cellular lipid catabolic process ACADL acyl-CoA dehydrogenase, long chain 0.0009700934142145933
GO:0044242: cellular lipid catabolic process SIRT2 sirtuin 2 -0.00084212566063987
GO:0044255: cellular lipid metabolic process ACADL acyl-CoA dehydrogenase, long chain 0.000

9737906064588633
GO:0044255: cellular lipid metabolic process    ACOT8   acyl-CoA thioesterase 8 0.001131112167355671
GO:0044255: cellular lipid metabolic process    AGT     angiotensinogen (serpin peptidase inhibitor, clade A, member 8)     -0.001121058767068387
GO:0044255: cellular lipid metabolic process    BDH1    3-hydroxybutyrate dehydrogenase, type 1 -0.00020683157912009212
GO:0044255: cellular lipid metabolic process    CPT1A   carnitine palmitoyltransferase 1A (liver) 0.0012664374856221055
GO:0044255: cellular lipid metabolic process    CROT    carnitine O-octanoyltransferase -0.0004339150449823121
GO:0044255: cellular lipid metabolic process    ELOVL2  ELOVL fatty acid elongase 2     0.0005201646931292289
GO:0044255: cellular lipid metabolic process    ELOVL5  ELOVL fatty acid elongase 5     0.0002074362068424032
GO:0044255: cellular lipid metabolic process    GPD1L   glycerol-3-phosphate dehydrogenase 1-like 0.000563632495905647
GO:0044255: cellular lipid metabolic process    HMGCL   3-hydroxymethyl-3-methylglutaryl-CoA lyase -0.0010552294126392032
GO:0044255: cellular lipid metabolic process    HSD17B4 hydroxysteroid (17-beta) dehydrogenase 4 -0.001153829470771509
GO:0044255: cellular lipid metabolic process    MED1    mediator complex subunit 1      0.001119088548138185
GO:0044255: cellular lipid metabolic process    MUT     methylmalonyl CoA mutase        0.0027415226852462293
GO:0044255: cellular lipid metabolic process    PEX11A  peroxisomal biogenesis factor 11 alpha  0.0007132308480431775
GO:0044255: cellular lipid metabolic process    RXRA    retinoid X receptor, alpha      0.0011076667250260507
GO:0044255: cellular lipid metabolic process    SMARCD3 SWI/SNF related, matrix associated, actin dependent regulator of chromatin, subfamily d, member 3    0.00034280319274689725
GO:0044255: cellular lipid metabolic process    TNFRSF21        tumor necrosis factor receptor superfamily, member 21        0.00034500180558703057
GO:0045717: negative regulation of fatty acid biosynthetic process      ACADL   acyl-CoA dehydrogenase, long chain   0.0009725879762096733
GO:0046322: negative regulation of fatty acid oxidation ACADL   acyl-CoA dehydrogenase, long chain 0.0009725879762096733
GO:0051289: protein homotetramerization ACADL   acyl-CoA dehydrogenase, long chain      0.0009790380904007192
GO:0051289: protein homotetramerization ALDH5A1 aldehyde dehydrogenase 5 family, member A1      0.0023240892942921554
GO:0051289: protein homotetramerization CTH     cystathionine gamma-lyase       0.0017440659159030833
GO:0051289: protein homotetramerization EVL     Enah/Vasp-like  0.001936933421172402
GO:0051289: protein homotetramerization GOLGA2  golgin A2       0.00016771156185921472
GO:0051289: protein homotetramerization HPRT1   hypoxanthine phosphoribosyltransferase 1        -0.00118488733145369
GO:0051289: protein homotetramerization IMPDH2  IMP (inosine 5'-monophosphate) dehydrogenase 2  -0.0003455513770533281
GO:0051289: protein homotetramerization KCNJ2   potassium inwardly-rectifying channel, subfamily J, member 2 -0.0007752722053270478
GO:0051289: protein homotetramerization RXRA    retinoid X receptor, alpha      0.0011134638179271618
GO:0051289: protein homotetramerization SHMT2   serine hydroxymethyltransferase 2 (mitochondrial) -0.0010981723116027266
GO:0055088: lipid homeostasis   ACADL   acyl-CoA dehydrogenase, long chain      0.0009540101374639424
GO:0055088: lipid homeostasis   GCDH    glutaryl-CoA dehydrogenase      -3.422879204465277e-6
GO:0055088: lipid homeostasis   IVD     isovaleryl-CoA dehydrogenase    -0.0009170692873098383
GO:0055088: lipid homeostasis   NR1H3   nuclear receptor subfamily 1, group H, member 3 0.0007877153867270803
GO:0055114: oxidation-reduction process ACADL   acyl-CoA dehydrogenase, long chain      0.0009832253573415456
GO:0055114: oxidation-reduction process AKR1C1  aldo-keto reductase family 1, member C1 -0.0014911606652435313
GO:0055114: oxidation-reduction process ALDH1A2 aldehyde dehydrogenase 1 family, member A2      -0.0030238006667679973
GO:0055114: oxidation-reduction process AOC1    amine oxidase, copper containing 1      -0.0014253945956716426
GO:0055114: oxidation-reduction process APEX1   APEX nuclease (multifunctional DNA repair enzyme) 1 0.0002005883040891177
GO:0055114: oxidation-reduction process BDH1    3-hydroxybutyrate dehydrogenase, type 1 -0.00020740454285367692
GO:0055114: oxidation-reduction process CBS     cystathionine-beta-synthase     0.0015192448254002203
GO:0055114: oxidation-reduction process CP      ceruloplasmin (ferroxidase)     0.001309523779875639
GO:0055114: oxidation-reduction process CPOX    coproporphyrinogen oxidase      0.0012756851172449329
GO:0055114: oxidation-reduction process CYB5R4  cytochrome b5 reductase 4       -2.843469780966511e-5
GO:0055114: oxidation-reduction process CYP1B1  cytochrome P450, family 1, subfamily B, polypeptide 1 0.00039822360454679496
GO:0055114: oxidation-reduction process CYP27B1 cytochrome P450, family 27, subfamily B, polypeptide 1      -0.0004721558687292171
GO:0055114: oxidation-reduction process CYP4B1  cytochrome P450, family 4, subfamily B, polypeptide 1 -0.0013589948123098427
GO:0055114: oxidation-reduction process CYP7B1  cytochrome P450, family 7, subfamily B, polypeptide 1 0.00016352255250777294

GO:0055114: oxidation-reduction process DCT     dopachrome tautomerase  -0.0023567373536330953
GO:0055114: oxidation-reduction process DHCR24  24-dehydrocholesterol reductase -0.001710259766274022
3
GO:0055114: oxidation-reduction process DHFR    dihydrofolate reductase -0.0003078650587550917
GO:0055114: oxidation-reduction process GLRX2   glutaredoxin 2  0.001117743886771678
GO:0055114: oxidation-reduction process GLUD2   glutamate dehydrogenase 2       -0.000659897177300448
8
GO:0055114: oxidation-reduction process GPD1L   glycerol-3-phosphate dehydrogenase 1-like       0.000
568364014623038
GO:0055114: oxidation-reduction process HSD17B4 hydroxysteroid (17-beta) dehydrogenase 4        -0.00
11645341890049057
GO:0055114: oxidation-reduction process HSD17B6 hydroxysteroid (17-beta) dehydrogenase 6        -0.00
08687761628787211
GO:0055114: oxidation-reduction process HSDL2   hydroxysteroid dehydrogenase like 2     0.00137043369
0322614
GO:0055114: oxidation-reduction process IDO1    indoleamine 2,3-dioxygenase 1   0.0012279669183532969
GO:0055114: oxidation-reduction process IMPDH1  IMP (inosine 5'-monophosphate) dehydrogenase 1  -0.00
18751854036111262
GO:0055114: oxidation-reduction process IMPDH2  IMP (inosine 5'-monophosphate) dehydrogenase 2  -0.00
034838345354934344
GO:0055114: oxidation-reduction process JMJD6   jumonji domain containing 6     0.0035963202880579183
GO:0055114: oxidation-reduction process LOX     lysyl oxidase   -0.0005774280777408367
GO:0055114: oxidation-reduction process MTHFD1  methylenetetrahydrofolate dehydrogenase (NADP+ depend
ent) 1, methenyltetrahydrofolate cyclohydrolase, formyltetrahydrofolate synthetase      -0.0006649171
409238845
GO:0055114: oxidation-reduction process PAX2    paired box 2    -0.0016168312553755796
GO:0055114: oxidation-reduction process PHGDH   phosphoglycerate dehydrogenase  0.0002934951681328274
4
GO:0055114: oxidation-reduction process PNPO    pyridoxamine 5'-phosphate oxidase       -0.0010931856
105589374
GO:0055114: oxidation-reduction process POR     P450 (cytochrome) oxidoreductase        0.00045800762
3863459
GO:0055114: oxidation-reduction process PRDX4   peroxiredoxin 4 0.001526965842019623
GO:0055114: oxidation-reduction process QDPR    quinoid dihydropteridine reductase      -0.0028603630
968169476
GO:0055114: oxidation-reduction process RRM1    ribonucleotide reductase M1     0.0019135618967990816
GO:0055114: oxidation-reduction process RRM2    ribonucleotide reductase M2     -0.000566635548573676
2
GO:0055114: oxidation-reduction process SORD    sorbitol dehydrogenase  -0.002022930934530212
GO:0055114: oxidation-reduction process SPR     sepiapterin reductase (7,8-dihydrobiopterin:NADP+ oxi
doreductase)    -0.0012347649997144635
GO:0055114: oxidation-reduction process SRD5A2  steroid-5-alpha-reductase, alpha polypeptide 2 (3-oxo
-5 alpha-steroid delta 4-dehydrogenase alpha 2) -0.0006443451086806785
GO:0055114: oxidation-reduction process STEAP3  STEAP family member 3, metalloreductase 0.00261307963
59889754
GO:0055114: oxidation-reduction process STEAP4  STEAP family member 4   -0.0027720278485580856
GO:0055114: oxidation-reduction process TH      tyrosine hydroxylase    -0.0003424965178086481
GO:0055114: oxidation-reduction process TXN2    thioredoxin 2   0.0013852052524674083
GO:0055114: oxidation-reduction process UGDH    UDP-glucose 6-dehydrogenase     0.0005023683750838108
GO:0090181: regulation of cholesterol metabolic process ACADL   acyl-CoA dehydrogenase, long chain
0.0009725879762096733
KEGG:00120: Primary bile acid biosynthesis      ACOT8   acyl-CoA thioesterase 8 0.0011250415158007095
KEGG:00120: Primary bile acid biosynthesis      CYP7B1  cytochrome P450, family 7, subfamily B, polyp
eptide 1        0.00016078807902458238
KEGG:00120: Primary bile acid biosynthesis      HSD17B4 hydroxysteroid (17-beta) dehydrogenase 4
-0.0011418915082748655
KEGG:04146: Peroxisome  ACOT8   acyl-CoA thioesterase 8 0.001126720298550438
KEGG:04146: Peroxisome  CROT    carnitine O-octanoyltransferase -0.00043112880764478 64
KEGG:04146: Peroxisome  HMGCL   3-hydroxymethyl-3-methylglutaryl-CoA lyase      -0.001049136670314674
KEGG:04146: Peroxisome  HSD17B4 hydroxysteroid (17-beta) dehydrogenase 4        -0.001147517066830305
4
KEGG:04146: Peroxisome  PEX11A  peroxisomal biogenesis factor 11 alpha  0.0007093964743551375
GO:0006637: acyl-CoA metabolic process  ACOT8   acyl-CoA thioesterase 8 0.0011296365421344244
GO:0006637: acyl-CoA metabolic process  HMGCL   3-hydroxymethyl-3-methylglutaryl-CoA lyase      -0.00
10515053781074918
GO:0006699: bile acid biosynthetic process      ACOT8   acyl-CoA thioesterase 8 0.0011264948182661994
GO:0006699: bile acid biosynthetic process      CYP7B1  cytochrome P450, family 7, subfamily B, polyp
eptide 1        0.00016106635738449642
GO:0006699: bile acid biosynthetic process      HSD17B4 hydroxysteroid (17-beta) dehydrogenase 4
-0.0011450626516497026
GO:0006699: bile acid biosynthetic process      STAR    steroidogenic acute regulatory protein  0.000
879700503651401
GO:0008206: bile acid metabolic process ACOT8   acyl-CoA thioesterase 8 0.0011400378260621646
GO:0008206: bile acid metabolic process AKR1C1  aldo-keto reductase family 1, member C1 -0.0014955608
771145693
GO:0008206: bile acid metabolic process CYP7B1  cytochrome P450, family 7, subfamily B, polypeptide 1
0.00016378248230008343
GO:0008206: bile acid metabolic process HSD17B4 hydroxysteroid (17-beta) dehydrogenase 4        -0.00
1168350972566201
GO:0008206: bile acid metabolic process LEP     leptin  0.003209837568461649
GO:0008206: bile acid metabolic process RXRA    retinoid X receptor, alpha      0.0011208793103175208

GO:0016032: viral process	ACOT8	acyl-CoA thioesterase 8 0.0011332087985093957
GO:0016032: viral process	BAX	BCL2-associated X protein	-0.0004239889194544546
GO:0016032: viral process	BICD1	bicaudal D homolog 1 (Drosophila)	0.0024310978956263865
GO:0016032: viral process	BNIP3	BCL2/adenovirus E1B 19kDa interacting protein 3 0.00289938193
3048571
GO:0016032: viral process	BNIP3L	BCL2/adenovirus E1B 19kDa interacting protein 3-like	-5.76
3728910336969e-5
GO:0016032: viral process	BRD4	bromodomain containing 4	0.0005528164370892127
GO:0016032: viral process	BUB1	BUB1 mitotic checkpoint serine/threonine kinase 0.00131793572
73544678
GO:0016032: viral process	CALCOCO2	calcium binding and coiled-coil domain 2	0.002
4467567733152218
GO:0016032: viral process	CCDC86	coiled-coil domain containing 86	0.0007146750595481014
GO:0016032: viral process	CREB1	cAMP responsive element binding protein 1	0.00065389482
45720485
GO:0016032: viral process	CUL7	cullin 7	-0.00011804496107927637
GO:0016032: viral process	CX3CR1	chemokine (C-X3-C motif) receptor 1	-0.000853631577973926
GO:0016032: viral process	CXCR4	chemokine (C-X-C motif) receptor 4	0.0007903798646155341
GO:0016032: viral process	DAXX	death-domain associated protein 0.0008866141034356596
GO:0016032: viral process	E4F1	E4F transcription factor 1	0.001477511507960713
GO:0016032: viral process	EIF4G1	eukaryotic translation initiation factor 4 gamma, 1	0.000
7391653691435424
GO:0016032: viral process	GFI1	growth factor independent 1 transcription repressor	0.001
658638564957058
GO:0016032: viral process	GRB2	growth factor receptor-bound protein 2	0.0004279285947750677
GO:0016032: viral process	HMGA1	high mobility group AT-hook 1	-0.00031825120185198025
GO:0016032: viral process	IL6ST	interleukin 6 signal transducer 0.001868790179750359
GO:0016032: viral process	IPO5	importin 5	0.00167576941341883
GO:0016032: viral process	KPNB1	karyopherin (importin) beta 1	0.0007693123028523802
GO:0016032: viral process	KRT18	keratin 18	-0.001012147259849316
GO:0016032: viral process	LYN	LYN proto-oncogene, Src family tyrosine kinase	-0.0014147386
365213096
GO:0016032: viral process	MAP3K5	mitogen-activated protein kinase kinase kinase 5	0.000
34071323975268605
GO:0016032: viral process	MFGE8	milk fat globule-EGF factor 8 protein	0.0017956025031863694
GO:0016032: viral process	MSH6	mutS homolog 6	0.001071519475507141
GO:0016032: viral process	NFX1	nuclear transcription factor, X-box binding 1	-7.1670514024
0043e-5
GO:0016032: viral process	NUP153	nucleoporin 153kDa	0.0008197281779380466
GO:0016032: viral process	PML	promyelocytic leukemia	-0.0006785609772968493
GO:0016032: viral process	POLA1	polymerase (DNA directed), alpha 1, catalytic subunit	0.001
2637540074285627
GO:0016032: viral process	POLR2D	polymerase (RNA) II (DNA directed) polypeptide D	0.001
8466217380317513
GO:0016032: viral process	PSMA5	proteasome (prosome, macropain) subunit, alpha type, 5	0.000
3300625245152298
GO:0016032: viral process	PSMD11	proteasome (prosome, macropain) 26S subunit, non-ATPase, 11
-0.0010919739958158618
GO:0016032: viral process	PSMD13	proteasome (prosome, macropain) 26S subunit, non-ATPase, 13
-6.048485203822167e-5
GO:0016032: viral process	RB1	retinoblastoma 1	-0.0014873904032028239
GO:0016032: viral process	RHOA	ras homolog family member A	0.0006670271923979145
GO:0016032: viral process	SEH1L	SEH1-like (S. cerevisiae)	-0.0007307140972634648
GO:0016032: viral process	SF3B2	splicing factor 3b, subunit 2, 145kDa	0.0006803728928588617
GO:0016032: viral process	SIRT1	sirtuin 1	-2.557371602793581e-6
GO:0016032: viral process	TFRC	transferrin receptor	0.0010804880289843422
GO:0016032: viral process	TP53	tumor protein p53	0.0011687103860240397
GO:0016032: viral process	TP73	tumor protein p73	0.001023504705947392
GO:0016032: viral process	TPR	translocated promoter region, nuclear basket protein	-0.00
0694545133445699
GO:0016032: viral process	UNG	uracil-DNA glycosylase	0.0005621207451607176
GO:0016032: viral process	VPS4A	vacuolar protein sorting 4 homolog A (S. cerevisiae)	-0.00
06818751482759128
GO:0016559: peroxisome fission	ACOT8	acyl-CoA thioesterase 8 0.0011242906316926771
GO:0016559: peroxisome fission	PEX11A	peroxisomal biogenesis factor 11 alpha	0.00070192948185206
GO:0033540: fatty acid beta-oxidation using acyl-CoA oxidase	ACOT8	acyl-CoA thioesterase 8 0.001
126342885732238
GO:0033540: fatty acid beta-oxidation using acyl-CoA oxidase	CROT	carnitine O-octanoyltransfera
se	-0.00043112441729309923
GO:0033540: fatty acid beta-oxidation using acyl-CoA oxidase	HSD17B4 hydroxysteroid (17-beta) dehy
drogenase 4	-0.0011472315828132777
GO:0033559: unsaturated fatty acid metabolic process	ACOT8	acyl-CoA thioesterase 8 0.00112413389
60080748
GO:0033559: unsaturated fatty acid metabolic process	ELOVL2	ELOVL fatty acid elongase 2	0.000
5126286214051828
GO:0033559: unsaturated fatty acid metabolic process	ELOVL5	ELOVL fatty acid elongase 5	0.000
20808309900323675
GO:0033559: unsaturated fatty acid metabolic process	HSD17B4 hydroxysteroid (17-beta) dehydrogenas
e 4	-0.0011418480529381604
GO:0036109: alpha-linolenic acid metabolic process	ACOT8	acyl-CoA thioesterase 8 0.00112413389
60080748

GO:0036109: alpha-linolenic acid metabolic process	ELOVL2	ELOVL fatty acid elongase 2	0.000
5126286214051828
GO:0036109: alpha-linolenic acid metabolic process	ELOVL5	ELOVL fatty acid elongase 5	0.000
20808309900323675
GO:0036109: alpha-linolenic acid metabolic process	HSD17B4	hydroxysteroid (17-beta) dehydrogenas
e 4	-0.0011418480529381604
GO:0043649: dicarboxylic acid catabolic process ACOT8	acyl-CoA thioesterase 8 0.0011250785466930266
GO:0007275: multicellular organismal development	ACRV1	acrosomal vesicle protein 1	-0.00
06026324800037688
GO:0007275: multicellular organismal development	AES	amino-terminal enhancer of split
0.0013608250886879664
GO:0007275: multicellular organismal development	ALX1	ALX homeobox 1	0.0023048546834939392
GO:0007275: multicellular organismal development	CENPE	centromere protein E, 312kDa	0.002
1641843690307526
GO:0007275: multicellular organismal development	DSPP	dentin sialophosphoprotein	0.000
8332002663454993
GO:0007275: multicellular organismal development	EGFL6	EGF-like-domain, multiple 6	0.000
1379128003375985
GO:0007275: multicellular organismal development	EYA3	EYA transcriptional coactivator and p
hosphatase 3	0.000841405296330342
GO:0007275: multicellular organismal development	GCNT2	glucosaminyl (N-acetyl) transferase
2, I-branching enzyme (I blood group)	0.002446769111629609
GO:0007275: multicellular organismal development	GNRHR	gonadotropin-releasing hormone recept
or	0.000744818180673966
GO:0007275: multicellular organismal development	HELLS	helicase, lymphoid-specific	0.003
484141576224596
GO:0007275: multicellular organismal development	HMGA2	high mobility group AT-hook 2	0.001
5176454559921598
GO:0007275: multicellular organismal development	HMGB3	high mobility group box 3	0.000
46591417626646643
GO:0007275: multicellular organismal development	HOXA10	homeobox A10	-0.002999580579960398
GO:0007275: multicellular organismal development	HOXB1	homeobox B1	0.0037064002506680916
GO:0007275: multicellular organismal development	HOXB2	homeobox B2	0.002788429492393047
GO:0007275: multicellular organismal development	HOXD13	homeobox D13	-0.000594838497653422
9
GO:0007275: multicellular organismal development	ID2	inhibitor of DNA binding 2, dominant
negative helix-loop-helix protein	5.999482185643313e-5
GO:0007275: multicellular organismal development	PAEP	progestagen-associated endometrial pr
otein	0.0010706068741512258
GO:0007275: multicellular organismal development	RREB1	ras responsive element binding protei
n 1	-0.0019922654238996524
GO:0007275: multicellular organismal development	SPDEF	SAM pointed domain containing ETS tra
nscription factor	0.003154445846453971
GO:0007275: multicellular organismal development	SUCO	SUN domain containing ossification fa
ctor	0.002261786124292338
GO:0007275: multicellular organismal development	TBX21	T-box 21	0.0004003452354227899
6
GO:0007275: multicellular organismal development	TP53	tumor protein p53	0.00118414751
6533695
GO:0001525: angiogenesis	ACVRL1	activin A receptor type II-like 1	0.0019380999107114758
GO:0001525: angiogenesis	APOD	apolipoprotein D	0.002619877964834698
GO:0001525: angiogenesis	ARHGAP24	Rho GTPase activating protein 24	-0.0009316971
671739718
GO:0001525: angiogenesis	CAV1	caveolin 1, caveolae protein, 22kDa	-0.000534296501562445
5
GO:0001525: angiogenesis	CCL2	chemokine (C-C motif) ligand 2	0.0008139262642777212
GO:0001525: angiogenesis	CCL8	chemokine (C-C motif) ligand 8	-0.0005893403229684583
GO:0001525: angiogenesis	CIB1	calcium and integrin binding 1 (calmyrin)	5.56054062538
9172e-5
GO:0001525: angiogenesis	CYP1B1	cytochrome P450, family 1, subfamily B, polypeptide 1	0.000
3967426204059147
GO:0001525: angiogenesis	DICER1	dicer 1, ribonuclease type III	-1.6371415982969285e-5
GO:0001525: angiogenesis	ECM1	extracellular matrix protein 1	-0.0015511685488916952
GO:0001525: angiogenesis	EFNA1	ephrin-A1	-0.0005195861981931489
GO:0001525: angiogenesis	EPHB1	EPH receptor B1 0.00195840660180056
GO:0001525: angiogenesis	EPHB2	EPH receptor B2 -0.0007209679109037155
GO:0001525: angiogenesis	EPHB3	EPH receptor B3 0.0008808647439006741
GO:0001525: angiogenesis	FAP	fibroblast activation protein, alpha	-0.000484703068957530
1
GO:0001525: angiogenesis	FGFR2	fibroblast growth factor receptor 2	0.0007605775669832328
GO:0001525: angiogenesis	HIF1A	hypoxia inducible factor 1, alpha subunit (basic helix-loop-h
elix transcription factor)	-0.0006448825921510365
GO:0001525: angiogenesis	HMOX1	heme oxygenase (decycling) 1	-0.0002167468167063793
GO:0001525: angiogenesis	HOXA3	homeobox A3	0.0009858971129960933
GO:0001525: angiogenesis	HOXB13	homeobox B13	0.0018204492376021157
GO:0001525: angiogenesis	JAG1	jagged 1	0.0017523636559871694
GO:0001525: angiogenesis	MED1	mediator complex subunit 1	0.0011248810657117855
GO:0001525: angiogenesis	MFGE8	milk fat globule-EGF factor 8 protein	0.001798406832602931
GO:0001525: angiogenesis	MMP2	matrix metallopeptidase 2 (gelatinase A, 72kDa gelatinase, 72
kDa type IV collagenase)	-0.0011267150998895677
GO:0001525: angiogenesis	NRP1	neuropilin 1	-0.0006407896487169695

GO:0001525: angiogenesis	PRKCA	protein kinase C, alpha -5.91854794812547e-6
GO:0001525: angiogenesis	PRKX	protein kinase, X-linked	0.0003925327430821656
GO:0001525: angiogenesis	PTEN	phosphatase and tensin homolog	1.7995857133031824e-5
GO:0001525: angiogenesis	RBPJ	recombination signal binding protein for immunoglobulin kappa J region	0.0009790331924208254
GO:0001525: angiogenesis	S100A7	S100 calcium binding protein A7 0.0015846662048894947
GO:0001525: angiogenesis	SERPINE1	serpin peptidase inhibitor, clade E (nexin, plasminogen activator inhibitor type 1), member 1	0.00011263528208728846
GO:0001525: angiogenesis	SIRT1	sirtuin 1	-2.1779228749137594e-6
GO:0001525: angiogenesis	SOX18	SRY (sex determining region Y)-box 18	0.001285129056028295
GO:0001525: angiogenesis	TGFB2	transforming growth factor, beta 2	-0.0010555947352951396
GO:0001525: angiogenesis	TGFBR1	transforming growth factor, beta receptor 1	0.0003384609672932745
GO:0001525: angiogenesis	VASH1	vasohibin 1	0.0005968959576848898
GO:0001525: angiogenesis	VEGFA	vascular endothelial growth factor A	0.0005921704392206562
GO:0001525: angiogenesis	VEGFC	vascular endothelial growth factor C	-0.0033536915302535245
GO:0001525: angiogenesis	WNT7A	wingless-type MMTV integration site family, member 7A	1.965404811143532e-5
GO:0001525: angiogenesis	XBP1	X-box binding protein 1 0.0002670899867725781
GO:0001701: in utero embryonic development	ACVRL1	activin A receptor type II-like 1	0.0019427990637215053
GO:0001701: in utero embryonic development	ADAR	adenosine deaminase, RNA-specific	-0.00013659547190262366
GO:0001701: in utero embryonic development	ANGPT1	angiopoietin 1	0.0009002207230357272
GO:0001701: in utero embryonic development	AXIN1	axin 1	-0.0007326864437604174
GO:0001701: in utero embryonic development	BCL2L11 BCL2-like 11 (apoptosis facilitator)	-0.0007743436337115884
GO:0001701: in utero embryonic development	CCNB1	cyclin B1	-0.0008817261174187466
GO:0001701: in utero embryonic development	CTNNB1	catenin (cadherin-associated protein), beta 1, 88kDa	-0.00011684514818058755
GO:0001701: in utero embryonic development	EPN1	epsin 1 -0.0005464726383597678
GO:0001701: in utero embryonic development	FGFR2	fibroblast growth factor receptor 2	0.0007627787719822484
GO:0001701: in utero embryonic development	FOXA2	forkhead box A2 -1.6718968042080125e-5
GO:0001701: in utero embryonic development	FOXC1	forkhead box C1 -2.2151633814181628e-5
GO:0001701: in utero embryonic development	GATA3	GATA binding protein 3	-3.9114628763145796e-5
GO:0001701: in utero embryonic development	GATA4	GATA binding protein 4	-0.0010941244136378353
GO:0001701: in utero embryonic development	GATA6	GATA binding protein 6	-2.8120494314859988e-5
GO:0001701: in utero embryonic development	GDF3	growth differentiation factor 3 -0.0018829331093555759
GO:0001701: in utero embryonic development	GJA1	gap junction protein, alpha 1, 43kDa	-0.0001614499533835122
GO:0001701: in utero embryonic development	GLI2	GLI family zinc finger 2	0.00185233581312177
GO:0001701: in utero embryonic development	GLI3	GLI family zinc finger 3	-0.0021503754049436827
GO:0001701: in utero embryonic development	GRHL2	grainyhead-like 2 (Drosophila)	0.0010062146366325907
GO:0001701: in utero embryonic development	HINFP	histone H4 transcription factor 0.0009920066048793143
GO:0001701: in utero embryonic development	IHH	indian hedgehog -0.002039318298078183
GO:0001701: in utero embryonic development	ITGB1	integrin, beta 1 (fibronectin receptor, beta polypeptide, antigen CD29 includes MDF2, MSK12)	0.002424991902474287
GO:0001701: in utero embryonic development	JAG2	jagged 2	-5.707070766657228e-6
GO:0001701: in utero embryonic development	KLF2	Kruppel-like factor 2	-0.0012315113992547777
GO:0001701: in utero embryonic development	KMT2D	lysine (K)-specific methyltransferase 2D	-0.0001992888189322397
GO:0001701: in utero embryonic development	MSH2	mutS homolog 2	0.0013914541109964563
GO:0001701: in utero embryonic development	MSX1	msh homeobox 1	-0.0027672491734704405
GO:0001701: in utero embryonic development	MYH10	myosin, heavy chain 10, non-muscle	-0.00032860395085510566
GO:0001701: in utero embryonic development	NOTCH1	notch 1 0.0005185223076809099
GO:0001701: in utero embryonic development	PCNT	pericentrin	0.001786291035467117
GO:0001701: in utero embryonic development	PDGFRB	platelet-derived growth factor receptor, beta polypeptide	-0.0003609376634283868
GO:0001701: in utero embryonic development	PITX2	paired-like homeodomain 2	0.002174714057885522
GO:0001701: in utero embryonic development	POLE	polymerase (DNA directed), epsilon, catalytic subunit 0.00294662192778463
GO:0001701: in utero embryonic development	PTCH1	patched 1	-6.590551620196469e-5
GO:0001701: in utero embryonic development	RNASEH2B	ribonuclease H2, subunit B	-4.3543304634598356e-5
GO:0001701: in utero embryonic development	RPA1	replication protein A1, 70kDa	0.00043890123848049404
GO:0001701: in utero embryonic development	RXRA	retinoid X receptor, alpha	0.00111439159

42975634
GO:0001701: in utero embryonic development SLIT2 slit homolog 2 (Drosophila) -0.0016191656
339764277
GO:0001701: in utero embryonic development SMO smoothened, frizzled class receptor 0.002
137949263826817
GO:0001701: in utero embryonic development SOX10 SRY (sex determining region Y)-box 10 0.000
19024190012834557
GO:0001701: in utero embryonic development SOX18 SRY (sex determining region Y)-box 18 0.001
2877585846430403
GO:0001701: in utero embryonic development TBX3 T-box 3 0.0012255975362851354
GO:0001701: in utero embryonic development TGFB3 transforming growth factor, beta 3 -0.00
18226037846556834
GO:0001701: in utero embryonic development TGFBR1 transforming growth factor, beta receptor 1
0.00033966513742213363
GO:0001701: in utero embryonic development TP53 tumor protein p53 0.0011748888646252158
GO:0001701: in utero embryonic development TWIST1 twist family bHLH transcription factor 1
-0.001340640539082981
GO:0001701: in utero embryonic development VEGFA vascular endothelial growth factor A 0.000
594572999930559
GO:0001701: in utero embryonic development WDR19 WD repeat domain 19 -0.000721730544462502
5
GO:0001701: in utero embryonic development YBX1 Y box binding protein 1 -0.000913341286919911
7
GO:0001701: in utero embryonic development ZBTB18 zinc finger and BTB domain containing 18
0.0012713024023252371
GO:0001701: in utero embryonic development ZNF335 zinc finger protein 335 -0.000364760986451832
16
GO:0001936: regulation of endothelial cell proliferation ACVRL1 activin A receptor type II-li
ke 1 0.00196396605902291
GO:0001936: regulation of endothelial cell proliferation ALDH1A2 aldehyde dehydrogenase 1 fami
ly, member A2 -0.003050478342022817
GO:0001937: negative regulation of endothelial cell proliferation ACVRL1 activin A receptor ty
pe II-like 1 0.0019250874938716665
GO:0001937: negative regulation of endothelial cell proliferation AGER advanced glycosylatio
n end product-specific receptor -0.0001732035223705776
GO:0001937: negative regulation of endothelial cell proliferation CAV1 caveolin 1, caveolae
protein, 22kDa -0.0005315075031537393
GO:0001937: negative regulation of endothelial cell proliferation ENG endoglin 0.000
8160904693519045
GO:0001937: negative regulation of endothelial cell proliferation GJA1 gap junction protein,
alpha 1, 43kDa -0.00015587520759072234
GO:0001937: negative regulation of endothelial cell proliferation SULF1 sulfatase 1 -0.00
07928164889860242
GO:0001937: negative regulation of endothelial cell proliferation TGFBR1 transforming growth f
actor, beta receptor 1 0.0003354764853332717
GO:0001937: negative regulation of endothelial cell proliferation THBS1 thrombospondin 1
-0.001025212600209794
GO:0001937: negative regulation of endothelial cell proliferation VASH1 vasohibin 1 0.000
5934720064626205
GO:0001938: positive regulation of endothelial cell proliferation ACVRL1 activin A receptor ty
pe II-like 1 0.001923187106381941
GO:0001938: positive regulation of endothelial cell proliferation AKT1 v-akt murine thymoma
viral oncogene homolog 1 0.0007213552958856503
GO:0001938: positive regulation of endothelial cell proliferation BMP4 bone morphogenetic pr
otein 4 -0.0003190363261614075
GO:0001938: positive regulation of endothelial cell proliferation CAV1 caveolin 1, caveolae
protein, 22kDa -0.000531205788571826
GO:0001938: positive regulation of endothelial cell proliferation CCL2 chemokine (C-C motif)
ligand 2 0.0008073470833861536
GO:0001938: positive regulation of endothelial cell proliferation CXCL12 chemokine (C-X-C moti
f) ligand 12 -0.0011941913984007727
GO:0001938: positive regulation of endothelial cell proliferation ECM1 extracellular matrix
protein 1 -0.0015363472780654832
GO:0001938: positive regulation of endothelial cell proliferation FGFR3 fibroblast growth fac
tor receptor 3 0.00021856251848388454
GO:0001938: positive regulation of endothelial cell proliferation HIF1A hypoxia inducible fac
tor 1, alpha subunit (basic helix-loop-helix transcription factor) -0.000640315571894329
GO:0001938: positive regulation of endothelial cell proliferation HMGB2 high mobility group b
ox 2 0.0002979610535235678
GO:0001938: positive regulation of endothelial cell proliferation NRP1 neuropilin 1 -0.00
06346855167324198
GO:0001938: positive regulation of endothelial cell proliferation PRKCA protein kinase C, alp
ha -5.930897761783614e-6
GO:0001938: positive regulation of endothelial cell proliferation PROX1 prospero homeobox 1
0.0011075362866752774
GO:0001938: positive regulation of endothelial cell proliferation TGFBR1 transforming growth f
actor, beta receptor 1 0.00033495335834605246
GO:0001938: positive regulation of endothelial cell proliferation THBS4 thrombospondin 4
-0.00044152113232423295
GO:0001938: positive regulation of endothelial cell proliferation VEGFA vascular endothelial
growth factor A 0.0005856396246766729

GO:0001938: positive regulation of endothelial cell proliferation WNT5A wingless-type MMTV in
tegration site family, member 5A -0.0006593814258707266
GO:0001946: lymphangiogenesis ACVRL1 activin A receptor type II-like 1 0.00194612909198142
GO:0001946: lymphangiogenesis FOXC2 forkhead box C2 (MFH-1, mesenchyme forkhead 1) 0.00175124628
23182425
GO:0001946: lymphangiogenesis PROX1 prospero homeobox 1 0.0011227685110076979
GO:0001946: lymphangiogenesis SOX18 SRY (sex determining region Y)-box 18 0.0012909123757203745
GO:0001955: blood vessel maturation ACVRL1 activin A receptor type II-like 1 0.00195271488
17602255
GO:0001955: blood vessel maturation MMP2 matrix metallopeptidase 2 (gelatinase A, 72kDa gelati
nase, 72kDa type IV collagenase) -0.0011387836895902072
GO:0001974: blood vessel remodeling ACVRL1 activin A receptor type II-like 1 0.00193709465
21689789
GO:0001974: blood vessel remodeling AGT angiotensinogen (serpin peptidase inhibitor, clade A,
member 8) -0.0011264545369124859
GO:0001974: blood vessel remodeling BAK1 BCL2-antagonist/killer 1 -0.001869782097675899
GO:0001974: blood vessel remodeling BAX BCL2-associated X protein -0.000424362714137155
46
GO:0001974: blood vessel remodeling FGF8 fibroblast growth factor 8 (androgen-induced) 0.000
9795387522410451
GO:0001974: blood vessel remodeling FOXC1 forkhead box C1 -2.3998618519730733e-5
GO:0001974: blood vessel remodeling FOXC2 forkhead box C2 (MFH-1, mesenchyme forkhead 1) 0.001
7423736182592224
GO:0001974: blood vessel remodeling HOXA3 homeobox A3 0.0009854992287546627
GO:0001974: blood vessel remodeling IGF1 insulin-like growth factor 1 (somatomedin C) 0.000
1297890536110021
GO:0001974: blood vessel remodeling JAG1 jagged 1 0.001752178132189106
GO:0001974: blood vessel remodeling MEF2C myocyte enhancer factor 2C 0.0009691194573440721
GO:0001974: blood vessel remodeling RBPJ recombination signal binding protein for immunoglobul
in kappa J region 0.0009788711086604438
GO:0001974: blood vessel remodeling SEMA3C sema domain, immunoglobulin domain (Ig), short basic
domain, secreted, (semaphorin) 3C -0.0002192360870652542
GO:0001974: blood vessel remodeling TGFB2 transforming growth factor, beta 2 -0.0010548733
572331346
GO:0002043: blood vessel endothelial cell proliferation involved in sprouting angiogenesis ACVRL
1 activin A receptor type II-like 1 0.001941089062057476
GO:0002043: blood vessel endothelial cell proliferation involved in sprouting angiogenesis BMP4
bone morphogenetic protein 4 -0.0003218816800786746
GO:0006275: regulation of DNA replication ACVRL1 activin A receptor type II-like 1 0.001
9557258365855866
GO:0006468: protein phosphorylation ACVRL1 activin A receptor type II-like 1 0.00194851571
38485387
GO:0006468: protein phosphorylation AKT1 v-akt murine thymoma viral oncogene homolog 1 0.000
7327183796848718
GO:0006468: protein phosphorylation AKT2 v-akt murine thymoma viral oncogene homolog 2 -0.00
0627221375344193
GO:0006468: protein phosphorylation AURKA aurora kinase A 0.0010107899453714246
GO:0006468: protein phosphorylation AURKB aurora kinase B 0.00035186325673515744
GO:0006468: protein phosphorylation AURKC aurora kinase C 0.0008881894274280079
GO:0006468: protein phosphorylation BIRC5 baculoviral IAP repeat containing 5 -0.0002606857
8214866954
GO:0006468: protein phosphorylation BRD4 bromodomain containing 4 0.0005545687792096688
GO:0006468: protein phosphorylation BUB1 BUB1 mitotic checkpoint serine/threonine kinase 0.001
3285965783585128
GO:0006468: protein phosphorylation BUB1B BUB1 mitotic checkpoint serine/threonine kinase B
-0.0012838557908560729
GO:0006468: protein phosphorylation CAMK1D calcium/calmodulin-dependent protein kinase ID -0.00
25945301030741792
GO:0006468: protein phosphorylation CCL2 chemokine (C-C motif) ligand 2 0.0008186401066323783
GO:0006468: protein phosphorylation CCL8 chemokine (C-C motif) ligand 8 -0.000591441696383413
4
GO:0006468: protein phosphorylation CCND1 cyclin D1 -0.0026407986233100127
GO:0006468: protein phosphorylation CCNE1 cyclin E1 0.00037201059294231085
GO:0006468: protein phosphorylation CDK8 cyclin-dependent kinase 8 0.002618452964160652
GO:0006468: protein phosphorylation CDKL5 cyclin-dependent kinase-like 5 0.0022902028514099307
GO:0006468: protein phosphorylation CFL1 cofilin 1 (non-muscle) -0.0011782671739032863
GO:0006468: protein phosphorylation CREB1 cAMP responsive element binding protein 1 0.000
6606053514895301
GO:0006468: protein phosphorylation CSNK2B casein kinase 2, beta polypeptide 0.00153287649
46920697
GO:0006468: protein phosphorylation GSK3B glycogen synthase kinase 3 beta 0.0015531071795888412
GO:0006468: protein phosphorylation GUCY2C guanylate cyclase 2C (heat stable enterotoxin recepto
r) -0.0016372526591578646
GO:0006468: protein phosphorylation HIPK2 homeodomain interacting protein kinase 2 0.000
77249390823163
GO:0006468: protein phosphorylation IGFBP3 insulin-like growth factor binding protein 3 0.000
8382644580709868
GO:0006468: protein phosphorylation IRAK1 interleukin-1 receptor-associated kinase 1 -0.00
16130541618468866
GO:0006468: protein phosphorylation JAK2 Janus kinase 2 -3.137260031916693e-5
GO:0006468: protein phosphorylation LYN LYN proto-oncogene, Src family tyrosine kinase -0.00

14239427431591298
GO:0006468: protein phosphorylation MAP3K5 mitogen-activated protein kinase kinase kinase 5 0.0003435018603542358
GO:0006468: protein phosphorylation MMD monocyte to macrophage differentiation-associated 0.00215373907974547 2
GO:0006468: protein phosphorylation MOK MOK protein kinase 0.00179391350799655 7
GO:0006468: protein phosphorylation NEK2 NIMA-related kinase 2 4.958374578080412e-5
GO:0006468: protein phosphorylation PAK1 p21 protein (Cdc42/Rac)-activated kinase 1 -0.00 2193587250243135
GO:0006468: protein phosphorylation PBK PDZ binding kinase 0.0008433623468465547
GO:0006468: protein phosphorylation PDK1 pyruvate dehydrogenase kinase, isozyme 1 0.001 4293432237704438
GO:0006468: protein phosphorylation PDK2 pyruvate dehydrogenase kinase, isozyme 2 0.001 4834715708303677
GO:0006468: protein phosphorylation PICK1 protein interacting with PRKCA 1 -0.0009574612 23837719
GO:0006468: protein phosphorylation PIK3CD phosphatidylinositol-4,5-bisphosphate 3-kinase, catal ytic subunit delta -0.0007828086515596969
GO:0006468: protein phosphorylation PKN1 protein kinase N1 -0.0019077501569560072
GO:0006468: protein phosphorylation PLK1 polo-like kinase 1 0.0010206881598149634
GO:0006468: protein phosphorylation PLK3 polo-like kinase 3 0.0025700428188143378
GO:0006468: protein phosphorylation PLK4 polo-like kinase 4 0.00299373214569410 7
GO:0006468: protein phosphorylation PRKCA protein kinase C, alpha -5.876725573191029e-6
GO:0006468: protein phosphorylation PRKCD protein kinase C, delta -0.0011315960280285288
GO:0006468: protein phosphorylation PRKCZ protein kinase C, zeta -0.0016046056222054341
GO:0006468: protein phosphorylation RAF1 Raf-1 proto-oncogene, serine/threonine kinase 0.001 5024938699221531
GO:0006468: protein phosphorylation RET ret proto-oncogene -0.0004922159180050824
GO:0006468: protein phosphorylation RPS6KA1 ribosomal protein S6 kinase, 90kDa, polypeptide 1 -0.002520030028593816
GO:0006468: protein phosphorylation STK24 serine/threonine kinase 24 0.00270133945196708 7
GO:0006468: protein phosphorylation TEX14 testis expressed 14 0.0017302251876234883
GO:0006468: protein phosphorylation TGFB1 transforming growth factor, beta 1 -7.3280217973 23936e-5
GO:0006468: protein phosphorylation TGFB2 transforming growth factor, beta 2 -0.0010623869 165459125
GO:0006468: protein phosphorylation TGFBR1 transforming growth factor, beta receptor 1 0.000 3409874916133343
GO:0007162: negative regulation of cell adhesion ACVRL1 activin A receptor type II-like 1 0.0019268955101691455
GO:0007162: negative regulation of cell adhesion AGER advanced glycosylation end product-sp ecific receptor -0.00017354573912856203
GO:0007162: negative regulation of cell adhesion ANGPT1 angiopoietin 1 0.0008935206257666982
GO:0007162: negative regulation of cell adhesion RASA1 RAS p21 protein activator (GTPase act ivating protein) 1 -0.0003440275050910197
GO:0007165: signal transduction ACVRL1 activin A receptor type II-like 1 0.001943488290606878
GO:0007165: signal transduction ADM adrenomedullin 0.0022374310578607325
GO:0007165: signal transduction AKT1 v-akt murine thymoma viral oncogene homolog 1 0.00073040738 70056509
GO:0007165: signal transduction AKT2 v-akt murine thymoma viral oncogene homolog 2 -0.0006257732 791294791
GO:0007165: signal transduction AMHR2 anti-Mullerian hormone receptor, type II -0.0002246880 3078219986
GO:0007165: signal transduction ANK3 ankyrin 3, node of Ranvier (ankyrin G) 0.0007000095618207148
GO:0007165: signal transduction AR androgen receptor 0.0026377324035076126
GO:0007165: signal transduction AVP arginine vasopressin -0.0009371013629622085
GO:0007165: signal transduction BCL11A B-cell CLL/lymphoma 11A (zinc finger protein) -0.0009769692 989772172
GO:0007165: signal transduction C3 complement component 3 0.0020241813149136
GO:0007165: signal transduction CASP8AP2 caspase 8 associated protein 2 0.0012716188181623072
GO:0007165: signal transduction CCL2 chemokine (C-C motif) ligand 2 0.0008164781588547727
GO:0007165: signal transduction CCL7 chemokine (C-C motif) ligand 7 -0.002402192590972942
GO:0007165: signal transduction CCL8 chemokine (C-C motif) ligand 8 -0.0005902866570882145
GO:0007165: signal transduction CHRNB1 cholinergic receptor, nicotinic, beta 1 (muscle) -6.97 4939505178814e-6
GO:0007165: signal transduction CHRNB2 cholinergic receptor, nicotinic, beta 2 (neuronal) -0.00 07782428189131891
GO:0007165: signal transduction CLOCK clock circadian regulator 0.00020015421509193593
GO:0007165: signal transduction CREB1 cAMP responsive element binding protein 1 0.00065806314 26769614
GO:0007165: signal transduction CSF3R colony stimulating factor 3 receptor (granulocyte) -8.33 914413561887e-5
GO:0007165: signal transduction CSNK2B casein kinase 2, beta polypeptide 0.001528290022274502
GO:0007165: signal transduction CXCL10 chemokine (C-X-C motif) ligand 10 6.252176489902889e-5
GO:0007165: signal transduction CXCL12 chemokine (C-X-C motif) ligand 12 -0.001208108420582850 5
GO:0007165: signal transduction DEK DEK proto-oncogene 0.002487075539820206
GO:0007165: signal transduction DOCK1 dedicator of cytokinesis 1 -0.0006840752277506568
GO:0007165: signal transduction ECM1 extracellular matrix protein 1 -0.0015567835148277815
GO:0007165: signal transduction EDA ectodysplasin A -0.0008142998396472185
GO:0007165: signal transduction EGFR epidermal growth factor receptor 0.00068555423265277 5

GO:0007165: signal transduction ERBB4 v-erb-b2 avian erythroblastic leukemia viral oncogene homolog 4 -0.00018282737676140516
GO:0007165: signal transduction ESR1 estrogen receptor 1 -0.0009479087845693361
GO:0007165: signal transduction FAS Fas cell surface death receptor -3.324798933360833e-5
GO:0007165: signal transduction FGF3 fibroblast growth factor 3 0.0015606321391585846
GO:0007165: signal transduction FGF7 fibroblast growth factor 7 0.0006335279489453931
GO:0007165: signal transduction GATA3 GATA binding protein 3 -3.902895825587389e-5
GO:0007165: signal transduction GDNF glial cell derived neurotrophic factor 0.0004446913272187222
GO:0007165: signal transduction GJA1 gap junction protein, alpha 1, 43kDa -0.00016145881297111378
GO:0007165: signal transduction GOLT1B golgi transport 1B -0.0018204313940101472
GO:0007165: signal transduction GREM1 gremlin 1, DAN family BMP antagonist -0.000827025232494862 8
GO:0007165: signal transduction HIF1A hypoxia inducible factor 1, alpha subunit (basic helix-loop-helix transcription factor) -0.0006463380068602735
GO:0007165: signal transduction IGBP1 immunoglobulin (CD79A) binding protein 1 0.0028487610213682314
GO:0007165: signal transduction IGF1 insulin-like growth factor 1 (somatomedin C) 0.00013073943590575065
GO:0007165: signal transduction IGF1R insulin-like growth factor 1 receptor 0.0010718343173586854
GO:0007165: signal transduction IGFBP1 insulin-like growth factor binding protein 1 0.0003878467873103585
GO:0007165: signal transduction IGFBP2 insulin-like growth factor binding protein 2, 36kDa 0.0001447531946209149
GO:0007165: signal transduction IGFBP4 insulin-like growth factor binding protein 4 -0.0012781286022579322
GO:0007165: signal transduction IGFBP6 insulin-like growth factor binding protein 6 -0.0020276698443934186
GO:0007165: signal transduction INHA inhibin, alpha 0.00021168128751260598
GO:0007165: signal transduction INPP4B inositol polyphosphate-4-phosphatase, type II, 105kDa 0.0002898240135355377 6
GO:0007165: signal transduction IRAK1 interleukin-1 receptor-associated kinase 1 -0.001608217676736459
GO:0007165: signal transduction ITPR1 inositol 1,4,5-trisphosphate receptor, type 1 -0.0007801932860876994
GO:0007165: signal transduction JAK2 Janus kinase 2 -3.184119929580907e-5
GO:0007165: signal transduction KIT v-kit Hardy-Zuckerman 4 feline sarcoma viral oncogene homolog 0.0002746279766027156
GO:0007165: signal transduction LGALS9 lectin, galactoside-binding, soluble, 9 -0.0001168634768572361 2
GO:0007165: signal transduction LRP8 low density lipoprotein receptor-related protein 8, apolipoprotein e receptor -0.0009315977289665422
GO:0007165: signal transduction LYN LYN proto-oncogene, Src family tyrosine kinase -0.0014201196210848553
GO:0007165: signal transduction MDK midkine (neurite growth-promoting factor 2) 0.0016645696032585094
GO:0007165: signal transduction MOK MOK protein kinase 0.0017891484430256099
GO:0007165: signal transduction NCK2 NCK adaptor protein 2 -0.00023207792304874
GO:0007165: signal transduction NDRG2 NDRG family member 2 0.002172917084249932
GO:0007165: signal transduction NMU neuromedin U 0.0018500546101785204
GO:0007165: signal transduction NRP1 neuropilin 1 -0.0006429554165685962
GO:0007165: signal transduction OR5I1 olfactory receptor, family 5, subfamily I, member 1 0.0018819441389188883
GO:0007165: signal transduction PDGFRB platelet-derived growth factor receptor, beta polypeptide -0.00036099984777882965
GO:0007165: signal transduction PEX11A peroxisomal biogenesis factor 11 alpha 0.0007172058370507031
GO:0007165: signal transduction PGR progesterone receptor 0.00035181374841912206
GO:0007165: signal transduction PIK3CD phosphatidylinositol-4,5-bisphosphate 3-kinase, catalytic subunit delta -0.0007806355534794371
GO:0007165: signal transduction PKN1 protein kinase N1 -0.0019025145368926843
GO:0007165: signal transduction PLCB1 phospholipase C, beta 1 (phosphoinositide-specific) 0.00015369129227155235
GO:0007165: signal transduction PRDM4 PR domain containing 4 0.000840152247260057
GO:0007165: signal transduction PRKCA protein kinase C, alpha -5.858715728466283e-6
GO:0007165: signal transduction PRKCD protein kinase C, delta -0.0011280425279524252
GO:0007165: signal transduction PRKCZ protein kinase C, zeta -0.0016003973826187452
GO:0007165: signal transduction PTK7 protein tyrosine kinase 7 -0.00022523708569734615
GO:0007165: signal transduction RAF1 Raf-1 proto-oncogene, serine/threonine kinase 0.0014984721965360791
GO:0007165: signal transduction RASA1 RAS p21 protein activator (GTPase activating protein) 1 -0.0003489944204112381
GO:0007165: signal transduction RASSF8 Ras association (RalGDS/AF-6) domain family (N-terminal) member 8 0.0033201926705127994
GO:0007165: signal transduction RET ret proto-oncogene -0.0004911562201324102
GO:0007165: signal transduction RPS6KA1 ribosomal protein S6 kinase, 90kDa, polypeptide 1 -0.0025137535542865577
GO:0007165: signal transduction SNX17 sorting nexin 17 0.0016412085197914953
GO:0007165: signal transduction SOX9 SRY (sex determining region Y)-box 9 -0.0005184320937996085
GO:0007165: signal transduction SPHK1 sphingosine kinase 1 0.0018104643880386781
GO:0007165: signal transduction STK24 serine/threonine kinase 24 0.00269410110934512

GO:0007165: signal transduction STMN1   stathmin 1      0.0005589882128292227
GO:0007165: signal transduction STX2    syntaxin 2      0.00030798354041124235
GO:0007165: signal transduction TGFBR1  transforming growth factor, beta receptor 1     0.00033977308
512114464
GO:0007165: signal transduction TNFRSF10C       tumor necrosis factor receptor superfamily, member 10
c, decoy without an intracellular domain        0.0003171586710983202
GO:0007165: signal transduction TNFRSF11A       tumor necrosis factor receptor superfamily, member 11
a, NFKB activator       0.0027751532878522835
GO:0007165: signal transduction TRAIP   TRAF interacting protein        0.0010222919425565634
GO:0007165: signal transduction VDR     vitamin D (1,25- dihydroxyvitamin D3) receptor  0.00056349640
89186081
GO:0007165: signal transduction VEGFC   vascular endothelial growth factor C    -0.00336328308155984l
GO:0007165: signal transduction VLDLR   very low density lipoprotein receptor   0.0009526634738594302
GO:0007165: signal transduction ZPR1    ZPR1 zinc finger        -0.0002094504814459663
GO:0007179: transforming growth factor beta receptor signaling pathway  ACVRL1  activin A receptor ty
pe II-like 1    0.0019477170205270315
GO:0007179: transforming growth factor beta receptor signaling pathway  AMHR2   anti-Mullerian hormon
e receptor, type II     -0.0002249287370729811
GO:0007179: transforming growth factor beta receptor signaling pathway  CCL2    chemokine (C-C motif)
ligand 2        0.0008183175303271573
GO:0007179: transforming growth factor beta receptor signaling pathway  CCNC    cyclin C        0.000
5688221525635015
GO:0007179: transforming growth factor beta receptor signaling pathway  CDK8    cyclin-dependent kina
se 8    0.0026173712241212847
GO:0007179: transforming growth factor beta receptor signaling pathway  CDKN2B  cyclin-dependent kina
se inhibitor 2B (p15, inhibits CDK4)    -0.0018392719383721055
GO:0007179: transforming growth factor beta receptor signaling pathway  CITED1  Cbp/p300-interacting
 transactivator, with Glu/Asp-rich carboxy-terminal domain, 1   0.0028244334171750652
GO:0007179: transforming growth factor beta receptor signaling pathway  CREB1   cAMP responsive eleme
nt binding protein 1    0.0006603604642300682
GO:0007179: transforming growth factor beta receptor signaling pathway  E2F4    E2F transcription fac
tor 4, p107/p130-binding        -0.0023502401060925245
GO:0007179: transforming growth factor beta receptor signaling pathway  ENG     endoglin        0.000
8287895959262676
GO:0007179: transforming growth factor beta receptor signaling pathway  FOXH1   forkhead box H1 -0.00
13747248438405284
GO:0007179: transforming growth factor beta receptor signaling pathway  GCNT2   glucosaminyl (N-acety
l) transferase 2, I-branching enzyme (I blood group)    0.002437106835718056
GO:0007179: transforming growth factor beta receptor signaling pathway  HIPK2   homeodomain interacti
ng protein kinase 2     0.0007721425017938069
GO:0007179: transforming growth factor beta receptor signaling pathway  MYC     v-myc avian myelocyto
matosis viral oncogene homolog  -0.0011384836277658148
GO:0007179: transforming growth factor beta receptor signaling pathway  PML     promyelocytic leukemi
a       -0.0006808554572287162
GO:0007179: transforming growth factor beta receptor signaling pathway  PRKCZ   protein kinase C, zet
a       -0.001603878967921764
GO:0007179: transforming growth factor beta receptor signaling pathway  RHOA    ras homolog family me
mber A  0.0006711381340593412
GO:0007179: transforming growth factor beta receptor signaling pathway  SERPINE1        serpin peptid
ase inhibitor, clade E (nexin, plasminogen activator inhibitor type 1), member 1        0.00011209206
002378356
GO:0007179: transforming growth factor beta receptor signaling pathway  SKI     SKI proto-oncogene
-0.0006716534248830596
GO:0007179: transforming growth factor beta receptor signaling pathway  TFDP1   transcription factor
 Dp-1   0.0014487226922070134
GO:0007179: transforming growth factor beta receptor signaling pathway  TGFB1   transforming growth f
actor, beta 1   -7.32868377945739e-5
GO:0007179: transforming growth factor beta receptor signaling pathway  TGFB2   transforming growth f
actor, beta 2   -0.0010619459331235477
GO:0007179: transforming growth factor beta receptor signaling pathway  TGFB3   transforming growth f
actor, beta 3   -0.0018281370084749257
GO:0007179: transforming growth factor beta receptor signaling pathway  TGFBR1  transforming growth f
actor, beta receptor 1  0.00034086779108912684
GO:0007179: transforming growth factor beta receptor signaling pathway  TP53    tumor protein p53
0.0011789064079598336
GO:0007179: transforming growth factor beta receptor signaling pathway  WWTR1   WW domain containing
 transcription regulator 1      0.000903050246202591
GO:0008015: blood circulation   ACVRL1  activin A receptor type II-like 1       0.0019546331310049602
GO:0008015: blood circulation   ADM     adrenomedullin  0.0022504405831868593
GO:0008015: blood circulation   CXCL10  chemokine (C-X-C motif) ligand 10       6.3652464121497e-5
GO:0008015: blood circulation   CXCL12  chemokine (C-X-C motif) ligand 12       -0.001215595278646423
7
GO:0008015: blood circulation   E2F4    E2F transcription factor 4, p107/p130-binding   -0.0023586915
419635014
GO:0008015: blood circulation   HOXB2   homeobox B2     0.0027878740896505228
GO:0008217: regulation of blood pressure        ACVRL1  activin A receptor type II-like 1       0.001
9582715530163924
GO:0008217: regulation of blood pressure        AGT     angiotensinogen (serpin peptidase inhibitor,
 clade A, member 8)     -0.0011463655435048638
GO:0008217: regulation of blood pressure        HMOX1   heme oxygenase (decycling) 1    -0.0002198384
5923779873

GO:0008217: regulation of blood pressure LEP leptin 0.0032197843419849363
GO:0008217: regulation of blood pressure LRP5 low density lipoprotein receptor-related protein 5 2.941578442795649e-5
GO:0008285: negative regulation of cell proliferation ACVRL1 activin A receptor type II-like 1 0.0019450423689848506
GO:0008285: negative regulation of cell proliferation ADM adrenomedullin 0.002239194025931009
GO:0008285: negative regulation of cell proliferation AGT angiotensinogen (serpin peptidase inhibitor, clade A, member 8) -0.0011340968087181543
GO:0008285: negative regulation of cell proliferation AIMP2 aminoacyl tRNA synthetase complex-interacting multifunctional protein 2 0.0012097239612722764
GO:0008285: negative regulation of cell proliferation ALDH1A2 aldehyde dehydrogenase 1 family, member A2 -0.0030209456747758013
GO:0008285: negative regulation of cell proliferation APC adenomatous polyposis coli 0.0006440335004019952
GO:0008285: negative regulation of cell proliferation AR androgen receptor 0.002639767820816289
GO:0008285: negative regulation of cell proliferation ATF5 activating transcription factor 5 -0.002736752461099586
GO:0008285: negative regulation of cell proliferation BAK1 BCL2-antagonist/killer 1 -0.0018772370398121956
GO:0008285: negative regulation of cell proliferation BCHE butyrylcholinesterase -6.976529494566937e-6
GO:0008285: negative regulation of cell proliferation BMP4 bone morphogenetic protein 4 -0.0003227789396435444
GO:0008285: negative regulation of cell proliferation CD9 CD9 molecule -0.002587709018700916
GO:0008285: negative regulation of cell proliferation CDC6 cell division cycle 6 0.001354879995944264
GO:0008285: negative regulation of cell proliferation CDKN2A cyclin-dependent kinase inhibitor 2A 0.0017593414912815564
GO:0008285: negative regulation of cell proliferation CDKN2B cyclin-dependent kinase inhibitor 2B (p15, inhibits CDK4) -0.0018366482265428058
GO:0008285: negative regulation of cell proliferation CHD5 chromodomain helicase DNA binding protein 5 -0.0005192094837780246
GO:0008285: negative regulation of cell proliferation CIB1 calcium and integrin binding 1 (calmyrin) 5.457377976086573e-5
GO:0008285: negative regulation of cell proliferation CSNK2B casein kinase 2, beta polypeptide 0.0015297135610614443
GO:0008285: negative regulation of cell proliferation CTNNB1 catenin (cadherin-associated protein), beta 1, 88kDa -0.00011700726060175805
GO:0008285: negative regulation of cell proliferation CYP1B1 cytochrome P450, family 1, subfamily B, polypeptide 1 0.00039787750542139347
GO:0008285: negative regulation of cell proliferation CYP27B1 cytochrome P450, family 27, subfamily B, polypeptide 1 -0.00047153962910640736
GO:0008285: negative regulation of cell proliferation DHCR24 24-dehydrocholesterol reductase -0.0017085568606880312
GO:0008285: negative regulation of cell proliferation DRD2 dopamine receptor D2 -0.00023414467350260133
GO:0008285: negative regulation of cell proliferation ERBB4 v-erb-b2 avian erythroblastic leukemia viral oncogene homolog 4 -0.00018313082969778505
GO:0008285: negative regulation of cell proliferation FGFR3 fibroblast growth factor receptor 3 0.00022285984370167556
GO:0008285: negative regulation of cell proliferation FLT3 fms-related tyrosine kinase 3 -0.0006819895209086497
GO:0008285: negative regulation of cell proliferation GATA3 GATA binding protein 3 -3.9013435158005835e-5
GO:0008285: negative regulation of cell proliferation GLI3 GLI family zinc finger 3 -0.0021530501276903267
GO:0008285: negative regulation of cell proliferation HMGA1 high mobility group AT-hook 1 -0.00031887073191870735
GO:0008285: negative regulation of cell proliferation IGF1 insulin-like growth factor 1 (somatomedin C) 0.00013110964500248759
GO:0008285: negative regulation of cell proliferation IGFBP3 insulin-like growth factor binding protein 3 0.0008371353197730286
GO:0008285: negative regulation of cell proliferation IGFBP6 insulin-like growth factor binding protein 6 -0.002029294690595642
GO:0008285: negative regulation of cell proliferation INHBA inhibin, beta A -0.0013541910655605754
GO:0008285: negative regulation of cell proliferation IRF6 interferon regulatory factor 6 -0.0016947300086202332
GO:0008285: negative regulation of cell proliferation JAK2 Janus kinase 2 -3.168009369485464e-5
GO:0008285: negative regulation of cell proliferation KLF11 Kruppel-like factor 11 0.0006172392148230504
GO:0008285: negative regulation of cell proliferation KMT2A lysine (K)-specific methyltransferase 2A 0.0006610398783569432
GO:0008285: negative regulation of cell proliferation LYN LYN proto-oncogene, Src family tyrosine kinase -0.0014213375854111867
GO:0008285: negative regulation of cell proliferation MSX1 msh homeobox 1 -0.0027708860856549456
GO:0008285: negative regulation of cell proliferation NCK2 NCK adaptor protein 2 -0.00023213668445202452
GO:0008285: negative regulation of cell proliferation NDRG1 N-myc downstream regulated 1 -0.00

09414853496860377
GO:0008285: negative regulation of cell proliferation   NF2     neurofibromin 2 (merlin)        -0.00
12725663301033133
GO:0008285: negative regulation of cell proliferation   NOTCH1  notch 1 0.0005190883491611527
GO:0008285: negative regulation of cell proliferation   PDX1    pancreatic and duodenal homeobox 1
0.0002575120281259301
GO:0008285: negative regulation of cell proliferation   PHOX2B  paired-like homeobox 2b 0.00039063016
1464768
GO:0008285: negative regulation of cell proliferation   PML     promyelocytic leukemia  -0.0006802973
368918826
GO:0008285: negative regulation of cell proliferation   POU1F1  POU class 1 homeobox 1  0.00013753675
60970426
GO:0008285: negative regulation of cell proliferation   PRKCA   protein kinase C, alpha -5.8807554021
953956e-6
GO:0008285: negative regulation of cell proliferation   PROX1   prospero homeobox 1     0.00112225812
74418985
GO:0008285: negative regulation of cell proliferation   PTEN    phosphatase and tensin homolog  1.699
1644797505496e-5
GO:0008285: negative regulation of cell proliferation   PTPN2   protein tyrosine phosphatase, non-rec
eptor type 2    -0.00032946570920118387
GO:0008285: negative regulation of cell proliferation   PTPRJ   protein tyrosine phosphatase, recepto
r type, J       -0.00033868018785245137
GO:0008285: negative regulation of cell proliferation   RAF1    Raf-1 proto-oncogene, serine/threonin
e kinase        0.001499718148365218
GO:0008285: negative regulation of cell proliferation   RARG    retinoic acid receptor, gamma   -0.00
23208919093867454
GO:0008285: negative regulation of cell proliferation   RBBP4   retinoblastoma binding protein 4
-0.0018615449201155015
GO:0008285: negative regulation of cell proliferation   RBPJ    recombination signal binding protein
 for immunoglobulin kappa J region      0.0009819875758064524
GO:0008285: negative regulation of cell proliferation   RXRA    retinoid X receptor, alpha      0.001
115788516632251
GO:0008285: negative regulation of cell proliferation   SERPINE2        serpin peptidase inhibitor, c
lade E (nexin, plasminogen activator inhibitor type 1), member 2        0.0025044221025937565
GO:0008285: negative regulation of cell proliferation   SFRP1   secreted frizzled-related protein 1
0.001282812940569684
GO:0008285: negative regulation of cell proliferation   SIRT2   sirtuin 2       -0.000852828402004903
4
GO:0008285: negative regulation of cell proliferation   SKI     SKI proto-oncogene      -0.0006708552
260220733
GO:0008285: negative regulation of cell proliferation   SLIT2   slit homolog 2 (Drosophila)     -0.00
1621271714509996
GO:0008285: negative regulation of cell proliferation   SOX4    SRY (sex determining region Y)-box 4
-3.8238102958755076e-5
GO:0008285: negative regulation of cell proliferation   TFAP2A  transcription factor AP-2 alpha (acti
vating enhancer binding protein 2 alpha)        0.0004801644901072094
GO:0008285: negative regulation of cell proliferation   TGFB1   transforming growth factor, beta 1
-7.306558685080601e-5
GO:0008285: negative regulation of cell proliferation   TGFB2   transforming growth factor, beta 2
-0.0010601243020513839
GO:0008285: negative regulation of cell proliferation   TGFB3   transforming growth factor, beta 3
-0.001824953092794578
GO:0008285: negative regulation of cell proliferation   TP53    tumor protein p53       0.00117659179
19464674
GO:0008285: negative regulation of cell proliferation   TP73    tumor protein p73       0.00102857626
28292005
GO:0008285: negative regulation of cell proliferation   VDR     vitamin D (1,25- dihydroxyvitamin D3)
receptor        0.0005638134434367348
GO:0008285: negative regulation of cell proliferation   VEGFC   vascular endothelial growth factor C
-0.003366077614319254
GO:0008285: negative regulation of cell proliferation   WT1     Wilms tumor 1   -0.000507278461243390
4
GO:0010596: negative regulation of endothelial cell migration   ACVRL1  activin A receptor type II-li
ke 1    0.001939254430979354
GO:0010596: negative regulation of endothelial cell migration   AGER    advanced glycosylation end pr
oduct-specific receptor -0.00017405620776148923
GO:0010596: negative regulation of endothelial cell migration   SLIT2   slit homolog 2 (Drosophila)
-0.0016155582912411354
GO:0010596: negative regulation of endothelial cell migration   THBS1   thrombospondin 1        -0.00
10327682943718805
GO:0010596: negative regulation of endothelial cell migration   VASH1   vasohibin 1     0.00059751929
73211141
GO:0010862: positive regulation of pathway-restricted SMAD protein phosphorylation      ACVRL1  activ
in A receptor type II-like 1    0.001942963068277838
GO:0010862: positive regulation of pathway-restricted SMAD protein phosphorylation      BMP4    bone
 morphogenetic protein 4        -0.0003224820909915349
GO:0010862: positive regulation of pathway-restricted SMAD protein phosphorylation      BMP7    bone
 morphogenetic protein 7        0.0008550989828034138
GO:0010862: positive regulation of pathway-restricted SMAD protein phosphorylation      CSNK2B  casei
n kinase 2, beta polypeptide    0.0015282231022148666
GO:0010862: positive regulation of pathway-restricted SMAD protein phosphorylation      ENG     endog

lin	0.0008261881775255028
GO:0010862: positive regulation of pathway-restricted SMAD protein phosphorylation	GDF3	growt
h differentiation factor 3	-0.0018830764182839463
GO:0010862: positive regulation of pathway-restricted SMAD protein phosphorylation	INHA	inhib
in, alpha	0.0002113348382898421
GO:0010862: positive regulation of pathway-restricted SMAD protein phosphorylation	INHBA	inhib
in, beta A	-0.001352833754871801
GO:0010862: positive regulation of pathway-restricted SMAD protein phosphorylation	TGFB1	trans
forming growth factor, beta 1	-7.314932938012392e-5
GO:0010862: positive regulation of pathway-restricted SMAD protein phosphorylation	TGFB2	trans
forming growth factor, beta 2	-0.0010589174293650278
GO:0010862: positive regulation of pathway-restricted SMAD protein phosphorylation	TGFB3	trans
forming growth factor, beta 3	-0.0018232911276634782
GO:0010862: positive regulation of pathway-restricted SMAD protein phosphorylation	TGFBR1	trans
forming growth factor, beta receptor 1	0.0003398629920313265
GO:0010862: positive regulation of pathway-restricted SMAD protein phosphorylation	TTK	TTK p
rotein kinase	0.0010856640624797387
GO:0023014: signal transduction by protein phosphorylation	ACVRL1	activin A receptor type II-li
ke 1	0.0019476480050772892
GO:0023014: signal transduction by protein phosphorylation	AMHR2	anti-Mullerian hormone recept
or, type II	-0.0002249630732012948
GO:0023014: signal transduction by protein phosphorylation	INSR	insulin receptor	-0.00
13661233431596818
GO:0023014: signal transduction by protein phosphorylation	LYN	LYN proto-oncogene, Src famil
y tyrosine kinase	-0.0014238942193058267
GO:0023014: signal transduction by protein phosphorylation	STK24	serine/threonine kinase 24
0.002700100931704591
GO:0023014: signal transduction by protein phosphorylation	TGFB2	transforming growth factor, b
eta 2	-0.0010619596012031703
GO:0023014: signal transduction by protein phosphorylation	TGFBR1	transforming growth factor, b
eta receptor 1	0.0003406119658438516
GO:0030308: negative regulation of cell growth	ACVRL1	activin A receptor type II-like 1	0.001
946593706219525
GO:0030308: negative regulation of cell growth	AGT	angiotensinogen (serpin peptidase inhibitor,
 clade A, member 8)	-0.0011366798436235892
GO:0030308: negative regulation of cell growth	BCL2	B-cell CLL/lymphoma 2	-5.2359613861494495e-
6
GO:0030308: negative regulation of cell growth	CDKN2A	cyclin-dependent kinase inhibitor 2A	0.001
7620138049313186
GO:0030308: negative regulation of cell growth	CRYAB	crystallin, alpha B	0.0009872064460255264
GO:0030308: negative regulation of cell growth	CYP27B1	cytochrome P450, family 27, subfamily B, poly
peptide 1	-0.00047250785522871103
GO:0030308: negative regulation of cell growth	GJA1	gap junction protein, alpha 1, 43kDa	-0.00
016379374024781917
GO:0030308: negative regulation of cell growth	GREM1	gremlin 1, DAN family BMP antagonist	-0.00
08299974383682902
GO:0030308: negative regulation of cell growth	INHBA	inhibin, beta A	-0.0013570205450785645
GO:0030308: negative regulation of cell growth	MSX1	msh homeobox 1	-0.002774196213296862
GO:0030308: negative regulation of cell growth	NF2	neurofibromin 2 (merlin)	-0.0012725211
664658487
GO:0030308: negative regulation of cell growth	PML	promyelocytic leukemia	-0.000679924255983231
GO:0030308: negative regulation of cell growth	PRDM4	PR domain containing 4	0.000840850758006366
GO:0030308: negative regulation of cell growth	PSRC1	proline/serine-rich coiled-coil 1	0.000
6490186907055042
GO:0030308: negative regulation of cell growth	PTPRJ	protein tyrosine phosphatase, receptor type,
 J	-0.0003385804136027843
GO:0030308: negative regulation of cell growth	SERPINE2	serpin peptidase inhibitor, clade E
 (nexin, plasminogen activator inhibitor type 1), member 2	0.0025067765308946824
GO:0030308: negative regulation of cell growth	SFRP1	secreted frizzled-related protein 1	0.001
2865837019925067
GO:0030308: negative regulation of cell growth	SIRT1	sirtuin 1	-1.0999163846877494e-6
GO:0030308: negative regulation of cell growth	SLIT2	slit homolog 2 (Drosophila)	-0.0016231315
668071007
GO:0030308: negative regulation of cell growth	TGFB1	transforming growth factor, beta 1	-7.35
2850406007193e-5
GO:0030308: negative regulation of cell growth	TGFB2	transforming growth factor, beta 2	-0.00
10613630616584163
GO:0030308: negative regulation of cell growth	TP53	tumor protein p53	0.0011786364252468068
GO:0030308: negative regulation of cell growth	WT1	Wilms tumor 1	-0.0005080295022154231
GO:0030336: negative regulation of cell migration	ACVRL1	activin A receptor type II-like 1
0.0019417931581083522
GO:0030336: negative regulation of cell migration	ARPIN	actin-related protein 2/3 complex inh
ibitor	-0.00014142792688151751
GO:0030336: negative regulation of cell migration	BCL2	B-cell CLL/lymphoma 2	-5.0980556268
56445e-6
GO:0030336: negative regulation of cell migration	CX3CR1	chemokine (C-X3-C motif) receptor 1
-0.0008561084499457647
GO:0030336: negative regulation of cell migration	CYP1B1	cytochrome P450, family 1, subfamily
 B, polypeptide 1	0.00039733772998888823
GO:0030336: negative regulation of cell migration	DACH1	dachshund family transcription factor
1	0.0020864773603336663

GO:0030336: negative regulation of cell migration DAG1 dystroglycan 1 (dystrophin-associated glycoprotein 1) 0.00036120985217686394
GO:0030336: negative regulation of cell migration DRD2 dopamine receptor D2 -0.00023385001518940208
GO:0030336: negative regulation of cell migration ENG endoglin 0.0008254162355259502
GO:0030336: negative regulation of cell migration KANK1 KN motif and ankyrin repeat domains 1 0.002530448940681982
GO:0030336: negative regulation of cell migration NF2 neurofibromin 2 (merlin) -0.0012705218028460686
GO:0030336: negative regulation of cell migration PTEN phosphatase and tensin homolog 1.72558522944505e-5
GO:0030336: negative regulation of cell migration PTPRJ protein tyrosine phosphatase, receptor type, J -0.00033823014031967403
GO:0030336: negative regulation of cell migration SERPINE1 serpin peptidase inhibitor, clade E (nexin, plasminogen activator inhibitor type 1), member 1 0.00011234506285016844
GO:0030336: negative regulation of cell migration SFRP1 secreted frizzled-related protein 1 0.0012798458679897998
GO:0030336: negative regulation of cell migration SHH sonic hedgehog 0.0005990921539456029
GO:0030336: negative regulation of cell migration SLIT2 slit homolog 2 (Drosophila) -0.0016182399277552256
GO:0030336: negative regulation of cell migration STK24 serine/threonine kinase 24 0.002691671557617778
GO:0030336: negative regulation of cell migration SULF1 sulfatase 1 -0.0008044807316197398
GO:0030336: negative regulation of cell migration WNT4 wingless-type MMTV integration site family, member 4 -0.00025158545905789003
GO:0030509: BMP signaling pathway ACVRL1 activin A receptor type II-like 1 0.0019435026586675326
GO:0030509: BMP signaling pathway BMP4 bone morphogenetic protein 4 -0.00032207533292147463
GO:0030509: BMP signaling pathway BMP7 bone morphogenetic protein 7 0.000856200155103951
GO:0030509: BMP signaling pathway DLX5 distal-less homeobox 5 -0.003285025452064135
GO:0030509: BMP signaling pathway EGR1 early growth response 1 0.0010933469015226144
GO:0030509: BMP signaling pathway ENG endoglin 0.0008263497207037055
GO:0030509: BMP signaling pathway FGF8 fibroblast growth factor 8 (androgen-induced) 0.0009834225103412014
GO:0030509: BMP signaling pathway GATA4 GATA binding protein 4 -0.0010940148851977716
GO:0030509: BMP signaling pathway GDF3 growth differentiation factor 3 -0.0018837184756084157
GO:0030509: BMP signaling pathway LEF1 lymphoid enhancer-binding factor 1 -9.961637110677028e-5
GO:0030509: BMP signaling pathway SKI SKI proto-oncogene -0.0006709456968314366
GO:0030513: positive regulation of BMP signaling pathway ACVRL1 activin A receptor type II-like 1 0.001942498688367108
GO:0030513: positive regulation of BMP signaling pathway BMP4 bone morphogenetic protein 4 -0.00032236645373537467
GO:0030513: positive regulation of BMP signaling pathway ENG endoglin 0.0008258856499784679
GO:0030513: positive regulation of BMP signaling pathway GATA4 GATA binding protein 4 -0.0010939836837037238
GO:0030513: positive regulation of BMP signaling pathway GATA6 GATA binding protein 6 -2.805840818749658e-5
GO:0030513: positive regulation of BMP signaling pathway HES1 hes family bHLH transcription factor 1 -0.0009102898038528629
GO:0030513: positive regulation of BMP signaling pathway MSX1 msh homeobox 1 -0.002766961922169556
GO:0030513: positive regulation of BMP signaling pathway NOTCH1 notch 1 0.0005184455403721943
GO:0030513: positive regulation of BMP signaling pathway RBPJ recombination signal binding protein for immunoglobulin kappa J region 0.0009807306261139035
GO:0030513: positive regulation of BMP signaling pathway SOX11 SRY (sex determining region Y)-box 11 -0.00020882416755528308
GO:0030513: positive regulation of BMP signaling pathway SULF1 sulfatase 1 -0.0008052128386622089
GO:0032332: positive regulation of chondrocyte differentiation ACVRL1 activin A receptor type II-like 1 0.0019536518325165403
GO:0032332: positive regulation of chondrocyte differentiation GLI3 GLI family zinc finger 3 -0.00216507680922713
GO:0032332: positive regulation of chondrocyte differentiation IHH indian hedgehog -0.002052110766871769
GO:0032332: positive regulation of chondrocyte differentiation POR P450 (cytochrome) oxidoreductase 0.00046015078850584313
GO:0032332: positive regulation of chondrocyte differentiation SOX9 SRY (sex determining region Y)-box 9 -0.0005210372975415026
GO:0032924: activin receptor signaling pathway ACVRL1 activin A receptor type II-like 1 0.0019500099733605403
GO:0032924: activin receptor signaling pathway INHBA inhibin, beta A -0.0013605193067040265
GO:0032924: activin receptor signaling pathway TGFBR1 transforming growth factor, beta receptor 1 0.00034160273668654585
GO:0035313: wound healing, spreading of epidermal cells ACVRL1 activin A receptor type II-like 1 0.0019396857516285296
GO:0035313: wound healing, spreading of epidermal cells ARHGAP24 Rho GTPase activating protein

24	-0.0009324411740603254
GO:0035313: wound healing, spreading of epidermal cells COL5A1	collagen, type V, alpha 1	-4.84
11186035843185e-5
GO:0043535: regulation of blood vessel endothelial cell migration	ACVRL1	activin A receptor ty
pe II-like 1	0.0019411176309128099
GO:0043535: regulation of blood vessel endothelial cell migration	EFNA1	ephrin-A1	-0.00
05207594575803087
GO:0043537: negative regulation of blood vessel endothelial cell migration	ACVRL1	activin A rec
eptor type II-like 1	0.0019350997676852316
GO:0043537: negative regulation of blood vessel endothelial cell migration	CSNK2B	casein kinase
2, beta polypeptide	0.001520208378070712
GO:0043537: negative regulation of blood vessel endothelial cell migration	HMGB1	high mobility
group box 1	-0.0007741255427552252
GO:0043537: negative regulation of blood vessel endothelial cell migration	TGFB1	transforming
growth factor, beta 1	-7.219485695222212e-5
GO:0043537: negative regulation of blood vessel endothelial cell migration	THBS1	thrombospondi
n 1	-0.0010305616462237105
GO:0043537: negative regulation of blood vessel endothelial cell migration	VASH1	vasohibin 1
0.0005964130896585947
GO:0045602: negative regulation of endothelial cell differentiation	ACVRL1	activin A receptor ty
pe II-like 1	0.0019557258365855866
GO:0045603: positive regulation of endothelial cell differentiation	ACVRL1	activin A receptor ty
pe II-like 1	0.0019386631618379944
GO:0045603: positive regulation of endothelial cell differentiation	ALOX12	arachidonate 12-lipox
ygenase -0.0019479874804598795
GO:0045603: positive regulation of endothelial cell differentiation	BMP4	bone morphogenetic pr
otein 4 -0.0003214854620348939
GO:0045603: positive regulation of endothelial cell differentiation	CTNNB1	catenin (cadherin-ass
ociated protein), beta 1, 88kDa -0.00011743315777938884
GO:0045603: positive regulation of endothelial cell differentiation	NOTCH1	notch 1 0.00051665745
87866227
GO:0045766: positive regulation of angiogenesis ACVRL1	activin A receptor type II-like 1	0.001
9484576661752935
GO:0045766: positive regulation of angiogenesis ADM	adrenomedullin	0.002243134031581997
GO:0045766: positive regulation of angiogenesis ALOX12	arachidonate 12-lipoxygenase	-0.0019585587
277376466
GO:0045766: positive regulation of angiogenesis C3	complement component 3	0.002029742039891387
GO:0045766: positive regulation of angiogenesis CX3CL1	chemokine (C-X3-C motif) ligand 1	0.002
1230040437213837
GO:0045766: positive regulation of angiogenesis CX3CR1	chemokine (C-X3-C motif) receptor 1	-0.00
0858762972197624
GO:0045766: positive regulation of angiogenesis CYP1B1	cytochrome P450, family 1, subfamily B, polyp
eptide 1	0.0003984165230839195
GO:0045766: positive regulation of angiogenesis ECM1	extracellular matrix protein 1	-0.0015619311
25469337
GO:0045766: positive regulation of angiogenesis GATA2	GATA binding protein 2	-0.000452982338543114
43
GO:0045766: positive regulation of angiogenesis GATA4	GATA binding protein 4	-0.001097592728648809
GO:0045766: positive regulation of angiogenesis GATA6	GATA binding protein 6	-2.7957635891901604e-
5
GO:0045766: positive regulation of angiogenesis GREM1	gremlin 1, DAN family BMP antagonist	-0.00
08303460376319974
GO:0045766: positive regulation of angiogenesis HIF1A	hypoxia inducible factor 1, alpha subunit (ba
sic helix-loop-helix transcription factor)	-0.0006478370845735985
GO:0045766: positive regulation of angiogenesis HIPK2	homeodomain interacting protein kinase 2
0.0007724192858138798
GO:0045766: positive regulation of angiogenesis HMOX1	heme oxygenase (decycling) 1	-0.0002182584
4159092643
GO:0045766: positive regulation of angiogenesis ISL1	ISL LIM homeobox 1	7.865100337366006e-5
GO:0045766: positive regulation of angiogenesis NR2E1	nuclear receptor subfamily 2, group E, member
1	-0.0029816642478921076
GO:0045766: positive regulation of angiogenesis PRKCA	protein kinase C, alpha -5.865913487281216e-6
GO:0045766: positive regulation of angiogenesis SERPINE1	serpin peptidase inhibitor, clade E
(nexin, plasminogen activator inhibitor type 1), member 1	0.00011210625265471011
GO:0045766: positive regulation of angiogenesis SPHK1	sphingosine kinase 1	0.0018153777778733007
GO:0045766: positive regulation of angiogenesis THBS1	thrombospondin 1	-0.001038422913631274
6
GO:0045766: positive regulation of angiogenesis TWIST1	twist family bHLH transcription factor 1
-0.001345317783625186
GO:0045766: positive regulation of angiogenesis VEGFA	vascular endothelial growth factor A	0.000
5970545210815664
GO:0045766: positive regulation of angiogenesis VEGFC	vascular endothelial growth factor C	-0.00
33721779956947967
GO:0045766: positive regulation of angiogenesis WNT5A	wingless-type MMTV integration site family, m
ember 5A	-0.0006692282805074454
GO:0045893: positive regulation of transcription, DNA-templated ACVRL1	activin A receptor type II-li
ke 1	0.0019424699523251578
GO:0045893: positive regulation of transcription, DNA-templated AGT	angiotensinogen (serpin pepti
dase inhibitor, clade A, member 8)	-0.0011319510858498062
GO:0045893: positive regulation of transcription, DNA-templated ALX1	ALX homeobox 1	0.00229043634
09177483

GO:0045893: positive regulation of transcription, DNA-templated AR      androgen receptor       0.002
636593340557387
GO:0045893: positive regulation of transcription, DNA-templated ATAD2   ATPase family, AAA domain con
taining 2       -4.7890110846955696e-5
GO:0045893: positive regulation of transcription, DNA-templated ATF5    activating transcription fact
or 5    -0.002733053347596575
GO:0045893: positive regulation of transcription, DNA-templated AXIN1   axin 1  -0.000732497399041859
9
GO:0045893: positive regulation of transcription, DNA-templated BLM     Bloom syndrome, RecQ helicase
-like   0.0005237542326881842
GO:0045893: positive regulation of transcription, DNA-templated BMP4    bone morphogenetic protein 4
-0.0003223391468437128
GO:0045893: positive regulation of transcription, DNA-templated BMP7    bone morphogenetic protein 7
0.0008551480720348396
GO:0045893: positive regulation of transcription, DNA-templated BRCA2   breast cancer 2, early onset
-1.0194446723873534e-5
GO:0045893: positive regulation of transcription, DNA-templated CCNE1   cyclin E1       0.00037051512
47849759
GO:0045893: positive regulation of transcription, DNA-templated CDKN2A  cyclin-dependent kinase inhib
itor 2A 0.0017565407423127842
GO:0045893: positive regulation of transcription, DNA-templated CITED1  Cbp/p300-interacting transact
ivator, with Glu/Asp-rich carboxy-terminal domain, 1    0.0028158909298131732
GO:0045893: positive regulation of transcription, DNA-templated CLOCK   clock circadian regulator
0.00020014388638220902
GO:0045893: positive regulation of transcription, DNA-templated COL1A1  collagen, type I, alpha 1
-0.0005254580146218143
GO:0045893: positive regulation of transcription, DNA-templated CREB1   cAMP responsive element bindi
ng protein 1    0.0006576257344525607
GO:0045893: positive regulation of transcription, DNA-templated CTNNB1  catenin (cadherin-associated
protein), beta 1, 88kDa        -0.00011693183698617328
GO:0045893: positive regulation of transcription, DNA-templated DLX5    distal-less homeobox 5  -0.00
32809466100890564
GO:0045893: positive regulation of transcription, DNA-templated DNAJC2  DnaJ (Hsp40) homolog, subfami
ly C, member 2  -0.00098306051359705
GO:0045893: positive regulation of transcription, DNA-templated E2F1    E2F transcription factor 1
0.002096015752039045
GO:0045893: positive regulation of transcription, DNA-templated E2F3    E2F transcription factor 3
0.0021602977746636724
GO:0045893: positive regulation of transcription, DNA-templated EGR1    early growth response 1 0.001
092277571488665
GO:0045893: positive regulation of transcription, DNA-templated EGR2    early growth response 2 0.001
437372938349719
GO:0045893: positive regulation of transcription, DNA-templated ERBB4   v-erb-b2 avian erythroblastic
leukemia viral oncogene homolog 4       -0.0001825305287042662
GO:0045893: positive regulation of transcription, DNA-templated ESR1    estrogen receptor 1     -0.00
09472384920453688
GO:0045893: positive regulation of transcription, DNA-templated ETS2    v-ets avian erythroblastosis
virus E26 oncogene homolog 2   -0.0010147518385566
GO:0045893: positive regulation of transcription, DNA-templated FGF7    fibroblast growth factor 7
0.0006332387776418407
GO:0045893: positive regulation of transcription, DNA-templated FOXA2   forkhead box A2 -1.6742221533
040816e-5
GO:0045893: positive regulation of transcription, DNA-templated FOXC1   forkhead box C1 -2.2422478273
078127e-5
GO:0045893: positive regulation of transcription, DNA-templated FOXC2   forkhead box C2 (MFH-1, mesen
chyme forkhead 1)       0.0017469206154163354
GO:0045893: positive regulation of transcription, DNA-templated FOXH1   forkhead box H1 -0.0013699111
435599696
GO:0045893: positive regulation of transcription, DNA-templated FOXM1   forkhead box M1 0.00019684151
868649375
GO:0045893: positive regulation of transcription, DNA-templated FOXO1   forkhead box O1 0.00179888866
80514542
GO:0045893: positive regulation of transcription, DNA-templated FOXO3   forkhead box O3 0.00119908040
52633506
GO:0045893: positive regulation of transcription, DNA-templated FZD7    frizzled class receptor 7
0.0010694453897585477
GO:0045893: positive regulation of transcription, DNA-templated GATA3   GATA binding protein 3  -3.87
9720197696244e-5
GO:0045893: positive regulation of transcription, DNA-templated GATA4   GATA binding protein 4  -0.00
1093886335399841
GO:0045893: positive regulation of transcription, DNA-templated GLI1    GLI family zinc finger 1
-0.0012959183367459673
GO:0045893: positive regulation of transcription, DNA-templated GLI2    GLI family zinc finger 2
0.0018520766205114224
GO:0045893: positive regulation of transcription, DNA-templated GLI3    GLI family zinc finger 3
-0.0021497679982375845
GO:0045893: positive regulation of transcription, DNA-templated HDAC2   histone deacetylase 2   -0.00
12065957095432295
GO:0045893: positive regulation of transcription, DNA-templated HIF1A   hypoxia inducible factor 1, a
lpha subunit (basic helix-loop-helix transcription factor)      -0.0006460986295605767
GO:0045893: positive regulation of transcription, DNA-templated HINFP   histone H4 transcription fact

or	0.0009917191211945696
GO:0045893: positive regulation of transcription, DNA-templated HIPK2	homeodomain interacting protein kinase 2	0.0007703154598295995
GO:0045893: positive regulation of transcription, DNA-templated HMGA1	high mobility group AT-hook 1	-0.0003185526900277931
GO:0045893: positive regulation of transcription, DNA-templated HMGA2	high mobility group AT-hook 2	0.001507866954171934
GO:0045893: positive regulation of transcription, DNA-templated HMGB2	high mobility group box 2	0.00030640789679783973
GO:0045893: positive regulation of transcription, DNA-templated ID2	inhibitor of DNA binding 2, dominant negative helix-loop-helix protein	6.0147338861706245e-5
GO:0045893: positive regulation of transcription, DNA-templated IFNA2	interferon, alpha 2	-0.0017160038228046426
GO:0045893: positive regulation of transcription, DNA-templated IGF1	insulin-like growth factor 1 (somatomedin C)	0.0001306756212132876
GO:0045893: positive regulation of transcription, DNA-templated IL4	interleukin 4	0.0002595043823436951
GO:0045893: positive regulation of transcription, DNA-templated INHBA	inhibin, beta A -0.001351585042189352
GO:0045893: positive regulation of transcription, DNA-templated INSR	insulin receptor	-0.0013621044597654384
GO:0045893: positive regulation of transcription, DNA-templated IRAK1	interleukin-1 receptor-associated kinase 1	-0.0016074433700917169
GO:0045893: positive regulation of transcription, DNA-templated IRF6	interferon regulatory factor 6	-0.0016923647209914836
GO:0045893: positive regulation of transcription, DNA-templated IRF7	interferon regulatory factor 7	-0.0013057533261672223
GO:0045893: positive regulation of transcription, DNA-templated KLF2	Kruppel-like factor 2	-0.0012312907135839464
GO:0045893: positive regulation of transcription, DNA-templated KMT2A	lysine (K)-specific methyltransferase 2A	0.0006600827015158212
GO:0045893: positive regulation of transcription, DNA-templated LEF1	lymphoid enhancer-binding factor 1	-0.00010027652822402453
GO:0045893: positive regulation of transcription, DNA-templated LGR4	leucine-rich repeat containing G protein-coupled receptor 4	0.0003082385651295196
GO:0045893: positive regulation of transcription, DNA-templated LHX1	LIM homeobox 1	-0.000758102673361387
GO:0045893: positive regulation of transcription, DNA-templated LRP5	low density lipoprotein receptor-related protein 5	3.0308567431594196e-5
GO:0045893: positive regulation of transcription, DNA-templated LRP6	low density lipoprotein receptor-related protein 6	0.00014935426329490184
GO:0045893: positive regulation of transcription, DNA-templated MDK	midkine (neurite growth-promoting factor 2)	0.001663442920024833
GO:0045893: positive regulation of transcription, DNA-templated MED1	mediator complex subunit 1	0.0011281820256609692
GO:0045893: positive regulation of transcription, DNA-templated MEF2C	myocyte enhancer factor 2C	0.0009725002200719218
GO:0045893: positive regulation of transcription, DNA-templated MYC	v-myc avian myelocytomatosis viral oncogene homolog -0.0011357788935995239
GO:0045893: positive regulation of transcription, DNA-templated NOTCH1	notch 1 0.0005183413877689854
GO:0045893: positive regulation of transcription, DNA-templated NPAT	nuclear protein, ataxia-telangiectasia locus -0.000709164538014839
GO:0045893: positive regulation of transcription, DNA-templated NR1H3	nuclear receptor subfamily 1, group H, member 3	0.0008004274557348049
GO:0045893: positive regulation of transcription, DNA-templated NRIP1	nuclear receptor interacting protein 1	0.0010704948635502874
GO:0045893: positive regulation of transcription, DNA-templated PAX2	paired box 2	-0.0016127127247679831
GO:0045893: positive regulation of transcription, DNA-templated PAX3	paired box 3	-0.0029431297390478763
GO:0045893: positive regulation of transcription, DNA-templated PAX6	paired box 6	0.0019341878361513528
GO:0045893: positive regulation of transcription, DNA-templated PAX8	paired box 8	0.0009476601432854526
GO:0045893: positive regulation of transcription, DNA-templated PLCB1	phospholipase C, beta 1 (phosphoinositide-specific)	0.0001535466408304212
GO:0045893: positive regulation of transcription, DNA-templated POU1F1	POU class 1 homeobox 1	0.00013727846243815698
GO:0045893: positive regulation of transcription, DNA-templated PROX1	prospero homeobox 1	0.0011205378911210578
GO:0045893: positive regulation of transcription, DNA-templated PSRC1	proline/serine-rich coiled-coil 1	0.000646659188442912
GO:0045893: positive regulation of transcription, DNA-templated PTCH1	patched 1	-6.600176165707593e-5
GO:0045893: positive regulation of transcription, DNA-templated RB1	retinoblastoma 1	-0.0014928297893234354
GO:0045893: positive regulation of transcription, DNA-templated RET	ret proto-oncogene	-0.0004908191583948984
GO:0045893: positive regulation of transcription, DNA-templated RNF187	ring finger protein 187 -0.00222445660235020З
GO:0045893: positive regulation of transcription, DNA-templated RREB1	ras responsive element bindin

g protein 1     -0.001980417869329789
GO:0045893: positive regulation of transcription, DNA-templated SALL1   spalt-like transcription fact
or 1    -0.0024350576906513022
GO:0045893: positive regulation of transcription, DNA-templated SFRP1   secreted frizzled-related pro
tein 1  0.001280139517605567
GO:0045893: positive regulation of transcription, DNA-templated SHH     sonic hedgehog  0.00059932514
29798957
GO:0045893: positive regulation of transcription, DNA-templated SIX1    SIX homeobox 1  -0.0018685348
885844179
GO:0045893: positive regulation of transcription, DNA-templated SMARCD3 SWI/SNF related, matrix assoc
iated, actin dependent regulator of chromatin, subfamily d, member 3    0.00034521483273301535
GO:0045893: positive regulation of transcription, DNA-templated SOX11   SRY (sex determining region
 Y)-box 11      -0.00020913766128075308
GO:0045893: positive regulation of transcription, DNA-templated SOX18   SRY (sex determining region
 Y)-box 18      0.0012876607654467279
GO:0045893: positive regulation of transcription, DNA-templated SOX4    SRY (sex determining region
 Y)-box 4       -3.847273016829247e-5
GO:0045893: positive regulation of transcription, DNA-templated SOX9    SRY (sex determining region
 Y)-box 9       -0.0005182236558763613
GO:0045893: positive regulation of transcription, DNA-templated TBX21   T-box 21        0.00039829661
30727617
GO:0045893: positive regulation of transcription, DNA-templated TBX3    T-box 3 0.0012256285385466954
GO:0045893: positive regulation of transcription, DNA-templated TFAP2A  transcription factor AP-2 alp
ha (activating enhancer binding protein 2 alpha)        0.00047961282795407864
GO:0045893: positive regulation of transcription, DNA-templated TGFB1   transforming growth factor, b
eta 1   -7.28896885272286e-5
GO:0045893: positive regulation of transcription, DNA-templated TGFB3   transforming growth factor, b
eta 3   -0.0018220658739433058
GO:0045893: positive regulation of transcription, DNA-templated TGFBR1  transforming growth factor, b
eta receptor 1  0.0003395534066790807
GO:0045893: positive regulation of transcription, DNA-templated TP53    tumor protein p53       0.001
1745536935249116
GO:0045893: positive regulation of transcription, DNA-templated TP73    tumor protein p73       0.001
0271237257381714
GO:0045893: positive regulation of transcription, DNA-templated TRIM16  tripartite motif containing 1
6       -0.001487624192465426
GO:0045893: positive regulation of transcription, DNA-templated WNT1    wingless-type MMTV integratio
n site family, member 1 0.0007863796247952716
GO:0045893: positive regulation of transcription, DNA-templated WNT4    wingless-type MMTV integratio
n site family, member 4 -0.0002516132220950705
GO:0045893: positive regulation of transcription, DNA-templated WNT5A   wingless-type MMTV integratio
n site family, member 5A        -0.0006668628482081898
GO:0045893: positive regulation of transcription, DNA-templated WNT7A   wingless-type MMTV integratio
n site family, member 7A        1.953676721406495e-5
GO:0045893: positive regulation of transcription, DNA-templated WT1     Wilms tumor 1   -0.0005064029
700403652
GO:0045944: positive regulation of transcription from RNA polymerase II promoter        ACVRL1  activ
in A receptor type II-like 1    0.0019443040642050959
GO:0045944: positive regulation of transcription from RNA polymerase II promoter        ADIRF   adipo
genesis regulatory factor       -0.0001664788414672766
GO:0045944: positive regulation of transcription from RNA polymerase II promoter        AKT1    v-akt
murine thymoma viral oncogene homolog 1 0.0007307987633961783
GO:0045944: positive regulation of transcription from RNA polymerase II promoter        AKT2    v-akt
murine thymoma viral oncogene homolog 2 -0.0006260107247967154
GO:0045944: positive regulation of transcription from RNA polymerase II promoter        ALX1    ALX h
omeobox 1       0.0022925475500006408
GO:0045944: positive regulation of transcription from RNA polymerase II promoter        AR      andro
gen receptor    0.002638822738909426
GO:0045944: positive regulation of transcription from RNA polymerase II promoter        ASCL1   achae
te-scute family bHLH transcription factor 1     -0.0014328599434172361
GO:0045944: positive regulation of transcription from RNA polymerase II promoter        ATAD2   ATPas
e family, AAA domain containing 2       -4.7625349177237585e-5
GO:0045944: positive regulation of transcription from RNA polymerase II promoter        ATAD2B  ATPas
e family, AAA domain containing 2B      -0.0006257229810351443
GO:0045944: positive regulation of transcription from RNA polymerase II promoter        ATF5    activ
ating transcription factor 5    -0.0027356679899733227
GO:0045944: positive regulation of transcription from RNA polymerase II promoter        BACH1   BTB a
nd CNC homology 1, basic leucine zipper transcription factor 1  0.0004854033082600651
GO:0045944: positive regulation of transcription from RNA polymerase II promoter        BATF    basic
leucine zipper transcription factor, ATF-like   0.0020395579562093196
GO:0045944: positive regulation of transcription from RNA polymerase II promoter        BCL11A  B-cel
l CLL/lymphoma 11A (zinc finger protein)        -0.000977247548991509
GO:0045944: positive regulation of transcription from RNA polymerase II promoter        BMP4    bone
 morphogenetic protein 4        -0.0003226663399600546
GO:0045944: positive regulation of transcription from RNA polymerase II promoter        BMP7    bone
 morphogenetic protein 7        0.0008558708714268644
GO:0045944: positive regulation of transcription from RNA polymerase II promoter        BRD4    bromo
domain containing 4     0.0005538715296433384
GO:0045944: positive regulation of transcription from RNA polymerase II promoter        CCNC    cycli
n C     0.0005675530115576571
GO:0045944: positive regulation of transcription from RNA polymerase II promoter        CDK5RAP3

CDK5 regulatory subunit associated protein 3    0.0011383878037459394
GO:0045944: positive regulation of transcription from RNA polymerase II promoter        CDK8    cycli
n-dependent kinase 8    0.002613265909024936
GO:0045944: positive regulation of transcription from RNA polymerase II promoter        CDKN2A  cycli
n-dependent kinase inhibitor 2A 0.0017585356679328027
GO:0045944: positive regulation of transcription from RNA polymerase II promoter        CDKN2B  cycli
n-dependent kinase inhibitor 2B (p15, inhibits CDK4)    -0.0018359023230596614
GO:0045944: positive regulation of transcription from RNA polymerase II promoter        CEBPB   CCAA
T/enhancer binding protein (C/EBP), beta        -0.00027402738146546456
GO:0045944: positive regulation of transcription from RNA polymerase II promoter        CKAP2   cytos
keleton associated protein 2    4.914027723800865e-5
GO:0045944: positive regulation of transcription from RNA polymerase II promoter        CLOCK   clock
circadian regulator     0.0002001297313635757
GO:0045944: positive regulation of transcription from RNA polymerase II promoter        CREB1   cAMP
 responsive element binding protein 1   0.0006584918070843077
GO:0045944: positive regulation of transcription from RNA polymerase II promoter        CTNNB1  caten
in (cadherin-associated protein), beta 1, 88kDa -0.00011698352942197965
GO:0045944: positive regulation of transcription from RNA polymerase II promoter        CXCL10  chemo
kine (C-X-C motif) ligand 10    6.257547612097433e-5
GO:0045944: positive regulation of transcription from RNA polymerase II promoter        DDX17   DEAD
 (Asp-Glu-Ala-Asp) box helicase 17      0.0017622599096275113
GO:0045944: positive regulation of transcription from RNA polymerase II promoter        DMRT1   doubl
esex and mab-3 related transcription factor 1   0.0015048980570308546
GO:0045944: positive regulation of transcription from RNA polymerase II promoter        DRD2    dopam
ine receptor D2 -0.0002340815848517177
GO:0045944: positive regulation of transcription from RNA polymerase II promoter        E2F1    E2F t
ranscription factor 1   0.002098146047396107
GO:0045944: positive regulation of transcription from RNA polymerase II promoter        E2F4    E2F t
ranscription factor 4, p107/p130-binding        -0.0023461456649955417
GO:0045944: positive regulation of transcription from RNA polymerase II promoter        E2F8    E2F t
ranscription factor 8   0.0017113278291275699
GO:0045944: positive regulation of transcription from RNA polymerase II promoter        EGFR    epide
rmal growth factor receptor     0.0006859607328492664
GO:0045944: positive regulation of transcription from RNA polymerase II promoter        EGR1    early
growth response 1       0.0010933264118176433
GO:0045944: positive regulation of transcription from RNA polymerase II promoter        EGR2    early
growth response 2       0.0014391647228194577
GO:0045944: positive regulation of transcription from RNA polymerase II promoter        ELF5    E74-l
ike factor 5 (ets domain transcription factor)  0.0011713153360883152
GO:0045944: positive regulation of transcription from RNA polymerase II promoter        EN1     engra
iled homeobox 1 -1.5794318398248988e-5
GO:0045944: positive regulation of transcription from RNA polymerase II promoter        ENG     endog
lin     0.0008267683638286391
GO:0045944: positive regulation of transcription from RNA polymerase II promoter        ESR1    estro
gen receptor 1  -0.00094834195459816
GO:0045944: positive regulation of transcription from RNA polymerase II promoter        ETS2    v-ets
avian erythroblastosis virus E26 oncogene homolog 2     -0.0010158235714558571
GO:0045944: positive regulation of transcription from RNA polymerase II promoter        ETV4    ets v
ariant 4        0.00019492280020143088
GO:0045944: positive regulation of transcription from RNA polymerase II promoter        EYA1    EYA t
ranscriptional coactivator and phosphatase 1    5.3112802003076616e-5
GO:0045944: positive regulation of transcription from RNA polymerase II promoter        FGFR2   fibro
blast growth factor receptor 2  0.0007634000106427034
GO:0045944: positive regulation of transcription from RNA polymerase II promoter        FOXA1   forkh
ead box A1      2.7747940359392077e-5
GO:0045944: positive regulation of transcription from RNA polymerase II promoter        FOXA2   forkh
ead box A2      -1.666760328064864e-5
GO:0045944: positive regulation of transcription from RNA polymerase II promoter        FOXC1   forkh
ead box C1      -2.223171321108802e-5
GO:0045944: positive regulation of transcription from RNA polymerase II promoter        FOXC2   forkh
ead box C2 (MFH-1, mesenchyme forkhead 1)       0.0017485811525600038
GO:0045944: positive regulation of transcription from RNA polymerase II promoter        FOXH1   forkh
ead box H1      -0.0013714971674223676
GO:0045944: positive regulation of transcription from RNA polymerase II promoter        FOXM1   forkh
ead box M1      0.0001972126308271497
GO:0045944: positive regulation of transcription from RNA polymerase II promoter        FOXO1   forkh
ead box O1      0.0018006087849650717
GO:0045944: positive regulation of transcription from RNA polymerase II promoter        FOXO3   forkh
ead box O3      0.0012003389763785048
GO:0045944: positive regulation of transcription from RNA polymerase II promoter        GATA2   GATA
 binding protein 2      -0.0004516560309394683
GO:0045944: positive regulation of transcription from RNA polymerase II promoter        GATA3   GATA
 binding protein 3      -3.9007046427181735e-5
GO:0045944: positive regulation of transcription from RNA polymerase II promoter        GATA4   GATA
 binding protein 4      -0.001095011394144148
GO:0045944: positive regulation of transcription from RNA polymerase II promoter        GATA6   GATA
 binding protein 6      -2.817318758581491e-5
GO:0045944: positive regulation of transcription from RNA polymerase II promoter        GDNF    glial
cell derived neurotrophic factor        0.00044485219823784316
GO:0045944: positive regulation of transcription from RNA polymerase II promoter        GLI1    GLI f
amily zinc finger 1     -0.001297082557194222

GO:0045944: positive regulation of transcription from RNA polymerase II promoter	GLI2	GLI family zinc finger 2	0.0018539204116941605
GO:0045944: positive regulation of transcription from RNA polymerase II promoter	GLI3	GLI family zinc finger 3	-0.002152105633066085З
GO:0045944: positive regulation of transcription from RNA polymerase II promoter	GREM1	gremlin 1, DAN family BMP antagonist -0.0008275668341616589
GO:0045944: positive regulation of transcription from RNA polymerase II promoter	GRHL2	grainyhead-like 2 (Drosophila)	0.0010069632332025888
GO:0045944: positive regulation of transcription from RNA polymerase II promoter	GSK3B	glycogen synthase kinase 3 beta	0.0015494892993057783
GO:0045944: positive regulation of transcription from RNA polymerase II promoter	HDAC2	histone deacetylase 2	-0.0012074866134420422
GO:0045944: positive regulation of transcription from RNA polymerase II promoter	HES1	hes family bHLH transcription factor 1	-0.0009111680624738234
GO:0045944: positive regulation of transcription from RNA polymerase II promoter	HEY2	hes-related family bHLH transcription factor with YRPW motif 2	0.0026995028122499573
GO:0045944: positive regulation of transcription from RNA polymerase II promoter	HEYL	hes-related family bHLH transcription factor with YRPW motif-like	0.0006777681632462486
GO:0045944: positive regulation of transcription from RNA polymerase II promoter	HIF1A	hypoxia inducible factor 1, alpha subunit (basic helix-loop-helix transcription factor)	-0.0006466180484311898
GO:0045944: positive regulation of transcription from RNA polymerase II promoter	HIPK2	homeodomain interacting protein kinase 2	0.0007709706169694361
GO:0045944: positive regulation of transcription from RNA polymerase II promoter	HMGA1	high mobility group AT-hook 1	-0.00031875032539067813
GO:0045944: positive regulation of transcription from RNA polymerase II promoter	HMGA2	high mobility group AT-hook 2	0.0015092633026494986
GO:0045944: positive regulation of transcription from RNA polymerase II promoter	HMGB1	high mobility group box 1	-0.0007765246540718912
GO:0045944: positive regulation of transcription from RNA polymerase II promoter	HMGB2	high mobility group box 2	0.00030719307885096193
GO:0045944: positive regulation of transcription from RNA polymerase II promoter	HOXA10	homeobox A10 -0.0029825409453013815
GO:0045944: positive regulation of transcription from RNA polymerase II promoter	HOXA5	homeobox A5	0.001066454837108327
GO:0045944: positive regulation of transcription from RNA polymerase II promoter	HOXB1	homeobox B1	0.0036872808076660038
GO:0045944: positive regulation of transcription from RNA polymerase II promoter	HOXD13	homeobox D13 -0.0005925971480680945
GO:0045944: positive regulation of transcription from RNA polymerase II promoter	IFNG	interferon, gamma	-4.8911649086303904e-5
GO:0045944: positive regulation of transcription from RNA polymerase II promoter	IGBP1	immunoglobulin (CD79A) binding protein 1	0.002849935930368595б
GO:0045944: positive regulation of transcription from RNA polymerase II promoter	IGF1	insulin-like growth factor 1 (somatomedin C) 0.00013093438469579288
GO:0045944: positive regulation of transcription from RNA polymerase II promoter	IHH	indian hedgehog	-0.002041046971787435
GO:0045944: positive regulation of transcription from RNA polymerase II promoter	IL4	interleukin 4	0.0002595770430440889
GO:0045944: positive regulation of transcription from RNA polymerase II promoter	INHBA	inhibin, beta A	-0.0013533643395976808
GO:0045944: positive regulation of transcription from RNA polymerase II promoter	IRF7	interferon regulatory factor 7	-0.0013069223190736588
GO:0045944: positive regulation of transcription from RNA polymerase II promoter	ISL1	ISL LIM homeobox 1	7.899059658237337e-5
GO:0045944: positive regulation of transcription from RNA polymerase II promoter	ITGA6	integrin, alpha 6	0.001669402506389009
GO:0045944: positive regulation of transcription from RNA polymerase II promoter	JAG1	jagged 1	0.0017569637542202388
GO:0045944: positive regulation of transcription from RNA polymerase II promoter	JAK2	Janus kinase 2	-3.1764008224896414e-5
GO:0045944: positive regulation of transcription from RNA polymerase II promoter	KLF15	Kruppel-like factor 15	0.00046526701612469186
GO:0045944: positive regulation of transcription from RNA polymerase II promoter	KLF2	Kruppel-like factor 2	-0.001232320746779287l
GO:0045944: positive regulation of transcription from RNA polymerase II promoter	KMT2A	lysine (K)-specific methyltransferase 2A	0.0006607590505708989
GO:0045944: positive regulation of transcription from RNA polymerase II promoter	KMT2D	lysine (K)-specific methyltransferase 2D	-0.00019942560142910436
GO:0045944: positive regulation of transcription from RNA polymerase II promoter	LEF1	lymphoid enhancer-binding factor 1	-0.00010029885240289677
GO:0045944: positive regulation of transcription from RNA polymerase II promoter	LMO4	LIM domain only 4	0.002111834998459686
GO:0045944: positive regulation of transcription from RNA polymerase II promoter	LRP5	low density lipoprotein receptor-related protein 5	3.0262688070737447e-5
GO:0045944: positive regulation of transcription from RNA polymerase II promoter	LRP6	low density lipoprotein receptor-related protein 6	0.00014955188373028678
GO:0045944: positive regulation of transcription from RNA polymerase II promoter	MED1	mediator complex subunit 1	0.0011294964355431272
GO:0045944: positive regulation of transcription from RNA polymerase II promoter	MEF2C	myocyte enhancer factor 2C	0.0009735755401232785

GO:0045944: positive regulation of transcription from RNA polymerase II promoter	MLLT10	myeloid/lymphoid or mixed-lineage leukemia (trithorax homolog, Drosophila); translocated to, 10	-0.000163285583128217
GO:0045944: positive regulation of transcription from RNA polymerase II promoter	MSX1	msh homeobox 1	-0.002769632461183464
GO:0045944: positive regulation of transcription from RNA polymerase II promoter	MYC	v-myc avian myelocytomatosis viral oncogene homolog	-0.00113671807787100
GO:0045944: positive regulation of transcription from RNA polymerase II promoter	NCK1	NCK adaptor protein 1	-0.0007844735590932211
GO:0045944: positive regulation of transcription from RNA polymerase II promoter	NCK2	NCK adaptor protein 2	-0.0002321168743430566
GO:0045944: positive regulation of transcription from RNA polymerase II promoter	NFIB	nuclear factor I/B	0.002919954450386312
GO:0045944: positive regulation of transcription from RNA polymerase II promoter	NOTCH1	notch 1	0.0005188996920113448
GO:0045944: positive regulation of transcription from RNA polymerase II promoter	NR1H3	nuclear receptor subfamily 1, group H, member 3	0.0008010554039037998
GO:0045944: positive regulation of transcription from RNA polymerase II promoter	NR2E1	nuclear receptor subfamily 2, group E, member 1	-0.0029756095966291317
GO:0045944: positive regulation of transcription from RNA polymerase II promoter	NRIP1	nuclear receptor interacting protein 1	0.0010713490281943922
GO:0045944: positive regulation of transcription from RNA polymerase II promoter	PAX2	paired box 2 -0.001614428983625367
GO:0045944: positive regulation of transcription from RNA polymerase II promoter	PAX3	paired box 3 -0.002945886157217531
GO:0045944: positive regulation of transcription from RNA polymerase II promoter	PAX6	paired box 6 0.0019361357343503501
GO:0045944: positive regulation of transcription from RNA polymerase II promoter	PAX8	paired box 8 0.0009483815195790168
GO:0045944: positive regulation of transcription from RNA polymerase II promoter	PDX1	pancreatic and duodenal homeobox 1	0.0002573361634943358
GO:0045944: positive regulation of transcription from RNA polymerase II promoter	PGR	progesterone receptor	0.00035194037278726695
GO:0045944: positive regulation of transcription from RNA polymerase II promoter	PHOX2B	paired-like homeobox 2b	0.000390547296770266
GO:0045944: positive regulation of transcription from RNA polymerase II promoter	PITX2	paired-like homeodomain 2	0.00217656487774886
GO:0045944: positive regulation of transcription from RNA polymerase II promoter	POU1F1	POU class 1 homeobox 1	0.0001374498773826738
GO:0045944: positive regulation of transcription from RNA polymerase II promoter	PRKDC	protein kinase, DNA-activated, catalytic polypeptide -0.001856019476409260
GO:0045944: positive regulation of transcription from RNA polymerase II promoter	PROX1	prospero homeobox 1	0.0011217509861606591
GO:0045944: positive regulation of transcription from RNA polymerase II promoter	RAD21	RAD21 homolog (S. pombe)	-0.00010233687057396999
GO:0045944: positive regulation of transcription from RNA polymerase II promoter	RAF1	Raf-1 proto-oncogene, serine/threonine kinase 0.001499129217626000
GO:0045944: positive regulation of transcription from RNA polymerase II promoter	RARG	retinoic acid receptor, gamma	-0.00231999291252978
GO:0045944: positive regulation of transcription from RNA polymerase II promoter	RB1	retinoblastoma 1	-0.001494326001697112
GO:0045944: positive regulation of transcription from RNA polymerase II promoter	RBPJ	recombination signal binding protein for immunoglobulin kappa J region	0.0009816834092332132
GO:0045944: positive regulation of transcription from RNA polymerase II promoter	RPS6KA1	ribosomal protein S6 kinase, 90kDa, polypeptide 1	-0.0025147744791885513
GO:0045944: positive regulation of transcription from RNA polymerase II promoter	RXRA	retinoid X receptor, alpha	0.001115341961476648
GO:0045944: positive regulation of transcription from RNA polymerase II promoter	SALL1	spalt-like transcription factor 1	-0.002437907997281629
GO:0045944: positive regulation of transcription from RNA polymerase II promoter	SERPINE1	serpin peptidase inhibitor, clade E (nexin, plasminogen activator inhibitor type 1), member 1	0.00011235290357465497
GO:0045944: positive regulation of transcription from RNA polymerase II promoter	SERPINF2	serpin peptidase inhibitor, clade F (alpha-2 antiplasmin, pigment epithelium derived factor), member 2	0.0006588716807296494
GO:0045944: positive regulation of transcription from RNA polymerase II promoter	SHH	sonic hedgehog	0.0006000688860940624
GO:0045944: positive regulation of transcription from RNA polymerase II promoter	SIRT1	sirtuin 1	-1.6533830443268e-6
GO:0045944: positive regulation of transcription from RNA polymerase II promoter	SIRT2	sirtuin 2	-0.0008524578540420826
GO:0045944: positive regulation of transcription from RNA polymerase II promoter	SIX1	SIX homeobox 1	-0.0018703894772010036
GO:0045944: positive regulation of transcription from RNA polymerase II promoter	SIX3	SIX homeobox 3	0.002037706898422995
GO:0045944: positive regulation of transcription from RNA polymerase II promoter	SKI	SKI proto-oncogene	-0.0006706217304763651
GO:0045944: positive regulation of transcription from RNA polymerase II promoter	SMO	smoothened, frizzled class receptor	0.0021397104584296336
GO:0045944: positive regulation of transcription from RNA polymerase II promoter	SOX10	SRY (sex determining region Y)-box 10	0.0001905016879148171

GO:0045944: positive regulation of transcription from RNA polymerase II promoter	SOX11	SRY (sex determining region Y)-box 11	-0.00020912628944135016
GO:0045944: positive regulation of transcription from RNA polymerase II promoter	SOX18	SRY (sex determining region Y)-box 18	0.0012887650551480564
GO:0045944: positive regulation of transcription from RNA polymerase II promoter	SOX4	SRY (sex determining region Y)-box 4	-3.830235497765471e-5
GO:0045944: positive regulation of transcription from RNA polymerase II promoter	SOX9	SRY (sex determining region Y)-box 9	-0.0005187298703103376
GO:0045944: positive regulation of transcription from RNA polymerase II promoter	SPDEF	SAM pointed domain containing ETS transcription factor	0.0031376050999923416
GO:0045944: positive regulation of transcription from RNA polymerase II promoter	STAT5A	signal transducer and activator of transcription 5A	0.001596658780054534
GO:0045944: positive regulation of transcription from RNA polymerase II promoter	TCF7L2	transcription factor 7-like 2 (T-cell specific, HMG-box)	0.0005738109595772473
GO:0045944: positive regulation of transcription from RNA polymerase II promoter	TFAP2A	transcription factor AP-2 alpha (activating enhancer binding protein 2 alpha)	0.0004799991172967249
GO:0045944: positive regulation of transcription from RNA polymerase II promoter	TFAP2C	transcription factor AP-2 gamma (activating enhancer binding protein 2 gamma)	0.001060333029437924
GO:0045944: positive regulation of transcription from RNA polymerase II promoter	TFDP1	transcription factor Dp-1	0.0014450484802161858
GO:0045944: positive regulation of transcription from RNA polymerase II promoter	TGFB1	transforming growth factor, beta 1	-7.302009650972837e-5
GO:0045944: positive regulation of transcription from RNA polymerase II promoter	TGFB3	transforming growth factor, beta 3	-0.0018241169280980704
GO:0045944: positive regulation of transcription from RNA polymerase II promoter	THRA	thyroid hormone receptor, alpha	0.0007633944560357849
GO:0045944: positive regulation of transcription from RNA polymerase II promoter	THRB	thyroid hormone receptor, beta	0.0019444257709734128
GO:0045944: positive regulation of transcription from RNA polymerase II promoter	TOP2A	topoisomerase (DNA) II alpha 170kDa	-8.251189842138416e-5
GO:0045944: positive regulation of transcription from RNA polymerase II promoter	TP53	tumor protein p53	0.0011759598638691271
GO:0045944: positive regulation of transcription from RNA polymerase II promoter	TP73	tumor protein p73	0.001028138191138277
GO:0045944: positive regulation of transcription from RNA polymerase II promoter	TRPS1	trichorhinophalangeal syndrome I	-0.00017778008644282671
GO:0045944: positive regulation of transcription from RNA polymerase II promoter	TWIST1	twist family bHLH transcription factor 1	-0.0013418321891895209
GO:0045944: positive regulation of transcription from RNA polymerase II promoter	VDR	vitamin D (1,25- dihydroxyvitamin D3) receptor	0.0005636709231557241
GO:0045944: positive regulation of transcription from RNA polymerase II promoter	VEGFA	vascular endothelial growth factor A	0.0005951064225557959
GO:0045944: positive regulation of transcription from RNA polymerase II promoter	WNT1	wingless-type MMTV integration site family, member 1	0.0007871140874052551
GO:0045944: positive regulation of transcription from RNA polymerase II promoter	WNT5A	wingless-type MMTV integration site family, member 5A	-0.0006675786331229682
GO:0045944: positive regulation of transcription from RNA polymerase II promoter	WNT7A	wingless-type MMTV integration site family, member 7A	1.950994738521406e-5
GO:0045944: positive regulation of transcription from RNA polymerase II promoter	WT1	Wilms tumor 1	-0.0005070002401581367
GO:0045944: positive regulation of transcription from RNA polymerase II promoter	WWP2	WW domain containing E3 ubiquitin protein ligase 2	-0.0009573922296456005
GO:0045944: positive regulation of transcription from RNA polymerase II promoter	WWTR1	WW domain containing transcription regulator 1	0.000900658297486782
GO:0045944: positive regulation of transcription from RNA polymerase II promoter	XBP1	X-box binding protein 1	0.0002673236707801498
GO:0045944: positive regulation of transcription from RNA polymerase II promoter	YAP1	Yes-associated protein 1	-0.00016437898165821553
GO:0045944: positive regulation of transcription from RNA polymerase II promoter	YBX1	Y box binding protein 1	-0.0009140689035934036
GO:0045944: positive regulation of transcription from RNA polymerase II promoter	ZNF148	zinc finger protein 148	0.0021044890548721582
GO:0051895: negative regulation of focal adhesion assembly	ACVRL1	activin A receptor type II-like 1	0.001953064965691358
GO:0051895: negative regulation of focal adhesion assembly	APOD	apolipoprotein D	0.0026421423650420162
GO:0051895: negative regulation of focal adhesion assembly	PTEN	phosphatase and tensin homolog	1.6114585803782166e-5
GO:0051895: negative regulation of focal adhesion assembly	THBS1	thrombospondin 1	-0.0010406856223369907
GO:0060836: lymphatic endothelial cell differentiation	ACVRL1	activin A receptor type II-like 1	0.001944657952279955
GO:0060836: lymphatic endothelial cell differentiation	PROX1	prospero homeobox 1	0.001121793651351741
GO:0060836: lymphatic endothelial cell differentiation	SOX18	SRY (sex determining region Y)-box 18	0.001289813463985067
GO:0060840: artery development	ACVRL1	activin A receptor type II-like 1	0.0019561802069634356
GO:0060840: artery development	GLI3	GLI family zinc finger 3	-0.0021687661501958704
GO:0060840: artery development	SHH	sonic hedgehog	0.0006054598683210004
GO:0060841: venous blood vessel development	ACVRL1	activin A receptor type II-like 1	0.0019557258365855866

GO:0061154: endothelial tube morphogenesis      ACVRL1  activin A receptor type II-like 1       0.001
9468136600092449
GO:0061154: endothelial tube morphogenesis      CSNK2B  casein kinase 2, beta polypeptide       0.001
5314226415575787
GO:0061154: endothelial tube morphogenesis      CTNNB1  catenin (cadherin-associated protein), beta
 1, 88kDa       -0.0001169837034554008
GO:0061298: retina vasculature development in camera-type eye   ACVRL1  activin A receptor type II-li
ke 1    0.0019461897695409879
GO:0061298: retina vasculature development in camera-type eye   HIF1A   hypoxia inducible factor 1, a
lpha subunit (basic helix-loop-helix transcription factor)      -0.0006476998418521948
GO:0061298: retina vasculature development in camera-type eye   PDGFRB  platelet-derived growth facto
r receptor, beta polypeptide    -0.0003609549333895535б
GO:0061298: retina vasculature development in camera-type eye   ROM1    retinal outer segment membran
e protein 1     0.002463912437196278
GO:0071560: cellular response to transforming growth factor beta stimulus       ACVRL1  activin A rec
eptor type II-like 1    0.0019262613974889422
GO:0071560: cellular response to transforming growth factor beta stimulus       CAV1    caveolin 1, c
aveolae protein, 22kDa  -0.0005316917174526755
GO:0071560: cellular response to transforming growth factor beta stimulus       COL1A1  collagen, typ
e I, alpha 1    -0.0005185334556705554
GO:0071560: cellular response to transforming growth factor beta stimulus       CX3CR1  chemokine (C-
X3-C motif) receptor 1  -0.0008498669779676429
GO:0071560: cellular response to transforming growth factor beta stimulus       HDAC2   histone deace
tylase 2        -0.001197809655568278
GO:0071560: cellular response to transforming growth factor beta stimulus       MEF2C   myocyte enhan
cer factor 2C   0.0009628080732897107
GO:0071560: cellular response to transforming growth factor beta stimulus       SFRP1   secreted friz
zled-related protein 1  0.0012641309059244078
GO:0071560: cellular response to transforming growth factor beta stimulus       SOX9    SRY (sex dete
rmining region Y)-box 9 -0.0005131145359244439
GO:0071560: cellular response to transforming growth factor beta stimulus       STAR    steroidogenic
acute regulatory protein        0.0008850365768218508
GO:0071560: cellular response to transforming growth factor beta stimulus       TGFB1   transforming
 growth factor, beta 1  -7.196941732189383e-5
GO:0071560: cellular response to transforming growth factor beta stimulus       TGFBR1  transforming
 growth factor, beta receptor 1 0.0003358247940326842
GO:0071560: cellular response to transforming growth factor beta stimulus       WNT4    wingless-type
MMTV integration site family, member 4  -0.0002474900637270028
GO:0071560: cellular response to transforming growth factor beta stimulus       WNT5A   wingless-type
MMTV integration site family, member 5A -0.0006607161693427856
GO:0071560: cellular response to transforming growth factor beta stimulus       WNT7A   wingless-type
MMTV integration site family, member 7A 2.001657918172286e-5
GO:0071773: cellular response to BMP stimulus   ACVRL1  activin A receptor type II-like 1       0.001
9469161414038606
GO:0071773: cellular response to BMP stimulus   BMP4    bone morphogenetic protein 4    -0.0003231866
008556083
GO:0071773: cellular response to BMP stimulus   BMP7    bone morphogenetic protein 7    0.00085672490
57354784
GO:0071773: cellular response to BMP stimulus   DLX5    distal-less homeobox 5  -0.003287992159580728
4
GO:0071773: cellular response to BMP stimulus   GATA3   GATA binding protein 3  -3.99441458484945e-5
GO:0071773: cellular response to BMP stimulus   GATA6   GATA binding protein 6  -2.773494593162445e-5
GO:0071773: cellular response to BMP stimulus   HEYL    hes-related family bHLH transcription factor
 with YRPW motif-like   0.0006787436791520901
GO:0071773: cellular response to BMP stimulus   PHOX2B  paired-like homeobox 2b 0.000390618616344778
GO:0071773: cellular response to BMP stimulus   SFRP1   secreted frizzled-related protein 1     0.001
2856317685286162
GO:2000279: negative regulation of DNA biosynthetic process     ACVRL1  activin A receptor type II-li
ke 1    0.0019437890735744718
GO:2000279: negative regulation of DNA biosynthetic process     DACH1   dachshund family transcriptio
n factor 1      0.002089578808858503
GO:2000279: negative regulation of DNA biosynthetic process     DNAJC2  DnaJ (Hsp40) homolog, subfami
ly C, member 2  -0.0009839168760175261
GO:2000279: negative regulation of DNA biosynthetic process     GJA1    gap junction protein, alpha
 1, 43kDa       -0.000160543432818284
GO:0006508: proteolysis ADAM2   ADAM metallopeptidase domain 2  0.0008093986193826224
GO:0006508: proteolysis AZU1    azurocidin 1    0.0001974591915609088
GO:0006508: proteolysis C2      complement component 2  -0.0003032388728206669
GO:0006508: proteolysis CHMP1A  charged multivesicular body protein 1A  -0.0007779082330697746
GO:0006508: proteolysis CPA4    carboxypeptidase A4     -0.0006964846600072998
GO:0006508: proteolysis CTSK    cathepsin K     -0.00036121391973878447
GO:0006508: proteolysis CUL7    cullin 7        -0.00011845549940195519
GO:0006508: proteolysis ESPL1   extra spindle pole bodies homolog 1 (S. cerevisiae)     0.00019570583
778176644
GO:0006508: proteolysis FAP     fibroblast activation protein, alpha    -0.00048527449866830793
GO:0006508: proteolysis GGH     gamma-glutamyl hydrolase (conjugase, folylpolygammaglutamyl hydrolas
e)      -0.00031457450643210364
GO:0006508: proteolysis HPN     hepsin  0.0031517195395803256
GO:0006508: proteolysis KLK5    kallikrein-related peptidase 5  0.0017293365197943113
GO:0006508: proteolysis KLK7    kallikrein-related peptidase 7  0.0016212867507967709
GO:0006508: proteolysis LRP8    low density lipoprotein receptor-related protein 8, apolipoprotein e

receptor	-0.0009299303650135814
GO:0006508: proteolysis METAP1	methionyl aminopeptidase 1	-0.001013992552932157
GO:0006508: proteolysis MMP2	matrix metallopeptidase 2 (gelatinase A, 72kDa gelatinase, 72kDa type IV collagenase) -0.0011276573466487822
GO:0006508: proteolysis MMP20	matrix metallopeptidase 20	-0.00242650013198353 1
GO:0006508: proteolysis MMP24	matrix metallopeptidase 24 (membrane-inserted)	0.001486161432591842
GO:0006508: proteolysis MMP7	matrix metallopeptidase 7 (matrilysin, uterine) 0.0004245508828506054
GO:0006508: proteolysis NAALADL1	N-acetylated alpha-linked acidic dipeptidase-like 1	-0.00 10165593282607669
GO:0006508: proteolysis NDEL1	nudE neurodevelopment protein 1-like 1	-0.0006025021567582331
GO:0006508: proteolysis PCSK2	proprotein convertase subtilisin/kexin type 2	0.0006040018243175647
GO:0006508: proteolysis PGC	progastricsin (pepsinogen C)	0.0013209284413645678
GO:0006508: proteolysis RELN	reelin	0.0015327924562527381
GO:0006508: proteolysis SFRP1	secreted frizzled-related protein 1	0.0012765291137464023
GO:0007155: cell adhesion	ADAM2	ADAM metallopeptidase domain 2	0.0008104134556117931
GO:0007155: cell adhesion	APC	adenomatous polyposis coli	0.0006434239199560317
GO:0007155: cell adhesion	CCL2	chemokine (C-C motif) ligand 2	0.0008159160900474185
GO:0007155: cell adhesion	CD22	CD22 molecule	0.0008367696149558019
GO:0007155: cell adhesion	CD9	CD9 molecule	-0.002583798372750069
GO:0007155: cell adhesion	CDH2	cadherin 2, type 1, N-cadherin (neuronal)	-0.0006440520 366830424
GO:0007155: cell adhesion	CDH3	cadherin 3, type 1, P-cadherin (placental)	-0.0012296514 424777675
GO:0007155: cell adhesion	CIB1	calcium and integrin binding 1 (calmyrin)	5.48341576874 9767e-5
GO:0007155: cell adhesion	COL5A1	collagen, type V, alpha 1	-4.900339602575988e-5
GO:0007155: cell adhesion	CSF3R	colony stimulating factor 3 receptor (granulocyte)	-8.31 2030252776025e-5
GO:0007155: cell adhesion	CTNNB1	catenin (cadherin-associated protein), beta 1, 88kDa	-0.00 011686020462413977
GO:0007155: cell adhesion	CX3CL1	chemokine (C-X3-C motif) ligand 1	0.002114675100483909
GO:0007155: cell adhesion	CX3CR1	chemokine (C-X3-C motif) receptor 1	-0.000856317316241553 7
GO:0007155: cell adhesion	CXCL12	chemokine (C-X-C motif) ligand 12	-0.001207260672067927 3
GO:0007155: cell adhesion	CYP1B1	cytochrome P450, family 1, subfamily B, polypeptide 1	0.000 3974579536730013
GO:0007155: cell adhesion	DSC2	desmocollin 2	0.00046989499160073105
GO:0007155: cell adhesion	DSG2	desmoglein 2	1.680558590932085e-5
GO:0007155: cell adhesion	EFS	embryonal Fyn-associated substrate	0.003904907051297463 2
GO:0007155: cell adhesion	ENG	endoglin	0.0008255748753880439
GO:0007155: cell adhesion	FAP	fibroblast activation protein, alpha	-0.000486835305309508 9
GO:0007155: cell adhesion	GRHL2	grainyhead-like 2 (Drosophila)	0.0010059665278473024
GO:0007155: cell adhesion	HES1	hes family bHLH transcription factor 1	-0.000910111658682948 3
GO:0007155: cell adhesion	ICAM1	intercellular adhesion molecule 1	0.0007288206912410563
GO:0007155: cell adhesion	ITGB4	integrin, beta 4	0.0005795781480714006
GO:0007155: cell adhesion	LAMB2	laminin, beta 2 (laminin S)	-0.001307743419665870 3
GO:0007155: cell adhesion	MFGE8	milk fat globule-EGF factor 8 protein	0.0018029424111920074
GO:0007155: cell adhesion	MYH10	myosin, heavy chain 10, non-muscle	-0.000328695499810490 6
GO:0007155: cell adhesion	PRKCA	protein kinase C, alpha -5.862354995876487e-6
GO:0007155: cell adhesion	PRKX	protein kinase, X-linked	0.0003942252178756825
GO:0007155: cell adhesion	PTK7	protein tyrosine kinase 7	-0.00022510246975984078
GO:0007155: cell adhesion	RELN	reelin	0.001535639956743589
GO:0007155: cell adhesion	ROBO1	roundabout, axon guidance receptor, homolog 1 (Drosophila) -0.002245569196689939
GO:0007155: cell adhesion	ROM1	retinal outer segment membrane protein 1	0.00245861614 3535573
GO:0007155: cell adhesion	SELPLG	selectin P ligand	0.0002865880411651113
GO:0007155: cell adhesion	SPAM1	sperm adhesion molecule 1 (PH-20 hyaluronidase, zona pellucid a binding)	-0.0018981817539297054
GO:0007155: cell adhesion	THBS1	thrombospondin 1	-0.0010349187003224824
GO:0007155: cell adhesion	TNC	tenascin C	0.0007335921996569873
GO:0007338: single fertilization	ADAM2	ADAM metallopeptidase domain 2	0.0008170994606222096
GO:0007338: single fertilization	CD9	CD9 molecule	-0.002608631029406045
GO:0007338: single fertilization	HOXA10	homeobox A10	-0.003008102117917001
GO:0007338: single fertilization	MFGE8	milk fat globule-EGF factor 8 protein	0.00182231378 48137059
GO:0007338: single fertilization	SPAM1	sperm adhesion molecule 1 (PH-20 hyaluronidase, zona pellucida binding)	-0.0019147568464803436
GO:0007339: binding of sperm to zona pellucida	ADAM2	ADAM metallopeptidase domain 2	0.00080521936 3178654
GO:0007339: binding of sperm to zona pellucida	CRISP1	cysteine-rich secretory protein 1	9.152 250557947092e-5
GO:0007339: binding of sperm to zona pellucida	SPAM1	sperm adhesion molecule 1 (PH-20 hyaluronidas e, zona pellucida binding)	-0.0018833847731023396
GO:0007342: fusion of sperm to egg plasma membrane	ADAM2	ADAM metallopeptidase domain 2	0.000 810674945915630 9
GO:0007342: fusion of sperm to egg plasma membrane	CD9	CD9 molecule	-0.002584966462837273

GO:0007342: fusion of sperm to egg plasma membrane      CRISP1  cysteine-rich secretory protein 1
9.054040874410055e-5
GO:0007342: fusion of sperm to egg plasma membrane      ROPN1B  rhophilin associated tail protein 1B
0.0005975338370428848
GO:0007342: fusion of sperm to egg plasma membrane      SERPINA5        serpin peptidase inhibitor, c
lade A (alpha-1 antiproteinase, antitrypsin), member 5  -0.0007106164040647416
GO:0007342: fusion of sperm to egg plasma membrane      SPAM1   sperm adhesion molecule 1 (PH-20 hyal
uronidase, zona pellucida binding)      -0.0018989810548254598
GO:0007342: fusion of sperm to egg plasma membrane      TPST2   tyrosylprotein sulfotransferase 2
0.0007132110022684664
GO:0008542: visual learning     ADAM2   ADAM metallopeptidase domain 2  0.0008098189728719657
GO:0008542: visual learning     CHRNB2  cholinergic receptor, nicotinic, beta 2 (neuronal)      -0.00
07769835682629722
GO:0008542: visual learning     DRD2    dopamine receptor D2    -0.00023370452311025203
GO:0008542: visual learning     HIF1A   hypoxia inducible factor 1, alpha subunit (basic helix-loop-h
elix transcription factor)      -0.0006456804283043314
GO:0008542: visual learning     HTT     huntingtin      -0.0009699089190292248
GO:0008542: visual learning     ITGB1   integrin, beta 1 (fibronectin receptor, beta polypeptide, ant
igen CD29 includes MDF2, MSK12) 0.0024216526595014862
GO:0008542: visual learning     KIT     v-kit Hardy-Zuckerman 4 feline sarcoma viral oncogene homolog
0.00027349569189479657
GO:0008542: visual learning     RGS14   regulator of G-protein signaling 14     -0.001580169586523755
3
GO:0030534: adult behavior      ADAM2   ADAM metallopeptidase domain 2  0.0007804693401182164
GO:0030534: adult behavior      BBS4    Bardet-Biedl syndrome 4 -0.0004923309987886047
GO:0030534: adult behavior      PTEN    phosphatase and tensin homolog  3.092046697754095e-5
GO:0032504: multicellular organism reproduction ADAM2   ADAM metallopeptidase domain 2  0.00081496505
17966669
GO:0032504: multicellular organism reproduction CD9     CD9 molecule    -0.0025997425372637378
GO:0032504: multicellular organism reproduction SPAM1   sperm adhesion molecule 1 (PH-20 hyaluronidas
e, zona pellucida binding)      -0.0019088849472196892
KEGG:05162: Measles     ADAR    adenosine deaminase, RNA-specific       -0.00013671288826628807
KEGG:05162: Measles     AKT1    v-akt murine thymoma viral oncogene homolog 1   0.0007287490684174984
KEGG:05162: Measles     AKT2    v-akt murine thymoma viral oncogene homolog 2   -0.000624665508244975
KEGG:05162: Measles     CCND1   cyclin D1       -0.002625302582796288
KEGG:05162: Measles     CCNE1   cyclin E1       0.00036954137069375513
KEGG:05162: Measles     CCNE2   cyclin E2       0.001103641397869948
KEGG:05162: Measles     CLEC4M  C-type lectin domain family 4, member M -0.0013869551595511633
KEGG:05162: Measles     CSNK2B  casein kinase 2, beta polypeptide       0.00152461196435286
KEGG:05162: Measles     FAS     Fas cell surface death receptor -3.321992477546166e-5
KEGG:05162: Measles     GSK3B   glycogen synthase kinase 3 beta 0.0015454716412320682
KEGG:05162: Measles     HSPA2   heat shock 70kDa protein 2      -0.00010593951113440409
KEGG:05162: Measles     IFNA2   interferon, alpha 2     -0.0017133409832278798
KEGG:05162: Measles     IFNG    interferon, gamma       -4.8915388042583256e-5
KEGG:05162: Measles     IL12B   interleukin 12B 0.001274568735310308
KEGG:05162: Measles     IL4     interleukin 4   0.00025943445894402944
KEGG:05162: Measles     IRAK1   interleukin-1 receptor-associated kinase 1      -0.001604958793645529
1
KEGG:05162: Measles     IRF7    interferon regulatory factor 7  -0.0013038436638979835
KEGG:05162: Measles     JAK2    Janus kinase 2  -3.2225395219683813e-5
KEGG:05162: Measles     PIK3CD  phosphatidylinositol-4,5-bisphosphate 3-kinase, catalytic subunit del
ta      -0.0007789979071440778
KEGG:05162: Measles     RCHY1   ring finger and CHY zinc finger domain containing 1, E3 ubiquitin pro
tein ligase     -0.0010955495125526656
KEGG:05162: Measles     STAT5A  signal transducer and activator of transcription 5A     0.00159278950
10371636
KEGG:05162: Measles     TNFAIP3 tumor necrosis factor, alpha-induced protein 3  0.0011138320160672033
KEGG:05162: Measles     TNFRSF10C       tumor necrosis factor receptor superfamily, member 10c, decoy
without an intracellular domain 0.00031694623567988476
KEGG:05162: Measles     TP53    tumor protein p53       0.0011722475970647732
KEGG:05162: Measles     TP73    tumor protein p73       0.001025530514262669
KEGG:05164: Influenza A ADAR    adenosine deaminase, RNA-specific       -0.00013830670650294928
KEGG:05164: Influenza A AKT1    v-akt murine thymoma viral oncogene homolog 1   0.0007154224829440941
KEGG:05164: Influenza A AKT2    v-akt murine thymoma viral oncogene homolog 2   -0.000617530756299901
9
KEGG:05164: Influenza A CCL2    chemokine (C-C motif) ligand 2  0.0008002299402729665
KEGG:05164: Influenza A CXCL10  chemokine (C-X-C motif) ligand 10       5.592860593354249e-5
KEGG:05164: Influenza A DNAJB1  DnaJ (Hsp40) homolog, subfamily B, member 1     -0.000233764990300689
4
KEGG:05164: Influenza A FAS     Fas cell surface death receptor -3.269710544663394e-5
KEGG:05164: Influenza A GSK3B   glycogen synthase kinase 3 beta 0.0015211235210667042
KEGG:05164: Influenza A HSPA2   heat shock 70kDa protein 2      -9.445156633811773e-5
KEGG:05164: Influenza A ICAM1   intercellular adhesion molecule 1       0.0007222394142386346
KEGG:05164: Influenza A IFNA2   interferon, alpha 2     -0.0016872938024025943
KEGG:05164: Influenza A IFNG    interferon, gamma       -4.922826925518966e-5
KEGG:05164: Influenza A IL12B   interleukin 12B 0.0012597054317034242
KEGG:05164: Influenza A IRF7    interferon regulatory factor 7  -0.0012855425305344482
KEGG:05164: Influenza A JAK2    Janus kinase 2  -3.703844849291685e-5
KEGG:05164: Influenza A PIK3CD  phosphatidylinositol-4,5-bisphosphate 3-kinase, catalytic subunit del
ta      -0.0007666088691241042
KEGG:05164: Influenza A PML     promyelocytic leukemia  -0.0006740708732245694

KEGG:05164: Influenza A PRKCA   protein kinase C, alpha -6.329865720102846e-6
KEGG:05164: Influenza A RAF1    Raf-1 proto-oncogene, serine/threonine kinase   0.00147224507592719
KEGG:05164: Influenza A TNFRSF10C       tumor necrosis factor receptor superfamily, member 10c, decoy without an intracellular domain 0.00031677110527670975
KEGG:04623: Cytosolic DNA-sensing pathway       ADAR    adenosine deaminase, RNA-specific       -0.00013757563618133957
KEGG:04623: Cytosolic DNA-sensing pathway       CXCL10  chemokine (C-X-C motif) ligand 10       5.9175677163762775e-5
KEGG:04623: Cytosolic DNA-sensing pathway       IFNA2   interferon, alpha 2     -0.0017025244418005946
KEGG:04623: Cytosolic DNA-sensing pathway       IRF7    interferon regulatory factor 7  -0.0012962997825249885
GO:0001649: osteoblast differentiation  ADAR    adenosine deaminase, RNA-specific       -0.0001364734564165748
GO:0001649: osteoblast differentiation  AKT1    v-akt murine thymoma viral oncogene homolog 1   0.0007310432064968364
GO:0001649: osteoblast differentiation  ASF1A   anti-silencing function 1A histone chaperone    0.0010506464179521327
GO:0001649: osteoblast differentiation  BMP4    bone morphogenetic protein 4    -0.0003227616967355372
GO:0001649: osteoblast differentiation  COL1A1  collagen, type I, alpha 1       -0.0005270021809212882
GO:0001649: osteoblast differentiation  DLX5    distal-less homeobox 5  -0.003284447664753777
GO:0001649: osteoblast differentiation  FBL     fibrillarin     -0.00027305741085551145
GO:0001649: osteoblast differentiation  GJA1    gap junction protein, alpha 1, 43kDa    -0.0001623649130394756
GO:0001649: osteoblast differentiation  GLI1    GLI family zinc finger 1        -0.0012973024908317563
GO:0001649: osteoblast differentiation  GLI2    GLI family zinc finger 2        0.0018543058393342199
GO:0001649: osteoblast differentiation  HSD17B4 hydroxysteroid (17-beta) dehydrogenase 4        -0.001163317815344932
GO:0001649: osteoblast differentiation  IGFBP3  insulin-like growth factor binding protein 3    0.0008368225771456331
GO:0001649: osteoblast differentiation  IHH     indian hedgehog -0.002041642297199191
GO:0001649: osteoblast differentiation  LEF1    lymphoid enhancer-binding factor 1      -0.00010026773340694056
GO:0001649: osteoblast differentiation  LGR4    leucine-rich repeat containing G protein-coupled receptor 4  0.0003086131415210505
GO:0001649: osteoblast differentiation  MEF2C   myocyte enhancer factor 2C      0.0009739339404381671
GO:0001649: osteoblast differentiation  SFRP1   secreted frizzled-related protein 1     0.001283292278490762
GO:0001649: osteoblast differentiation  SMO     smoothened, frizzled class receptor     0.00214021025579662
GO:0001649: osteoblast differentiation  TNC     tenascin C      0.0007346243974645873
GO:0001649: osteoblast differentiation  TWIST1  twist family bHLH transcription factor 1        -0.0013423248367231628
GO:0001649: osteoblast differentiation  WWTR1   WW domain containing transcription regulator 1  0.0009013842858677854
GO:0002244: hematopoietic progenitor cell differentiation       ADAR    adenosine deaminase, RNA-specific    -0.00013602220375612045
GO:0002244: hematopoietic progenitor cell differentiation       BMP4    bone morphogenetic protein 4 -0.00032074253230548365
GO:0002244: hematopoietic progenitor cell differentiation       INHBA   inhibin, beta A -0.0013437987085223022
GO:0002244: hematopoietic progenitor cell differentiation       PLEK    pleckstrin      0.0009786768375722964
GO:0002244: hematopoietic progenitor cell differentiation       SFRP1   secreted frizzled-related protein 1  0.0012736910229649459
GO:0002244: hematopoietic progenitor cell differentiation       TGFB1   transforming growth factor, beta 1   -7.285023274196379e-5
GO:0002244: hematopoietic progenitor cell differentiation       TOP2A   topoisomerase (DNA) II alpha 170kDa -8.170358880283567e-5
GO:0002566: somatic diversification of immune receptors via somatic mutation    ADAR    adenosine deaminase, RNA-specific    -0.00013990644784975787
GO:0006382: adenosine to inosine editing        ADAR    adenosine deaminase, RNA-specific       -0.0013990644784975787
GO:0006397: mRNA processing     ADAR    adenosine deaminase, RNA-specific       -0.00013680736123699474
GO:0006397: mRNA processing     JMJD6   jumonji domain containing 6     0.0036057524520759325
GO:0006397: mRNA processing     SF3B2   splicing factor 3b, subunit 2, 145kDa   0.0006868879876014886
GO:0006397: mRNA processing     ZPR1    ZPR1 zinc finger        -0.00021114347508795607
GO:0006606: protein import into nucleus ADAR    adenosine deaminase, RNA-specific       -0.0001393602364594814
GO:0006606: protein import into nucleus HTT     huntingtin      -0.0009425434904552681
GO:0006606: protein import into nucleus KPNB1   karyopherin (importin) beta 1   0.0007528784128011072
GO:0006611: protein export from nucleus ADAR    adenosine deaminase, RNA-specific       -0.00013675380177956796
GO:0006611: protein export from nucleus EGR2    early growth response 2 0.0014334529998805967
GO:0006611: protein export from nucleus GSK3B   glycogen synthase kinase 3 beta 0.0015447044502712784
GO:0006611: protein export from nucleus TGFB1   transforming growth factor, beta 1      -7.245090681910452e-5

GO:0009615: response to virus	ADAR	adenosine deaminase, RNA-specific	-0.00013723254133641102
GO:0009615: response to virus	CCL8	chemokine (C-C motif) ligand 8	-0.0005850600306212153
GO:0009615: response to virus	CFL1	cofilin 1 (non-muscle)	-0.0011566900270993236
GO:0009615: response to virus	CXCL12	chemokine (C-X-C motif) ligand 12	-0.00119025062443047
GO:0009615: response to virus	CXCR4	chemokine (C-X-C motif) receptor 4	0.0007819109697478865
GO:0009615: response to virus	GATA3	GATA binding protein 3	-3.676280958412622e-5
GO:0009615: response to virus	HMGA1	high mobility group AT-hook 1	-0.00031544907302057996
GO:0009615: response to virus	HMGA2	high mobility group AT-hook 2	0.0014888663935048638
GO:0009615: response to virus	IFNG	interferon, gamma	-4.88179650128264e-5
GO:0009615: response to virus	IRF7	interferon regulatory factor 7	-0.001290264334461148
GO:0009615: response to virus	MEF2C	myocyte enhancer factor 2C	0.0009571999464746801
GO:0009615: response to virus	ODC1	ornithine decarboxylase 1	0.0008129549933178983
GO:0009615: response to virus	STMN1	stathmin 1	0.0005468524571078876
GO:0009615: response to virus	TBX21	T-box 21	0.00039470139805897577
GO:0010467: gene expression	ADAR	adenosine deaminase, RNA-specific	-0.000137150401353772
8
GO:0010467: gene expression	AIMP2	aminoacyl tRNA synthetase complex-interacting multifunctional protein 2	0.001207413818874052
GO:0010467: gene expression	AKT1	v-akt murine thymoma viral oncogene homolog 1	0.0007279500947189412
GO:0010467: gene expression	AR	androgen receptor	0.002633079968865364
GO:0010467: gene expression	CCNC	cyclin C	0.0005647110472129126
GO:0010467: gene expression	CDC40	cell division cycle 40	0.004153556308255369
GO:0010467: gene expression	CDK8	cyclin-dependent kinase 8	0.0026058946731098618
GO:0010467: gene expression	CDKN2B	cyclin-dependent kinase inhibitor 2B (p15, inhibits CDK4)	-0.0018302426117802646
GO:0010467: gene expression	DCP1A	decapping mRNA 1A	-0.0011889563362459158
GO:0010467: gene expression	DICER1	dicer 1, ribonuclease type III	-1.6280070019297425e-5
GO:0010467: gene expression	E2F4	E2F transcription factor 4, p107/p130-binding	-0.002339110389203165
GO:0010467: gene expression	EDA	ectodysplasin A	-0.000811106771902646
GO:0010467: gene expression	EIF4G1	eukaryotic translation initiation factor 4 gamma, 1	0.0007399557932371728
GO:0010467: gene expression	ESR1	estrogen receptor 1	-0.0009437023007935864
GO:0010467: gene expression	EZH2	enhancer of zeste 2 polycomb repressive complex 2 subunit	-0.000138227025409417
GO:0010467: gene expression	HDAC2	histone deacetylase 2	-0.0012059666510790795
GO:0010467: gene expression	IGF2BP2	insulin-like growth factor 2 mRNA binding protein 2	-0.0009345954717638299
GO:0010467: gene expression	IGF2BP3	insulin-like growth factor 2 mRNA binding protein 3	-0.00038405624373259354
GO:0010467: gene expression	LARS2	leucyl-tRNA synthetase 2, mitochondrial	0.00046596367780559926
GO:0010467: gene expression	MED1	mediator complex subunit 1	0.0011243407578243
GO:0010467: gene expression	MTERF1	mitochondrial transcription termination factor 1	0.0002856379806586486
GO:0010467: gene expression	MYC	v-myc avian myelocytomatosis viral oncogene homolog	-0.0011346788857881215
GO:0010467: gene expression	NOTCH1	notch 1	0.0005167143356663517
GO:0010467: gene expression	NR1H3	nuclear receptor subfamily 1, group H, member 3	0.000799496232748564
GO:0010467: gene expression	NR2E1	nuclear receptor subfamily 2, group E, member 1	-0.002967461502036658
GO:0010467: gene expression	NRBF2	nuclear receptor binding factor 2	-0.0009485175135698681
GO:0010467: gene expression	NUP153	nucleoporin 153kDa	0.0008206700334699573
GO:0010467: gene expression	PGR	progesterone receptor	0.0003525480759175042
GO:0010467: gene expression	POLR2D	polymerase (RNA) II (DNA directed) polypeptide D	0.0018487785823618483
GO:0010467: gene expression	PRKCA	protein kinase C, alpha	-6.002875484826653e-6
GO:0010467: gene expression	PRKCD	protein kinase C, delta	-0.0011237942402427216
GO:0010467: gene expression	PSMA5	proteasome (prosome, macropain) subunit, alpha type, 5	0.00033036532193111474
GO:0010467: gene expression	PSMD11	proteasome (prosome, macropain) 26S subunit, non-ATPase, 11	-0.0010933626387221026
GO:0010467: gene expression	PSMD13	proteasome (prosome, macropain) 26S subunit, non-ATPase, 13	-6.052002956505356e-5
GO:0010467: gene expression	PTEN	phosphatase and tensin homolog	1.8723213306911515e-5
GO:0010467: gene expression	RARG	retinoic acid receptor, gamma	-0.0023127320494180426
GO:0010467: gene expression	RBBP4	retinoblastoma binding protein 4	-0.001858325761041683
GO:0010467: gene expression	RBPJ	recombination signal binding protein for immunoglobulin kappa J region	0.000979469324246348
GO:0010467: gene expression	RXRA	retinoid X receptor, alpha	0.0011121463300903572
GO:0010467: gene expression	SEH1L	SEH1-like (S. cerevisiae)	-0.0007317701174542198
GO:0010467: gene expression	SERPINE1	serpin peptidase inhibitor, clade E (nexin, plasminogen activator inhibitor type 1), member 1	0.00011305298860409826
GO:0010467: gene expression	SF3B2	splicing factor 3b, subunit 2, 145kDa	0.0006811560016530261
GO:0010467: gene expression	SIRT1	sirtuin 1	-2.5139683459378703e-6
GO:0010467: gene expression	SKI	SKI proto-oncogene	-0.000668966627975757
GO:0010467: gene expression	SMC1A	structural maintenance of chromosomes 1A	-0.0010173157

563004769
GO:0010467: gene expression	SNRPB	small nuclear ribonucleoprotein polypeptides B and B1	0.0006619905665319625
GO:0010467: gene expression	TFDP1	transcription factor Dp-1	0.0014375611998349556
GO:0010467: gene expression	THRA	thyroid hormone receptor, alpha 0.0007613593629614999
GO:0010467: gene expression	THRB	thyroid hormone receptor, beta	0.0019392091276832673
GO:0010467: gene expression	TP53	tumor protein p53	0.0011701994104631075
GO:0010467: gene expression	TPR	translocated promoter region, nuclear basket protein	-0.0006954732631280886
GO:0010467: gene expression	VDR	vitamin D (1,25- dihydroxyvitamin D3) receptor	0.0005629512267216589
GO:0010467: gene expression	WDR77	WD repeat domain 77	0.00013144749149961163
GO:0010467: gene expression	WWTR1	WW domain containing transcription regulator 1	0.0008953087042605957
GO:0010467: gene expression	YAP1	Yes-associated protein 1	-0.00016484178373847886
GO:0010467: gene expression	YBX1	Y box binding protein 1 -0.0009133835980316578
GO:0016553: base conversion or substitution editing	ADAR	adenosine deaminase, RNA-specific	-0.00013990644784975787
GO:0016556: mRNA modification	ADAR	adenosine deaminase, RNA-specific	-0.00013990644784975787
GO:0019221: cytokine-mediated signaling pathway ADAR	adenosine deaminase, RNA-specific	-0.0001372456577960013
GO:0019221: cytokine-mediated signaling pathway CCL2	chemokine (C-C motif) ligand 2	0.0008103532508516601
GO:0019221: cytokine-mediated signaling pathway CSF3R	colony stimulating factor 3 receptor (granulocyte)	-7.926335450691344e-5
GO:0019221: cytokine-mediated signaling pathway CX3CL1	chemokine (C-X3-C motif) ligand 1	0.0020986239607392234
GO:0019221: cytokine-mediated signaling pathway EGR1	early growth response 1 0.0010857561953761554
GO:0019221: cytokine-mediated signaling pathway EIF4G1	eukaryotic translation initiation factor 4 gamma, 1	0.000737172288195492
GO:0019221: cytokine-mediated signaling pathway FLT3	fms-related tyrosine kinase 3	-0.0006749201034360919
GO:0019221: cytokine-mediated signaling pathway ICAM1	intercellular adhesion molecule 1	0.000726516115879625
GO:0019221: cytokine-mediated signaling pathway IFNA2	interferon, alpha 2	-0.0017057107594057289
GO:0019221: cytokine-mediated signaling pathway IFNG	interferon, gamma	-4.9015339485445686e-5
GO:0019221: cytokine-mediated signaling pathway IL12B	interleukin 12B 0.001270339482466138
GO:0019221: cytokine-mediated signaling pathway IL20RA	interleukin 20 receptor, alpha	0.00017454558999419168
GO:0019221: cytokine-mediated signaling pathway IL6ST	interleukin 6 signal transducer 0.001864674808726246
GO:0019221: cytokine-mediated signaling pathway IRF6	interferon regulatory factor 6	-0.0016815704969173608
GO:0019221: cytokine-mediated signaling pathway IRF7	interferon regulatory factor 7	-0.0012985258191372685
GO:0019221: cytokine-mediated signaling pathway JAK2	Janus kinase 2	-3.375622037933417e-5
GO:0019221: cytokine-mediated signaling pathway KIT	v-kit Hardy-Zuckerman 4 feline sarcoma viral oncogene homolog	0.0002689952556001819
GO:0019221: cytokine-mediated signaling pathway KPNA3	karyopherin alpha 3 (importin alpha 4)	-0.000813957505852581
GO:0019221: cytokine-mediated signaling pathway KPNB1	karyopherin (importin) beta 1	0.0007677265884137688
GO:0019221: cytokine-mediated signaling pathway LRP8	low density lipoprotein receptor-related protein 8, apolipoprotein e receptor	-0.0009269519506827206
GO:0019221: cytokine-mediated signaling pathway NEDD4	neural precursor cell expressed, developmentally down-regulated 4, E3 ubiquitin protein ligase	0.0022499782277308183
GO:0019221: cytokine-mediated signaling pathway NUP153	nucleoporin 153kDa	0.0008163710379417002
GO:0019221: cytokine-mediated signaling pathway PML	promyelocytic leukemia	-0.0006776433942854825
GO:0019221: cytokine-mediated signaling pathway PRKCD	protein kinase C, delta -0.0011187482464835319
GO:0019221: cytokine-mediated signaling pathway PTPN1	protein tyrosine phosphatase, non-receptor type 1	-0.0010340136506469674
GO:0019221: cytokine-mediated signaling pathway PTPN2	protein tyrosine phosphatase, non-receptor type 2	-0.00032803946915602357
GO:0019221: cytokine-mediated signaling pathway SEH1L	SEH1-like (S. cerevisiae)	-0.0007295901577404033
GO:0019221: cytokine-mediated signaling pathway TPR	translocated promoter region, nuclear basket protein	-0.00069302240210558
GO:0030218: erythrocyte differentiation ADAR	adenosine deaminase, RNA-specific	-0.0001370087877979017
GO:0030218: erythrocyte differentiation BMP4	bone morphogenetic protein 4	-0.0003172322672196642
GO:0030218: erythrocyte differentiation GATA3	GATA binding protein 3	-3.632909010990918e-5
GO:0030218: erythrocyte differentiation HIPK2	homeodomain interacting protein kinase 2	0.0007597017919034311
GO:0030218: erythrocyte differentiation INHA	inhibin, alpha	0.00021091052704461406
GO:0030218: erythrocyte differentiation INHBA	inhibin, beta A -0.0013218216895317595

GO:0030218: erythrocyte differentiation JAK2    Janus kinase 2  -3.458261394232068e-5
GO:0030218: erythrocyte differentiation KIT     v-kit Hardy-Zuckerman 4 feline sarcoma viral oncogene homolog 0.0002644140262405659
GO:0030218: erythrocyte differentiation LYN     LYN proto-oncogene, Src family tyrosine kinase  -0.0013975350218323366
GO:0030218: erythrocyte differentiation PTPN2   protein tyrosine phosphatase, non-receptor type 2 -0.00032479907528572857
GO:0030218: erythrocyte differentiation THRA    thyroid hormone receptor, alpha 0.0007511726389885196
GO:0031054: pre-miRNA processing        ADAR    adenosine deaminase, RNA-specific       -0.0001365135916260654
GO:0031054: pre-miRNA processing        DICER1  dicer 1, ribonuclease type III  -3.7208271204632496e-5
GO:0035280: miRNA loading onto RISC involved in gene silencing by miRNA ADAR    adenosine deaminase, RNA-specific   -0.0001365135916260654
GO:0035280: miRNA loading onto RISC involved in gene silencing by miRNA DICER1  dicer 1, ribonuclease type III        -3.7208271204632496e-5
GO:0035455: response to interferon-alpha        ADAR    adenosine deaminase, RNA-specific       -0.00013990644784975787
GO:0043066: negative regulation of apoptotic process    ADAR    adenosine deaminase, RNA-specific -0.0001365630742551135
GO:0043066: negative regulation of apoptotic process    AKT1    v-akt murine thymoma viral oncogene homolog 1        0.0007318832915288952
GO:0043066: negative regulation of apoptotic process    ALOX12  arachidonate 12-lipoxygenase    -0.0019565985768180486
GO:0043066: negative regulation of apoptotic process    ANGPT1  angiopoietin 1  0.0009019132492647957
GO:0043066: negative regulation of apoptotic process    ASCL1   achaete-scute family bHLH transcription factor 1     -0.0014349478609823755
GO:0043066: negative regulation of apoptotic process    ASIC2   acid-sensing (proton-gated) ion channel 2    0.0022830086044381874
GO:0043066: negative regulation of apoptotic process    ATF5    activating transcription factor 5 -0.002739075703281903
GO:0043066: negative regulation of apoptotic process    AURKA   aurora kinase A 0.0010097033297251838
GO:0043066: negative regulation of apoptotic process    AVP     arginine vasopressin    -0.0009390430798288734
GO:0043066: negative regulation of apoptotic process    AZU1    azurocidin 1    0.00019823895219863463
GO:0043066: negative regulation of apoptotic process    BARD1   BRCA1 associated RING domain 1  0.001746074914910911
GO:0043066: negative regulation of apoptotic process    BCL2    B-cell CLL/lymphoma 2   -4.964459510846905e-6
GO:0043066: negative regulation of apoptotic process    BFAR    bifunctional apoptosis regulator -0.0003708538608130186
GO:0043066: negative regulation of apoptotic process    BIRC5   baculoviral IAP repeat containing 5 -0.00026048410182925483
GO:0043066: negative regulation of apoptotic process    BMP4    bone morphogenetic protein 4    -0.00032307350129527
GO:0043066: negative regulation of apoptotic process    BNIP3   BCL2/adenovirus E1B 19kDa interacting protein 3       0.002917156946463021
GO:0043066: negative regulation of apoptotic process    BNIP3L  BCL2/adenovirus E1B 19kDa interacting protein 3-like  -5.759533705013386e-5
GO:0043066: negative regulation of apoptotic process    CAMK1D  calcium/calmodulin-dependent protein kinase ID      -0.002591789934411612
GO:0043066: negative regulation of apoptotic process    CDK1    cyclin-dependent kinase 1       0.00031198844568769
GO:0043066: negative regulation of apoptotic process    CFL1    cofilin 1 (non-muscle)  -0.001177005737954231
GO:0043066: negative regulation of apoptotic process    CIB1    calcium and integrin binding 1 (calmyrin)    5.4286365686468054e-5
GO:0043066: negative regulation of apoptotic process    CRYAA   crystallin, alpha A     -0.0006763605736025916
GO:0043066: negative regulation of apoptotic process    CRYAB   crystallin, alpha B     0.000986279586138894
GO:0043066: negative regulation of apoptotic process    DHCR24  24-dehydrocholesterol reductase -0.001710098145002904
GO:0043066: negative regulation of apoptotic process    EGFR    epidermal growth factor receptor 0.0006872254158491331
GO:0043066: negative regulation of apoptotic process    EGR1    early growth response 1 0.0010946950640368608
GO:0043066: negative regulation of apoptotic process    EGR2    early growth response 2 0.0014416586602442342
GO:0043066: negative regulation of apoptotic process    ERBB4   v-erb-b2 avian erythroblastic leukemia viral oncogene homolog 4      -0.00018360934112878727
GO:0043066: negative regulation of apoptotic process    FAS     Fas cell surface death receptor -3.3344210576554394e-5
GO:0043066: negative regulation of apoptotic process    FOXE3   forkhead box E3 0.0010523252216038269
GO:0043066: negative regulation of apoptotic process    FOXO1   forkhead box O1 0.0018028519283239265
GO:0043066: negative regulation of apoptotic process    GATA6   GATA binding protein 6  -2.8040580375928203e-5
GO:0043066: negative regulation of apoptotic process    GDNF    glial cell derived neurotrophic factor       0.00044527566529955603
GO:0043066: negative regulation of apoptotic process    GLI3    GLI family zinc finger 3        -0.00

21552163228767017
GO:0043066: negative regulation of apoptotic process GREM1 gremlin 1, DAN family BMP antagonist
-0.0008291584871450937
GO:0043066: negative regulation of apoptotic process GSK3B glycogen synthase kinase 3 beta 0.001
5515058164287821
GO:0043066: negative regulation of apoptotic process GSTP1 glutathione S-transferase pi 1 0.000
23224553744555254
GO:0043066: negative regulation of apoptotic process HDAC2 histone deacetylase 2 -0.0012085838
811785432
GO:0043066: negative regulation of apoptotic process HMGA2 high mobility group AT-hook 2 0.001
5110520694431187
GO:0043066: negative regulation of apoptotic process HPN hepsin 0.003162522412723068
GO:0043066: negative regulation of apoptotic process IGF1 insulin-like growth factor 1 (somatom
edin C) 0.0001313927588091416
GO:0043066: negative regulation of apoptotic process IGF1R insulin-like growth factor 1 receptor
0.0010731641395217626
GO:0043066: negative regulation of apoptotic process IHH indian hedgehog -0.002043848834352691
GO:0043066: negative regulation of apoptotic process IL4 interleukin 4 0.0002596022801489127
GO:0043066: negative regulation of apoptotic process IL6ST interleukin 6 signal transducer 0.001
8780714790041745
GO:0043066: negative regulation of apoptotic process IL7 interleukin 7 0.0008728000895207232
GO:0043066: negative regulation of apoptotic process IRAK1 interleukin-1 receptor-associated kin
ase 1 -0.001611254775255239
GO:0043066: negative regulation of apoptotic process KIF14 kinesin family member 14 0.000
4867689475413458
GO:0043066: negative regulation of apoptotic process KRT18 keratin 18 -0.001019567446735863
4
GO:0043066: negative regulation of apoptotic process LEF1 lymphoid enhancer-binding factor 1
-0.00010025273476476155
GO:0043066: negative regulation of apoptotic process LEP leptin 0.003196334262655334
GO:0043066: negative regulation of apoptotic process MAD2L1 MAD2 mitotic arrest deficient-like 1
(yeast) 0.00031933451556359144
GO:0043066: negative regulation of apoptotic process MED1 mediator complex subunit 1 0.001
1312947550757776
GO:0043066: negative regulation of apoptotic process MSX1 msh homeobox 1 -0.002773633304305703
GO:0043066: negative regulation of apoptotic process MYC v-myc avian myelocytomatosis viral on
cogene homolog -0.0011379328297025273
GO:0043066: negative regulation of apoptotic process NR2E1 nuclear receptor subfamily 2, group
E, member 1 -0.002979045549123312
GO:0043066: negative regulation of apoptotic process PAX2 paired box 2 -0.001616701654041186
3
GO:0043066: negative regulation of apoptotic process PCNT pericentrin 0.0017896678743523186
GO:0043066: negative regulation of apoptotic process PDGFRB platelet-derived growth factor recept
or, beta polypeptide -0.00036219737673812076
GO:0043066: negative regulation of apoptotic process PHB2 prohibitin 2 -0.000737110001924458
9
GO:0043066: negative regulation of apoptotic process PLK1 polo-like kinase 1 0.00101960923
1091452
GO:0043066: negative regulation of apoptotic process PLK3 polo-like kinase 3 0.00256763538
4412102
GO:0043066: negative regulation of apoptotic process PRKCZ protein kinase C, zeta -0.0016030255
100953586
GO:0043066: negative regulation of apoptotic process PRKDC protein kinase, DNA-activated, cataly
tic polypeptide -0.0018581700433447817
GO:0043066: negative regulation of apoptotic process PRLR prolactin receptor 0.00218635787
5551232
GO:0043066: negative regulation of apoptotic process PSMA5 proteasome (prosome, macropain) subun
it, alpha type, 5 0.00033248937418795086
GO:0043066: negative regulation of apoptotic process PSMD11 proteasome (prosome, macropain) 26S s
ubunit, non-ATPase, 11 -0.0010961632989426244
GO:0043066: negative regulation of apoptotic process PSMD13 proteasome (prosome, macropain) 26S s
ubunit, non-ATPase, 13 -5.9517092312380985e-5
GO:0043066: negative regulation of apoptotic process PTEN phosphatase and tensin homolog 1.672
7331316222732e-5
GO:0043066: negative regulation of apoptotic process RAF1 Raf-1 proto-oncogene, serine/threonin
e kinase 0.0015010131652367302
GO:0043066: negative regulation of apoptotic process RARG retinoic acid receptor, gamma -0.00
23228513270552547
GO:0043066: negative regulation of apoptotic process RPS6KA1 ribosomal protein S6 kinase, 90kDa, p
olypeptide 1 -0.002517688388686801
GO:0043066: negative regulation of apoptotic process SFRP1 secreted frizzled-related protein 1
0.0012846742657332073
GO:0043066: negative regulation of apoptotic process SHH sonic hedgehog 0.0006011234781037242
GO:0043066: negative regulation of apoptotic process SIRT1 sirtuin 1 -1.4205056609002592e-
6
GO:0043066: negative regulation of apoptotic process SMO smoothened, frizzled class receptor
0.002142486239564675
GO:0043066: negative regulation of apoptotic process SOX10 SRY (sex determining region Y)-box 10
0.00019152662733506733
GO:0043066: negative regulation of apoptotic process SOX9 SRY (sex determining region Y)-box 9
-0.0005194356866524715

GO:0043066: negative regulation of apoptotic process    SPHK1   sphingosine kinase 1    0.00181361813
5050493
GO:0043066: negative regulation of apoptotic process    TBX3    T-box 3 0.0012276793959356835
GO:0043066: negative regulation of apoptotic process    TFAP2A  transcription factor AP-2 alpha (acti
vating enhancer binding protein 2 alpha)        0.0004804813256791423
GO:0043066: negative regulation of apoptotic process    THBS1   thrombospondin 1        -0.0010374064
289076868
GO:0043066: negative regulation of apoptotic process    TMBIM6  transmembrane BAX inhibitor motif con
taining 6       -0.0001479160926199905
GO:0043066: negative regulation of apoptotic process    TP53    tumor protein p53       0.00117793383
36137708
GO:0043066: negative regulation of apoptotic process    TWIST1  twist family bHLH transcription facto
r 1     -0.0013438239194713568
GO:0043066: negative regulation of apoptotic process    VEGFA   vascular endothelial growth factor A
0.0005962379960733927
GO:0043066: negative regulation of apoptotic process    WNT5A   wingless-type MMTV integration site f
amily, member 5A        -0.0006685224561495124
GO:0043066: negative regulation of apoptotic process    WNT7A   wingless-type MMTV integration site f
amily, member 7A        1.9431653555381167e-5
GO:0043066: negative regulation of apoptotic process    WT1     Wilms tumor 1   -0.000507846692805235
4
GO:0043066: negative regulation of apoptotic process    XBP1    X-box binding protein 1 0.00026745362
41916085
GO:0044387: negative regulation of protein kinase activity by regulation of protein phosphorylation
ADAR    adenosine deaminase, RNA-specific       -0.00014172340387966556
GO:0044387: negative regulation of protein kinase activity by regulation of protein phosphorylation
CDK5RAP3        CDK5 regulatory subunit associated protein 3    0.0011148389451219034
GO:0045070: positive regulation of viral genome replication     ADAR    adenosine deaminase, RNA-spec
ific    -0.00012748806385473493
GO:0045070: positive regulation of viral genome replication     TOP2A   topoisomerase (DNA) II alpha
 170kDa -0.00010353514325020871
GO:0045071: negative regulation of viral genome replication     ADAR    adenosine deaminase, RNA-spec
ific    -0.00013782742167199429
GO:0045071: negative regulation of viral genome replication     PROX1   prospero homeobox 1     0.001
102813701165674
GO:0045087: innate immune response      ADAR    adenosine deaminase, RNA-specific       -0.0001369097
3890795742
GO:0045087: innate immune response      AGER    advanced glycosylation end product-specific receptor
-0.00017408994298191997
GO:0045087: innate immune response      AKT1    v-akt murine thymoma viral oncogene homolog 1   0.000
729131294067227
GO:0045087: innate immune response      ANGPT1  angiopoietin 1  0.0008992085477490441
GO:0045087: innate immune response      BCL2    B-cell CLL/lymphoma 2   -4.639779174334028e-6
GO:0045087: innate immune response      C1QA    complement component 1, q subcomponent, A chain 0.000
5413367481662142
GO:0045087: innate immune response      C2      complement component 2  -0.0003033928982477652
GO:0045087: innate immune response      C3      complement component 3  0.002020819095584043
GO:0045087: innate immune response      C8B     complement component 8, beta polypeptide        0.000
23004215933241607
GO:0045087: innate immune response      C8G     complement component 8, gamma polypeptide       -0.00
11927590870437687
GO:0045087: innate immune response      CDC42   cell division cycle 42  0.0012380849751142218
GO:0045087: innate immune response      CFL1    cofilin 1 (non-muscle)  -0.0011725829120191704
GO:0045087: innate immune response      CLEC4M  C-type lectin domain family 4, member M -0.0013876179
810787335
GO:0045087: innate immune response      CREB1   cAMP responsive element binding protein 1       0.000
6563420940543394
GO:0045087: innate immune response      CTNNB1  catenin (cadherin-associated protein), beta 1, 88kDa
-0.00011721946459964427
GO:0045087: innate immune response      CTSK    cathepsin K     -0.00036104001928485266
GO:0045087: innate immune response      DOCK1   dedicator of cytokinesis 1      -0.000682492963446371
1
GO:0045087: innate immune response      EGFR    epidermal growth factor receptor        0.00068350996
30099178
GO:0045087: innate immune response      ERBB4   v-erb-b2 avian erythroblastic leukemia viral oncogene
homolog 4       -0.000181180349062987
GO:0045087: innate immune response      FGF3    fibroblast growth factor 3      0.0015582032755426124
GO:0045087: innate immune response      FGF5    fibroblast growth factor 5      -0.001092025283465080
3
GO:0045087: innate immune response      FGF7    fibroblast growth factor 7      0.0006327941327195195
GO:0045087: innate immune response      FGF8    fibroblast growth factor 8 (androgen-induced)   0.000
9816754327888503
GO:0045087: innate immune response      FGFR2   fibroblast growth factor receptor 2     0.00076147340
31902966
GO:0045087: innate immune response      FGFR3   fibroblast growth factor receptor 3     0.00022228057
360437905
GO:0045087: innate immune response      FOXO1   forkhead box O1 0.0017971524130897447
GO:0045087: innate immune response      FOXO3   forkhead box O3 0.0011976005448450793
GO:0045087: innate immune response      FRS2    fibroblast growth factor receptor substrate 2   0.000
927735698684054
GO:0045087: innate immune response      GATA3   GATA binding protein 3  -3.7500962696795244e-5

GO:0045087: innate immune response      GRB2    growth factor receptor-bound protein 2  0.00042900931
60448253
GO:0045087: innate immune response      GSK3B   glycogen synthase kinase 3 beta 0.0015463276968181818
GO:0045087: innate immune response      HMGB1   high mobility group box 1       -0.000775852096919558
8
GO:0045087: innate immune response      IFNA2   interferon, alpha 2     -0.0017142513169112919
GO:0045087: innate immune response      IRAK1   interleukin-1 receptor-associated kinase 1      -0.00
16061162290777725
GO:0045087: innate immune response      IRF7    interferon regulatory factor 7  -0.001304373328927445
8
GO:0045087: innate immune response      ITPR1   inositol 1,4,5-trisphosphate receptor, type 1   -0.00
07779355211256254
GO:0045087: innate immune response      JAK2    Janus kinase 2  -3.2508702579955e-5
GO:0045087: innate immune response      KIT     v-kit Hardy-Zuckerman 4 feline sarcoma viral oncogene
homolog 0.00027281533716681447
GO:0045087: innate immune response      LGR4    leucine-rich repeat containing G protein-coupled rece
ptor 4  0.0003078166954871878
GO:0045087: innate immune response      LYN     LYN proto-oncogene, Src family tyrosine kinase  -0.00
1418218271748839
GO:0045087: innate immune response      MAP3K5  mitogen-activated protein kinase kinase kinase 5
0.00034169417563195177
GO:0045087: innate immune response      MEF2C   myocyte enhancer factor 2C      0.0009711998910311992
GO:0045087: innate immune response      NCK1    NCK adaptor protein 1   -0.0007843203023177385
GO:0045087: innate immune response      PADI4   peptidyl arginine deiminase, type IV    -0.0008246749
297307712
GO:0045087: innate immune response      PAK1    p21 protein (Cdc42/Rac)-activated kinase 1      -0.00
21837200619320824
GO:0045087: innate immune response      PAK3    p21 protein (Cdc42/Rac)-activated kinase 3      -0.00
12570743350347785
GO:0045087: innate immune response      PDGFRB  platelet-derived growth factor receptor, beta polypep
tide    -0.00035921679789667
GO:0045087: innate immune response      PIK3CD  phosphatidylinositol-4,5-bisphosphate 3-kinase, catal
ytic subunit delta      -0.0007793959194915439
GO:0045087: innate immune response      PML     promyelocytic leukemia  -0.0006796648975348977
GO:0045087: innate immune response      PRKCA   protein kinase C, alpha -6.014805301749354e-6
GO:0045087: innate immune response      PRKCD   protein kinase C, delta -0.001125627376266381
GO:0045087: innate immune response      PRKDC   protein kinase, DNA-activated, catalytic polypeptide
-0.001853069054887686
GO:0045087: innate immune response      PSMA5   proteasome (prosome, macropain) subunit, alpha type,
 5      0.0003309004949413003
GO:0045087: innate immune response      PSMD11  proteasome (prosome, macropain) 26S subunit, non-ATPa
se, 11  -0.0010939357184463609
GO:0045087: innate immune response      PSMD13  proteasome (prosome, macropain) 26S subunit, non-ATPa
se, 13  -6.014618912810836e-5
GO:0045087: innate immune response      PTEN    phosphatase and tensin homolog  1.798925006716674e-5
GO:0045087: innate immune response      RAF1    Raf-1 proto-oncogene, serine/threonine kinase   0.001
4960551789201786
GO:0045087: innate immune response      RASA1   RAS p21 protein activator (GTPase activating protein)
1       -0.00034784005484039766
GO:0045087: innate immune response      RPS6KA1 ribosomal protein S6 kinase, 90kDa, polypeptide 1
-0.0025100788340712003
GO:0045087: innate immune response      S100A7  S100 calcium binding protein A7 0.0015866887479776808
GO:0045087: innate immune response      S100B   S100 calcium binding protein B  0.005174118983814036
GO:0045087: innate immune response      SIRT2   sirtuin 2       -0.0008504675302753776
GO:0045087: innate immune response      SPTBN1  spectrin, beta, non-erythrocytic 1      0.00160441979
51686748
GO:0045087: innate immune response      TNFAIP3 tumor necrosis factor, alpha-induced protein 3  0.001
1142374994470715
GO:0051607: defense response to virus   ADAR    adenosine deaminase, RNA-specific       -0.0001373714
845315713
GO:0051607: defense response to virus   AZU1    azurocidin 1    0.00019661682879495258
GO:0051607: defense response to virus   BCL2    B-cell CLL/lymphoma 2   -4.80010353877158e-6
GO:0051607: defense response to virus   BNIP3   BCL2/adenovirus E1B 19kDa interacting protein 3 0.002
889094167213789
GO:0051607: defense response to virus   BNIP3L  BCL2/adenovirus E1B 19kDa interacting protein 3-like
-5.756032665825826e-5
GO:0051607: defense response to virus   CXADR   coxsackie virus and adenovirus receptor 0.00043004343
585807366
GO:0051607: defense response to virus   CXCL10  chemokine (C-X-C motif) ligand 10       5.96967268539
8529e-5
GO:0051607: defense response to virus   DICER1  dicer 1, ribonuclease type III  -1.6544603795708045e-
5
GO:0051607: defense response to virus   IFNA2   interferon, alpha 2     -0.001704552535092798
GO:0051607: defense response to virus   IFNG    interferon, gamma       -4.905082573239482e-5
GO:0051607: defense response to virus   IL12B   interleukin 12B 0.0012697272945275384
GO:0051607: defense response to virus   PML     promyelocytic leukemia  -0.0006775239138134026
GO:0060216: definitive hemopoiesis      ADAR    adenosine deaminase, RNA-specific       -0.0001386256
656119203
GO:0060216: definitive hemopoiesis      GATA2   GATA binding protein 2  -0.00042300767220347917
GO:0060216: definitive hemopoiesis      KMT2A   lysine (K)-specific methyltransferase 2A        0.000
6264229801728788

GO:0060216: definitive hemopoiesis	ZFP36L2 ZFP36 ring finger protein-like 2	-0.0003555471
4451408424
GO:0060337: type I interferon signaling pathway ADAR	adenosine deaminase, RNA-specific	-0.00
01373960988728698
GO:0060337: type I interferon signaling pathway EGR1	early growth response 1 0.0010902037090911388
GO:0060337: type I interferon signaling pathway IFNA2	interferon, alpha 2	-0.001712599673081712
7
GO:0060337: type I interferon signaling pathway IRF6	interferon regulatory factor 6	-0.0016882263
485073642
GO:0060337: type I interferon signaling pathway IRF7	interferon regulatory factor 7	-0.0013029546
957386527
GO:0060337: type I interferon signaling pathway PTPN1	protein tyrosine phosphatase, non-receptor ty
pe 1	-0.0010385328019515674
GO:0060339: negative regulation of type I interferon-mediated signaling pathway ADAR	adenosine dea
minase, RNA-specific	-0.00014405564065521343
GO:0060339: negative regulation of type I interferon-mediated signaling pathway PTPN2	protein tyros
ine phosphatase, non-receptor type 2	-0.0002988267418296444
GO:0061484: hematopoietic stem cell homeostasis ADAR	adenosine deaminase, RNA-specific	-0.00
013990644784975787
GO:1900369: negative regulation of RNA interference	ADAR	adenosine deaminase, RNA-specific
-0.00013990644784975787
GO:0006351: transcription, DNA-templated	ADIRF	adipogenesis regulatory factor	-0.0001675617
2091968017
GO:0006351: transcription, DNA-templated	AES	amino-terminal enhancer of split	0.001
35775690530536
GO:0006351: transcription, DNA-templated	APEX1	APEX nuclease (multifunctional DNA repair enz
yme) 1	0.00020166743038709683
GO:0006351: transcription, DNA-templated	AR	androgen receptor	0.0026462465827891235
GO:0006351: transcription, DNA-templated	ASCL1	achaete-scute family bHLH transcription facto
r 1	-0.0014379690025452753
GO:0006351: transcription, DNA-templated	ASF1A	anti-silencing function 1A histone chaperone
0.0010541688723845265
GO:0006351: transcription, DNA-templated	ATAD2	ATPase family, AAA domain containing 2	-4.68
00766832629655e-5
GO:0006351: transcription, DNA-templated	BCL11A	B-cell CLL/lymphoma 11A (zinc finger protein)
-0.0009789705885269245
GO:0006351: transcription, DNA-templated	BIRC5	baculoviral IAP repeat containing 5	-0.00
02607198766295168
GO:0006351: transcription, DNA-templated	BRD4	bromodomain containing 4	0.00055487229
54290831
GO:0006351: transcription, DNA-templated	CASP8AP2	caspase 8 associated protein 2	0.001
277721511183039
GO:0006351: transcription, DNA-templated	CCNC	cyclin C	0.0005696399025622206
GO:0006351: transcription, DNA-templated	CCND1	cyclin D1	-0.0026437378838172427
GO:0006351: transcription, DNA-templated	CDK8	cyclin-dependent kinase 8	0.00262061623
86631115
GO:0006351: transcription, DNA-templated	CDKN2A	cyclin-dependent kinase inhibitor 2A	0.001
7650103453912002
GO:0006351: transcription, DNA-templated	CDKN2B	cyclin-dependent kinase inhibitor 2B (p15, in
hibits CDK4)	-0.0018417462822618975
GO:0006351: transcription, DNA-templated	CHAF1B	chromatin assembly factor 1, subunit B (p60)
0.006306137171951332
GO:0006351: transcription, DNA-templated	CHD3	chromodomain helicase DNA binding protein 3
0.00011887975520542777
GO:0006351: transcription, DNA-templated	CHMP1A	charged multivesicular body protein 1A	-0.00
0782524372196498
GO:0006351: transcription, DNA-templated	CITED1	Cbp/p300-interacting transactivator, with Gl
u/Asp-rich carboxy-terminal domain, 1	0.0028283906085487726
GO:0006351: transcription, DNA-templated	CTNNB1	catenin (cadherin-associated protein), beta
1, 88kDa	-0.00011719238664682538
GO:0006351: transcription, DNA-templated	DAXX	death-domain associated protein 0.00089522331
62214329
GO:0006351: transcription, DNA-templated	DDX17	DEAD (Asp-Glu-Ala-Asp) box helicase 17	0.001
7684564306945573
GO:0006351: transcription, DNA-templated	DNAJC2	DnaJ (Hsp40) homolog, subfamily C, member 2
-0.0009869002119678738
GO:0006351: transcription, DNA-templated	E2F1	E2F transcription factor 1	0.00210506912
7653711
GO:0006351: transcription, DNA-templated	E2F4	E2F transcription factor 4, p107/p130-binding
-0.0023533151198579854
GO:0006351: transcription, DNA-templated	E2F6	E2F transcription factor 6	0.00072026672
71313002
GO:0006351: transcription, DNA-templated	E2F8	E2F transcription factor 8	0.00171622044
14668026
GO:0006351: transcription, DNA-templated	E4F1	E4F transcription factor 1	0.00148559140
3271661
GO:0006351: transcription, DNA-templated	ERBB4	v-erb-b2 avian erythroblastic leukemia viral
oncogene homolog 4	-0.00018432545847019087
GO:0006351: transcription, DNA-templated	ESR1	estrogen receptor 1	-0.000951786834163365
5
GO:0006351: transcription, DNA-templated	ETS2	v-ets avian erythroblastosis virus E26 oncoge

ne homolog 2	-0.0010194748802169618
GO:0006351: transcription, DNA-templated	EYA1	EYA transcriptional coactivator and phosphatase 1	5.3591435563177826e-5
GO:0006351: transcription, DNA-templated	EYA3	EYA transcriptional coactivator and phosphatase 3	0.0008393394739330917
GO:0006351: transcription, DNA-templated	EZH2	enhancer of zeste 2 polycomb repressive complex 2 subunit	-0.00013623578626891606
GO:0006351: transcription, DNA-templated	FOXM1	forkhead box M1 0.00019821275763450404
GO:0006351: transcription, DNA-templated	FOXO1	forkhead box O1 0.0018062678046832812
GO:0006351: transcription, DNA-templated	FOXO3	forkhead box O3 0.0012044867691304317
GO:0006351: transcription, DNA-templated	GATA2	GATA binding protein 2	-0.0004534173767054O646
GO:0006351: transcription, DNA-templated	GFI1	growth factor independent 1 transcription repressor	0.0016694925462020502
GO:0006351: transcription, DNA-templated	GLI1	GLI family zinc finger 1	-0.0013007656246102246
GO:0006351: transcription, DNA-templated	HDAC2	histone deacetylase 2	-0.0012105156916769365
GO:0006351: transcription, DNA-templated	HELLS	helicase, lymphoid-specific	0.0034758686176730234
GO:0006351: transcription, DNA-templated	HES1	hes family bHLH transcription factor 1	-0.0009141658046763737
GO:0006351: transcription, DNA-templated	HEXIM1	hexamethylene bis-acetamide inducible 1 0.0001532764170697497
GO:0006351: transcription, DNA-templated	HEY2	hes-related family bHLH transcription factor with YRPW motif 2	0.0027093792983095464
GO:0006351: transcription, DNA-templated	HINFP	histone H4 transcription factor 0.0009959325316793047
GO:0006351: transcription, DNA-templated	HIPK2	homeodomain interacting protein kinase 2	0.0007731044956616217
GO:0006351: transcription, DNA-templated	HOXA3	homeobox A3	0.0009918027629435857
GO:0006351: transcription, DNA-templated	HOXB1	homeobox B1	0.0036978126034359054
GO:0006351: transcription, DNA-templated	HOXB13	homeobox B13	0.0018316255175505555
GO:0006351: transcription, DNA-templated	HOXB2	homeobox B2	0.0027820391586367067
GO:0006351: transcription, DNA-templated	ID2	inhibitor of DNA binding 2, dominant negative helix-loop-helix protein	6.002927244346824e-5
GO:0006351: transcription, DNA-templated	IRF6	interferon regulatory factor 6	-0.0016995131237588447
GO:0006351: transcription, DNA-templated	JMJD6	jumonji domain containing 6	0.0036023216885157764
GO:0006351: transcription, DNA-templated	KANK1	KN motif and ankyrin repeat domains 1	0.002542335029496424
GO:0006351: transcription, DNA-templated	KLF2	Kruppel-like factor 2	-0.0012354829826068461
GO:0006351: transcription, DNA-templated	KMT2B	lysine (K)-specific methyltransferase 2B	0.0004632856997464389
GO:0006351: transcription, DNA-templated	KMT2D	lysine (K)-specific methyltransferase 2D	-0.00020011455246932128
GO:0006351: transcription, DNA-templated	LHX6	LIM homeobox 6	0.001349292479343373
GO:0006351: transcription, DNA-templated	MLLT10	myeloid/lymphoid or mixed-lineage leukemia (trithorax homolog, Drosophila); translocated to, 10	-0.00016387801880077656
GO:0006351: transcription, DNA-templated	MLXIP	MLX interacting protein 0.0027297553060307398
GO:0006351: transcription, DNA-templated	MYC	v-myc avian myelocytomatosis viral oncogene homolog	-0.0011399598187555912
GO:0006351: transcription, DNA-templated	NEUROD4	neuronal differentiation 4	0.0015115037634845423
GO:0006351: transcription, DNA-templated	NOC2L	nucleolar complex associated 2 homolog (S. cerevisiae)	-0.00015144586939234875
GO:0006351: transcription, DNA-templated	NPAT	nuclear protein, ataxia-telangiectasia locus	-0.0007116665604177847
GO:0006351: transcription, DNA-templated	NRIP1	nuclear receptor interacting protein 1	0.0010742903363907155
GO:0006351: transcription, DNA-templated	PADI4	peptidyl arginine deiminase, type IV	-0.0008298530598771133
GO:0006351: transcription, DNA-templated	PAX2	paired box 2	-0.0016200068037620424
GO:0006351: transcription, DNA-templated	PAX8	paired box 8	0.0009506601049226411
GO:0006351: transcription, DNA-templated	PER2	period circadian clock 2	0.00124245990140614
GO:0006351: transcription, DNA-templated	PHB2	prohibitin 2	-0.0007384633744968914
GO:0006351: transcription, DNA-templated	PKN1	protein kinase N1	-0.0019094981210912747
GO:0006351: transcription, DNA-templated	PML	promyelocytic leukemia	-0.0006815997443380529
GO:0006351: transcription, DNA-templated	PPP1R13L	protein phosphatase 1, regulatory subunit 13 like	-0.001478946538092876
GO:0006351: transcription, DNA-templated	PROX1	prospero homeobox 1	0.0011257513484109634
GO:0006351: transcription, DNA-templated	PURA	purine-rich element binding protein A	-0.0001712873539289803
GO:0006351: transcription, DNA-templated	RB1	retinoblastoma 1	-0.0014992189923968763
GO:0006351: transcription, DNA-templated	RBBP4	retinoblastoma binding protein 4	-0.00

18653430843826372
GO:0006351: transcription, DNA-templated	RFC1	replication factor C (activator 1) 1, 145kDa
0.0010631081672241947
GO:0006351: transcription, DNA-templated	SALL1	spalt-like transcription factor 1	-0.00
2447497162819614
GO:0006351: transcription, DNA-templated	SERPINE1	serpin peptidase inhibitor, clade E
(nexin, plasminogen activator inhibitor type 1), member 1	0.00011211078648559634
GO:0006351: transcription, DNA-templated	SIRT1	sirtuin 1	-1.1689012488517732e-6
GO:0006351: transcription, DNA-templated	SIRT2	sirtuin 2	-0.0008553464471506224
GO:0006351: transcription, DNA-templated	SKI	SKI proto-oncogene	-0.00067246506414285
GO:0006351: transcription, DNA-templated	SMARCD3	SWI/SNF related, matrix associated, actin dep
endent regulator of chromatin, subfamily d, member 3	0.00034685432064183303
GO:0006351: transcription, DNA-templated	SPDEF	SAM pointed domain containing ETS transcripti
on factor	0.0031466516320485228
GO:0006351: transcription, DNA-templated	STAT5A	signal transducer and activator of transcript
ion 5A	0.0016015566768442433
GO:0006351: transcription, DNA-templated	TBX21	T-box 21	0.00039949044251085653
GO:0006351: transcription, DNA-templated	TBX3	T-box 3 0.001229611724893063
GO:0006351: transcription, DNA-templated	TCF15	transcription factor 15 (basic helix-loop-hel
ix)	-0.0026633036080798435
GO:0006351: transcription, DNA-templated	TCF7L2	transcription factor 7-like 2 (T-cell specifi
c, HMG-box)	0.000576051457913259
GO:0006351: transcription, DNA-templated	TFCP2L1 transcription factor CP2-like 1 0.00045065901
260167763
GO:0006351: transcription, DNA-templated	TFDP1	transcription factor Dp-1	0.00145099395
41695938
GO:0006351: transcription, DNA-templated	THRB	thyroid hormone receptor, beta	0.00195010915
83035255
GO:0006351: transcription, DNA-templated	TWIST1	twist family bHLH transcription factor 1
-0.0013467142954703025
GO:0006351: transcription, DNA-templated	TXLNG	taxilin gamma	-0.0008396131646636603
GO:0006351: transcription, DNA-templated	VPS72	vacuolar protein sorting 72 homolog (S. cerev
isiae)	0.0005604743224801483
GO:0006351: transcription, DNA-templated	WWTR1	WW domain containing transcription regulator
1	0.0009044947774101581
GO:0006351: transcription, DNA-templated	YY1	YY1 transcription factor	0.00134542575
2509013
GO:0006351: transcription, DNA-templated	ZBTB18	zinc finger and BTB domain containing 18
0.001276055256972934
GO:0006351: transcription, DNA-templated	ZNF205	zinc finger protein 205 0.0005971545050382174
GO:0006351: transcription, DNA-templated	ZNF24	zinc finger protein 24	0.0012956758041332134
GO:0006351: transcription, DNA-templated	ZNF266	zinc finger protein 266 0.0004135283287241028
7
GO:0006351: transcription, DNA-templated	ZNF335	zinc finger protein 335 -0.000365896095415836
GO:0006351: transcription, DNA-templated	ZNF442	zinc finger protein 442 0.0010621231350483454
GO:0030154: cell differentiation	ADIRF	adipogenesis regulatory factor	-0.000166818871948602
92
GO:0030154: cell differentiation	AKT1	v-akt murine thymoma viral oncogene homolog 1	0.000
7306761542897612
GO:0030154: cell differentiation	ARHGAP24	Rho GTPase activating protein 24	-0.00
09343006482775353
GO:0030154: cell differentiation	AXIN1	axin 1	-0.0007333463540473517
GO:0030154: cell differentiation	CPLX2	complexin 2	0.002031753329072328
GO:0030154: cell differentiation	EDA	ectodysplasin A -0.0008146890664469242
GO:0030154: cell differentiation	EDAR	ectodysplasin A receptor	0.0005936239813974122
GO:0030154: cell differentiation	EGFL6	EGF-like-domain, multiple 6	0.0001366426792905652
8
GO:0030154: cell differentiation	ELF5	E74-like factor 5 (ets domain transcription factor)
0.0011718001958474476
GO:0030154: cell differentiation	ETS2	v-ets avian erythroblastosis virus E26 oncogene homol
og 2	-0.0010155851001721265
GO:0030154: cell differentiation	ETV4	ets variant 4	0.00019503508883322862
GO:0030154: cell differentiation	FOXA1	forkhead box A1 2.7263894893049888e-5
GO:0030154: cell differentiation	FOXA2	forkhead box A2 -1.660561017841773e-5
GO:0030154: cell differentiation	FOXC1	forkhead box C1 -2.185149261822642e-5
GO:0030154: cell differentiation	FOXC2	forkhead box C2 (MFH-1, mesenchyme forkhead 1)	0.001
7480670298103474
GO:0030154: cell differentiation	FOXE3	forkhead box E3 0.0010507447838200954
GO:0030154: cell differentiation	FOXH1	forkhead box H1 -0.00137142561380663
GO:0030154: cell differentiation	FOXM1	forkhead box M1 0.00019748677887642313
GO:0030154: cell differentiation	FOXO1	forkhead box O1 0.0018001303307412376
GO:0030154: cell differentiation	FOXO3	forkhead box O3 0.0012001196054617592
GO:0030154: cell differentiation	GLRX2	glutaredoxin 2	0.0011160844527823418
GO:0030154: cell differentiation	INHA	inhibin, alpha	0.0002114755901390126
GO:0030154: cell differentiation	INHBA	inhibin, beta A -0.0013534974399642872
GO:0030154: cell differentiation	JAG2	jagged 2	-5.620467313112915e-6
GO:0030154: cell differentiation	JAK2	Janus kinase 2	-3.152112797203177e-5
GO:0030154: cell differentiation	KIF2A	kinesin heavy chain member 2A	-0.00078439840897598
GO:0030154: cell differentiation	MDK	midkine (neurite growth-promoting factor 2)	0.001
6657587701324886
GO:0030154: cell differentiation	NDRG2	NDRG family member 2	0.002174139695071476

GO:0030154: cell differentiation        NELL1   NEL-like 1 (chicken)    0.0015329098517756214
GO:0030154: cell differentiation        PURA    purine-rich element binding protein A   -0.0001705929
2069549633
GO:0030154: cell differentiation        SLC7A5  solute carrier family 7 (amino acid transporter light
chain, L system), member 5      -0.0025740751474991504
GO:0030154: cell differentiation        SPDEF   SAM pointed domain containing ETS transcription facto
r       0.0031363775898288313
GO:0030154: cell differentiation        SRD5A2  steroid-5-alpha-reductase, alpha polypeptide 2 (3-oxo
-5 alpha-steroid delta 4-dehydrogenase alpha 2) -0.0006432281117309362
GO:0030154: cell differentiation        TP53    tumor protein p53       0.001175980698785729
GO:0030154: cell differentiation        YY1     YY1 transcription factor        0.0013402815459465913
GO:0045600: positive regulation of fat cell differentiation     ADIRF   adipogenesis regulatory facto
r       -0.00016418273076696852
GO:0045600: positive regulation of fat cell differentiation     AKT1    v-akt murine thymoma viral on
cogene homolog 1        0.0007232830270269224
GO:0045600: positive regulation of fat cell differentiation     CEBPB   CCAAT/enhancer binding protei
n (C/EBP), beta -0.00027106293107750267
GO:0045600: positive regulation of fat cell differentiation     CREB1   cAMP responsive element bindi
ng protein 1    0.0006504646253352423
GO:0045600: positive regulation of fat cell differentiation     HTR2C   5-hydroxytryptamine (serotoni
n) receptor 2C, G protein-coupled       -0.0005203800227167745
GO:0045600: positive regulation of fat cell differentiation     ID2     inhibitor of DNA binding 2, d
ominant negative helix-loop-helix protein       6.035294622757794e-5
GO:0045600: positive regulation of fat cell differentiation     LRP5    low density lipoprotein recep
tor-related protein 5   3.132876831481795e-5
GO:0045600: positive regulation of fat cell differentiation     SAV1    salvador family WW domain con
taining protein 1       -0.002320787296476295
GO:0045600: positive regulation of fat cell differentiation     SFRP1   secreted frizzled-related pro
tein 1  0.0012656508995616946
GO:0045600: positive regulation of fat cell differentiation     XBP1    X-box binding protein 1 0.000
26508061877325784
GO:0071478: cellular response to radiation      ADIRF   adipogenesis regulatory factor  -0.0001435353
1936957796
GO:0072719: cellular response to cisplatin      ADIRF   adipogenesis regulatory factor  -0.0001633631
8119645045
GO:0072719: cellular response to cisplatin      RAD51   RAD51 recombinase       -0.001848593006405684
GO:0072719: cellular response to cisplatin      SLC31A1 solute carrier family 31 (copper transporte
r), member 1    -2.7586334276709638e-5
GO:2001023: regulation of response to drug      ADIRF   adipogenesis regulatory factor  -0.0001648556
9311122314
GO:2001023: regulation of response to drug      GABRA4  gamma-aminobutyric acid (GABA) A receptor, al
pha 4   -0.001474518371420807
GO:0001570: vasculogenesis      ADM     adrenomedullin  0.002243407888577193
GO:0001570: vasculogenesis      CAV1    caveolin 1, caveolae protein, 22kDa     -0.000536389671852577
3
GO:0001570: vasculogenesis      CITED1  Cbp/p300-interacting transactivator, with Glu/Asp-rich carbox
y-terminal domain, 1    0.00282634847978138
GO:0001570: vasculogenesis      CUL7    cullin 7        -0.00011962825521386781
GO:0001570: vasculogenesis      ENG     endoglin        0.0008293706626310093
GO:0001570: vasculogenesis      FOXM1   forkhead box M1 0.00019753488647465225
GO:0001570: vasculogenesis      HEY2    hes-related family bHLH transcription factor with YRPW motif
 2      0.002706969051319108
GO:0001570: vasculogenesis      PITX2   paired-like homeodomain 2       0.0021820487507270485
GO:0001570: vasculogenesis      PTPRJ   protein tyrosine phosphatase, receptor type, J  -0.0003391328
524076696
GO:0001570: vasculogenesis      RASA1   RAS p21 protein activator (GTPase activating protein) 1 -0.00
034999822780314285
GO:0001570: vasculogenesis      SHH     sonic hedgehog  0.0006020355406372965
GO:0001570: vasculogenesis      SMO     smoothened, frizzled class receptor     0.002144905640206822
GO:0001570: vasculogenesis      SOX18   SRY (sex determining region Y)-box 18   0.0012914910099911384
GO:0001570: vasculogenesis      VEGFA   vascular endothelial growth factor A    0.0005968811974446953
GO:0001570: vasculogenesis      WT1     Wilms tumor 1   -0.0005086154486246668
GO:0001570: vasculogenesis      YAP1    Yes-associated protein 1        -0.00016470097204287595
GO:0001843: neural tube closure ADM     adrenomedullin  0.00224464289403796
GO:0001843: neural tube closure ALX1    ALX homeobox 1  0.0022988626457801646
GO:0001843: neural tube closure BBS4    Bardet-Biedl syndrome 4 -0.0005105909852871573
GO:0001843: neural tube closure BMP4    bone morphogenetic protein 4    -0.00032355394167502943
GO:0001843: neural tube closure FZD3    frizzled class receptor 3       0.00038046660546738623
GO:0001843: neural tube closure GRHL2   grainyhead-like 2 (Drosophila)  0.0010095235843284257
GO:0001843: neural tube closure IFT122  intraflagellar transport 122 homolog (Chlamydomonas)    -0.00
16152546694120256
GO:0001843: neural tube closure LIAS    lipoic acid synthetase  -0.0005556733559023037
GO:0001843: neural tube closure LMO4    LIM domain only 4       0.002117965519340124
GO:0001843: neural tube closure LRP6    low density lipoprotein receptor-related protein 6      0.000
14996070998739782
GO:0001843: neural tube closure MTHFD1  methylenetetrahydrofolate dehydrogenase (NADP+ dependent) 1,
 methenyltetrahydrofolate cyclohydrolase, formyltetrahydrofolate synthetase     -0.000665730350697600
4
GO:0001843: neural tube closure PAX2    paired box 2    -0.0016195239316690046
GO:0001843: neural tube closure PAX3    paired box 3    -0.002954136339905277
GO:0001843: neural tube closure PHACTR4 phosphatase and actin regulator 4       -0.001796271394250020

1
GO:0001843: neural tube closure PTCH1 patched 1 -6.577755257534144e-5
GO:0001843: neural tube closure RARG retinoic acid receptor, gamma -0.0023266427241055336
GO:0001843: neural tube closure SALL1 spalt-like transcription factor 1 -0.002446828641577958
6
GO:0001843: neural tube closure SKI SKI proto-oncogene -0.0006723450428151537
GO:0001843: neural tube closure TULP3 tubby like protein 3 0.0009591934635432494
GO:0001843: neural tube closure TWIST1 twist family bHLH transcription factor 1 -0.0013463727
26287962
GO:0002026: regulation of the force of heart contraction ADM adrenomedullin 0.00222994670
9411426
GO:0002026: regulation of the force of heart contraction IFNG interferon, gamma -4.91
7862048942022e-5
GO:0002026: regulation of the force of heart contraction PRKCA protein kinase C, alpha -6.13
0061387185331e-6
GO:0002031: G-protein coupled receptor internalization ADM adrenomedullin 0.002241989129939734
GO:0002031: G-protein coupled receptor internalization DRD2 dopamine receptor D2 -0.0002343534
6128204245
GO:0006171: cAMP biosynthetic process ADM adrenomedullin 0.0022574560122283513
GO:0006701: progesterone biosynthetic process ADM adrenomedullin 0.002247342140914187
GO:0006701: progesterone biosynthetic process STAR steroidogenic acute regulatory protein 0.000
8989063754188105
GO:0007204: positive regulation of cytosolic calcium ion concentration ADM adrenomedullin 0.002
23819578503534
GO:0007204: positive regulation of cytosolic calcium ion concentration AGTR1 angiotensin II recept
or, type 1 0.00034695014327386164
GO:0007204: positive regulation of cytosolic calcium ion concentration AVP arginine vasopressin
-0.0009377923473621331
GO:0007204: positive regulation of cytosolic calcium ion concentration CD24 CD24 molecule 0.001
0723046314722062
GO:0007204: positive regulation of cytosolic calcium ion concentration CXCL13 chemokine (C-X-C moti
f) ligand 13 0.002993858781088234
GO:0007204: positive regulation of cytosolic calcium ion concentration CXCR4 chemokine (C-X-C moti
f) receptor 4 0.0007952823600648643
GO:0007204: positive regulation of cytosolic calcium ion concentration ESR1 estrogen receptor 1
-0.0009489711330357444
GO:0007204: positive regulation of cytosolic calcium ion concentration GATA2 GATA binding protein
2 -0.000452190396797567
GO:0007204: positive regulation of cytosolic calcium ion concentration GJA1 gap junction protein,
alpha 1, 43kDa -0.00016247681117484487
GO:0007204: positive regulation of cytosolic calcium ion concentration HMGB1 high mobility group b
ox 1 -0.0007759442251216441
GO:0007204: positive regulation of cytosolic calcium ion concentration JAK2 Janus kinase 2 -3.13
9059991722579e-5
GO:0007267: cell-cell signaling ADM adrenomedullin 0.002242158358743023
GO:0007267: cell-cell signaling AGT angiotensinogen (serpin peptidase inhibitor, clade A, member
8) -0.0011363402136848188
GO:0007267: cell-cell signaling AR androgen receptor 0.0026432879501507873
GO:0007267: cell-cell signaling AREG amphiregulin 0.002462287461729004
GO:0007267: cell-cell signaling AVP arginine vasopressin -0.0009395884492577824
GO:0007267: cell-cell signaling C1QA complement component 1, q subcomponent, A chain 0.00054351884
46500535
GO:0007267: cell-cell signaling CCL7 chemokine (C-C motif) ligand 7 -0.002406873097617114
GO:0007267: cell-cell signaling CCL8 chemokine (C-C motif) ligand 8 -0.0005913692449657672
GO:0007267: cell-cell signaling CXCL10 chemokine (C-X-C motif) ligand 10 6.277374234536331e-5
GO:0007267: cell-cell signaling CXCL13 chemokine (C-X-C motif) ligand 13 0.0029995297624913412
GO:0007267: cell-cell signaling EFNA1 ephrin-A1 -0.0005215606381449763
GO:0007267: cell-cell signaling FGF3 fibroblast growth factor 3 0.0015642588656650518
GO:0007267: cell-cell signaling FGF5 fibroblast growth factor 5 -0.0010961366567452285
GO:0007267: cell-cell signaling FGFR2 fibroblast growth factor receptor 2 0.0007649176784101619
GO:0007267: cell-cell signaling GATA4 GATA binding protein 4 -0.0010970487169320359
GO:0007267: cell-cell signaling GJA1 gap junction protein, alpha 1, 43kDa -0.000162700920972711
34
GO:0007267: cell-cell signaling GRB2 growth factor receptor-bound protein 2 0.0004305622912941412
GO:0007267: cell-cell signaling GREM1 gremlin 1, DAN family BMP antagonist -0.000829799277102856
3
GO:0007267: cell-cell signaling IFNA2 interferon, alpha 2 -0.001720762184470687
GO:0007267: cell-cell signaling IHH indian hedgehog -0.002045057184720435
GO:0007267: cell-cell signaling IL7 interleukin 7 0.0008733124452919574
GO:0007267: cell-cell signaling INHA inhibin, alpha 0.00021174228440315032
GO:0007267: cell-cell signaling INHBA inhibin, beta A -0.0013570390310194821
GO:0007267: cell-cell signaling LHX1 LIM homeobox 1 -0.0007604609081624376
GO:0007267: cell-cell signaling NRP1 neuropilin 1 -0.0006448268696486021
GO:0007267: cell-cell signaling PGR progesterone receptor 0.00035247299751778603
GO:0007267: cell-cell signaling SHH sonic hedgehog 0.000601598000356959
GO:0007267: cell-cell signaling SRD5A2 steroid-5-alpha-reductase, alpha polypeptide 2 (3-oxo-5 alpha
-steroid delta 4-dehydrogenase alpha 2) -0.0006447623364778302
GO:0007267: cell-cell signaling TFAP2C transcription factor AP-2 gamma (activating enhancer binding
protein 2 gamma) 0.0010633530740946505
GO:0007267: cell-cell signaling TGFB2 transforming growth factor, beta 2 -0.001061919689039696
5

GO:0007267: cell-cell signaling TNFRSF11A tumor necrosis factor receptor superfamily, member 11
a, NFKB activator 0.0027819203611569665
GO:0007267: cell-cell signaling WNT1 wingless-type MMTV integration site family, member 1 0.000
7884694180261598
GO:0007507: heart development ADM adrenomedullin 0.00224039765412097
GO:0007507: heart development CRIP1 cysteine-rich protein 1 (intestinal) 0.0017234078132703319
GO:0007507: heart development CXADR coxsackie virus and adenovirus receptor 0.0004359879102501645
6
GO:0007507: heart development ERBB4 v-erb-b2 avian erythroblastic leukemia viral oncogene homolog
4 -0.00018362642045981697
GO:0007507: heart development FOXC1 forkhead box C1 -2.1815688535006202e-5
GO:0007507: heart development FOXC2 forkhead box C2 (MFH-1, mesenchyme forkhead 1) 0.00175008872
52007932
GO:0007507: heart development GATA2 GATA binding protein 2 -0.00045234520014068516
GO:0007507: heart development GATA3 GATA binding protein 3 -3.946625380090387e-5
GO:0007507: heart development GJA1 gap junction protein, alpha 1, 43kDa -0.000162487435923002
78
GO:0007507: heart development GLI2 GLI family zinc finger 2 0.0018556407034636836
GO:0007507: heart development GLI3 GLI family zinc finger 3 -0.0021545456124907633
GO:0007507: heart development HEXIM1 hexamethylene bis-acetamide inducible 1 0.0001527033173223712
2
GO:0007507: heart development ITGA3 integrin, alpha 3 (antigen CD49C, alpha 3 subunit of VLA-3 re
ceptor) 0.0014011512107650489
GO:0007507: heart development JMJD6 jumonji domain containing 6 0.0035947509921381428
GO:0007507: heart development MEF2C myocyte enhancer factor 2C 0.0009746357047119459
GO:0007507: heart development MTHFD1 methylenetetrahydrofolate dehydrogenase (NADP+ dependent) 1,
methenyltetrahydrofolate cyclohydrolase, formyltetrahydrofolate synthetase -0.000664543679672547
1
GO:0007507: heart development NOTCH1 notch 1 0.0005195080009200778
GO:0007507: heart development PAX3 paired box 3 -0.0029484446967786592
GO:0007507: heart development PCNA proliferating cell nuclear antigen 0.0012280391624806327
GO:0007507: heart development PRKDC protein kinase, DNA-activated, catalytic polypeptide -0.00
18574484148168396
GO:0007507: heart development PTEN phosphatase and tensin homolog 1.6771831926168876e-5
GO:0007507: heart development PTPRJ protein tyrosine phosphatase, receptor type, J -0.0003387748
230284295
GO:0007507: heart development RAF1 Raf-1 proto-oncogene, serine/threonine kinase 0.00150050381
99883138
GO:0007507: heart development SALL1 spalt-like transcription factor 1 -0.002440794807058593
GO:0007507: heart development SHH sonic hedgehog 0.0006008317048758141
GO:0007507: heart development SOX4 SRY (sex determining region Y)-box 4 -3.797852553967947e-5
GO:0007507: heart development STRA6 stimulated by retinoic acid 6 -0.0016824461670900401
GO:0007507: heart development TGFB2 transforming growth factor, beta 2 -0.001060737018336623
GO:0007507: heart development TGFBR1 transforming growth factor, beta receptor 1 0.00034043058
182679726
GO:0007507: heart development TH tyrosine hydroxylase -0.0003423539449118166
GO:0007507: heart development TRPS1 trichorhinophalangeal syndrome I -0.000178312805524067
5
GO:0007507: heart development VLDLR very low density lipoprotein receptor 0.0009544257474276611
GO:0007507: heart development WT1 Wilms tumor 1 -0.0005075979069531802
GO:0007565: female pregnancy ADM adrenomedullin 0.0022416185676936698
GO:0007565: female pregnancy AGT angiotensinogen (serpin peptidase inhibitor, clade A, member
8) -0.0011362942712785187
GO:0007565: female pregnancy BCL2 B-cell CLL/lymphoma 2 -4.9559769533563885e-6
GO:0007565: female pregnancy EPN1 epsin 1 -0.0005480359649646315
GO:0007565: female pregnancy IDO1 indoleamine 2,3-dioxygenase 1 0.001228343150625145
GO:0007565: female pregnancy IGFBP2 insulin-like growth factor binding protein 2, 36kDa 0.000
14538508354895374
GO:0007565: female pregnancy IL4 interleukin 4 0.0002594868369637105
GO:0007565: female pregnancy LEP leptin 0.003197561759565936
GO:0007565: female pregnancy LGALS9 lectin, galactoside-binding, soluble, 9 -0.000116647187943979
58
GO:0007565: female pregnancy MAPT microtubule-associated protein tau 0.0015532975118281046
GO:0007565: female pregnancy PSG4 pregnancy specific beta-1-glycoprotein 4 -0.0010328661
238374205
GO:0007565: female pregnancy STAT5A signal transducer and activator of transcription 5A 0.001
5989561575371666
GO:0007565: female pregnancy TFCP2L1 transcription factor CP2-like 1 0.0004499474064500434
GO:0007565: female pregnancy TGFB1 transforming growth factor, beta 1 -7.329792419287216e-5
GO:0007565: female pregnancy TGFB3 transforming growth factor, beta 3 -0.001827661027896623
3
GO:0007568: aging ADM adrenomedullin 0.002243579445825092
GO:0007568: aging AGT angiotensinogen (serpin peptidase inhibitor, clade A, member 8) -0.00
11377192141996972
GO:0007568: aging AKT1 v-akt murine thymoma viral oncogene homolog 1 0.0007328830645346387
GO:0007568: aging ALOX12 arachidonate 12-lipoxygenase -0.001958980650874002
GO:0007568: aging APOD apolipoprotein D 0.002635286312649951
GO:0007568: aging ASS1 argininosuccinate synthase 1 0.00014394037307516772
GO:0007568: aging BAK1 BCL2-antagonist/killer 1 -0.0018808797564946193
GO:0007568: aging CAST calpastatin -0.003041868365457816
GO:0007568: aging CCL2 chemokine (C-C motif) ligand 2 0.0008189559148773362

GO:0007568: aging CDKN2B cyclin-dependent kinase inhibitor 2B (p15, inhibits CDK4) -0.0018402570942091766
GO:0007568: aging CRYAB crystallin, alpha B 0.0009879638542821011
GO:0007568: aging CX3CL1 chemokine (C-X3-C motif) ligand 1 0.00212376964791682
GO:0007568: aging GLRX2 glutaredoxin 2 0.0011187928960393282
GO:0007568: aging GPX1 glutathione peroxidase 1 0.00038770150745406505
GO:0007568: aging GRB2 growth factor receptor-bound protein 2 0.00043072339321570765
GO:0007568: aging IGFBP2 insulin-like growth factor binding protein 2, 36kDa 0.00014553747740461188
GO:0007568: aging IRAK1 interleukin-1 receptor-associated kinase 1 -0.001613150040502562
GO:0007568: aging KCNE2 potassium voltage-gated channel, Isk-related family, member 2 -0.0005766565756590681
GO:0007568: aging KRT33B keratin 33B 0.0009447727301874563
GO:0007568: aging MMP7 matrix metallopeptidase 7 (matrilysin, uterine) 0.00042639368531000347
GO:0007568: aging PAX2 paired box 2 -0.0016188877523914803
GO:0007568: aging PTEN phosphatase and tensin homolog 1.623914354362556e-5
GO:0007568: aging RXRA retinoid X receptor, alpha 0.0011178851661669655
GO:0007568: aging TFRC transferrin receptor 0.0010869358788891507
GO:0007568: aging TGFB1 transforming growth factor, beta 1 -7.344736873760854e-5
GO:0007568: aging TGFB3 transforming growth factor, beta 3 -0.0018296143229937802
GO:0007568: aging TYMS thymidylate synthetase 0.0015670155233409745
GO:0008209: androgen metabolic process ADM adrenomedullin 0.002235753116548338
GO:0008209: androgen metabolic process ESR1 estrogen receptor 1 -0.0009473512583995906
GO:0008209: androgen metabolic process HSD17B4 hydroxysteroid (17-beta) dehydrogenase 4 -0.0011616394572610343
GO:0008209: androgen metabolic process SHH sonic hedgehog 0.0005990696765356115
GO:0008209: androgen metabolic process SRD5A2 steroid-5-alpha-reductase, alpha polypeptide 2 (3-oxo-5 alpha-steroid delta 4-dehydrogenase alpha 2) -0.0006422839767221326
GO:0008284: positive regulation of cell proliferation ADM adrenomedullin 0.002241095099308058
GO:0008284: positive regulation of cell proliferation ALDH1A2 aldehyde dehydrogenase 1 family, member A2 -0.0030233719864110917
GO:0008284: positive regulation of cell proliferation ALOX12 arachidonate 12-lipoxygenase -0.001956588532168184
GO:0008284: positive regulation of cell proliferation AR androgen receptor 0.0026412302672169123
GO:0008284: positive regulation of cell proliferation AREG amphiregulin 0.0024605282476191062
GO:0008284: positive regulation of cell proliferation AVP arginine vasopressin -0.0009391515786076598
GO:0008284: positive regulation of cell proliferation BIRC5 baculoviral IAP repeat containing 5 -0.00026023622568528557
GO:0008284: positive regulation of cell proliferation CDC20 cell division cycle 20 -0.0001883575178449535
GO:0008284: positive regulation of cell proliferation CDC7 cell division cycle 7 -0.0006803484279616096
GO:0008284: positive regulation of cell proliferation CIB1 calcium and integrin binding 1 (calmyrin) 5.403418133617749e-5
GO:0008284: positive regulation of cell proliferation CST3 cystatin C -7.092712328068067e-5
GO:0008284: positive regulation of cell proliferation CXCL10 chemokine (C-X-C motif) ligand 10 6.311418221515935e-5
GO:0008284: positive regulation of cell proliferation E2F3 E2F transcription factor 3 0.0021651822739979293
GO:0008284: positive regulation of cell proliferation EGFR epidermal growth factor receptor 0.0006873945507094657
GO:0008284: positive regulation of cell proliferation ERBB4 v-erb-b2 avian erythroblastic leukemia viral oncogene homolog 4 -0.00018389225564362292
GO:0008284: positive regulation of cell proliferation FGF3 fibroblast growth factor 3 0.0015633065651605806
GO:0008284: positive regulation of cell proliferation FGF5 fibroblast growth factor 5 -0.0010955850081850222
GO:0008284: positive regulation of cell proliferation FGF7 fibroblast growth factor 7 0.0006345221184297736
GO:0008284: positive regulation of cell proliferation FGF8 fibroblast growth factor 8 (androgen-induced) 0.000985730356116512
GO:0008284: positive regulation of cell proliferation FGFR2 fibroblast growth factor receptor 2 0.0007645573325409215
GO:0008284: positive regulation of cell proliferation FGFR3 fibroblast growth factor receptor 3 0.00022307374408331044
GO:0008284: positive regulation of cell proliferation FLT3 fms-related tyrosine kinase 3 -0.0006828990140541775
GO:0008284: positive regulation of cell proliferation FOXM1 forkhead box M1 0.00019796836757311295
GO:0008284: positive regulation of cell proliferation FZR1 fizzy/cell division cycle 20 related 1 (Drosophila) 0.0009496567670515858
GO:0008284: positive regulation of cell proliferation GCNT2 glucosaminyl (N-acetyl) transferase 2, I-branching enzyme (I blood group) 0.0024357318828497183
GO:0008284: positive regulation of cell proliferation GDNF glial cell derived neurotrophic factor 0.00044520541170774364
GO:0008284: positive regulation of cell proliferation GLI1 GLI family zinc finger 1 -0.0012985550423469138
GO:0008284: positive regulation of cell proliferation GREM1 gremlin 1, DAN family BMP antagonist

-0.000829320446677592
GO:0008284: positive regulation of cell proliferation HDAC2 histone deacetylase 2 -0.0012082371845297465
GO:0008284: positive regulation of cell proliferation HES1 hes family bHLH transcription factor 1 -0.0009123981761961905
GO:0008284: positive regulation of cell proliferation HIPK2 homeodomain interacting protein kinase 2 0.0007717085127869683
GO:0008284: positive regulation of cell proliferation HOXA3 homeobox A3 0.0009900277491969953
GO:0008284: positive regulation of cell proliferation IFNG interferon, gamma -4.88745699783481e-5
GO:0008284: positive regulation of cell proliferation IGF1 insulin-like growth factor 1 (somatomedin C) 0.00013135316324580068
GO:0008284: positive regulation of cell proliferation IGF1R insulin-like growth factor 1 receptor 0.001072870770111451
GO:0008284: positive regulation of cell proliferation IL6ST interleukin 6 signal transducer 0.0018777446167258067
GO:0008284: positive regulation of cell proliferation IL7 interleukin 7 0.0008729160679010594
GO:0008284: positive regulation of cell proliferation INSR insulin receptor -0.001365485686901853
GO:0008284: positive regulation of cell proliferation ISL1 ISL LIM homeobox 1 7.867174807322098e-5
GO:0008284: positive regulation of cell proliferation ITGB1 integrin, beta 1 (fibronectin receptor, beta polypeptide, antigen CD29 includes MDF2, MSK12) 0.00243078676658753
GO:0008284: positive regulation of cell proliferation JAK2 Janus kinase 2 -3.133823986738434e-5
GO:0008284: positive regulation of cell proliferation KIF14 kinesin family member 14 0.0004869357289194405
GO:0008284: positive regulation of cell proliferation KIT v-kit Hardy-Zuckerman 4 feline sarcoma viral oncogene homolog 0.0002760787871846932
GO:0008284: positive regulation of cell proliferation KMT2D lysine (K)-specific methyltransferase 2D -0.00019977270008911634
GO:0008284: positive regulation of cell proliferation LEF1 lymphoid enhancer-binding factor 1 -0.00010029784314462732
GO:0008284: positive regulation of cell proliferation LEP leptin 0.003196366515619117
GO:0008284: positive regulation of cell proliferation LRP5 low density lipoprotein receptor-related protein 5 3.0206857642998193e-5
GO:0008284: positive regulation of cell proliferation LYN LYN proto-oncogene, Src family tyrosine kinase -0.0014224121197098168
GO:0008284: positive regulation of cell proliferation MAS1 MAS1 proto-oncogene, G protein-coupled receptor 0.0002911621625667219
GO:0008284: positive regulation of cell proliferation MFGE8 milk fat globule-EGF factor 8 protein 0.0018078374057839488
GO:0008284: positive regulation of cell proliferation MYC v-myc avian myelocytomatosis viral oncogene homolog -0.001137632466996802
GO:0008284: positive regulation of cell proliferation NOTCH1 notch 1 0.0005197075762901453
GO:0008284: positive regulation of cell proliferation ODC1 ornithine decarboxylase 1 0.0008268257255228951
GO:0008284: positive regulation of cell proliferation PAX3 paired box 3 -0.0029493475218467114
GO:0008284: positive regulation of cell proliferation PDGFRB platelet-derived growth factor receptor, beta polypeptide -0.0003624394164056987
GO:0008284: positive regulation of cell proliferation PDX1 pancreatic and duodenal homeobox 1 0.00025791783621007214
GO:0008284: positive regulation of cell proliferation POU1F1 POU class 1 homeobox 1 0.0001376925741848538
GO:0008284: positive regulation of cell proliferation PRC1 protein regulator of cytokinesis 1 0.0012311469731749356
GO:0008284: positive regulation of cell proliferation PRKCZ protein kinase C, zeta -0.0016029215175104086
GO:0008284: positive regulation of cell proliferation PROX1 prospero homeobox 1 0.0011234164177144784
GO:0008284: positive regulation of cell proliferation PTEN phosphatase and tensin homolog 1.6580784823382373e-5
GO:0008284: positive regulation of cell proliferation PTN pleiotrophin 0.00030270899571554523
GO:0008284: positive regulation of cell proliferation PURA purine-rich element binding protein A -0.00017096419545517539
GO:0008284: positive regulation of cell proliferation RARG retinoic acid receptor, gamma -0.0023228080482035345
GO:0008284: positive regulation of cell proliferation RNF187 ring finger protein 187 -0.0022298669757898812
GO:0008284: positive regulation of cell proliferation RPA1 replication protein A1, 70kDa 0.0004406417422636791
GO:0008284: positive regulation of cell proliferation S100B S100 calcium binding protein B 0.005193531837173987
GO:0008284: positive regulation of cell proliferation SFRP1 secreted frizzled-related protein 1 0.0012851051968391038
GO:0008284: positive regulation of cell proliferation SHH sonic hedgehog 0.0006011691842453297
GO:0008284: positive regulation of cell proliferation SHMT2 serine hydroxymethyltransferase 2 (mitochondrial) -0.0010984777529084399
GO:0008284: positive regulation of cell proliferation SIRT1 sirtuin 1 -1.3705262127160919e-6

GO:0008284: positive regulation of cell proliferation   SOX11   SRY (sex determining region Y)-box 11
-0.0002088266702245989
GO:0008284: positive regulation of cell proliferation   SOX4    SRY (sex determining region Y)-box 4
-3.7824390763041414e-5
GO:0008284: positive regulation of cell proliferation   SOX9    SRY (sex determining region Y)-box 9
-0.000519283733273082
GO:0008284: positive regulation of cell proliferation   STAMBP  STAM binding protein    -0.0001705258
2419290952
GO:0008284: positive regulation of cell proliferation   TBX3    T-box 3 0.0012274221775040344
GO:0008284: positive regulation of cell proliferation   TGFB1   transforming growth factor, beta 1
-7.329146074140705e-5
GO:0008284: positive regulation of cell proliferation   TGFB2   transforming growth factor, beta 2
-0.0010612035139734974
GO:0008284: positive regulation of cell proliferation   TGFBR1  transforming growth factor, beta rece
ptor 1  0.00034063030559820453
GO:0008284: positive regulation of cell proliferation   TIPIN   TIMELESS interacting protein    -0.00
05408522058055692
GO:0008284: positive regulation of cell proliferation   TNC     tenascin C      0.0007354561470014529
GO:0008284: positive regulation of cell proliferation   TNFRSF11A       tumor necrosis factor recepto
r superfamily, member 11a, NFKB activator       0.0027803592949542616
GO:0008284: positive regulation of cell proliferation   TTK     TTK protein kinase      0.00108792083
1025738
GO:0008284: positive regulation of cell proliferation   VEGFA   vascular endothelial growth factor A
0.0005964425569342984
GO:0008284: positive regulation of cell proliferation   VEGFC   vascular endothelial growth factor C
-0.003368988555231368
GO:0008284: positive regulation of cell proliferation   WDR77   WD repeat domain 77     0.00013368823
95141296
GO:0008284: positive regulation of cell proliferation   WNT1    wingless-type MMTV integration site f
amily, member 1 0.0007880109313703513
GO:0008284: positive regulation of cell proliferation   WNT10B  wingless-type MMTV integration site f
amily, member 10B       -0.0013909401968136056
GO:0008284: positive regulation of cell proliferation   WWTR1   WW domain containing transcription re
gulator 1       0.0009025750375793533
GO:0008284: positive regulation of cell proliferation   YAP1    Yes-associated protein 1        -0.00
016426358315464287
GO:0009409: response to cold    ADM     adrenomedullin  0.0022397904488824037
GO:0009409: response to cold    AGT     angiotensinogen (serpin peptidase inhibitor, clade A, member
 8)     -0.0011351167577243067
GO:0009409: response to cold    CXCL10  chemokine (C-X-C motif) ligand 10       6.31008925387069e-5
GO:0009409: response to cold    HSPA2   heat shock 70kDa protein 2      -0.00010820053490843672
GO:0009409: response to cold    IMPDH1  IMP (inosine 5'-monophosphate) dehydrogenase 1  -0.0018739381
381500835
GO:0009409: response to cold    THRA    thyroid hormone receptor, alpha 0.0007637331060292849
GO:0009611: response to wounding        ADM     adrenomedullin  0.002236324725947605
GO:0009611: response to wounding        AGER    advanced glycosylation end product-specific receptor
-0.00017421177489628436
GO:0009611: response to wounding        CCL2    chemokine (C-C motif) ligand 2  0.000815580296128885
GO:0009611: response to wounding        CX3CR1  chemokine (C-X3-C motif) receptor 1     -0.0008561588
55469583
GO:0009611: response to wounding        FGF7    fibroblast growth factor 7      0.0006335254785793171
GO:0009611: response to wounding        GAP43   growth associated protein 43    -0.000129617682620767
02
GO:0009611: response to wounding        HOXB13  homeobox B13    0.001824699375745667
GO:0009611: response to wounding        ITGB4   integrin, beta 4        0.000579406653867322
GO:0009611: response to wounding        MDK     midkine (neurite growth-promoting factor 2)     0.001
6628219498300999
GO:0009611: response to wounding        PAX6    paired box 6    0.0019339674064337217
GO:0009611: response to wounding        PDX1    pancreatic and duodenal homeobox 1      0.00025689463
20263222
GO:0009611: response to wounding        SLC1A3  solute carrier family 1 (glial high affinity glutamat
e transporter), member 3        -0.0031195166092617046
GO:0009611: response to wounding        TGFB1   transforming growth factor, beta 1      -7.2615426280
97544e-5
GO:0009611: response to wounding        TGFB2   transforming growth factor, beta 2      -0.0010585455
482918812
GO:0009611: response to wounding        TNC     tenascin C      0.0007340740888897686
GO:0009611: response to wounding        VASH1   vasohibin 1     0.0005983295308603422
GO:0009611: response to wounding        WNT1    wingless-type MMTV integration site family, member 1
0.0007865486151669957
GO:0010460: positive regulation of heart rate   ADM     adrenomedullin  0.00225989832402309
GO:0010460: positive regulation of heart rate   HEY2    hes-related family bHLH transcription factor
 with YRPW motif 2      0.002731516827600815
GO:0010460: positive regulation of heart rate   SLC1A1  solute carrier family 1 (neuronal/epithelial
 high affinity glutamate transporter, system Xag), member 1     -0.0009115214102514169
GO:0019731: antibacterial humoral response      ADM     adrenomedullin  0.0022574560122283513
GO:0019933: cAMP-mediated signaling     ADM     adrenomedullin  0.0022353920621425434
GO:0019933: cAMP-mediated signaling     EPHA5   EPH receptor A5 0.0005981340292903148
GO:0019933: cAMP-mediated signaling     SOX9    SRY (sex determining region Y)-box 9    -0.0005187964
341177326
GO:0030819: positive regulation of cAMP biosynthetic process    ADM     adrenomedullin  0.00224866908

88087217
GO:0030819: positive regulation of cAMP biosynthetic process AVP arginine vasopressin -0.0009429485082957511
GO:0031100: organ regeneration ADM adrenomedullin 0.0022459042879211613
GO:0031100: organ regeneration BAK1 BCL2-antagonist/killer 1 -0.0018825654073903223
GO:0031100: organ regeneration CAST calpastatin -0.0030452372340490086
GO:0031100: organ regeneration CAV1 caveolin 1, caveolae protein, 22kDa -0.0005366414608921823
GO:0031100: organ regeneration CCL2 chemokine (C-C motif) ligand 2 0.0008201448042826116
GO:0031100: organ regeneration CCND1 cyclin D1 -0.0026460934235707904
GO:0031100: organ regeneration CCNE1 cyclin E1 0.0003736165667641785
GO:0031100: organ regeneration CXCL12 chemokine (C-X-C motif) ligand 12 -0.001213591761001856
GO:0031100: organ regeneration LEF1 lymphoid enhancer-binding factor 1 -0.00010032116575230784
GO:0031100: organ regeneration MED1 mediator complex subunit 1 0.001134772260258472
GO:0031100: organ regeneration MKI67 marker of proliferation Ki-67 0.0015788570868711287
GO:0031100: organ regeneration NOTCH1 notch 1 0.0005211581069085078
GO:0031100: organ regeneration PDX1 pancreatic and duodenal homeobox 1 0.00025893995986892785
GO:0031100: organ regeneration TGFB1 transforming growth factor, beta 1 -7.379122373448561e-5
GO:0031100: organ regeneration TYMS thymidylate synthetase 0.001569715464068976
GO:0031100: organ regeneration WNT1 wingless-type MMTV integration site family, member 1 0.0007895807975310406
GO:0031102: neuron projection regeneration ADM adrenomedullin 0.0022574560122283513
GO:0031623: receptor internalization ADM adrenomedullin 0.0022461762578210613
GO:0031623: receptor internalization GRB2 growth factor receptor-bound protein 2 0.0004316806068749084
GO:0031623: receptor internalization NEDD4 neural precursor cell expressed, developmentally down-regulated 4, E3 ubiquitin protein ligase 0.0022725957418444046
GO:0032496: response to lipopolysaccharide ADM adrenomedullin 0.0022475306611521566
GO:0032496: response to lipopolysaccharide CEBPB CCAAT/enhancer binding protein (C/EBP), beta -0.00027498273941970185
GO:0032496: response to lipopolysaccharide CITED1 Cbp/p300-interacting transactivator, with Glu/Asp-rich carboxy-terminal domain, 1 0.002831618417272932
GO:0032496: response to lipopolysaccharide CXCL13 chemokine (C-X-C motif) ligand 13 0.0030073685792690883
GO:0032496: response to lipopolysaccharide CYP27B1 cytochrome P450, family 27, subfamily B, polypeptide 1 -0.0004743644212478377
GO:0032496: response to lipopolysaccharide HDAC2 histone deacetylase 2 -0.0012109459890452521
GO:0032496: response to lipopolysaccharide HMGB2 high mobility group box 2 0.00031142181429570285
GO:0032496: response to lipopolysaccharide IDO1 indoleamine 2,3-dioxygenase 1 0.001232111705335171
GO:0032496: response to lipopolysaccharide IRAK1 interleukin-1 receptor-associated kinase 1 -0.0016163172384537738
GO:0032496: response to lipopolysaccharide JAK2 Janus kinase 2 -3.0720347629982354e-5
GO:0032496: response to lipopolysaccharide LGALS9 lectin, galactoside-binding, soluble, 9 -0.0011650656355459592
GO:0032496: response to lipopolysaccharide LIAS lipoic acid synthetase -0.0005563796193691328
GO:0032496: response to lipopolysaccharide NOTCH1 notch 1 0.0005213228839302018
GO:0032496: response to lipopolysaccharide S100A14 S100 calcium binding protein A14 -0.0021712557788089526
GO:0032496: response to lipopolysaccharide S100A7 S100 calcium binding protein A7 0.0015981945312622558
GO:0032496: response to lipopolysaccharide TFAP2A transcription factor AP-2 alpha (activating enhancer binding protein 2 alpha) 0.000481564113528641
GO:0032496: response to lipopolysaccharide TH tyrosine hydroxylase -0.00034304691041342326
GO:0032496: response to lipopolysaccharide TNFRSF11A tumor necrosis factor receptor superfamily, member 11a, NFKB activator 0.0027893462064071664
GO:0032868: response to insulin ADM adrenomedullin 0.0022571304207396298
GO:0032868: response to insulin AGRP agouti related protein homolog (mouse) -0.0025931567832848624
GO:0032868: response to insulin CITED1 Cbp/p300-interacting transactivator, with Glu/Asp-rich carboxy-terminal domain, 1 0.002845316550106531
GO:0032868: response to insulin EGR2 early growth response 2 0.0014564520686788356
GO:0032868: response to insulin GCNT1 glucosaminyl (N-acetyl) transferase 1, core 2 0.0007052307700962995
GO:0032868: response to insulin GGH gamma-glutamyl hydrolase (conjugase, folylpolygammaglutamyl hydrolase) -0.0003151803611604784
GO:0032868: response to insulin LEP leptin 0.0032240763975226907
GO:0032868: response to insulin LYN LYN proto-oncogene, Src family tyrosine kinase -0.001432791039888424
GO:0032868: response to insulin SIRT1 sirtuin 1 1.6888794087047267e-9
GO:0042475: odontogenesis of dentin-containing tooth ADM adrenomedullin 0.002237900733244428
GO:0042475: odontogenesis of dentin-containing tooth BAX BCL2-associated X protein -0.00042534434938371106
GO:0042475: odontogenesis of dentin-containing tooth BCL2L11 BCL2-like 11 (apoptosis facilitator) -0.0007749680304709949

GO:0042475: odontogenesis of dentin-containing tooth    BMP4    bone morphogenetic protein 4    -0.00
032256605062526487
GO:0042475: odontogenesis of dentin-containing tooth    BMP7    bone morphogenetic protein 7    0.000
8558030048578341
GO:0042475: odontogenesis of dentin-containing tooth    CTNNB1  catenin (cadherin-associated protei
n), beta 1, 88kDa       -0.00011709937218164794
GO:0042475: odontogenesis of dentin-containing tooth    EDA     ectodysplasin A -0.000814413405726350
4
GO:0042475: odontogenesis of dentin-containing tooth    EDAR    ectodysplasin A receptor        0.000
5940033102101945
GO:0042475: odontogenesis of dentin-containing tooth    FOXC1   forkhead box C1 -2.2580485670959423e-
5
GO:0042475: odontogenesis of dentin-containing tooth    GLI2    GLI family zinc finger 2        0.001
853532447217153
GO:0042475: odontogenesis of dentin-containing tooth    GLI3    GLI family zinc finger 3        -0.00
2151267354800685
GO:0042475: odontogenesis of dentin-containing tooth    HDAC2   histone deacetylase 2   -0.0012076448
724121487
GO:0042475: odontogenesis of dentin-containing tooth    JAG2    jagged 2        -5.740313696177113e-6
GO:0042475: odontogenesis of dentin-containing tooth    LEF1    lymphoid enhancer-binding factor 1
-0.00010029132528500807
GO:0042475: odontogenesis of dentin-containing tooth    LRP6    low density lipoprotein receptor-rela
ted protein 6   0.0001493770263094102
GO:0042475: odontogenesis of dentin-containing tooth    MSX1    msh homeobox 1  -0.002768754521346487
7
GO:0042475: odontogenesis of dentin-containing tooth    NF2     neurofibromin 2 (merlin)        -0.00
127214579751921
GO:0042475: odontogenesis of dentin-containing tooth    PITX2   paired-like homeodomain 2       0.002
176081270647936
GO:0042475: odontogenesis of dentin-containing tooth    SHH     sonic hedgehog  0.0005998055592181085
GO:0042475: odontogenesis of dentin-containing tooth    SMO     smoothened, frizzled class receptor
0.0021392087362287985
GO:0042475: odontogenesis of dentin-containing tooth    SOSTDC1 sclerostin domain containing 1  -0.00
15248043678562436
GO:0042594: response to starvation      ADM     adrenomedullin  0.0022490741292883693
GO:0042594: response to starvation      HMGCL   3-hydroxymethyl-3-methylglutaryl-CoA lyase      -0.00
10697342762814774
GO:0043065: positive regulation of apoptotic process    ADM     adrenomedullin  0.0022498471391765914
GO:0043065: positive regulation of apoptotic process    ALDH1A2 aldehyde dehydrogenase 1 family, memb
er A2   -0.0030351067118519898
GO:0043065: positive regulation of apoptotic process    APC     adenomatous polyposis coli      0.000
6454770127754742
GO:0043065: positive regulation of apoptotic process    ARL6IP5 ADP-ribosylation factor-like 6 intera
cting protein 5 0.0001698951896098909
GO:0043065: positive regulation of apoptotic process    BAK1    BCL2-antagonist/killer 1        -0.00
18861589047304058
GO:0043065: positive regulation of apoptotic process    BARD1   BRCA1 associated RING domain 1  0.001
7548024382458183
GO:0043065: positive regulation of apoptotic process    BAX     BCL2-associated X protein       -0.00
04269250851690821
GO:0043065: positive regulation of apoptotic process    BCL2L11 BCL2-like 11 (apoptosis facilitator)
-0.0007779063600343191
GO:0043065: positive regulation of apoptotic process    BMP4    bone morphogenetic protein 4    -0.00
032443722104737437
GO:0043065: positive regulation of apoptotic process    BMP7    bone morphogenetic protein 7    0.000
8594945586820191
GO:0043065: positive regulation of apoptotic process    BNIP3   BCL2/adenovirus E1B 19kDa interacting
protein 3       0.002929639518528973
GO:0043065: positive regulation of apoptotic process    BNIP3L  BCL2/adenovirus E1B 19kDa interacting
protein 3-like  -5.759519515641894e-5
GO:0043065: positive regulation of apoptotic process    CAMK1D  calcium/calmodulin-dependent protein
 kinase ID      -0.0026028522913662815
GO:0043065: positive regulation of apoptotic process    CDKN2A  cyclin-dependent kinase inhibitor 2A
0.0017700596267689462
GO:0043065: positive regulation of apoptotic process    CTNNB1  catenin (cadherin-associated protei
n), beta 1, 88kDa       -0.00011699420714832131
GO:0043065: positive regulation of apoptotic process    CYP1B1  cytochrome P450, family 1, subfamily
 B, polypeptide 1       0.00039937548439569987
GO:0043065: positive regulation of apoptotic process    DIABLO  diablo, IAP-binding mitochondrial pro
tein    -0.0005983876797996911
GO:0043065: positive regulation of apoptotic process    ECT2    epithelial cell transforming 2  0.001
0689481078891745
GO:0043065: positive regulation of apoptotic process    EEF1A2  eukaryotic translation elongation fac
tor 1 alpha 2   -0.0018661239554174094
GO:0043065: positive regulation of apoptotic process    FAS     Fas cell surface death receptor -3.34
9400984284591e-5
GO:0043065: positive regulation of apoptotic process    FOXO1   forkhead box O1 0.0018099178475307666
GO:0043065: positive regulation of apoptotic process    HMGA2   high mobility group AT-hook 2   0.001
5165756674686435
GO:0043065: positive regulation of apoptotic process    HMGB1   high mobility group box 1       -0.00
07791148104959676

GO:0043065: positive regulation of apoptotic process    HOXA5   homeobox A5     0.00107329093703349
GO:0043065: positive regulation of apoptotic process    IGFBP3  insulin-like growth factor binding pr
otein 3 0.0008397833620326271
GO:0043065: positive regulation of apoptotic process    ITGA6   integrin, alpha 6       0.00168251559
53329328
GO:0043065: positive regulation of apoptotic process    ITGB1   integrin, beta 1 (fibronectin recepto
r, beta polypeptide, antigen CD29 includes MDF2, MSK12) 0.0024421811923970907
GO:0043065: positive regulation of apoptotic process    KCNMA1  potassium large conductance calcium-a
ctivated channel, subfamily M, alpha member 1   -0.0006647765072681674
GO:0043065: positive regulation of apoptotic process    KLF11   Kruppel-like factor 11  0.00062282367
85852953
GO:0043065: positive regulation of apoptotic process    MAP3K5  mitogen-activated protein kinase kina
se kinase 5     0.000344806027095111
GO:0043065: positive regulation of apoptotic process    MELK    maternal embryonic leucine zipper kin
ase     0.002082269148067369
GO:0043065: positive regulation of apoptotic process    NOTCH1  notch 1 0.0005220117098999353
GO:0043065: positive regulation of apoptotic process    PRKDC   protein kinase, DNA-activated, cataly
tic polypeptide -0.0018647529353817878
GO:0043065: positive regulation of apoptotic process    RARG    retinoic acid receptor, gamma   -0.00
23319501212653655
GO:0043065: positive regulation of apoptotic process    RXRA    retinoid X receptor, alpha      0.001
1209440490071174
GO:0043065: positive regulation of apoptotic process    S100B   S100 calcium binding protein B  0.005
216725688016133
GO:0043065: positive regulation of apoptotic process    SAV1    salvador family WW domain containing
 protein 1      -0.002348211193288586
GO:0043065: positive regulation of apoptotic process    SFRP1   secreted frizzled-related protein 1
0.0012937941625074984
GO:0043065: positive regulation of apoptotic process    SHQ1    SHQ1, H/ACA ribonucleoprotein assembl
y factor        -0.00144300012592386
GO:0043065: positive regulation of apoptotic process    SIRT1   sirtuin 1       -6.269227723448736e-7
GO:0043065: positive regulation of apoptotic process    SLIT2   slit homolog 2 (Drosophila)     -0.00
16302965686240574
GO:0043065: positive regulation of apoptotic process    SOX4    SRY (sex determining region Y)-box 4
-3.684691026260636e-5
GO:0043065: positive regulation of apoptotic process    SPDEF   SAM pointed domain containing ETS tra
nscription factor       0.0031516094141339477
GO:0043065: positive regulation of apoptotic process    STEAP3  STEAP family member 3, metalloreducta
se      0.0026225694808750906
GO:0043065: positive regulation of apoptotic process    TGFB1   transforming growth factor, beta 1
-7.391976541094113e-5
GO:0043065: positive regulation of apoptotic process    TGFB3   transforming growth factor, beta 3
-0.001836027210878173
GO:0043065: positive regulation of apoptotic process    TGM2    transglutaminase 2      -0.0001824927
2792951072
GO:0043065: positive regulation of apoptotic process    TOP2A   topoisomerase (DNA) II alpha 170kDa
-8.027966129823008e-5
GO:0043065: positive regulation of apoptotic process    TP53    tumor protein p53       0.00118440814
16860876
GO:0043065: positive regulation of apoptotic process    WNT10B  wingless-type MMTV integration site f
amily, member 10B       -0.0013974797399692572
GO:0043065: positive regulation of apoptotic process    WT1     Wilms tumor 1   -0.000510557175069480
9
GO:0043116: negative regulation of vascular permeability        ADM     adrenomedullin  0.00224006065
84065475
GO:0043116: negative regulation of vascular permeability        ANGPT1  angiopoietin 1  0.00090158506
85944627
GO:0043116: negative regulation of vascular permeability        PTPRJ   protein tyrosine phosphatase,
receptor type, J        -0.00033895407554470367
GO:0043116: negative regulation of vascular permeability        SLIT2   slit homolog 2 (Drosophila)
-0.0016217891238139064
GO:0045906: negative regulation of vasoconstriction     ADM     adrenomedullin  0.0022669326078214743
GO:0045906: negative regulation of vasoconstriction     CX3CL1  chemokine (C-X3-C motif) ligand 1
0.0021518927541464596
GO:0045906: negative regulation of vasoconstriction     LEP     leptin  0.0032405431344564323
GO:0045909: positive regulation of vasodilation ADM     adrenomedullin  0.0022443629968690025
GO:0045909: positive regulation of vasodilation AGT     angiotensinogen (serpin peptidase inhibitor,
 clade A, member 8)     -0.0011387460881769527
GO:0045909: positive regulation of vasodilation ALOX12  arachidonate 12-lipoxygenase    -0.0019597816
912600816
GO:0045909: positive regulation of vasodilation GJA1    gap junction protein, alpha 1, 43kDa    -0.00
01642964009464253
GO:0045909: positive regulation of vasodilation HMOX1   heme oxygenase (decycling) 1    -0.0002185043
8694969522
GO:0046879: hormone secretion   ADM     adrenomedullin  0.0022574560122283513
GO:0048589: developmental growth        ADM     adrenomedullin  0.0022482873183828447
GO:0048589: developmental growth        ASPM    asp (abnormal spindle) homolog, microcephaly associat
ed (Drosophila) 1.8428209697634484e-5
GO:0048589: developmental growth        GATA3   GATA binding protein 3  -4.1866391024640146e-5
GO:0048589: developmental growth        GLI2    GLI family zinc finger 2        0.0018623475678098947
GO:0048589: developmental growth        GLI3    GLI family zinc finger 3        -0.002164746941102460

2
GO:0048589: developmental growth SOX10 SRY (sex determining region Y)-box 10 0.00019614568
829672119
GO:0048589: developmental growth STRA6 stimulated by retinoic acid 6 -0.001689000152505710
5
GO:0048589: developmental growth TYMS thymidylate synthetase 0.00157247530154634
GO:0050829: defense response to Gram-negative bacterium ADM adrenomedullin 0.002229597701253052
GO:0050829: defense response to Gram-negative bacterium AZU1 azurocidin 1 0.0001971389329927297
GO:0050829: defense response to Gram-negative bacterium HMGB2 high mobility group box 2 0.000
30346669977967156
GO:0050829: defense response to Gram-negative bacterium IL12B interleukin 12B 0.0012732256978753313
GO:0050829: defense response to Gram-negative bacterium MMP7 matrix metallopeptidase 7 (matrilysi
n, uterine) 0.0004242195853160619
GO:0050829: defense response to Gram-negative bacterium S100A7 S100 calcium binding protein A7 0.001
5833155525608468
GO:0050829: defense response to Gram-negative bacterium SERPINE1 serpin peptidase inhibitor, c
lade E (nexin, plasminogen activator inhibitor type 1), member 1 0.0001128146421674076
GO:0050830: defense response to Gram-positive bacterium ADM adrenomedullin 0.002225749123551417
GO:0050830: defense response to Gram-positive bacterium HMGB2 high mobility group box 2 0.000
3023446602309773
GO:0050830: defense response to Gram-positive bacterium IL27RA interleukin 27 receptor, alpha -0.00
04817948427053054
GO:0050830: defense response to Gram-positive bacterium MMP7 matrix metallopeptidase 7 (matrilysi
n, uterine) 0.000423897028710161
GO:0050830: defense response to Gram-positive bacterium SEH1L SEH1-like (S. cerevisiae) -0.00
07290475215124663
GO:0051384: response to glucocorticoid ADM adrenomedullin 0.002254255245069569
GO:0051384: response to glucocorticoid AREG amphiregulin 0.002475830499200363
GO:0051384: response to glucocorticoid BCHE butyrylcholinesterase -8.878500389435835e-6
GO:0051384: response to glucocorticoid BCL2 B-cell CLL/lymphoma 2 -4.979173179272878e-6
GO:0051384: response to glucocorticoid BMP4 bone morphogenetic protein 4 -0.000325111336767044
75
GO:0051384: response to glucocorticoid C3 complement component 3 0.002040502706986943
GO:0051384: response to glucocorticoid CAV1 caveolin 1, caveolae protein, 22kDa -0.0005380439
279837387
GO:0051384: response to glucocorticoid CTSV cathepsin V 0.0006611152623379375
GO:0051384: response to glucocorticoid FAS Fas cell surface death receptor -3.358561388999537e-5
GO:0051384: response to glucocorticoid IGFBP2 insulin-like growth factor binding protein 2, 36kDa
0.00014678157828740092
GO:0051384: response to glucocorticoid MDK midkine (neurite growth-promoting factor 2) 0.001
683973947861748
GO:0051384: response to glucocorticoid PDX1 pancreatic and duodenal homeobox 1 0.00026055980
019655653
GO:0051384: response to glucocorticoid RXRA retinoid X receptor, alpha 0.0011231159560783001
GO:0051384: response to glucocorticoid S100B S100 calcium binding protein B 0.005228628183890329
GO:0051384: response to glucocorticoid TAT tyrosine aminotransferase 0.0010617844211096926
GO:0051384: response to glucocorticoid TYMS thymidylate synthetase 0.001577414410666402
GO:0060670: branching involved in labyrinthine layer morphogenesis ADM adrenomedullin 0.002
235840336474218
GO:0060670: branching involved in labyrinthine layer morphogenesis FGFR2 fibroblast growth fac
tor receptor 2 0.0007620538491925126
GO:0060670: branching involved in labyrinthine layer morphogenesis GRB2 growth factor recepto
r-bound protein 2 0.00042949293077006694
GO:0060712: spongiotrophoblast layer development ADM adrenomedullin 0.0022633438948317305
GO:0060712: spongiotrophoblast layer development CITED1 Cbp/p300-interacting transactivator,
with Glu/Asp-rich carboxy-terminal domain, 1 0.002854266054516458
GO:0097084: vascular smooth muscle cell development ADM adrenomedullin 0.0022597339534806216
GO:0097084: vascular smooth muscle cell development HES1 hes family bHLH transcription factor
1 -0.0009206495187076086
GO:0097084: vascular smooth muscle cell development HEY2 hes-related family bHLH transcription
factor with YRPW motif 2 0.0027312608271300967
GO:2001214: positive regulation of vasculogenesis ADM adrenomedullin 0.0022574560122283513
GO:0000122: negative regulation of transcription from RNA polymerase II promoter AES amino
-terminal enhancer of split 0.0013551182258712234
GO:0000122: negative regulation of transcription from RNA polymerase II promoter ALX1 ALX h
omeobox 1 0.002294359415347118
GO:0000122: negative regulation of transcription from RNA polymerase II promoter ASCL1 achae
te-scute family bHLH transcription factor 1 -0.0014341810788094532
GO:0000122: negative regulation of transcription from RNA polymerase II promoter AURKB auror
a kinase B 0.00035102425374751103
GO:0000122: negative regulation of transcription from RNA polymerase II promoter BACH1 BTB a
nd CNC homology 1, basic leucine zipper transcription factor 1 0.0004858791321776827
GO:0000122: negative regulation of transcription from RNA polymerase II promoter BCL11A B-cel
l CLL/lymphoma 11A (zinc finger protein) -0.0009778037521593697
GO:0000122: negative regulation of transcription from RNA polymerase II promoter BMP4 bone
morphogenetic protein 4 -0.00032916655418938
GO:0000122: negative regulation of transcription from RNA polymerase II promoter CAV1 caveo
lin 1, caveolae protein, 22kDa -0.0005358515151663146
GO:0000122: negative regulation of transcription from RNA polymerase II promoter CCND1 cycli
n D1 -0.002636169611349596
GO:0000122: negative regulation of transcription from RNA polymerase II promoter CTNNB1 caten

in (cadherin-associated protein), beta 1, 88kDa -0.00011707202416043353
GO:0000122: negative regulation of transcription from RNA polymerase II promoter DACH1 dachshund family transcription factor 1 0.002090612041893036
GO:0000122: negative regulation of transcription from RNA polymerase II promoter DAXX death-domain associated protein 0.000892676060272952
GO:0000122: negative regulation of transcription from RNA polymerase II promoter DICER1 dicer 1, ribonuclease type III -1.608524241915997e-5
GO:0000122: negative regulation of transcription from RNA polymerase II promoter DKK1 dickkopf WNT signaling pathway inhibitor 1 -0.0001808628076458805
GO:0000122: negative regulation of transcription from RNA polymerase II promoter DMRT1 doublesex and mab-3 related transcription factor 1 0.0015062826956418272
GO:0000122: negative regulation of transcription from RNA polymerase II promoter E2F1 E2F transcription factor 1 0.0020999519103675314
GO:0000122: negative regulation of transcription from RNA polymerase II promoter E2F6 E2F transcription factor 6 0.0007183936207001569
GO:0000122: negative regulation of transcription from RNA polymerase II promoter E2F8 E2F transcription factor 8 0.0017125554910919414
GO:0000122: negative regulation of transcription from RNA polymerase II promoter E4F1 E4F transcription factor 1 0.0014829497580189296
GO:0000122: negative regulation of transcription from RNA polymerase II promoter EFNA1 ephrin-A1 -0.0005211819739195806
GO:0000122: negative regulation of transcription from RNA polymerase II promoter EGR1 early growth response 1 0.0010942465833379373
GO:0000122: negative regulation of transcription from RNA polymerase II promoter EN1 engrailed homeobox 1 -1.5604905726076602e-5
GO:0000122: negative regulation of transcription from RNA polymerase II promoter ENG endoglin 0.000827663470796291
GO:0000122: negative regulation of transcription from RNA polymerase II promoter ETS2 v-ets avian erythroblastosis virus E26 oncogene homolog 2 -0.001016821930638603
GO:0000122: negative regulation of transcription from RNA polymerase II promoter EZH2 enhancer of zeste 2 polycomb repressive complex 2 subunit -0.00013652193455570446
GO:0000122: negative regulation of transcription from RNA polymerase II promoter FGFR2 fibroblast growth factor receptor 2 0.000764087648215591
GO:0000122: negative regulation of transcription from RNA polymerase II promoter FGFR3 fibroblast growth factor receptor 3 0.00022303140380648315
GO:0000122: negative regulation of transcription from RNA polymerase II promoter FOXC2 forkhead box C2 (MFH-1, mesenchyme forkhead 1) 0.0017500454485836897
GO:0000122: negative regulation of transcription from RNA polymerase II promoter FOXM1 forkhead box M1 0.00019738474514374832
GO:0000122: negative regulation of transcription from RNA polymerase II promoter FOXO1 forkhead box O1 0.0018021228107439754
GO:0000122: negative regulation of transcription from RNA polymerase II promoter FOXO3 forkhead box O3 0.0012014358019426347
GO:0000122: negative regulation of transcription from RNA polymerase II promoter GATA2 GATA binding protein 2 -0.0004520628084075148
GO:0000122: negative regulation of transcription from RNA polymerase II promoter GATA3 GATA binding protein 3 -3.89969609605361e-5
GO:0000122: negative regulation of transcription from RNA polymerase II promoter GATA6 GATA binding protein 6 -2.816422090276687e-5
GO:0000122: negative regulation of transcription from RNA polymerase II promoter GFI1 growth factor independent 1 transcription repressor 0.001666098553113379
GO:0000122: negative regulation of transcription from RNA polymerase II promoter GLI2 GLI family zinc finger 2 0.0018555577160233195
GO:0000122: negative regulation of transcription from RNA polymerase II promoter GLI3 GLI family zinc finger 3 -0.002154025316445987
GO:0000122: negative regulation of transcription from RNA polymerase II promoter HDAC2 histone deacetylase 2 -0.0012083777929821993
GO:0000122: negative regulation of transcription from RNA polymerase II promoter HES1 hes family bHLH transcription factor 1 -0.0009119510641972799
GO:0000122: negative regulation of transcription from RNA polymerase II promoter HEXIM1 hexamethylene bis-acetamide inducible 1 0.00015296059642953415
GO:0000122: negative regulation of transcription from RNA polymerase II promoter HEY2 hes-related family bHLH transcription factor with YRPW motif 2 0.0027021025193000405
GO:0000122: negative regulation of transcription from RNA polymerase II promoter HINFP histone H4 transcription factor 0.0009935279983226885
GO:0000122: negative regulation of transcription from RNA polymerase II promoter HIPK2 homeodomain interacting protein kinase 2 0.0007715493022325743
GO:0000122: negative regulation of transcription from RNA polymerase II promoter HMGA2 high mobility group AT-hook 2 0.0015105061269616291
GO:0000122: negative regulation of transcription from RNA polymerase II promoter HMGB1 high mobility group box 1 -0.0007770585873115468
GO:0000122: negative regulation of transcription from RNA polymerase II promoter ID2 inhibitor of DNA binding 2, dominant negative helix-loop-helix protein 6.0102725365222885e-5
GO:0000122: negative regulation of transcription from RNA polymerase II promoter IFNG interferon, gamma -4.8930795400853005e-5
GO:0000122: negative regulation of transcription from RNA polymerase II promoter IGBP1 immunoglobulin (CD79A) binding protein 1 0.002852166382103166
GO:0000122: negative regulation of transcription from RNA polymerase II promoter IRF7 interferon regulatory factor 7 -0.001307882305971927З
GO:0000122: negative regulation of transcription from RNA polymerase II promoter ISL1 ISL LIM homeobox 1 7.891835877104978e-5

GO:0000122: negative regulation of transcription from RNA polymerase II promoter	KLF11	Kruppel-like factor 11	0.0006176055842431099
GO:0000122: negative regulation of transcription from RNA polymerase II promoter	LEF1	lymphoid enhancer-binding factor 1	-0.00010024651916415763
GO:0000122: negative regulation of transcription from RNA polymerase II promoter	LEP	leptin	0.0031947153487910497
GO:0000122: negative regulation of transcription from RNA polymerase II promoter	LRP8	low density lipoprotein receptor-related protein 8, apolipoprotein e receptor	-0.0009327662659727047
GO:0000122: negative regulation of transcription from RNA polymerase II promoter	MED1	mediator complex subunit 1	0.0011306282915268355
GO:0000122: negative regulation of transcription from RNA polymerase II promoter	MEF2C	myocyte enhancer factor 2C	0.0009745291680395773
GO:0000122: negative regulation of transcription from RNA polymerase II promoter	MSX1	msh homeobox 1	-0.00277207992396805
GO:0000122: negative regulation of transcription from RNA polymerase II promoter	MYC	v-myc avian myelocytomatosis viral oncogene homolog	-0.001137664461376058
GO:0000122: negative regulation of transcription from RNA polymerase II promoter	NFIB	nuclear factor I/B	0.0029227627622900342
GO:0000122: negative regulation of transcription from RNA polymerase II promoter	NFX1	nuclear transcription factor, X-box binding 1	-6.977924080879207e-5
GO:0000122: negative regulation of transcription from RNA polymerase II promoter	NOC2L	nucleolar complex associated 2 homolog (S. cerevisiae)	-0.00015122390490234492
GO:0000122: negative regulation of transcription from RNA polymerase II promoter	NOTCH1	notch 1	0.0005193083282218355
GO:0000122: negative regulation of transcription from RNA polymerase II promoter	NR1H3	nuclear receptor subfamily 1, group H, member 3	0.0008015691707731484
GO:0000122: negative regulation of transcription from RNA polymerase II promoter	NR2E1	nuclear receptor subfamily 2, group E, member 1	-0.0029779697158289634
GO:0000122: negative regulation of transcription from RNA polymerase II promoter	NRIP1	nuclear receptor interacting protein 1	0.0010722096159011536
GO:0000122: negative regulation of transcription from RNA polymerase II promoter	ORC2	origin recognition complex, subunit 2	-0.0006128708749429621
GO:0000122: negative regulation of transcription from RNA polymerase II promoter	PAX3	paired box 3 -0.00294824899374306
GO:0000122: negative regulation of transcription from RNA polymerase II promoter	PAX6	paired box 6 0.0019377480187608097
GO:0000122: negative regulation of transcription from RNA polymerase II promoter	PDX1	pancreatic and duodenal homeobox 1	0.0002576710689291967
GO:0000122: negative regulation of transcription from RNA polymerase II promoter	PER2	period circadian clock 2	0.0012395282488476761
GO:0000122: negative regulation of transcription from RNA polymerase II promoter	PHF14	PHD finger protein 14	0.003228289177588465
GO:0000122: negative regulation of transcription from RNA polymerase II promoter	PITX2	paired-like homeodomain 2	0.0021784794792241856
GO:0000122: negative regulation of transcription from RNA polymerase II promoter	PLK1	polo-like kinase 1	0.0010189579927959973
GO:0000122: negative regulation of transcription from RNA polymerase II promoter	PLK3	polo-like kinase 3	0.002566737133402384
GO:0000122: negative regulation of transcription from RNA polymerase II promoter	POU1F1	POU class 1 homeobox 1	0.00013762064483145787
GO:0000122: negative regulation of transcription from RNA polymerase II promoter	PPP1R13L	protein phosphatase 1, regulatory subunit 13 like	-0.001475378327993628
GO:0000122: negative regulation of transcription from RNA polymerase II promoter	PROX1	prospero homeobox 1	0.0011228040046543481
GO:0000122: negative regulation of transcription from RNA polymerase II promoter	PTCH1	patched 1	-6.590360048554083e-5
GO:0000122: negative regulation of transcription from RNA polymerase II promoter	RARG	retinoic acid receptor, gamma	-0.0023218968713357028
GO:0000122: negative regulation of transcription from RNA polymerase II promoter	RBPJ	recombination signal binding protein for immunoglobulin kappa J region	0.0009823683903396022
GO:0000122: negative regulation of transcription from RNA polymerase II promoter	RREB1	ras responsive element binding protein 1	-0.0019836430271720504
GO:0000122: negative regulation of transcription from RNA polymerase II promoter	RXRA	retinoid X receptor, alpha	0.001116294931667694
GO:0000122: negative regulation of transcription from RNA polymerase II promoter	S100A1	S100 calcium binding protein A1	0.001587087287117167
GO:0000122: negative regulation of transcription from RNA polymerase II promoter	SALL1	spalt-like transcription factor 1	-0.002440445299273784
GO:0000122: negative regulation of transcription from RNA polymerase II promoter	SHH	sonic hedgehog	0.0006007415421319319
GO:0000122: negative regulation of transcription from RNA polymerase II promoter	SIRT1	sirtuin 1	-1.5243308058532593e-6
GO:0000122: negative regulation of transcription from RNA polymerase II promoter	SIRT2	sirtuin 2	-0.000853211656384486
GO:0000122: negative regulation of transcription from RNA polymerase II promoter	SIX1	SIX homeobox 1	-0.0018719885232860087
GO:0000122: negative regulation of transcription from RNA polymerase II promoter	SKI	SKI proto-oncogene	-0.0006711169904449968
GO:0000122: negative regulation of transcription from RNA polymerase II promoter	SMO	smoothened, frizzled class receptor	0.002141537306566368

GO:0000122: negative regulation of transcription from RNA polymerase II promoter SOX11 SRY (sex determining region Y)-box 11 -0.00020923557087659575
GO:0000122: negative regulation of transcription from RNA polymerase II promoter SOX18 SRY (sex determining region Y)-box 18 0.0012897248055904994
GO:0000122: negative regulation of transcription from RNA polymerase II promoter SOX9 SRY (sex determining region Y)-box 9 -0.000519268949292811
GO:0000122: negative regulation of transcription from RNA polymerase II promoter SPDEF SAM pointed domain containing ETS transcription factor 0.0031401075821517353
GO:0000122: negative regulation of transcription from RNA polymerase II promoter TBX3 T-box 3 0.0012274167587069324
GO:0000122: negative regulation of transcription from RNA polymerase II promoter TCF7L2 transcription factor 7-like 2 (T-cell specific, HMG-box) 0.0005743624922343588
GO:0000122: negative regulation of transcription from RNA polymerase II promoter TFAP2A transcription factor AP-2 alpha (activating enhancer binding protein 2 alpha) 0.00048035891781791836
GO:0000122: negative regulation of transcription from RNA polymerase II promoter TFAP2C transcription factor AP-2 gamma (activating enhancer binding protein 2 gamma) 0.0010616603099897152
GO:0000122: negative regulation of transcription from RNA polymerase II promoter TFCP2L1 transcription factor CP2-like 1 0.0004495484372106236
GO:0000122: negative regulation of transcription from RNA polymerase II promoter TGFB1 transforming growth factor, beta 1 -7.309557659658689e-5
GO:0000122: negative regulation of transcription from RNA polymerase II promoter THRB thyroid hormone receptor, beta 0.0019459537768192002
GO:0000122: negative regulation of transcription from RNA polymerase II promoter TP53 tumor protein p53 0.0011771957338800397
GO:0000122: negative regulation of transcription from RNA polymerase II promoter TP73 tumor protein p73 0.0010290644601768275
GO:0000122: negative regulation of transcription from RNA polymerase II promoter TPR translocated promoter region, nuclear basket protein -0.0006977416686542843
GO:0000122: negative regulation of transcription from RNA polymerase II promoter TRPS1 trichorhinophalangeal syndrome I -0.00017814533556740556
GO:0000122: negative regulation of transcription from RNA polymerase II promoter TWIST1 twist family bHLH transcription factor 1 -0.0013431266966506335
GO:0000122: negative regulation of transcription from RNA polymerase II promoter VDR vitamin D (1,25- dihydroxyvitamin D3) receptor 0.0005640252179659062
GO:0000122: negative regulation of transcription from RNA polymerase II promoter VEGFA vascular endothelial growth factor A 0.0005957531111647731
GO:0000122: negative regulation of transcription from RNA polymerase II promoter VLDLR very low density lipoprotein receptor 0.0009540857561877538
GO:0000122: negative regulation of transcription from RNA polymerase II promoter VPS72 vacuolar protein sorting 72 homolog (S. cerevisiae) 0.0005591059954160047
GO:0000122: negative regulation of transcription from RNA polymerase II promoter WFS1 Wolfram syndrome 1 (wolframin) 0.0005773100865995263
GO:0000122: negative regulation of transcription from RNA polymerase II promoter WNT10B wingless-type MMTV integration site family, member 10B -0.0013901551851193837
GO:0000122: negative regulation of transcription from RNA polymerase II promoter WT1 Wilms tumor 1 -0.0005075491450663939
GO:0000122: negative regulation of transcription from RNA polymerase II promoter WWP2 WW domain containing E3 ubiquitin protein ligase 2 -0.0009581368781906831
GO:0000122: negative regulation of transcription from RNA polymerase II promoter WWTR1 WW domain containing transcription regulator 1 0.0009016010907040487
GO:0000122: negative regulation of transcription from RNA polymerase II promoter XBP1 X-box binding protein 1 0.0002675704031140032
GO:0000122: negative regulation of transcription from RNA polymerase II promoter YBX1 Y box binding protein 1 -0.0009146851717373776
GO:0000122: negative regulation of transcription from RNA polymerase II promoter YY1 YY1 transcription factor 0.0013417771859454137
GO:0000122: negative regulation of transcription from RNA polymerase II promoter ZBTB18 zinc finger and BTB domain containing 18 0.001273315113422191
GO:0000122: negative regulation of transcription from RNA polymerase II promoter ZNF148 zinc finger protein 148 0.0021063360980877795
GO:0000122: negative regulation of transcription from RNA polymerase II promoter ZNF205 zinc finger protein 205 0.0005964780761849501
GO:0009887: organ morphogenesis AES amino-terminal enhancer of split 0.0013551700108189613
GO:0009887: organ morphogenesis CCL2 chemokine (C-C motif) ligand 2 0.0008169353018639718
GO:0009887: organ morphogenesis E2F4 E2F transcription factor 4, p107/p130-binding -0.0023475824803697344
GO:0009887: organ morphogenesis EVL Enah/Vasp-like 0.0019399869451577245
GO:0009887: organ morphogenesis FGFR2 fibroblast growth factor receptor 2 0.0007636188960153763
GO:0009887: organ morphogenesis GATA2 GATA binding protein 2 -0.0004513740507206312
GO:0009887: organ morphogenesis GATA3 GATA binding protein 3 -3.8121562623495844e-5
GO:0009887: organ morphogenesis GSK3B glycogen synthase kinase 3 beta 0.0015504609697309835
GO:0009887: organ morphogenesis IL7 interleukin 7 0.0008715709758289297
GO:0009887: organ morphogenesis LHX1 LIM homeobox 1 -0.0007592971182673821
GO:0009887: organ morphogenesis NRP1 neuropilin 1 -0.0006436307598317075
GO:0009887: organ morphogenesis PAX3 paired box 3 -0.00294760827572801
GO:0009887: organ morphogenesis PAX6 paired box 6 0.0019368754649154137
GO:0009887: organ morphogenesis PDX1 pancreatic and duodenal homeobox 1 0.00025736213537804305
GO:0009887: organ morphogenesis PTCH1 patched 1 -6.626513105950112e-5
GO:0009887: organ morphogenesis STX2 syntaxin 2 0.00030844837225686976

GO:0009887: organ morphogenesis TBX3    T-box 3 0.0012278099074311084
GO:0009887: organ morphogenesis TH      tyrosine hydroxylase    -0.0003424167802420946
GO:0009887: organ morphogenesis THRB    thyroid hormone receptor, beta  0.0019457902119361342
GO:0009887: organ morphogenesis TRPS1   trichorhinophalangeal syndrome I        -0.00017755481586765787
GO:0009887: organ morphogenesis VEGFC   vascular endothelial growth factor C    -0.003366471390167560
4
GO:0010629: negative regulation of gene expression      AES     amino-terminal enhancer of split
0.001351914001093741З
GO:0010629: negative regulation of gene expression      BAK1    BCL2-antagonist/killer 1        -0.00
18730431728721155
GO:0010629: negative regulation of gene expression      BBS4    Bardet-Biedl syndrome 4 -0.0005090853
651926168
GO:0010629: negative regulation of gene expression      C1QTNF3 C1q and tumor necrosis factor related
protein 3       0.0001389293275212743
GO:0010629: negative regulation of gene expression      CCNB1   cyclin B1       -0.000880933288522791
7
GO:0010629: negative regulation of gene expression      CDC42   cell division cycle 42  0.00123828978
89685952
GO:0010629: negative regulation of gene expression      CRYAB   crystallin, alpha B     0.00098302226
52376516
GO:0010629: negative regulation of gene expression      ESR1    estrogen receptor 1     -0.0009468039
846844984
GO:0010629: negative regulation of gene expression      GJA1    gap junction protein, alpha 1, 43kDa
-0.00016121412881582508
GO:0010629: negative regulation of gene expression      HEY2    hes-related family bHLH transcription
factor with YRPW motif 2        0.002694421435013309
GO:0010629: negative regulation of gene expression      HINFP   histone H4 transcription factor 0.000
9911689595940656
GO:0010629: negative regulation of gene expression      ID2     inhibitor of DNA binding 2, dominant
negative helix-loop-helix protein      6.0115906696228944e-5
GO:0010629: negative regulation of gene expression      IFNA2   interferon, alpha 2     -0.0017146287
008386837
GO:0010629: negative regulation of gene expression      IFNG    interferon, gamma       -4.8832332093
85635e-5
GO:0010629: negative regulation of gene expression      LGALS9  lectin, galactoside-binding, soluble,
9       -0.00011678216342476742
GO:0010629: negative regulation of gene expression      MEF2C   myocyte enhancer factor 2C      0.000
9717040659472457
GO:0010629: negative regulation of gene expression      PGR     progesterone receptor   0.00035122284
7724556
GO:0010629: negative regulation of gene expression      RNASEH2B        ribonuclease H2, subunit B
-4.346778081713118e-5
GO:0010629: negative regulation of gene expression      SFRP1   secreted frizzled-related protein 1
0.0012794387325746083
GO:0010629: negative regulation of gene expression      SIRT1   sirtuin 1       -1.7852213432193776e-
6
GO:0010629: negative regulation of gene expression      SLIT2   slit homolog 2 (Drosophila)     -0.00
16175920743850968
GO:0010629: negative regulation of gene expression      SMO     smoothened, frizzled class receptor
0.0021359331492284035
GO:0010629: negative regulation of gene expression      SOX11   SRY (sex determining region Y)-box 11
-0.000208673276676616
GO:0010629: negative regulation of gene expression      STC2    stanniocalcin 2 -0.001395134547043584
6
GO:0010629: negative regulation of gene expression      TGFB1   transforming growth factor, beta 1
-7.294610457040552e-5
GO:0010629: negative regulation of gene expression      WNT4    wingless-type MMTV integration site f
amily, member 4 -0.00025154255592147
GO:0010629: negative regulation of gene expression      WWP2    WW domain containing E3 ubiquitin pro
tein ligase 2   -0.0009557896463014892
GO:0010629: negative regulation of gene expression      YY1     YY1 transcription factor        0.001
3378788044023022
GO:0010629: negative regulation of gene expression      ZNF148  zinc finger protein 148 0.00210067804
8886488
GO:0016055: Wnt signaling pathway       AES     amino-terminal enhancer of split        0.00135273951
66038423
GO:0016055: Wnt signaling pathway       CCNE1   cyclin E1       0.00037081171272804143
GO:0016055: Wnt signaling pathway       CD24    CD24 molecule   0.0010707816370809054
GO:0016055: Wnt signaling pathway       CELSR2  cadherin, EGF LAG seven-pass G-type receptor 2  -0.00
11340597826993827
GO:0016055: Wnt signaling pathway       CPE     carboxypeptidase E      -0.0025422313126101308
GO:0016055: Wnt signaling pathway       CSNK2B  casein kinase 2, beta polypeptide       0.00152735385
0485443
GO:0016055: Wnt signaling pathway       CTNNB1  catenin (cadherin-associated protein), beta 1, 88kDa
-0.0001167708523579361
GO:0016055: Wnt signaling pathway       CTNNBIP1        catenin, beta interacting protein 1     0.000
5851476176684198
GO:0016055: Wnt signaling pathway       DRD2    dopamine receptor D2    -0.0002339062974386067
GO:0016055: Wnt signaling pathway       LEF1    lymphoid enhancer-binding factor 1      -0.0001003597
2892575306

GO:0016055: Wnt signaling pathway LRP5 low density lipoprotein receptor-related protein 5 3.0445528481599446e-5
GO:0016055: Wnt signaling pathway LRP6 low density lipoprotein receptor-related protein 6 0.00014952042784598294
GO:0016055: Wnt signaling pathway NDRG2 NDRG family member 2 0.0021715200009711014
GO:0016055: Wnt signaling pathway PITX2 paired-like homeodomain 2 0.0021740623016134678
GO:0016055: Wnt signaling pathway SOSTDC1 sclerostin domain containing 1 -0.0015229194321765072
GO:0016055: Wnt signaling pathway WNT1 wingless-type MMTV integration site family, member 1 0.0007862487173985866
GO:0016055: Wnt signaling pathway WNT10B wingless-type MMTV integration site family, member 10B -0.0013872120620972372
GO:0016055: Wnt signaling pathway WNT5A wingless-type MMTV integration site family, member 5A -0.0006668920517706576
GO:0032091: negative regulation of protein binding AES amino-terminal enhancer of split 0.0013463307260487638
GO:0032091: negative regulation of protein binding AURKA aurora kinase A 0.0009991882076638276
GO:0032091: negative regulation of protein binding AURKB aurora kinase B 0.0003460323517961098
GO:0032091: negative regulation of protein binding BAX BCL2-associated X protein -0.00042289228848572014
GO:0032091: negative regulation of protein binding CAV1 caveolin 1, caveolae protein, 22kDa -0.0005329481437877453
GO:0032091: negative regulation of protein binding CTNNBIP1 catenin, beta interacting protein 1 0.0005811182393533245
GO:0032091: negative regulation of protein binding GOLGA2 golgin A2 0.00016704976704624974
GO:0032091: negative regulation of protein binding GSK3B glycogen synthase kinase 3 beta 0.0015382435686406635
GO:0032091: negative regulation of protein binding NES nestin 0.00136080464634235
GO:0032091: negative regulation of protein binding PRKCD protein kinase C, delta -0.0011190671462786529
GO:0032091: negative regulation of protein binding TEX14 testis expressed 14 0.0017153611552606792
GO:0032091: negative regulation of protein binding TMBIM6 transmembrane BAX inhibitor motif containing 6 -0.00014697587107788531
GO:0045892: negative regulation of transcription, DNA-templated AES amino-terminal enhancer of split 0.0013517847305740715
GO:0045892: negative regulation of transcription, DNA-templated ASCL1 achaete-scute family bHLH transcription factor 1 -0.001429550917306822
GO:0045892: negative regulation of transcription, DNA-templated ATF5 activating transcription factor 5 -0.0027301283776263673
GO:0045892: negative regulation of transcription, DNA-templated BASP1 brain abundant, membrane attached signal protein 1 0.0004775931535688067
GO:0045892: negative regulation of transcription, DNA-templated BIRC5 baculoviral IAP repeat containing 5 -0.0002603185906516552
GO:0045892: negative regulation of transcription, DNA-templated BMP4 bone morphogenetic protein 4 -0.00032199024647346504
GO:0045892: negative regulation of transcription, DNA-templated BMP7 bone morphogenetic protein 7 0.0008543523023319434
GO:0045892: negative regulation of transcription, DNA-templated CDKN2A cyclin-dependent kinase inhibitor 2A 0.0017543369588370777
GO:0045892: negative regulation of transcription, DNA-templated CEBPB CCAAT/enhancer binding protein (C/EBP), beta -0.0002735178826884574
GO:0045892: negative regulation of transcription, DNA-templated CHMP1A charged multivesicular body protein 1A -0.0007783748268909477
GO:0045892: negative regulation of transcription, DNA-templated CITED1 Cbp/p300-interacting transactivator, with Glu/Asp-rich carboxy-terminal domain, 1 0.0028126282879064957
GO:0045892: negative regulation of transcription, DNA-templated CLOCK clock circadian regulator 0.0002001716583007587
GO:0045892: negative regulation of transcription, DNA-templated CTNNB1 catenin (cadherin-associated protein), beta 1, 88kDa -0.0001168607762602405
GO:0045892: negative regulation of transcription, DNA-templated DACH1 dachshund family transcription factor 1 0.002085597153487456
GO:0045892: negative regulation of transcription, DNA-templated DAXX death-domain associated protein 0.0008895562116497331
GO:0045892: negative regulation of transcription, DNA-templated E2F1 E2F transcription factor 1 0.002093653136243879
GO:0045892: negative regulation of transcription, DNA-templated EZH2 enhancer of zeste 2 polycomb repressive complex 2 subunit -0.00013672358495650795
GO:0045892: negative regulation of transcription, DNA-templated FOXM1 forkhead box M1 0.00019648954157836876
GO:0045892: negative regulation of transcription, DNA-templated FOXO1 forkhead box O1 0.0017969593486101645
GO:0045892: negative regulation of transcription, DNA-templated GATA3 GATA binding protein 3 -3.864292371758217e-5
GO:0045892: negative regulation of transcription, DNA-templated GATA6 GATA binding protein 6 -2.8335332695295194e-5
GO:0045892: negative regulation of transcription, DNA-templated GFI1 growth factor independent 1 transcription repressor 0.0016618497178077427
GO:0045892: negative regulation of transcription, DNA-templated GLI3 GLI family zinc finger 3 -0.0021471929500517495

GO:0045892: negative regulation of transcription, DNA-templated GREM1 gremlin 1, DAN family BMP antagonist -0.0008252087915197292
GO:0045892: negative regulation of transcription, DNA-templated HDAC2 histone deacetylase 2 -0.0012055610861570983
GO:0045892: negative regulation of transcription, DNA-templated HES1 hes family bHLH transcription factor 1 -0.0009092210165938036
GO:0045892: negative regulation of transcription, DNA-templated HEXIM1 hexamethylene bis-acetamide inducible 1 0.0001524773705220633
GO:0045892: negative regulation of transcription, DNA-templated HEY2 hes-related family bHLH transcription factor with YRPW motif 2 0.00269317760151747 8
GO:0045892: negative regulation of transcription, DNA-templated HEYL hes-related family bHLH transcription factor with YRPW motif-like 0.0006763193724865658
GO:0045892: negative regulation of transcription, DNA-templated HINFP histone H4 transcription factor 0.000990620591567639
GO:0045892: negative regulation of transcription, DNA-templated HMGA1 high mobility group AT-hook 1 -0.0003182893538899945
GO:0045892: negative regulation of transcription, DNA-templated HMGA2 high mobility group AT-hook 2 0.001506300927691984
GO:0045892: negative regulation of transcription, DNA-templated HMGB2 high mobility group box 2 0.00030554420025999315
GO:0045892: negative regulation of transcription, DNA-templated ID2 inhibitor of DNA binding 2, dominant negative helix-loop-helix protein 6.017573893963391e-5
GO:0045892: negative regulation of transcription, DNA-templated IFNA2 interferon, alpha 2 -0.0017141527027006906
GO:0045892: negative regulation of transcription, DNA-templated IL4 interleukin 4 0.0002594261121362926
GO:0045892: negative regulation of transcription, DNA-templated LANCL2 LanC lantibiotic synthetase component C-like 2 (bacterial) 0.0011831124860059632
GO:0045892: negative regulation of transcription, DNA-templated LEF1 lymphoid enhancer-binding factor 1 -0.00010028864894957717
GO:0045892: negative regulation of transcription, DNA-templated LGR4 leucine-rich repeat containing G protein-coupled receptor 4 0.00030794274591630817
GO:0045892: negative regulation of transcription, DNA-templated LHX1 LIM homeobox 1 -0.0007571991675023071
GO:0045892: negative regulation of transcription, DNA-templated NOTCH1 notch 1 0.0005177525193474505
GO:0045892: negative regulation of transcription, DNA-templated PAX2 paired box 2 -0.001610811635700988
GO:0045892: negative regulation of transcription, DNA-templated PER2 period circadian clock 2 0.0012358609273226844
GO:0045892: negative regulation of transcription, DNA-templated PHB2 prohibitin 2 -0.0007348650885099741
GO:0045892: negative regulation of transcription, DNA-templated PLCB1 phospholipase C, beta 1 (phosphoinositide-specific) 0.0001531790128339127
GO:0045892: negative regulation of transcription, DNA-templated PML promyelocytic leukemia -0.000679144432353243
GO:0045892: negative regulation of transcription, DNA-templated PROX1 prospero homeobox 1 0.0011191792502334794
GO:0045892: negative regulation of transcription, DNA-templated PURA purine-rich element binding protein A -0.0001699504896944985
GO:0045892: negative regulation of transcription, DNA-templated RB1 retinoblastoma 1 -0.0014911611868604366
GO:0045892: negative regulation of transcription, DNA-templated RBPJ recombination signal binding protein for immunoglobulin kappa J region 0.0009799788864599793
GO:0045892: negative regulation of transcription, DNA-templated SALL1 spalt-like transcription factor 1 -0.002431822842432363
GO:0045892: negative regulation of transcription, DNA-templated SFRP1 secreted frizzled-related protein 1 0.001278059449732701
GO:0045892: negative regulation of transcription, DNA-templated SIRT1 sirtuin 1 -1.9291066704486976e-6
GO:0045892: negative regulation of transcription, DNA-templated SIRT2 sirtuin 2 -0.0008505944997300519
GO:0045892: negative regulation of transcription, DNA-templated SIX3 SIX homeobox 3 0.0020329987152231705
GO:0045892: negative regulation of transcription, DNA-templated SOX10 SRY (sex determining region Y)-box 10 0.00018912903029736052
GO:0045892: negative regulation of transcription, DNA-templated SOX18 SRY (sex determining region Y)-box 18 0.001286436170074658
GO:0045892: negative regulation of transcription, DNA-templated SOX9 SRY (sex determining region Y)-box 9 -0.0005176144539026881
GO:0045892: negative regulation of transcription, DNA-templated TBX3 T-box 3 0.0012245786504059763
GO:0045892: negative regulation of transcription, DNA-templated TCF7L2 transcription factor 7-like 2 (T-cell specific, HMG-box) 0.0005723457182564315
GO:0045892: negative regulation of transcription, DNA-templated TFAP2A transcription factor AP-2 alpha (activating enhancer binding protein 2 alpha) 0.00047917332791790187
GO:0045892: negative regulation of transcription, DNA-templated TGFB1 transforming growth factor, beta 1 -7.275729444127835e-5
GO:0045892: negative regulation of transcription, DNA-templated THRA thyroid hormone receptor, alpha 0.0007619017160781876
GO:0045892: negative regulation of transcription, DNA-templated TP53 tumor protein p53 0.0011729599534958286
GO:0045892: negative regulation of transcription, DNA-templated TWIST1 twist family bHLH transcripti

on factor 1	-0.001338717112133744
GO:0045892: negative regulation of transcription, DNA-templated VDR	vitamin D (1,25- dihydroxyvitamin D3) receptor	0.000562819871141773
GO:0045892: negative regulation of transcription, DNA-templated WNT4	wingless-type MMTV integration site family, member 4 -0.0002510665715639827
GO:0045892: negative regulation of transcription, DNA-templated WNT5A	wingless-type MMTV integration site family, member 5A	-0.0006660780260055668
GO:0045892: negative regulation of transcription, DNA-templated WT1	Wilms tumor 1	-0.0005057158283539181
GO:0045892: negative regulation of transcription, DNA-templated WWP2	WW domain containing E3 ubiquitin protein ligase 2	-0.0009555623803290933
GO:0045892: negative regulation of transcription, DNA-templated ZBTB18	zinc finger and BTB domain containing 18	0.001269882572929039
GO:0045892: negative regulation of transcription, DNA-templated ZNF148	zinc finger protein 148 0.0021000348961236655
GO:0045892: negative regulation of transcription, DNA-templated ZNF24	zinc finger protein 24	0.0012894295712332707
GO:0060761: negative regulation of response to cytokine stimulus	AES	amino-terminal enhancer of split	0.0013463150483353046
GO:0070555: response to interleukin-1	AES	amino-terminal enhancer of split	0.0013629143706802712
GO:0070555: response to interleukin-1	CITED1	Cbp/p300-interacting transactivator, with Glu/Asp-rich carboxy-terminal domain, 1	0.002841623051359837
GO:0070555: response to interleukin-1	IGBP1	immunoglobulin (CD79A) binding protein 1	0.0028698158549162107
GO:0070555: response to interleukin-1	IRAK1	interleukin-1 receptor-associated kinase 1	-0.001622894614721151
GO:0070555: response to interleukin-1	LGALS9	lectin, galactoside-binding, soluble, 9 -0.0001166480465430497
GO:0070555: response to interleukin-1	PRKCA	protein kinase C, alpha -6.143989571452003e-6
GO:0070555: response to interleukin-1	TNFRSF11A	tumor necrosis factor receptor superfamily, member 11a, NFKB activator	0.0027976703073307307
GO:0090090: negative regulation of canonical Wnt signaling pathway	AES	amino-terminal enhancer of split	0.0013446295997899033
GO:0090090: negative regulation of canonical Wnt signaling pathway	APC	adenomatous polyposis coli	0.0006405561210755343
GO:0090090: negative regulation of canonical Wnt signaling pathway	AXIN1	axin 1	-0.0007259334725522165
GO:0090090: negative regulation of canonical Wnt signaling pathway	CAV1	caveolin 1, caveolae protein, 22kDa -0.0005323052824182281
GO:0090090: negative regulation of canonical Wnt signaling pathway	CDH2	cadherin 2, type 1, N-cadherin (neuronal)	-0.0006365518356656022
GO:0090090: negative regulation of canonical Wnt signaling pathway	DKK1	dickkopf WNT signaling pathway inhibitor 1	-0.0001796534172234697
GO:0090090: negative regulation of canonical Wnt signaling pathway	EGR1	early growth response 1	0.0010843701458420835
GO:0090090: negative regulation of canonical Wnt signaling pathway	FOXO1	forkhead box O1 0.001785897385355353
GO:0090090: negative regulation of canonical Wnt signaling pathway	FOXO3	forkhead box O3 0.0011896221452105983
GO:0090090: negative regulation of canonical Wnt signaling pathway	GLI1	GLI family zinc finger 1	-0.0012874867377596444
GO:0090090: negative regulation of canonical Wnt signaling pathway	GLI3	GLI family zinc finger 3	-0.002132751476636087
GO:0090090: negative regulation of canonical Wnt signaling pathway	GREM1	gremlin 1, DAN family BMP antagonist	-0.0008181393414796463
GO:0090090: negative regulation of canonical Wnt signaling pathway	GSK3B	glycogen synthase kinase 3 beta	0.0015363067664930818
GO:0090090: negative regulation of canonical Wnt signaling pathway	ISL1	ISL LIM homeobox 1	7.974583911044472e-5
GO:0090090: negative regulation of canonical Wnt signaling pathway	LEF1	lymphoid enhancer-binding factor 1	-0.00010044236999972487
GO:0090090: negative regulation of canonical Wnt signaling pathway	LRP6	low density lipoprotein receptor-related protein 6	0.00014822228708482926
GO:0090090: negative regulation of canonical Wnt signaling pathway	NOTCH1	notch 1 0.0005145056565424624
GO:0090090: negative regulation of canonical Wnt signaling pathway	PPP2R3A protein phosphatase 2, regulatory subunit B'', alpha	-3.40172345663179e-5
GO:0090090: negative regulation of canonical Wnt signaling pathway	PSMA5	proteasome (prosome, macropain) subunit, alpha type, 5	0.000329073166588321
GO:0090090: negative regulation of canonical Wnt signaling pathway	PSMD11	proteasome (prosome, macropain) 26S subunit, non-ATPase, 11 -0.0010879595894439224
GO:0090090: negative regulation of canonical Wnt signaling pathway	PSMD13	proteasome (prosome, macropain) 26S subunit, non-ATPase, 13 -6.047868398404916e-5
GO:0090090: negative regulation of canonical Wnt signaling pathway	SFRP1	secreted frizzled-related protein 1	0.0012664764144225064
GO:0090090: negative regulation of canonical Wnt signaling pathway	SHH	sonic hedgehog	0.0005936221018870158
GO:0090090: negative regulation of canonical Wnt signaling pathway	SOSTDC1 sclerostin domain containing 1	-0.001516383593033325
GO:0090090: negative regulation of canonical Wnt signaling pathway	SOX10	SRY (sex determining

region Y)-box 10 0.0001853021654686115
GO:0090090: negative regulation of canonical Wnt signaling pathway SOX9 SRY (sex determining region Y)-box 9 -0.0005139641789025874
GO:0090090: negative regulation of canonical Wnt signaling pathway TCF7L2 transcription factor 7-like 2 (T-cell specific, HMG-box) 0.0005681147050651933
GO:0090090: negative regulation of canonical Wnt signaling pathway WNT4 wingless-type MMTV integration site family, member 4 -0.00024806568642926863
GO:0090090: negative regulation of canonical Wnt signaling pathway WNT5A wingless-type MMTV integration site family, member 5A -0.0006616863540872482
GO:0090090: negative regulation of canonical Wnt signaling pathway WWTR1 WW domain containing transcription regulator 1 0.0008909372223944112
GO:2000210: positive regulation of anoikis AES amino-terminal enhancer of split 0.0013463150483353046
GO:0051180: vitamin transport AFM afamin -0.0016939199422974886
GO:0001933: negative regulation of protein phosphorylation AGER advanced glycosylation end product-specific receptor -0.00017358060487045516
GO:0001933: negative regulation of protein phosphorylation ANGPT1 angiopoietin 1 0.0008942613539592709
GO:0001933: negative regulation of protein phosphorylation CCNB1 cyclin B1 -0.0008789843383837513
GO:0001933: negative regulation of protein phosphorylation CDK5RAP3 CDK5 regulatory subunit associated protein 3 0.001130476766689401
GO:0001933: negative regulation of protein phosphorylation CIB1 calcium and integrin binding 1 (calmyrin) 5.769625484162519e-5
GO:0001933: negative regulation of protein phosphorylation IGFBP3 insulin-like growth factor binding protein 3 0.000832480956160039
GO:0001933: negative regulation of protein phosphorylation LRP6 low density lipoprotein receptor-related protein 6 0.00014759079330597496
GO:0001933: negative regulation of protein phosphorylation LYN LYN proto-oncogene, Src family tyrosine kinase -0.001409547889793704
GO:0001933: negative regulation of protein phosphorylation MAS1 MAS1 proto-oncogene, G protein-coupled receptor 0.0002936659677934921
GO:0001933: negative regulation of protein phosphorylation PAX6 paired box 6 0.0019193292556878086
GO:0001933: negative regulation of protein phosphorylation PRKDC protein kinase, DNA-activated, catalytic polypeptide -0.0018424092921994434
GO:0001933: negative regulation of protein phosphorylation PTEN phosphatase and tensin homolog 1.98489635151551e-5
GO:0001933: negative regulation of protein phosphorylation SLIT2 slit homolog 2 (Drosophila) -0.0016053719849884749
GO:0001933: negative regulation of protein phosphorylation TGFB1 transforming growth factor, beta 1 -7.162701065306363e-5
GO:0001933: negative regulation of protein phosphorylation WWTR1 WW domain containing transcription regulator 1 0.0008893523319973961
GO:0006954: inflammatory response AGER advanced glycosylation end product-specific receptor -0.00017416051933252468
GO:0006954: inflammatory response AKT1 v-akt murine thymoma viral oncogene homolog 1 0.0007293226205558227
GO:0006954: inflammatory response AZU1 azurocidin 1 0.00019770024192330228
GO:0006954: inflammatory response C3 complement component 3 0.0020214935294109795
GO:0006954: inflammatory response CAMK1D calcium/calmodulin-dependent protein kinase ID -0.0025835848378670543
GO:0006954: inflammatory response CCL2 chemokine (C-C motif) ligand 2 0.0008151701822056872
GO:0006954: inflammatory response CCL7 chemokine (C-C motif) ligand 7 -0.002400020929641093
GO:0006954: inflammatory response CCL8 chemokine (C-C motif) ligand 8 -0.0005901406027813148
GO:0006954: inflammatory response CEBPB CCAAT/enhancer binding protein (C/EBP), beta -0.0002738361398219049
GO:0006954: inflammatory response CXCL10 chemokine (C-X-C motif) ligand 10 6.185463254846114e-5
GO:0006954: inflammatory response CXCL13 chemokine (C-X-C motif) ligand 13 0.002988368130941916
GO:0006954: inflammatory response CXCR4 chemokine (C-X-C motif) receptor 4 0.0007930653805844551
GO:0006954: inflammatory response ECM1 extracellular matrix protein 1 -0.0015538446464899178
GO:0006954: inflammatory response HMGB1 high mobility group box 1 -0.0007758328482250261
GO:0006954: inflammatory response IFNA2 interferon, alpha 2 -0.0017147744852922713
GO:0006954: inflammatory response IGFBP4 insulin-like growth factor binding protein 4 -0.0012759153273434404
GO:0006954: inflammatory response KIT v-kit Hardy-Zuckerman 4 feline sarcoma viral oncogene homolog 0.00027329672935830077
GO:0006954: inflammatory response LGALS9 lectin, galactoside-binding, soluble, 9 -0.00011702440917983022
GO:0006954: inflammatory response LIAS lipoic acid synthetase -0.000553396380485162
GO:0006954: inflammatory response NFX1 nuclear transcription factor, X-box binding 1 -7.071783108679612e-5
GO:0006954: inflammatory response ORM1 orosomucoid 1 -0.00047229648387088675
GO:0006954: inflammatory response PIK3CD phosphatidylinositol-4,5-bisphosphate 3-kinase, catalytic subunit delta -0.000779620615391681

GO:0006954: inflammatory response	PRKCZ	protein kinase C, zeta	-0.001598584458870639
GO:0006954: inflammatory response	RXRA	retinoid X receptor, alpha	0.0011135967399531438
GO:0006954: inflammatory response	SPHK1	sphingosine kinase 1	0.0018079082150883497
GO:0006954: inflammatory response	TGFB1	transforming growth factor, beta 1	-7.2650896599
43394e-5
GO:0006954: inflammatory response	THBS1	thrombospondin 1	-0.0010339970738101815
GO:0006954: inflammatory response	TNFAIP3 tumor necrosis factor, alpha-induced protein 3	0.001
1147675776687195
GO:0006954: inflammatory response	TP73	tumor protein p73	0.001026395025305485
GO:0007166: cell surface receptor signaling pathway	AGER	advanced glycosylation end product-sp
ecific receptor -0.00017473212348881833
GO:0007166: cell surface receptor signaling pathway	AGT	angiotensinogen (serpin peptidase inh
ibitor, clade A, member 8)	-0.0011409484028379242
GO:0007166: cell surface receptor signaling pathway	CCL2	chemokine (C-C motif) ligand 2	0.000
8207016857744452
GO:0007166: cell surface receptor signaling pathway	CD9	CD9 molecule	-0.002598572049718699
GO:0007166: cell surface receptor signaling pathway	CXCL10	chemokine (C-X-C motif) ligand 10
6.367702381689456e-5
GO:0007166: cell surface receptor signaling pathway	CXCL13	chemokine (C-X-C motif) ligand 13
0.0030085060487865323
GO:0007166: cell surface receptor signaling pathway	EGFR	epidermal growth factor receptor
0.0006904557236067014
GO:0007166: cell surface receptor signaling pathway	EVL	Enah/Vasp-like	0.0019458134711758197
GO:0007166: cell surface receptor signaling pathway	FAS	Fas cell surface death receptor -3.35
1252345089456e-5
GO:0007166: cell surface receptor signaling pathway	IFNA2	interferon, alpha 2	-0.0017255790
80846506
GO:0007166: cell surface receptor signaling pathway	IFNG	interferon, gamma	-4.8943590677
42938e-5
GO:0007166: cell surface receptor signaling pathway	IL27RA	interleukin 27 receptor, alpha	-0.00
048582361139973805
GO:0007166: cell surface receptor signaling pathway	INHA	inhibin, alpha	0.0002117948921175704
5
GO:0007166: cell surface receptor signaling pathway	INHBA	inhibin, beta A -0.001362513948571460
4
GO:0007166: cell surface receptor signaling pathway	JMJD6	jumonji domain containing 6	0.003
6071729847609794
GO:0007166: cell surface receptor signaling pathway	PRLR	prolactin receptor	0.00219347152
55802865
GO:0007166: cell surface receptor signaling pathway	TACSTD2 tumor-associated calcium signal trans
ducer 2 -0.0022709673817425417
GO:0007166: cell surface receptor signaling pathway	TSPAN6	tetraspanin 6	0.0013747101455425753
GO:0007259: JAK-STAT cascade	AGER	advanced glycosylation end product-specific receptor	-0.00
0172341741706043
GO:0007259: JAK-STAT cascade	CCL2	chemokine (C-C motif) ligand 2	0.0007961782341408855
GO:0007259: JAK-STAT cascade	FGFR3	fibroblast growth factor receptor 3	0.0002148664894982138
GO:0007259: JAK-STAT cascade	JAK2	Janus kinase 2	-3.8072571645887247e-5
GO:0007259: JAK-STAT cascade	STAMBP	STAM binding protein	-0.00016823482695193614
GO:0007259: JAK-STAT cascade	STAT5A	signal transducer and activator of transcription 5A	0.001
5624547873845593
GO:0007420: brain development	AGER	advanced glycosylation end product-specific receptor	-0.00
017446850428290962
GO:0007420: brain development	APOD	apolipoprotein D	0.0026341666368316263
GO:0007420: brain development	BAK1	BCL2-antagonist/killer 1	-0.0018801523234930593
GO:0007420: brain development	BBS7	Bardet-Biedl syndrome 7 -0.0009791393949996829
GO:0007420: brain development	BCL2L11 BCL2-like 11 (apoptosis facilitator)	-0.000776140215065747
5
GO:0007420: brain development	BRCA2	breast cancer 2, early onset	-1.0011110254309197e-5
GO:0007420: brain development	CAST	calpastatin	-0.003040365683747042
GO:0007420: brain development	CD9	CD9 molecule	-0.0025917006430470958
GO:0007420: brain development	CDK5RAP3	CDK5 regulatory subunit associated protein 3	0.001
1403608568741012
GO:0007420: brain development	CITED1	Cbp/p300-interacting transactivator, with Glu/Asp-rich carbox
y-terminal domain, 1	0.0028247663778074544
GO:0007420: brain development	CST3	cystatin C	-7.082053828187326e-5
GO:0007420: brain development	EGR2	early growth response 2 0.001442978107336196
GO:0007420: brain development	FOXC1	forkhead box C1 -2.1904968841335597e-5
GO:0007420: brain development	GRHL2	grainyhead-like 2 (Drosophila)	0.00100864526181659
GO:0007420: brain development	IGF1R	insulin-like growth factor 1 receptor	0.0010737533514887517
GO:0007420: brain development	MAP1S	microtubule-associated protein 1S	0.0005311389918535143
GO:0007420: brain development	MAPT	microtubule-associated protein tau	0.0015541305624082935
GO:0007420: brain development	MED1	mediator complex subunit 1	0.001132247148194274
GO:0007420: brain development	NES	nestin	0.0013775043968719625
GO:0007420: brain development	PHGDH	phosphoglycerate dehydrogenase	0.00029394755769203743
GO:0007420: brain development	PRKDC	protein kinase, DNA-activated, catalytic polypeptide	-0.00
1859400602884387
GO:0007420: brain development	PROX1	prospero homeobox 1	0.0011242561510562117
GO:0007420: brain development	PTCH1	patched 1	-6.575980054019736e-5
GO:0007420: brain development	RELN	reelin	0.0015412381043283837
GO:0007420: brain development	SHROOM2 shroom family member 2	0.0011148750556103646
GO:0007420: brain development	SIX3	SIX homeobox 3	0.002042279524246117

GO:0007420: brain development   SPHK1   sphingosine kinase 1    0.0018148802383100918
GO:0007420: brain development   STAR    steroidogenic acute regulatory protein  0.0008967393806491162
GO:0007420: brain development   STMN1   stathmin 1      0.0005610502424353875
GO:0007420: brain development   THRA    thyroid hormone receptor, alpha 0.0007648297005168201
GO:0007420: brain development   TULP3   tubby like protein 3    0.0009584191021333232
GO:0007420: brain development   ZNF335  zinc finger protein 335 -0.000365528325018728
GO:0009100: glycoprotein metabolic process      AGER    advanced glycosylation end product-specific r
eceptor -0.00014834649160886423
GO:0009750: response to fructose        AGER    advanced glycosylation end product-specific receptor
-0.00014834649160886423
GO:0010508: positive regulation of autophagy    AGER    advanced glycosylation end product-specific r
eceptor -0.00017480707487236733
GO:0010508: positive regulation of autophagy    BNIP3   BCL2/adenovirus E1B 19kDa interacting protein
3       0.0029297983611412813
GO:0010508: positive regulation of autophagy    FOXO1   forkhead box O1 0.001810035042835207
GO:0010508: positive regulation of autophagy    XBP1    X-box binding protein 1 0.0002673288898503276
5
GO:0010718: positive regulation of epithelial to mesenchymal transition AGER    advanced glycosylatio
n end product-specific receptor -0.00017412770780643645
GO:0010718: positive regulation of epithelial to mesenchymal transition ALX1    ALX homeobox 1  0.002
289675380527129
GO:0010718: positive regulation of epithelial to mesenchymal transition COL1A1  collagen, type I, alp
ha 1    -0.0005260914185390755
GO:0010718: positive regulation of epithelial to mesenchymal transition CTNNB1  catenin (cadherin-ass
ociated protein), beta 1, 88kDa -0.00011654993116169162
GO:0010718: positive regulation of epithelial to mesenchymal transition EZH2    enhancer of zeste 2 p
olycomb repressive complex 2 subunit    -0.00013567220937800198
GO:0010718: positive regulation of epithelial to mesenchymal transition GCNT2   glucosaminyl (N-acety
l) transferase 2, I-branching enzyme (I blood group)    0.0024294293566783534
GO:0010718: positive regulation of epithelial to mesenchymal transition HDAC2   histone deacetylase 2
-0.0012052943575696275
GO:0010718: positive regulation of epithelial to mesenchymal transition LEF1    lymphoid enhancer-bin
ding factor 1   -0.00010043258594003536
GO:0010718: positive regulation of epithelial to mesenchymal transition NOTCH1  notch 1 0.00051854533
71369046
GO:0010718: positive regulation of epithelial to mesenchymal transition TGFB1   transforming growth f
actor, beta 1   -7.313794013420126e-5
GO:0010718: positive regulation of epithelial to mesenchymal transition TGFB2   transforming growth f
actor, beta 2   -0.0010581970171750204
GO:0010718: positive regulation of epithelial to mesenchymal transition TGFB3   transforming growth f
actor, beta 3   -0.00182229364780218
GO:0010718: positive regulation of epithelial to mesenchymal transition TWIST1  twist family bHLH tra
nscription factor 1     -0.001340089402570323
GO:0010718: positive regulation of epithelial to mesenchymal transition WWTR1   WW domain containing
transcription regulator 1      0.0009000166833594054
GO:0010763: positive regulation of fibroblast migration AGER    advanced glycosylation end product-sp
ecific receptor -0.00017177467981926528
GO:0010763: positive regulation of fibroblast migration AKT1    v-akt murine thymoma viral oncogene h
omolog 1        0.0007084604504139824
GO:0010763: positive regulation of fibroblast migration TGFB1   transforming growth factor, beta 1
-6.900388222676314e-5
GO:0010763: positive regulation of fibroblast migration THBS1   thrombospondin 1        -0.0010069743
726467083
GO:0014823: response to activity        AGER    advanced glycosylation end product-specific receptor
-0.00017291152542450695
GO:0014823: response to activity        CCL2    chemokine (C-C motif) ligand 2  0.0008024190714658063
GO:0014823: response to activity        MAS1    MAS1 proto-oncogene, G protein-coupled receptor 0.000
29584200617529925
GO:0014823: response to activity        PRKDC   protein kinase, DNA-activated, catalytic polypeptide
-0.0018321034931001404
GO:0014823: response to activity        PTN     pleiotrophin    0.00029075271927588557
GO:0014823: response to activity        STAR    steroidogenic acute regulatory protein  0.00087860704
66180652
GO:0014823: response to activity        TH      tyrosine hydroxylase    -0.000339160978269231
GO:0014911: positive regulation of smooth muscle cell migration AGER    advanced glycosylation end pr
oduct-specific receptor -0.00016892625099548168
GO:0014911: positive regulation of smooth muscle cell migration BCL2    B-cell CLL/lymphoma 2   -6.25
35302081431585e-6
GO:0014911: positive regulation of smooth muscle cell migration IGF1    insulin-like growth factor 1
(somatomedin C)        0.00011656591892340843
GO:0014911: positive regulation of smooth muscle cell migration NRP1    neuropilin 1    -0.0006070586
606722715
GO:0014911: positive regulation of smooth muscle cell migration PDGFRB  platelet-derived growth facto
r receptor, beta polypeptide    -0.0003310873939760621
GO:0030324: lung development    AGER    advanced glycosylation end product-specific receptor    -0.00
017443942649393208
GO:0030324: lung development    ALDH1A2 aldehyde dehydrogenase 1 family, member A2      -0.0030238104
6331798
GO:0030324: lung development    DICER1  dicer 1, ribonuclease type III  -1.598508287682775e-5
GO:0030324: lung development    FGFR2   fibroblast growth factor receptor 2     0.0007644304721028299
GO:0030324: lung development    GLI1    GLI family zinc finger 1        -0.00129858945259542

GO:0030324: lung development    GLI2    GLI family zinc finger 2        0.0018565882658731472
GO:0030324: lung development    GLI3    GLI family zinc finger 3        -0.0021548768134108704
GO:0030324: lung development    HES1    hes family bHLH transcription factor 1  -0.000912387133289415
4
GO:0030324: lung development    ITGA3   integrin, alpha 3 (antigen CD49C, alpha 3 subunit of VLA-3 re
ceptor) 0.0014022278178806892
GO:0030324: lung development    JMJD6   jumonji domain containing 6     0.003596253880307907
GO:0030324: lung development    LOX     lysyl oxidase   -0.0005774279246892741
GO:0030324: lung development    NOTCH1  notch 1 0.0005194326424966049
GO:0030324: lung development    PROX1   prospero homeobox 1     0.0011233759559207209
GO:0030324: lung development    PTN     pleiotrophin    0.00030253182270621607
GO:0030324: lung development    SHH     sonic hedgehog  0.000601079743806589
GO:0030324: lung development    STRA6   stimulated by retinoic acid 6   -0.0016831186198885432
GO:0030324: lung development    TMBIM6  transmembrane BAX inhibitor motif containing 6  -0.0001476083
6380781412
GO:0030324: lung development    VEGFA   vascular endothelial growth factor A    0.000595900729728088
GO:0030324: lung development    WNT5A   wingless-type MMTV integration site family, member 5A   -0.00
06684151812011536
GO:0031175: neuron projection development       AGER    advanced glycosylation end product-specific r
eceptor -0.00017444118644246637
GO:0031175: neuron projection development       AREG    amphiregulin    0.00246011882975356 4
GO:0031175: neuron projection development       GDNF    glial cell derived neurotrophic factor  0.000
4454944683215134
GO:0031175: neuron projection development       HMGB1   high mobility group box 1       -0.0007779385
934992563
GO:0031175: neuron projection development       LYN     LYN proto-oncogene, Src family tyrosine kinas
e       -0.0014223060883479441
GO:0031175: neuron projection development       NEDD4   neural precursor cell expressed, developmenta
lly down-regulated 4, E3 ubiquitin protein ligase       0.0022661973448634093
GO:0031175: neuron projection development       PHGDH   phosphoglycerate dehydrogenase  0.00029135468
48188862
GO:0031175: neuron projection development       RB1     retinoblastoma 1        -0.001495524972351423
8
GO:0031175: neuron projection development       STMN1   stathmin 1      0.0005586829782879243
GO:0031175: neuron projection development       STMN2   stathmin 2      -0.0019494785260562404
GO:0031175: neuron projection development       STMN3   stathmin-like 3 0.0025934341869785107
GO:0031175: neuron projection development       STMN4   stathmin-like 4 0.00034856226811197686
GO:0032966: negative regulation of collagen biosynthetic process        AGER    advanced glycosylatio
n end product-specific receptor -0.00014834649160886423
GO:0033189: response to vitamin A       AGER    advanced glycosylation end product-specific receptor
-0.0001748276179846603
GO:0033189: response to vitamin A       ALDH1A2 aldehyde dehydrogenase 1 family, member A2      -0.00
3037151243199917
GO:0033189: response to vitamin A       GATA4   GATA binding protein 4  -0.001101936753420912
GO:0033189: response to vitamin A       PITX2   paired-like homeodomain 2       0.0021901196203252 26
GO:0033189: response to vitamin A       RXRA    retinoid X receptor, alpha      0.0011218503108525503
GO:0033189: response to vitamin A       TYMS    thymidylate synthetase  0.001573841341160684
GO:0033595: response to genistein       AGER    advanced glycosylation end product-specific receptor
-0.00014834649160886423
GO:0033689: negative regulation of osteoblast proliferation     AGER    advanced glycosylation end pr
oduct-specific receptor -0.00017376064785373045
GO:0033689: negative regulation of osteoblast proliferation     BCL2    B-cell CLL/lymphoma 2   -5.53
1878170583768e-6
GO:0033689: negative regulation of osteoblast proliferation     GREM1   gremlin 1, DAN family BMP ant
agonist -0.0008254190077890396
GO:0033689: negative regulation of osteoblast proliferation     NELL1   NEL-like 1 (chicken)    0.001
5282466465045426
GO:0033689: negative regulation of osteoblast proliferation     SFRP1   secreted frizzled-related pro
tein 1  0.0012792945079847498
GO:0035690: cellular response to drug   AGER    advanced glycosylation end product-specific receptor
-0.00017404904762181105
GO:0035690: cellular response to drug   CCL2    chemokine (C-C motif) ligand 2  0.0008138434299684035
GO:0035690: cellular response to drug   EGR1    early growth response 1 0.001090024476089523
GO:0035690: cellular response to drug   KCNE2   potassium voltage-gated channel, Isk-related family,
 member 2       -0.0005740932946616277
GO:0035690: cellular response to drug   MEF2C   myocyte enhancer factor 2C      0.0009698909105021106
GO:0035690: cellular response to drug   MYC     v-myc avian myelocytomatosis viral oncogene homolog
-0.0011344313889561496
GO:0035690: cellular response to drug   QDPR    quinoid dihydropteridine reductase      -0.0028475046
788878403
GO:0035690: cellular response to drug   TFRC    transferrin receptor    0.0010817403016909531
GO:0035690: cellular response to drug   TH      tyrosine hydroxylase    -0.00034158854759942233
GO:0035690: cellular response to drug   TP53    tumor protein p53       0.0011707613367156178
GO:0043507: positive regulation of JUN kinase activity  AGER    advanced glycosylation end product-sp
ecific receptor -0.00017483046779262425
GO:0043507: positive regulation of JUN kinase activity  MAP3K5  mitogen-activated protein kinase kina
se kinase 5     0.00034513897915360785
GO:0043507: positive regulation of JUN kinase activity  PAK1    p21 protein (Cdc42/Rac)-activated kin
ase 1   -0.0022036882015784493
GO:0043507: positive regulation of JUN kinase activity  TNFRSF11A       tumor necrosis factor recepto
r superfamily, member 11a, NFKB activator       0.002795834307888436 6

GO:0043525: positive regulation of neuron apoptotic process     AGER    advanced glycosylation end pr
oduct-specific receptor -0.00017316224415798748
GO:0043525: positive regulation of neuron apoptotic process     ASCL1   achaete-scute family bHLH tra
nscription factor 1     -0.0014125259486000288
GO:0043525: positive regulation of neuron apoptotic process     BAX     BCL2-associated X protein
-0.000422477019074834
GO:0043525: positive regulation of neuron apoptotic process     BCL2L11 BCL2-like 11 (apoptosis facil
itator) -0.0007689878285913832
GO:0043525: positive regulation of neuron apoptotic process     CDC34   cell division cycle 34  -0.00
01600339100319368
GO:0043525: positive regulation of neuron apoptotic process     CDC42   cell division cycle 42  0.001
2258436174132365
GO:0043525: positive regulation of neuron apoptotic process     EGR1    early growth response 1 0.001
0810191041238104
GO:0043525: positive regulation of neuron apoptotic process     FGFR3   fibroblast growth factor rece
ptor 3  0.00021964963809514151
GO:0043525: positive regulation of neuron apoptotic process     FOXO3   forkhead box O3 0.00118482002
39213138
GO:0043525: positive regulation of neuron apoptotic process     GRIK5   glutamate receptor, ionotropi
c, kainate 5    -0.00023169200191268326
GO:0043525: positive regulation of neuron apoptotic process     PAK3    p21 protein (Cdc42/Rac)-activ
ated kinase 3   -0.0012475001292670774
GO:0043525: positive regulation of neuron apoptotic process     TFAP2A  transcription factor AP-2 alp
ha (activating enhancer binding protein 2 alpha)        0.00047631841377539143
GO:0043525: positive regulation of neuron apoptotic process     TGFB2   transforming growth factor, b
eta 2   -0.0010453288708284097
GO:0043525: positive regulation of neuron apoptotic process     TP53    tumor protein p53       0.001
1571306736293347
GO:0048146: positive regulation of fibroblast proliferation     AGER    advanced glycosylation end pr
oduct-specific receptor -0.0001737589078074863
GO:0048146: positive regulation of fibroblast proliferation     AGT     angiotensinogen (serpin pepti
dase inhibitor, clade A, member 8)      -0.0011236518161791577
GO:0048146: positive regulation of fibroblast proliferation     CCNB1   cyclin B1       -0.0008795881
028420067
GO:0048146: positive regulation of fibroblast proliferation     E2F1    E2F transcription factor 1
0.0020851488695950425
GO:0048146: positive regulation of fibroblast proliferation     EGFR    epidermal growth factor recep
tor     0.0006800807786013382
GO:0048146: positive regulation of fibroblast proliferation     ESR1    estrogen receptor 1     -0.00
09415233693240736
GO:0048146: positive regulation of fibroblast proliferation     IGF1    insulin-like growth factor 1
 (somatomedin C)        0.00012893953284377692
GO:0048146: positive regulation of fibroblast proliferation     MYC     v-myc avian myelocytomatosis
 viral oncogene homolog -0.0011312139237075823
GO:0048146: positive regulation of fibroblast proliferation     PML     promyelocytic leukemia  -0.00
06776744080157948
GO:0048146: positive regulation of fibroblast proliferation     PRKDC   protein kinase, DNA-activate
d, catalytic polypeptide        -0.0018460968198891934
GO:0048146: positive regulation of fibroblast proliferation     RNASEH2B        ribonuclease H2, subu
nit B   -4.2662503416772365e-5
GO:0048146: positive regulation of fibroblast proliferation     SPHK1   sphingosine kinase 1    0.001
8002259895043599
GO:0048146: positive regulation of fibroblast proliferation     TGFB1   transforming growth factor, b
eta 1   -7.214004938491884e-5
GO:0048146: positive regulation of fibroblast proliferation     WNT1    wingless-type MMTV integratio
n site family, member 1 0.0007827611742660041
GO:0048146: positive regulation of fibroblast proliferation     WNT5A   wingless-type MMTV integratio
n site family, member 5A        -0.0006631535226664452
GO:0048661: positive regulation of smooth muscle cell proliferation     AGER    advanced glycosylatio
n end product-specific receptor -0.0001736169086333462
GO:0048661: positive regulation of smooth muscle cell proliferation     ALOX12  arachidonate 12-lipox
ygenase -0.0019405398748443717
GO:0048661: positive regulation of smooth muscle cell proliferation     BMP4    bone morphogenetic pr
otein 4 -0.00032047514972448023
GO:0048661: positive regulation of smooth muscle cell proliferation     CX3CL1  chemokine (C-X3-C mot
if) ligand 1    0.002100798079705573
GO:0048661: positive regulation of smooth muscle cell proliferation     FGFR2   fibroblast growth fac
tor receptor 2  0.00075776608380576
GO:0048661: positive regulation of smooth muscle cell proliferation     HMOX1   heme oxygenase (decyc
ling) 1 -0.00021587041829301994
GO:0048661: positive regulation of smooth muscle cell proliferation     ID2     inhibitor of DNA bind
ing 2, dominant negative helix-loop-helix protein       6.0341930686014384e-5
GO:0048661: positive regulation of smooth muscle cell proliferation     IGF1    insulin-like growth f
actor 1 (somatomedin C) 0.0001288956100438221
GO:0048661: positive regulation of smooth muscle cell proliferation     IRAK1   interleukin-1 recepto
r-associated kinase 1   -0.0015978646104060348
GO:0048661: positive regulation of smooth muscle cell proliferation     ORC1    origin recognition co
mplex, subunit 1        -0.0007351319920685379
GO:0048661: positive regulation of smooth muscle cell proliferation     PDGFRB  platelet-derived grow
th factor receptor, beta polypeptide    -0.0003567015459709957
GO:0048661: positive regulation of smooth muscle cell proliferation     SERPINF2        serpin peptid

ase inhibitor, clade F (alpha-2 antiplasmin, pigment epithelium derived factor), member 2 0.000
6549047756364494
GO:0048661: positive regulation of smooth muscle cell proliferation SKP2 S-phase kinase-associ
ated protein 2, E3 ubiquitin protein ligase -0.0005360494011624169
GO:0050729: positive regulation of inflammatory response AGER advanced glycosylation end pr
oduct-specific receptor -0.00017368653090787866
GO:0050729: positive regulation of inflammatory response AGT angiotensinogen (serpin pepti
dase inhibitor, clade A, member 8) -0.0011252073986677223
GO:0050729: positive regulation of inflammatory response AGTR1 angiotensin II receptor, type
1 0.00034747408201561285
GO:0050729: positive regulation of inflammatory response CX3CL1 chemokine (C-X3-C motif) liga
nd 1 0.0021041179405726623
GO:0050729: positive regulation of inflammatory response IL12B interleukin 12B 0.00127146959
73251732
GO:0050729: positive regulation of inflammatory response JAK2 Janus kinase 2 -3.2577995143
77627e-5
GO:0050729: positive regulation of inflammatory response PRKCA protein kinase C, alpha -5.92
5663619042989e-6
GO:0050729: positive regulation of inflammatory response SERPINE1 serpin peptidase inhi
bitor, clade E (nexin, plasminogen activator inhibitor type 1), member 1 0.0001126160737699245
6
GO:0050729: positive regulation of inflammatory response STAT5A signal transducer and activat
or of transcription 5A 0.001588238638304995
GO:0050729: positive regulation of inflammatory response WNT5A wingless-type MMTV integratio
n site family, member 5A -0.0006636463894865769
GO:0050930: induction of positive chemotaxis AGER advanced glycosylation end product-specific r
eceptor -0.0001744705746370809
GO:0050930: induction of positive chemotaxis AZU1 azurocidin 1 0.000198359811675336
GO:0050930: induction of positive chemotaxis CXCL12 chemokine (C-X-C motif) ligand 12 -0.00
1210938400321844
GO:0050930: induction of positive chemotaxis PRKCA protein kinase C, alpha -5.875526875742755e-6
GO:0050930: induction of positive chemotaxis VEGFA vascular endothelial growth factor A 0.000
5965945992619952
GO:0050930: induction of positive chemotaxis VEGFC vascular endothelial growth factor C -0.00
3370773012919577
GO:0051092: positive regulation of NF-kappaB transcription factor activity AGER advanced glyc
osylation end product-specific receptor -0.00017432335669368253
GO:0051092: positive regulation of NF-kappaB transcription factor activity AGT angiotensinog
en (serpin peptidase inhibitor, clade A, member 8) -0.0011337690988374245
GO:0051092: positive regulation of NF-kappaB transcription factor activity AR androgen rece
ptor 0.0026406868806194946
GO:0051092: positive regulation of NF-kappaB transcription factor activity CIB1 calcium and i
ntegrin binding 1 (calmyrin) 5.504244238658197e-5
GO:0051092: positive regulation of NF-kappaB transcription factor activity CLOCK clock circadi
an regulator 0.00020029395010825264
GO:0051092: positive regulation of NF-kappaB transcription factor activity CTH cystathionine
gamma-lyase 0.0017494893888893995
GO:0051092: positive regulation of NF-kappaB transcription factor activity EDA ectodysplasin
A -0.0008150419386035191
GO:0051092: positive regulation of NF-kappaB transcription factor activity GREM1 gremlin 1, DA
N family BMP antagonist -0.0008278792419378839
GO:0051092: positive regulation of NF-kappaB transcription factor activity ICAM1 intercellular
adhesion molecule 1 0.0007295545166121159
GO:0051092: positive regulation of NF-kappaB transcription factor activity IRAK1 interleukin-1
receptor-associated kinase 1 -0.001610292102245563
GO:0051092: positive regulation of NF-kappaB transcription factor activity LGALS9 lectin, galac
toside-binding, soluble, 9 -0.00011687984728779948
GO:0051092: positive regulation of NF-kappaB transcription factor activity PRKCZ protein kinas
e C, zeta -0.0016019594428588928
GO:0051092: positive regulation of NF-kappaB transcription factor activity SPHK1 sphingosine k
inase 1 0.001811821086001211
GO:0051092: positive regulation of NF-kappaB transcription factor activity TGFB1 transforming
growth factor, beta 1 -7.28818014547426e-5
GO:0051092: positive regulation of NF-kappaB transcription factor activity TNFRSF11A tumor
necrosis factor receptor superfamily, member 11a, NFKB activator 0.002777704844494479
GO:0051092: positive regulation of NF-kappaB transcription factor activity WNT5A wingless-type
MMTV integration site family, member 5A -0.00066770787962612
GO:0051101: regulation of DNA binding AGER advanced glycosylation end product-specific receptor
-0.00017282634813972492
GO:0051101: regulation of DNA binding CSNK2B casein kinase 2, beta polypeptide 0.00150495023
58291905
GO:0051101: regulation of DNA binding HJURP Holliday junction recognition protein -7.6928380050
05383e-5
GO:0051101: regulation of DNA binding TGFB1 transforming growth factor, beta 1 -7.1443911215
37731e-5
GO:0051595: response to methylglyoxal AGER advanced glycosylation end product-specific receptor
-0.00014834649160886423
GO:0055074: calcium ion homeostasis AGER advanced glycosylation end product-specific receptor
-0.00017381943443857043
GO:0055074: calcium ion homeostasis CAV1 caveolin 1, caveolae protein, 22kDa -0.0005338583
631218384

GO:0055074: calcium ion homeostasis     CYP27B1 cytochrome P450, family 27, subfamily B, polypeptide 1      -0.00046865746350316677
GO:0055074: calcium ion homeostasis     S100A14 S100 calcium binding protein A14        -0.0021505132543200908
GO:0055074: calcium ion homeostasis     WFS1    Wolfram syndrome 1 (wolframin)  0.0005753734761084317
GO:0055093: response to hyperoxia       AGER    advanced glycosylation end product-specific receptor -0.0001744937618179712
GO:0055093: response to hyperoxia       BNIP3   BCL2/adenovirus E1B 19kDa interacting protein 3 0.0029210754344249153
GO:0055093: response to hyperoxia       COL1A1  collagen, type I, alpha 1       -0.0005294384627999856
GO:0055093: response to hyperoxia       HDAC2   histone deacetylase 2   -0.0012086430895868053
GO:0055093: response to hyperoxia       PDGFRB  platelet-derived growth factor receptor, beta polypeptide    -0.0003637642548400947
GO:0060100: positive regulation of phagocytosis, engulfment     AGER    advanced glycosylation end product-specific receptor -0.00017437269042957555
GO:0060100: positive regulation of phagocytosis, engulfment     GATA2   GATA binding protein 2  -0.00045139068839606154
GO:0060100: positive regulation of phagocytosis, engulfment     STAP1   signal transducing adaptor family member 1   0.00244564877336297
GO:0060290: transdifferentiation        AGER    advanced glycosylation end product-specific receptor -0.0001718688689624252
GO:0060290: transdifferentiation        GATA4   GATA binding protein 4  -0.0010650683605545053
GO:0060290: transdifferentiation        PDX1    pancreatic and duodenal homeobox 1      0.00024676608728110686
GO:0070301: cellular response to hydrogen peroxide      AGER    advanced glycosylation end product-specific receptor -0.00017398449921313607
GO:0070301: cellular response to hydrogen peroxide      BNIP3   BCL2/adenovirus E1B 19kDa interacting protein 3       0.00290176859723207
GO:0070301: cellular response to hydrogen peroxide      CST3    cystatin C      -6.895543295744787e-5
GO:0070301: cellular response to hydrogen peroxide      CYP1B1  cytochrome P450, family 1, subfamily B, polypeptide 1       0.0003967770016374729
GO:0070301: cellular response to hydrogen peroxide      ECT2    epithelial cell transforming 2  0.0010582300901999433
GO:0070301: cellular response to hydrogen peroxide      EZH2    enhancer of zeste 2 polycomb repressive complex 2 subunit    -0.00013681638465076702
GO:0070301: cellular response to hydrogen peroxide      HDAC2   histone deacetylase 2   -0.0012039100219929334
GO:0070301: cellular response to hydrogen peroxide      KLF2    Kruppel-like factor 2   -0.001228736506686079
GO:0070301: cellular response to hydrogen peroxide      MAP3K5  mitogen-activated protein kinase kinase kinase 5     0.0003410789784105755
GO:0070301: cellular response to hydrogen peroxide      PAX2    paired box 2    -0.0016077238925806033
GO:0070301: cellular response to hydrogen peroxide      SIRT1   sirtuin 1       -2.353779916703928e-6
GO:0070301: cellular response to hydrogen peroxide      TNFAIP3 tumor necrosis factor, alpha-induced protein 3      0.0011126139106054412
GO:0071333: cellular response to glucose stimulus       AGER    advanced glycosylation end product-specific receptor -0.0001727993677298939
GO:0071333: cellular response to glucose stimulus       GATA4   GATA binding protein 4  -0.0010778050662791766
GO:0071333: cellular response to glucose stimulus       GRIK5   glutamate receptor, ionotropic, kainate 5    -0.00023142460422644488
GO:0071333: cellular response to glucose stimulus       ICAM1   intercellular adhesion molecule 1 0.0007224109980621391
GO:0071333: cellular response to glucose stimulus       PAX2    paired box 2    -0.0015880765090634655
GO:0071333: cellular response to glucose stimulus       PDK3    pyruvate dehydrogenase kinase, isozyme 3     0.0007545067307062337
GO:0071333: cellular response to glucose stimulus       SOX4    SRY (sex determining region Y)-box 4 -4.0989990021899445e-5
GO:0071333: cellular response to glucose stimulus       STAR    steroidogenic acute regulatory protein       0.0008794869527143406
GO:0071333: cellular response to glucose stimulus       TH      tyrosine hydroxylase    -0.0003388087754680805
GO:0071333: cellular response to glucose stimulus       XBP1    X-box binding protein 1 0.00026540600671913316
GO:0071398: cellular response to fatty acid     AGER    advanced glycosylation end product-specific receptor -0.00017403648752573972
GO:0071398: cellular response to fatty acid     CCNB1   cyclin B1       -0.0008811124135056452
GO:0071398: cellular response to fatty acid     CPT1A   carnitine palmitoyltransferase 1A (liver) 0.0012703301162785693
GO:0071398: cellular response to fatty acid     E2F1    E2F transcription factor 1      0.0020919373161603747
GO:0071398: cellular response to fatty acid     PDK3    pyruvate dehydrogenase kinase, isozyme 3 0.0007692072096432324
GO:0071407: cellular response to organic cyclic compound        AGER    advanced glycosylation end product-specific receptor -0.00017355911864847437
GO:0071407: cellular response to organic cyclic compound        AKT1    v-akt murine thymoma viral oncogene homolog 1        0.0007233967173948215
GO:0071407: cellular response to organic cyclic compound        AKT2    v-akt murine thymoma viral on

cogene homolog 2	-0.0006222397836267628
GO:0071407: cellular response to organic cyclic compound	AXIN1	axin 1	-0.000724465466645846
5
GO:0071407: cellular response to organic cyclic compound	CCL2	chemokine (C-C motif) ligand
 2	0.0008079755974389152
GO:0071407: cellular response to organic cyclic compound	CCNB1	cyclin B1	-0.0008807212
614018306
GO:0071407: cellular response to organic cyclic compound	CYP1B1	cytochrome P450, family 1, su
bfamily B, polypeptide 1	0.0003948647056333418
GO:0071407: cellular response to organic cyclic compound	GLI2	GLI family zinc finger 2
0.0018375816077707456
GO:0071407: cellular response to organic cyclic compound	MGMT	O-6-methylguanine-DNA methylt
ransferase	0.0004081079628331253
GO:0071407: cellular response to organic cyclic compound	PAK1	p21 protein (Cdc42/Rac)-activ
ated kinase 1	-0.0021684321580586093
GO:0071407: cellular response to organic cyclic compound	PAK3	p21 protein (Cdc42/Rac)-activ
ated kinase 3	-0.0012506631005173507
GO:0071407: cellular response to organic cyclic compound	TGFB1	transforming growth factor, b
eta 1	-7.087309899284887e-5
GO:0072657: protein localization to membrane	AGER	advanced glycosylation end product-specific r
eceptor -0.00017484498155257284
GO:0072657: protein localization to membrane	CPE	carboxypeptidase E	-0.002559902518296101
GO:0072714: response to selenite ion	AGER	advanced glycosylation end product-specific receptor
-0.00014834649160886423
GO:1901018: positive regulation of potassium ion transmembrane transporter activity	AGER	advan
ced glycosylation end product-specific receptor -0.00017284645634956528
GO:1901018: positive regulation of potassium ion transmembrane transporter activity	ATP1B3	ATPas
e, Na+/K+ transporting, beta 3 polypeptide	-0.0014828321350051446
GO:2000353: positive regulation of endothelial cell apoptotic process	AGER	advanced glycosylatio
n end product-specific receptor -0.0001710091813436243
GO:2000353: positive regulation of endothelial cell apoptotic process	BMP4	bone morphogenetic pr
otein 4 -0.0003112667840613703
GO:2000353: positive regulation of endothelial cell apoptotic process	THBS1	thrombospondin 1
-0.000998148501082753
GO:2000353: positive regulation of endothelial cell apoptotic process	XBP1	X-box binding protein
1	0.0002619672047020387
GO:2000379: positive regulation of reactive oxygen species metabolic process	AGER	advanced glyc
osylation end product-specific receptor -0.0001744571595647961
GO:2000379: positive regulation of reactive oxygen species metabolic process	AGT	angiotensinog
en (serpin peptidase inhibitor, clade A, member 8)	-0.0011398484349049046
GO:2000379: positive regulation of reactive oxygen species metabolic process	AGTR1	angiotensin I
I receptor, type 1	0.0003468076530572346
GO:2000379: positive regulation of reactive oxygen species metabolic process	GRB2	growth factor
receptor-bound protein 2	0.0004309935682787311
GO:2000379: positive regulation of reactive oxygen species metabolic process	LEP	leptin	0.003
2042520517062507
GO:2000379: positive regulation of reactive oxygen species metabolic process	PDGFRB	platelet-deri
ved growth factor receptor, beta polypeptide	-0.0003644626908305289
GO:2000379: positive regulation of reactive oxygen species metabolic process	THBS1	thrombospondi
n 1	-0.0010398131143783793
GO:2000379: positive regulation of reactive oxygen species metabolic process	TP53	tumor protein
p53	0.0011816822614750904
GO:2000676: positive regulation of type B pancreatic cell apoptotic process	AGER	advanced glyc
osylation end product-specific receptor -0.00014834649160886423
GO:0010628: positive regulation of gene expression	AGR2	anterior gradient 2	0.00163102585
40125494
GO:0010628: positive regulation of gene expression	ALDH1A2	aldehyde dehydrogenase 1 family, memb
er A2	-0.0030214035597184888
GO:0010628: positive regulation of gene expression	ALOX12	arachidonate 12-lipoxygenase	-0.00
19549918846050434
GO:0010628: positive regulation of gene expression	ANK3	ankyrin 3, node of Ranvier (ankyrin
 G)	0.0007006326936388545
GO:0010628: positive regulation of gene expression	AR	androgen receptor	0.00264130702
7378681
GO:0010628: positive regulation of gene expression	AVP	arginine vasopressin	-0.0009377555
621954946
GO:0010628: positive regulation of gene expression	AZU1	azurocidin 1	0.0001982533945105531
5
GO:0010628: positive regulation of gene expression	CAV1	caveolin 1, caveolae protein, 22kDa
-0.000535831446425033
GO:0010628: positive regulation of gene expression	CDC42	cell division cycle 42	0.00124141809
09225776
GO:0010628: positive regulation of gene expression	CDH3	cadherin 3, type 1, P-cadherin (place
ntal)	-0.0012325089245543428
GO:0010628: positive regulation of gene expression	CITED1	Cbp/p300-interacting transactivator,
 with Glu/Asp-rich carboxy-terminal domain, 1	0.002820644403418464
GO:0010628: positive regulation of gene expression	DICER1	dicer 1, ribonuclease type III	-1.59
3903553030115e-5
GO:0010628: positive regulation of gene expression	E2F1	E2F transcription factor 1	0.002
0989260280933333
GO:0010628: positive regulation of gene expression	FGF8	fibroblast growth factor 8 (androgen-

induced)        0.0009845874591857982
GO:0010628: positive regulation of gene expression      GJA1    gap junction protein, alpha 1, 43kDa
-0.00016104021761663886
GO:0010628: positive regulation of gene expression      HINFP   histone H4 transcription factor 0.000
9927224392016236
GO:0010628: positive regulation of gene expression      HMGA2   high mobility group AT-hook 2   0.001
510370069653 8387
GO:0010628: positive regulation of gene expression      HPN     hepsin  0.0031614623473557807
GO:0010628: positive regulation of gene expression      ID2     inhibitor of DNA binding 2, dominant
negative helix-loop-helix protein      6.018731043526227e-5
GO:0010628: positive regulation of gene expression      IFNG    interferon, gamma       -4.9042182046
04564e-5
GO:0010628: positive regulation of gene expression      INHBA   inhibin, beta A -0.001353712520955763
7
GO:0010628: positive regulation of gene expression      ITGA3   integrin, alpha 3 (antigen CD49C, alp
ha 3 subunit of VLA-3 receptor) 0.0014014776966537075
GO:0010628: positive regulation of gene expression      LEF1    lymphoid enhancer-binding factor 1
-0.00010009965043653985
GO:0010628: positive regulation of gene expression      MED1    mediator complex subunit 1      0.001
12979237132437
GO:0010628: positive regulation of gene expression      MEF2C   myocyte enhancer factor 2C      0.000
974040835236969 7
GO:0010628: positive regulation of gene expression      PAX6    paired box 6    0.0019366682096019544
GO:0010628: positive regulation of gene expression      PIK3CD  phosphatidylinositol-4,5-bisphosphate
3-kinase, catalytic subunit delta       -0.0007814128934088678
GO:0010628: positive regulation of gene expression      SMO     smoothened, frizzled class receptor
0.002140706813498219
GO:0010628: positive regulation of gene expression      SOX11   SRY (sex determining region Y)-box 11
-0.00021004944823715377
GO:0010628: positive regulation of gene expression      STAP1   signal transducing adaptor family mem
ber 1   0.002445488441460362
GO:0010628: positive regulation of gene expression      STAR    steroidogenic acute regulatory protei
n       0.0008950555723609028
GO:0010628: positive regulation of gene expression      TFAP2A  transcription factor AP-2 alpha (acti
vating enhancer binding protein 2 alpha)        0.0004804717661061146
GO:0010628: positive regulation of gene expression      TGFB1   transforming growth factor, beta 1
-7.277166689019998e-5
GO:0010628: positive regulation of gene expression      TGFB2   transforming growth factor, beta 2
-0.001060160127169235
GO:0010628: positive regulation of gene expression      TGFBR1  transforming growth factor, beta rece
ptor 1  0.0003400085653103005
GO:0010628: positive regulation of gene expression      TNC     tenascin C      0.0007351709405452432
GO:0010628: positive regulation of gene expression      TP53    tumor protein p53       0.00117625292
07385666
GO:0010628: positive regulation of gene expression      TWIST1  twist family bHLH transcription facto
r 1     -0.0013424544095350136
GO:0010628: positive regulation of gene expression      VDR     vitamin D (1,25- dihydroxyvitamin D3)
receptor        0.0005641977550405359
GO:0010628: positive regulation of gene expression      VEGFA   vascular endothelial growth factor A
0.0005948186584939899
GO:0010628: positive regulation of gene expression      ZPR1    ZPR1 zinc finger        -0.0002100850
5472527552
GO:0010811: positive regulation of cell-substrate adhesion      AGR2    anterior gradient 2     0.001
6160594067224788
GO:0010811: positive regulation of cell-substrate adhesion      EGFL6   EGF-like-domain, multiple 6
0.00013387785099819072
GO:0010811: positive regulation of cell-substrate adhesion      ITGA3   integrin, alpha 3 (antigen CD
49C, alpha 3 subunit of VLA-3 receptor) 0.001385132937098839
GO:0010811: positive regulation of cell-substrate adhesion      JAK2    Janus kinase 2  -3.5414436133
2172e-5
GO:0010811: positive regulation of cell-substrate adhesion      PTN     pleiotrophin    0.00029182850
013118413
GO:0010811: positive regulation of cell-substrate adhesion      THBS1   thrombospondin 1        -0.00
10220888368528101
GO:0045742: positive regulation of epidermal growth factor receptor signaling pathway   AGR2    anter
ior gradient 2  0.0016287095961735913
GO:0045742: positive regulation of epidermal growth factor receptor signaling pathway   AGT     angio
tensinogen (serpin peptidase inhibitor, clade A, member 8)      -0.0011385117083971214
GO:0048546: digestive tract morphogenesis       AGR2    anterior gradient 2     0.001620494430298279
GO:0048546: digestive tract morphogenesis       BBS7    Bardet-Biedl syndrome 7 -0.000971161426819931
5
GO:0048546: digestive tract morphogenesis       BCL2    B-cell CLL/lymphoma 2   -5.614563355390786e-6
GO:0048546: digestive tract morphogenesis       EGFR    epidermal growth factor receptor        0.000
6825754042497431
GO:0048546: digestive tract morphogenesis       EPHB3   EPH receptor B3 0.0008813007896211691
GO:0048546: digestive tract morphogenesis       FGFR3   fibroblast growth factor receptor 3     0.000
2207993340683321
GO:0048546: digestive tract morphogenesis       GLI1    GLI family zinc finger 1        -0.0012917663
020571228
GO:0048546: digestive tract morphogenesis       HIF1A   hypoxia inducible factor 1, alpha subunit (ba
sic helix-loop-helix transcription factor)      -0.0006434311090543012

GO:0048546: digestive tract morphogenesis       SFRP1   secreted frizzled-related protein 1     0.001
2754926175704388
GO:0048546: digestive tract morphogenesis       SOX10   SRY (sex determining region Y)-box 10   0.000
18939330145281634
GO:0048546: digestive tract morphogenesis       STRA6   stimulated by retinoic acid 6   -0.0016729989
147031131
GO:0048546: digestive tract morphogenesis       STX2    syntaxin 2      0.00030624891537237665
GO:0048546: digestive tract morphogenesis       TP73    tumor protein p73       0.0010229919354482917
GO:0048639: positive regulation of developmental growth AGR2    anterior gradient 2     0.00163791501
8435804
GO:0048639: positive regulation of developmental growth C3      complement component 3  0.00204172878
78284726
GO:0048639: positive regulation of developmental growth INSR    insulin receptor        -0.0013755111
282643966
GO:0048639: positive regulation of developmental growth LEP     leptin  0.0032210207512611143
GO:0048639: positive regulation of developmental growth PLCB1   phospholipase C, beta 1 (phosphoinosi
tide-specific)  0.00015698768375104608
GO:0048639: positive regulation of developmental growth PRKDC   protein kinase, DNA-activated, cataly
tic polypeptide -0.001869449519710636
GO:0060480: lung goblet cell differentiation    AGR2    anterior gradient 2     0.0016381287197342277
GO:0060480: lung goblet cell differentiation    HOXA5   homeobox A5     0.0010735335675550143
GO:0060480: lung goblet cell differentiation    SPDEF   SAM pointed domain containing ETS transcripti
on factor       0.0031559374562677048
GO:0060548: negative regulation of cell death   AGR2    anterior gradient 2     0.0015923902205138088
GO:0060548: negative regulation of cell death   BMP4    bone morphogenetic protein 4    -0.0003125633
803041753
GO:0060548: negative regulation of cell death   BMP7    bone morphogenetic protein 7    0.00083436548
48161876
GO:0060548: negative regulation of cell death   CST3    cystatin C      -5.692310563403836e-5
GO:0060548: negative regulation of cell death   MGMT    O-6-methylguanine-DNA methyltransferase 0.000
408629385464537
GO:0060548: negative regulation of cell death   NCK1    NCK adaptor protein 1   -0.000771942476618261
2
GO:0060548: negative regulation of cell death   SOX11   SRY (sex determining region Y)-box 11   -0.00
021052977470417664
GO:0060548: negative regulation of cell death   SOX4    SRY (sex determining region Y)-box 4    -4.63
5715176203974e-5
GO:0070254: mucus secretion     AGR2    anterior gradient 2     0.001616559967177163
GO:0070254: mucus secretion     VAMP8   vesicle-associated membrane protein 8   -5.739824683197586e-5
GO:0090004: positive regulation of establishment of protein localization to plasma membrane     AGR2
anterior gradient 2     0.0016314391645785946
GO:0090004: positive regulation of establishment of protein localization to plasma membrane     AKT1
v-akt murine thymoma viral oncogene homolog 1   0.0007330372891685905
GO:0090004: positive regulation of establishment of protein localization to plasma membrane     CIB1
calcium and integrin binding 1 (calmyrin)       5.3716006980066744e-5
GO:0090004: positive regulation of establishment of protein localization to plasma membrane     ITGA3
integrin, alpha 3 (antigen CD49C, alpha 3 subunit of VLA-3 receptor)    0.0014033841205646704
GO:0090004: positive regulation of establishment of protein localization to plasma membrane     ITGB1
integrin, beta 1 (fibronectin receptor, beta polypeptide, antigen CD29 includes MDF2, MSK12)    0.002
4347740464617847
GO:1903896: positive regulation of IRE1-mediated unfolded protein response      AGR2    anterior grad
ient 2  0.0016261012102319805
GO:1903896: positive regulation of IRE1-mediated unfolded protein response      BAK1    BCL2-antagoni
st/killer 1     -0.0018674303700608123
GO:1903896: positive regulation of IRE1-mediated unfolded protein response      BAX     BCL2-associat
ed X protein    -0.0004245322910266187
GO:1903896: positive regulation of IRE1-mediated unfolded protein response      BCL2L11 BCL2-like 11
(apoptosis facilitator)        -0.0007730341545850338
GO:1903896: positive regulation of IRE1-mediated unfolded protein response      PTPN1   protein tyros
ine phosphatase, non-receptor type 1    -0.0010358127098714785
GO:1903899: positive regulation of PERK-mediated unfolded protein response      AGR2    anterior grad
ient 2  0.001615409903912475
GO:1903899: positive regulation of PERK-mediated unfolded protein response      PTPN2   protein tyros
ine phosphatase, non-receptor type 2    -0.0003276632763198136
KEGG:04920: Adipocytokine signaling pathway     AGRP    agouti related protein homolog (mouse)  -0.00
25845829348576086
KEGG:04920: Adipocytokine signaling pathway     AKT1    v-akt murine thymoma viral oncogene homolog 1
0.000735593243065051
KEGG:04920: Adipocytokine signaling pathway     AKT2    v-akt murine thymoma viral oncogene homolog 2
-0.0006290229838738279
KEGG:04920: Adipocytokine signaling pathway     CPT1A   carnitine palmitoyltransferase 1A (liver)
0.0012785223405717785
KEGG:04920: Adipocytokine signaling pathway     JAK2    Janus kinase 2  -3.113227062993063e-5
KEGG:04920: Adipocytokine signaling pathway     LEP     leptin  0.0032118289187881366
KEGG:04920: Adipocytokine signaling pathway     RXRA    retinoid X receptor, alpha      0.00112171768
46465623
GO:0007218: neuropeptide signaling pathway      AGRP    agouti related protein homolog (mouse)  -0.00
25980799544318943
GO:0007218: neuropeptide signaling pathway      CPE     carboxypeptidase E      -0.002572735144835655
GO:0007218: neuropeptide signaling pathway      NMU     neuromedin U    0.0018733421957252024
GO:0007623: circadian rhythm    AGRP    agouti related protein homolog (mouse)  -0.002581485207481716

4
GO:0007623: circadian rhythm    ATF5    activating transcription factor 5       -0.002746917064340821
5
GO:0007623: circadian rhythm    AVP     arginine vasopressin    -0.0009420770898513261
GO:0007623: circadian rhythm    CLOCK   clock circadian regulator       0.00019997548570510737
GO:0007623: circadian rhythm    CPT1A   carnitine palmitoyltransferase 1A (liver)       0.00127703879
22196992
GO:0007623: circadian rhythm    CREB1   cAMP responsive element binding protein 1       0.00066239449
54809945
GO:0007623: circadian rhythm    DRD4    dopamine receptor D4    -0.0019527946500199067
GO:0007623: circadian rhythm    EGR1    early growth response 1 0.001097860031630588
GO:0007623: circadian rhythm    FAS     Fas cell surface death receptor -3.360411498469052e-5
GO:0007623: circadian rhythm    GSK3B   glycogen synthase kinase 3 beta 0.0015561253915316107
GO:0007623: circadian rhythm    LEP     leptin  0.003207028097573989
GO:0007623: circadian rhythm    NRIP1   nuclear receptor interacting protein 1  0.0010756727878886594
GO:0007623: circadian rhythm    PER2    period circadian clock 2        0.0012439650648792555
GO:0007623: circadian rhythm    PROX1   prospero homeobox 1     0.0011269489669413157
GO:0007623: circadian rhythm    SERPINE1        serpin peptidase inhibitor, clade E (nexin, plasminog
en activator inhibitor type 1), member 1        0.00011194746317624237
GO:0007623: circadian rhythm    TYMS    thymidylate synthetase  0.0015696074884617008
GO:0007631: feeding behavior    AGRP    agouti related protein homolog (mouse)  -0.002576312040367613
GO:0007631: feeding behavior    DRD2    dopamine receptor D2    -0.00023435807408985764
GO:0007631: feeding behavior    HTR2C   5-hydroxytryptamine (serotonin) receptor 2C, G protein-couple
d       -0.0005261961742939731
GO:0007631: feeding behavior    STRA6   stimulated by retinoic acid 6   -0.0016843392582535418
GO:0008343: adult feeding behavior      AGRP    agouti related protein homolog (mouse)  -0.0026068701
655804663
GO:0008343: adult feeding behavior      LEP     leptin  0.0032441430554579808
GO:0009648: photoperiodism      AGRP    agouti related protein homolog (mouse)  -0.002586827397754355
GO:0009648: photoperiodism      CLOCK   clock circadian regulator       0.0001995736604176251
GO:0009648: photoperiodism      DRD4    dopamine receptor D4    -0.0019562601533895537
GO:0009648: photoperiodism      NMU     neuromedin U    0.0018635970788099961
GO:0009755: hormone-mediated signaling pathway  AGRP    agouti related protein homolog (mouse)  -0.00
25752002813014094
GO:0009755: hormone-mediated signaling pathway  JAK2    Janus kinase 2  -3.209043029876431e-5
GO:0009755: hormone-mediated signaling pathway  THRA    thyroid hormone receptor, alpha 0.00076507448
11826133
GO:0042755: eating behavior     AGRP    agouti related protein homolog (mouse)  -0.002594858428084232
GO:0042755: eating behavior     CPT1A   carnitine palmitoyltransferase 1A (liver)       0.00128159287
13301437
GO:0042755: eating behavior     LEP     leptin  0.0032263055093409323
GO:0042755: eating behavior     NMU     neuromedin U    0.0018697811514943202
GO:0042755: eating behavior     TCF15   transcription factor 15 (basic helix-loop-helix)        -0.00
26763268812153925
GO:0042755: eating behavior     TH      tyrosine hydroxylase    -0.00034417826032922096
GO:0060135: maternal process involved in female pregnancy       AGRP    agouti related protein homolo
g (mouse)       -0.002571362204035075
GO:0060135: maternal process involved in female pregnancy       CCL2    chemokine (C-C motif) ligand
 2      0.0008167085215575097
GO:0060135: maternal process involved in female pregnancy       DSG2    desmoglein 2    1.66297648020
01453e-5
GO:0060135: maternal process involved in female pregnancy       IHH     indian hedgehog -0.0020411997
21887619
GO:0060135: maternal process involved in female pregnancy       ITGA3   integrin, alpha 3 (antigen CD
49C, alpha 3 subunit of VLA-3 receptor) 0.0014006125361972638
GO:0060135: maternal process involved in female pregnancy       LGALS9  lectin, galactoside-binding,
 soluble, 9     -0.00011694111963838872
GO:0060135: maternal process involved in female pregnancy       MMP7    matrix metallopeptidase 7 (ma
trilysin, uterine)      0.00042535684990099355
GO:0060259: regulation of feeding behavior      AGRP    agouti related protein homolog (mouse)  -0.00
25994695453300212
GO:2000253: positive regulation of feeding behavior     AGRP    agouti related protein homolog (mous
e)      -0.0025994695453300212
KEGG:04614: Renin-angiotensin system    AGT     angiotensinogen (serpin peptidase inhibitor, clade A,
member 8)       -0.0011169922119025706
KEGG:04614: Renin-angiotensin system    AGTR1   angiotensin II receptor, type 1 0.0003452163775896156
KEGG:04614: Renin-angiotensin system    MAS1    MAS1 proto-oncogene, G protein-coupled receptor 0.000
29097664979938224
KEGG:04924: Renin secretion     AGT     angiotensinogen (serpin peptidase inhibitor, clade A, member
 8)     -0.0011139206084095313
KEGG:04924: Renin secretion     AGTR1   angiotensin II receptor, type 1 0.0003453941297338615
KEGG:04924: Renin secretion     CREB1   cAMP responsive element binding protein 1       0.00064678059
74784756
KEGG:04924: Renin secretion     ITPR1   inositol 1,4,5-trisphosphate receptor, type 1   -0.0007686695
942128233
KEGG:04924: Renin secretion     KCNJ2   potassium inwardly-rectifying channel, subfamily J, member 2
-0.0007667497713812747
KEGG:04924: Renin secretion     KCNMA1  potassium large conductance calcium-activated channel, subfam
ily M, alpha member 1   -0.0006468055333396447
KEGG:04924: Renin secretion     PLCB1   phospholipase C, beta 1 (phosphoinositide-specific)     0.000
14994329191748572

```
KEGG:04924: Renin secretion     PRKX    protein kinase, X-linked        0.00038800348559060315
GO:0001543: ovarian follicle rupture    AGT     angiotensinogen (serpin peptidase inhibitor, clade A,
member 8)       -0.0011335364167528725
GO:0001543: ovarian follicle rupture    NRIP1   nuclear receptor interacting protein 1  0.00106908393
781542
GO:0001558: regulation of cell growth   AGT     angiotensinogen (serpin peptidase inhibitor, clade A,
member 8)       -0.0011239591957926942
GO:0001558: regulation of cell growth   AGTR1   angiotensin II receptor, type 1 0.0003464629597979704
GO:0001558: regulation of cell growth   FOXM1   forkhead box M1 0.00019628552953102402
GO:0001558: regulation of cell growth   IGFBP2  insulin-like growth factor binding protein 2, 36kDa
0.00014265375258051436
GO:0001558: regulation of cell growth   IGFBP3  insulin-like growth factor binding protein 3    0.000
8327229314907617
GO:0001558: regulation of cell growth   IGFBP4  insulin-like growth factor binding protein 4    -0.00
12706687737660473
GO:0001558: regulation of cell growth   IGFBP6  insulin-like growth factor binding protein 6    -0.00
20158103178430824
GO:0001558: regulation of cell growth   KIF14   kinesin family member 14        0.0004840104428949066
7
GO:0001568: blood vessel development    AGT     angiotensinogen (serpin peptidase inhibitor, clade A,
member 8)       -0.0011386435333327549
GO:0001568: blood vessel development    ALDH1A2 aldehyde dehydrogenase 1 family, member A2      -0.00
3028229487703539
GO:0001568: blood vessel development    COL1A1  collagen, type I, alpha 1       -0.000529444988126817
7
GO:0001568: blood vessel development    COL5A1  collagen, type V, alpha 1       -5.199023414011716e-5
GO:0001568: blood vessel development    FOXO1   forkhead box O1 0.0018057365213942013
GO:0001568: blood vessel development    LOX     lysyl oxidase   -0.0005785892056026905
GO:0001568: blood vessel development    MEF2C   myocyte enhancer factor 2C      0.000977010443855564
GO:0001568: blood vessel development    PAX6    paired box 6    0.0019421263098318656
GO:0001568: blood vessel development    SPHK1   sphingosine kinase 1    0.0018168231948020308
GO:0001568: blood vessel development    STRA6   stimulated by retinoic acid 6   -0.001686052197201494
2
GO:0001568: blood vessel development    TBX3    T-box 3 0.0012289109230176056
GO:0001568: blood vessel development    TCF7L2  transcription factor 7-like 2 (T-cell specific, HMG-b
ox)     0.000576176223437502
GO:0001658: branching involved in ureteric bud morphogenesis    AGT     angiotensinogen (serpin pepti
dase inhibitor, clade A, member 8)      -0.0011324215955683156
GO:0001658: branching involved in ureteric bud morphogenesis    BCL2    B-cell CLL/lymphoma 2   -4.78
3028522101644e-6
GO:0001658: branching involved in ureteric bud morphogenesis    BMP4    bone morphogenetic protein 4
-0.00032231007745216515
GO:0001658: branching involved in ureteric bud morphogenesis    CITED1  Cbp/p300-interacting transact
ivator, with Glu/Asp-rich carboxy-terminal domain, 1    0.0028166717397520523
GO:0001658: branching involved in ureteric bud morphogenesis    CTNNB1  catenin (cadherin-associated
 protein), beta 1, 88kDa        -0.00011696429615207505
GO:0001658: branching involved in ureteric bud morphogenesis    CTNNBIP1        catenin, beta interac
ting protein 1  0.0005853909251136399
GO:0001658: branching involved in ureteric bud morphogenesis    EYA1    EYA transcriptional coactivat
or and phosphatase 1    5.310144828986275e-5
GO:0001658: branching involved in ureteric bud morphogenesis    FGF8    fibroblast growth factor 8 (a
ndrogen-induced)        0.0009833334907729443
GO:0001658: branching involved in ureteric bud morphogenesis    GDNF    glial cell derived neurotroph
ic factor       0.0004445285581730751
GO:0001658: branching involved in ureteric bud morphogenesis    GLI3    GLI family zinc finger 3
-0.0021500712390826915
GO:0001658: branching involved in ureteric bud morphogenesis    GREM1   gremlin 1, DAN family BMP ant
agonist -0.0008268716043978844
GO:0001658: branching involved in ureteric bud morphogenesis    LHX1    LIM homeobox 1  -0.0007582861
704903153
GO:0001658: branching involved in ureteric bud morphogenesis    MYC     v-myc avian myelocytomatosis
 viral oncogene homolog -0.0011360947770484148
GO:0001658: branching involved in ureteric bud morphogenesis    PAX2    paired box 2    -0.0016129605
463831643
GO:0001658: branching involved in ureteric bud morphogenesis    PAX8    paired box 8    0.00094763224
99311977
GO:0001658: branching involved in ureteric bud morphogenesis    PTCH1   patched 1       -6.5969178795
42922e-5
GO:0001658: branching involved in ureteric bud morphogenesis    SALL1   spalt-like transcription fact
or 1    -0.0024359193832538743
GO:0001658: branching involved in ureteric bud morphogenesis    SHH     sonic hedgehog  0.00059959412
96707862
GO:0001658: branching involved in ureteric bud morphogenesis    SIX1    SIX homeobox 1  -0.0018688091
832661283
GO:0001658: branching involved in ureteric bud morphogenesis    SOX9    SRY (sex determining region
 Y)-box 9       -0.000518501685048646
GO:0001658: branching involved in ureteric bud morphogenesis    WNT1    wingless-type MMTV integratio
n site family, member 1 0.0007864630843607673
GO:0001658: branching involved in ureteric bud morphogenesis    WNT4    wingless-type MMTV integratio
n site family, member 4 -0.0002517234722073028
GO:0001658: branching involved in ureteric bud morphogenesis    WT1     Wilms tumor 1   -0.0005066333
```

996672473
GO:0001819: positive regulation of cytokine production AGT angiotensinogen (serpin peptidase inhibitor, clade A, member 8) -0.0011514506577944977
GO:0001819: positive regulation of cytokine production FLOT1 flotillin 1 -0.000770043146373940
4
GO:0001819: positive regulation of cytokine production LEP leptin 0.00323030203018037l
GO:0001822: kidney development AGT angiotensinogen (serpin peptidase inhibitor, clade A, member 8) -0.0011322885087761368
GO:0001822: kidney development AGTR1 angiotensin II receptor, type 1 0.00034764348513865453
GO:0001822: kidney development ALDH1A2 aldehyde dehydrogenase 1 family, member A2 -0.0030174404
523699142
GO:0001822: kidney development APC adenomatous polyposis coli 0.0006435382371252014
GO:0001822: kidney development ARL3 ADP-ribosylation factor-like 3 -0.00118026928474688l
GO:0001822: kidney development ASS1 argininosuccinate synthase 1 0.0001423666715683510 3
GO:0001822: kidney development BAX BCL2-associated X protein -0.00042495573157465396
GO:0001822: kidney development BCL2L11 BCL2-like 11 (apoptosis facilitator) -0.000774409498806054
6
GO:0001822: kidney development BMP4 bone morphogenetic protein 4 -0.00032241516797292723
GO:0001822: kidney development GATA3 GATA binding protein 3 -3.8969050880196824e-5
GO:0001822: kidney development GLI2 GLI family zinc finger 2 0.001852388754220484
GO:0001822: kidney development JMJD6 jumonji domain containing 6 0.003588983878481648
GO:0001822: kidney development LEF1 lymphoid enhancer-binding factor 1 -0.000100306752723667
15
GO:0001822: kidney development LHX1 LIM homeobox 1 -0.0007582482598129829
GO:0001822: kidney development ODC1 ornithine decarboxylase 1 0.0008248122290811506
GO:0001822: kidney development PAX8 paired box 8 0.0009477606784070615
GO:0001822: kidney development PROX1 prospero homeobox 1 0.0011207695483854566
GO:0001822: kidney development SALL1 spalt-like transcription factor 1 -0.002435606551412037
GO:0001822: kidney development SIX1 SIX homeobox 1 -0.001868872554158794
GO:0001822: kidney development SOX11 SRY (sex determining region Y)-box 11 -0.000209040638474912
34
GO:0001822: kidney development STRA6 stimulated by retinoic acid 6 -0.0016794296292797035
GO:0001822: kidney development SULF1 sulfatase 1 -0.0008049564743280628
GO:0001822: kidney development TFAP2A transcription factor AP-2 alpha (activating enhancer binding protein 2 alpha) 0.0004796553809574216
GO:0001822: kidney development TGFBR1 transforming growth factor, beta receptor 1 0.00033964589
359753404
GO:0001822: kidney development THRA thyroid hormone receptor, alpha 0.0007627917589666247
GO:0001822: kidney development TP73 tumor protein p73 0.0010272788843812654
GO:0001822: kidney development VEGFA vascular endothelial growth factor A 0.0005944993942241
GO:0001822: kidney development WFS1 Wolfram syndrome 1 (wolframin) 0.0005765835697601829
GO:0001822: kidney development WNT4 wingless-type MMTV integration site family, member 4 -0.00
02517607146785744
GO:0001822: kidney development WT1 Wilms tumor 1 -0.0005065112946419222
GO:0001998: angiotensin mediated vasoconstriction involved in regulation of systemic arterial blood pressure AGT angiotensinogen (serpin peptidase inhibitor, clade A, member 8) -0.00114132446189895 3
GO:0001999: renal response to blood flow involved in circulatory renin-angiotensin regulation of systemic arterial blood pressure AGT angiotensinogen (serpin peptidase inhibitor, clade A, member 8) -0.00114132446189895 3
GO:0002003: angiotensin maturation AGT angiotensinogen (serpin peptidase inhibitor, clade A, member 8) -0.001141324461898953
GO:0002016: regulation of blood volume by renin-angiotensin AGT angiotensinogen (serpin peptidase inhibitor, clade A, member 8) -0.001141324461898953
GO:0002018: renin-angiotensin regulation of aldosterone production AGT angiotensinogen (serpin peptidase inhibitor, clade A, member 8) -0.0011273061613599606
GO:0002018: renin-angiotensin regulation of aldosterone production AGTR1 angiotensin II receptor, type 1 0.00034518432154975775
GO:0002019: regulation of renal output by angiotensin AGT angiotensinogen (serpin peptidase inhibitor, clade A, member 8) -0.001141324461898953
GO:0002027: regulation of heart rate AGT angiotensinogen (serpin peptidase inhibitor, clade A, member 8) -0.0011190207888007173
GO:0002027: regulation of heart rate DRD2 dopamine receptor D2 -0.00023222127721904368
GO:0002027: regulation of heart rate GPD1L glycerol-3-phosphate dehydrogenase 1-like 0.000
5611155218810469
GO:0002034: regulation of blood vessel size by renin-angiotensin AGT angiotensinogen (serpin peptidase inhibitor, clade A, member 8) -0.0011197730280088998
GO:0002034: regulation of blood vessel size by renin-angiotensin AGTR1 angiotensin II receptor, type 1 0.0003459028828390447
GO:0002034: regulation of blood vessel size by renin-angiotensin SERPINF2 serpin peptidase inhibitor, clade F (alpha-2 antiplasmin, pigment epithelium derived factor), member 2 0.000
652590444932185
GO:0003014: renal system process AGT angiotensinogen (serpin peptidase inhibitor, clade A, member 8) -0.0011242494876576785
GO:0003014: renal system process BCL2 B-cell CLL/lymphoma 2 -4.80962211010082e-6
GO:0003014: renal system process BMP4 bone morphogenetic protein 4 -0.000320652952761324
9
GO:0003014: renal system process FAS Fas cell surface death receptor -3.3127978881144846e-
5
GO:0003014: renal system process PKN1 protein kinase N1 -0.0018923139906094533
GO:0003051: angiotensin-mediated drinking behavior AGT angiotensinogen (serpin peptidase inhibitor, clade A, member 8) -0.001141324461898953

GO:0003331: positive regulation of extracellular matrix constituent secretion AGT angiotensinog
en (serpin peptidase inhibitor, clade A, member 8) -0.001141324461898953
GO:0006883: cellular sodium ion homeostasis AGT angiotensinogen (serpin peptidase inhibitor,
clade A, member 8) -0.001137812587929324
GO:0006883: cellular sodium ion homeostasis ATP1B3 ATPase, Na+/K+ transporting, beta 3 polypepti
de -0.0015001052654050246
GO:0007160: cell-matrix adhesion AGT angiotensinogen (serpin peptidase inhibitor, clade A,
member 8) -0.0011379943618682912
GO:0007160: cell-matrix adhesion BCL2L11 BCL2-like 11 (apoptosis facilitator) -0.0007757746
956300362
GO:0007160: cell-matrix adhesion CD63 CD63 molecule -0.0010646265330701342
GO:0007160: cell-matrix adhesion CTNNB1 catenin (cadherin-associated protein), beta 1, 88kDa
-0.00011670494850022758
GO:0007160: cell-matrix adhesion EDA ectodysplasin A -0.0008172001685629282
GO:0007160: cell-matrix adhesion ITGA3 integrin, alpha 3 (antigen CD49C, alpha 3 subunit of
VLA-3 receptor) 0.0014023025785692765
GO:0007160: cell-matrix adhesion ITGA6 integrin, alpha 6 0.0016758569954095257
GO:0007160: cell-matrix adhesion ITGB1 integrin, beta 1 (fibronectin receptor, beta polypept
ide, antigen CD29 includes MDF2, MSK12) 0.0024336332480607284
GO:0007160: cell-matrix adhesion ITGB4 integrin, beta 4 0.0005828388908821711
GO:0007160: cell-matrix adhesion THBS3 thrombospondin 3 0.0016203013163478294
GO:0007186: G-protein coupled receptor signaling pathway AGT angiotensinogen (serpin pepti
dase inhibitor, clade A, member 8) -0.0011356818867707846
GO:0007186: G-protein coupled receptor signaling pathway AGTR1 angiotensin II receptor, type
1 0.00034727750241947874
GO:0007186: G-protein coupled receptor signaling pathway AKT1 v-akt murine thymoma viral on
cogene homolog 1 0.0007317522629380935
GO:0007186: G-protein coupled receptor signaling pathway AREG amphiregulin 0.00246009128
5627331
GO:0007186: G-protein coupled receptor signaling pathway C3 complement component 3 0.002
027451858856668
GO:0007186: G-protein coupled receptor signaling pathway CCL2 chemokine (C-C motif) ligand
2 0.0008179630959293955
GO:0007186: G-protein coupled receptor signaling pathway CCL7 chemokine (C-C motif) ligand
7 -0.00240494535122658
GO:0007186: G-protein coupled receptor signaling pathway CCL8 chemokine (C-C motif) ligand
8 -0.0005905841499930065
GO:0007186: G-protein coupled receptor signaling pathway CELSR2 cadherin, EGF LAG seven-pass
G-type receptor 2 -0.0011365478355988014
GO:0007186: G-protein coupled receptor signaling pathway CX3CL1 chemokine (C-X3-C motif) liga
nd 1 0.00212059621953148
GO:0007186: G-protein coupled receptor signaling pathway CX3CR1 chemokine (C-X3-C motif) rece
ptor 1 -0.0008580729258881208
GO:0007186: G-protein coupled receptor signaling pathway CXCL10 chemokine (C-X-C motif) ligan
d 10 6.3194193373997e-5
GO:0007186: G-protein coupled receptor signaling pathway CXCL12 chemokine (C-X-C motif) ligan
d 12 -0.0012103273851138334
GO:0007186: G-protein coupled receptor signaling pathway CXCL13 chemokine (C-X-C motif) ligan
d 13 0.0029976064731152066
GO:0007186: G-protein coupled receptor signaling pathway CXCR4 chemokine (C-X-C motif) recep
tor 4 0.0007961901815883602
GO:0007186: G-protein coupled receptor signaling pathway FRS2 fibroblast growth factor rece
ptor substrate 2 0.0009311209785255436
GO:0007186: G-protein coupled receptor signaling pathway FZD3 frizzled class receptor 3
0.0003795844270815237
GO:0007186: G-protein coupled receptor signaling pathway FZD7 frizzled class receptor 7
0.0010725725389893167
GO:0007186: G-protein coupled receptor signaling pathway GNRHR gonadotropin-releasing hormon
e receptor 0.0007414722958120309
GO:0007186: G-protein coupled receptor signaling pathway GPR87 G protein-coupled receptor 87
-2.4972515873510258e-5
GO:0007186: G-protein coupled receptor signaling pathway GPSM2 G-protein signaling modulator
2 0.001775301350673212
GO:0007186: G-protein coupled receptor signaling pathway INSR insulin receptor -0.00
13653012317385407
GO:0007186: G-protein coupled receptor signaling pathway JAK2 Janus kinase 2 -3.1294157820
48628e-5
GO:0007186: G-protein coupled receptor signaling pathway LGR4 leucine-rich repeat containin
g G protein-coupled receptor 4 0.00030888653092908897
GO:0007186: G-protein coupled receptor signaling pathway MAS1 MAS1 proto-oncogene, G protei
n-coupled receptor 0.0002911040070759541
GO:0007186: G-protein coupled receptor signaling pathway NMU neuromedin U 0.00185347482
34731637
GO:0007186: G-protein coupled receptor signaling pathway OR5I1 olfactory receptor, family 5,
subfamily I, member 1 0.0018843550241663886
GO:0007186: G-protein coupled receptor signaling pathway PDGFRB platelet-derived growth facto
r receptor, beta polypeptide -0.000362477823094391
GO:0007186: G-protein coupled receptor signaling pathway SMO smoothened, frizzled class re
ceptor 0.0021421930109414007
GO:0007186: G-protein coupled receptor signaling pathway TULP3 tubby like protein 3 0.000
9579090710494537

GO:0007199: G-protein coupled receptor signaling pathway coupled to cGMP nucleotide second messenger
AGT     angiotensinogen (serpin peptidase inhibitor, clade A, member 8) -0.001141324461898953
GO:0007200: phospholipase C-activating G-protein coupled receptor signaling pathway     AGT     angio
tensinogen (serpin peptidase inhibitor, clade A, member 8)      -0.0011148533931377087
GO:0007200: phospholipase C-activating G-protein coupled receptor signaling pathway     AGTR1   angio
tensin II receptor, type 1      0.00034567159193022926
GO:0007200: phospholipase C-activating G-protein coupled receptor signaling pathway     ESR1    estro
gen receptor 1  -0.0009352668420446904
GO:0007200: phospholipase C-activating G-protein coupled receptor signaling pathway     HTR2C   5-hyd
roxytryptamine (serotonin) receptor 2C, G protein-coupled       -0.0005181719878916908
GO:0007200: phospholipase C-activating G-protein coupled receptor signaling pathway     TGM2    trans
glutaminase 2   -0.0001769681504918573
GO:0007202: activation of phospholipase C activity      AGT     angiotensinogen (serpin peptidase inh
ibitor, clade A, member 8)      -0.0011236894533061513
GO:0007202: activation of phospholipase C activity      CREB1   cAMP responsive element binding prote
in 1    0.0006526255386096972
GO:0007202: activation of phospholipase C activity      EGFR    epidermal growth factor receptor
0.0006800670113925344
GO:0007202: activation of phospholipase C activity      ITPR1   inositol 1,4,5-trisphosphate recepto
r, type 1       -0.0007744885115874012
GO:0007202: activation of phospholipase C activity      PRKCA   protein kinase C, alpha -5.8315058233
737055e-6
GO:0007202: activation of phospholipase C activity      PRKCD   protein kinase C, delta -0.0011195788
348135893
GO:0007250: activation of NF-kappaB-inducing kinase activity    AGT     angiotensinogen (serpin pepti
dase inhibitor, clade A, member 8)      -0.0011342443589731632
GO:0007250: activation of NF-kappaB-inducing kinase activity    IRAK1   interleukin-1 receptor-associ
ated kinase 1   -0.0016090896509 2749
GO:0007250: activation of NF-kappaB-inducing kinase activity    MAS1    MAS1 proto-oncogene, G protei
n-coupled receptor      0.0002907531420779391
GO:0007263: nitric oxide mediated signal transduction   AGT     angiotensinogen (serpin peptidase inh
ibitor, clade A, member 8)      -0.0011272561800915576
GO:0007263: nitric oxide mediated signal transduction   PDX1    pancreatic and duodenal homeobox 1
0.0002558342245161702
GO:0007588: excretion   AGT     angiotensinogen (serpin peptidase inhibitor, clade A, member 8) -0.00
1119416453216491
GO:0007588: excretion   HMOX1   heme oxygenase (decycling) 1    -0.00021522995210703457
GO:0007588: excretion   SLC22A18        solute carrier family 22, member 18     0.000727406454103647
GO:0008065: establishment of blood-nerve barrier        AGT     angiotensinogen (serpin peptidase inh
ibitor, clade A, member 8)      -0.001136418554814798
GO:0008065: establishment of blood-nerve barrier        GSTM3   glutathione S-transferase mu 3 (brai
n)      -0.0012904510104553937
GO:0009651: response to salt stress     AGT     angiotensinogen (serpin peptidase inhibitor, clade A,
member 8)       -0.001125037157788183
GO:0009651: response to salt stress     BAX     BCL2-associated X protein       -0.000422612395972620
74
GO:0009651: response to salt stress     TH      tyrosine hydroxylase    -0.0003404987448474476
GO:0009651: response to salt stress     TP53    tumor protein p53       0.0011676634794120422
GO:0010535: positive regulation of activation of JAK2 kinase activity   AGT     angiotensinogen (serp
in peptidase inhibitor, clade A, member 8)      -0.0011340970972180376
GO:0010535: positive regulation of activation of JAK2 kinase activity   IL12B   interleukin 12B 0.001
2764517062932865
GO:0010595: positive regulation of endothelial cell migration   AGT     angiotensinogen (serpin pepti
dase inhibitor, clade A, member 8)      -0.0011220269800965622
GO:0010595: positive regulation of endothelial cell migration   ALOX12  arachidonate 12-lipoxygenase
-0.0019394077472052165
GO:0010595: positive regulation of endothelial cell migration   ANGPT1  angiopoietin 1  0.00089504716
83722529
GO:0010595: positive regulation of endothelial cell migration   BMP4    bone morphogenetic protein 4
-0.0003203270222288023
GO:0010595: positive regulation of endothelial cell migration   FOXC2   forkhead box C2 (MFH-1, mesen
chyme forkhead 1)       0.0017363448333599623
GO:0010595: positive regulation of endothelial cell migration   GATA3   GATA binding protein 3  -3.76
63198222700246e-5
GO:0010595: positive regulation of endothelial cell migration   NRP1    neuropilin 1    -0.0006378237
944011305
GO:0010595: positive regulation of endothelial cell migration   PRKCA   protein kinase C, alpha -5.93
64492213861e-6
GO:0010595: positive regulation of endothelial cell migration   PROX1   prospero homeobox 1     0.001
1126813719566889
GO:0010595: positive regulation of endothelial cell migration   THBS1   thrombospondin 1        -0.00
10283456165712283
GO:0010595: positive regulation of endothelial cell migration   VEGFA   vascular endothelial growth f
actor A 0.0005889216945167236
GO:0010595: positive regulation of endothelial cell migration   WNT5A   wingless-type MMTV integratio
n site family, member 5A        -0.000662291012087647 7
GO:0010595: positive regulation of endothelial cell migration   WNT7A   wingless-type MMTV integratio
n site family, member 7A        1.9848836577096883e-5
GO:0010613: positive regulation of cardiac muscle hypertrophy   AGT     angiotensinogen (serpin pepti
dase inhibitor, clade A, member 8)      -0.001127706138226641
GO:0010613: positive regulation of cardiac muscle hypertrophy   IGF1    insulin-like growth factor 1

(somatomedin C)        0.00012999061425373693
GO:0010613: positive regulation of cardiac muscle hypertrophy   IL6ST   interleukin 6 signal transducer      0.001870096656151251
GO:0010613: positive regulation of cardiac muscle hypertrophy   PRKCA   protein kinase C, alpha -5.9528413322747466e-6
GO:0010744: positive regulation of macrophage derived foam cell differentiation AGT     angiotensinogen (serpin peptidase inhibitor, clade A, member 8)      -0.0011273061613599606
GO:0010744: positive regulation of macrophage derived foam cell differentiation AGTR1   angiotensin II receptor, type 1      0.00034518432154975775
GO:0010873: positive regulation of cholesterol esterification   AGT     angiotensinogen (serpin peptidase inhibitor, clade A, member 8)      -0.0011273061613599606
GO:0010873: positive regulation of cholesterol esterification   AGTR1   angiotensin II receptor, type 1       0.00034518432154975775
GO:0010951: negative regulation of endopeptidase activity       AGT     angiotensinogen (serpin peptidase inhibitor, clade A, member 8)      -0.001142786796267062
GO:0010951: negative regulation of endopeptidase activity       AKT1    v-akt murine thymoma viral oncogene homolog 1        0.000735711899928649
GO:0010951: negative regulation of endopeptidase activity       C3      complement component 3  0.002037330251145309
GO:0010951: negative regulation of endopeptidase activity       CAST    calpastatin     -0.003052321195622298
GO:0010951: negative regulation of endopeptidase activity       CST3    cystatin C      -7.225017747036895e-5
GO:0010951: negative regulation of endopeptidase activity       PI3     peptidase inhibitor 3, skin-derived  -0.0010467915172388358
GO:0010951: negative regulation of endopeptidase activity       PTTG1   pituitary tumor-transforming 1      -0.0003621731364624223
GO:0010951: negative regulation of endopeptidase activity       SERPINA5        serpin peptidase inhibitor, clade A (alpha-1 antiproteinase, antitrypsin), member 5  -0.000713912592934043
GO:0010951: negative regulation of endopeptidase activity       SERPINB5        serpin peptidase inhibitor, clade B (ovalbumin), member 5    -0.004556683309996608
GO:0010951: negative regulation of endopeptidase activity       SERPINE1        serpin peptidase inhibitor, clade E (nexin, plasminogen activator inhibitor type 1), member 1        0.00011187084480742906
GO:0010951: negative regulation of endopeptidase activity       SERPINE2        serpin peptidase inhibitor, clade E (nexin, plasminogen activator inhibitor type 1), member 2        0.0025179901756495976
GO:0010951: negative regulation of endopeptidase activity       SERPINF2        serpin peptidase inhibitor, clade F (alpha-2 antiplasmin, pigment epithelium derived factor), member 2       0.0006623869558177233
GO:0010951: negative regulation of endopeptidase activity       SERPINH1        serpin peptidase inhibitor, clade H (heat shock protein 47), member 1, (collagen binding protein 1)  -0.0010955084418645005
GO:0014061: regulation of norepinephrine secretion      AGT     angiotensinogen (serpin peptidase inhibitor, clade A, member 8)      -0.001141324461898953
GO:0014068: positive regulation of phosphatidylinositol 3-kinase signaling      AGT     angiotensinogen (serpin peptidase inhibitor, clade A, member 8)      -0.0011254763501075066
GO:0014068: positive regulation of phosphatidylinositol 3-kinase signaling      ANGPT1  angiopoietin 1      0.0008966105857940089
GO:0014068: positive regulation of phosphatidylinositol 3-kinase signaling      FLT3    fms-related tyrosine kinase 3        -0.0006772422850948269
GO:0014068: positive regulation of phosphatidylinositol 3-kinase signaling      IGF1    insulin-like growth factor 1 (somatomedin C)        0.0001295003642962163
GO:0014068: positive regulation of phosphatidylinositol 3-kinase signaling      JAK2    Janus kinase 2      -3.2625643350948375e-5
GO:0014068: positive regulation of phosphatidylinositol 3-kinase signaling      KIT     v-kit Hardy-Zuckerman 4 feline sarcoma viral oncogene homolog       0.00027163259255019273
GO:0014068: positive regulation of phosphatidylinositol 3-kinase signaling      NEDD4   neural precursor cell expressed, developmentally down-regulated 4, E3 ubiquitin protein ligase      0.002252824626825657
GO:0014068: positive regulation of phosphatidylinositol 3-kinase signaling      PDGFRB  platelet-derived growth factor receptor, beta polypeptide    -0.0003578912793950707
GO:0014068: positive regulation of phosphatidylinositol 3-kinase signaling      RELN    reelin  0.0015284506475392243
GO:0014068: positive regulation of phosphatidylinositol 3-kinase signaling      SOX9    SRY (sex determining region Y)-box 9 -0.0005160082523193129
GO:0014068: positive regulation of phosphatidylinositol 3-kinase signaling      TGFB2   transforming growth factor, beta 2  -0.0010534663100429142
GO:0014824: artery smooth muscle contraction    AGT     angiotensinogen (serpin peptidase inhibitor, clade A, member 8)     -0.001141324461898953
GO:0014873: response to muscle activity involved in regulation of muscle adaptation     AGT     angiotensinogen (serpin peptidase inhibitor, clade A, member 8)      -0.001141324461898953
GO:0016525: negative regulation of angiogenesis AGT     angiotensinogen (serpin peptidase inhibitor, clade A, member 8)     -0.0011142345282872878
GO:0016525: negative regulation of angiogenesis CCL2    chemokine (C-C motif) ligand 2  0.0008062675702981251
GO:0016525: negative regulation of angiogenesis CX3CR1  chemokine (C-X3-C motif) receptor 1     -0.0008474072551148822
GO:0016525: negative regulation of angiogenesis CXCL10  chemokine (C-X-C motif) ligand 10       6.0373743432934394e-5
GO:0016525: negative regulation of angiogenesis FOXC1   forkhead box C1 -2.313858152175273e-5
GO:0016525: negative regulation of angiogenesis HOXA5   homeobox A5     0.0010503760111728365

GO:0016525: negative regulation of angiogenesis PML promyelocytic leukemia -0.0006733407439434915
GO:0016525: negative regulation of angiogenesis SULF1 sulfatase 1 -0.0007903565640150295
GO:0016525: negative regulation of angiogenesis THBS1 thrombospondin 1 -0.0010227421262162888
GO:0016525: negative regulation of angiogenesis THBS4 thrombospondin 4 -0.00044121451627804196
GO:0016525: negative regulation of angiogenesis VASH1 vasohibin 1 0.0005917174396845662
GO:0019229: regulation of vasoconstriction AGT angiotensinogen (serpin peptidase inhibitor, clade A, member 8) -0.0011377222488670722
GO:0019229: regulation of vasoconstriction AGTR1 angiotensin II receptor, type 1 0.00034766174511833127
GO:0019229: regulation of vasoconstriction ASIC2 acid-sensing (proton-gated) ion channel 2 0.0022858783346506183
GO:0019229: regulation of vasoconstriction PER2 period circadian clock 2 0.0012416736147635248
GO:0030198: extracellular matrix organization AGT angiotensinogen (serpin peptidase inhibitor, clade A, member 8) -0.0011353962802214247
GO:0030198: extracellular matrix organization BMP4 bone morphogenetic protein 4 -0.0003230041778459228
GO:0030198: extracellular matrix organization BMP7 bone morphogenetic protein 7 0.0008564480838143905
GO:0030198: extracellular matrix organization COL11A1 collagen, type XI, alpha 1 -0.00044233237692936205
GO:0030198: extracellular matrix organization COL1A1 collagen, type I, alpha 1 -0.0005275772332481822
GO:0030198: extracellular matrix organization COL5A1 collagen, type V, alpha 1 -5.0651252001064865e-5
GO:0030198: extracellular matrix organization COL5A2 collagen, type V, alpha 2 -0.0003465949792463343
GO:0030198: extracellular matrix organization CTSK cathepsin K -0.00036403213030256103
GO:0030198: extracellular matrix organization CTSV cathepsin V 0.0006550577223109953
GO:0030198: extracellular matrix organization DAG1 dystroglycan 1 (dystrophin-associated glycoprotein 1) 0.0003620444556816928
GO:0030198: extracellular matrix organization DSPP dentin sialophosphoprotein 0.0008306362325807082
GO:0030198: extracellular matrix organization EGFL6 EGF-like-domain, multiple 6 0.00013687380078647782
GO:0030198: extracellular matrix organization FBN2 fibrillin 2 -0.00074239444549566
GO:0030198: extracellular matrix organization ICAM1 intercellular adhesion molecule 1 0.000729635439617389
GO:0030198: extracellular matrix organization ITGA3 integrin, alpha 3 (antigen CD49C, alpha 3 subunit of VLA-3 receptor) 0.0014011452838385526
GO:0030198: extracellular matrix organization ITGA6 integrin, alpha 6 0.0016721877762541249
GO:0030198: extracellular matrix organization ITGB1 integrin, beta 1 (fibronectin receptor, beta polypeptide, antigen CD29 includes MDF2, MSK12) 0.0024300089939523427
GO:0030198: extracellular matrix organization ITGB4 integrin, beta 4 0.0005814874489974711
GO:0030198: extracellular matrix organization KLK7 kallikrein-related peptidase 7 0.0016280003757965932
GO:0030198: extracellular matrix organization LAMB2 laminin, beta 2 (laminin S) -0.0013108738780589946
GO:0030198: extracellular matrix organization LOX lysyl oxidase -0.0005772349323089477
GO:0030198: extracellular matrix organization MMP2 matrix metallopeptidase 2 (gelatinase A, 72kDa gelatinase, 72kDa type IV collagenase) -0.0011332886696424156
GO:0030198: extracellular matrix organization MMP20 matrix metallopeptidase 20 -0.002434669398792978
GO:0030198: extracellular matrix organization MMP7 matrix metallopeptidase 7 (matrilysin, uterine) 0.0004259156495676094
GO:0030198: extracellular matrix organization NR2E1 nuclear receptor subfamily 2, group E, member 1 -0.0029780332048168334
GO:0030198: extracellular matrix organization PRDX4 peroxiredoxin 4 0.0015264475327929482
GO:0030198: extracellular matrix organization PRKCA protein kinase C, alpha -5.839797129212219e-6
GO:0030198: extracellular matrix organization SERPINB5 serpin peptidase inhibitor, clade B (ovalbumin), member 5 -0.004536539408318589
GO:0030198: extracellular matrix organization SERPINE1 serpin peptidase inhibitor, clade E (nexin, plasminogen activator inhibitor type 1), member 1 0.00011213224138966569
GO:0030198: extracellular matrix organization SERPINH1 serpin peptidase inhibitor, clade H (heat shock protein 47), member 1, (collagen binding protein 1) -0.0010902762951661925
GO:0030198: extracellular matrix organization SOX9 SRY (sex determining region Y)-box 9 -0.0005191319035850547
GO:0030198: extracellular matrix organization TGFB1 transforming growth factor, beta 1 -7.32614485706903e-5
GO:0030198: extracellular matrix organization TGFB2 transforming growth factor, beta 2 -0.0010608832620807034
GO:0030198: extracellular matrix organization TGFB3 transforming growth factor, beta 3 -0.0018265300195037517
GO:0030198: extracellular matrix organization THBS1 thrombospondin 1 -0.0010372329481276883
GO:0030198: extracellular matrix organization TNC tenascin C 0.0007352193829639208
GO:0030432: peristalsis AGT angiotensinogen (serpin peptidase inhibitor, clade A, member 8) -0.0011158610057580539

GO:0030432: peristalsis DRD2    dopamine receptor D2    -0.00023187057536279692
GO:0030432: peristalsis GDNF    glial cell derived neurotrophic factor  0.00044020063148044017
GO:0032270: positive regulation of cellular protein metabolic process   AGT     angiotensinogen (serpin peptidase inhibitor, clade A, member 8)      -0.0011252869951244595
GO:0032270: positive regulation of cellular protein metabolic process   AGTR1   angiotensin II receptor, type 1      0.0003465033700135899
GO:0032270: positive regulation of cellular protein metabolic process   AKT1    v-akt murine thymoma viral oncogene homolog 1       0.0007258870202631821
GO:0032270: positive regulation of cellular protein metabolic process   INHBA   inhibin, beta A -0.0013433361947508917
GO:0032270: positive regulation of cellular protein metabolic process   NR1H3   nuclear receptor subfamily 1, group H, member 3      0.0007965623366097777
GO:0032270: positive regulation of cellular protein metabolic process   TGFB1   transforming growth factor, beta 1   -7.252275201797459e-5
GO:0032930: positive regulation of superoxide anion generation  AGT     angiotensinogen (serpin peptidase inhibitor, clade A, member 8)      -0.0011230354599038475
GO:0032930: positive regulation of superoxide anion generation  GSTP1   glutathione S-transferase pi 1      0.0002302086985985101
GO:0032930: positive regulation of superoxide anion generation  PRKCD   protein kinase C, delta -0.0011189176103720967
GO:0032930: positive regulation of superoxide anion generation  TGFB1   transforming growth factor, beta 1   -7.253484562525232e-5
GO:0033138: positive regulation of peptidyl-serine phosphorylation      AGT     angiotensinogen (serpin peptidase inhibitor, clade A, member 8)      -0.0011282006662961485
GO:0033138: positive regulation of peptidyl-serine phosphorylation      AKT1    v-akt murine thymoma viral oncogene homolog 1       0.0007282841687977957
GO:0033138: positive regulation of peptidyl-serine phosphorylation      AKT2    v-akt murine thymoma viral oncogene homolog 2       -0.0006244734032888663
GO:0033138: positive regulation of peptidyl-serine phosphorylation      ANGPT1  angiopoietin 1  0.0008983012903031153
GO:0033138: positive regulation of peptidyl-serine phosphorylation      AVP     arginine vasopressin -0.0009339310067446518
GO:0033138: positive regulation of peptidyl-serine phosphorylation      AXIN1   axin 1  -0.0007304631354543684
GO:0033138: positive regulation of peptidyl-serine phosphorylation      BCL2    B-cell CLL/lymphoma 2 -4.701486539408031e-6
GO:0033138: positive regulation of peptidyl-serine phosphorylation      CAV1    caveolin 1, caveolae protein, 22kDa -0.0005343319103674759
GO:0033138: positive regulation of peptidyl-serine phosphorylation      CDC42   cell division cycle 42       0.0012366886409960769
GO:0033138: positive regulation of peptidyl-serine phosphorylation      GSK3B   glycogen synthase kinase 3 beta      0.0015446040341675611
GO:0033138: positive regulation of peptidyl-serine phosphorylation      PAK1    p21 protein (Cdc42/Rac)-activated kinase 1   -0.002181236255591259
GO:0033138: positive regulation of peptidyl-serine phosphorylation      PFN2    profilin 2      0.0019989673366821257
GO:0033138: positive regulation of peptidyl-serine phosphorylation      RAF1    Raf-1 proto-oncogene, serine/threonine kinase 0.0014944590290901214
GO:0033138: positive regulation of peptidyl-serine phosphorylation      TGFB1   transforming growth factor, beta 1   -7.240759311739791e-5
GO:0033138: positive regulation of peptidyl-serine phosphorylation      VEGFA   vascular endothelial growth factor A        0.0005920034529647972
GO:0033138: positive regulation of peptidyl-serine phosphorylation      WNT5A   wingless-type MMTV integration site family, member 5A        -0.0006650898199579928
GO:0033864: positive regulation of NAD(P)H oxidase activity     AGT     angiotensinogen (serpin peptidase inhibitor, clade A, member 8)      -0.0011273061613599606
GO:0033864: positive regulation of NAD(P)H oxidase activity     AGTR1   angiotensin II receptor, type 1       0.00034518432154975775
GO:0034104: negative regulation of tissue remodeling    AGT     angiotensinogen (serpin peptidase inhibitor, clade A, member 8)      -0.001141324461898953
GO:0035411: catenin import into nucleus AGT     angiotensinogen (serpin peptidase inhibitor, clade A, member 8)      -0.001141324461898953
GO:0035813: regulation of renal sodium excretion        AGT     angiotensinogen (serpin peptidase inhibitor, clade A, member 8)      -0.0011266866601655772
GO:0035813: regulation of renal sodium excretion        AGTR1   angiotensin II receptor, type 1 0.0003457120126052984
GO:0035813: regulation of renal sodium excretion        AVP     arginine vasopressin    -0.0009321890790038753
GO:0035815: positive regulation of renal sodium excretion       AGT     angiotensinogen (serpin peptidase inhibitor, clade A, member 8)      -0.0011260364256689736
GO:0035815: positive regulation of renal sodium excretion       DRD2    dopamine receptor D2    -0.00023290838230142068
GO:0040018: positive regulation of multicellular organism growth        AGT     angiotensinogen (serpin peptidase inhibitor, clade A, member 8)      -0.001125341647488298
GO:0040018: positive regulation of multicellular organism growth        BBS4    Bardet-Biedl syndrome 4       -0.0005073218178665523
GO:0040018: positive regulation of multicellular organism growth        BCL2    B-cell CLL/lymphoma 2 -4.803184511103303e-6
GO:0040018: positive regulation of multicellular organism growth        CREB1   cAMP responsive element binding protein 1    0.000653817210767881
GO:0040018: positive regulation of multicellular organism growth        DRD2    dopamine receptor D2

-0.00023319793886676708
GO:0040018: positive regulation of multicellular organism growth POU1F1 POU class 1 homeobox 1 0.00013657207900539276
GO:0040018: positive regulation of multicellular organism growth SMO smoothened, frizzled class receptor 0.002128862632710565
GO:0040018: positive regulation of multicellular organism growth STAT5A signal transducer and activator of transcription 5A 0.0015892017426617899
GO:0042127: regulation of cell proliferation AGT angiotensinogen (serpin peptidase inhibitor, clade A, member 8) -0.0011341826209634676
GO:0042127: regulation of cell proliferation AGTR1 angiotensin II receptor, type 1 0.0003471994036480126
GO:0042127: regulation of cell proliferation CHEK1 checkpoint kinase 1 0.0009009769811606014
GO:0042127: regulation of cell proliferation CIB1 calcium and integrin binding 1 (calmyrin) 5.4114351718589844e-5
GO:0042127: regulation of cell proliferation CXCL10 chemokine (C-X-C motif) ligand 10 6.302211511482038e-5
GO:0042127: regulation of cell proliferation CXCL13 chemokine (C-X-C motif) ligand 13 0.0029945756728374942
GO:0042127: regulation of cell proliferation E2F4 E2F transcription factor 4, p107/p130-binding -0.0023463311704491627
GO:0042127: regulation of cell proliferation ENG endoglin 0.0008269777790662978
GO:0042127: regulation of cell proliferation EZH2 enhancer of zeste 2 polycomb repressive complex 2 subunit -0.0001358448631660165
GO:0042127: regulation of cell proliferation FOXM1 forkhead box M1 0.000197832730867238
GO:0042127: regulation of cell proliferation GUCY2C guanylate cyclase 2C (heat stable enterotoxin receptor) -0.0016341243245748662
GO:0042127: regulation of cell proliferation HOXD13 homeobox D13 -0.0005924310398957534
GO:0042127: regulation of cell proliferation INHA inhibin, alpha 0.00021149484023045935
GO:0042127: regulation of cell proliferation JAG1 jagged 1 0.0017568436223542353
GO:0042127: regulation of cell proliferation JAG2 jagged 2 -5.598829809758457e-6
GO:0042127: regulation of cell proliferation JAK2 Janus kinase 2 -3.144520313963049e-5
GO:0042127: regulation of cell proliferation JUP junction plakoglobin -0.001479485583929458 8
GO:0042127: regulation of cell proliferation KIT v-kit Hardy-Zuckerman 4 feline sarcoma viral oncogene homolog 0.00027555034989152954
GO:0042127: regulation of cell proliferation MMP7 matrix metallopeptidase 7 (matrilysin, uterine) 0.0004257983501875646
GO:0042127: regulation of cell proliferation RPA3 replication protein A3, 14kDa 0.003138171179304925
GO:0042127: regulation of cell proliferation SHH sonic hedgehog 0.0006002971515527103
GO:0042127: regulation of cell proliferation SIRT1 sirtuin 1 -1.5183824789954705e-6
GO:0042127: regulation of cell proliferation SIX3 SIX homeobox 3 0.002038510990589221
GO:0042127: regulation of cell proliferation SOX9 SRY (sex determining region Y)-box 9 -0.0005185173871042271
GO:0042127: regulation of cell proliferation TFAP2C transcription factor AP-2 gamma (activating enhancer binding protein 2 gamma) 0.0010611623802875966
GO:0042127: regulation of cell proliferation TGFB3 transforming growth factor, beta 3 -0.0018249526898884544
GO:0042311: vasodilation AGT angiotensinogen (serpin peptidase inhibitor, clade A, member 8) -0.0011265381176853435
GO:0042311: vasodilation GPX1 glutathione peroxidase 1 0.0003857047065419783
GO:0043410: positive regulation of MAPK cascade AGT angiotensinogen (serpin peptidase inhibitor, clade A, member 8) -0.0011327409337338345
GO:0043410: positive regulation of MAPK cascade CDH2 cadherin 2, type 1, N-cadherin (neuronal) -0.0006448969576984282
GO:0043410: positive regulation of MAPK cascade CTNNB1 catenin (cadherin-associated protein), beta 1, 88kDa -0.00011675481917323443
GO:0043410: positive regulation of MAPK cascade FGFR2 fibroblast growth factor receptor 2 0.000762806861838605
GO:0043410: positive regulation of MAPK cascade FGFR3 fibroblast growth factor receptor 3 0.00022251376683993446
GO:0043410: positive regulation of MAPK cascade FLT3 fms-related tyrosine kinase 3 -0.0006811615422099966
GO:0043410: positive regulation of MAPK cascade HMGB1 high mobility group box 1 -0.0007756731930923221
GO:0043410: positive regulation of MAPK cascade IGF1 insulin-like growth factor 1 (somatomedin C) 0.0001309116690577524 2
GO:0043410: positive regulation of MAPK cascade IGF1R insulin-like growth factor 1 receptor 0.0010710913076569983
GO:0043410: positive regulation of MAPK cascade IGFBP3 insulin-like growth factor binding protein 3 0.0008360134930686309
GO:0043410: positive regulation of MAPK cascade IGFBP4 insulin-like growth factor binding protein 4 -0.001277635403252776
GO:0043410: positive regulation of MAPK cascade INSR insulin receptor -0.001362366543641768
GO:0043410: positive regulation of MAPK cascade KIT v-kit Hardy-Zuckerman 4 feline sarcoma viral oncogene homolog 0.000274918950177969 8
GO:0043410: positive regulation of MAPK cascade LEP leptin 0.003188983622543391
GO:0043524: negative regulation of neuron apoptotic process AGT angiotensinogen (serpin peptidase inhibitor, clade A, member 8) -0.0011332376252700877
GO:0043524: negative regulation of neuron apoptotic process ANGPT1 angiopoietin 1 0.000900635825464509

GO:0043524: negative regulation of neuron apoptotic process BAX BCL2-associated X protein
-0.0004251634550346603
GO:0043524: negative regulation of neuron apoptotic process BCL2 B-cell CLL/lymphoma 2 -4.95
3281541930809e-6
GO:0043524: negative regulation of neuron apoptotic process CCL2 chemokine (C-C motif) ligand
2 0.00081653700332012
GO:0043524: negative regulation of neuron apoptotic process CEBPB CCAAT/enhancer binding protei
n (C/EBP), beta -0.0002740039177823731
GO:0043524: negative regulation of neuron apoptotic process CITED1 Cbp/p300-interacting transact
ivator, with Glu/Asp-rich carboxy-terminal domain, 1 0.002818100246927244З
GO:0043524: negative regulation of neuron apoptotic process FGF8 fibroblast growth factor 8 (a
ndrogen-induced) 0.0009839431073587384
GO:0043524: negative regulation of neuron apoptotic process GCLM glutamate-cysteine ligase, mo
difier subunit -0.003761309374100509З
GO:0043524: negative regulation of neuron apoptotic process GDNF glial cell derived neurotroph
ic factor 0.00044471905434896906
GO:0043524: negative regulation of neuron apoptotic process HIPK2 homeodomain interacting prote
in kinase 2 0.0007707519811739051
GO:0043524: negative regulation of neuron apoptotic process HMOX1 heme oxygenase (decycling) 1
-0.0002176040508603668
GO:0043524: negative regulation of neuron apoptotic process HTT huntingtin -0.0009715676
832402418
GO:0043524: negative regulation of neuron apoptotic process ISL1 ISL LIM homeobox 1 7.892
673650974066e-5
GO:0043524: negative regulation of neuron apoptotic process JAK2 Janus kinase 2 -3.1710086875
264245e-5
GO:0043524: negative regulation of neuron apoptotic process KIF14 kinesin family member 14
0.00048599383726066
GO:0043524: negative regulation of neuron apoptotic process MDK midkine (neurite growth-promo
ting factor 2) 0.0016652841063607665
GO:0043524: negative regulation of neuron apoptotic process MEF2C myocyte enhancer factor 2C
0.0009733500044987472
GO:0043524: negative regulation of neuron apoptotic process MSH2 mutS homolog 2 0.00139227906
1904727
GO:0043524: negative regulation of neuron apoptotic process NES nestin 0.0013735926438886218
GO:0043524: negative regulation of neuron apoptotic process NRP1 neuropilin 1 -0.0006432072
180678693
GO:0043524: negative regulation of neuron apoptotic process RASA1 RAS p21 protein activator (GT
Pase activating protein) 1 -0.0003490480491911307
GO:0043524: negative regulation of neuron apoptotic process SIX1 SIX homeobox 1 -0.0018698843
56182417
GO:0043524: negative regulation of neuron apoptotic process STAMBP STAM binding protein -0.00
017040826720829692
GO:0043524: negative regulation of neuron apoptotic process STAR steroidogenic acute regulator
y protein 0.0008945065528124635
GO:0043524: negative regulation of neuron apoptotic process TGFB3 transforming growth factor, b
eta 3 -0.001823656656031718
GO:0043524: negative regulation of neuron apoptotic process TP73 tumor protein p73 0.001
0278837325965117
GO:0043524: negative regulation of neuron apoptotic process WFS1 Wolfram syndrome 1 (wolframi
n) 0.000576775225110592
GO:0044267: cellular protein metabolic process AGT angiotensinogen (serpin peptidase inhibitor,
clade A, member 8) -0.001129864686643366
GO:0044267: cellular protein metabolic process BCHE butyrylcholinesterase -6.363623417655952e-6
GO:0044267: cellular protein metabolic process BLM Bloom syndrome, RecQ helicase-like 0.000
5233192652441638
GO:0044267: cellular protein metabolic process CCL2 chemokine (C-C motif) ligand 2 0.00081509745
70626104
GO:0044267: cellular protein metabolic process CDKN2A cyclin-dependent kinase inhibitor 2A 0.001
7540359393327823
GO:0044267: cellular protein metabolic process CPE carboxypeptidase E -0.002539351420838333
GO:0044267: cellular protein metabolic process CUL7 cullin 7 -0.00011849200300347696
GO:0044267: cellular protein metabolic process EIF4G1 eukaryotic translation initiation factor 4 ga
mma, 1 0.0007410971874029054
GO:0044267: cellular protein metabolic process GALNT6 polypeptide N-acetylgalactosaminyltransferase
6 -0.001771917485097236
GO:0044267: cellular protein metabolic process GCNT1 glucosaminyl (N-acetyl) transferase 1, core 2
0.0006921481954517551
GO:0044267: cellular protein metabolic process GCNT3 glucosaminyl (N-acetyl) transferase 3, mucin
type -0.0023071710945766865
GO:0044267: cellular protein metabolic process GCNT4 glucosaminyl (N-acetyl) transferase 4, core 2
0.00045128242337401706
GO:0044267: cellular protein metabolic process IGF1 insulin-like growth factor 1 (somatomedin C)
0.00012999108687664667
GO:0044267: cellular protein metabolic process IGFBP1 insulin-like growth factor binding protein 1
0.00038737283713590815
GO:0044267: cellular protein metabolic process IGFBP2 insulin-like growth factor binding protein 2,
36kDa 0.0001441149318450229
GO:0044267: cellular protein metabolic process IGFBP3 insulin-like growth factor binding protein 3
0.0008356939504056861
GO:0044267: cellular protein metabolic process IGFBP4 insulin-like growth factor binding protein 4

-0.001275839278567085
GO:0044267: cellular protein metabolic process  IGFBP6  insulin-like growth factor binding protein 6
-0.0020241788444113123
GO:0044267: cellular protein metabolic process  MMP2    matrix metallopeptidase 2 (gelatinase A, 72kD
a gelatinase, 72kDa type IV collagenase)        -0.0011283739831991324
GO:0044267: cellular protein metabolic process  MUC1    mucin 1, cell surface associated       0.001
2824971077652508
GO:0044267: cellular protein metabolic process  MUC7    mucin 7, secreted       -0.00315468030030830
4
GO:0044267: cellular protein metabolic process  NUP153  nucleoporin 153kDa      0.000823156837922047
GO:0044267: cellular protein metabolic process  PCSK2   proprotein convertase subtilisin/kexin type 2
0.0006041548261440689
GO:0044267: cellular protein metabolic process  PGM3    phosphoglucomutase 3    0.00198326723008385
GO:0044267: cellular protein metabolic process  PML     promyelocytic leukemia  -0.000679006418601918
3
GO:0044267: cellular protein metabolic process  RAD21   RAD21 homolog (S. pombe)        -0.0001020872
2922996512
GO:0044267: cellular protein metabolic process  RPA1    replication protein A1, 70kDa   0.00043766112
91702859
GO:0044267: cellular protein metabolic process  SEH1L   SEH1-like (S. cerevisiae)       -0.0007311997
696591838
GO:0044267: cellular protein metabolic process  SMC1A   structural maintenance of chromosomes 1A
-0.0010182806749436466
GO:0044267: cellular protein metabolic process  SMC5    structural maintenance of chromosomes 5 -0.00
0701326191098389
GO:0044267: cellular protein metabolic process  SMC6    structural maintenance of chromosomes 6 -0.00
17478535236534368
GO:0044267: cellular protein metabolic process  SPHK1   sphingosine kinase 1    0.001807209072782821
GO:0044267: cellular protein metabolic process  THBS1   thrombospondin 1        -0.00103381994620243
2
GO:0044267: cellular protein metabolic process  TPR     translocated promoter region, nuclear basket
 protein        -0.0006956505200481419
GO:0044267: cellular protein metabolic process  WFS1    Wolfram syndrome 1 (wolframin)  0.00057600371
62688468
GO:0044267: cellular protein metabolic process  XBP1    X-box binding protein 1 0.0002666677820414090
6
GO:0045429: positive regulation of nitric oxide biosynthetic process    AGT     angiotensinogen (serp
in peptidase inhibitor, clade A, member 8)      -0.001114094899517808
GO:0045429: positive regulation of nitric oxide biosynthetic process    AKT1    v-akt murine thymoma
 viral oncogene homolog 1       0.0007201755139836967
GO:0045429: positive regulation of nitric oxide biosynthetic process    AKT2    v-akt murine thymoma
 viral oncogene homolog 2       -0.000618964829837747
GO:0045429: positive regulation of nitric oxide biosynthetic process    ASS1    argininosuccinate syn
thase 1 0.00013797008423045544
GO:0045429: positive regulation of nitric oxide biosynthetic process    EGFR    epidermal growth fact
or receptor     0.0006742966701912763
GO:0045429: positive regulation of nitric oxide biosynthetic process    ESR1    estrogen receptor 1
-0.0009346559442157988
GO:0045429: positive regulation of nitric oxide biosynthetic process    ICAM1   intercellular adhesio
n molecule 1    0.0007229834608800637
GO:0045429: positive regulation of nitric oxide biosynthetic process    IFNG    interferon, gamma
-4.8715390411869144e-5
GO:0045429: positive regulation of nitric oxide biosynthetic process    INSR    insulin receptor
-0.0013447000675839108
GO:0045429: positive regulation of nitric oxide biosynthetic process    JAK2    Janus kinase 2  -3.36
32384777810186e-5
GO:0045723: positive regulation of fatty acid biosynthetic process      AGT     angiotensinogen (serp
in peptidase inhibitor, clade A, member 8)      -0.0011295254163965211
GO:0045723: positive regulation of fatty acid biosynthetic process      NR1H3   nuclear receptor subf
amily 1, group H, member 3      0.0007982188691035322
GO:0046622: positive regulation of organ growth AGT     angiotensinogen (serpin peptidase inhibitor,
 clade A, member 8)     -0.0011348383041610303
GO:0046622: positive regulation of organ growth FGF8    fibroblast growth factor 8 (androgen-induced)
0.0009848521483020668
GO:0046622: positive regulation of organ growth IL7     interleukin 7   0.0008721384041544568
GO:0046622: positive regulation of organ growth SMO     smoothened, frizzled class receptor     0.002
1405929497700935
GO:0046622: positive regulation of organ growth YAP1    Yes-associated protein 1        -0.0001642044
5902064844
GO:0048143: astrocyte activation        AGT     angiotensinogen (serpin peptidase inhibitor, clade A,
member 8)       -0.0011464001484450565
GO:0048143: astrocyte activation        SMO     smoothened, frizzled class receptor     0.00215561688
5929775
GO:0048144: fibroblast proliferation    AGT     angiotensinogen (serpin peptidase inhibitor, clade A,
member 8)       -0.00114132446189895
GO:0048169: regulation of long-term neuronal synaptic plasticity        AGT     angiotensinogen (serp
in peptidase inhibitor, clade A, member 8)      -0.001124766421243505
GO:0048169: regulation of long-term neuronal synaptic plasticity        DRD2    dopamine receptor D2
-0.00023294888617112795
GO:0048169: regulation of long-term neuronal synaptic plasticity        EGR1    early growth response
1       0.0010866701092259233

GO:0048659: smooth muscle cell proliferation    AGT     angiotensinogen (serpin peptidase inhibitor, clade A, member 8)     -0.001141324461898953
GO:0050663: cytokine secretion  AGT     angiotensinogen (serpin peptidase inhibitor, clade A, member 8)     -0.0011379983167102844
GO:0050663: cytokine secretion  LYN     LYN proto-oncogene, Src family tyrosine kinase  -0.0014235085791756516
GO:0050731: positive regulation of peptidyl-tyrosine phosphorylation    AGT     angiotensinogen (serpin peptidase inhibitor, clade A, member 8)      -0.0011283689576810588
GO:0050731: positive regulation of peptidyl-tyrosine phosphorylation    ANGPT1  angiopoietin 1  0.0008983233655464458
GO:0050731: positive regulation of peptidyl-tyrosine phosphorylation    EFNA1   ephrin-A1       -0.000519714684684735
GO:0050731: positive regulation of peptidyl-tyrosine phosphorylation    FGF7    fibroblast growth factor 7   0.0006320722858868139
GO:0050731: positive regulation of peptidyl-tyrosine phosphorylation    ICAM1   intercellular adhesion molecule 1    0.0007278150678050749
GO:0050731: positive regulation of peptidyl-tyrosine phosphorylation    IGF1    insulin-like growth factor 1 (somatomedin C) 0.00013018510362595778
GO:0050731: positive regulation of peptidyl-tyrosine phosphorylation    JAK2    Janus kinase 2  -3.246110063992826e-5
GO:0050731: positive regulation of peptidyl-tyrosine phosphorylation    LRP8    low density lipoprotein receptor-related protein 8, apolipoprotein e receptor        -0.0009298787235284279
GO:0050731: positive regulation of peptidyl-tyrosine phosphorylation    NRP1    neuropilin 1    -0.0006410515846548176
GO:0050731: positive regulation of peptidyl-tyrosine phosphorylation    RELN    reelin  0.0015320091530040385
GO:0050731: positive regulation of peptidyl-tyrosine phosphorylation    TGFB1   transforming growth factor, beta 1   -7.247029046501873e-5
GO:0050731: positive regulation of peptidyl-tyrosine phosphorylation    THBS4   thrombospondin 4        -0.00044362526692384274
GO:0050731: positive regulation of peptidyl-tyrosine phosphorylation    TP53    tumor protein p53       0.0011712953670444946
GO:0050731: positive regulation of peptidyl-tyrosine phosphorylation    VEGFA   vascular endothelial growth factor A        0.0005921379831008293
GO:0050731: positive regulation of peptidyl-tyrosine phosphorylation    VEGFC   vascular endothelial growth factor C        -0.0033543365913086013
GO:0051145: smooth muscle cell differentiation  AGT     angiotensinogen (serpin peptidase inhibitor, clade A, member 8)     -0.0011385839497855875
GO:0051145: smooth muscle cell differentiation  CTNNB1  catenin (cadherin-associated protein), beta 1, 88kDa       -0.0001166925282289397
GO:0051145: smooth muscle cell differentiation  GATA6   GATA binding protein 6  -2.74480381997768e-5
GO:0051145: smooth muscle cell differentiation  HEY2    hes-related family bHLH transcription factor with YRPW motif 2      0.0027079462603562822
GO:0051145: smooth muscle cell differentiation  MEF2C   myocyte enhancer factor 2C      0.000976600518996167
GO:0051145: smooth muscle cell differentiation  WNT4    wingless-type MMTV integration site family, member 4 -0.00025416908849661737
GO:0051387: negative regulation of neurotrophin TRK receptor signaling pathway  AGT     angiotensinogen (serpin peptidase inhibitor, clade A, member 8)      -0.001141324461898953
GO:0051403: stress-activated MAPK cascade       AGT     angiotensinogen (serpin peptidase inhibitor, clade A, member 8)     -0.0011381000621126763
GO:0051403: stress-activated MAPK cascade       CREB1   cAMP responsive element binding protein 1       0.0006612426418666482
GO:0051403: stress-activated MAPK cascade       CRYAB   crystallin, alpha B     0.0009881431402928252
GO:0051403: stress-activated MAPK cascade       IRAK1   interleukin-1 receptor-associated kinase 1      -0.0016138378058296805
GO:0051403: stress-activated MAPK cascade       MAP3K19 mitogen-activated protein kinase kinase kinase 19    -0.0016542761039534373
GO:0051403: stress-activated MAPK cascade       MEF2C   myocyte enhancer factor 2C      0.0009767451355542315
GO:0051403: stress-activated MAPK cascade       RPS6KA1 ribosomal protein S6 kinase, 90kDa, polypeptide 1    -0.0025209549577330007
GO:0051924: regulation of calcium ion transport AGT     angiotensinogen (serpin peptidase inhibitor, clade A, member 8)     -0.0011194082351256931
GO:0051924: regulation of calcium ion transport BCL2    B-cell CLL/lymphoma 2   -5.88124121249246e-6
GO:0051924: regulation of calcium ion transport GJA1    gap junction protein, alpha 1, 43kDa    -0.0015850971419528766
GO:0051969: regulation of transmission of nerve impulse AGT     angiotensinogen (serpin peptidase inhibitor, clade A, member 8)      -0.001141324461898953
GO:0061049: cell growth involved in cardiac muscle cell development     AGT     angiotensinogen (serpin peptidase inhibitor, clade A, member 8)      -0.0011356652797173752
GO:0061049: cell growth involved in cardiac muscle cell development     GATA4   GATA binding protein 4      -0.0010955738267244865
GO:0061098: positive regulation of protein tyrosine kinase activity     AGT     angiotensinogen (serpin peptidase inhibitor, clade A, member 8)      -0.0011336556975768699
GO:0061098: positive regulation of protein tyrosine kinase activity     CD24    CD24 molecule   0.001071639702462681
GO:0061098: positive regulation of protein tyrosine kinase activity     GREM1   gremlin 1, DAN family BMP antagonist  -0.00082775270678709
GO:0061098: positive regulation of protein tyrosine kinase activity     LRP8    low density lipoprotein receptor-related protein 8, apolipoprotein e receptor        -0.0009308798552369751

GO:0061098: positive regulation of protein tyrosine kinase activity     PTPN1   protein tyrosine phosphatase, non-receptor type 1    -0.0010418241030069751
GO:0061098: positive regulation of protein tyrosine kinase activity     RELN    reelin  0.0015369846024346164
GO:0070371: ERK1 and ERK2 cascade       AGT     angiotensinogen (serpin peptidase inhibitor, clade A, member 8)       -0.001124565031875697
GO:0070371: ERK1 and ERK2 cascade       AVP     arginine vasopressin    -0.0009309082047241992
GO:0070371: ERK1 and ERK2 cascade       IGF1    insulin-like growth factor 1 (somatomedin C)    0.00012932958021281903
GO:0070371: ERK1 and ERK2 cascade       MED1    mediator complex subunit 1      0.001121218119224792
GO:0070371: ERK1 and ERK2 cascade       SOX9    SRY (sex determining region Y)-box 9    -0.000515046260078393
GO:0070471: uterine smooth muscle contraction   AGT     angiotensinogen (serpin peptidase inhibitor, clade A, member 8)     -0.001141324461898953
GO:0071260: cellular response to mechanical stimulus    AGT     angiotensinogen (serpin peptidase inhibitor, clade A, member 8)      -0.0011308490019734615
GO:0071260: cellular response to mechanical stimulus    AKT1    v-akt murine thymoma viral oncogene homolog 1        0.0007290802436596671
GO:0071260: cellular response to mechanical stimulus    BAK1    BCL2-antagonist/killer 1        -0.0018722361088507142
GO:0071260: cellular response to mechanical stimulus    BMP4    bone morphogenetic protein 4    -0.0003220266518879278
GO:0071260: cellular response to mechanical stimulus    BNIP3   BCL2/adenovirus E1B 19kDa interacting protein 3       0.0029070189298056157
GO:0071260: cellular response to mechanical stimulus    CASP8AP2        caspase 8 associated protein 2      0.0012695237955783517
GO:0071260: cellular response to mechanical stimulus    CAV1    caveolin 1, caveolae protein, 22kDa -0.0005345261633046917
GO:0071260: cellular response to mechanical stimulus    CHEK1   checkpoint kinase 1     0.0008983420885444519
GO:0071260: cellular response to mechanical stimulus    COL1A1  collagen, type I, alpha 1       -0.0005252066452635591
GO:0071260: cellular response to mechanical stimulus    EGR1    early growth response 1 0.0010909384885159697
GO:0071260: cellular response to mechanical stimulus    FAS     Fas cell surface death receptor -3.315298479412203e-5
GO:0071260: cellular response to mechanical stimulus    GJA1    gap junction protein, alpha 1, 43kDa -0.00016123313902589019
GO:0071260: cellular response to mechanical stimulus    KCNJ2   potassium inwardly-rectifying channel, subfamily J, member 2        -0.0007752898729378253
GO:0071260: cellular response to mechanical stimulus    MMP7    matrix metallopeptidase 7 (matrilysin, uterine)     0.0004249327201853575
GO:0071260: cellular response to mechanical stimulus    SOX9    SRY (sex determining region Y)-box 9 -0.0005172766424310748
GO:0090190: positive regulation of branching involved in ureteric bud morphogenesis     AGT     angiotensinogen (serpin peptidase inhibitor, clade A, member 8)      -0.0011358793941080557
GO:0090190: positive regulation of branching involved in ureteric bud morphogenesis     GDNF    glial cell derived neurotrophic factor        0.00044512131423449333
GO:0090190: positive regulation of branching involved in ureteric bud morphogenesis     GREM1   gremlin 1, DAN family BMP antagonist -0.0008294600210623033
GO:0090190: positive regulation of branching involved in ureteric bud morphogenesis     LGR4    leucine-rich repeat containing G protein-coupled receptor 4  0.0003088071821391137
GO:0090190: positive regulation of branching involved in ureteric bud morphogenesis     LHX1    LIM homeobox 1       -0.0007599456780626498
GO:0090190: positive regulation of branching involved in ureteric bud morphogenesis     PAX2    paired box 2 -0.001616570649938755
GO:0090190: positive regulation of branching involved in ureteric bud morphogenesis     PAX8    paired box 8 0.0009488374160705636
GO:0090190: positive regulation of branching involved in ureteric bud morphogenesis     SALL1   spalt-like transcription factor 1    -0.002442120875766333
GO:0090190: positive regulation of branching involved in ureteric bud morphogenesis     SIX1    SIX homeobox 1       -0.001872528142639171
GO:0090190: positive regulation of branching involved in ureteric bud morphogenesis     SMO     smoothened, frizzled class receptor  0.0021421752555515613
GO:0090190: positive regulation of branching involved in ureteric bud morphogenesis     SOX9    SRY (sex determining region Y)-box 9       -0.0005194276940881328
GO:0090190: positive regulation of branching involved in ureteric bud morphogenesis     TGFB1   transforming growth factor, beta 1   -7.32421990089243e-5
GO:0090190: positive regulation of branching involved in ureteric bud morphogenesis     VEGFA   vascular endothelial growth factor A 0.0005962983015804185
GO:1900020: positive regulation of protein kinase C activity    AGT     angiotensinogen (serpin peptidase inhibitor, clade A, member 8)      -0.0011306837922656992
GO:1900020: positive regulation of protein kinase C activity    WNT5A   wingless-type MMTV integration site family, member 5A        -0.0006657721693316113
GO:1901201: regulation of extracellular matrix assembly AGT     angiotensinogen (serpin peptidase inhibitor, clade A, member 8)      -0.00112896000986938
GO:1901201: regulation of extracellular matrix assembly NOTCH1  notch 1 0.0005170083157855369
GO:1902632: positive regulation of membrane hyperpolarization   AGT     angiotensinogen (serpin peptidase inhibitor, clade A, member 8)      -0.001141324461898953
GO:1903598: positive regulation of gap junction assembly        AGT     angiotensinogen (serpin peptidase inhibitor, clade A, member 8)      -0.0011274210960150427

GO:1903598: positive regulation of gap junction assembly	CAV1	caveolin 1, caveolae protein, 22kDa	-0.0005330003027876892
GO:1903779: regulation of cardiac conduction	AGT	angiotensinogen (serpin peptidase inhibitor, clade A, member 8)	-0.00114132446189895 3
GO:2000650: negative regulation of sodium ion transmembrane transporter activity	AGT	angiotensinogen (serpin peptidase inhibitor, clade A, member 8)	-0.0011440163609418425
GO:2000650: negative regulation of sodium ion transmembrane transporter activity	NEDD4	neural precursor cell expressed, developmentally down-regulated 4, E3 ubiquitin protein ligase	0.0022756847164011945
GO:2001238: positive regulation of extrinsic apoptotic signaling pathway	AGT	angiotensinogen (serpin peptidase inhibitor, clade A, member 8)	-0.001118859106551077
GO:2001238: positive regulation of extrinsic apoptotic signaling pathway	CAV1	caveolin 1, caveolae protein, 22kDa	-0.0005313378843129165
GO:2001238: positive regulation of extrinsic apoptotic signaling pathway	PML	promyelocytic leukemia	-0.0006749960214677706
KEGG:04080: Neuroactive ligand-receptor interaction	AGTR1	angiotensin II receptor, type 1	0.00035059797124114054
KEGG:04080: Neuroactive ligand-receptor interaction	CHRNB1	cholinergic receptor, nicotinic, beta 1 (muscle)	-7.4441822803154206e-6
KEGG:04080: Neuroactive ligand-receptor interaction	CHRNB2	cholinergic receptor, nicotinic, beta 2 (neuronal)	-0.0007775952737220379
KEGG:04080: Neuroactive ligand-receptor interaction	CSH1	chorionic somatomammotropin hormone 1 (placental lactogen)	-0.0002574152848051913
KEGG:04080: Neuroactive ligand-receptor interaction	DRD2	dopamine receptor D2	-0.0002336862061888899
KEGG:04080: Neuroactive ligand-receptor interaction	DRD4	dopamine receptor D4	-0.0019435849857920122
KEGG:04080: Neuroactive ligand-receptor interaction	GABRA4	gamma-aminobutyric acid (GABA) A receptor, alpha 4	-0.0014826296661433036
KEGG:04080: Neuroactive ligand-receptor interaction	GNRHR	gonadotropin-releasing hormone receptor	0.0007407863776446287
KEGG:04080: Neuroactive ligand-receptor interaction	GRIK5	glutamate receptor, ionotropic, kainate 5	-0.0002329090563643155
KEGG:04080: Neuroactive ligand-receptor interaction	HTR2C	5-hydroxytryptamine (serotonin) receptor 2C, G protein-coupled	-0.0005246970516044354
KEGG:04080: Neuroactive ligand-receptor interaction	HTR6	5-hydroxytryptamine (serotonin) receptor 6, G protein-coupled	-0.0010813633042108218
KEGG:04080: Neuroactive ligand-receptor interaction	LEP	leptin	0.0031865997634206895
KEGG:04080: Neuroactive ligand-receptor interaction	MAS1	MAS1 proto-oncogene, G protein-coupled receptor	0.00029318047410087497
KEGG:04080: Neuroactive ligand-receptor interaction	PRLR	prolactin receptor	0.0021826627020479807
KEGG:04080: Neuroactive ligand-receptor interaction	THRA	thyroid hormone receptor, alpha	0.0007633707382364811
KEGG:04080: Neuroactive ligand-receptor interaction	THRB	thyroid hormone receptor, beta	0.001943070177755303
KEGG:04270: Vascular smooth muscle contraction	AGTR1	angiotensin II receptor, type 1	0.0003477717624472632
KEGG:04270: Vascular smooth muscle contraction	ITPR1	inositol 1,4,5-trisphosphate receptor, type 1	-0.0007726285830895634
KEGG:04270: Vascular smooth muscle contraction	KCNMA1	potassium large conductance calcium-activated channel, subfamily M, alpha member 1	-0.0006502154596969323
KEGG:04270: Vascular smooth muscle contraction	PLA2G5	phospholipase A2, group V	0.002063440001743988
KEGG:04270: Vascular smooth muscle contraction	PLCB1	phospholipase C, beta 1 (phosphoinositide-specific)	0.0001507684537455336
KEGG:04270: Vascular smooth muscle contraction	PRKCA	protein kinase C, alpha	-5.907479652625633e-6
KEGG:04270: Vascular smooth muscle contraction	PRKCD	protein kinase C, delta	-0.0011175627256523007
KEGG:04270: Vascular smooth muscle contraction	PRKX	protein kinase, X-linked	0.00038938186318328
KEGG:04270: Vascular smooth muscle contraction	RAF1	Raf-1 proto-oncogene, serine/threonine kinase	0.001486784599444832
KEGG:04270: Vascular smooth muscle contraction	RHOA	ras homolog family member A	0.0006644111183773569
KEGG:05200: Pathways in cancer	AGTR1	angiotensin II receptor, type 1	0.0003475353240859817
KEGG:05200: Pathways in cancer	AKT1	v-akt murine thymoma viral oncogene homolog 1	0.0007292479628088177
KEGG:05200: Pathways in cancer	AKT2	v-akt murine thymoma viral oncogene homolog 2	-0.0006249550526819738
KEGG:05200: Pathways in cancer	APC	adenomatous polyposis coli	0.0006430765155879746
KEGG:05200: Pathways in cancer	AR	androgen receptor	0.002634242820837509
KEGG:05200: Pathways in cancer	AXIN1	axin 1	-0.0007317074748467242
KEGG:05200: Pathways in cancer	BAX	BCL2-associated X protein	-0.0004245212723179918
KEGG:05200: Pathways in cancer	BCL2	B-cell CLL/lymphoma 2	-5.1169649692408316e-6
KEGG:05200: Pathways in cancer	BIRC5	baculoviral IAP repeat containing 5	-0.0002601839706338838
KEGG:05200: Pathways in cancer	BMP4	bone morphogenetic protein 4	-0.00032207511799027863
KEGG:05200: Pathways in cancer	BRCA2	breast cancer 2, early onset	-1.017752536150188e-5
KEGG:05200: Pathways in cancer	CCNA1	cyclin A1	-0.0017566511546665902
KEGG:05200: Pathways in cancer	CCND1	cyclin D1	-0.002627716223086891 4

KEGG:05200: Pathways in cancer  CCNE1   cyclin E1       0.0003702560262258883
KEGG:05200: Pathways in cancer  CCNE2   cyclin E2       0.0011049610661534753
KEGG:05200: Pathways in cancer  CDC42   cell division cycle 42  0.0012381436869618636
KEGG:05200: Pathways in cancer  CDKN2A  cyclin-dependent kinase inhibitor 2A    0.0017548340934393533
KEGG:05200: Pathways in cancer  CDKN2B  cyclin-dependent kinase inhibitor 2B (p15, inhibits CDK4)
-0.0018324229590384973
KEGG:05200: Pathways in cancer  CKS2    CDC28 protein kinase regulatory subunit 2       -4.8289909527
43842e-5
KEGG:05200: Pathways in cancer  CSF3R   colony stimulating factor 3 receptor (granulocyte)      -8.28
503106724201e-5
KEGG:05200: Pathways in cancer  CTNNB1  catenin (cadherin-associated protein), beta 1, 88kDa    -0.00
01168028193652878
KEGG:05200: Pathways in cancer  CXCL12  chemokine (C-X-C motif) ligand 12       -0.001206308219044905
2
KEGG:05200: Pathways in cancer  CXCR4   chemokine (C-X-C motif) receptor 4      0.0007932901940236525
KEGG:05200: Pathways in cancer  E2F1    E2F transcription factor 1      0.002094094751964618
KEGG:05200: Pathways in cancer  E2F3    E2F transcription factor 3      0.0021585216052612214
KEGG:05200: Pathways in cancer  EGFR    epidermal growth factor receptor        0.00068430548878953
KEGG:05200: Pathways in cancer  FAS     Fas cell surface death receptor -3.3187967708277944e-5
KEGG:05200: Pathways in cancer  FGF3    fibroblast growth factor 3      0.001558362316629871
KEGG:05200: Pathways in cancer  FGF5    fibroblast growth factor 5      -0.0010922603319737502
KEGG:05200: Pathways in cancer  FGF7    fibroblast growth factor 7      0.0006326358006618654
KEGG:05200: Pathways in cancer  FGF8    fibroblast growth factor 8 (androgen-induced)   0.00098205648
2524673
KEGG:05200: Pathways in cancer  FGFR2   fibroblast growth factor receptor 2     0.0007618709844360252
KEGG:05200: Pathways in cancer  FGFR3   fibroblast growth factor receptor 3     0.0002219821041618055
5
KEGG:05200: Pathways in cancer  FLT3    fms-related tyrosine kinase 3   -0.0006801293417308778
KEGG:05200: Pathways in cancer  FOXO1   forkhead box O1 0.001797241350210865
KEGG:05200: Pathways in cancer  FZD3    frizzled class receptor 3       0.00037754096218346343
KEGG:05200: Pathways in cancer  FZD7    frizzled class receptor 7       0.0010684524267841444
KEGG:05200: Pathways in cancer  GLI1    GLI family zinc finger 1        -0.001294887557244521
KEGG:05200: Pathways in cancer  GLI2    GLI family zinc finger 2        0.0018503085215558054
KEGG:05200: Pathways in cancer  GLI3    GLI family zinc finger 3        -0.002147785799533418
KEGG:05200: Pathways in cancer  GRB2    growth factor receptor-bound protein 2  0.0004289053722929083
KEGG:05200: Pathways in cancer  GSK3B   glycogen synthase kinase 3 beta 0.0015464803123016698
KEGG:05200: Pathways in cancer  GSTP1   glutathione S-transferase pi 1  0.0002311562579136978
KEGG:05200: Pathways in cancer  HDAC2   histone deacetylase 2   -0.0012055335931622567
KEGG:05200: Pathways in cancer  HIF1A   hypoxia inducible factor 1, alpha subunit (basic helix-loop-h
elix transcription factor)      -0.0006455051398532944
KEGG:05200: Pathways in cancer  IGF1    insulin-like growth factor 1 (somatomedin C)    0.00013034899
531345472
KEGG:05200: Pathways in cancer  IGF1R   insulin-like growth factor 1 receptor   0.0010705675369078605
KEGG:05200: Pathways in cancer  ITGA3   integrin, alpha 3 (antigen CD49C, alpha 3 subunit of VLA-3 re
ceptor) 0.0013975734579825935
KEGG:05200: Pathways in cancer  ITGA6   integrin, alpha 6       0.001665443181661152
KEGG:05200: Pathways in cancer  ITGB1   integrin, beta 1 (fibronectin receptor, beta polypeptide, ant
igen CD29 includes MDF2, MSK12) 0.002422057221873879
KEGG:05200: Pathways in cancer  JUP     junction plakoglobin    -0.001476930825441092
KEGG:05200: Pathways in cancer  KIT     v-kit Hardy-Zuckerman 4 feline sarcoma viral oncogene homolog
0.00027388354277701126
KEGG:05200: Pathways in cancer  LAMB2   laminin, beta 2 (laminin S)     -0.0013066990581547023
KEGG:05200: Pathways in cancer  LEF1    lymphoid enhancer-binding factor 1      -0.000100325958478382
57
KEGG:05200: Pathways in cancer  MLH1    mutL homolog 1  0.002332503326640001
KEGG:05200: Pathways in cancer  MMP2    matrix metallopeptidase 2 (gelatinase A, 72kDa gelatinase, 72
kDa type IV collagenase)        -0.0011290305320316343
KEGG:05200: Pathways in cancer  MSH2    mutS homolog 2  0.0013896006625863002
KEGG:05200: Pathways in cancer  MSH3    mutS homolog 3  -0.0005297582184598427
KEGG:05200: Pathways in cancer  MSH6    mutS homolog 6  0.0010758355738002222
KEGG:05200: Pathways in cancer  MYC     v-myc avian myelocytomatosis viral oncogene homolog     -0.00
11346829922679578
KEGG:05200: Pathways in cancer  PAX8    paired box 8    0.0009469430853739511
KEGG:05200: Pathways in cancer  PDGFRB  platelet-derived growth factor receptor, beta polypeptide
-0.00036020480106274574
KEGG:05200: Pathways in cancer  PIK3CD  phosphatidylinositol-4,5-bisphosphate 3-kinase, catalytic sub
unit delta      -0.0007795017904443303
KEGG:05200: Pathways in cancer  PLCB1   phospholipase C, beta 1 (phosphoinositide-specific)     0.000
1532798270354626
KEGG:05200: Pathways in cancer  PML     promyelocytic leukemia  -0.0006791343357943938
KEGG:05200: Pathways in cancer  PRKCA   protein kinase C, alpha -5.867986599925455e-6
KEGG:05200: Pathways in cancer  PRKX    protein kinase, X-linked        0.0003937998854287266
KEGG:05200: Pathways in cancer  PTCH1   patched 1       -6.599530310629082e-5
KEGG:05200: Pathways in cancer  PTEN    phosphatase and tensin homolog  1.744518293943311e-5
KEGG:05200: Pathways in cancer  RAD51   RAD51 recombinase       -0.0018582796598198695
KEGG:05200: Pathways in cancer  RAF1    Raf-1 proto-oncogene, serine/threonine kinase   0.00149632993
09306973
KEGG:05200: Pathways in cancer  RB1     retinoblastoma 1        -0.0014914358284641807
KEGG:05200: Pathways in cancer  RET     ret proto-oncogene      -0.0004905200765679361
KEGG:05200: Pathways in cancer  RHOA    ras homolog family member A     0.0006686522994744406
KEGG:05200: Pathways in cancer  RXRA    retinoid X receptor, alpha      0.0011132437576497674

KEGG:05200: Pathways in cancer  SHH     sonic hedgehog  0.0005986343010761837
KEGG:05200: Pathways in cancer  SKP2    S-phase kinase-associated protein 2, E3 ubiquitin protein ligase     -0.0005362216081415751
KEGG:05200: Pathways in cancer  SMO     smoothened, frizzled class receptor     0.00213562171191383
KEGG:05200: Pathways in cancer  STAT5A  signal transducer and activator of transcription 5A     0.0015937308840572007
KEGG:05200: Pathways in cancer  TCF7L2  transcription factor 7-like 2 (T-cell specific, HMG-box)     0.0005725618128288142
KEGG:05200: Pathways in cancer  TGFB1   transforming growth factor, beta 1      -7.283372914293476e-5
KEGG:05200: Pathways in cancer  TGFB2   transforming growth factor, beta 2      -0.001057378518212652
KEGG:05200: Pathways in cancer  TGFB3   transforming growth factor, beta 3      -0.001820325400538048 3
KEGG:05200: Pathways in cancer  TGFBR1  transforming growth factor, beta receptor 1     0.0003391849923124031
KEGG:05200: Pathways in cancer  TP53    tumor protein p53       0.0011732916887733887
KEGG:05200: Pathways in cancer  TPR     translocated promoter region, nuclear basket protein    -0.0006959176399215365
KEGG:05200: Pathways in cancer  VEGFA   vascular endothelial growth factor A    0.0005936513709879257
KEGG:05200: Pathways in cancer  VEGFC   vascular endothelial growth factor C    -0.003358503626733516 3
KEGG:05200: Pathways in cancer  WNT1    wingless-type MMTV integration site family, member 1    0.0007856803371034066
KEGG:05200: Pathways in cancer  WNT10B  wingless-type MMTV integration site family, member 10B  -0.001385949165637525
KEGG:05200: Pathways in cancer  WNT4    wingless-type MMTV integration site family, member 4    -0.0025125883013474786
KEGG:05200: Pathways in cancer  WNT5A   wingless-type MMTV integration site family, member 5A   -0.0006662623515509101
KEGG:05200: Pathways in cancer  WNT7A   wingless-type MMTV integration site family, member 7A   1.960930410129098e-5
KEGG:04022: cGMP-PKG signaling pathway  AGTR1   angiotensin II receptor, type 1 0.0003485926894849319 3
KEGG:04022: cGMP-PKG signaling pathway  AKT1    v-akt murine thymoma viral oncogene homolog 1   0.0007225817719238704
KEGG:04022: cGMP-PKG signaling pathway  AKT2    v-akt murine thymoma viral oncogene homolog 2   -0.0006210979171807567
KEGG:04022: cGMP-PKG signaling pathway  ATP1B3  ATPase, Na+/K+ transporting, beta 3 polypeptide -0.0014853337829943084
KEGG:04022: cGMP-PKG signaling pathway  CREB1   cAMP responsive element binding protein 1       0.000648961978164158
KEGG:04022: cGMP-PKG signaling pathway  GATA4   GATA binding protein 4  -0.0010834492017024245
KEGG:04022: cGMP-PKG signaling pathway  INSR    insulin receptor        -0.0013487715813365355
KEGG:04022: cGMP-PKG signaling pathway  ITPR1   inositol 1,4,5-trisphosphate receptor, type 1   -0.0007704561243640074
KEGG:04022: cGMP-PKG signaling pathway  KCNMA1  potassium large conductance calcium-activated channel, subfamily M, alpha member 1  -0.0006479979619600897
KEGG:04022: cGMP-PKG signaling pathway  MEF2C   myocyte enhancer factor 2C      0.0009623584081566344
KEGG:04022: cGMP-PKG signaling pathway  PIK3CD  phosphatidylinositol-4,5-bisphosphate 3-kinase, catalytic subunit delta      -0.0007731994922275433
KEGG:04022: cGMP-PKG signaling pathway  PLCB1   phospholipase C, beta 1 (phosphoinositide-specific)     0.00015005615254737385
KEGG:04022: cGMP-PKG signaling pathway  RAF1    Raf-1 proto-oncogene, serine/threonine kinase   0.0014845086216911884
KEGG:04022: cGMP-PKG signaling pathway  RHOA    ras homolog family member A     0.0006635702649900567
KEGG:04261: Adrenergic signaling in cardiomyocytes      AGTR1   angiotensin II receptor, type 1 0.0003485667211912763
KEGG:04261: Adrenergic signaling in cardiomyocytes      AKT1    v-akt murine thymoma viral oncogene homolog 1        0.0007080662896574126
KEGG:04261: Adrenergic signaling in cardiomyocytes      AKT2    v-akt murine thymoma viral oncogene homolog 2        -0.0006121077292545785
KEGG:04261: Adrenergic signaling in cardiomyocytes      ATP1B3  ATPase, Na+/K+ transporting, beta 3 polypeptide      -0.0014621576289393062
KEGG:04261: Adrenergic signaling in cardiomyocytes      BCL2    B-cell CLL/lymphoma 2   -5.544687975875021e-6
KEGG:04261: Adrenergic signaling in cardiomyocytes      CACNB3  calcium channel, voltage-dependent, beta 3 subunit   0.0002085321789024473
KEGG:04261: Adrenergic signaling in cardiomyocytes      CREB1   cAMP responsive element binding protein 1    0.0006326820546893797
KEGG:04261: Adrenergic signaling in cardiomyocytes      PIK3CD  phosphatidylinositol-4,5-bisphosphate 3-kinase, catalytic subunit delta       -0.0007594704790757687
KEGG:04261: Adrenergic signaling in cardiomyocytes      PLCB1   phospholipase C, beta 1 (phosphoinositide-specific)  0.00014373582046099716
KEGG:04261: Adrenergic signaling in cardiomyocytes      PPP2R3A protein phosphatase 2, regulatory subunit B'', alpha -4.2160544074152406e-5
KEGG:04261: Adrenergic signaling in cardiomyocytes      PRKCA   protein kinase C, alpha -6.153864745555016e-6
KEGG:04261: Adrenergic signaling in cardiomyocytes      PRKX    protein kinase, X-linked        0.0003762661135962695
KEGG:04020: Calcium signaling pathway   AGTR1   angiotensin II receptor, type 1 0.0003466102480739935
KEGG:04020: Calcium signaling pathway   EGFR    epidermal growth factor receptor        0.0006719619679738328

KEGG:04020: Calcium signaling pathway   ERBB4   v-erb-b2 avian erythroblastic leukemia viral oncogene homolog 4       -0.00017706689218756352
KEGG:04020: Calcium signaling pathway   HTR2C   5-hydroxytryptamine (serotonin) receptor 2C, G protein-coupled       -0.0005172444913433183
KEGG:04020: Calcium signaling pathway   HTR6    5-hydroxytryptamine (serotonin) receptor 6, G protein-coupled        -0.0010658225355745846
KEGG:04020: Calcium signaling pathway   ITPR1   inositol 1,4,5-trisphosphate receptor, type 1   -0.0007666911504247498
KEGG:04020: Calcium signaling pathway   ORAI2   ORAI calcium release-activated calcium modulator 2 0.0007224552284418565
KEGG:04020: Calcium signaling pathway   ORAI3   ORAI calcium release-activated calcium modulator 3 -0.0008751577028992492
KEGG:04020: Calcium signaling pathway   PDGFRB  platelet-derived growth factor receptor, beta polypeptide    -0.0003518927572837581
KEGG:04020: Calcium signaling pathway   PLCB1   phospholipase C, beta 1 (phosphoinositide-specific) 0.00014868911896114464
KEGG:04020: Calcium signaling pathway   PRKCA   protein kinase C, alpha -5.844379105646571e-6
KEGG:04020: Calcium signaling pathway   PRKX    protein kinase, X-linked        0.00038599883480925165
KEGG:04020: Calcium signaling pathway   SPHK1   sphingosine kinase 1    0.001783148414863169
GO:0003081: regulation of systemic arterial blood pressure by renin-angiotensin AGTR1   angiotensin II receptor, type 1      0.00034087272072047314
GO:0007266: Rho protein signal transduction     AGTR1   angiotensin II receptor, type 1 0.00034745936097046185
GO:0007266: Rho protein signal transduction     CFL1    cofilin 1 (non-muscle)  -0.0011751921834503032
GO:0007266: Rho protein signal transduction     CNKSR1  connector enhancer of kinase suppressor of Ras 1     -0.0011061531138089678
GO:0007266: Rho protein signal transduction     PAK1    p21 protein (Cdc42/Rac)-activated kinase 1 -0.002187846403390787
GO:0007266: Rho protein signal transduction     PAK3    p21 protein (Cdc42/Rac)-activated kinase 3 -0.0012586585921412232
GO:0007266: Rho protein signal transduction     PHACTR4 phosphatase and actin regulator 4       -0.001790958980079619
GO:0007266: Rho protein signal transduction     RHOA    ras homolog family member A     0.0006697706205168874
GO:0007266: Rho protein signal transduction     ROPN1B  rhophilin associated tail protein 1B    0.0005978015550404622
GO:0019722: calcium-mediated signaling  AGTR1   angiotensin II receptor, type 1 0.0003470865204972391
GO:0019722: calcium-mediated signaling  CXCR4   chemokine (C-X-C motif) receptor 4      0.0007919562854707345
GO:0019722: calcium-mediated signaling  SPHK1   sphingosine kinase 1    0.0018046616123086462
GO:0032430: positive regulation of phospholipase A2 activity    AGTR1   angiotensin II receptor, type 1       0.00034087272072047314
GO:0038166: angiotensin-activated signaling pathway     AGTR1   angiotensin II receptor, type 1 0.0003348582837193237
GO:0038166: angiotensin-activated signaling pathway     MAS1    MAS1 proto-oncogene, G protein-coupled receptor      0.0003017651563656742
GO:0042312: regulation of vasodilation  AGTR1   angiotensin II receptor, type 1 0.00034087272072047314
GO:0050727: regulation of inflammatory response AGTR1   angiotensin II receptor, type 1 0.0003496533503864794
GO:0050727: regulation of inflammatory response BRD4    bromodomain containing 4        0.0005486325697231977
GO:0050727: regulation of inflammatory response JAK2    Janus kinase 2  -3.6790578713862576e-5
GO:0050727: regulation of inflammatory response LYN     LYN proto-oncogene, Src family tyrosine kinase       -0.0013931861534899263
GO:0050727: regulation of inflammatory response MAS1    MAS1 proto-oncogene, G protein-coupled receptor      0.0002963608384649917
GO:0050727: regulation of inflammatory response TNFAIP3 tumor necrosis factor, alpha-induced protein 3      0.0010936282909919455
GO:0051482: positive regulation of cytosolic calcium ion concentration involved in phospholipase C-activating G-protein coupled signaling pathway    AGTR1   angiotensin II receptor, type 1 0.0003495600405393911
GO:0051482: positive regulation of cytosolic calcium ion concentration involved in phospholipase C-activating G-protein coupled signaling pathway    DRD2    dopamine receptor D2    -0.0002216842067900547
GO:0051482: positive regulation of cytosolic calcium ion concentration involved in phospholipase C-activating G-protein coupled signaling pathway    HTR2C   5-hydroxytryptamine (serotonin) receptor 2C, G protein-coupled      -0.00048738072849223907
GO:0060326: cell chemotaxis     AGTR1   angiotensin II receptor, type 1 0.0003469201288968819
GO:0060326: cell chemotaxis     AZU1    azurocidin 1    0.00019664738122811815
GO:0060326: cell chemotaxis     BIN2    bridging integrator 2   -0.0016348003649213076
GO:0060326: cell chemotaxis     CXCL12  chemokine (C-X-C motif) ligand 12       -0.00120147170364253
GO:0060326: cell chemotaxis     ENG     endoglin        0.0008205391086668359
GO:0060326: cell chemotaxis     EPHB1   EPH receptor B1 0.0019525787321272945
GO:0060326: cell chemotaxis     HMGB2   high mobility group box 2       0.0003031835318071074
GO:0060326: cell chemotaxis     KIT     v-kit Hardy-Zuckerman 4 feline sarcoma viral oncogene homolog 0.00027175530725491063
GO:0060326: cell chemotaxis     LEF1    lymphoid enhancer-binding factor 1      -0.00010041531808374979

GO:0060326: cell chemotaxis     PDGFRB  platelet-derived growth factor receptor, beta polypeptide -0.0003579372517893109
GO:0060326: cell chemotaxis     PRKCD   protein kinase C, delta -0.0011208776123495456
GO:0086097: phospholipase C-activating angiotensin-activated signaling pathway  AGTR1   angiotensin II receptor, type 1      0.00034087272072047314
GO:0006418: tRNA aminoacylation for protein translation AIMP2   aminoacyl tRNA synthetase complex-interacting multifunctional protein 2      0.0011874080425433107
GO:0006418: tRNA aminoacylation for protein translation LARS2   leucyl-tRNA synthetase 2, mitochondrial      0.00044953839655748263
GO:0006915: apoptotic process   AIMP2   aminoacyl tRNA synthetase complex-interacting multifunctional protein 2       0.0012090167990556283
GO:0006915: apoptotic process   AKT1    v-akt murine thymoma viral oncogene homolog 1   0.000730274175376861
GO:0006915: apoptotic process   APC     adenomatous polyposis coli      0.0006437735456922245
GO:0006915: apoptotic process   AXIN1   axin 1  -0.0007327393983158619
GO:0006915: apoptotic process   BAK1    BCL2-antagonist/killer 1        -0.001875396163227535
GO:0006915: apoptotic process   BAX     BCL2-associated X protein       -0.00042514058779966475
GO:0006915: apoptotic process   BCL2    B-cell CLL/lymphoma 2   -4.977765350639308e-6
GO:0006915: apoptotic process   BCL2L11 BCL2-like 11 (apoptosis facilitator)    -0.0007746608587026223
GO:0006915: apoptotic process   BFAR    bifunctional apoptosis regulator        -0.0003693625766074068
GO:0006915: apoptotic process   BNIP3   BCL2/adenovirus E1B 19kDa interacting protein 3 0.002911376222599086
GO:0006915: apoptotic process   BUB1    BUB1 mitotic checkpoint serine/threonine kinase 0.0013241946823426413
GO:0006915: apoptotic process   BUB1B   BUB1 mitotic checkpoint serine/threonine kinase B       -0.0012811083549150622
GO:0006915: apoptotic process   CDK1    cyclin-dependent kinase 1       0.00031063400550536885
GO:0006915: apoptotic process   CIB1    calcium and integrin binding 1 (calmyrin)       5.498901725151012e-5
GO:0006915: apoptotic process   CITED1  Cbp/p300-interacting transactivator, with Glu/Asp-rich carboxy-terminal domain, 1    0.0028169464176882207
GO:0006915: apoptotic process   CKAP2   cytoskeleton associated protein 2       4.881591577572256e-5
GO:0006915: apoptotic process   CST3    cystatin C      -6.975577470499743e-5
GO:0006915: apoptotic process   CTNNB1  catenin (cadherin-associated protein), beta 1, 88kDa    -0.0011702062396451214
GO:0006915: apoptotic process   CXCR4   chemokine (C-X-C motif) receptor 4      0.0007942356023770184
GO:0006915: apoptotic process   DAXX    death-domain associated protein 0.0008910378855435982
GO:0006915: apoptotic process   DHCR24  24-dehydrocholesterol reductase -0.0017067530777823052
GO:0006915: apoptotic process   DIABLO  diablo, IAP-binding mitochondrial protein       -0.0005955950874973309
GO:0006915: apoptotic process   DOCK1   dedicator of cytokinesis 1      -0.0006838636074784753
GO:0006915: apoptotic process   DSG2    desmoglein 2    1.668456633629186e-5
GO:0006915: apoptotic process   DSP     desmoplakin     -5.666357773000118e-5
GO:0006915: apoptotic process   E2F1    E2F transcription factor 1      0.0020967558273057435
GO:0006915: apoptotic process   EDAR    ectodysplasin A receptor        0.0005937664990297262
GO:0006915: apoptotic process   ESPL1   extra spindle pole bodies homolog 1 (S. cerevisiae)     0.00019668379559849593
GO:0006915: apoptotic process   FAS     Fas cell surface death receptor -3.326835424946711e-5
GO:0006915: apoptotic process   FEM1B   fem-1 homolog b (C. elegans)    -0.004129437588366252
GO:0006915: apoptotic process   FGF8    fibroblast growth factor 8 (androgen-induced)   0.000983422489568303
GO:0006915: apoptotic process   FGFR2   fibroblast growth factor receptor 2     0.0007628372960817823
GO:0006915: apoptotic process   FOXO1   forkhead box O1 0.0017995590026224568
GO:0006915: apoptotic process   GJA1    gap junction protein, alpha 1, 43kDa    -0.00016112545416969727
GO:0006915: apoptotic process   GLRX2   glutaredoxin 2  0.0011156950979251806
GO:0006915: apoptotic process   GREM1   gremlin 1, DAN family BMP antagonist    -0.0008267059407351256
GO:0006915: apoptotic process   HMGB1   high mobility group box 1       -0.000776293451810886
GO:0006915: apoptotic process   HMGB2   high mobility group box 2       0.00030641277187692343
GO:0006915: apoptotic process   IFNA2   interferon, alpha 2     -0.0017166446112798208
GO:0006915: apoptotic process   IFNG    interferon, gamma       -4.893234267981272e-5
GO:0006915: apoptotic process   IGFBP3  insulin-like growth factor binding protein 3    0.0008366433195386395
GO:0006915: apoptotic process   JAK2    Janus kinase 2  -3.197876945434216e-5
GO:0006915: apoptotic process   KLF11   Kruppel-like factor 11  0.0006160522158830204
GO:0006915: apoptotic process   KMT2A   lysine (K)-specific methyltransferase 2A        0.0006603287159805296
GO:0006915: apoptotic process   KPNB1   karyopherin (importin) beta 1   0.0007716663649921329
GO:0006915: apoptotic process   MAP1S   microtubule-associated protein 1S       0.0005305226458520359
GO:0006915: apoptotic process   MAPT    microtubule-associated protein tau      0.001551211951391722
GO:0006915: apoptotic process   MEF2C   myocyte enhancer factor 2C      0.0009728470760646272
GO:0006915: apoptotic process   MELK    maternal embryonic leucine zipper kinase        0.002068095947441584
GO:0006915: apoptotic process   NOC2L   nucleolar complex associated 2 homolog (S. cerevisiae)  -0.0015109743019154446
GO:0006915: apoptotic process   PAEP    progestagen-associated endometrial protein      0.0010649492454120446
GO:0006915: apoptotic process   PAK1    p21 protein (Cdc42/Rac)-activated kinase 1      -0.0021868631

81408708
GO:0006915: apoptotic process   PAX3    paired box 3    -0.0029442138311813787
GO:0006915: apoptotic process   PLK3    polo-like kinase 3      0.002563436347137538
GO:0006915: apoptotic process   PML     promyelocytic leukemia  -0.0006799470690519058
GO:0006915: apoptotic process   PPP1R13L        protein phosphatase 1, regulatory subunit 13 like
-0.0014731505524553804
GO:0006915: apoptotic process   PRKCD   protein kinase C, delta -0.0011277401215461836
GO:0006915: apoptotic process   PSMA5   proteasome (prosome, macropain) subunit, alpha type, 5  0.000
3317158097982079
GO:0006915: apoptotic process   PSMD11  proteasome (prosome, macropain) 26S subunit, non-ATPase, 11
-0.001094786897076217
GO:0006915: apoptotic process   PSMD13  proteasome (prosome, macropain) 26S subunit, non-ATPase, 13
-5.983052028420809e-5
GO:0006915: apoptotic process   PTEN    phosphatase and tensin homolog  1.7370802439597952e-5
GO:0006915: apoptotic process   RAD21   RAD21 homolog (S. pombe)        -0.00010240308952538412
GO:0006915: apoptotic process   RAF1    Raf-1 proto-oncogene, serine/threonine kinase   0.00149822185
51294764
GO:0006915: apoptotic process   S100A14 S100 calcium binding protein A14        -0.00215858573866971
7
GO:0006915: apoptotic process   SIX1    SIX homeobox 1  -0.0018692210724397707
GO:0006915: apoptotic process   STEAP3  STEAP family member 3, metalloreductase 0.0026082397026490163
GO:0006915: apoptotic process   STK24   serine/threonine kinase 24      0.002693624030796801
GO:0006915: apoptotic process   SULF1   sulfatase 1     -0.0008048270968838142
GO:0006915: apoptotic process   TFDP1   transcription factor Dp-1       0.0014437072999589982
GO:0006915: apoptotic process   TGFBR1  transforming growth factor, beta receptor 1     0.00033965305
776263195
GO:0006915: apoptotic process   TNFAIP3 tumor necrosis factor, alpha-induced protein 3  0.00111607527
93497309
GO:0006915: apoptotic process   TNFRSF10C       tumor necrosis factor receptor superfamily, member 10
c, decoy without an intracellular domain        0.0003172603557685242
GO:0006915: apoptotic process   TNFRSF21        tumor necrosis factor receptor superfamily, member 21
0.00034825319608522437
GO:0006915: apoptotic process   TP53    tumor protein p53       0.0011749295130844954
GO:0006915: apoptotic process   TPX2    TPX2, microtubule-associated    0.0013408903692152202
GO:0006915: apoptotic process   TRAIP   TRAF interacting protein        0.001022035919147696
GO:0031398: positive regulation of protein ubiquitination       AIMP2   aminoacyl tRNA synthetase com
plex-interacting multifunctional protein 2      0.0012010069598741675
GO:0031398: positive regulation of protein ubiquitination       ANGPT1  angiopoietin 1  0.00089136580
74095579
GO:0031398: positive regulation of protein ubiquitination       AXIN1   axin 1  -0.000721306845065099
9
GO:0031398: positive regulation of protein ubiquitination       CAV1    caveolin 1, caveolae protein,
22kDa   -0.0005316868040034789
GO:0031398: positive regulation of protein ubiquitination       CDK5RAP3        CDK5 regulatory subun
it associated protein 3 0.0011275586233072874
GO:0031398: positive regulation of protein ubiquitination       FGFR3   fibroblast growth factor rece
ptor 3  0.00021868572785721461
GO:0031398: positive regulation of protein ubiquitination       RCHY1   ring finger and CHY zinc fing
er domain containing 1, E3 ubiquitin protein ligase     -0.0010856415125516392
GO:0031398: positive regulation of protein ubiquitination       SPHK1   sphingosine kinase 1    0.001
7881180246662413
GO:0031398: positive regulation of protein ubiquitination       WFS1    Wolfram syndrome 1 (wolframi
n)      0.0005759734548774605
GO:0060510: Type II pneumocyte differentiation  AIMP2   aminoacyl tRNA synthetase complex-interacting
multifunctional protein 2       0.001213658882964944
GO:0060510: Type II pneumocyte differentiation  GATA6   GATA binding protein 6  -2.6558144610594352e-
5
GO:0060510: Type II pneumocyte differentiation  IGF1    insulin-like growth factor 1 (somatomedin C)
0.0001329820850048354
GO:0060510: Type II pneumocyte differentiation  NFIB    nuclear factor I/B      0.002944762326837304
GO:1901216: positive regulation of neuron death AIMP2   aminoacyl tRNA synthetase complex-interacting
multifunctional protein 2       0.0011963101836747186
GO:1901216: positive regulation of neuron death DAXX    death-domain associated protein 0.00087052662
13619487
GO:1901216: positive regulation of neuron death MAP3K5  mitogen-activated protein kinase kinase kinas
e 5     0.0003345509255520174
KEGG:00980: Metabolism of xenobiotics by cytochrome P450        AKR1C1  aldo-keto reductase family 1,
member C1       -0.0014842345117490703
KEGG:00980: Metabolism of xenobiotics by cytochrome P450        CYP1B1  cytochrome P450, family 1, su
bfamily B, polypeptide 1        0.00039667067958173464
KEGG:00980: Metabolism of xenobiotics by cytochrome P450        GSTM1   glutathione S-transferase mu
 1      -0.0014679242835137231
KEGG:00980: Metabolism of xenobiotics by cytochrome P450        GSTM2   glutathione S-transferase mu
 2 (muscle)     -0.0019064539301062594
KEGG:00980: Metabolism of xenobiotics by cytochrome P450        GSTM3   glutathione S-transferase mu
 3 (brain)      -0.0012838890629065193
KEGG:00980: Metabolism of xenobiotics by cytochrome P450        GSTP1   glutathione S-transferase pi
 1      0.00022946726349868944
KEGG:00140: Steroid hormone biosynthesis        AKR1C1  aldo-keto reductase family 1, member C1 -0.00
14712069940209108
KEGG:00140: Steroid hormone biosynthesis        CYP1B1  cytochrome P450, family 1, subfamily B, polyp

eptide 1 0.000392570391013102
KEGG:00140: Steroid hormone biosynthesis CYP7B1 cytochrome P450, family 7, subfamily B, polypeptide 1 0.00016022857801800203
KEGG:00140: Steroid hormone biosynthesis HSD17B6 hydroxysteroid (17-beta) dehydrogenase 6 -0.0008473314212231181
KEGG:00140: Steroid hormone biosynthesis SRD5A2 steroid-5-alpha-reductase, alpha polypeptide 2 (3-oxo-5 alpha-steroid delta 4-dehydrogenase alpha 2) -0.0006299596753869803
GO:0001523: retinoid metabolic process AKR1C1 aldo-keto reductase family 1, member C1 -0.0014878829230930406
GO:0001523: retinoid metabolic process LRP8 low density lipoprotein receptor-related protein 8, apolipoprotein e receptor -0.0009319875632310964
GO:0001523: retinoid metabolic process STRA6 stimulated by retinoic acid 6 -0.0016765788660167856
GO:0006805: xenobiotic metabolic process AKR1C1 aldo-keto reductase family 1, member C1 -0.00148942202367127
GO:0006805: xenobiotic metabolic process CYP1B1 cytochrome P450, family 1, subfamily B, polypeptide 1 0.0003978334256150409
GO:0006805: xenobiotic metabolic process CYP27B1 cytochrome P450, family 27, subfamily B, polypeptide 1 -0.0004709361534989215
GO:0006805: xenobiotic metabolic process CYP4B1 cytochrome P450, family 4, subfamily B, polypeptide 1 -0.0013561574475498966
GO:0006805: xenobiotic metabolic process CYP7B1 cytochrome P450, family 7, subfamily B, polypeptide 1 0.00016330216790402924
GO:0006805: xenobiotic metabolic process GCLM glutamate-cysteine ligase, modifier subunit -0.0037621196683362518
GO:0006805: xenobiotic metabolic process GSTM1 glutathione S-transferase mu 1 -0.0014741932946486188
GO:0006805: xenobiotic metabolic process GSTM2 glutathione S-transferase mu 2 (muscle) -0.0019143488737018634
GO:0006805: xenobiotic metabolic process GSTM3 glutathione S-transferase mu 3 (brain) -0.0012889978412383453
GO:0006805: xenobiotic metabolic process GSTP1 glutathione S-transferase pi 1 0.0002314408387097846
GO:0006805: xenobiotic metabolic process SLC35D1 solute carrier family 35 (UDP-GlcA/UDP-GalNAc transporter), member D1 -6.69650537166446e-5
GO:0006805: xenobiotic metabolic process SULT4A1 sulfotransferase family 4A, member 1 -0.0011690724189763392
GO:0006805: xenobiotic metabolic process UGDH UDP-glucose 6-dehydrogenase 0.0005021831791150163
GO:0006805: xenobiotic metabolic process UGP2 UDP-glucose pyrophosphorylase 2 0.0007852680716861068
GO:0007586: digestion AKR1C1 aldo-keto reductase family 1, member C1 -0.0014971109160566137
GO:0007586: digestion NMU neuromedin U 0.0018630596659245924
GO:0007586: digestion PGC progastricsin (pepsinogen C) 0.0013304868788324758
GO:0007586: digestion SLC15A1 solute carrier family 15 (oligopeptide transporter), member 1 -0.002777325068263749
GO:0007603: phototransduction, visible light AKR1C1 aldo-keto reductase family 1, member C1 -0.0014807883341618937
GO:0007603: phototransduction, visible light LRP8 low density lipoprotein receptor-related protein 8, apolipoprotein e receptor -0.0009278360939371799
GO:0007603: phototransduction, visible light METAP1 methionyl aminopeptidase 1 -0.001011936479930692
GO:0007603: phototransduction, visible light PRKCA protein kinase C, alpha -6.423218744170265e-6
GO:0007603: phototransduction, visible light STRA6 stimulated by retinoic acid 6 -0.0016651468689401101
GO:0015721: bile acid and bile salt transport AKR1C1 aldo-keto reductase family 1, member C1 -0.0014814815180658955
GO:0015721: bile acid and bile salt transport RXRA retinoid X receptor, alpha 0.0011089479183730645
GO:0030299: intestinal cholesterol absorption AKR1C1 aldo-keto reductase family 1, member C1 -0.0014835116889152422
GO:0030855: epithelial cell differentiation AKR1C1 aldo-keto reductase family 1, member C1 -0.0014884134398433027
GO:0030855: epithelial cell differentiation BMP7 bone morphogenetic protein 7 0.0008552728225342675
GO:0030855: epithelial cell differentiation CDK1 cyclin-dependent kinase 1 0.00030944164835295686
GO:0030855: epithelial cell differentiation CPT1A carnitine palmitoyltransferase 1A (liver) 0.0012723765196986756
GO:0030855: epithelial cell differentiation DLX5 distal-less homeobox 5 -0.0032809231137978915
GO:0030855: epithelial cell differentiation DLX6 distal-less homeobox 6 -0.0001799230800021182 8
GO:0030855: epithelial cell differentiation FGFR2 fibroblast growth factor receptor 2 0.0007621441874261383
GO:0030855: epithelial cell differentiation PCNA proliferating cell nuclear antigen 0.001223721972125872
GO:0030855: epithelial cell differentiation SIX1 SIX homeobox 1 -0.0018681328167706032
GO:0030855: epithelial cell differentiation STX2 syntaxin 2 0.00030777425472991286
GO:0030855: epithelial cell differentiation VEGFA vascular endothelial growth factor A 0.000593340280970033 3

GO:0030855: epithelial cell differentiation WT1 Wilms tumor 1 -0.0005061393445866929
GO:0042448: progesterone metabolic process AKR1C1 aldo-keto reductase family 1, member C1 -0.00
14835116889152422
GO:0042574: retinal metabolic process AKR1C1 aldo-keto reductase family 1, member C1 -0.0014965655
797961744
GO:0042574: retinal metabolic process ALDH1A2 aldehyde dehydrogenase 1 family, member A2 -0.00
3034857237374307
GO:0042574: retinal metabolic process CYP1B1 cytochrome P450, family 1, subfamily B, polypeptide 1
0.00039888720936128553
GO:0044597: daunorubicin metabolic process AKR1C1 aldo-keto reductase family 1, member C1 -0.00
14835116889152422
GO:0044598: doxorubicin metabolic process AKR1C1 aldo-keto reductase family 1, member C1 -0.00
14835116889152422
GO:0046683: response to organophosphorus AKR1C1 aldo-keto reductase family 1, member C1 -0.00
14904468159268127
GO:0046683: response to organophosphorus RFC3 replication factor C (activator 1) 3, 38kDa
0.0003143934576300518
GO:0046683: response to organophosphorus TRIM16 tripartite motif containing 16 -0.0014908189
289572127
GO:0046683: response to organophosphorus TYMS thymidylate synthetase 0.0015651544575019053
GO:0051260: protein homooligomerization AKR1C1 aldo-keto reductase family 1, member C1 -0.0014765187
659678439
GO:0051260: protein homooligomerization AXIN1 axin 1 -0.0007229657882528065
GO:0051260: protein homooligomerization BAX BCL2-associated X protein -0.000421813608639165
27
GO:0051260: protein homooligomerization CAV1 caveolin 1, caveolae protein, 22kDa -0.0005317576
552947029
GO:0051260: protein homooligomerization CEP57 centrosomal protein 57kDa -0.000535008865069088
7
GO:0051260: protein homooligomerization CPT1A carnitine palmitoyltransferase 1A (liver) 0.001
263172456800l816
GO:0051260: protein homooligomerization CRYAA crystallin, alpha A -0.000666426903659395
GO:0051260: protein homooligomerization CRYAB crystallin, alpha B 0.0009705595613955782
GO:0051260: protein homooligomerization ECT2 epithelial cell transforming 2 0.0010495059974709273
GO:0051260: protein homooligomerization FAS Fas cell surface death receptor -3.284126954160665e-5
GO:0051260: protein homooligomerization FLOT1 flotillin 1 -0.0007567730086071728
GO:0051260: protein homooligomerization HMOX1 heme oxygenase (decycling) 1 -0.00021458075627884O
74
GO:0051260: protein homooligomerization KCNMA1 potassium large conductance calcium-activated channe
l, subfamily M, alpha member 1 -0.000646759362272597
GO:0051260: protein homooligomerization KCNV2 potassium channel, subfamily V, member 2 0.000
9177516646332517
GO:0051260: protein homooligomerization RAD51 RAD51 recombinase -0.001842998967944961
GO:0051260: protein homooligomerization SLC1A1 solute carrier family 1 (neuronal/epithelial high aff
inity glutamate transporter, system Xag), member 1 -0.0008918154514029545
GO:0051260: protein homooligomerization SYT1 synaptotagmin I -0.0014261697188205774
GO:0071395: cellular response to jasmonic acid stimulus AKR1C1 aldo-keto reductase family 1, member
C1 -0.0014835116889152422
KEGG:04510: Focal adhesion AKT1 v-akt murine thymoma viral oncogene homolog 1 0.00073111196
3575552
KEGG:04510: Focal adhesion AKT2 v-akt murine thymoma viral oncogene homolog 2 -0.0006259514
437814398
KEGG:04510: Focal adhesion BCL2 B-cell CLL/lymphoma 2 -5.087034901944398e-6
KEGG:04510: Focal adhesion CAV1 caveolin 1, caveolae protein, 22kDa -0.000535454747021665
4
KEGG:04510: Focal adhesion CCND1 cyclin D1 -0.0026352127440208276
KEGG:04510: Focal adhesion CDC42 cell division cycle 42 0.0012409586708569862
KEGG:04510: Focal adhesion CHAD chondroadherin -0.0021796141370551162
KEGG:04510: Focal adhesion COL11A1 collagen, type XI, alpha 1 -0.0004419448188928467
KEGG:04510: Focal adhesion COL1A1 collagen, type I, alpha 1 -0.0005271888644141816
KEGG:04510: Focal adhesion COL5A1 collagen, type V, alpha 1 -5.0553277993472204e-5
KEGG:04510: Focal adhesion COL5A2 collagen, type V, alpha 2 -0.00034630787067053227
KEGG:04510: Focal adhesion CTNNB1 catenin (cadherin-associated protein), beta 1, 88kDa -0.00
011676958739873388
KEGG:04510: Focal adhesion DOCK1 dedicator of cytokinesis 1 -0.0006848151025178808
KEGG:04510: Focal adhesion EGFR epidermal growth factor receptor 0.0006866066273333798
KEGG:04510: Focal adhesion GRB2 growth factor receptor-bound protein 2 0.0004297740014662271
6
KEGG:04510: Focal adhesion GSK3B glycogen synthase kinase 3 beta 0.0015498469809238434
KEGG:04510: Focal adhesion IGF1 insulin-like growth factor 1 (somatomedin C) 0.00013115624
351227068
KEGG:04510: Focal adhesion IGF1R insulin-like growth factor 1 receptor 0.0010719019030154214
KEGG:04510: Focal adhesion ITGA3 integrin, alpha 3 (antigen CD49C, alpha 3 subunit of VLA-3 re
ceptor) 0.0014001321876838258
KEGG:04510: Focal adhesion ITGA6 integrin, alpha 6 0.0016709391283725884
KEGG:04510: Focal adhesion ITGB1 integrin, beta 1 (fibronectin receptor, beta polypeptide, ant
igen CD29 includes MDF2, MSK12) 0.0024282030285735774
KEGG:04510: Focal adhesion ITGB4 integrin, beta 4 0.0005810363399926138
KEGG:04510: Focal adhesion LAMB2 laminin, beta 2 (laminin S) -0.0013099544265133157
KEGG:04510: Focal adhesion PAK1 p21 protein (Cdc42/Rac)-activated kinase 1 -0.0021888679
20935422

```
KEGG:04510: Focal adhesion      PAK3    p21 protein (Cdc42/Rac)-activated kinase 3      -0.0012590159
858141346
KEGG:04510: Focal adhesion      PDGFRB  platelet-derived growth factor receptor, beta polypeptide
-0.0003620155184959438
KEGG:04510: Focal adhesion      PIK3CD  phosphatidylinositol-4,5-bisphosphate 3-kinase, catalytic sub
unit delta      -0.0007812225179908655
KEGG:04510: Focal adhesion      PRKCA   protein kinase C, alpha -5.841197382942685e-6
KEGG:04510: Focal adhesion      PTEN    phosphatase and tensin homolog  1.6638750585002486e-5
KEGG:04510: Focal adhesion      RAF1    Raf-1 proto-oncogene, serine/threonine kinase   0.00149950669
9584488
KEGG:04510: Focal adhesion      RELN    reelin  0.0015384521456824544
KEGG:04510: Focal adhesion      RHOA    ras homolog family member A     0.0006700457466109884
KEGG:04510: Focal adhesion      THBS1   thrombospondin 1        -0.0010365297244895536
KEGG:04510: Focal adhesion      THBS3   thrombospondin 3        0.0016180337455121826
KEGG:04510: Focal adhesion      THBS4   thrombospondin 4        -0.00044493624999140485
KEGG:04510: Focal adhesion      TNC     tenascin C      0.0007346498478172675
KEGG:04510: Focal adhesion      VEGFA   vascular endothelial growth factor A    0.000595741455227481
KEGG:04510: Focal adhesion      VEGFC   vascular endothelial growth factor C    -0.003365676955124798
KEGG:05166: HTLV-I infection    AKT1    v-akt murine thymoma viral oncogene homolog 1   0.00073014872
65280059
KEGG:05166: HTLV-I infection    AKT2    v-akt murine thymoma viral oncogene homolog 2   -0.0006254672
127444562
KEGG:05166: HTLV-I infection    APC     adenomatous polyposis coli      0.0006433803976048463
KEGG:05166: HTLV-I infection    BAX     BCL2-associated X protein       -0.0004248209935957699

Excessive output truncated after 524318 bytes.
```

KEGG:05166: HTLV-I infection	BUB1B	BUB1 mitotic checkpoint serine/threonine kinase B	-0.00
12803290688684185
KEGG:05166: HTLV-I infection	BUB3	BUB3 mitotic checkpoint protein 0.0016214780334313297
KEGG:05166: HTLV-I infection	CCND1	cyclin D1	-0.0026314383248254762
KEGG:05166: HTLV-I infection	CDC20	cell division cycle 20	-0.00018846746314222143
KEGG:05166: HTLV-I infection	CDC23	cell division cycle 23	-0.0015523879891386671
KEGG:05166: HTLV-I infection	CDC27	cell division cycle 27	0.0005361343533335339
KEGG:05166: HTLV-I infection	CDKN2A	cyclin-dependent kinase inhibitor 2A	0.0017572163952829754
KEGG:05166: HTLV-I infection	CDKN2B	cyclin-dependent kinase inhibitor 2B (p15, inhibits CDK4)
-0.0018343261379406253
KEGG:05166: HTLV-I infection	CHEK1	checkpoint kinase 1	0.0008996601738664551
KEGG:05166: HTLV-I infection	CREB1	cAMP responsive element binding protein 1	0.00065793162
82892347
KEGG:05166: HTLV-I infection	CTNNB1	catenin (cadherin-associated protein), beta 1, 88kDa	-0.00
01167776571928328
KEGG:05166: HTLV-I infection	E2F1	E2F transcription factor 1	0.002096507746263104
KEGG:05166: HTLV-I infection	E2F3	E2F transcription factor 3	0.002160892745362072
KEGG:05166: HTLV-I infection	EGR1	early growth response 1 0.0010924201527298136
KEGG:05166: HTLV-I infection	EGR2	early growth response 2 0.0014379525470036324
KEGG:05166: HTLV-I infection	ETS2	v-ets avian erythroblastosis virus E26 oncogene homolog 2
-0.0010148671813836455
KEGG:05166: HTLV-I infection	FZD3	frizzled class receptor 3	0.00037831878734940306
KEGG:05166: HTLV-I infection	FZD7	frizzled class receptor 7	0.001070010419455324
KEGG:05166: HTLV-I infection	GSK3B	glycogen synthase kinase 3 beta 0.0015481661412175241
KEGG:05166: HTLV-I infection	ICAM1	intercellular adhesion molecule 1	0.0007288551674977717
KEGG:05166: HTLV-I infection	MAD2L1	MAD2 mitotic arrest deficient-like 1 (yeast)	0.00031861014
213334566
KEGG:05166: HTLV-I infection	MSX1	msh homeobox 1	-0.002767385544638794
KEGG:05166: HTLV-I infection	MYC	v-myc avian myelocytomatosis viral oncogene homolog	-0.00
11355383056244046
KEGG:05166: HTLV-I infection	NRP1	neuropilin 1	-0.0006427489251350105
KEGG:05166: HTLV-I infection	PCNA	proliferating cell nuclear antigen	0.0012255293610770976
KEGG:05166: HTLV-I infection	PDGFRB	platelet-derived growth factor receptor, beta polypeptide
-0.00036110174672602147
KEGG:05166: HTLV-I infection	PIK3CD	phosphatidylinositol-4,5-bisphosphate 3-kinase, catalytic sub
unit delta	-0.0007803532808301466
KEGG:05166: HTLV-I infection	POLD1	polymerase (DNA directed), delta 1, catalytic subunit	-0.00
04444761587691851
KEGG:05166: HTLV-I infection	POLD2	polymerase (DNA directed), delta 2, accessory subunit	0.000
7395023232667378
KEGG:05166: HTLV-I infection	POLD3	polymerase (DNA-directed), delta 3, accessory subunit	0.001
739501097365773
KEGG:05166: HTLV-I infection	POLD4	polymerase (DNA-directed), delta 4, accessory subunit	-0.00
06380253171875901
KEGG:05166: HTLV-I infection	POLE	polymerase (DNA directed), epsilon, catalytic subunit	0.002
9465906056635703
KEGG:05166: HTLV-I infection	POLE2	polymerase (DNA directed), epsilon 2, accessory subunit 0.001
3251314623131934
KEGG:05166: HTLV-I infection	POLE3	polymerase (DNA directed), epsilon 3, accessory subunit 0.000
8044456814438906
KEGG:05166: HTLV-I infection	PRKX	protein kinase, X-linked	0.0003947014380226403
KEGG:05166: HTLV-I infection	PTTG1	pituitary tumor-transforming 1	-0.00036139932312809366
KEGG:05166: HTLV-I infection	PTTG2	pituitary tumor-transforming 2	-0.001723714808519096
KEGG:05166: HTLV-I infection	RB1	retinoblastoma 1	-0.0014930913695895957
KEGG:05166: HTLV-I infection	STAT5A	signal transducer and activator of transcription 5A	0.001
5953326831280425
KEGG:05166: HTLV-I infection	TBPL1	TBP-like 1	-0.0007881631366525118
KEGG:05166: HTLV-I infection	TGFB1	transforming growth factor, beta 1	-7.303893692018511e-5
KEGG:05166: HTLV-I infection	TGFB2	transforming growth factor, beta 2	-0.001058701643025118
6
KEGG:05166: HTLV-I infection	TGFB3	transforming growth factor, beta 3	-0.001822789925395239
4
KEGG:05166: HTLV-I infection	TGFBR1	transforming growth factor, beta receptor 1	0.00033971200
41990668
KEGG:05166: HTLV-I infection	TP53	tumor protein p53	0.0011750040561670676
KEGG:05166: HTLV-I infection	WNT1	wingless-type MMTV integration site family, member 1	0.000
7864609236476779
KEGG:05166: HTLV-I infection	WNT10B	wingless-type MMTV integration site family, member 10B	-0.00
13876971599892575
KEGG:05166: HTLV-I infection	WNT4	wingless-type MMTV integration site family, member 4	-0.00
025196024583135583
KEGG:05166: HTLV-I infection	WNT5A	wingless-type MMTV integration site family, member 5A	-0.00
06671137872325666
KEGG:05166: HTLV-I infection	WNT7A	wingless-type MMTV integration site family, member 7A	1.957
9561150038034e-5
KEGG:05166: HTLV-I infection	XBP1	X-box binding protein 1 0.0002668001478054871
KEGG:05169: Epstein-Barr virus infection	AKT1	v-akt murine thymoma viral oncogene homolog 1
0.0007262566190756881
KEGG:05169: Epstein-Barr virus infection	AKT2	v-akt murine thymoma viral oncogene homolog 2
-0.0006235672077896339
KEGG:05169: Epstein-Barr virus infection	BCL2	B-cell CLL/lymphoma 2	-4.882937985058629e-6

KEGG:05169: Epstein-Barr virus infection CCNA1 cyclin A1 -0.001750148799974855
KEGG:05169: Epstein-Barr virus infection CDK1 cyclin-dependent kinase 1 0.00030695291
37300434
KEGG:05169: Epstein-Barr virus infection CSNK2B casein kinase 2, beta polypeptide 0.001
5192769043707265
KEGG:05169: Epstein-Barr virus infection GSK3B glycogen synthase kinase 3 beta 0.00154119024
95898234
KEGG:05169: Epstein-Barr virus infection HDAC2 histone deacetylase 2 -0.001203898890181675
2
KEGG:05169: Epstein-Barr virus infection HSPA2 heat shock 70kDa protein 2 -0.0001034582
414893158
KEGG:05169: Epstein-Barr virus infection ICAM1 intercellular adhesion molecule 1 0.000
7273626760522081
KEGG:05169: Epstein-Barr virus infection IFNG interferon, gamma -4.900796379268888e-5
KEGG:05169: Epstein-Barr virus infection IRAK1 interleukin-1 receptor-associated kinase 1
-0.0016002937427628528
KEGG:05169: Epstein-Barr virus infection LYN LYN proto-oncogene, Src family tyrosine kinas
e -0.0014136847532094168
KEGG:05169: Epstein-Barr virus infection MYC v-myc avian myelocytomatosis viral oncogene h
omolog -0.0011325117414121158
KEGG:05169: Epstein-Barr virus infection NEDD4 neural precursor cell expressed, developmenta
lly down-regulated 4, E3 ubiquitin protein ligase 0.002253522858241044
KEGG:05169: Epstein-Barr virus infection PIK3CD phosphatidylinositol-4,5-bisphosphate 3-kinas
e, catalytic subunit delta -0.0007767642901220774
KEGG:05169: Epstein-Barr virus infection POLR2D polymerase (RNA) II (DNA directed) polypeptid
e D 0.0018450093234024662
KEGG:05169: Epstein-Barr virus infection PRKX protein kinase, X-linked 0.00039030241
681122646
KEGG:05169: Epstein-Barr virus infection PSMD11 proteasome (prosome, macropain) 26S subunit,
non-ATPase, 11 -0.001091548683016147
KEGG:05169: Epstein-Barr virus infection PSMD13 proteasome (prosome, macropain) 26S subunit,
non-ATPase, 13 -6.068925883060733e-5
KEGG:05169: Epstein-Barr virus infection RB1 retinoblastoma 1 -0.001486167860636004
6
KEGG:05169: Epstein-Barr virus infection RBPJ recombination signal binding protein for immu
noglobulin kappa J region 0.0009777741038018578
KEGG:05169: Epstein-Barr virus infection SKP2 S-phase kinase-associated protein 2, E3 ubiqu
itin protein ligase -0.0005374559859162735
KEGG:05169: Epstein-Barr virus infection TBPL1 TBP-like 1 -0.0007871806823993505
KEGG:05169: Epstein-Barr virus infection TNFAIP3 tumor necrosis factor, alpha-induced protein
3 0.0011106622138456054
KEGG:05169: Epstein-Barr virus infection TP53 tumor protein p53 0.0011672564642736677
KEGG:04630: Jak-STAT signaling pathway AKT1 v-akt murine thymoma viral oncogene homolog 1 0.000
7322366576941183
KEGG:04630: Jak-STAT signaling pathway AKT2 v-akt murine thymoma viral oncogene homolog 2 -0.00
06269139625152389
KEGG:04630: Jak-STAT signaling pathway CCND1 cyclin D1 -0.002638355521223424
KEGG:04630: Jak-STAT signaling pathway CSF3R colony stimulating factor 3 receptor (granulocyte)
-8.433393498168491e-5
KEGG:04630: Jak-STAT signaling pathway CSH1 chorionic somatomammotropin hormone 1 (placental lact
ogen) -0.00025766423016498137
KEGG:04630: Jak-STAT signaling pathway GRB2 growth factor receptor-bound protein 2 0.00043051485
571086605
KEGG:04630: Jak-STAT signaling pathway IFNA2 interferon, alpha 2 -0.0017203925871120368
KEGG:04630: Jak-STAT signaling pathway IFNG interferon, gamma -4.899191761680744e-5
KEGG:04630: Jak-STAT signaling pathway IL12B interleukin 12B 0.0012788481986337784
KEGG:04630: Jak-STAT signaling pathway IL20RA interleukin 20 receptor, alpha 0.0001745061911577262
3
KEGG:04630: Jak-STAT signaling pathway IL4 interleukin 4 0.00025976679277823594
KEGG:04630: Jak-STAT signaling pathway IL6ST interleukin 6 signal transducer 0.0018793482781852779
KEGG:04630: Jak-STAT signaling pathway IL7 interleukin 7 0.0008729639337432425
KEGG:04630: Jak-STAT signaling pathway JAK2 Janus kinase 2 -3.159219747417667e-5
KEGG:04630: Jak-STAT signaling pathway LEP leptin 0.0031976777282567794
KEGG:04630: Jak-STAT signaling pathway MYC v-myc avian myelocytomatosis viral oncogene homolog
-0.0011388686705106022
KEGG:04630: Jak-STAT signaling pathway PIK3CD phosphatidylinositol-4,5-bisphosphate 3-kinase, catal
ytic subunit delta -0.0007822987779466499
KEGG:04630: Jak-STAT signaling pathway PRLR prolactin receptor 0.002187381210529882
KEGG:04630: Jak-STAT signaling pathway STAT5A signal transducer and activator of transcription 5A
0.0015991927405241135
KEGG:04071: Sphingolipid signaling pathway AKT1 v-akt murine thymoma viral oncogene homolog 1
0.0007202122493181 33
KEGG:04071: Sphingolipid signaling pathway AKT2 v-akt murine thymoma viral oncogene homolog 2
-0.000619913285639517
KEGG:04071: Sphingolipid signaling pathway BAX BCL2-associated X protein -0.0004216900
039775024
KEGG:04071: Sphingolipid signaling pathway BCL2 B-cell CLL/lymphoma 2 -4.414007247247561e-6

In [58]:

```
for (v, β) in zip(variable_names[β_orig .!= 0], β_orig[β_orig .!= 0])
    println("$v\t$β")
end
```

| | |
|---|---|
| RFC2 | 0.017828370354349206 |
| PAX8 | 0.03684757874519626 |
| THRA | 0.028067107007347464 |
| EPHB3 | 0.019420774902517746 |
| CFL1 | -0.030460550648403983 |
| YY1 | 0.013402782876598919 |
| ZPR1 | -0.004170459413053396 |
| RHOA | 0.04322099220670631 |
| GUK1 | 0.05284161532839051 |
| DSP | -0.0008924333259685127 |
| RAD21 | -0.0016314685844209594 |
| SF3B2 | 0.004117778492081898 |
| NDRG1 | -0.006543628992920167 |
| CD63 | -0.015885903765799948 |
| DNAJB1 | -0.00207402357079152 8 |
| XBP1 | 0.021161233616516416 |
| SPTBN1 | 0.03684346496032823 |
| HMGB1 | -0.03075659605031252 |
| GPX1 | 0.021234007378015077 |
| NUMA1 | 0.021341452238237472 |
| ARL6IP5 | 0.0010924161799196034 |
| STMN1 | 0.007247922509085316 |
| ODC1 | 0.00984689697218672 |
| DAZAP2 | -0.0007855012627521639 |
| TMBIM6 | -0.0022818769304895916 |
| GSTP1 | 0.008155197247241442 |
| ZNF207 | -0.0027205566179267695 |
| DHCR24 | -0.03582951324451189 |
| DEK | 0.017525131494012178 |
| HIF1A | -0.049801244025156705 |
| SNX17 | 0.011478189129344106 |
| CD9 | -0.038892803436755025 |
| JUP | -0.038198768007394066 |
| TGM2 | -0.0017271186611303253 |
| MMP2 | -0.02598193263059218 |
| THBS1 | -0.08446415723750327 |
| TUFM | -0.006526015058534712 |
| POLD2 | 0.017044076906068118 |
| CPE | -0.02298547947437439 |
| SLC7A5 | -0.04388439401899304 |
| AMD1 | -0.025402082120591245 |
| PCNA | 0.04177675913977362 |
| PSMD13 | -0.0024832832828435856 |
| RAF1 | 0.14338085024453623 |
| PSMA5 | 0.01305134817715597 |
| OGDH | -0.005067398489416658 |
| TOP2A | -0.0015644814822285936 |
| ETS2 | -0.012155698340348462 |
| CST3 | -0.0020043270870368723 |
| ZFP36L2 | -0.004307263131398863 |
| CSNK2B | 0.03354336593236935 |
| PHGDH | 0.005731129900934964 |
| HSD17B4 | -0.02195477179188622 |
| SOX4 | -0.0018435626169123894 |
| WDR77 | 0.0010212673455483959 |
| BUB3 | 0.02098069229083203 |
| ITGA3 | 0.05313965779313374 |
| RRM1 | 0.03261837436083388 |
| IGFBP4 | -0.015294959879804445 |
| APOD | 0.052813903449321326 |
| RPA1 | 0.01670607000190569 |
| CTNNB1 | -0.01669235255966663 |
| ZNHIT1 | -0.0016925861371024221 |
| MCM3 | 0.02142799502691732 |
| SORD | -0.03855195661545189 |
| ID2 | 0.0030436585873623 7 |
| IRAK1 | -0.08686643583075766 |
| SMC1A | -0.02133148172546458 |
| KRT18 | -0.010141434375582896 |
| PHB2 | -0.0123826123009966 5 |
| TNC | 0.010270550679614054 |
| ITGA6 | 0.06202647497027872 |
| SMC4 | 0.008291635589654026 |
| GJA1 | -0.009687495934931687 |
| EGR1 | 0.04890522631472147 |
| TPR | -0.028977825234667652 |
| TP53 | 0.17001586712848318 |
| MCM5 | 0.013891019627766394 |
| RPA2 | 0.008798613227269441 |
| DAXX | 0.022950864698423112 |
| AZIN1 | -0.0008413038683729089 |

| | |
|---|---|
| NCAPD2 | -0.006694478531962177 |
| ADAR | -0.004273578061227502 |
| ENG | 0.0303531751783837 |
| HDAC2 | -0.07936953927783139 |
| BNIP3 | 0.10258521554357351 |
| RRM2 | -0.009046537510581779 |
| IMPDH2 | -0.004533325675427474 |
| PSRC1 | 0.00515087891989595 |
| PRDX4 | 0.015233898271638829 |
| MCM6 | 0.013085227098779626 |
| CHMP1A | -0.011579771180144555 |
| CIB1 | 0.002494086200296384 |
| CCNC | 0.005668826446434941 |
| NASP | 0.01679481485255259 |
| EGFR | 0.05923885482145961 |
| LANCL1 | 0.0007339826494670798 |
| EFNA1 | -0.01241014380812595 |
| SFRP1 | 0.10537629932164161 |
| SLC39A6 | 0.0011622219237774859 |
| BIRC5 | -0.006728392679664581 |
| NUP153 | 0.020401330786399767 |
| BRD4 | 0.0098786158197804 |
| IGBP1 | 0.03716659575857557 |
| MCM2 | 0.02964701901069167 |
| NOC2L | -0.0013018167154264441 |
| WWTR1 | 0.024221399551931384 |
| AIMP2 | 0.008424538648259098 |
| PKN1 | -0.038089434810030715 |
| PRKCZ | -0.08299626615623115 |
| MVP | -0.00503993936641827 |
| SLC6A8 | -0.010418167114822331 |
| PLK1 | 0.044418889686983744 |
| E2F4 | -0.04000230602549144 |
| VPS72 | 0.002791817778844994 |
| PDGFRB | -0.030899955754132916 |
| TACSTD2 | -0.02726592640576279 |
| MTHFD1 | -0.015106671256448267 |
| COL1A1 | -0.02774187855127703 |
| UNG | 0.0050402466853051615 |
| UBE2K | -0.002633814755276221 |
| HTT | -0.0433308657325134 |
| BASP1 | 0.004749137833453016 |
| IGSF3 | 0.00028465425193114957 |
| RXRA | 0.05780812177993815 |
| MYC | -0.07254841230400205 |
| CYP1B1 | 0.016791485175176302 |
| CTSK | -0.005372887927588193 |
| MLXIP | 0.01097144116387061 |
| MLH1 | 0.049206926689411956 |
| DHFR | -0.007368555092904131 |
| PRKCD | -0.06848826247053864 |
| VAMP8 | -0.0006123414941763332 |
| GSTM3 | -0.01677195290960026 |
| FOXM1 | 0.005470530397087538 |
| NFX1 | -0.00036010966259641177 |
| TYMS | 0.05804193843777034 |
| PDK2 | 0.022218697269260047 |
| SSBP1 | 0.0033581919245183277 |
| ENSA | 0.011071869332661386 |
| IRF6 | -0.018578500771458274 |
| NRIP1 | 0.010689665804530209 |
| CTPS1 | -0.005033769164073506 |
| LYN | -0.13602000880799608 |
| SERPINE1 | 0.004549817149767903 |
| TOPBP1 | 0.00826258318265347 |
| ICAM1 | 0.046988132805018984 |
| ARL3 | -0.012944671028492987 |
| TNFAIP3 | 0.06196838444377323 |
| RASA1 | -0.010347310860967336 |
| PTPN1 | -0.03400858029035727 |
| IGFBP2 | 0.002158294645734256 |
| FOXO1 | 0.08981646049044094 |
| LIG1 | 0.0193507999466856 |
| USP8 | -0.004295447026511487 |
| SLC7A8 | -0.007029917025023931 |
| HMGCL | -0.01792059645370268 |
| UBE2S | 0.004517544081043286 |
| SLC1A3 | -0.07846231096017796 |
| DBN1 | -0.007958079888046345 |
| WBP1L | -0.0006812879912496229 |
| STAMBP | -0.0013439330003021685 |

| | |
|---|---|
| HEXIM1 | 0.0009146807422728782 |
| AGT | -0.11777825302279477 |
| HPRT1 | -0.03429130520873595 |
| CDC20 | -0.004158031714099344 |
| CDC23 | -0.0292898490145349 |
| E2F1 | 0.09452257827698173 |
| WFS1 | 0.016589595131934934 |
| MSH6 | 0.01827567569356198 |
| ADM | 0.09880822122504294 |
| SOX9 | -0.0553437515959798 |
| UBE2C | -0.005246903745117184 |
| MUT | 0.035950521811880666 |
| POLD4 | -0.015987318033880763 |
| STMN2 | -0.01556383248822085 |
| STAT5A | 0.09242498840939786 |
| RNASEH2A | 0.00413588001927989 |
| C2 | -0.0033094363111025545 |
| CAV1 | -0.06306570615282248 |
| CHST15 | -0.0057826912306122085 |
| CTNNBIP1 | 0.008138176628460052 |
| TGFB1 | -0.012869806540407442 |
| KIF2A | -0.009405835015435746 |
| CCDC86 | 0.0007146750595481014 |
| SLC11A2 | -0.025404068219129067 |
| RB1 | -0.0850807506072826 |
| KIAA0040 | -0.0022389706808229127 |
| SPAG5 | -0.0032009057004588675 |
| PVRL2 | -0.0006298298280750445 |
| RNF8 | 0.014322281504453253 |
| KATNB1 | -0.0011054373409471162 |
| FBN2 | -0.007380182720434392 |
| DOCK1 | -0.011621186923791518 |
| RFC5 | 0.04044914529319225 |
| MTMR2 | -0.008937113364945702 |
| CDK1 | 0.011067291839721289 |
| LGALS9 | -0.004539435216978999 |
| ZNF24 | 0.0038769440123150216 |
| CDH3 | -0.0281774043815424 |
| DTYMK | 0.05614040994978363 |
| HS2ST1 | -0.006412566664718715 |
| NCK2 | -0.006297849220518807 |
| ZNF148 | 0.01687101223878772 |
| COL5A1 | -0.0011032697296370568 |
| UGDH | 0.006978188918291195 |
| ORC4 | -0.0003733583483527787 |
| EZH2 | -0.004516562690241722 |
| MAD2L1 | 0.0056195618553048785 |
| CDC40 | 0.03781642763801979 |
| RPS6KA1 | -0.09063268000400476 |
| HES1 | -0.06412769702404401 |
| PSMG1 | 0.00040652018292425306 |
| MMD | 0.010754803225344929 |
| KMT2B | 0.005044953078137121 |
| POLD1 | -0.014189772998580038 |
| ASF1A | 0.009452157844126933 |
| SUCO | 0.01123697897827811 |
| STC2 | -0.01671840339994169 |
| CDH2 | -0.017260462882438505 |
| SPR | -0.01968662444539362 |
| PLEK | 0.024370121612437633 |
| CEP57 | -0.004804746283049292 |
| MED1 | 0.0675752988782279 |
| GCDH | 7.034724478452054e-6 |
| SPG11 | -8.779000084344101e-5 |
| APC | 0.04037734300954118 |
| PTTG1 | -0.005785185428833486 |
| CUL7 | -0.001874500267223692 |
| AOC1 | -0.01847043164444562 |
| GGH | -0.0024980678538852955 |
| ADIRF | -0.0011367963587797798 |
| CSF3R | -0.0010011980500108369 |
| ALDH5A1 | 0.04921593880047607 |
| SKP2 | -0.011781180780861291 |
| IGF1R | 0.05658983955277871 |
| CPT1A | 0.027930537006637578 |
| FGFR2 | 0.09289228548238233 |
| PCNT | 0.0286016211618618 |
| POLR2D | 0.04460874361078859 |
| HMOX1 | -0.010639419074505253 |
| CXCL12 | -0.0577420790375968 |
| IVD | -0.0074381991637425815 |

| | |
|---|---|
| BCL2 | -0.0008945318307243503 |
| CX3CL1 | 0.08062568872404573 |
| PI3 | -0.002078090528169931 |
| E2F3 | 0.04115371621305719 |
| RREB1 | -0.021820656476435024 |
| FZD7 | 0.02875299261893593 |
| ITPR1 | -0.04164910864273341 |
| KIAA0020 | 0.0009336617440814803 |
| BAK1 | -0.11224104241962284 |
| HMGB3 | 0.003201531061351816 |
| BUB1B | -0.020432339950024473 |
| DYNC2LI1 | 0.004614918362283695 |
| DLGAP5 | -0.007949800760510885 |
| MANBA | -0.019025815224275856 |
| SEMA3C | -0.003073579063100939 |
| AKT2 | -0.06589440234688392 |
| IGF2BP3 | -0.0022821883184598094 |
| MAP3K5 | 0.00973399305748306 |
| IGFBP6 | -0.010108693926009852 |
| PIK3CD | -0.08446872456456452 |
| CXADR | 0.008125836938706712 |
| NR1H3 | 0.029204949713111033 |
| GCLM | -0.07189730152604519 |
| MAPT | 0.035560439470454265 |
| VASH1 | 0.005336994188221941 |
| KIF3B | -0.002474744514527213 |
| E2F6 | 0.00215732876329845 |
| CDC6 | 0.021677609174670775 |
| SLC31A1 | -0.0002675017652726711 |
| CHAF1A | 0.002173170247103242 |
| SYT1 | -0.0231985911200916 |
| LARS2 | 0.002685547772540582 |
| PURA | -0.0015289193774571034 |
| WWP2 | -0.01810959141480461 |
| RFC4 | 0.013094847932800593 |
| ZWINT | 0.005037357692006729 |
| CELSR2 | -0.015828809359290333 |
| PTEN | 0.002703874590459573 |
| PRKX | 0.026310909182115184 |
| TPST2 | 0.002124847987139037 |
| AURKA | 0.026072099828434364 |
| RBMX2 | 0.0011525304047793957 |
| SMARCD3 | 0.005056510060143434 |
| LIG3 | 0.017937525776897882 |
| CDC45 | 0.035128743764696796 |
| RFC3 | 0.006289485493524517 |
| FOXO3 | 0.04304335440765908 |
| TFDP1 | 0.024597855220075616 |
| AKR1C1 | -0.028235242102568604 |
| NDC80 | -0.01299434899650587 |
| IMPDH1 | -0.0225262430651548 |
| CKS2 | -0.0002767823749429342 |
| CPOX | 0.015309998121761632 |
| RARG | -0.07205944464497151 |
| BACH1 | 0.0038557769403125743 |
| ACOT8 | 0.014671699445802378 |
| SMC2 | 0.003475792884709932 |
| VDR | 0.015094705098909537 |
| MMP7 | 0.00671835148729826 |
| SKI | -0.02532169897096145 |
| RGS14 | -0.03319822659144861 |
| LOX | -0.005739266799934819 |
| PROM1 | -0.007435736621323743 |
| CREB1 | 0.05291850511267864 |
| COL11A1 | -0.009683886337841511 |
| TRIM16 | -0.013286678121464168 |
| BCAS1 | -0.00012129172027097893 |
| FGFR3 | 0.015061013692887143 |
| GOLGA2 | 0.002426158179985119 |
| EFS | 0.007825864377930104 |
| GSTM2 | -0.03453019093217697 |
| SLC2A5 | 0.002611578059423238 |
| POLA2 | -0.02095432318344106 |
| KIF11 | 0.012877878211679263 |
| GAP43 | -0.0010497450042150282 |
| CDC7 | -0.009474360240546223 |
| PHF14 | 0.012996775839253325 |
| CXCL10 | 0.0022540377037781716 |
| EEF1A2 | -0.011206457515607324 |
| KIAA0513 | -0.002633608574740678 |
| STAR | 0.048759191204857645 |

| | |
|---|---|
| GSTM1 | -0.016224872433555543 |
| FKBP5 | 0.000934779712501054 |
| ABCG1 | -0.06107660423264804 |
| CROT | -0.004733522036937512 |
| CD22 | 0.003342834303618192 |
| SLC7A7 | -0.008350703535127244 |
| DKK1 | -0.004376640259688258 |
| EXO1 | -0.007169600632849004 |
| NEK2 | 0.0008735531837337482 |
| TFAP2A | 0.014741557686044909 |
| FOXA1 | 0.0007464072405028407 |
| KIF23 | -0.003814023656309869 |
| NCK1 | -0.023238340655216336 |
| CNKSR1 | -0.003309506835284202 |
| BICD1 | 0.029467848470741357 |
| PICK1 | -0.018798454080332132 |
| DSC2 | 0.0036990888717844854 |
| NUDT1 | 0.016584887220270798 |
| FEN1 | 0.039280027632950985 |
| CHAF1B | 0.06336904561093555 |
| THBS4 | -0.007936586153612108 |
| FAS | -0.0018303251038774466 |
| ZW10 | -0.009789089406096782 |
| ESPL1 | 0.0026349483067022485 |
| TTK | 0.00865778441087693 |
| MELK | 0.020732613609057232 |
| CCNF | 0.0017514255657294694 |
| RAD9A | 0.007814524589593192 |
| CDK8 | 0.026158579700205582 |
| POLA1 | 0.028022750740118868 |
| CP | 0.006502042373259053 |
| ORC2 | -0.0036782166622163805 |
| SERPINB5 | -0.03646908129424964 |
| IL6ST | 0.05064993589383435 |
| GCHFR | -0.00664382491807731 |
| PCSK2 | 0.004188397886511956 |
| MTERF1 | 0.002183754335489942 |
| MGMT | 0.0060875087326060125 |
| UGCG | 0.003266556069896875 |
| PLK4 | 0.027056234063494857 |
| SOX11 | -0.010375705499242788 |
| INHBA | -0.07031194038416438 |
| HPN | 0.06343944613246816 |
| PTPN2 | -0.011850499157072163 |
| CARD8 | 0.0009737029049671212 |
| MTAP | 0.026794832993818404 |
| ORC5 | -0.0008066718761636687 |
| PLK3 | 0.056517760847479115 |
| SHROOM2 | 0.016570941054944707 |
| CLOCK | 0.005225377238792192 |
| SLC22A18 | 0.0036381804080160784 |
| ITGB4 | 0.013350845909100321 |
| NF2 | -0.036644670398708716 |
| PFN2 | 0.030182653760940353 |
| ATF5 | -0.04388951833889739 |
| RAD51 | -0.05959231669664289 |
| CCNE2 | 0.018853635423271606 |
| ORM1 | -0.002784695993913156 |
| CENPE | 0.03231574820149773 |
| PSPH | -0.021255067676917743 |
| KIT | 0.02330305444052017 |
| AUH | -0.008742765931883351 |
| PRIM1 | 0.007640062297202148 |
| SLC22A5 | -0.024090201409722133 |
| SERPINF2 | 0.014325953337234937 |
| CRIP1 | 0.017160963848149774 |
| ORC1 | -0.005887435280780704 |
| MOK | 0.005371912559628551 |
| PEX11A | 0.0063813019014951405 |
| GLI3 | -0.14256324992470193 |
| KAL1 | -0.0011795132247511004 |
| GALK2 | 0.00012695444364892622 |
| ESR1 | -0.04057493084726801 |
| AREG | 0.051696773694660275 |
| GPSM2 | 0.007115593911870687 |
| CXCL13 | 0.07837155826906521 |
| EGR2 | 0.03901184423000849 |
| PER2 | 0.03325074136646313 |
| PGC | 0.005304266152110369 |
| LRP8 | -0.020463024120600193 |
| TFAP2C | 0.020189519687844052 |

| | |
|---|---|
| CDC14A | 0.002323084411661581 6 |
| IGFBP1 | 0.003091742143176988 |
| SLC15A2 | -0.0010016510545315768 |
| S100A1 | 0.007943911280445217 |
| DCT | -0.03064933540650469 |
| BARD1 | 0.020946437515230745 |
| AGTR1 | 0.012107886892131919 |
| INPP4B | 0.004323559735075347 |
| PDZK1 | -0.001581735302411606 |
| CHEK1 | 0.019745754982267764 |
| MLLT10 | -0.0003271636019289943 |
| DAG1 | 0.009834514052789574 |
| BCHE | 4.2763232913350035e-5 |
| OCEL1 | -7.907603710843584e-5 |
| HOXB2 | 0.03343632574992887 |
| C6orf106 | 0.0014717327485321869 |
| DACH1 | 0.027188465761183736 |
| UGP2 | 0.010926257825547094 |
| GCNT1 | 0.008369281276925853 |
| GATA4 | -0.06217314871779477 |
| SLC44A4 | 0.02573706775945674 |
| TRAIP | 0.007132092222931832 |
| LRP6 | 0.005758161221066203 |
| ANGPT1 | 0.04651626867644292 |
| PRIM2 | -0.010671924399342713 |
| PAX6 | 0.10863461454752399 |
| GAL3ST1 | 0.013072702833115058 |
| CYP27B1 | -0.01250224047819641 |
| HSD17B6 | -0.005185492295727337 |
| BLM | 0.012447397255563736 |
| KLK7 | 0.008128490790703463 |
| FGF7 | 0.027689500012265028 |
| TCP11L1 | 0.002348702682587310 7 |
| ROM1 | 0.012298719107757091 |
| SIX1 | -0.09910802932249513 |
| CPA4 | -0.001363436260613098 |
| JAK2 | -0.0037682467699033874 |
| TULP3 | 0.018105928783140265 |
| GREB1 | -0.000685505651094436 |
| RET | -0.01794275462279358 6 |
| MSH3 | -0.005832871205581587 |
| SLC22A4 | -0.005223516220411505 |
| CX3CR1 | -0.01877762630860521 3 |
| CCNA1 | -0.029898620261494986 |
| POLE2 | 0.022591169092778675 |
| S100A7 | 0.01911226924543656 |
| RELN | 0.0693005409065772 |
| IL27RA | -0.0038227634199274746 |
| MSX1 | -0.10305141919523 |
| BATF | 0.028505086668223368 |
| WNT5A | -0.07847281242458237 |
| GPR64 | 0.0018646637234725897 |
| NMU | 0.022327964446469235 |
| DICER1 | -0.0011654068895853738 |
| WT1 | -0.02453096159075084 |
| ACADL | 0.017504038742022124 |
| HMGA1 | -0.005375502601779681 |
| LRRC23 | 0.0008850087968634606 |
| CTH | 0.0368370720162585 5 |
| NELL1 | 0.01071026244982359 |
| GINS1 | 0.005782639831933587 |
| ISL1 | 0.0042831521179754915 |
| PLA2G5 | 0.06866360274356277 |
| WNT10B | -0.05960610425415587 |
| EDA | -0.012953896218986433 |
| TNFRSF10C | 0.0022202442847365058 |
| PAX2 | -0.1065520693269734 3 |
| LHX1 | -0.04795659968052898 |
| GUCY2C | -0.011412771717607462 |
| PRLR | 0.04369735823572912 |
| PDK3 | 0.009255883892137948 |
| KIF14 | 0.015300681464566696 |
| CPLX2 | 0.016297126542175237 |
| EYA3 | 0.008291707317188424 |
| ANK3 | 0.016523807369716526 |
| SLC1A1 | -0.012628381153536341 |
| ZNF205 | 0.002972237203413029 |
| TBCCD1 | 0.002407925925447312 5 |
| NDRG2 | 0.01967272037070913 |
| KIAA0125 | 0.0004132393741945018 |
| HINFP | 0.014878738381224676 |

| | |
|---|---|
| PML | -0.04791240195835318 |
| MGAM | -0.011773852786785862 |
| ABAT | 0.01909954045691702 |
| KLHL24 | 0.00032070457449530636 |
| SMA4 | 0.0020387403396643564 |
| CDKL5 | 0.018288948441041876 |
| GFI1 | 0.031571796504911286 |
| DRD2 | -0.021851368214878313 |
| SIX3 | 0.06346645664152692 |
| CHRNB2 | -0.03847303987011225 |
| GLI1 | -0.03497752184708723 |
| FLT3 | -0.020259913217113883 |
| PDK1 | 0.025638035768205728 |
| ASIC2 | 0.05260758220733848 |
| IL7 | 0.015592135087047685 |
| CHRNB1 | -0.00019164268514129993 |
| KCNJ2 | -0.01756720575902059 |
| ACRV1 | -0.0006026324800037688 |
| ERBB4 | -0.0084868856195354 |
| SCGB1D2 | 0.0004387758538985074 |
| STATH | -0.0053950355600486 |
| ALX1 | 0.02986215295181513 |
| AFM | -0.0016939199422974886 |
| PAEP | 0.006354664338546462 |
| CHAD | -0.013078713959233528 |
| AMHR2 | -0.002368030217064604 |
| SALL1 | -0.08840567092421867 |
| TAT | 0.016862964075962113 |
| PRKCA | -0.0006614508433366013 |
| SRD5A2 | -0.015277224904423726 |
| TGFBR1 | 0.025894693239995895 |
| HTR6 | -0.009662179726478119 |
| C8B | 0.0025303041239801514 |
| PHOX2B | 0.011495585806956119 |
| PTK7 | -0.0035529964314248742 |
| ALDH1A2 | -0.12135910958184191 |
| LDHC | 0.016357644900862892 |
| CRISP1 | 0.000264454942651869 |
| GLI2 | 0.10206865475980201 |
| TNFRSF11A | 0.07533420474772554 |
| SLC16A6 | 0.002083699361257855 |
| CDKN2A | 0.10917112540001164 |
| THRB | 0.029168583684075385 |
| ASS1 | 0.004830149327111394 |
| LEP | 0.1998378331370313 |
| CITED1 | 0.11631010981891542 |
| AKT1 | 0.13258238507628145 |
| ZBTB18 | 0.0139589612891684 |
| AGRP | -0.031067674487839272 |
| ALOX12 | -0.07043025237679759 |
| SLC15A1 | -0.02213285966238262 |
| SLC17A3 | -0.013743047633786646 |
| TCF15 | -0.0425778237859497 |
| HTR2C | -0.011836730934854097 |
| TFRC | 0.022648351025412853 |
| STX2 | 0.005044391242978731 |
| CYP7B1 | 0.0019296241757945803 |
| HOXD13 | -0.008245711200176871 |
| PROX1 | 0.06596269150182503 |
| RAD17 | -0.0012069700278087017 |
| MSMB | -0.0013331384342858382 |
| NR2E1 | -0.09852641803918796 |
| CAST | -0.03052701044183323 |
| ATP5G3 | 0.0018423725159665725 |
| CDKN2B | -0.05876492166517278 |
| IL4 | 0.018822780983768387 |
| PITX2 | 0.10480238529079346 |
| MAGEB3 | -0.006916581004888721 |
| SHH | 0.08366559408643312 |
| MMP20 | -0.017020842905839707 |
| SLC4A7 | 0.010467453451719635 |
| FOXH1 | -0.024641509894376825 |
| ADAM2 | 0.006458059209497759 |
| PAX3 | -0.05906825353319958 |
| KIF25 | -0.009609396255554063 |
| SERPINH1 | -0.007563636479709535 |
| PPP2R3A | -0.0006220842487928135 |
| RASSF8 | 0.0033201926705127994 |
| RBPJ | 0.05165924746472582 |
| KRT33B | 0.0018854914919091375 |
| POU1F1 | 0.00196846360803228 |

| | |
|---|---|
| MUC1 | 0.015340453178736262 |
| AVP | -0.036239740177752014 |
| INSR | -0.07200559687285481 |
| NAALADL1 | -0.0010165593282607669 |
| IL12B | 0.09369902304859438 |
| GATA2 | -0.019232401633833166 |
| MEF2C | 0.09777138637302664 |
| CLEC4M | -0.024936805204337182 |
| RFC1 | 0.023320146009614223 |
| CDC14B | 0.0001570744684361436 |
| HMGA2 | 0.07233033972660001 |
| LAIR1 | 0.001294624618107643 |
| CCL7 | -0.05525290613251585 |
| ZNF442 | 0.0021217095248059524 |
| NDEL1 | -0.014938650791860787 |
| ANP32E | 0.0008414262211640951 |
| DDX17 | 0.010624504900571069 |
| CHIT1 | 0.0017042028202982392 |
| PSG4 | -0.0010328661238374205 |
| TTN | 0.03969133562208975 |
| MAS1 | 0.005871102423210317 |
| DRD4 | -0.07577515006143414 |
| HOXB1 | 0.048177605227734295 |
| TH | -0.022989804685408042 |
| PGR | 0.005606482706949607 |
| CSH1 | -0.0010294405113189218 |
| UGT8 | 0.021797239011992684 |
| BRCA2 | -0.0004001008524303959 |
| FGF5 | -0.02945446553198055 |
| MMP24 | 0.010426711820916278 |
| TBPL1 | -0.009405012825298755 |
| PTN | 0.007133781784054908 |
| IRF7 | -0.054667593900032764 |
| FGF8 | 0.0714011127898124 |
| GABRA4 | -0.023644228919724872 |
| BAX | -0.04740219544026608 |
| PTTG3P | -0.00020876180747159113 |
| OR5I1 | 0.009398362467484081 |
| PTCH1 | -0.0035436404931479233 |
| BCL2L11 | -0.03770305268840599 |
| PDX1 | 0.010516918867804344 |
| WNT1 | 0.04369680334753987 |
| HOXA3 | 0.013754711326618266 |
| WNT4 | -0.016164899247737858 |
| EIF4G1 | 0.011039070602603436 |
| YBX1 | -0.010053564233932747 |
| CD24 | 0.02988863062484949 |
| MAGED2 | 0.001145939693081225 |
| PRKDC | -0.07789656076747947 |
| NRD1 | -0.001053191850308878 |
| CCND1 | -0.18493934739300474 |
| CDC42 | 0.11480020475125494 |
| FLOT1 | -0.0202378900509657 |
| PSMD11 | -0.04560858541280287 |
| ELOVL5 | 0.002689234967436743 |
| MCM7 | -0.023035440617418333 |
| CHD3 | 0.0010863652782815024 |
| HMGB2 | 0.010195853792630813 |
| SNRPB | 0.009292643196300647 |
| POLE3 | 0.008058861478672534 |
| ATP1B3 | -0.04910506135762837 |
| STK24 | 0.04315211111121438 |
| SLC1A5 | 0.013826124416096994 |
| POR | 0.009350598710947199 |
| KPNB1 | 0.01603367391667606 |
| MDK | 0.031790169158598905 |
| TXN2 | 0.020657983182013912 |
| JAG1 | 0.06489289364190193 |
| TSPAN6 | 0.005498471709099 |
| QDPR | -0.04019139695326161 |
| AGR2 | 0.017854233491790952 |
| CXCR4 | 0.019727755787297355 |
| LMO4 | 0.023297350079814507 |
| CRYAB | 0.02158200226853595 |
| NFIB | 0.05601209399912987 |
| ECM1 | -0.021813657033172763 |
| STXBP2 | -0.009931038665850008 |
| KIF2C | 0.007483289154081434 |
| FZR1 | 0.017831600102073526 |
| MSH2 | 0.041758053283648636 |
| SERPINA5 | -0.004965822186561536 |

| | |
|---|---|
| AURKB | 0.0075737352529248025 |
| LRP5 | 0.0014000670266215848 |
| RPA3 | 0.09428235665230202 |
| HSDL2 | 0.002733916507979596 |
| CACNB3 | 0.0033579975839442134 |
| IGF1 | 0.014117033104510663 |
| YIPF4 | 0.0010880978793895205 |
| THBS3 | 0.01615380344497765 |
| EPHB2 | -0.017254484462277564 |
| BMP7 | 0.058579874415387936 |
| GATA3 | -0.003547293725863641 |
| PAK1 | -0.12024530596624507 |
| MCCC2 | -0.007785154522843757 |
| BUB1 | 0.023772293746745807 |
| KIFC1 | 0.008559426097992033 |
| S100B | 0.08900178023831676 |
| SLC35D1 | -0.0007554871546431536 |
| TGFB3 | -0.13674655463203483 |
| LLPH | -0.00021432664927591781 |
| JAG2 | -0.00016145912642093926 |
| PADI2 | 0.0051776286151684 |
| NPAT | -0.0035409649949677555 |
| VLDLR | 0.02188999182983962 |
| CDT1 | 0.013044432987764098 |
| SOX10 | 0.0037792197723009184 |
| HOXB13 | 0.016446173550618 |
| SELPLG | 0.0022892769697621758 |
| SLIT2 | -0.07791903893529609 |
| TGFB2 | -0.11937808737609407 |
| PARP3 | 0.01543425994457865 |
| FBXL4 | 0.004265644090267212 |
| GSK3B | 0.13906893656117805 |
| VEGFC | -0.12818045716407228 |
| FAP | -0.007189592876954761 |
| ASCL1 | -0.06877170297567123 |
| GATA6 | -0.001112533276770195 |
| APEX1 | 0.0035320394450440943 |
| IDO1 | 0.02684206122285139 |
| PGM3 | 0.02388703108559418 |
| TPX2 | 0.009385015564002138 |
| CTSV | 0.012273549678511544 |
| AGER | -0.008157913176116416 |
| IGFBP3 | 0.016694774973335498 |
| CYP4B1 | -0.008157465159450511 |
| FOXA2 | -0.0008266461929169126 |
| INHA | 0.005488558464491146 |
| PTPRJ | -0.00701621521197242 |
| CRYAA | -0.00400758932177829 |
| WNT7A | 0.0010529178437344018 |
| CYP2B7P | 0.00013132082203898846 |
| RECQL5 | 0.012637809951031562 |
| C8G | -0.010678030681020806 |
| BCL11A | -0.01459751416609687 |
| IFNG | -0.004231728952820178 |
| RBBP4 | -0.03534281334268238 |
| GLUD2 | -0.007176789123447408 |
| GPR125 | 0.0012335524835453977 |
| NRP1 | -0.04242006248869886 |
| VEGFA | 0.06666330837534155 |
| MSH4 | -0.0014911312037883808 |
| SPAM1 | -0.01712007318515972 |
| POLDIP3 | -0.0003304417171146347 |
| MFGE8 | 0.016281240676370122 |
| EPHB1 | 0.045318873159804755 |
| CALCOCO2 | 0.00987327046847262 |
| ACVRL1 | 0.0855696402292806 |
| LEF1 | -0.008738760148707034 |
| GCNT2 | 0.0365829825538686 |
| AURKC | 0.009679931011797385 |
| AR | 0.08189387341717852 |
| IFNA2 | -0.07366269960649005 |
| PADI4 | -0.00904727123774782 |
| BMP4 | -0.051948548495902636 |
| GNRHR | 0.0036862891801634633 |
| HSPA2 | -0.0021826014130113995 |
| ETV4 | 0.0017136601329301399 |
| FBL | -0.0018366177387389563 |
| BDH1 | -0.001844661543438061 |
| PRG2 | -0.007673598049155522 |
| GINS4 | -0.011529877940197964 |
| KMT2D | -0.0025206311151364153 |

| | |
|---|---|
| IGLJ3 | 0.003248875669633096 |
| IGK | 0.002947273319160409 |
| PLCB1 | 0.010705113862137893 |
| ITGB1 | 0.15327054269447865 |
| IPO5 | 0.010054180206643164 |
| TMEM123 | -0.001369198395808352 |
| TOP2B | 0.003444803971710198 |
| MKI67 | 0.014176230928602424 |
| TPGS2 | 0.0018304301414036241 |
| KMT2A | 0.01548594762858203 |
| SLC39A14 | 0.0009186088282803245 |
| MCM4 | -0.01213181177975491 |
| RNF187 | -0.011075139468168136 |
| SERPINE2 | 0.06795055647096448 |
| MAP1B | -0.005304206896318236 |
| SEL1L3 | 0.001482943805671272 |
| YIPF6 | 0.0004924847107853045 |
| SULF1 | -0.017637187191465113 |
| FEM1B | -0.029090201626783607 |
| MYH10 | -0.009530578464474467 |
| ZCCHC24 | -0.0006336174544859327 |
| KIAA0232 | -0.0025749317470091728 |
| TENC1 | -0.0016540678942851379 |
| CEBPB | -0.00914299377376715 |
| GPD1L | 0.01115513164353617 |
| SPIDR | -0.01805494699474201 |
| CDC34 | -0.0021652700354158005 |
| ZNF609 | -0.002093260493643199 |
| METAP1 | -0.00802137483893706 |
| SLC25A44 | 0.0003453198541108319 |
| JMJD6 | 0.07225247109460835 |
| C16orf45 | 0.0015129853607803165 |
| RCHY1 | -0.012047180085251727 |
| BBS4 | -0.02449128618535784 |
| TCF7L2 | 0.022737387684181265 |
| FAM171A1 | -0.0008474461993496482 |
| OBSL1 | 0.0004996753392981063 |
| NCAPD3 | -0.006291945324096311 |
| CBS | 0.027408147568601854 |
| CKAP5 | -0.007844218411740494 |
| POLD3 | 0.04009422205891675 |
| AXIN1 | -0.04008779072142446 |
| DNAJC16 | -0.00010774179655897079 |
| SMC5 | -0.008404355246720136 |
| NCAPH | 0.0005077986713116514 |
| KANK1 | 0.03570418486402972 |
| NEDD4 | 0.08611271769809645 |
| LOC100272216 | -0.0004901303998456606 |
| DNAJC2 | -0.00785480274112632 |
| HOXA10 | -0.03585348281798796 |
| ROBO1 | -0.03381507510547673 |
| TNNT1 | 0.0023616702764739823 |
| FOXC1 | -0.0010032433396977912 |
| YAP1 | -0.005210933274387357 |
| ATAD2B | -0.0012403210541986126 |
| SPDEF | 0.03149979170441675 |
| SOSTDC1 | -0.012182241980776633 |
| CLCN2 | 0.008450306755612699 |
| CCNE1 | 0.01335756392164103 |
| DNA2 | -0.0005924515358255836 |
| DLX5 | -0.06904625505431025 |
| ELOVL2 | 0.0060805461246128895 |
| HIPK2 | 0.028394484848613805 |
| HOXA5 | 0.02651939251917732 |
| TWIST1 | -0.07355870775008258 |
| CHD5 | -0.004060582690920924 |
| YPEL1 | 0.0028447557066546337 |
| CCL8 | -0.014706434092778392 |
| LIAS | -0.004366434702725852 |
| SHMT2 | -0.013105990524783962 |
| XIST | -0.0013837068271472977 |
| GMPS | 0.017799732080132098 |
| FOXC2 | 0.06632063111815427 |
| PTTG2 | -0.013710164253373864 |
| FGF3 | 0.04819322200362268 |
| AZU1 | 0.005302910090948995 |
| TNFRSF21 | 0.006502653335445798 |
| PAK3 | -0.04013955338735343 |
| EYA1 | 0.0017480075922933968 |
| ZNF266 | 0.0008265747472587705 |
| CCNB1 | -0.035987139340594845 |

| | |
|---|---|
| PP14571 | -0.0025847593046036026 |
| KCNMA1 | -0.02152134514870762 |
| GRIK5 | -0.0028890162313301585 |
| RNASEH2B | -0.0003486624368721745 |
| GRB2 | 0.028244434541645335 |
| DNM1 | -0.02218182942553515 |
| CEP152 | 0.01624295019809501 |
| STOML2 | 0.005173659861751104 |
| IHH | -0.0981336737412036 |
| EPHA5 | 0.006530393686639467 |
| RAD21L1 | -0.011228435521741091 |
| POLE | 0.05615385969439243 |
| RAB3GAP2 | 0.013865146208784956 |
| LAMB2 | -0.02355652281477593 |
| MUC7 | -0.015812317014986945 |
| CCL2 | 0.07777608367830821 |
| IFT122 | -0.03543911313873815 |
| LOC100293211 | 0.0022504739740209054 |
| DISC1 | -0.03731677569455142 |
| KCNV2 | 0.003696900978180594 |
| ORAI2 | 0.002164600637607095 |
| ARPIN | -0.000581488877706825 |
| STEAP1B | 0.002804840397185504 |
| SEC14L1P1 | 0.004847184538895705 |
| AES | 0.016231813272144714 |
| PERP | 0.014624321373756953 |
| NAA50 | 0.000696110791108242 |
| HN1 | -5.8127791173796244e-5 |
| C3 | 0.08302685884379522 |
| UFC1 | 0.002081909240303292 |
| EVL | 0.02326562508306125 |
| CDC27 | 0.007420671443608753 |
| DSG2 | 0.00017329429395627894 |
| KLHDC2 | -0.00037589909397841464 |
| VPS4A | -0.01888574164846359 |
| FAM127B | 0.0020688261001252576 |
| PRC1 | 0.004887640416565871 |
| PPDPF | -0.004333499725805429 |
| FAM65A | -0.0033104341627444417 |
| RBM47 | 0.0008135753629212928 |
| NUSAP1 | -0.0031989576398166906 |
| BFAR | -0.0032646454557970625 |
| WDR11 | 0.0004084790987555992 |
| C21orf59 | 0.0012766631140000372 |
| SLC25A37 | -0.0011019483415305995 |
| SMAP1 | 0.004010691731589972 |
| GSDMD | 0.0023281873418337165 |
| GOLT1B | -0.009110027865084445 |
| STMN3 | 0.020792249166952208 |
| TMEM258 | 0.0036313222830984962 |
| LANCL2 | 0.0035635361194808896 |
| C1QA | 0.005377251754758938 |
| TSKU | 0.0065141041458630896 |
| CKAP2 | 0.00018595795606970813 |
| MKL2 | -0.0007206805073324869 |
| BCCIP | 0.005814671919895755 |
| SAV1 | -0.03272071156694 |
| LGR4 | 0.00657108329594273 |
| PRDM4 | 0.008370680347239902 |
| KIF4A | 0.015508808710484125 |
| PGLS | 0.010472817234260473 |
| FAM204A | -0.0044788576096735245 |
| ABT1 | 0.01796166914155394 |
| NDE1 | -0.03188184230872025 |
| STEAP3 | 0.028820763187179237 |
| MCCC1 | 0.0007363760398943214 |
| TTC4 | 0.0003864077187552613 |
| CMC2 | -0.00024523649873989695 |
| GREM1 | -0.036280446583260524 |
| KLF11 | 0.0037117930552162124 |
| RNASEH1 | 0.0016941513093079103 |
| TRPS1 | -0.002804754380042559 |
| DCP1A | -0.0059072595504989014 |
| LPPR2 | -0.0013568981277233031 |
| PNPO | -0.01199231896941335 |
| MAP1S | 0.0042251920556763654 |
| E4F1 | 0.017773349853668394 |
| MRS2 | 0.0019480424105514202 |
| C8orf4 | -0.0010783892034500846 |
| CEP55 | 0.00018269415358942805 |
| KBTBD4 | 0.002194444134936169 |

| | |
|---|---|
| RINT1 | -0.006919338654133498 |
| SMO | 0.1371278839293916 |
| NCAPG | 0.000693116427805255 |
| S100A14 | -0.0151651506219373 |
| NES | 0.015184277222533588 |
| GINS3 | -0.00027924086714144783 |
| HJURP | -0.00045926730263408903 |
| CDK5RAP3 | 0.02262447700144694 |
| KIF20A | -0.0016866338094267347 |
| SMC6 | -0.013984717819969966 |
| ATAD2 | -0.00019412537390888472 |
| FCGRT | 0.0017651946099501248 |
| IGF2BP2 | -0.004613114653713913 |
| PPP1R13L | -0.017696251377432606 |
| FBXO5 | 0.00750373341738888 |
| SIRT1 | -0.0009599762338105295 |
| NOTCH1 | 0.06908067138798715 |
| INTS8 | -0.004526390346986949 |
| RAB17 | 0.0023503069475402905 |
| HSPB7 | 0.001584431842469641 |
| KRT23 | -0.0007346504263966152 |
| DNAJC12 | 0.0007439037392301426 |
| METRN | -0.005624192028500556 |
| CYB5R4 | -0.00021192099129544977 |
| CTPS2 | 0.014098873505983605 |
| SHQ1 | -0.0043061549669067166 |
| IL20RA | 0.0008584941824028738 |
| PBK | 0.0050207360634157705 |
| BIN2 | -0.0065614115670257645 |
| SCUBE2 | 0.0025221661308145404 |
| HSPA14 | 0.00027877884879170975 |
| RBKS | 0.000886240999496021 |
| PHACTR4 | -0.01607924402078718 |
| CEP63 | -0.0001574280868425614 |
| SPHK1 | 0.06878627113250542 |
| TIPIN | -0.0048284433759789 |
| C1orf159 | -0.0013948275277618292 |
| DIABLO | -0.00534403595986967 |
| ATHL1 | -0.000990180753650577 |
| KLF2 | -0.023289426706930974 |
| GRHL2 | 0.0250921327001324 |
| TTYH1 | 0.009566840787928563 |
| COA7 | 0.0005966881548589191 |
| TTC33 | -0.0026210948750966644 |
| SULT4A1 | -0.005814538701167441 |
| EGFL6 | 0.0008221974015097802 |
| C7orf63 | 0.0012350401404626569 |
| LAMP5 | -0.0035792461217451187 |
| LRFN4 | 0.0013939386408319413 |
| GCNT3 | -0.02550658691485919 |
| CEP72 | -0.016425670499335236 |
| BORA | 0.011964177974982485 |
| ROBO3 | 0.004799815360220844 |
| SOX18 | 0.029500021112874815 |
| KIF16B | 0.035238187510967627 |
| PIEZO2 | 0.002089829323134258 |
| CEP83 | -0.0007677935602652968 |
| PTPLA | 0.002554874314263294 |
| TBX3 | 0.050105796897260475 |
| FZD3 | 0.009331988846620856 |
| BBS7 | -0.020445120694412334 |
| TFCP2L1 | 0.004859367748266087 |
| HEY2 | 0.13883585381515862 |
| VTCN1 | -0.011060391877614666 |
| GIMAP6 | -0.0007352876517308432 |
| MTL5 | -0.0025622556198957322 |
| ECT2 | 0.024320114192369503 |
| ARL15 | -0.00453672821623362 |
| NARF | 0.0014994215563336724 |
| FAM198B | 9.679876491163104e-5 |
| LHX6 | 0.009407866917839706 |
| SPAG4 | 0.0006137799805040559 |
| ASPM | 0.0002548203396439115 |
| GLRX2 | 0.01667586375117496 |
| GPR87 | -7.288109362732303e-5 |
| GALNT6 | -0.010690878808872862 |
| KIZ | 3.3117504151390165e-5 |
| TXLNG | -0.005025118529839372 |
| E2F8 | 0.020587127716812 |
| AUNIP | -0.0004402857743541961 |
| ERO1LB | 0.0018389095331322225 |

```
MFI2    0.004359643072220507
EDAR    0.005933150194892734
GDF3    -0.0394937194432559
STAP1   0.0319230290662234 5
HELLS   0.027898557808903234
STEAP4  -0.013913880365065035
CAMK1D  -0.031182194982635202
ARPP21  -0.0008943828850588782
ALLC    -0.002765043249267739
ROPN1B  0.004169631433242043
DMRT1   0.02853845282955852
EN1     -0.0002179432341088694
CCDC170 -0.0009410021876603598
SIRT2   -0.048816855988439746
FOXE3   0.014651722139499572
ELF5    0.011752955688516508
MCM10   -0.004191810376388577
KIF24   -0.011283208386569685
HEYL    0.013492537971960204
TBX21   0.003869400748971832 3
SLC5A7  0.020065233487922305
CCDC7   -0.0014199489523593887
EDDM3B  0.0010086661202008467
TP73    0.04204531239272019
GCNT4   0.005380114461413647
C1orf112        0.0015332318190178 08
PSAT1   -0.005765858129630002
WDR19   -0.010047564609988222
C1QTNF3 0.0015438782131631148
ARHGAP24        -0.007478400689910492
TEX13B  -0.0009161916424618884
TEX14   0.01549261154539974
KCNE2   -0.011773142899107384
EPN1    -0.004333698632650038
RPA4    -0.02490832447198216
MAP3K19 -0.008239112962222229
IQCG    0.0014698666492631834
LOC100131532    -0.005151893057862085
STMN4   0.001395983811242588
KIF18A  0.01285037659610607
DLX6    -0.001800012326516208
KLF15   0.003681754231953001
FRS2    0.02771376092040482
NEUROD4 0.015071128029595659
C6orf25 -0.000498361078864335
GDNF    0.014056253615913419
TMEM14B 0.003590952595578689
BNIP3L  -0.00046372092071295264
KPNA3   -0.0040731287551446956
GINS2   0.0036502344793744894
SOAT1   -0.003584009922054709
FAM64A  0.00022582903076745383
DSPP    0.004961435490698742
SPDL1   0.02250710639363425
STRA6   -0.06393712883327846
COL5A2  -0.005812223762505951
NRBF2   -0.006598199026985762
GPR124  -0.0024255751217839277
KLHL22  0.003054254639106805
ORAI3   -0.0026187771148240358
FAM174B -6.72904177286392e-5
ZNF335  -0.004675023254393562
SEH1L   -0.02255406508872177
LOC389906       0.005430567767702915
KIF18B  0.015702773273153527
RACGAP1 0.026271373792290352
CASP8AP2        0.012679791553399915
KLK5    0.006951607495603109
OR7E47P 0.000576342576718726
SCAF4   -0.00032868048497175506
SNHG17  -0.0011774470329610938
Age                     -0.002246735353315669
intercept                       -0.06328011266889626
```

In [ ]:
